# A sexual dimorphic role of the *miR-465* cluster in placental development

**DOI:** 10.1101/2021.01.16.426973

**Authors:** Zhuqing Wang, Nan Meng, Yue Wang, Tong Zhou, Musheng Li, Shawn Wang, Sheng Chen, Huili Zheng, Shuangbo Kong, Haibin Wang, Wei Yan

## Abstract

A sexually dimorphic role of miRNAs in placental development has never been reported. Here, we show that ablation of the *miR-465* cluster caused selective degeneration of female conceptuses as early as embryonic day (E)8.5, leading to a male-biased sex ratio (60% males) among *miR-465* KO mice. Given that the *miR-465* cluster miRNAs were predominantly expressed in the extraembryonic tissue and ablation of these miRNAs led to dysregulation of numerous critical placental genes, our data strongly suggest the *miR-465* cluster is required for full developmental potential of the female, but not the male, extraembryonic tissue/placenta.

## Background

miRNAs are ~22 nucleotide small non-coding RNAs that regulate gene expression at post-transcriptional levels [1]. Inactivation of either DICER or DROSHA, the two enzymes required for miRNA biogenesis, leads to early embryonic lethality in mice, indicating an essential role of miRNAs in early development [1]. Our previous studies have shown that an X-linked *miR-465* cluster, which encodes 6 pre-miRNAs and 12 mature miRNAs, belongs to a large X-linked *miR-506* family [2]. High abundance of these miRNAs in the testis, sperm, newborn ovary as well as in 8-16-cell embryos and blastocysts [2–5] suggests a potential role of this miRNA cluster in gametogenesis and early embryonic development in mice. However, their physiological role has not been investigated *in vivo*. Here, we report that the *miR-465* cluster miRNAs are also abundantly expressed in developing placenta, and ablation of the *miR-465* cluster does not affect fertility, but causes a skewed sex ratio favoring males due to selective degradation of the female placenta during early embryonic development.

## Results and Discussion

The *miR-465* cluster consists of 6 miRNA genes encompassing a ~16.4 kb region on the X chromosome (Fig. 1A). Although 6 pre-miRNAs and 12 mature miRNAs are produced, only 6 mature miRNAs can be distinguished based on their sequences, including *miR-465a-5p, miR-465b-5p, miR-465c-5p, miR-465d-5p, miR-465a/b/c-3p* and *miR-465d-3p* (Fig. 1A). To define their physiological roles, we deleted the entire *miR-465* cluster using CRISPR-Cas9 (Fig. S1), as previously described [2, 6, 7]. The *miR-465* KO mice were fertile, have normal testis size (Fig. S1D), and both the litter size (8.4 ± 0.85, n=35) and interval (25.4 ± 0.86, n=35) of the KO mice were comparable to those of wild type (WT) controls (Litter size: 8.6 ± 1.59; litter interval: 26.6 ± 1.42, n=23) (Fig. 1B), suggesting that these miRNAs are dispensable for both spermatogenesis and folliculogenesis.

**Figure 1.**
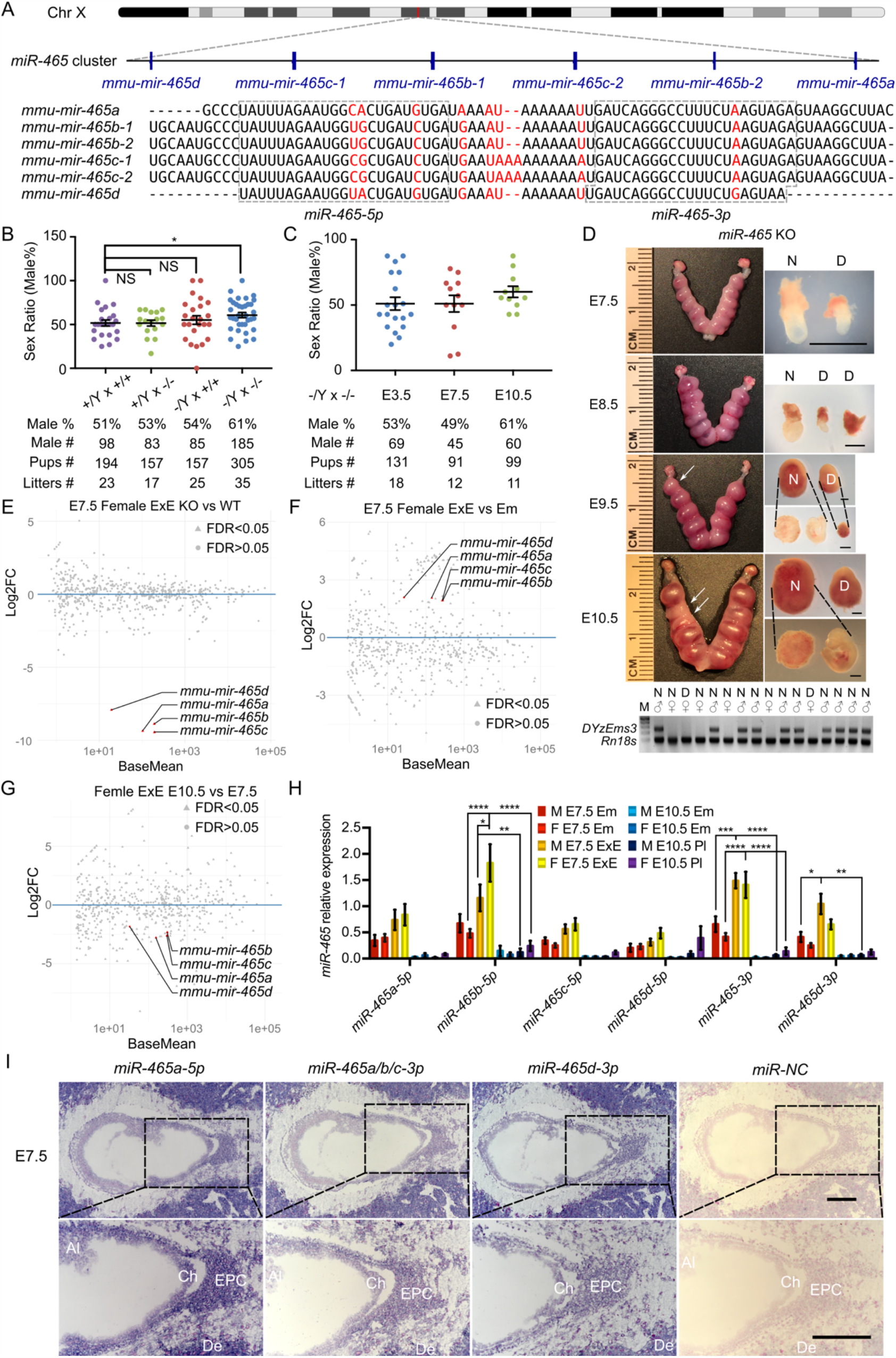
Phenotype of *miR-465* KO mice and expression pattern of the *miR-465* cluster. A. The genomic location and sequences of the *miR-465* cluster on the X chromosome. B. The sex ratios among pups from different breeding schemes. *, p<0.05; NS, statistically not significant. C. The sex ratios of pups from homozygous inbreeding (−/Y × −/−) at E3.5, E7.5, and E10.5. D. Representative images of the KO uteri and conceptuses collected between E7.5 and E10.5 (upper panels) and PCR-based genotyping results (lower panels). Arrows point to the degenerating/degenerated conceptuses (D) among the normal-looking (N) ones. Scale bars = 1mm. *DYzEms3*, a Y-linked genomic fragment, was amplified to identify male conceptuses, and *Rn18s*, which encodes 18s ribosomal RNA, was used as a loading control in the PCR-based genotyping analyses. E-G. Differentially expressed miRNAs between the following comparisons: WT vs. KO female extraembryonic tissues at E7.5 (E), WT female extraembryonic tissues vs. embryos at E7.5 (F), WT female extraembryonic tissues/placentas at E7.5 vs. E10.5 (G). Data points representing the *miR-465* cluster miRNAs are highlighted in red. sRNA-seq analyses were conducted in biological triplicates (n=3). H. TaqMan qPCR analyses of expression levels of the *miR-465* cluster miRNAs in extraembryonic tissues/placenta and embryos at E7.5 and E10.5. M, male; F, female; Em, embryo; ExE, extraembryonic tissue; Pl, placenta. *, p<0.05; **, p<0.01; *** p<0.001. I. Representative miRNA-ISH results showing localization of the *miR-465* cluster miRNAs in female conceptuses at E7.5. Ch: chorion; EPC: ectoplacental core; Al: allantois; De: decidua. Scale bars = 200 µm.

Interestingly, unlike equal distribution of both sexes (~50%) among pups from the WT breeding pairs (+/Y × +/+), the sex ratio is significantly skewed towards males (185 males out of 305 mice, t-Test, p<0.05) among the *miR-465* KO pups derived from the homozygous breeding pairs (−/Y × −/−) (Fig. 1B). It has been reported previously that certain dietary changes, *in vitro* fertilization procedures and genetic mutations can cause biased sex ratio [8–13], and of interest, 60% appears to be the most common sex ratios observed in these studies (Table S1).

The skewed sex ratio could result from either a skewed X/Y sperm ratio or loss of female embryos/fetuses during development. If the sex ratio is already skewed in X/Y sperm, the bias should be observed among pups from the breeding pairs of KO males (−/Y) and WT females (+/+), but not in those from the breeding pairs of WT males (+/Y) and homozygous KO females (−/−). However, the sex ratio among the pups from the −/Y × +/+ breeding scheme was slightly, but not significantly, skewed to males (54%) (Fig. 1B), suggesting that the significantly skewed sex ratio likely occurs during development. Males accounted for ~50% among all of the KO embryos at E3.5 and E7.5, whereas the ratio of the males increased to ~61% at E10.5 (Fig. 1C), suggesting that some female embryos are lost between E7.5 and E10.5. Indeed, we observed that on average 1-2 conceptuses per uterus were either being absorbed or had already been absorbed between E8.5 and E10.5, and more intriguingly, 6 out of 7 resorbed conceptuses were all female KOs (Fig. 1D). Earlier studies have shown that 1-2 embryos of equal sex distribution undergo spontaneous resorption between E7.5-E10.5 in a pregnant female mouse of the C57BL/6J strain [14, 15]. Our observation that the resorbed conceptuses are mostly KO females strongly suggests that the lack of *miR-465* cluster specifically causes female conceptuses to degenerate. A similar number of early embryonic loss (1-2 conceptuses per pregnancy) during E7.5 and E10.5 between wild type and KO pregnancies may explain why we observed a similar litter size in both wild-type and *miR-465* cluster KO breeding pairs.

Although loss of the *miR-465* cluster leads to female-biased lethality, it remains unknown whether the primary defects lie in embryos or extraembryonic/placental tissue. To address this question, we collected both WT and KO embryos and extraembryonic/placental tissue from both sexes at E7.5 and E10.5, and performed small RNA deep sequencing (sRNA-seq) (Fig. 1E-G). sRNA-seq data confirmed that the *miR-465* cluster miRNAs were truly absent in the KO embryos and extraembryonic/placental tissue (Fig. 1E, S2A-S2C). While no significant sex differences in miRNA levels were observed in WT embryos and extraembryonic tissues at E7.5 (Fig. S2D, S2E), the *miR-465* cluster miRNAs were predominantly expressed in extraembryonic tissues, as compared to embryos of both sexes at E7.5 (Fig. 1F, S2F), and these miRNAs were significantly downregulated from E7.5 to E10.5 (Fig. 1G, S2G-S2I). Indeed, the TaqMan real-time PCR analyses further confirmed the sRNA-seq results (Fig. 1H). To determine the localization of the *miR-465* cluster, we further performed miRNA *in situ* hybridization (ISH) assays (Fig. 1I).

Consistent with the sRNA-seq and qPCR data, miRNA ISH results showed that the *miR-465* cluster miRNAs were predominantly expressed in extraembryonic tissues, especially in the ectoplacental core and chorion (Fig. 1I). Although the *miR-465* cluster miRNAs were also detected in maternal decidua (Fig. 1I), potential decidual defects are highly unlikely based on our breeding data showing normal sex ratio among offspring of the +/Y × −/− breeding pairs (Fig. 1B). Given the predominant expression of the *miR-465* cluster in the extraembryonic tissues, it is highly likely that the loss of some female embryos was secondary to placental defects.

To identify targets genes of the *miR-465* cluster, we conducted RNA-seq on WT and KO embryos and extraembryonic tissues of both sexes at E7.5. We chose E7.5 because obvious degeneration and resorption were rare at this timepoint, but transcriptomic alterations should have accumulated (Fig. 1D). Principal component analyses (PCA) identified two major clusters, each containing either embryos or extraembryonic tissue of both WT and most of the KO of both sexes except two outliers (Fig. 2A). The two outliers turned out to be one female KO embryo and its extraembryonic tissue, suggesting that this conceptus most likely represents a “to-be-degenerating” KO female. While WT and non-degenerating KO embryos and extraembryonic tissue of both sexes displayed similar mRNA transcriptomes (Fig. 2B, Table S2), numerous differentially expressed genes (DEGs) were identified between the extraembryonic tissue from the “to-be-degenerating” KO female and that from non-degenerating KO females (Fig. 2B, Table S2). Gene ontology (GO) term analyses identified that the DEGs were mostly involved in extraembryonic/placental development (Fig. 2C). Among the DEGs, two genes, *Rlim* (also known as *Rnf12*) and *Alkbh1*, were noted because of their known functions. *Rlim* is an X-linked gene highly expressed in extraembryonic tissues at E7.5 [16]. *Rlim* appears to be responsible for imprinted XCI by maintaining *Xist* expression [17] and its dysregulation leads to female-biased lethality [8, 17]. *Alkbh1* is a tRNA demethylation enzyme [18] highly expressed in chorion and the ectoplacental cone at E8.5 [19]; its ablation also induces female-biased lethality [12]. Luciferase reporter assays suggest that *Rlim* is not a direct target of *miR-465*, whereas *Alkbh1* is directly repressed by *miR-465* (Fig. 2D). Consistently, *Alkbh1* was upregulated in the extraembryonic tissue of the “to-be-degenerating” KO females (Fig. 2B, Table S2).

**Figure 2.**
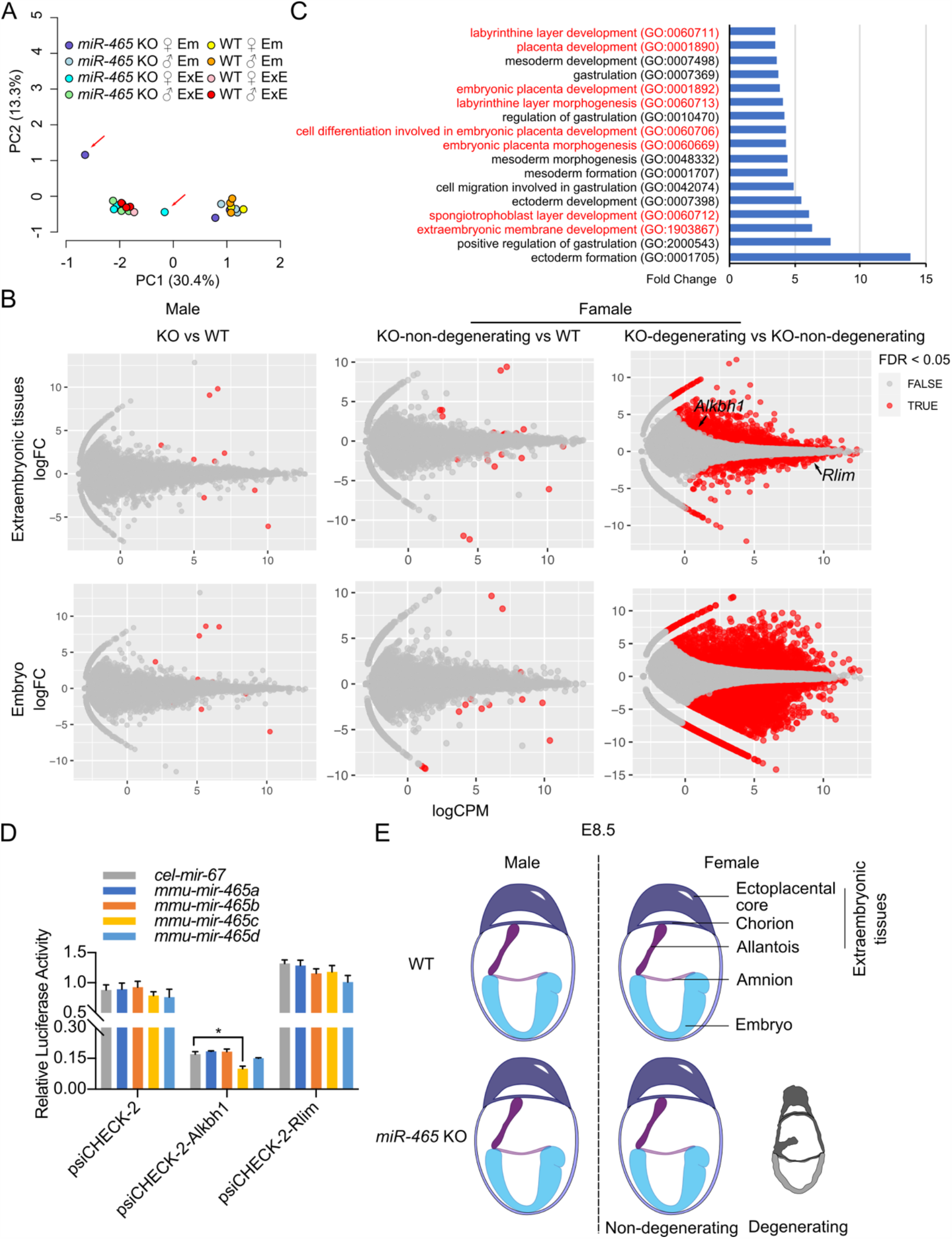
RNA-seq analyses of *miR-465* KO and WT conceptuses. A. Principal component analyses of RNA-seq data from embryonic (Em) and extraembryonic (ExE) tissues of *miR-465* KO and WT mice. The red arrows indicate the degenerating embryo and extraembryonic tissues from a *miR-465* KO female conceptus. B. Differentially expressed genes (DEGs) identified in the following three comparisons: between the *miR-465* KO and WT males (left), between the *miR-465* KO non-degenerating and WT females (middle), and between *miR-465* KO degenerating and non-degenerating females (right). C. GO term enrichment analyses of DEGs between the degenerating and non-degenerating *miR-465* KO female extraembryonic tissues. GO terms related to extraembryonic development are highlighted in red. D. Luciferase reporter assays for *Alkbh1* and *Rlim*. psiCHECK-2 is the empty vector used; the *cel-mir-67* served as a negative control, and firefly luciferase activity was used as the internal control. *, p<0.05. E. Schematics showing a critical role of the *miR-465* cluster in supporting full developmental potential of the female placenta and embryos.

## Conclusions

Taken together, our study uncovered an important role of the *miR-465* cluster in supporting full developmental potential of the female, but not the male, extraembryonic tissue/placenta (Fig. 2E). The male-biased sex ratio among *miR-465* KO mice results from selective degeneration of the female placenta and resorption of the female embryos in the absence of the *miR-465* cluster (Fig. 2E).

## Methods

### The aim, design and setting of the study

The aim of the present study is to reveal the physiological roles of the *miR-465* cluster by deleting the entire *miR-465* cluster in the mouse genome using CRISPR-Cas9. Different breeding pairs (+/+ × +/+, +/Y × −/−, −/Y × +/+, and −/Y × −/−) were set up to evaluate fertility and to analyze the sex ratio among offspring. To determine when the sex ratio bias commences, the KO breeding pairs (−/Y × −/−) were set up, and uteri at E3.5, E7.5, E8.5, E9.5 and E10.5 were collected for gross examination. Both embryos and extraembryonic/placental tissue were collected at E7.5 and/or E10.5 to assess miRNA expression profiles and cellular localization using sRNA-seq, qPCR, and RNA-ISH. Embryos and extraembryonic/placental tissue of both sexes at E7.5 were also used to perform RNA-seq and luciferase-based reporter assays to identify the target genes of the *miR-465* cluster miRNAs.

### Generation of global knockout mice and mouse genotyping

Generation of global KO mice and mouse genotyping were performed as described [2, 6, 7]. All gRNA and genotyping primers were listed in Table S3.

### DNA and RNA isolation, library construction and qPCR analyses

DNA and RNA were extracted from WT and KO embryos using the mirVana™ miRNA Isolation Kit as previously described [7]. The sexes of the conceptuses were determined based on PCR amplification of *DYzEms3* (a Y chromosome-specific repetitive sequence) and *Rn18s* (a house-keeping transcript as the internal control). Males display two bands (*DYzEms3 & Rn18s*), while females only show one band (*Rn18s*). Large RNA libraries were constructed using KAPA Stranded RNA-Seq Kits with RiboErase (Cat. # 07962282001, Roche) according to the manufacturer’s instructions. Small RNA libraries were constructed using NEBNext® Small RNA Library Prep Set for Illumina® (Cat. # E7330L, NEB) according to the manufacturer’s instructions. miRNA qPCR was performed as previously described [2]. All oligos for sex determination and qPCR were listed in Table S3.

### *In situ* hybridization

Cryosections (10 μm) were adhered to poly-L-lysine-coated slides and fixed in 4 % paraformaldehyde (Cat. # P6148, Sigma-Aldrich) solution in PBS for 1 h at room temperature. The sections were then washed 3 times in PBS for 5 min each, acetylated for 10 minutes (0.25% acetic anhydride), washed 2 times in PBS for 5 min each, and hybridized with DIG-labeled probes overnight at 50°C. Hybridization buffer contained 1×salts (200 mM NaCl, 13 mM Tris, 5 mM sodium phosphate monobasic, 5mM sodium phosphate dibasic, 5 mM EDTA), 50% formamide, 10% (w/v) dextran sulfate, 1 mg/ml yeast tRNA (Cat. # 10109509001, Roche), 1×Denhardt’s [1% (w/v) bovine serum albumin, 1% (w/v) Ficoll, 1% (w/v) polyvinylpyrrolidone], and RNA probe (final concentration: 1 μM). Post-hybridization washes were followed by an RNase treatment (20 μg/ml RNase A).

After blocking in 20% heat-inactivated sheep serum (Cat. # ZLI-9021, Beijing Zhongshan Jinqiao Biotechnology Company) and 2% blocking reagent (Cat. # 12039672910, Roche) for 1 h, sections were incubated overnight in blocking solution containing anti-DIG antibody (1:2500 dilution; Cat. # 11093274910, Roche) at room temperature. After washing, color was developed using NBT/BCIP according to the manufacturer’s instructions (NBT: Cat. # N1332, Gentihold; BCIP: Cat. # B1360, Gentihold). Sections were counterstained in Nuclear Fast Red (Cat. # G1321, Solarbio), dehydrated in gradient alcohol, cleared in xylene, and mounted in neutral resins. All oligos used for RNA ISH were listed in Table S3.

### RNA-Seq data analysis

The Sailfish [20] and SPORTS1.0 [21] pipelines were used to quantify the large RNA expression and small RNA expression, respectively. Transcript per million reads (TPM) was used as the unit of gene expression level. Groupwise differential expression was estimated by the likelihood ratio test and the RNAs with a false discovery rate < 5% were deemed differentially expressed.

### Luciferase assay

Luciferase assay was performed as previously described [22]. *cel-mir-67* was used as a negative control miRNA. *Renilla* luciferase signals were normalized to *Firefly* luciferase signal to correct the transfection efficiency. All oligos for constructing 3’UTR luciferase vectors were listed in Table S3.

### Statistical analyses

Data are presented as mean ± SEM, and statistical differences between datasets were assessed by two samples t-test unless stated otherwise. p < 0.05, 0.01, and 0.001 are considered as statistically significant and indicated with *, **, and ***, respectively.

## Abbreviations

WT: wild-type
KO: knockout
M: male
F: female
Em: embryo
ExE: extraembryonic tissue
Pl: placenta
Ch: chorion
EPC: ectoplacental core
Al: allantois
De: decidua
ISH: *in situ* hybridization
NS: not significant
N: normal-looking conceptuses
D: delayed/degenerating/degenerated conceptuses
DEGs: differentially expressed genes
PCA: principal component analyses
GO: Gene ontology
XCI: X chromosome inactivation
TPM: Transcript per million reads

## Declarations

### Ethics approval and consent to participate

The animal use protocol was approved by the Institutional Animal Care and Use Committee (IACUC) of the University of Nevada, Reno (protocol number 00494)

### Consent for publication

Not applicable

### Availability of data and materials

The datasets generated and/or analyzed during the current study are available in the Sequence Read Archive (SRA) under BioProject ID: PRJNA669325, https://www.ncbi.nlm.nih.gov/bioproject/PRJNA669325/

### Authors’ contributions

Z.W. and W.Y. designed the research. Z. W., N. M., Y. W., S. W., S.C., and H. Z. performed bench experiments. Z.W., T.Z., and M.L. performed bioinformatics analysis. S. K., and H.W., contributed reagents and protocols. Z.W., and W.Y. wrote the manuscript.

## Acknowledgements

Not applicable

