## Supplementary material for "A sexual dimorphic role of the *miR-465* cluster in placental development": Figure S1, S2, Table S1, S2, and S3

[1]Department of Physiology and Cell Biology, University of Nevada, Reno School of Medicine, Reno, NV 89557, USA; [2]Reproductive Medical Center, The First Affiliated Hospital of Xiamen University, 361003 Xiamen, Fujian, People's Republic of China; [3]Fujian Provincial Key Laboratory of Reproductive Health Research, School of Medicine Xiamen University, 361102 Xiamen, Fujian, People's Republic of China;[4]The Lundquist Institute for Biomedical Innovation at Harbor-UCLA Medical Center, Torrance, CA 90502, USA; [5]Department of Medicine, David Geffen School of Medicine at UCLA, Los Angeles, CA 90095, USA

This file includes Figure S1, S2, Table S1, S2, and S3.

Correspondence:

Wei Yan M.D., Ph.D.
*Investigator*
The Lundquist Institute at Harbor-UCLA Medical Center
*Professor of Medicine*
David Geffen School of Medicine at UCLA
1124 West Carson Street, Torrance, CA 90502


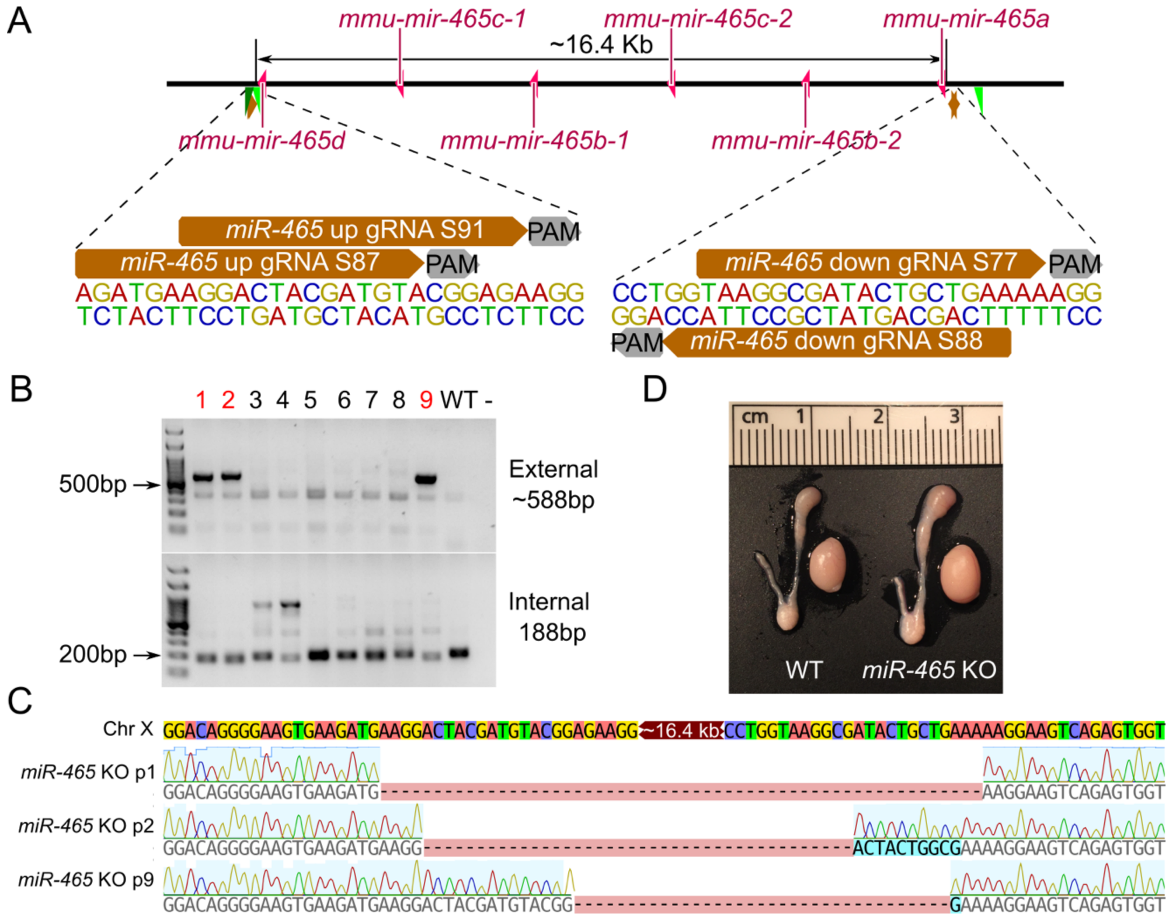

**Figure S1.** Generation of *miR-465* KO mice. A. Schematic illustration showing strategy used to generate the *miR-465* KO mice using CRISPR/Cas9. The brown triangle represents the gRNA used, and its right and left orientations indicate the gRNA targeting the reverse and forward strands of the genomic DNA, respectively. The green triangles show the position of primers for genotyping. B. A representative genotyping result of the *miR-465* KO founder mice. WT, wild-type. The expected size of PCR products is indicated on the right of the gel, the animal numbers in red represent the positive founders. C. Sanger sequencing of the founder mice. D. Normal morphology of WT and miR-465 KO testes and epididymides. One unit on the ruler is 1 mm.

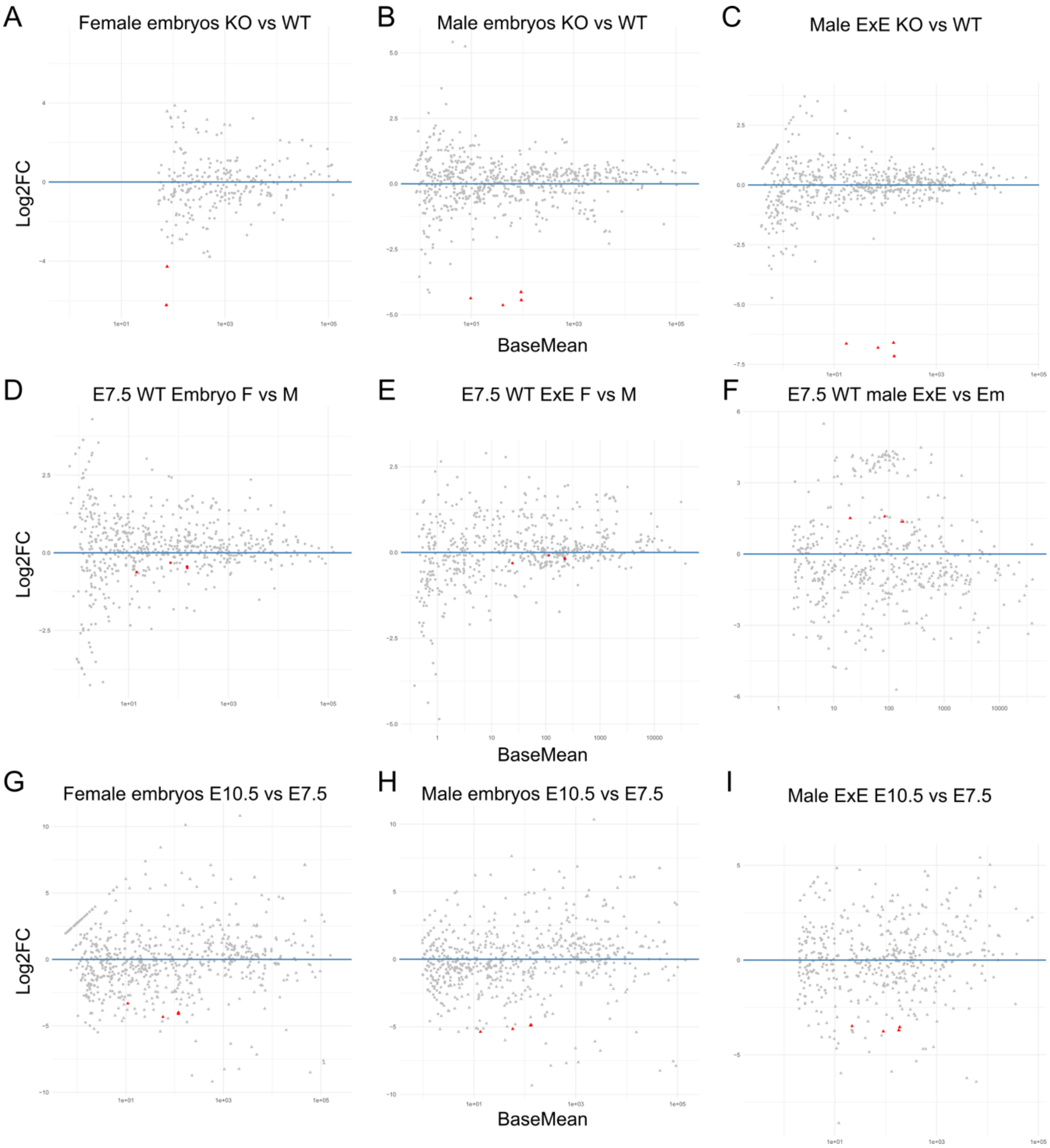
A
Female embryos KO vs WT
B
Male embryos KO vs WT
C
Male ExE KO vs WT
D
E7.5 WT Embryo F vs M
E
E7.5 WT ExE F vs M
F
E7.5 WT male ExE vs Em
G
Female embryos E10.5 vs E7.5
H
Male embryos E10.5 vs E7.5
I
Male ExE E10.5 vs E7.5
Log2FC
BaseMean

**Figure S2.** sRNA-seq of embryos and extraembryonic tissues/placenta in mice. A-C. Differentially expressed miRNAs between WT and KO female embryos (A), male embryos (B), and male extraembryonic tissues (C) at E7.5. D. Differentially expressed miRNAs between female and male embryos at E7.5. E. Differentially expressed miRNAs between female and male extraembryonic tissues in wild-type mice at E7.5. F. Differentially expressed miRNAs between extraembryonic tissues and embryos in wild-type males at E7.5. G-I. Differentially expressed miRNAs between E10.5 and E7.5 female embryos (G), male embryos (H), and male extraembryonic tissues (I). sRNA-seq analyses were conducted in biological triplicates (n=3), ExE, extraembryonic tissues. Triangle and circle indicate FDR < 0.05 and FDR > 0.05, respectively. Dots in red indicate the *miR-465* cluster miRNAs.

**Table S1. Sex ratio bias in literature**

| Sex ratio (male %) | Condition | Species | References (PMID) |
|---|---|---|---|
| 60% | *in vitro* fertilization | Mice | 26951653 |
| 67% | Very high in saturated fat | Mice | 12672968 |
| 39% | Low in saturated fat | Mice | 12672968 |
| 58%~76% | *Mcm2/4* mutation | Mice | 30787433 |
| 52%~59% | *Alkbh1* mutation | Mice | 21072209 |
| 60% | Excess fructose consumption | Rats | 23759309 |

**Table S2A. DEGs between KO and WT male extraembryonic tissues.**

| | logFC | logCPM | F | PValue | FDR < 0.05 | microrna.org | mirdb | mir wal | target scan |
|---|---|---|---|---|---|---|---|---|---|
| Rps2 | -6.0678791 | 10.0297078 | 3105.16874 | 7.24E-13 | 1.57E-08 | F | F | F | F |
| Gm10260 | 9.81099704 | 6.60960908 | 495.247971 | 2.93E-09 | 3.18E-05 | F | F | F | F |
| Rps18-ps5 | 9.07670214 | 6.05149944 | 292.324508 | 3.08E-08 | 0.0002227 | F | F | F | F |
| Dynlt1b | -2.7607997 | 5.68446508 | 251.363758 | 6.01E-08 | 0.0003257 | T | F | T | F |
| Nr1h5 | 3.30115277 | 2.77198039 | 102.201613 | 2.96E-06 | 0.0128228 | F | F | T | F |
| mt-Co3 | 2.37947752 | 7.06705344 | 76.5186418 | 9.88E-06 | 0.0357213 | F | F | F | F |
| Cd209c | 1.65766702 | 4.99016364 | 69.8782523 | 1.43E-05 | 0.0403404 | F | F | F | T |
| Gm8797 | 1.44748417 | 6.39873991 | 69.2430957 | 1.49E-05 | 0.0403404 | F | F | F | F |
| Hmga1 | -1.9067357 | 9.1236858 | 64.3352994 | 2.01E-05 | 0.0483282 | F | F | F | F |

**Table S2B. DEGs between KO and WT male embryos.**

| | logFC | logCPM | F | PValue | FDR < 0.05 | microrna.org | mirdb | mir walk | target scan |
|---|---|---|---|---|---|---|---|---|---|
| Rps2 | -5.9950039 | 10.2266789 | 3672.85048 | 5.15E-12 | 1.12E-07 | F | F | F | F |
| Nr1h5 | 7.28071904 | 5.16060307 | 502.453421 | 1.49E-08 | 0.00016178 | F | F | T | F |
| Dynlt1b | -2.9018182 | 5.31181792 | 356.814029 | 5.80E-08 | 0.00041376 | T | F | T | F |
| Gm10260 | 8.52603679 | 6.58215883 | 332.790888 | 7.63E-08 | 0.00041376 | F | F | F | F |
| Rps18-ps5 | 8.60233603 | 5.63103609 | 220.322441 | 3.85E-07 | 0.00167169 | F | F | F | F |
| Rims4 | -2.1394385 | 3.3383218 | 126.761418 | 3.26E-06 | 0.01178805 | F | F | F | F |
| Clec16a | 1.24548466 | 5.05736577 | 114.266995 | 4.84E-06 | 0.01324589 | T | F | T | T |
| Olfr1307 | 0.90066514 | 6.10279458 | 113.979923 | 4.89E-06 | 0.01324589 | F | F | F | F |
| Rps7 | -0.7565723 | 8.50255099 | 95.2572019 | 9.61E-06 | 0.0231634 | F | F | F | T |
| Bglap3 | 3.6828149 | 2.01732084 | 85.9668516 | 1.41E-05 | 0.03058378 | F | F | F | F |

**Table S2C. DEGs between non-degenerating and WT female extraembryonic tissues.**

| | logFC | logCPM | F | PValue | FDR < 0.05 | microrna.org | mirdb | mir walk | target scan |
|---|---|---|---|---|---|---|---|---|---|
| Rps2 | -6.0808554 | 10.1022545 | 2932.5274 | 8.06E-12 | 6.87E-08 | F | F | F | F |
| Rps18-ps5 | 8.93208344 | 6.63548849 | 2827.02832 | 9.37E-12 | 6.87E-08 | F | F | F | F |
| Gm10260 | 9.39403198 | 7.07038264 | 2817.10788 | 9.50E-12 | 6.87E-08 | F | F | F | F |
| Mid1 | -3.2146037 | 6.20188014 | 483.553473 | 1.29E-08 | 7.01E-05 | F | F | F | F |
| Srp54b | -12.433006 | 4.40569464 | 718.161756 | 1.62E-08 | 7.02E-05 | F | F | F | T |
| Rps18 | -2.146089 | 8.14415987 | 371.325209 | 3.77E-08 | 0.00013628 | F | F | F | F |
| Dynlt1b | -2.4322692 | 5.70717689 | 349.3203 | 4.83E-08 | 0.00014952 | T | F | T | F |
| Rps13 | 1.49053804 | 8.28635571 | 266.118226 | 1.44E-07 | 0.00039114 | F | F | F | F |
| Gm49450 | -11.989268 | 3.96363079 | 337.059727 | 2.42E-07 | 0.00058335 | F | F | F | F |
| Ube2l6 | -1.560661 | 5.62997025 | 133.949742 | 2.20E-06 | 0.00476039 | F | F | T | F |
| Gsn | 0.97061214 | 7.75998606 | 115.757636 | 3.87E-06 | 0.00763832 | F | F | F | F |
| Gapdh | -0.6653078 | 11.1230064 | 109.612782 | 4.79E-06 | 0.00864726 | F | F | T | F |
| Hist1h2br | -1.694337 | 6.71295442 | 85.9792698 | 1.21E-05 | 0.02024081 | F | F | T | T |
| Rsad2 | 3.13553293 | 2.45262235 | 81.7031565 | 1.47E-05 | 0.02280402 | T | F | T | T |
| Rpl10l | 3.92138704 | 2.42664819 | 75.8285584 | 1.95E-05 | 0.02819884 | F | F | F | F |
| Gm8797 | 1.072867 | 6.13976258 | 73.5611673 | 2.19E-05 | 0.02962688 | F | F | F | F |
| Syna | 3.92761994 | 2.27986736 | 70.7050936 | 2.54E-05 | 0.03233707 | F | F | F | T |
| Epas1 | 0.81031549 | 6.82697378 | 65.6516656 | 3.34E-05 | 0.04024589 | F | F | F | F |
| H2-K1 | -2.3237766 | 3.53178428 | 60.9779198 | 4.39E-05 | 0.04792843 | F | F | F | F |
| Ide | 0.92894938 | 6.93957151 | 60.8576536 | 4.42E-05 | 0.04792843 | F | F | T | F |

**Table S2D. DEGs between non-degenerating and WT female embryos.**

| | logFC | logCPM | F | PValue | FDR < 0.05 | micror na.org | mirdb | mir walk | target scan |
|---|---|---|---|---|---|---|---|---|---|
| Rps2 | -6.1966089 | 10.4002944 | 2065.58828 | 1.30E-12 | 2.82E-08 | F | F | F | F |
| Gm10260 | 8.23278729 | 6.93554358 | 782.96622 | 1.38E-10 | 1.49E-06 | F | F | F | F |
| Rps18-ps5 | 9.63483544 | 6.133021 | 443.998628 | 2.06E-09 | 1.49E-05 | F | F | F | F |
| Rps18 | -1.7005495 | 8.37884887 | 143.340296 | 4.04E-07 | 0.00218996 | F | F | F | F |
| Dynlt1b | -2.7164773 | 5.41202016 | 133.152003 | 5.65E-07 | 0.0024504 | T | F | T | F |
| Gfer | -2.2790495 | 4.22910723 | 100.196392 | 2.03E-06 | 0.00733555 | T | F | T | T |
| Rims4 | -3.0332973 | 3.75255142 | 86.452886 | 3.90E-06 | 0.0120745 | F | F | F | F |
| Rps13 | 1.26803262 | 8.26263005 | 83.3474832 | 4.58E-06 | 0.0124043 | F | F | F | F |
| A2m | -2.2949638 | 5.95951538 | 74.2431072 | 7.57E-06 | 0.01824676 | F | F | F | F |
| Dagla | -9.2772371 | 1.2938512 | 81.1693604 | 1.08E-05 | 0.023365 | F | F | T | F |
| Gm8797 | 1.07584898 | 6.39465719 | 63.1828921 | 1.51E-05 | 0.0298306 | F | F | F | F |
| Hmga1 | -2.0721869 | 9.85899269 | 61.2114271 | 1.73E-05 | 0.0300417 | F | F | F | F |
| H2-T22 | -9.2042148 | 1.22379593 | 71.2393406 | 1.80E-05 | 0.0300417 | F | F | F | F |
| Gm12666 | -9.0196033 | 1.04888597 | 67.9920793 | 2.16E-05 | 0.03152814 | F | F | F | F |
| Wfdc2 | -1.6622952 | 4.62215939 | 57.9513857 | 2.18E-05 | 0.03152814 | F | F | F | F |

**Table S2E. DEGs between degenerating and non-degenerating female extraembryonic tissues.**

| | logFC | logCPM | F | PValue | FDR < 0.05 | microrna.org | mirdb | mir walk | target scan |
|---|---|---|---|---|---|---|---|---|---|
| Lefty2 | 8.97531835 | 4.70513448 | 770.889969 | 1.86E-23 | 4.02E-19 | T | F | T | F |
| Trh | 6.66372422 | 5.43681229 | 713.083327 | 5.92E-23 | 6.42E-19 | F | F | F | F |
| Cer1 | 7.33353205 | 4.5473825 | 641.587547 | 2.84E-22 | 2.05E-18 | T | F | T | T |
| Pim2 | 3.52570075 | 6.17537783 | 518.692814 | 6.51E-21 | 3.53E-17 | F | F | F | F |
| Pcdh19 | 4.94154224 | 5.1123767 | 485.283247 | 1.72E-20 | 7.47E-17 | T | F | T | T |
| Tenm4 | 3.60616792 | 5.88975181 | 463.548356 | 3.36E-20 | 1.21E-16 | T | F | T | T |
| Sfrp1 | 5.23210449 | 4.69539891 | 446.530764 | 5.78E-20 | 1.79E-16 | F | F | T | T |
| Gm49450 | 12.3982617 | 3.71736615 | 442.029044 | 1.68E-19 | 4.55E-16 | F | F | F | F |
| Dll1 | 4.15190087 | 4.78056433 | 383.126406 | 5.29E-19 | 1.27E-15 | F | F | F | F |
| Cachd1 | 2.99219998 | 6.04789944 | 369.859057 | 8.76E-19 | 1.90E-15 | T | F | F | F |
| T | 4.24989367 | 6.43984104 | 356.913316 | 1.46E-18 | 2.88E-15 | F | F | F | F |
| Pdzd4 | 3.8068071 | 4.89169731 | 344.441715 | 2.43E-18 | 4.38E-15 | F | F | F | F |
| Otx2 | 7.99944768 | 3.44158091 | 337.951729 | 3.18E-18 | 5.30E-15 | T | F | T | T |
| Gjb3 | -2.0318858 | 8.62623862 | 333.202717 | 3.89E-18 | 6.02E-15 | F | F | T | F |
| Fzd2 | 3.50415853 | 5.19282715 | 329.71077 | 4.52E-18 | 6.53E-15 | F | F | F | F |
| Pou5f1 | 5.09202158 | 6.44060026 | 326.051166 | 5.29E-18 | 7.17E-15 | T | F | F | F |
| Pfpl | 2.77752941 | 7.68359412 | 315.987036 | 8.25E-18 | 1.05E-14 | F | F | F | F |
| Gjb5 | -2.4175638 | 7.40462763 | 314.153404 | 8.96E-18 | 1.08E-14 | T | F | F | T |
| Fst | 3.71513094 | 6.15478939 | 311.324368 | 1.02E-17 | 1.16E-14 | T | F | T | T |
| Pcdh8 | 6.80767683 | 3.52358387 | 309.030907 | 1.13E-17 | 1.23E-14 | F | F | F | F |
| Mmp9 | 2.18364349 | 7.12222786 | 305.737108 | 1.32E-17 | 1.36E-14 | F | F | F | F |
| Gm28635 | 11.8610194 | 3.18807973 | 316.4927 | 1.75E-17 | 1.73E-14 | F | F | F | F |
| Col4a1 | 1.50686206 | 11.2764183 | 283.314028 | 3.84E-17 | 3.47E-14 | T | F | T | T |
| Dpysl5 | 3.37905996 | 4.8794738 | 283.270313 | 3.85E-17 | 3.47E-14 | F | F | T | F |
| Pla2g2f | -5.4800817 | 4.67837859 | 282.477215 | 4.00E-17 | 3.47E-14 | F | F | F | F |
| Ascl2 | -2.5278258 | 6.76913907 | 274.365369 | 6.01E-17 | 4.97E-14 | F | F | T | F |
| Slc7a3 | 5.82673931 | 3.93293241 | 273.791819 | 6.19E-17 | 4.97E-14 | F | F | F | F |
| Dll3 | 4.72078541 | 3.95959356 | 264.511958 | 1.00E-16 | 7.76E-14 | F | F | F | F |
| Foxj1 | 4.80175168 | 3.76647722 | 254.239587 | 1.74E-16 | 1.30E-13 | T | F | T | F |
| Zic3 | 4.85619662 | 4.45914617 | 250.722238 | 2.11E-16 | 1.44E-13 | T | F | T | T |
| Sox11 | 2.89536144 | 6.28233179 | 250.582375 | 2.12E-16 | 1.44E-13 | T | F | T | T |
| Peg10 | -1.2278783 | 11.8835244 | 250.520257 | 2.13E-16 | 1.44E-13 | T | F | F | T |
| Sp5 | 4.60883219 | 4.07865005 | 243.805719 | 3.10E-16 | 2.04E-13 | F | F | F | F |
| Ror1 | 4.63812219 | 3.76517659 | 243.062081 | 3.23E-16 | 2.06E-13 | F | F | F | F |
| Jph4 | 6.56699418 | 3.15093067 | 242.580035 | 3.32E-16 | 2.06E-13 | T | F | F | F |
| Dnmt3b | 1.30409854 | 9.76040577 | 241.551738 | 3.52E-16 | 2.12E-13 | F | F | F | F |
| Lamb1 | 1.45877338 | 9.5856001 | 232.31193 | 6.03E-16 | 3.53E-13 | F | F | F | T |
| Sp8 | 4.52457116 | 3.74811978 | 231.591352 | 6.29E-16 | 3.59E-13 | F | F | T | F |
| Sox4 | 2.47650833 | 5.71079315 | 229.060848 | 7.31E-16 | 4.06E-13 | F | F | T | T |

| | | | | | | | | | |
|---|---|---|---|---|---|---|---|---|---|
| Ror2 | 2.98678851 | 4.8626092 | 228.167133 | 7.71E-16 | 4.18E-13 | T | F | T | F |
| Gpc4 | 3.24093391 | 4.56914634 | 227.264378 | 8.14E-16 | 4.31E-13 | F | F | F | T |
| Kif1a | 4.76009455 | 3.59482719 | 221.845047 | 1.13E-15 | 5.85E-13 | F | F | F | F |
| Crmp1 | 4.59698051 | 3.9078907 | 220.321083 | 1.24E-15 | 6.27E-13 | F | F | F | F |
| Prune2 | -2.0429069 | 7.05465173 | 219.047142 | 1.35E-15 | 6.64E-13 | T | F | T | F |
| Rgma | 3.63247845 | 4.36612678 | 217.318788 | 1.50E-15 | 7.23E-13 | T | F | T | T |
| Msi1 | 2.99466948 | 5.45716861 | 216.951842 | 1.54E-15 | 7.24E-13 | F | F | F | F |
| Robo1 | 2.83835239 | 5.8065183 | 214.266688 | 1.82E-15 | 8.39E-13 | T | F | F | T |
| Prl3d1 | 1.88911479 | 7.18297878 | 212.408857 | 2.05E-15 | 9.12E-13 | F | F | F | F |
| Kif26a | -1.7101243 | 7.89071004 | 212.308017 | 2.06E-15 | 9.12E-13 | F | F | F | F |
| Tenm3 | 2.81095583 | 5.23978058 | 209.835201 | 2.42E-15 | 1.05E-12 | T | F | T | T |
| Sulf1 | 2.55746783 | 5.45414712 | 207.972477 | 2.73E-15 | 1.16E-12 | T | F | T | T |
| Gm42742 | -12.125731 | 4.42995848 | 204.843429 | 3.35E-15 | 1.40E-12 | F | F | F | F |
| Krt19 | -1.7302885 | 8.57331767 | 201.159418 | 4.28E-15 | 1.74E-12 | F | F | T | F |
| Sall2 | 2.64727816 | 5.09533992 | 200.946406 | 4.34E-15 | 1.74E-12 | F | F | T | F |
| Phc1 | 2.09771692 | 6.09985314 | 199.121004 | 4.91E-15 | 1.93E-12 | F | F | F | F |
| Rassf4 | -3.7577877 | 4.83079146 | 198.084639 | 5.26E-15 | 2.04E-12 | T | F | T | T |
| Lgals1 | -1.7862766 | 7.97624921 | 196.924594 | 5.69E-15 | 2.17E-12 | F | F | T | F |
| Tdgf1 | 5.96557583 | 3.06803175 | 195.023969 | 6.49E-15 | 2.43E-12 | T | F | T | F |
| Lama1 | 1.82573202 | 10.0874725 | 194.341329 | 6.80E-15 | 2.50E-12 | F | F | F | F |
| Col4a2 | 1.34514234 | 10.39473 | 192.146337 | 7.92E-15 | 2.86E-12 | F | F | T | F |
| Dlx3 | -2.7579252 | 5.69261635 | 191.175391 | 8.48E-15 | 2.99E-12 | T | F | T | T |
| Lamc1 | 1.58271292 | 10.1092111 | 190.903468 | 8.64E-15 | 2.99E-12 | F | F | T | F |
| Crispld2 | -4.1263601 | 4.52363436 | 190.814037 | 8.69E-15 | 2.99E-12 | F | F | T | F |
| Notch3 | 2.73288261 | 4.82643219 | 190.191014 | 9.08E-15 | 3.08E-12 | F | F | F | F |
| Tspan7 | 3.81795957 | 3.93763159 | 189.089918 | 9.82E-15 | 3.28E-12 | T | T | T | T |
| Tfap2c | -1.5244845 | 8.53323337 | 188.272911 | 1.04E-14 | 3.42E-12 | T | F | F | F |
| Fzd7 | 1.75555991 | 6.66310359 | 181.432534 | 1.70E-14 | 5.52E-12 | T | F | T | T |
| Wnt3a | 6.08486288 | 2.82539786 | 181.077758 | 1.75E-14 | 5.58E-12 | T | F | T | F |
| Zscan10 | 2.61244395 | 4.98321929 | 180.559899 | 1.82E-14 | 5.59E-12 | F | F | F | F |
| Irx5 | 6.48795961 | 2.72247105 | 180.44092 | 1.83E-14 | 5.59E-12 | F | F | F | F |
| Fn1 | 1.07473409 | 10.2557 | 180.360212 | 1.84E-14 | 5.59E-12 | T | F | F | F |
| Cxcl12 | 4.84928557 | 3.79248724 | 180.27587 | 1.86E-14 | 5.59E-12 | T | F | T | T |
| Nfasc | 5.93551294 | 2.81979095 | 179.866774 | 1.91E-14 | 5.68E-12 | F | F | T | F |
| Sbk1 | 2.73597066 | 4.87299566 | 178.89788 | 2.05E-14 | 6.02E-12 | T | F | T | T |
| Ldoc1 | -2.0153297 | 6.96464182 | 177.699712 | 2.25E-14 | 6.49E-12 | T | F | F | F |
| Nptx2 | 3.55422496 | 4.13401774 | 177.433918 | 2.29E-14 | 6.54E-12 | F | F | T | F |
| Palm | -1.6683411 | 7.72605041 | 176.881515 | 2.39E-14 | 6.72E-12 | F | F | F | F |
| Foxc1 | 4.84564473 | 3.15022748 | 173.47457 | 3.09E-14 | 8.59E-12 | F | F | T | F |
| Ina | 4.95564697 | 3.45106588 | 172.921432 | 3.22E-14 | 8.76E-12 | T | F | T | F |
| Rimbp2 | 5.2017891 | 2.9773164 | 172.883167 | 3.23E-14 | 8.76E-12 | T | F | T | F |
| Car14 | 2.20475839 | 5.49147107 | 169.402858 | 4.23E-14 | 1.13E-11 | T | F | F | F |

| | | | | | | | | | |
|---|---|---|---|---|---|---|---|---|---|
| Zic2 | 2.66249961 | 5.555095 | 165.560141 | 5.71E-14 | 1.51E-11 | F | F | F | F |
| Gas1 | 3.38433033 | 3.92985681 | 165.18725 | 5.89E-14 | 1.54E-11 | T | F | T | T |
| Flna | 1.40346513 | 9.09250219 | 164.806778 | 6.07E-14 | 1.57E-11 | F | F | F | F |
| Foxi3 | 8.627261 | 2.31257821 | 160.391617 | 8.65E-14 | 2.21E-11 | F | F | F | F |
| Fstl3 | -1.7016099 | 7.03938274 | 159.949118 | 8.97E-14 | 2.26E-11 | F | F | F | F |
| Ero1l | -1.6763173 | 7.78776342 | 159.813362 | 9.07E-14 | 2.26E-11 | F | F | T | F |
| Tubb4a | 4.5925666 | 3.11185319 | 159.338873 | 9.43E-14 | 2.32E-11 | F | F | T | F |
| Nectin1 | 1.85637055 | 6.35543751 | 158.83372 | 9.83E-14 | 2.40E-11 | T | F | T | T |
| Entpd1 | -1.6817449 | 7.32991092 | 155.075032 | 1.34E-13 | 3.22E-11 | T | T | T | T |
| Itgb4 | -1.9346151 | 6.51566669 | 154.943033 | 1.36E-13 | 3.22E-11 | F | F | F | F |
| Foxb1 | 6.6056587 | 2.487356 | 154.849231 | 1.37E-13 | 3.22E-11 | F | T | T | T |
| Serpinb9e | -2.479964 | 5.59785369 | 154.181621 | 1.45E-13 | 3.37E-11 | T | T | F | T |
| Frzb | 3.04372631 | 4.11500585 | 151.142695 | 1.87E-13 | 4.32E-11 | T | F | T | T |
| Klhl13 | -1.5269595 | 7.67674246 | 150.799766 | 1.93E-13 | 4.40E-11 | T | F | T | T |
| Nrk | -1.541039 | 8.89516328 | 150.49177 | 1.98E-13 | 4.47E-11 | T | F | T | T |
| AU018091 | -1.5966868 | 7.54333201 | 150.219499 | 2.03E-13 | 4.53E-11 | F | F | F | F |
| Dbn1 | 2.29281805 | 5.94665914 | 148.892637 | 2.27E-13 | 5.03E-11 | F | F | F | F |
| Aplnr | 1.87287373 | 5.93360172 | 148.506775 | 2.35E-13 | 5.15E-11 | T | F | T | T |
| Cdh2 | 2.96790404 | 5.18120424 | 147.41048 | 2.59E-13 | 5.61E-11 | T | F | F | T |
| Pxdn | 1.24048487 | 8.06333008 | 146.784633 | 2.73E-13 | 5.87E-11 | T | F | F | F |
| Lrp1 | 1.24817031 | 8.02513594 | 146.60369 | 2.78E-13 | 5.90E-11 | T | T | F | T |
| Lgals9 | -2.0378006 | 6.19161791 | 146.40132 | 2.83E-13 | 5.95E-11 | F | F | F | T |
| Sct | -3.3646891 | 4.49793126 | 144.961876 | 3.21E-13 | 6.69E-11 | F | F | F | F |
| Ptk7 | 1.45103077 | 7.67300008 | 144.343719 | 3.39E-13 | 7.00E-11 | F | F | F | F |
| Igfbp2 | 2.32013908 | 5.6490437 | 142.699951 | 3.93E-13 | 8.03E-11 | F | F | F | F |
| Prtg | 1.24370453 | 8.84417486 | 141.672489 | 4.31E-13 | 8.73E-11 | T | F | T | F |
| Usp44 | 4.18649279 | 3.25482065 | 140.80983 | 4.66E-13 | 9.34E-11 | T | F | T | T |
| H2bu2 | 2.80097186 | 4.35258153 | 140.719215 | 4.70E-13 | 9.34E-11 | F | F | F | F |
| Frem2 | 1.63046931 | 6.42036627 | 140.413841 | 4.83E-13 | 9.52E-11 | T | T | F | T |
| Armcx2 | 2.27855035 | 5.19571932 | 139.890545 | 5.06E-13 | 9.89E-11 | F | F | T | F |
| Celsr2 | 2.95488417 | 4.05043365 | 139.541546 | 5.23E-13 | 1.01E-10 | F | F | F | F |
| Mbnl3 | -2.6240866 | 5.33347729 | 139.287817 | 5.35E-13 | 1.03E-10 | T | T | T | T |
| Fgfr1 | 1.3886983 | 7.08591854 | 138.827325 | 5.58E-13 | 1.06E-10 | F | F | F | F |
| Zc3h7b | 1.60643411 | 6.44375515 | 138.615403 | 5.69E-13 | 1.07E-10 | T | F | T | T |
| Foxc2 | 4.93846274 | 3.05591704 | 138.491754 | 5.76E-13 | 1.08E-10 | F | F | F | F |
| Rragd | 2.34833854 | 4.79452988 | 137.726717 | 6.18E-13 | 1.14E-10 | T | F | T | T |
| Cts7 | 2.30786849 | 5.17737587 | 137.649573 | 6.22E-13 | 1.14E-10 | F | F | F | F |
| Fthl17a | -2.8265419 | 4.87583894 | 137.376807 | 6.38E-13 | 1.16E-10 | F | F | F | F |
| Lef1 | 2.39290053 | 5.31634231 | 135.861607 | 7.35E-13 | 1.33E-10 | T | F | F | F |
| Col2a1 | 5.05170215 | 2.65719481 | 134.949763 | 8.00E-13 | 1.43E-10 | T | F | T | F |
| Map1a | 2.59724022 | 4.37602889 | 134.542406 | 8.32E-13 | 1.48E-10 | T | F | F | F |
| Dsp | -1.0942246 | 9.48909405 | 134.138486 | 8.64E-13 | 1.52E-10 | F | F | F | T |

| | | | | | | | | | |
|---|---|---|---|---|---|---|---|---|---|
| Tns1 | -2.2581011 | 5.70190097 | 133.905367 | 8.83E-13 | 1.54E-10 | T | T | T | T |
| Tmem132a | 1.56966297 | 6.46432852 | 133.631897 | 9.06E-13 | 1.57E-10 | F | F | F | F |
| Mrc2 | 2.64025356 | 4.27792723 | 132.616714 | 9.98E-13 | 1.72E-10 | F | F | F | F |
| Gata2 | -1.2833956 | 7.98713576 | 130.041656 | 1.28E-12 | 2.18E-10 | F | F | F | F |
| Tpm2 | -1.8711886 | 6.27428353 | 129.984449 | 1.29E-12 | 2.18E-10 | F | F | T | F |
| Cdkn1c | -1.2293939 | 9.11072682 | 129.503624 | 1.35E-12 | 2.26E-10 | T | T | F | T |
| Wnt2b | 3.52587987 | 3.62120565 | 128.519186 | 1.48E-12 | 2.47E-10 | T | F | T | T |
| Plac1 | -1.4268771 | 7.44374353 | 128.290915 | 1.52E-12 | 2.51E-10 | T | F | F | F |
| Arid3a | -1.2644466 | 8.06854838 | 127.648401 | 1.61E-12 | 2.65E-10 | F | F | T | F |
| Slc6a2 | 3.12966212 | 3.74832913 | 127.424601 | 1.65E-12 | 2.69E-10 | F | F | T | F |
| Serpinb9g | -2.6504253 | 4.93688955 | 126.921238 | 1.74E-12 | 2.81E-10 | T | T | F | T |
| Zbtb10 | -1.3806954 | 7.9903207 | 126.397276 | 1.83E-12 | 2.94E-10 | T | F | T | T |
| Smo | 1.48008387 | 6.56251695 | 124.156348 | 2.29E-12 | 3.64E-10 | F | F | F | F |
| Notch1 | 1.81195616 | 5.96520095 | 124.037457 | 2.31E-12 | 3.66E-10 | T | F | F | F |
| Vrtn | 2.54399384 | 5.53350229 | 123.961489 | 2.33E-12 | 3.66E-10 | F | F | F | F |
| Armcx4 | 2.00756907 | 5.57749424 | 122.664903 | 2.66E-12 | 4.15E-10 | F | F | F | F |
| Fes | -2.7810182 | 4.70935434 | 121.382293 | 3.03E-12 | 4.69E-10 | F | F | F | F |
| Arnt2 | 3.95043619 | 3.10841242 | 120.968304 | 3.16E-12 | 4.86E-10 | T | F | T | F |
| Epha1 | 1.66286434 | 5.96603125 | 119.539356 | 3.66E-12 | 5.59E-10 | F | F | F | F |
| Fads2 | 1.86483401 | 5.43698569 | 118.520392 | 4.07E-12 | 6.18E-10 | T | F | T | F |
| Dclk1 | 5.40961145 | 2.32399548 | 116.796311 | 4.88E-12 | 7.35E-10 | T | F | T | T |
| Tead2 | 1.26517989 | 7.42344935 | 116.471945 | 5.05E-12 | 7.56E-10 | F | F | T | F |
| Kdm5b | 1.15198167 | 7.72575495 | 116.397728 | 5.09E-12 | 7.56E-10 | T | F | T | T |
| Grik3 | 2.19415774 | 4.98329233 | 115.65302 | 5.51E-12 | 8.13E-10 | F | F | T | T |
| Sez6l2 | -2.4743837 | 5.10818337 | 115.308714 | 5.72E-12 | 8.38E-10 | F | F | F | F |
| Mcf2 | 6.90967394 | 2.0188252 | 115.119258 | 5.83E-12 | 8.45E-10 | T | F | F | F |
| Nmnat2 | 3.76474801 | 3.07777235 | 115.067008 | 5.87E-12 | 8.45E-10 | T | F | F | T |
| Pbx1 | 1.89287418 | 6.03911721 | 115.040204 | 5.88E-12 | 8.45E-10 | T | F | T | T |
| Pde4dip | -1.3543782 | 7.49493295 | 114.854875 | 6.00E-12 | 8.56E-10 | T | F | T | F |
| Nrxn2 | 5.04937963 | 2.63309623 | 114.731018 | 6.08E-12 | 8.59E-10 | F | F | T | T |
| Trp53i11 | 2.59643566 | 4.11983658 | 114.700419 | 6.10E-12 | 8.59E-10 | T | F | T | T |
| Adm | 1.35361618 | 8.77429412 | 114.515218 | 6.22E-12 | 8.71E-10 | T | F | T | F |
| Arhgef26 | 3.5127289 | 3.25617907 | 114.399086 | 6.30E-12 | 8.76E-10 | F | F | F | F |
| C1qtnf6 | -2.7353224 | 4.60525618 | 113.866917 | 6.67E-12 | 9.22E-10 | T | F | T | T |
| Satb1 | -1.3589033 | 7.64730167 | 113.524626 | 6.93E-12 | 9.41E-10 | F | T | T | T |
| Mex3a | 1.70178527 | 6.09840436 | 113.518047 | 6.93E-12 | 9.41E-10 | T | F | T | T |
| Dact1 | 3.57755843 | 3.20467255 | 113.503902 | 6.94E-12 | 9.41E-10 | T | F | T | F |
| Sfmbt2 | -1.0217369 | 9.44296089 | 112.958261 | 7.36E-12 | 9.92E-10 | T | F | T | T |
| Tgif1 | 2.03344522 | 4.92405317 | 112.671027 | 7.60E-12 | 1.02E-09 | T | F | T | F |
| Tbx6 | 3.6791601 | 3.11243911 | 112.032625 | 8.14E-12 | 1.08E-09 | F | F | F | F |
| Sin3b | -1.2567022 | 8.00704224 | 111.417272 | 8.71E-12 | 1.15E-09 | F | F | F | F |
| Serpinh1 | 0.95843917 | 9.04008616 | 110.762409 | 9.36E-12 | 1.22E-09 | F | F | T | F |

| | | | | | | | | | |
|---|---|---|---|---|---|---|---|---|---|
| Hand1 | -1.1897103 | 8.84422854 | 110.756496 | 9.37E-12 | 1.22E-09 | T | T | F | T |
| Sox3 | 3.80282801 | 3.23959343 | 109.418918 | 1.09E-11 | 1.41E-09 | F | F | F | F |
| Nkx1-2 | 2.61985849 | 4.66431767 | 108.278163 | 1.23E-11 | 1.59E-09 | T | F | T | F |
| Col23a1 | 3.93303779 | 2.85994074 | 107.76386 | 1.31E-11 | 1.68E-09 | T | F | T | T |
| Ccn2 | 2.49346812 | 4.32020737 | 107.575137 | 1.34E-11 | 1.70E-09 | T | F | T | F |
| Zdbf2 | 2.1477712 | 5.03838449 | 107.52123 | 1.34E-11 | 1.70E-09 | F | T | F | T |
| Arpc1b | -1.2317803 | 7.73549544 | 107.257854 | 1.38E-11 | 1.75E-09 | T | F | F | T |
| Ralbp1 | -1.2651902 | 7.55548197 | 107.061548 | 1.42E-11 | 1.77E-09 | F | F | T | F |
| Ccnd3 | -1.0962389 | 8.4464695 | 106.438291 | 1.52E-11 | 1.89E-09 | T | F | T | F |
| Stox2 | 2.73483417 | 3.97191959 | 105.943611 | 1.61E-11 | 1.99E-09 | T | F | T | T |
| Bahcc1 | 1.42776613 | 6.40800071 | 105.771678 | 1.64E-11 | 2.02E-09 | T | F | F | T |
| Rlim | -1.168757 | 9.19914555 | 104.983806 | 1.79E-11 | 2.20E-09 | T | F | T | F |
| Celsr3 | 2.96729873 | 3.60342286 | 103.847433 | 2.05E-11 | 2.49E-09 | F | F | T | F |
| Clip2 | -1.2477576 | 7.615097 | 103.434693 | 2.15E-11 | 2.59E-09 | F | F | T | F |
| Lpl | -2.2323237 | 5.654877 | 103.419853 | 2.15E-11 | 2.59E-09 | T | F | F | F |
| Wnt8a | 5.49356853 | 2.14791778 | 103.059503 | 2.24E-11 | 2.69E-09 | F | F | F | F |
| Dapk1 | 1.55354593 | 5.93982362 | 102.919223 | 2.28E-11 | 2.72E-09 | T | F | F | T |
| Chsy1 | -1.2651067 | 7.33968673 | 102.706307 | 2.34E-11 | 2.77E-09 | T | F | T | F |
| Nnat | 3.24977551 | 3.59289063 | 102.591064 | 2.37E-11 | 2.79E-09 | F | F | F | F |
| Prl8a9 | 3.89088663 | 3.03550251 | 102.051938 | 2.52E-11 | 2.96E-09 | T | F | F | F |
| Astn1 | 6.07858075 | 2.16500369 | 101.85651 | 2.58E-11 | 3.01E-09 | T | F | T | T |
| Vash2 | 2.8193088 | 3.68015307 | 101.049797 | 2.84E-11 | 3.29E-09 | F | F | T | F |
| Gja1 | 1.04289378 | 7.96796457 | 100.704789 | 2.96E-11 | 3.41E-09 | F | F | F | F |
| Syna | -10.247479 | 2.57034418 | 100.217093 | 3.13E-11 | 3.60E-09 | F | F | F | T |
| Elf4 | -1.4647542 | 6.64637354 | 100.169485 | 3.15E-11 | 3.60E-09 | T | F | T | T |
| Pla2g5 | 2.4522019 | 4.36665434 | 99.9988408 | 3.22E-11 | 3.65E-09 | T | T | T | T |
| Liph | -2.7897347 | 4.32234641 | 99.8475347 | 3.28E-11 | 3.70E-09 | T | F | T | T |
| Pls3 | 1.73262767 | 5.55640526 | 99.7087556 | 3.33E-11 | 3.74E-09 | T | F | F | T |
| Zfp266 | 1.46028353 | 6.330304 | 99.5830708 | 3.38E-11 | 3.78E-09 | F | F | T | T |
| Krt8 | -0.9117665 | 9.96702513 | 99.2121391 | 3.54E-11 | 3.93E-09 | F | F | F | F |
| Ptp4a3 | -2.0240711 | 5.35731679 | 98.3803821 | 3.91E-11 | 4.32E-09 | F | F | T | T |
| Pcdh12 | -3.7596947 | 3.82927941 | 98.2433823 | 3.97E-11 | 4.37E-09 | F | F | F | T |
| Slc38a4 | -1.0830967 | 8.17521809 | 98.166016 | 4.01E-11 | 4.39E-09 | F | F | T | T |
| Tnfrsf19 | 1.99620869 | 5.30215487 | 98.0137135 | 4.09E-11 | 4.45E-09 | T | F | F | T |
| Tmem132e | -3.0215984 | 4.15752176 | 97.6731137 | 4.26E-11 | 4.61E-09 | T | F | F | F |
| Sort1 | 1.7374457 | 5.33986536 | 97.6481523 | 4.27E-11 | 4.61E-09 | T | F | T | T |
| Aff1 | 1.83031868 | 5.09696381 | 96.7369721 | 4.77E-11 | 5.12E-09 | T | F | F | T |
| Nedd4 | 0.80882801 | 10.063965 | 96.6133187 | 4.85E-11 | 5.18E-09 | T | F | T | T |
| Car2 | -1.3235931 | 7.3351914 | 96.1485856 | 5.13E-11 | 5.45E-09 | T | F | F | F |
| Mrps24 | -2.9067427 | 4.27896288 | 95.843821 | 5.33E-11 | 5.62E-09 | T | F | T | T |
| Chdh | -2.027957 | 5.31465474 | 95.8180261 | 5.34E-11 | 5.62E-09 | T | T | F | T |
| Adap1 | -1.7925376 | 5.74647319 | 95.3717874 | 5.65E-11 | 5.90E-09 | T | F | F | T |

| | | | | | | | | | |
|---|---|---|---|---|---|---|---|---|---|
| Ppard | -1.2592354 | 7.50010877 | 95.3559162 | 5.66E-11 | 5.90E-09 | F | F | F | T |
| AI661453 | -1.3458758 | 7.31889108 | 95.2416002 | 5.74E-11 | 5.95E-09 | T | F | T | F |
| Nbea | 2.21536709 | 4.27912443 | 95.1542192 | 5.80E-11 | 5.99E-09 | T | F | F | T |
| Pnma2 | 4.21937361 | 2.57104051 | 94.7161731 | 6.13E-11 | 6.26E-09 | T | F | T | T |
| Lhx1 | 2.6232302 | 3.80690347 | 94.7161349 | 6.13E-11 | 6.26E-09 | F | F | T | T |
| C530008M1 | 2.10094204 | 4.45702706 | 93.7062957 | 6.95E-11 | 7.07E-09 | T | F | F | T |
| Efhd2 | -1.2398998 | 7.18492431 | 92.8962976 | 7.70E-11 | 7.80E-09 | T | F | T | T |
| Adcy7 | -1.4114958 | 6.68433076 | 92.7878843 | 7.80E-11 | 7.87E-09 | T | F | T | F |
| Nav3 | -1.5774473 | 6.18186733 | 92.1225119 | 8.49E-11 | 8.52E-09 | T | F | T | T |
| Irs4 | 5.68891916 | 1.96294931 | 92.0623318 | 8.55E-11 | 8.55E-09 | T | F | T | T |
| Cep164 | -1.2036147 | 7.25945574 | 91.6153385 | 9.05E-11 | 9.01E-09 | F | F | T | F |
| Ajuba | 1.13570646 | 7.04234979 | 90.6163841 | 1.03E-10 | 1.02E-08 | F | F | F | F |
| Dnmt3a | 1.14760728 | 6.99408525 | 90.5801765 | 1.03E-10 | 1.02E-08 | T | T | T | T |
| Dusp9 | -0.9189245 | 9.31161936 | 89.7659061 | 1.15E-10 | 1.13E-08 | F | F | T | F |
| Ccnd1 | 1.65782191 | 6.16304095 | 89.5911239 | 1.18E-10 | 1.15E-08 | T | F | T | T |
| P4ha2 | 1.83907383 | 5.12775817 | 89.2563579 | 1.23E-10 | 1.19E-08 | T | F | F | F |
| Arrb1 | 1.39798993 | 6.20547274 | 88.935591 | 1.28E-10 | 1.24E-08 | T | F | T | T |
| H1f10 | 1.71201158 | 5.19235242 | 88.844675 | 1.29E-10 | 1.25E-08 | T | F | F | T |
| Hic2 | -1.0102127 | 8.2800492 | 88.8193887 | 1.30E-10 | 1.25E-08 | T | F | T | T |
| Ldlr | -1.1380485 | 7.51945779 | 88.5371345 | 1.35E-10 | 1.28E-08 | T | F | T | T |
| Kif5c | 2.2708674 | 4.09440654 | 88.5232003 | 1.35E-10 | 1.28E-08 | F | F | T | T |
| Efna5 | 3.39551822 | 3.28942099 | 88.3029683 | 1.39E-10 | 1.32E-08 | T | F | T | T |
| Zfp423 | 2.09264441 | 4.46753745 | 88.2567384 | 1.40E-10 | 1.32E-08 | F | F | F | F |
| Myo18a | -1.0146055 | 8.26484443 | 88.1669013 | 1.42E-10 | 1.33E-08 | T | F | F | T |
| Jup | -1.0339472 | 8.17275187 | 88.123869 | 1.42E-10 | 1.33E-08 | F | F | T | T |
| Afap1l2 | -2.4795001 | 4.42638496 | 88.0837838 | 1.43E-10 | 1.33E-08 | T | F | T | T |
| Adamts15 | 3.22918037 | 3.06380684 | 87.9996018 | 1.45E-10 | 1.34E-08 | T | F | F | T |
| Foxh1 | 1.53097164 | 5.63368095 | 87.8427616 | 1.48E-10 | 1.36E-08 | F | F | F | F |
| Sema6a | 1.23147246 | 6.62873732 | 87.3521189 | 1.58E-10 | 1.45E-08 | F | F | T | F |
| Ldhb | 1.66511086 | 5.27887248 | 86.8613523 | 1.68E-10 | 1.53E-08 | T | F | T | T |
| Greb1 | 1.47295939 | 5.94949485 | 86.8192412 | 1.69E-10 | 1.53E-08 | F | F | F | F |
| Col18a1 | 1.00911392 | 7.76745169 | 86.8173345 | 1.69E-10 | 1.53E-08 | T | F | F | F |
| Rcn3 | 1.38526019 | 6.05897993 | 86.7934249 | 1.70E-10 | 1.53E-08 | T | F | F | F |
| Prkch | -1.6554262 | 6.18692162 | 86.344724 | 1.80E-10 | 1.62E-08 | F | F | F | F |
| Gtf2i | 0.9371538 | 8.11530847 | 85.9772078 | 1.89E-10 | 1.70E-08 | F | F | F | F |
| Prl3d2 | 2.41749039 | 3.90315153 | 85.8264229 | 1.93E-10 | 1.72E-08 | F | F | F | F |
| Btbd2 | 1.68376797 | 5.15218711 | 85.6216664 | 1.98E-10 | 1.76E-08 | T | F | F | F |
| Sept5 | -1.9366306 | 5.22211706 | 85.5236179 | 2.01E-10 | 1.78E-08 | F | F | F | F |
| Rgcc | -3.1539066 | 3.86249404 | 85.440793 | 2.03E-10 | 1.79E-08 | T | F | F | F |
| Clip3 | 2.213002 | 4.13420211 | 84.4342408 | 2.33E-10 | 2.05E-08 | T | F | T | T |
| Gata3 | -1.629666 | 6.3107282 | 83.8730061 | 2.51E-10 | 2.20E-08 | F | F | F | F |
| Plxna2 | 1.33167535 | 6.1905225 | 83.7205659 | 2.57E-10 | 2.24E-08 | T | F | T | T |

| | | | | | | | | | |
|---|---|---|---|---|---|---|---|---|---|
| Cdh3 | -1.0698938 | 7.65513968 | 83.2387863 | 2.74E-10 | 2.38E-08 | F | F | T | F |
| Tgfb3 | -4.5430939 | 3.03517424 | 82.7902438 | 2.92E-10 | 2.52E-08 | F | F | T | F |
| Fam131b | 4.11047424 | 2.34950144 | 82.4693936 | 3.05E-10 | 2.62E-08 | F | F | T | F |
| Zfp334 | 1.73261257 | 5.07055135 | 82.4562456 | 3.05E-10 | 2.62E-08 | T | F | T | T |
| Lrig3 | 1.84812199 | 4.73076277 | 82.3640469 | 3.09E-10 | 2.64E-08 | T | F | F | F |
| Plec | -0.8145502 | 9.88909342 | 82.2770576 | 3.13E-10 | 2.66E-08 | T | F | F | T |
| Scin | 1.69068711 | 5.26585295 | 81.4313975 | 3.52E-10 | 2.98E-08 | F | F | F | F |
| Pkd1l1 | 4.9390699 | 2.00471917 | 80.9001453 | 3.79E-10 | 3.20E-08 | F | F | F | F |
| Myo1c | -0.9685191 | 8.3988061 | 80.7973753 | 3.85E-10 | 3.23E-08 | T | F | T | T |
| Gli3 | 1.51665777 | 5.65085023 | 80.5145522 | 4.00E-10 | 3.34E-08 | T | F | F | T |
| Zfp629 | 2.04374536 | 4.45255305 | 80.5054193 | 4.01E-10 | 3.34E-08 | T | F | T | F |
| Pkp2 | -1.193794 | 7.05217674 | 80.2161729 | 4.17E-10 | 3.47E-08 | T | F | F | T |
| Bmerb1 | 4.45899749 | 2.38017387 | 80.1891939 | 4.19E-10 | 3.47E-08 | T | F | F | F |
| Cd24a | 1.50170376 | 6.82645886 | 80.1176689 | 4.23E-10 | 3.49E-08 | F | F | F | F |
| Mtcl1 | 1.18440022 | 6.63819114 | 79.442384 | 4.66E-10 | 3.81E-08 | F | F | F | T |
| Eps8l2 | -1.219534 | 6.9094772 | 79.4417721 | 4.66E-10 | 3.81E-08 | F | F | F | F |
| Sept6 | 2.22993698 | 3.9849433 | 79.2057985 | 4.82E-10 | 3.92E-08 | T | F | T | T |
| Was | -1.9273022 | 5.06409526 | 79.1933681 | 4.83E-10 | 3.92E-08 | T | F | T | F |
| Myc | -1.0140834 | 8.11353666 | 78.977963 | 4.98E-10 | 4.03E-08 | F | F | F | F |
| Fstl1 | 1.58827445 | 5.25744217 | 78.3577848 | 5.44E-10 | 4.38E-08 | T | F | T | T |
| Nfat5 | -1.809485 | 6.30707695 | 77.5619074 | 6.10E-10 | 4.90E-08 | T | F | T | T |
| Cdh5 | -0.8537516 | 9.08868804 | 77.3173849 | 6.32E-10 | 5.05E-08 | T | T | F | T |
| Lyn | -1.7005113 | 5.49149325 | 77.269006 | 6.36E-10 | 5.07E-08 | T | T | F | T |
| Bcl9 | 1.24966547 | 6.30591891 | 77.2317959 | 6.39E-10 | 5.08E-08 | F | F | T | F |
| Shisa6 | 5.15493924 | 1.88427736 | 76.9280849 | 6.68E-10 | 5.29E-08 | T | F | T | T |
| Gli2 | 1.99538735 | 4.26052127 | 76.291579 | 7.33E-10 | 5.76E-08 | T | F | T | F |
| Meis2 | 2.24880141 | 3.92214227 | 76.280996 | 7.34E-10 | 5.76E-08 | T | F | F | F |
| Sfrp2 | 4.0186997 | 2.39854659 | 76.2643915 | 7.36E-10 | 5.76E-08 | T | F | T | F |
| Macroh2a2 | 1.86665977 | 4.71640227 | 76.1438178 | 7.49E-10 | 5.84E-08 | F | F | T | F |
| Uchl1 | 5.72318538 | 2.20971429 | 75.2657422 | 8.52E-10 | 6.60E-08 | F | F | F | F |
| Fgf5 | 5.19422828 | 1.93755845 | 75.2648974 | 8.53E-10 | 6.60E-08 | T | F | T | F |
| Stmn2 | 4.778549 | 1.96354793 | 75.2339838 | 8.56E-10 | 6.61E-08 | T | F | T | T |
| Phlda2 | -0.8987743 | 8.54024782 | 75.1941651 | 8.61E-10 | 6.62E-08 | T | F | F | F |
| Zfp36l3 | -2.1472182 | 4.57092251 | 75.0297686 | 8.83E-10 | 6.76E-08 | F | F | F | F |
| Galnt16 | 4.5661017 | 2.05765538 | 74.8599104 | 9.05E-10 | 6.91E-08 | T | F | T | T |
| Rgl3 | -9.9423486 | 2.25781785 | 74.7712254 | 9.17E-10 | 6.97E-08 | F | F | T | F |
| Thbs3 | 3.10407953 | 2.93440908 | 74.7529742 | 9.20E-10 | 6.97E-08 | T | F | F | T |
| Llgl1 | 1.20363853 | 6.57344528 | 74.4671606 | 9.60E-10 | 7.25E-08 | F | F | F | F |
| Igf2 | -0.707217 | 10.595689 | 74.0832733 | 1.02E-09 | 7.65E-08 | T | F | T | F |
| Sema4d | -1.1186284 | 7.21553507 | 73.976629 | 1.03E-09 | 7.75E-08 | F | F | F | F |
| Tro | 1.67293944 | 4.9091508 | 73.9155579 | 1.04E-09 | 7.79E-08 | F | F | F | F |
| Arhgap35 | 0.96193189 | 7.45682457 | 73.6695773 | 1.08E-09 | 8.05E-08 | T | F | T | F |

| | | | | | | | | | |
|---|---|---|---|---|---|---|---|---|---|
| Itga6 | 1.44204138 | 5.65867538 | 73.2739895 | 1.15E-09 | 8.52E-08 | T | F | T | T |
| Lrp4 | 1.89231778 | 4.37700116 | 73.0957875 | 1.18E-09 | 8.72E-08 | T | T | T | T |
| Celsr1 | 1.45174037 | 5.5757118 | 72.7217314 | 1.25E-09 | 9.19E-08 | T | F | T | F |
| Pmaip1 | 2.53151821 | 3.51021295 | 72.6328172 | 1.26E-09 | 9.29E-08 | F | F | T | F |
| Nfkbiz | -1.3700497 | 6.65953929 | 72.6006344 | 1.27E-09 | 9.30E-08 | T | F | T | T |
| Antxr1 | 3.72426644 | 2.78609677 | 72.5630413 | 1.28E-09 | 9.32E-08 | T | F | T | T |
| Jade2 | 1.92123944 | 4.27774677 | 72.0170234 | 1.39E-09 | 1.01E-07 | T | F | F | F |
| Nxf7 | -1.0659345 | 7.28346188 | 71.7755381 | 1.44E-09 | 1.04E-07 | F | F | F | F |
| Lfng | 3.81395392 | 2.33971714 | 71.6944927 | 1.46E-09 | 1.05E-07 | F | F | T | F |
| Pclaf | 1.45748218 | 5.75195336 | 71.6588375 | 1.47E-09 | 1.06E-07 | F | F | T | F |
| Bcor | -0.8331739 | 9.08036175 | 71.6192856 | 1.47E-09 | 1.06E-07 | T | F | F | F |
| Cadm1 | 1.48995241 | 5.31617072 | 71.5664578 | 1.49E-09 | 1.06E-07 | F | F | T | F |
| Elf3 | -1.1117604 | 7.01485556 | 71.3301325 | 1.54E-09 | 1.10E-07 | T | F | T | F |
| Fat3 | 3.32013799 | 2.66238993 | 71.1227031 | 1.59E-09 | 1.13E-07 | T | F | T | F |
| Adcy3 | 2.36758656 | 3.63469569 | 70.8766181 | 1.65E-09 | 1.17E-07 | F | F | F | T |
| Sparc | 0.80402966 | 9.07758709 | 70.8482199 | 1.66E-09 | 1.17E-07 | T | F | T | F |
| Igdcc3 | 1.7257648 | 4.65404666 | 70.8467084 | 1.66E-09 | 1.17E-07 | F | F | F | F |
| Adgrb2 | 2.81858965 | 3.3200526 | 70.5589314 | 1.74E-09 | 1.22E-07 | F | F | F | F |
| Ptbp2 | 1.65185955 | 4.80701533 | 70.4125982 | 1.78E-09 | 1.24E-07 | F | F | T | F |
| Coch | 4.32798908 | 2.14169899 | 70.4103994 | 1.78E-09 | 1.24E-07 | F | F | F | F |
| Akt1 | -0.8789794 | 8.4157751 | 70.1510731 | 1.85E-09 | 1.28E-07 | T | F | F | F |
| Lbh | 2.93796879 | 3.01349411 | 69.9699063 | 1.90E-09 | 1.32E-07 | F | F | T | T |
| Mpdz | 1.30169783 | 5.86753547 | 69.9379562 | 1.91E-09 | 1.32E-07 | T | F | F | T |
| Fxyd6 | 2.13600666 | 3.9725519 | 69.616838 | 2.01E-09 | 1.38E-07 | F | F | F | T |
| Hdgfl3 | 2.7336459 | 3.31891988 | 69.5887549 | 2.02E-09 | 1.38E-07 | T | F | T | F |
| Ifitm1 | 2.69709134 | 3.32224729 | 69.35962 | 2.09E-09 | 1.43E-07 | F | F | F | F |
| Lima1 | -1.0150871 | 7.39014636 | 69.3271145 | 2.10E-09 | 1.43E-07 | F | F | T | F |
| Tcf3 | 0.93899913 | 7.40577605 | 69.2202021 | 2.14E-09 | 1.45E-07 | F | F | T | F |
| Mcc | 1.65831877 | 4.710387 | 68.9849941 | 2.22E-09 | 1.50E-07 | T | F | T | T |
| mt-Co3 | -1.0610394 | 7.19077552 | 68.9341292 | 2.24E-09 | 1.51E-07 | F | F | F | F |
| Pard6g | -1.3691312 | 6.0402167 | 67.5218707 | 2.80E-09 | 1.88E-07 | F | F | T | F |
| Foxd4 | 9.75519317 | 1.1527806 | 68.9526083 | 2.88E-09 | 1.93E-07 | F | F | F | F |
| Sall3 | 2.55493542 | 5.25096857 | 67.1694528 | 2.96E-09 | 1.98E-07 | T | F | T | T |
| Islr2 | 2.76294571 | 3.13537214 | 67.1100262 | 2.99E-09 | 1.99E-07 | T | F | T | T |
| Kctd17 | -1.3552347 | 6.26116337 | 66.8185459 | 3.13E-09 | 2.08E-07 | F | F | F | F |
| Dchs1 | 3.02246316 | 2.86410225 | 66.7872275 | 3.14E-09 | 2.08E-07 | F | F | F | F |
| Trib2 | 3.46352383 | 2.63040746 | 66.6625523 | 3.21E-09 | 2.11E-07 | T | T | T | T |
| Extl1 | -1.7590339 | 5.02537143 | 66.6620877 | 3.21E-09 | 2.11E-07 | F | F | F | T |
| Magel2 | 3.40458101 | 2.47835082 | 66.4038962 | 3.34E-09 | 2.20E-07 | T | F | F | F |
| B4galnt4 | 2.41569663 | 3.59723586 | 66.0519694 | 3.54E-09 | 2.32E-07 | F | F | F | T |
| Ntn1 | 1.45132099 | 5.47332525 | 65.9353857 | 3.61E-09 | 2.35E-07 | T | F | T | T |
| Dock11 | 1.3057617 | 5.77949651 | 65.9308751 | 3.61E-09 | 2.35E-07 | F | F | F | T |

| | | | | | | | | | |
|---|---|---|---|---|---|---|---|---|---|
| Gab2 | -1.9039815 | 4.72437553 | 65.8444022 | 3.66E-09 | 2.38E-07 | F | F | T | T |
| Plcg2 | 1.94162922 | 4.20879502 | 65.3486802 | 3.97E-09 | 2.57E-07 | T | T | F | T |
| Tubb6 | -1.0119501 | 7.25942456 | 65.1313123 | 4.11E-09 | 2.65E-07 | F | F | T | T |
| Tcfl5 | 2.3871428 | 3.52908238 | 64.9542433 | 4.23E-09 | 2.72E-07 | T | F | T | F |
| Nek6 | -1.3882448 | 6.04627786 | 64.8793378 | 4.28E-09 | 2.74E-07 | F | F | T | F |
| Dio3 | -3.4731312 | 3.4029327 | 64.8733977 | 4.29E-09 | 2.74E-07 | F | F | F | F |
| Tbc1d16 | 1.28039793 | 5.80351667 | 64.7748848 | 4.36E-09 | 2.78E-07 | F | F | T | T |
| Gprc5b | 2.97513127 | 2.79401914 | 64.2945204 | 4.72E-09 | 2.99E-07 | T | F | T | T |
| Setbp1 | 2.221457 | 3.69359938 | 64.2871689 | 4.72E-09 | 2.99E-07 | T | F | T | T |
| St6gal1 | 2.12045428 | 3.9319733 | 64.172389 | 4.81E-09 | 3.04E-07 | T | T | F | T |
| Cald1 | -1.1133663 | 7.71750982 | 63.9692112 | 4.98E-09 | 3.14E-07 | T | T | T | T |
| Snrpn | 1.18222515 | 6.33049509 | 63.9389879 | 5.00E-09 | 3.14E-07 | T | F | F | F |
| Hist1h2br | 1.33350101 | 6.21473633 | 63.9202315 | 5.02E-09 | 3.14E-07 | F | F | T | T |
| Tmem47 | 2.42454375 | 3.72443938 | 63.8341501 | 5.09E-09 | 3.18E-07 | F | F | T | F |
| Zic5 | 1.73885893 | 4.61488121 | 63.698884 | 5.21E-09 | 3.24E-07 | F | F | F | T |
| Chga | 4.15843702 | 2.03955515 | 63.685363 | 5.22E-09 | 3.24E-07 | F | F | F | F |
| Slitrk5 | 4.51119417 | 1.98891665 | 63.603978 | 5.29E-09 | 3.27E-07 | T | F | T | T |
| Lin28b | 1.50098951 | 4.99344447 | 63.5935251 | 5.30E-09 | 3.27E-07 | T | T | T | T |
| Gsn | -0.888019 | 7.91679672 | 63.4392394 | 5.44E-09 | 3.34E-07 | F | F | F | F |
| Ppp1r9b | -0.960254 | 7.47032362 | 63.4353119 | 5.44E-09 | 3.34E-07 | T | F | T | T |
| Ttyh3 | 1.05154138 | 6.63359095 | 63.2532213 | 5.61E-09 | 3.43E-07 | T | F | F | F |
| Emilin1 | -1.0702799 | 7.12709156 | 63.2221639 | 5.64E-09 | 3.44E-07 | F | F | F | F |
| Mmp14 | 1.45667857 | 5.32473216 | 63.2033707 | 5.65E-09 | 3.44E-07 | F | F | T | T |
| Atp1b2 | 3.80205901 | 2.17666528 | 63.180626 | 5.68E-09 | 3.45E-07 | T | F | T | T |
| Zfp219 | 1.43251695 | 5.25093159 | 63.1555406 | 5.70E-09 | 3.45E-07 | F | F | F | F |
| Ets2 | -0.9786589 | 7.4989815 | 62.7577269 | 6.09E-09 | 3.68E-07 | F | F | T | F |
| Sema6d | 1.54181784 | 5.02115139 | 62.5945332 | 6.26E-09 | 3.77E-07 | T | F | F | T |
| Trim28 | 0.66970513 | 9.84398076 | 62.4928742 | 6.37E-09 | 3.82E-07 | F | F | F | F |
| Mdk | 0.9514046 | 7.19743577 | 62.4747103 | 6.39E-09 | 3.83E-07 | T | T | T | T |
| Sirpa | 2.2804661 | 3.60437994 | 62.4304648 | 6.44E-09 | 3.84E-07 | T | F | F | T |
| Pstpip1 | -4.0036008 | 2.80044261 | 62.4048862 | 6.46E-09 | 3.85E-07 | F | F | F | F |
| Vat1l | 9.65734753 | 1.06068731 | 63.794697 | 6.49E-09 | 3.86E-07 | T | F | T | T |
| Fzd3 | 1.70146685 | 4.80647644 | 62.3031546 | 6.57E-09 | 3.90E-07 | F | F | T | T |
| Map1b | 0.96291436 | 6.99840964 | 62.2300961 | 6.66E-09 | 3.93E-07 | T | F | F | T |
| Sdk2 | -1.6126584 | 5.42876282 | 62.0798886 | 6.83E-09 | 4.02E-07 | F | F | F | F |
| Nacc1 | 0.79218628 | 8.20207318 | 62.0418313 | 6.87E-09 | 4.03E-07 | T | F | T | T |
| Myrf | 1.3433196 | 5.60329896 | 62.0357175 | 6.88E-09 | 4.03E-07 | F | F | T | T |
| Ltbp1 | 2.28252804 | 3.65591454 | 61.857811 | 7.09E-09 | 4.14E-07 | F | F | T | T |
| Fgfbp1 | -1.9249495 | 4.83474676 | 61.8504181 | 7.10E-09 | 4.14E-07 | F | F | F | F |
| Rragb | 2.89906516 | 2.81736377 | 61.6917626 | 7.29E-09 | 4.24E-07 | F | F | T | F |
| Mpzl1 | -0.9676434 | 7.36279279 | 61.6588564 | 7.33E-09 | 4.25E-07 | T | F | T | F |
| F5 | 2.71108604 | 2.99667212 | 61.6198368 | 7.38E-09 | 4.27E-07 | F | F | T | F |

| | | | | | | | | | |
|---|---|---|---|---|---|---|---|---|---|
| Bin2 | -1.7215233 | 4.97887444 | 61.5177051 | 7.51E-09 | 4.33E-07 | F | F | F | T |
| Sall4 | 0.70349708 | 9.43509896 | 61.2002849 | 7.93E-09 | 4.56E-07 | T | F | T | T |
| Jag2 | -1.5535176 | 5.94051756 | 60.7270248 | 8.60E-09 | 4.93E-07 | F | F | F | F |
| Hmgn5 | -1.638059 | 5.60642659 | 60.5064157 | 8.93E-09 | 5.11E-07 | F | F | T | T |
| Cabp7 | 6.18027338 | 1.33792846 | 60.4485212 | 9.02E-09 | 5.14E-07 | F | F | T | T |
| Mapk8ip2 | 2.45630296 | 3.47527669 | 60.1350123 | 9.52E-09 | 5.42E-07 | F | F | F | F |
| Wipf3 | -1.4858471 | 5.46933113 | 60.1046602 | 9.57E-09 | 5.43E-07 | F | F | T | F |
| Pitpnc1 | 2.3543843 | 3.40285559 | 59.8421454 | 1.00E-08 | 5.67E-07 | T | T | T | T |
| Tspan6 | 1.75058866 | 4.27426347 | 59.7710444 | 1.01E-08 | 5.72E-07 | T | F | T | F |
| Tex19.1 | -1.1035688 | 6.65281724 | 59.6508294 | 1.03E-08 | 5.83E-07 | T | F | F | F |
| Grb10 | 0.74767961 | 8.6539955 | 59.2928517 | 1.10E-08 | 6.19E-07 | T | F | T | T |
| Zfp521 | 2.21373931 | 3.60561717 | 59.2407591 | 1.11E-08 | 6.23E-07 | T | F | T | T |
| Man2a2 | -1.2324774 | 6.16763216 | 59.1156649 | 1.14E-08 | 6.35E-07 | T | F | T | T |
| Csrp1 | -1.0496758 | 7.01078972 | 58.900325 | 1.18E-08 | 6.57E-07 | F | F | F | T |
| Serpinb9f | -1.9825047 | 4.37673111 | 58.8552437 | 1.19E-08 | 6.61E-07 | T | T | F | T |
| Pde5a | 1.85458165 | 4.07957374 | 58.5972941 | 1.24E-08 | 6.90E-07 | F | F | T | T |
| Pramef12 | -1.5929603 | 5.31568226 | 58.5007546 | 1.27E-08 | 7.00E-07 | F | F | T | F |
| Adamts9 | 0.95152974 | 7.08726585 | 58.2632102 | 1.32E-08 | 7.28E-07 | F | F | F | F |
| Laptm4b | 1.20417427 | 5.91162237 | 57.9059185 | 1.41E-08 | 7.72E-07 | F | F | F | F |
| Slc16a2 | 3.37577117 | 2.58007147 | 57.8996809 | 1.41E-08 | 7.72E-07 | T | F | F | T |
| Axin2 | 1.27297401 | 6.00172915 | 57.8351864 | 1.42E-08 | 7.79E-07 | F | F | F | F |
| Dyrk2 | -1.1019696 | 6.75870241 | 57.755622 | 1.44E-08 | 7.88E-07 | T | F | F | T |
| Mapk4 | 2.30955582 | 3.37950142 | 57.4217908 | 1.53E-08 | 8.34E-07 | T | F | F | T |
| Mvp | -1.1539621 | 6.53382734 | 57.3677931 | 1.55E-08 | 8.40E-07 | F | F | F | F |
| Mta1 | 0.88877157 | 7.13648503 | 57.0988538 | 1.62E-08 | 8.79E-07 | F | F | T | F |
| Fos | -1.7209365 | 4.75647064 | 57.0324977 | 1.64E-08 | 8.87E-07 | T | F | F | T |
| Fam57b | 2.8389461 | 2.74982528 | 57.0206465 | 1.64E-08 | 8.87E-07 | F | F | T | F |
| Slit3 | 1.60669671 | 4.49533775 | 56.9895776 | 1.65E-08 | 8.90E-07 | F | F | T | T |
| Megf6 | 2.39833722 | 3.80629509 | 56.9781671 | 1.66E-08 | 8.90E-07 | F | F | T | T |
| Spon1 | 2.57466243 | 3.09002139 | 56.9315914 | 1.67E-08 | 8.95E-07 | T | F | T | T |
| Wnk2 | -0.9441854 | 7.2042256 | 56.8503603 | 1.70E-08 | 9.04E-07 | T | F | F | F |
| Ldb2 | 5.49515422 | 1.53593942 | 56.8454144 | 1.70E-08 | 9.04E-07 | T | F | T | F |
| Cntfr | 3.57551342 | 2.16436084 | 56.7744663 | 1.72E-08 | 9.14E-07 | F | F | T | F |
| Gng3 | 3.537852 | 2.17951649 | 56.7439948 | 1.73E-08 | 9.16E-07 | T | F | F | T |
| Krt7 | -1.2052928 | 6.57414171 | 56.7275823 | 1.73E-08 | 9.17E-07 | F | F | F | F |
| St8sia3 | 5.78782667 | 1.38049629 | 56.6996324 | 1.74E-08 | 9.19E-07 | T | F | T | T |
| BC067074 | 3.14347649 | 2.70455906 | 56.2182466 | 1.90E-08 | 1.00E-06 | F | F | F | F |
| Cpe | 2.88458105 | 2.76748357 | 55.9925865 | 1.98E-08 | 1.04E-06 | F | F | T | F |
| Pank2 | 1.83967109 | 4.08517062 | 55.9862421 | 1.98E-08 | 1.04E-06 | F | F | F | T |
| Fam129a | -1.3244317 | 5.85980802 | 55.944709 | 2.00E-08 | 1.04E-06 | F | F | F | T |
| Arhgap23 | -1.0763671 | 6.75637442 | 55.8092538 | 2.05E-08 | 1.07E-06 | F | F | T | F |
| Noto | 9.51566804 | 0.927923 | 56.8881061 | 2.08E-08 | 1.08E-06 | F | F | T | F |

| | | | | | | | | | |
|---|---|---|---|---|---|---|---|---|---|
| Rbm24 | 3.64317909 | 2.51993314 | 55.7146698 | 2.08E-08 | 1.08E-06 | T | F | F | T |
| Flnb | -0.6000389 | 10.6390889 | 55.6335564 | 2.11E-08 | 1.09E-06 | T | F | T | T |
| Cdk2ap1 | 0.8807994 | 7.08391284 | 55.5932486 | 2.13E-08 | 1.10E-06 | F | F | T | F |
| Zfp532 | 1.20567527 | 5.74794679 | 55.4665126 | 2.18E-08 | 1.12E-06 | T | F | T | F |
| Vangl2 | 1.20094082 | 5.70286387 | 55.3470484 | 2.23E-08 | 1.14E-06 | F | F | T | T |
| Tbrg1 | -0.7816173 | 8.33651951 | 55.0604457 | 2.35E-08 | 1.20E-06 | F | F | F | F |
| Rfx3 | 1.44068008 | 5.0352251 | 54.848867 | 2.44E-08 | 1.25E-06 | T | T | T | T |
| BC016579 | -4.1598098 | 2.57237623 | 54.7958894 | 2.46E-08 | 1.25E-06 | F | F | F | F |
| Rhobtb3 | 1.52468748 | 4.76302997 | 54.7905886 | 2.46E-08 | 1.25E-06 | T | F | T | F |
| Usp28 | 1.38986796 | 5.00632332 | 54.6243106 | 2.54E-08 | 1.29E-06 | F | F | F | F |
| Adgrv1 | 3.75827494 | 1.99602642 | 54.3896592 | 2.65E-08 | 1.34E-06 | F | F | F | F |
| Tagln2 | -0.8196587 | 7.97952173 | 54.2889176 | 2.70E-08 | 1.37E-06 | T | F | F | T |
| Mpzl2 | -1.0322832 | 6.70983299 | 54.1229844 | 2.79E-08 | 1.41E-06 | T | F | F | F |
| Map7d3 | 3.43168481 | 2.17471249 | 53.7704071 | 2.98E-08 | 1.50E-06 | F | F | T | F |
| Sdc1 | -0.742501 | 8.84399441 | 53.7549921 | 2.98E-08 | 1.50E-06 | T | F | T | T |
| Pcdh18 | 1.75309468 | 4.55888484 | 53.6802492 | 3.03E-08 | 1.52E-06 | T | F | T | T |
| Ephb4 | 0.85346202 | 7.18655351 | 53.6664694 | 3.03E-08 | 1.52E-06 | T | F | F | F |
| Rasgrp3 | 1.66061605 | 4.27390355 | 53.6462095 | 3.04E-08 | 1.52E-06 | T | F | T | F |
| Ank3 | 1.53411771 | 4.54479278 | 53.4932378 | 3.13E-08 | 1.56E-06 | T | T | T | T |
| Fzd5 | 1.39000436 | 5.18813873 | 53.4368453 | 3.17E-08 | 1.57E-06 | T | F | T | T |
| Gm29695 | 9.46300567 | 0.87876152 | 54.4837594 | 3.19E-08 | 1.58E-06 | F | F | F | F |
| Fscn1 | 0.69503155 | 8.71239175 | 53.3326295 | 3.23E-08 | 1.59E-06 | F | F | T | F |
| Ptpn13 | 0.9164871 | 6.96319957 | 53.2502543 | 3.28E-08 | 1.62E-06 | T | F | F | F |
| Gabbr2 | 5.39548142 | 1.78877668 | 53.2386191 | 3.29E-08 | 1.62E-06 | F | F | F | F |
| Fam83b | -1.9321253 | 4.42017324 | 53.2257079 | 3.29E-08 | 1.62E-06 | F | F | T | T |
| Tmem145 | 4.01761608 | 1.87730878 | 53.1474094 | 3.34E-08 | 1.64E-06 | F | F | F | F |
| Ezr | -0.651407 | 9.74320203 | 53.01489 | 3.43E-08 | 1.67E-06 | F | F | T | F |
| Col17a1 | -2.6514471 | 3.49625786 | 52.951666 | 3.47E-08 | 1.69E-06 | T | F | F | F |
| Zfp618 | 2.62961036 | 2.93705002 | 52.7130376 | 3.63E-08 | 1.76E-06 | T | F | T | F |
| Stra6 | 1.92246614 | 4.16845317 | 52.6115169 | 3.70E-08 | 1.79E-06 | T | F | F | F |
| Serpinb9b | -1.5468541 | 5.0037256 | 52.5979428 | 3.71E-08 | 1.79E-06 | T | F | F | T |
| Syt11 | 1.63584527 | 4.35294185 | 52.5692753 | 3.73E-08 | 1.80E-06 | T | F | T | T |
| Mesp1 | 3.73770034 | 2.99331361 | 52.5649318 | 3.73E-08 | 1.80E-06 | F | F | F | F |
| Slc39a6 | -1.209411 | 6.04321451 | 52.4579625 | 3.80E-08 | 1.83E-06 | F | F | F | T |
| Rhbg | 9.43847643 | 0.85589907 | 53.393506 | 3.89E-08 | 1.87E-06 | F | F | T | F |
| Fkbp10 | 1.73356086 | 4.11664446 | 52.2574162 | 3.95E-08 | 1.89E-06 | F | F | T | F |
| Col13a1 | 3.4368278 | 2.42730906 | 52.1644072 | 4.02E-08 | 1.92E-06 | T | F | F | T |
| Ceacam9 | -2.9661654 | 4.76461007 | 51.8677349 | 4.26E-08 | 2.03E-06 | T | F | F | T |
| Zfp64 | -0.9868284 | 6.93011102 | 51.8567169 | 4.26E-08 | 2.03E-06 | F | F | T | F |
| Epn3 | -1.425592 | 5.3338962 | 51.7890356 | 4.32E-08 | 2.05E-06 | F | F | F | T |
| Nf1 | 1.14533831 | 5.97475952 | 51.4240483 | 4.63E-08 | 2.19E-06 | T | F | F | F |
| Prl8a2 | 2.28825332 | 3.47620439 | 51.3875144 | 4.66E-08 | 2.20E-06 | F | F | F | F |

| | | | | | | | | | |
|---|---|---|---|---|---|---|---|---|---|
| Dpysl2 | 1.057927 | 6.42398267 | 51.3405018 | 4.71E-08 | 2.22E-06 | T | F | T | T |
| Mical2 | -1.1853953 | 6.35216621 | 51.167881 | 4.86E-08 | 2.29E-06 | T | F | T | T |
| Anxa4 | -1.3638243 | 5.44396701 | 51.1339838 | 4.90E-08 | 2.30E-06 | T | F | T | F |
| Tmod2 | 2.10190876 | 3.48805113 | 50.9913325 | 5.03E-08 | 2.36E-06 | T | F | T | T |
| Slit1 | 2.3461453 | 3.13218167 | 50.8891519 | 5.13E-08 | 2.40E-06 | F | F | F | T |
| Rai1 | 1.08000811 | 6.12656052 | 50.6535226 | 5.37E-08 | 2.50E-06 | T | F | F | T |
| Irs3 | -1.6117955 | 4.74578202 | 50.6401573 | 5.38E-08 | 2.51E-06 | F | F | F | T |
| Sox9 | 3.23187082 | 2.70378929 | 50.5956617 | 5.43E-08 | 2.52E-06 | F | F | F | F |
| Svil | -1.0793625 | 6.37680993 | 50.5851749 | 5.44E-08 | 2.52E-06 | T | T | F | T |
| Elmo1 | 1.68314134 | 4.19349671 | 50.5357687 | 5.49E-08 | 2.54E-06 | T | F | T | F |
| Rtl8b | 1.28435252 | 5.21181225 | 50.5104366 | 5.52E-08 | 2.55E-06 | T | F | F | F |
| Stmn1 | 0.8457236 | 7.04467185 | 50.4622843 | 5.57E-08 | 2.57E-06 | T | F | T | T |
| Socs2 | 2.13115043 | 3.44442607 | 50.3963412 | 5.65E-08 | 2.59E-06 | T | F | T | T |
| Irx2 | -1.1657071 | 6.17748987 | 50.2795043 | 5.77E-08 | 2.65E-06 | T | F | T | F |
| Nr2f2 | -1.5110944 | 4.98224406 | 50.1392436 | 5.93E-08 | 2.71E-06 | T | F | T | F |
| Vmn2r4 | -9.3626856 | 1.69195115 | 50.0295497 | 6.06E-08 | 2.77E-06 | F | F | F | F |
| Nacad | -2.5600901 | 3.5341429 | 49.9833885 | 6.12E-08 | 2.79E-06 | F | F | F | F |
| Gpc1 | 1.06789836 | 6.16256738 | 49.8834596 | 6.24E-08 | 2.84E-06 | F | F | T | F |
| Shisa2 | 2.27393513 | 3.41895514 | 49.8525116 | 6.27E-08 | 2.85E-06 | T | F | F | T |
| Zfhx2 | 1.67622169 | 4.13252587 | 49.8186916 | 6.32E-08 | 2.86E-06 | F | F | F | T |
| Sfn | -1.0378607 | 6.80895586 | 49.8059635 | 6.33E-08 | 2.86E-06 | F | F | F | F |
| Irx1 | -1.460623 | 5.86562656 | 49.7556004 | 6.39E-08 | 2.88E-06 | T | F | F | F |
| Pth1r | 0.79603734 | 7.43038317 | 49.7506089 | 6.40E-08 | 2.88E-06 | F | F | F | F |
| Mmp2 | 1.69913527 | 4.23955067 | 49.6649033 | 6.51E-08 | 2.92E-06 | F | F | T | F |
| Pcsk9 | 3.92172203 | 1.91216031 | 49.6640323 | 6.51E-08 | 2.92E-06 | T | F | T | T |
| Zbtb12 | 1.74166245 | 4.01625134 | 49.5881229 | 6.61E-08 | 2.95E-06 | F | F | T | F |
| Rxra | -0.9906439 | 6.76416945 | 49.5331783 | 6.68E-08 | 2.98E-06 | T | F | F | F |
| Grik5 | 1.95055718 | 3.86345572 | 49.5260263 | 6.69E-08 | 2.98E-06 | F | F | T | F |
| Unc80 | 5.3796511 | 1.26240712 | 49.4638731 | 6.77E-08 | 3.01E-06 | T | F | F | T |
| Rufy4 | -6.3957279 | 1.87953135 | 49.3947927 | 6.86E-08 | 3.04E-06 | T | F | F | T |
| Igsf9 | 1.27717291 | 5.21505464 | 49.2947352 | 7.00E-08 | 3.10E-06 | T | F | F | F |
| Sos1 | -1.1251335 | 6.50944328 | 49.2455839 | 7.07E-08 | 3.12E-06 | T | F | T | F |
| Mt2 | -1.0549308 | 7.74391217 | 49.2136009 | 7.11E-08 | 3.13E-06 | F | F | F | F |
| Dkk1 | 2.03660865 | 3.55401918 | 49.0461532 | 7.35E-08 | 3.23E-06 | T | F | F | F |
| Tor1aip1 | -1.1180348 | 6.15973205 | 49.0321672 | 7.37E-08 | 3.24E-06 | T | F | F | T |
| Piezo2 | 3.07565391 | 2.45495933 | 48.7333341 | 7.82E-08 | 3.43E-06 | F | F | T | F |
| Spred3 | 2.62417026 | 2.80197879 | 48.707104 | 7.86E-08 | 3.44E-06 | T | F | T | T |
| Neo1 | 1.00995048 | 6.44094008 | 48.6083969 | 8.01E-08 | 3.50E-06 | T | F | T | F |
| Plxna1 | 0.70783029 | 8.35793446 | 48.5890688 | 8.05E-08 | 3.50E-06 | F | F | T | T |
| Jam3 | 1.24421321 | 5.25183001 | 48.4590997 | 8.26E-08 | 3.59E-06 | T | T | T | T |
| Pkp3 | -1.4373707 | 5.09969285 | 48.2213553 | 8.66E-08 | 3.75E-06 | F | F | F | F |
| Rgs12 | 1.55345666 | 4.3662406 | 47.4927797 | 1.00E-07 | 4.33E-06 | T | F | F | T |

| | | | | | | | | | |
|---|---|---|---|---|---|---|---|---|---|
| Ncan | 4.59351536 | 1.51185213 | 47.4782029 | 1.00E-07 | 4.34E-06 | T | F | T | T |
| Gbx2 | 2.65564667 | 2.66492187 | 47.4685381 | 1.01E-07 | 4.34E-06 | T | T | T | T |
| Crybg3 | -1.0088041 | 6.60443484 | 47.4576088 | 1.01E-07 | 4.34E-06 | F | F | F | F |
| Olr1 | -2.3549752 | 3.71175963 | 47.3834974 | 1.02E-07 | 4.40E-06 | T | F | T | T |
| Tln1 | -0.6495967 | 9.24803793 | 47.1855614 | 1.07E-07 | 4.57E-06 | F | F | F | F |
| Fzd8 | 1.77431073 | 3.87656798 | 47.089149 | 1.09E-07 | 4.65E-06 | F | F | F | F |
| Trim8 | -0.8424396 | 7.61316399 | 46.7731115 | 1.16E-07 | 4.94E-06 | F | F | F | T |
| Pecam1 | 1.16103248 | 5.7241524 | 46.7516476 | 1.16E-07 | 4.95E-06 | F | F | F | F |
| Myd88 | -1.4546953 | 4.97202565 | 46.7460335 | 1.16E-07 | 4.95E-06 | F | F | T | F |
| Cxadr | 1.1736359 | 5.49176345 | 46.7270544 | 1.17E-07 | 4.96E-06 | T | T | T | T |
| Bmper | 2.51563309 | 3.85101323 | 46.6648318 | 1.18E-07 | 5.01E-06 | T | F | T | F |
| Camta1 | 2.54160737 | 2.74710259 | 46.6573634 | 1.19E-07 | 5.01E-06 | T | F | T | T |
| Cyfip2 | 1.72899016 | 4.00024841 | 46.5479272 | 1.21E-07 | 5.12E-06 | T | F | T | F |
| Ctsk | 4.21507572 | 1.60608886 | 46.2954236 | 1.28E-07 | 5.38E-06 | F | F | T | F |
| Hip1 | 0.91188469 | 6.6555805 | 46.2414454 | 1.29E-07 | 5.42E-06 | T | F | T | T |
| Peli2 | 2.36715992 | 3.0969889 | 46.1934072 | 1.30E-07 | 5.47E-06 | T | F | T | T |
| Zc3hav1 | -0.9041038 | 6.95594977 | 45.5584589 | 1.49E-07 | 6.22E-06 | T | F | T | F |
| Six3 | 7.09787041 | 0.85774804 | 45.2436748 | 1.59E-07 | 6.62E-06 | F | F | T | F |
| Ccdc184 | -2.1887303 | 3.71038424 | 45.2303532 | 1.59E-07 | 6.63E-06 | F | F | F | F |
| Tyro3 | 0.97979904 | 6.20189382 | 45.2084019 | 1.60E-07 | 6.63E-06 | T | F | F | T |
| Rnf213 | 1.39042682 | 5.90935504 | 45.2049247 | 1.60E-07 | 6.63E-06 | F | F | F | F |
| Itprid2 | -1.0632266 | 6.19395209 | 45.2008851 | 1.60E-07 | 6.63E-06 | F | F | F | F |
| Serpinb9c | -2.4183311 | 3.43584279 | 45.049198 | 1.65E-07 | 6.83E-06 | T | F | F | F |
| Foxa1 | 4.2313642 | 1.57398469 | 44.9759144 | 1.68E-07 | 6.92E-06 | T | F | F | T |
| Pcdh7 | 1.39332986 | 4.58668561 | 44.8546671 | 1.72E-07 | 7.09E-06 | T | F | T | T |
| Prickle1 | 1.32177385 | 4.96531738 | 44.6286846 | 1.80E-07 | 7.41E-06 | T | F | T | F |
| Pparg | -3.2879547 | 2.73979843 | 44.5221919 | 1.84E-07 | 7.57E-06 | F | F | F | T |
| Atp11a | -1.0049455 | 6.5798789 | 44.4873485 | 1.86E-07 | 7.61E-06 | F | F | F | F |
| Tln2 | 0.89430042 | 7.14546638 | 44.4605057 | 1.87E-07 | 7.64E-06 | T | F | T | T |
| Cd82 | -1.0478138 | 6.24434413 | 44.3690708 | 1.90E-07 | 7.77E-06 | F | F | T | F |
| Wasf1 | 1.72173245 | 4.1621188 | 44.3117928 | 1.93E-07 | 7.85E-06 | T | F | F | T |
| Ccdc40 | 3.1091035 | 2.39976557 | 44.2922166 | 1.93E-07 | 7.87E-06 | F | F | T | F |
| Rsad2 | -3.1354877 | 2.76248183 | 44.1285613 | 2.00E-07 | 8.13E-06 | T | F | T | T |
| Slc4a4 | 1.70373129 | 3.99149681 | 44.0943352 | 2.02E-07 | 8.17E-06 | T | F | T | T |
| Nlgn2 | 1.06609997 | 5.74208408 | 43.9973849 | 2.06E-07 | 8.32E-06 | T | F | T | F |
| Plet1 | -0.7146704 | 8.30657223 | 43.9307857 | 2.09E-07 | 8.43E-06 | T | F | F | T |
| Sri | -1.2669393 | 5.51914639 | 43.8476009 | 2.12E-07 | 8.56E-06 | F | F | F | F |
| Psrc1 | 1.62039268 | 4.04408674 | 43.8331135 | 2.13E-07 | 8.57E-06 | F | F | T | F |
| Prrt3 | 4.3428971 | 1.47068796 | 43.7420707 | 2.17E-07 | 8.72E-06 | F | F | F | F |
| Wnt3 | 3.76356821 | 1.78533541 | 43.6922101 | 2.19E-07 | 8.80E-06 | T | F | T | F |
| Pde3b | 1.95393965 | 3.69751511 | 43.6452548 | 2.22E-07 | 8.87E-06 | T | F | T | T |
| Hoxb1 | 3.58019157 | 3.1080679 | 43.5999742 | 2.24E-07 | 8.93E-06 | F | F | T | F |

| | | | | | | | | | |
|---|---|---|---|---|---|---|---|---|---|
| Apoc2 | -0.6997165 | 8.23617172 | 43.5934228 | 2.24E-07 | 8.93E-06 | T | F | F | F |
| Porcn | 5.22941898 | 1.42416665 | 43.5426687 | 2.27E-07 | 9.01E-06 | F | F | F | T |
| Crybg1 | -1.4258662 | 4.8844124 | 43.5366898 | 2.27E-07 | 9.01E-06 | F | F | T | F |
| Amz1 | -3.0872657 | 2.77716225 | 43.5266402 | 2.27E-07 | 9.01E-06 | T | F | F | T |
| Basp1 | -0.8161609 | 7.68980934 | 43.5027558 | 2.28E-07 | 9.04E-06 | T | F | F | F |
| Dusp8 | -2.37741 | 3.48563363 | 43.3124076 | 2.38E-07 | 9.40E-06 | F | F | T | T |
| Rimklb | 2.4014049 | 2.85342281 | 43.2860811 | 2.39E-07 | 9.43E-06 | T | F | T | T |
| Nrcam | 3.76843697 | 2.45195958 | 43.1084851 | 2.48E-07 | 9.78E-06 | T | F | T | T |
| Syne2 | 0.987057 | 6.15580707 | 42.9507045 | 2.57E-07 | 1.01E-05 | T | F | F | F |
| Pcsk2 | 9.1978669 | 0.63293637 | 43.6668139 | 2.59E-07 | 1.01E-05 | T | F | F | F |
| Hmga2 | 0.87499383 | 6.90218509 | 42.8655108 | 2.62E-07 | 1.02E-05 | T | F | T | F |
| Lrrn1 | 3.12786189 | 2.24461455 | 42.7765484 | 2.67E-07 | 1.04E-05 | F | F | F | F |
| Arhgef17 | 0.87631118 | 6.60425275 | 42.7505827 | 2.68E-07 | 1.05E-05 | T | F | T | T |
| Nog | 2.1924215 | 3.74807439 | 42.6910025 | 2.72E-07 | 1.06E-05 | F | F | T | F |
| Spata13 | 0.80861304 | 6.93176193 | 42.6521105 | 2.74E-07 | 1.06E-05 | T | F | T | F |
| Gnas | 0.66770432 | 8.35954495 | 42.5444683 | 2.80E-07 | 1.09E-05 | T | F | T | F |
| Grem2 | 1.97575899 | 3.67445423 | 42.4952375 | 2.83E-07 | 1.10E-05 | T | F | T | T |
| Jag1 | 1.21924146 | 5.02217258 | 42.2368718 | 3.00E-07 | 1.16E-05 | F | F | F | F |
| Col9a3 | 4.02625762 | 1.76718263 | 42.166897 | 3.04E-07 | 1.17E-05 | T | F | F | F |
| Arap1 | -1.1426572 | 5.72789329 | 42.1049705 | 3.08E-07 | 1.19E-05 | F | F | F | F |
| Mboat2 | 1.67131754 | 3.88221709 | 42.0838117 | 3.10E-07 | 1.19E-05 | F | F | T | T |
| Serpinb9 | -1.6890914 | 4.25142225 | 41.6831448 | 3.38E-07 | 1.30E-05 | T | F | T | T |
| Zswim6 | 2.07993428 | 3.48454219 | 41.674037 | 3.39E-07 | 1.30E-05 | T | F | F | F |
| Prss8 | -0.8115484 | 7.30130351 | 41.6575347 | 3.40E-07 | 1.30E-05 | T | F | F | F |
| Gjc1 | 2.35578111 | 3.01293077 | 41.5175481 | 3.50E-07 | 1.34E-05 | T | F | F | T |
| Thbd | 0.86729016 | 6.55803687 | 41.5118895 | 3.51E-07 | 1.34E-05 | T | F | F | T |
| Ccdc194 | 6.19566388 | 0.83999943 | 41.3618312 | 3.62E-07 | 1.38E-05 | F | F | T | F |
| Tuba1c | -0.6397191 | 9.00741505 | 41.2346445 | 3.73E-07 | 1.42E-05 | F | F | T | F |
| Plat | 1.754216 | 3.77096416 | 41.217069 | 3.74E-07 | 1.42E-05 | T | F | F | F |
| Pip4k2c | -0.7998758 | 7.17043987 | 41.1521369 | 3.79E-07 | 1.44E-05 | T | F | T | F |
| Gm8797 | -0.9849266 | 6.3095021 | 41.0941365 | 3.84E-07 | 1.45E-05 | F | F | F | F |
| Maged2 | 1.25963178 | 5.07655315 | 40.9809348 | 3.94E-07 | 1.48E-05 | F | F | F | F |
| Igdcc4 | 3.49056401 | 1.80738891 | 40.9798193 | 3.94E-07 | 1.48E-05 | T | F | T | T |
| Gpc3 | 0.61094581 | 8.7019384 | 40.9740543 | 3.95E-07 | 1.48E-05 | F | F | F | F |
| Dtx3 | 1.32007546 | 4.61059673 | 40.9712138 | 3.95E-07 | 1.48E-05 | F | F | T | F |
| Rtn1 | -1.2922009 | 5.42305709 | 40.9153075 | 4.00E-07 | 1.50E-05 | T | F | T | F |
| Ly6e | 1.14444833 | 5.25135217 | 40.8985061 | 4.01E-07 | 1.50E-05 | F | F | T | F |
| Mmp25 | 9.13968433 | 0.5793978 | 41.5680496 | 4.03E-07 | 1.50E-05 | F | F | F | T |
| Nptx1 | 3.73635365 | 1.65582611 | 40.8323317 | 4.07E-07 | 1.52E-05 | T | F | T | T |
| Snn | 1.75301529 | 3.79356712 | 40.8294233 | 4.07E-07 | 1.52E-05 | T | F | T | T |
| Tafa5 | 2.26558329 | 2.88147092 | 40.760647 | 4.14E-07 | 1.54E-05 | T | F | T | F |
| Vav3 | -1.936887 | 3.92282166 | 40.7458544 | 4.15E-07 | 1.54E-05 | T | F | T | F |

| | | | | | | | | | |
|---|---|---|---|---|---|---|---|---|---|
| Mgat3 | 2.21638989 | 2.98147688 | 40.6853731 | 4.21E-07 | 1.56E-05 | F | F | T | F |
| Tmbim1 | -1.0245654 | 6.14594716 | 40.6744808 | 4.22E-07 | 1.56E-05 | F | F | T | F |
| Lrmp | 5.8800545 | 1.01516435 | 40.6655981 | 4.22E-07 | 1.56E-05 | F | F | F | F |
| Cd164 | -0.7666155 | 8.07059031 | 40.6429565 | 4.25E-07 | 1.56E-05 | T | F | T | F |
| Trim6 | -0.796216 | 7.15402922 | 40.5586596 | 4.33E-07 | 1.59E-05 | F | F | T | F |
| Gab1 | -0.7358816 | 7.80013178 | 40.5581274 | 4.33E-07 | 1.59E-05 | F | F | T | F |
| Spred1 | 1.06172399 | 5.86365446 | 40.4239391 | 4.46E-07 | 1.63E-05 | T | F | F | T |
| Ttc3 | 0.75135674 | 7.09977472 | 40.4025017 | 4.48E-07 | 1.64E-05 | T | F | F | F |
| Arpc2 | -0.7196981 | 7.99574776 | 40.3780961 | 4.50E-07 | 1.64E-05 | T | F | F | F |
| Kcnj3 | 6.96395761 | 0.76443495 | 40.3107909 | 4.57E-07 | 1.67E-05 | T | F | T | F |
| Phf6 | 1.44086788 | 4.49135915 | 40.1893492 | 4.69E-07 | 1.71E-05 | T | F | F | F |
| Cntn2 | 3.50739388 | 1.92232005 | 40.1613679 | 4.72E-07 | 1.72E-05 | T | F | T | F |
| Gm17067 | -1.3723943 | 4.86806974 | 39.9683395 | 4.93E-07 | 1.79E-05 | F | F | F | T |
| Wnk3 | -0.9525289 | 6.45153682 | 39.9122166 | 4.99E-07 | 1.81E-05 | T | F | T | T |
| Sox12 | 1.00317805 | 5.78509441 | 39.787648 | 5.13E-07 | 1.86E-05 | F | F | F | T |
| Cdh1 | 0.61416539 | 8.8844594 | 39.7803148 | 5.14E-07 | 1.86E-05 | T | F | F | F |
| Itgb5 | 0.93936846 | 6.15997661 | 39.656545 | 5.29E-07 | 1.90E-05 | F | F | F | T |
| Trib1 | 1.69526791 | 3.74915452 | 39.4291236 | 5.57E-07 | 2.00E-05 | T | F | F | F |
| Tub | 4.04783723 | 1.57882558 | 39.4125442 | 5.59E-07 | 2.00E-05 | T | F | T | F |
| Cdca7 | 0.79258193 | 6.88169895 | 39.4108983 | 5.59E-07 | 2.00E-05 | F | F | F | F |
| Ralgds | -0.8999819 | 6.61874457 | 39.3880219 | 5.62E-07 | 2.01E-05 | T | F | T | F |
| Myo1d | -1.1285564 | 5.6473979 | 39.36032 | 5.65E-07 | 2.02E-05 | T | F | T | F |
| Plekho2 | 1.63492091 | 3.99196148 | 39.3186276 | 5.71E-07 | 2.03E-05 | T | F | T | F |
| Zmym2 | 0.99671953 | 5.91692666 | 39.2559405 | 5.79E-07 | 2.06E-05 | F | F | T | T |
| Rcc2 | 0.59079097 | 8.7872274 | 39.1015027 | 5.99E-07 | 2.13E-05 | F | F | T | F |
| Glcci1 | 1.46219743 | 4.17766578 | 39.0200586 | 6.10E-07 | 2.17E-05 | T | T | T | T |
| Pdzrn3 | 1.57941382 | 3.9372432 | 38.9720957 | 6.17E-07 | 2.19E-05 | T | F | F | T |
| Pltp | -1.1122054 | 5.84386892 | 38.9041221 | 6.27E-07 | 2.22E-05 | F | F | F | F |
| Jph3 | 6.34275555 | 0.81999171 | 38.8493099 | 6.34E-07 | 2.24E-05 | T | F | T | T |
| Gse1 | -0.7614835 | 7.37472815 | 38.8155326 | 6.39E-07 | 2.25E-05 | T | F | T | F |
| Ksr1 | -1.0017019 | 6.08810785 | 38.7698692 | 6.46E-07 | 2.27E-05 | F | F | T | F |
| Inf2 | -0.9678027 | 6.27103622 | 38.7602028 | 6.47E-07 | 2.28E-05 | F | F | F | F |
| Sgk2 | -4.5730845 | 1.94875571 | 38.6097298 | 6.70E-07 | 2.35E-05 | F | F | F | T |
| Dysf | -0.9367421 | 6.36176719 | 38.4517243 | 6.95E-07 | 2.43E-05 | F | F | F | F |
| Shtn1 | -1.3185783 | 5.04667497 | 38.2394087 | 7.29E-07 | 2.55E-05 | F | F | T | F |
| Adgra3 | 0.7927826 | 6.81629515 | 38.1785939 | 7.39E-07 | 2.58E-05 | F | F | T | F |
| Fam168a | -0.9258987 | 6.5229021 | 38.1363348 | 7.47E-07 | 2.60E-05 | T | F | T | T |
| Sall1 | 0.91315986 | 6.77240443 | 38.10975 | 7.51E-07 | 2.61E-05 | F | F | F | T |
| Tacc1 | -0.8066114 | 6.96455909 | 38.1009861 | 7.53E-07 | 2.61E-05 | T | F | T | T |
| Snph | 4.17839466 | 1.4187736 | 38.0948011 | 7.54E-07 | 2.61E-05 | F | F | F | F |
| Fbln1 | 0.72041488 | 7.18994796 | 38.0900514 | 7.55E-07 | 2.61E-05 | F | F | T | F |
| Pcbp4 | 0.67280481 | 7.7585484 | 37.9548185 | 7.78E-07 | 2.69E-05 | F | F | F | F |

| | | | | | | | | | |
|---|---|---|---|---|---|---|---|---|---|
| Mgat5b | 4.24065535 | 1.32055387 | 37.9437998 | 7.80E-07 | 2.69E-05 | F | F | F | F |
| Robo3 | 9.0434177 | 0.491163 | 38.514925 | 7.86E-07 | 2.71E-05 | F | F | T | F |
| Ide | -0.8144977 | 7.09785529 | 37.8733819 | 7.93E-07 | 2.73E-05 | F | F | T | F |
| Cnn3 | 0.72941093 | 7.23497368 | 37.8274338 | 8.02E-07 | 2.75E-05 | T | F | F | T |
| Zfp667 | -1.3639676 | 4.82830924 | 37.7500463 | 8.16E-07 | 2.80E-05 | T | F | F | T |
| Lmo2 | -1.6645981 | 4.70336599 | 37.7085124 | 8.24E-07 | 2.82E-05 | T | F | F | T |
| Loxl2 | 0.70857641 | 7.28197428 | 37.7009389 | 8.25E-07 | 2.82E-05 | F | F | T | T |
| Ncs1 | 1.18310675 | 4.91610411 | 37.6929233 | 8.27E-07 | 2.82E-05 | T | F | T | T |
| Prr36 | 3.7354403 | 1.62706823 | 37.6892881 | 8.28E-07 | 2.82E-05 | F | F | T | F |
| Cnn2 | -0.7719473 | 7.12193417 | 37.652738 | 8.35E-07 | 2.84E-05 | T | F | T | T |
| Pkm | 0.43634109 | 11.3833522 | 37.6119054 | 8.42E-07 | 2.86E-05 | F | F | F | F |
| Tmsb4x | -1.0601823 | 5.931734 | 37.5999912 | 8.45E-07 | 2.87E-05 | F | F | T | F |
| Prl7a1 | 1.82454184 | 5.03170268 | 37.5910635 | 8.47E-07 | 2.87E-05 | F | F | F | F |
| Fbn2 | 1.15714866 | 5.69577772 | 37.5356076 | 8.57E-07 | 2.90E-05 | T | F | T | F |
| Evc | 1.93893043 | 3.23388429 | 37.419289 | 8.81E-07 | 2.98E-05 | T | F | T | T |
| Ppp1r13b | -0.9965303 | 6.11278394 | 37.3966366 | 8.86E-07 | 2.99E-05 | F | F | F | F |
| Nasp | 0.59549646 | 8.53337 | 37.3494588 | 8.95E-07 | 3.01E-05 | T | F | F | T |
| Hmga1b | 0.51690075 | 9.85376775 | 37.2743984 | 9.11E-07 | 3.06E-05 | F | F | F | T |
| Dtx3l | 1.78324527 | 3.53751851 | 37.2461148 | 9.17E-07 | 3.08E-05 | T | F | T | F |
| mt-Co1 | -0.431907 | 12.1033374 | 37.1270323 | 9.43E-07 | 3.16E-05 | F | F | F | F |
| Arap2 | -1.1495306 | 5.43851575 | 37.115975 | 9.45E-07 | 3.16E-05 | F | F | T | F |
| Jak1 | -0.7762798 | 7.1705534 | 37.0049111 | 9.70E-07 | 3.24E-05 | F | F | T | F |
| Plod1 | 1.01593931 | 5.56358011 | 36.8968948 | 9.95E-07 | 3.32E-05 | T | F | T | T |
| Hap1 | -1.6899148 | 4.07245102 | 36.8540655 | 1.00E-06 | 3.35E-05 | F | F | T | F |
| Myl12a | -0.7842989 | 7.02735473 | 36.8480044 | 1.01E-06 | 3.35E-05 | F | F | F | T |
| Csf1r | 2.08862654 | 2.96707101 | 36.7058991 | 1.04E-06 | 3.45E-05 | T | F | F | F |
| Fabp5 | 1.06084372 | 5.38731948 | 36.6529636 | 1.05E-06 | 3.49E-05 | F | F | F | F |
| Klhl22 | -0.7256819 | 7.44281755 | 36.570511 | 1.07E-06 | 3.56E-05 | T | F | F | T |
| Gm9 | -1.8951145 | 3.77300079 | 36.520937 | 1.09E-06 | 3.59E-05 | T | F | F | F |
| Anp32a | 0.91274792 | 6.28991451 | 36.4937014 | 1.09E-06 | 3.61E-05 | F | F | T | T |
| Fads1 | 1.12581874 | 5.09088077 | 36.4562978 | 1.10E-06 | 3.64E-05 | T | F | T | T |
| Gfra3 | 5.62564422 | 0.80872256 | 36.4183846 | 1.11E-06 | 3.66E-05 | F | F | F | F |
| Lpar4 | 3.06870801 | 2.24127691 | 36.4084499 | 1.12E-06 | 3.67E-05 | T | F | T | T |
| Arpc5 | -0.7791626 | 7.32094629 | 36.3554436 | 1.13E-06 | 3.71E-05 | F | F | F | F |
| Atp2c2 | -1.3257855 | 4.80094971 | 36.3391068 | 1.13E-06 | 3.71E-05 | F | F | F | F |
| Ubxn10 | 8.98432608 | 0.43722563 | 36.8136039 | 1.16E-06 | 3.78E-05 | F | F | T | F |
| Spry2 | 1.18210301 | 4.8114745 | 36.2543079 | 1.16E-06 | 3.78E-05 | F | F | T | F |
| Mgat4b | -0.7513486 | 7.29193752 | 36.2114417 | 1.17E-06 | 3.81E-05 | F | F | F | F |
| Cbx2 | 0.8742134 | 6.24675158 | 36.156886 | 1.18E-06 | 3.85E-05 | T | F | T | T |
| Nnt | 1.01121621 | 5.61608487 | 36.1359978 | 1.19E-06 | 3.87E-05 | T | F | T | T |
| Upp1 | -1.1130287 | 5.59050425 | 36.0820627 | 1.21E-06 | 3.91E-05 | F | F | F | T |
| Casp3 | 1.05956809 | 5.46527671 | 36.0530793 | 1.21E-06 | 3.93E-05 | T | F | F | F |

| | | | | | | | | | |
|---|---|---|---|---|---|---|---|---|---|
| Gm49394 | -0.7043495 | 7.79651592 | 36.0108556 | 1.23E-06 | 3.97E-05 | F | F | F | F |
| Cdh20 | 5.18780103 | 0.93235621 | 36.0010895 | 1.23E-06 | 3.97E-05 | T | F | T | T |
| Vgll3 | -1.5612219 | 4.47940235 | 35.9587982 | 1.24E-06 | 4.00E-05 | T | F | F | T |
| Heph | 3.13811416 | 2.2940437 | 35.9463471 | 1.24E-06 | 4.01E-05 | T | F | F | T |
| Chrd | 3.64300624 | 3.01472417 | 35.9419574 | 1.25E-06 | 4.01E-05 | T | F | T | F |
| Wdfy1 | -1.3568056 | 4.67161115 | 35.9324886 | 1.25E-06 | 4.01E-05 | F | F | T | T |
| Cdon | 1.23819371 | 4.7507288 | 35.9159729 | 1.25E-06 | 4.02E-05 | T | F | T | T |
| Crim1 | -0.9149557 | 6.49937372 | 35.9150749 | 1.25E-06 | 4.02E-05 | F | F | T | F |
| Runx1 | -1.017931 | 5.85557605 | 35.8810367 | 1.26E-06 | 4.04E-05 | T | F | T | T |
| Nub1 | -0.8871881 | 6.87985129 | 35.6873006 | 1.32E-06 | 4.23E-05 | T | F | T | T |
| Etv1 | 2.14800076 | 2.94391429 | 35.6474095 | 1.34E-06 | 4.26E-05 | T | F | T | T |
| Shroom4 | -0.8917875 | 6.40583424 | 35.6349473 | 1.34E-06 | 4.26E-05 | F | F | T | T |
| Il13ra1 | -2.220755 | 3.28631352 | 35.6321224 | 1.34E-06 | 4.26E-05 | T | F | F | T |
| Smarcd1 | 0.66936095 | 7.46456676 | 35.6106113 | 1.35E-06 | 4.28E-05 | F | F | F | F |
| 3830417A1 | -1.2034449 | 5.39618187 | 35.5909214 | 1.35E-06 | 4.29E-05 | F | F | F | F |
| H4c11 | 0.61632306 | 8.01620717 | 35.5875896 | 1.36E-06 | 4.29E-05 | F | F | F | T |
| Mmp28 | -2.721424 | 2.75629687 | 35.5828045 | 1.36E-06 | 4.29E-05 | F | F | T | F |
| Jak3 | -1.6524558 | 4.18026168 | 35.5714831 | 1.36E-06 | 4.29E-05 | T | T | F | T |
| Psg29 | 1.52182501 | 6.81028892 | 35.5190496 | 1.38E-06 | 4.34E-05 | T | F | T | F |
| Arhgap30 | -8.88375 | 1.23929556 | 35.4958921 | 1.39E-06 | 4.36E-05 | T | F | F | T |
| Kif26b | 1.8155693 | 3.77552245 | 35.488475 | 1.39E-06 | 4.36E-05 | F | F | F | T |
| Chd6 | 0.81165931 | 6.52401415 | 35.3294676 | 1.44E-06 | 4.52E-05 | T | F | F | F |
| Atp10a | 1.04815444 | 5.71635996 | 35.231958 | 1.48E-06 | 4.62E-05 | F | F | F | T |
| Pik3c2b | 1.18373811 | 4.79857989 | 35.1581806 | 1.50E-06 | 4.70E-05 | T | F | T | F |
| Serpinb6b | -2.069285 | 3.78045743 | 35.1498536 | 1.51E-06 | 4.70E-05 | F | F | F | T |
| Gpr50 | 8.93935147 | 0.39629269 | 35.5543364 | 1.55E-06 | 4.83E-05 | F | F | T | T |
| E330009J07 | 1.35914267 | 4.37589359 | 35.0095752 | 1.56E-06 | 4.85E-05 | T | F | T | F |
| Xkr5 | 2.8945966 | 2.02901616 | 34.9957841 | 1.56E-06 | 4.86E-05 | F | F | F | F |
| Dll4 | 1.31544436 | 4.35782053 | 34.9901338 | 1.56E-06 | 4.86E-05 | F | F | F | F |
| Wdr1 | -0.6357764 | 8.36713093 | 34.9800205 | 1.57E-06 | 4.86E-05 | T | F | T | F |
| Scn8a | 3.4601377 | 1.65054315 | 34.8933752 | 1.60E-06 | 4.96E-05 | T | F | T | F |
| Tram2 | -0.6966616 | 7.6626956 | 34.8769521 | 1.61E-06 | 4.97E-05 | T | F | F | F |
| Sept3 | 2.53839827 | 2.31052939 | 34.8021779 | 1.64E-06 | 5.05E-05 | T | F | T | T |
| Rhox5 | -0.6447162 | 8.04402707 | 34.8001595 | 1.64E-06 | 5.05E-05 | F | F | F | F |
| Nfatc4 | 1.70012732 | 3.64407374 | 34.7691986 | 1.65E-06 | 5.08E-05 | T | F | F | T |
| Aplp2 | 0.58821632 | 8.31357135 | 34.7615477 | 1.65E-06 | 5.08E-05 | F | F | F | F |
| Kif1c | -0.6480014 | 8.06580172 | 34.6045812 | 1.72E-06 | 5.27E-05 | T | F | T | T |
| Arhgef6 | -2.0178512 | 3.5444777 | 34.6011103 | 1.72E-06 | 5.27E-05 | T | F | T | T |
| Rusc2 | -1.2331948 | 4.98218863 | 34.5439271 | 1.74E-06 | 5.34E-05 | F | F | T | F |
| Vwa5b1 | 5.32270735 | 0.81517362 | 34.4851912 | 1.77E-06 | 5.40E-05 | F | F | F | T |
| H1f6 | 1.98531632 | 3.09726347 | 34.3877044 | 1.81E-06 | 5.53E-05 | F | F | F | F |
| Asxl1 | 0.68159428 | 7.49756612 | 34.3713778 | 1.82E-06 | 5.54E-05 | T | F | F | F |

| | | | | | | | | | |
|---|---|---|---|---|---|---|---|---|---|
| Zxdb | -1.5581929 | 4.19969035 | 34.3622599 | 1.82E-06 | 5.55E-05 | T | F | T | T |
| Elf5 | -0.9966169 | 6.43576962 | 34.3398505 | 1.83E-06 | 5.57E-05 | T | F | F | T |
| Rnf144a | 2.07149654 | 3.15514135 | 34.2806402 | 1.86E-06 | 5.64E-05 | T | F | T | F |
| Mt1 | -0.840548 | 7.35332955 | 34.2625815 | 1.87E-06 | 5.66E-05 | F | F | F | T |
| Lrfn4 | 1.03446808 | 5.33142218 | 34.2487684 | 1.87E-06 | 5.67E-05 | F | F | F | F |
| Pknox2 | 2.2164843 | 2.66936827 | 34.1687656 | 1.91E-06 | 5.77E-05 | T | F | T | F |
| Fat4 | 2.34464412 | 2.50204011 | 34.1618703 | 1.91E-06 | 5.77E-05 | T | F | F | F |
| Gys1 | -0.8550785 | 6.66416618 | 34.0926384 | 1.94E-06 | 5.86E-05 | T | F | F | T |
| Myh9 | -0.4471135 | 11.1411072 | 34.087557 | 1.95E-06 | 5.86E-05 | T | F | T | F |
| Skida1 | 2.01972655 | 2.94583329 | 34.0251126 | 1.98E-06 | 5.95E-05 | T | F | F | T |
| Arhgef40 | 1.64699547 | 3.61324426 | 34.0064232 | 1.99E-06 | 5.96E-05 | T | F | T | T |
| Ccdc177 | 6.77257857 | 0.5914959 | 34.0031396 | 1.99E-06 | 5.96E-05 | F | F | T | F |
| Cnksr3 | 1.2704692 | 4.42186328 | 33.9568878 | 2.01E-06 | 6.02E-05 | F | F | F | T |
| Rnf122 | 3.55857605 | 1.7078115 | 33.9312347 | 2.02E-06 | 6.05E-05 | T | F | T | T |
| Rap1a | -1.1112564 | 5.38978923 | 33.9211933 | 2.03E-06 | 6.06E-05 | T | F | T | T |
| Procr | 0.92889172 | 5.77515344 | 33.891921 | 2.04E-06 | 6.09E-05 | T | F | F | T |
| Marcks | 0.81726132 | 7.13248553 | 33.8899848 | 2.04E-06 | 6.09E-05 | T | T | T | T |
| Adcyap1r1 | 2.70123862 | 2.13310806 | 33.8762706 | 2.05E-06 | 6.10E-05 | T | F | T | T |
| Slc35d3 | 4.42254469 | 1.13967918 | 33.8410325 | 2.07E-06 | 6.14E-05 | F | F | T | F |
| Fzd10 | 1.43217372 | 4.02238569 | 33.7092129 | 2.14E-06 | 6.34E-05 | T | F | T | T |
| Dbi | -1.0307862 | 5.68459205 | 33.6935964 | 2.14E-06 | 6.35E-05 | F | F | F | F |
| Rps27 | -3.5066641 | 3.26093619 | 33.6700848 | 2.16E-06 | 6.38E-05 | T | F | F | F |
| Shroom3 | 0.7992901 | 6.59114063 | 33.5091694 | 2.24E-06 | 6.63E-05 | T | F | T | T |
| Trp53 | 0.68611652 | 7.11624173 | 33.479676 | 2.26E-06 | 6.67E-05 | F | F | F | F |
| Grb2 | -0.729854 | 7.33285825 | 33.4603713 | 2.27E-06 | 6.69E-05 | T | F | T | F |
| Dsc2 | -1.1074776 | 5.54871651 | 33.4552736 | 2.27E-06 | 6.69E-05 | T | F | T | F |
| Tgm2 | -0.8012599 | 6.72394084 | 33.4446799 | 2.28E-06 | 6.70E-05 | F | F | T | T |
| Nostrin | -1.2977054 | 4.85636458 | 33.4022767 | 2.30E-06 | 6.76E-05 | F | F | F | F |
| Eif1ax | 0.9073183 | 5.85313357 | 33.3774029 | 2.32E-06 | 6.79E-05 | F | F | T | T |
| Gpr20 | -3.4634013 | 2.1554387 | 33.339599 | 2.34E-06 | 6.85E-05 | F | F | F | F |
| Cacng7 | 1.7336522 | 3.59062234 | 33.3189989 | 2.35E-06 | 6.87E-05 | F | F | F | F |
| Scd2 | 0.54266844 | 8.83642287 | 33.2567389 | 2.39E-06 | 6.97E-05 | T | F | T | T |
| Capns1 | -0.837966 | 6.53195599 | 33.1391413 | 2.46E-06 | 7.17E-05 | T | F | T | T |
| Sgpp1 | -1.1541839 | 5.15697944 | 33.0583219 | 2.51E-06 | 7.30E-05 | T | T | F | T |
| Rtl8c | 2.59960513 | 2.18363512 | 33.0355664 | 2.52E-06 | 7.33E-05 | F | F | F | F |
| Nfkb1 | 1.25289116 | 4.41945819 | 33.0029765 | 2.54E-06 | 7.38E-05 | T | F | F | T |
| Atp2a3 | -1.4241443 | 4.36201228 | 32.8952255 | 2.61E-06 | 7.57E-05 | T | F | F | T |
| Lhfpl2 | 1.10311518 | 5.05930811 | 32.8616042 | 2.63E-06 | 7.63E-05 | T | T | T | T |
| Dpysl3 | 1.96763829 | 3.00409455 | 32.7969479 | 2.68E-06 | 7.74E-05 | F | F | T | T |
| Wdr86 | 1.77352288 | 3.37404289 | 32.7839508 | 2.69E-06 | 7.76E-05 | F | F | F | T |
| Sptbn2 | -0.8814127 | 7.03304077 | 32.7708703 | 2.70E-06 | 7.77E-05 | F | F | T | F |
| Arl4c | 0.87993283 | 5.93788504 | 32.6421806 | 2.78E-06 | 8.01E-05 | F | F | T | F |

| | | | | | | | | | |
|---|---|---|---|---|---|---|---|---|---|
| Gdf1 | 2.73494121 | 2.07138398 | 32.6247773 | 2.80E-06 | 8.04E-05 | T | F | F | F |
| Itgav | 0.68086545 | 7.05616876 | 32.5569506 | 2.84E-06 | 8.17E-05 | T | F | T | F |
| Dnm2 | -0.6428099 | 7.7890678 | 32.4865994 | 2.89E-06 | 8.30E-05 | F | F | T | F |
| Grin2c | 2.72453091 | 2.05642819 | 32.4750074 | 2.90E-06 | 8.31E-05 | F | F | F | F |
| Ptov1 | 0.74055967 | 6.75816223 | 32.468257 | 2.91E-06 | 8.32E-05 | F | F | T | F |
| Zhx2 | 2.24898289 | 2.63788758 | 32.4605714 | 2.91E-06 | 8.32E-05 | F | F | F | F |
| Evx1 | 1.8772128 | 3.98671691 | 32.3913875 | 2.96E-06 | 8.46E-05 | T | F | T | F |
| Dsg2 | -0.6380476 | 7.99719787 | 32.3715057 | 2.98E-06 | 8.49E-05 | T | F | T | F |
| Nup62cl | -0.9886866 | 5.97381524 | 32.3089513 | 3.03E-06 | 8.61E-05 | F | F | F | F |
| Nr4a3 | 2.68441464 | 2.1736464 | 32.2806359 | 3.05E-06 | 8.66E-05 | F | F | T | T |
| Patz1 | 0.79330671 | 6.41439334 | 32.2590639 | 3.06E-06 | 8.70E-05 | T | F | T | T |
| Psat1 | -0.5912793 | 8.37847712 | 32.2331758 | 3.08E-06 | 8.74E-05 | T | F | T | T |
| Klf6 | -0.8213046 | 7.22027509 | 32.1857062 | 3.12E-06 | 8.84E-05 | T | F | F | T |
| Lrp11 | 1.37803401 | 4.07088219 | 32.1787636 | 3.13E-06 | 8.84E-05 | T | F | T | T |
| Gpc6 | 1.66080837 | 3.49499951 | 32.1387787 | 3.16E-06 | 8.92E-05 | T | F | T | T |
| Arhgap10 | -0.7779697 | 6.78966541 | 32.1217032 | 3.17E-06 | 8.94E-05 | F | F | T | F |
| Anxa7 | -0.7562972 | 6.86614716 | 32.1205329 | 3.17E-06 | 8.94E-05 | T | F | T | T |
| Kcnk6 | -0.8880689 | 6.22231418 | 32.1060447 | 3.18E-06 | 8.96E-05 | T | F | T | F |
| Slc7a5 | -0.5902036 | 8.38228584 | 32.0981696 | 3.19E-06 | 8.96E-05 | F | F | T | F |
| Pbx3 | 1.1093431 | 4.82169928 | 32.0808169 | 3.21E-06 | 8.98E-05 | T | F | T | T |
| Casz1 | -0.9006778 | 6.16421178 | 32.0790522 | 3.21E-06 | 8.98E-05 | F | F | T | F |
| Las1l | -0.6863667 | 7.64077977 | 32.072383 | 3.21E-06 | 8.99E-05 | T | F | F | T |
| Wdr90 | 1.12074695 | 5.14057546 | 32.018074 | 3.26E-06 | 9.10E-05 | F | F | F | F |
| Acta2 | -2.877812 | 3.88323553 | 31.9355075 | 3.33E-06 | 9.28E-05 | F | F | T | F |
| Mn1 | 1.04285378 | 5.1142301 | 31.8933035 | 3.36E-06 | 9.37E-05 | T | F | T | F |
| Adra2b | 1.14125634 | 4.70737613 | 31.863373 | 3.39E-06 | 9.43E-05 | T | F | F | T |
| Flywch1 | 0.90851406 | 5.75503329 | 31.7439452 | 3.49E-06 | 9.69E-05 | F | F | T | F |
| Prps2 | 1.08884125 | 4.88390605 | 31.7423304 | 3.49E-06 | 9.69E-05 | T | F | T | T |
| Slc16a3 | 0.55925874 | 8.3824305 | 31.6873728 | 3.54E-06 | 9.81E-05 | T | F | F | T |
| Gtf2ird1 | 0.92231143 | 5.65243767 | 31.6838281 | 3.54E-06 | 9.81E-05 | T | F | T | F |
| Zfp217 | 0.85314209 | 6.13857014 | 31.660088 | 3.57E-06 | 9.86E-05 | T | F | F | T |
| Plk2 | -1.4464817 | 4.25988591 | 31.6302976 | 3.59E-06 | 9.92E-05 | F | F | F | F |
| N4bp2 | -0.6558576 | 8.25827318 | 31.6136063 | 3.61E-06 | 9.95E-05 | F | F | F | T |
| Lipg | -0.7423957 | 7.08017233 | 31.6097389 | 3.61E-06 | 9.95E-05 | T | F | T | F |
| Myh10 | 0.5618893 | 8.50425697 | 31.602914 | 3.62E-06 | 9.95E-05 | T | F | T | F |
| Adamts4 | -1.4223876 | 4.37751634 | 31.601698 | 3.62E-06 | 9.95E-05 | F | F | F | F |
| Snx9 | -0.70721 | 7.48052082 | 31.5915817 | 3.63E-06 | 9.96E-05 | F | F | T | F |
| Cdca7l | 1.04536724 | 5.14488373 | 31.5638742 | 3.65E-06 | 0.0001 | T | F | F | T |
| AU015836 | -1.490867 | 4.59483683 | 31.234668 | 3.97E-06 | 0.000109 | T | F | F | T |
| Dock8 | -0.9157651 | 5.98575388 | 31.1686343 | 4.04E-06 | 0.00011 | T | F | F | T |
| Ninj1 | -1.00368 | 5.6007161 | 31.1667928 | 4.04E-06 | 0.00011 | F | F | F | F |
| Hspb1 | -0.7608296 | 7.06858752 | 31.1399165 | 4.07E-06 | 0.000111 | T | F | F | T |

| | | | | | | | | | |
|---|---|---|---|---|---|---|---|---|---|
| Irx4 | -2.0254378 | 4.79789369 | 31.1249198 | 4.09E-06 | 0.000111 | T | F | T | F |
| Nars | -0.6938675 | 7.48226747 | 31.10836 | 4.11E-06 | 0.000112 | F | F | F | F |
| Gadd45g | -1.28434 | 4.59485503 | 31.0985202 | 4.12E-06 | 0.000112 | F | F | T | F |
| Ccnjl | 1.83052569 | 3.47576996 | 31.0929885 | 4.12E-06 | 0.000112 | F | F | F | F |
| Optn | -1.3887501 | 4.34715615 | 30.997786 | 4.22E-06 | 0.000114 | F | F | T | T |
| Ptpn14 | 0.69603485 | 6.86529841 | 30.9874257 | 4.24E-06 | 0.000115 | T | F | T | T |
| Gpr161 | 2.07140261 | 2.78250917 | 30.941618 | 4.29E-06 | 0.000116 | F | F | T | T |
| Igsf10 | 2.00120226 | 2.82611535 | 30.8727608 | 4.36E-06 | 0.000118 | F | F | T | F |
| Gfod1 | -2.0131013 | 3.3140126 | 30.8691108 | 4.37E-06 | 0.000118 | T | F | F | F |
| Magi1 | 0.92916059 | 5.56183222 | 30.8200286 | 4.42E-06 | 0.000119 | T | F | T | T |
| Smpdl3b | -1.4987473 | 4.12044521 | 30.8067409 | 4.44E-06 | 0.000119 | F | F | F | F |
| Ank2 | 0.90320764 | 5.77412575 | 30.8006141 | 4.44E-06 | 0.000119 | T | F | T | T |
| Tmod3 | -0.7060413 | 7.09550167 | 30.7708143 | 4.48E-06 | 0.00012 | T | F | T | T |
| Hbb-bh1 | -4.619652 | 5.87026242 | 30.7504942 | 4.50E-06 | 0.000121 | F | F | F | F |
| Rab8b | -0.9445081 | 5.90923072 | 30.74845 | 4.50E-06 | 0.000121 | T | F | T | F |
| Tpm4 | -0.6398956 | 8.53438876 | 30.6314913 | 4.64E-06 | 0.000124 | T | F | T | T |
| Zfp36 | -1.2205802 | 4.83189831 | 30.5664845 | 4.72E-06 | 0.000126 | F | F | F | F |
| Crabp1 | 2.10992982 | 2.93806887 | 30.5127201 | 4.79E-06 | 0.000128 | F | F | F | F |
| Reg2 | -0.7412052 | 6.85175017 | 30.4863303 | 4.82E-06 | 0.000128 | F | F | F | F |
| Tnfrsf1b | -1.1501637 | 5.0001162 | 30.4806294 | 4.83E-06 | 0.000128 | T | F | T | F |
| Mns1 | 1.05487725 | 5.0758363 | 30.4449665 | 4.87E-06 | 0.000129 | F | F | T | F |
| Smagp | -1.1383508 | 5.0174206 | 30.416232 | 4.91E-06 | 0.00013 | F | F | T | F |
| Plxna3 | 1.50499143 | 3.77137143 | 30.4101691 | 4.92E-06 | 0.00013 | T | F | T | F |
| Srgn | 1.30521858 | 4.81818842 | 30.4088441 | 4.92E-06 | 0.00013 | F | F | F | T |
| Hmcn1 | 2.30857164 | 2.38485934 | 30.3035831 | 5.05E-06 | 0.000134 | T | T | T | T |
| Sorl1 | 0.8263093 | 6.13148901 | 30.2548353 | 5.12E-06 | 0.000135 | F | F | T | T |
| Lama5 | 0.54395041 | 8.56536074 | 30.2089154 | 5.18E-06 | 0.000137 | F | F | F | F |
| Tmc7 | 1.60740176 | 3.49369084 | 30.192187 | 5.20E-06 | 0.000137 | T | F | T | T |
| Psg22 | 1.42630021 | 4.50305531 | 30.173528 | 5.23E-06 | 0.000138 | T | F | F | T |
| Dkkl1 | -2.2480063 | 2.98035668 | 30.1588708 | 5.25E-06 | 0.000138 | F | F | F | F |
| Gabbr1 | 2.05253972 | 2.70930609 | 30.1550032 | 5.25E-06 | 0.000138 | T | F | T | T |
| Ripor1 | -0.8379388 | 6.2873725 | 30.1323282 | 5.29E-06 | 0.000139 | T | F | F | F |
| Ugp2 | -1.034906 | 5.87844072 | 30.1290209 | 5.29E-06 | 0.000139 | F | F | T | T |
| Ifitm2 | 1.14473694 | 4.62388422 | 30.1131503 | 5.31E-06 | 0.000139 | F | F | F | T |
| Cyb5b | -0.6131575 | 8.01795222 | 30.1088908 | 5.32E-06 | 0.000139 | T | F | T | F |
| Rxrg | 3.90633638 | 1.18408777 | 30.0965067 | 5.34E-06 | 0.000139 | F | F | F | F |
| Slc7a1 | -0.7340109 | 7.18696446 | 30.073987 | 5.37E-06 | 0.00014 | T | F | F | F |
| Chst3 | 2.5056649 | 2.17086352 | 29.948806 | 5.55E-06 | 0.000144 | T | F | T | F |
| Slc4a8 | 1.50442576 | 4.21991351 | 29.8833594 | 5.64E-06 | 0.000147 | T | F | T | T |
| Kif7 | 1.46147875 | 3.7749695 | 29.8770369 | 5.65E-06 | 0.000147 | F | F | F | F |
| Tceal8 | 0.83520439 | 5.97611381 | 29.846504 | 5.70E-06 | 0.000148 | F | F | F | F |
| Atm | 0.74256704 | 6.57234827 | 29.8323387 | 5.72E-06 | 0.000148 | T | F | F | T |

| | | | | | | | | | |
|---|---|---|---|---|---|---|---|---|---|
| Galnt6 | -2.2341035 | 3.04783445 | 29.7965315 | 5.77E-06 | 0.000149 | T | F | T | T |
| Ubb | 0.48413555 | 9.4838798 | 29.7912566 | 5.78E-06 | 0.000149 | F | F | F | F |
| Gsta4 | 1.31003515 | 4.13752672 | 29.7759926 | 5.80E-06 | 0.00015 | F | F | F | F |
| Selenon | 0.99884877 | 5.38625855 | 29.7634586 | 5.82E-06 | 0.00015 | T | T | T | T |
| Dgkk | 0.96922748 | 5.50403972 | 29.7547414 | 5.83E-06 | 0.00015 | T | F | T | T |
| Cyp11a1 | 1.43591791 | 4.35318286 | 29.7519948 | 5.84E-06 | 0.00015 | T | F | F | F |
| Eps8 | -0.7897027 | 6.51019772 | 29.7334875 | 5.87E-06 | 0.000151 | F | F | F | T |
| Gng12 | -0.8310458 | 6.31676421 | 29.715318 | 5.90E-06 | 0.000151 | T | F | T | T |
| Pfn1 | -0.4950229 | 9.5656686 | 29.6689417 | 5.97E-06 | 0.000153 | T | F | F | F |
| Dach1 | 5.2434641 | 0.89023254 | 29.6414271 | 6.01E-06 | 0.000154 | T | F | T | F |
| Gabrb3 | 2.8295467 | 1.8726253 | 29.6091172 | 6.06E-06 | 0.000155 | T | T | T | T |
| Cited4 | -1.8283732 | 4.63277912 | 29.5895163 | 6.09E-06 | 0.000155 | F | F | F | F |
| Syngap1 | 1.21903582 | 4.32843275 | 29.5880864 | 6.10E-06 | 0.000155 | F | F | F | T |
| Tal1 | -1.1811642 | 5.14600186 | 29.5866033 | 6.10E-06 | 0.000155 | T | T | T | T |
| Asns | -0.7028059 | 7.00788873 | 29.4806838 | 6.27E-06 | 0.00016 | F | F | F | T |
| Gm14698 | -8.9091318 | 1.30283639 | 29.462964 | 6.30E-06 | 0.00016 | F | F | F | T |
| Sphkap | 8.71952997 | 0.19779869 | 29.8457472 | 6.31E-06 | 0.00016 | T | F | F | F |
| Sap30 | 1.42389922 | 3.86594902 | 29.4486746 | 6.32E-06 | 0.00016 | F | F | T | F |
| Pcnx2 | 4.45406723 | 0.90367608 | 29.4261465 | 6.36E-06 | 0.000161 | F | F | T | F |
| Psip1 | 0.61364001 | 7.47991316 | 29.4247208 | 6.36E-06 | 0.000161 | T | T | T | F |
| Actr2 | -0.5799357 | 8.2831165 | 29.4186332 | 6.38E-06 | 0.000161 | T | T | T | T |
| Maneal | 4.88254151 | 0.84352883 | 29.4060086 | 6.40E-06 | 0.000161 | F | F | T | F |
| Stat3 | -0.7689736 | 6.59764891 | 29.3962981 | 6.41E-06 | 0.000162 | F | F | T | T |
| Senp1 | -0.6697535 | 7.27579041 | 29.3918285 | 6.42E-06 | 0.000162 | T | F | T | T |
| Slit2 | 2.00870142 | 2.96552248 | 29.3912098 | 6.42E-06 | 0.000162 | T | F | F | T |
| Ndrg1 | -0.7525399 | 6.6823922 | 29.3332034 | 6.52E-06 | 0.000164 | T | F | T | F |
| Pcdhgc3 | 0.98993734 | 5.22683832 | 29.3247342 | 6.54E-06 | 0.000164 | F | F | F | F |
| Msi2 | 0.85505689 | 6.0396769 | 29.2706664 | 6.63E-06 | 0.000166 | T | T | T | T |
| Fras1 | 0.84854709 | 6.04966293 | 29.2539583 | 6.66E-06 | 0.000167 | F | F | F | F |
| Gm4737 | -1.1875585 | 5.14168139 | 29.2434161 | 6.68E-06 | 0.000167 | F | F | T | F |
| Mid1ip1 | 1.28645711 | 4.16922183 | 29.2261836 | 6.71E-06 | 0.000168 | T | F | F | F |
| Galnt10 | 0.82725073 | 5.99113987 | 29.1936045 | 6.77E-06 | 0.000169 | F | F | F | F |
| Slc12a2 | 0.90669885 | 5.55092827 | 29.1719099 | 6.81E-06 | 0.00017 | T | F | T | F |
| Mybl2 | -0.5706078 | 8.21191149 | 29.1375715 | 6.87E-06 | 0.000171 | F | F | F | F |
| Amotl2 | 0.61210217 | 7.39593917 | 29.084223 | 6.97E-06 | 0.000173 | F | F | T | F |
| Sh3pxd2b | 0.785366 | 6.20399615 | 29.0459497 | 7.04E-06 | 0.000175 | T | F | F | T |
| Tmem59l | 2.23275119 | 2.3939104 | 28.9743807 | 7.17E-06 | 0.000178 | T | F | F | T |
| Samd3 | 5.81393336 | 0.49730868 | 28.952575 | 7.21E-06 | 0.000179 | T | F | T | F |
| Arhgap32 | 1.11620525 | 4.84263799 | 28.9445924 | 7.23E-06 | 0.000179 | F | F | T | T |
| Olfr1506 | -8.9994662 | 1.31048777 | 28.9319865 | 7.25E-06 | 0.000179 | F | F | F | F |
| Cbfa2t3 | -1.1047415 | 5.02815595 | 28.9021707 | 7.31E-06 | 0.000181 | F | F | T | F |
| Nanog | 2.28086936 | 2.82499096 | 28.865598 | 7.38E-06 | 0.000182 | F | F | F | F |

| | | | | | | | | | |
|---|---|---|---|---|---|---|---|---|---|
| Lin28a | 0.46057794 | 9.74689659 | 28.8131834 | 7.49E-06 | 0.000184 | F | F | F | F |
| Klhl14 | 8.69052353 | 0.17181286 | 29.1478327 | 7.56E-06 | 0.000186 | F | F | T | T |
| Adcy6 | 0.96045328 | 5.30033665 | 28.7436159 | 7.63E-06 | 0.000188 | F | F | T | F |
| Nectin2 | 0.5921605 | 7.57813311 | 28.7194241 | 7.68E-06 | 0.000188 | T | F | F | F |
| Lrpap1 | 0.61202289 | 7.37258571 | 28.7159584 | 7.68E-06 | 0.000188 | T | F | T | T |
| Rasl11a | 2.32796035 | 2.273564 | 28.7083249 | 7.70E-06 | 0.000189 | F | F | F | F |
| Sgpl1 | 0.6324732 | 7.12824626 | 28.6816043 | 7.76E-06 | 0.00019 | T | F | T | F |
| Prl3d3 | 2.11112356 | 2.70492797 | 28.649945 | 7.82E-06 | 0.000191 | F | F | F | F |
| Srrm4 | 3.69454324 | 1.26573272 | 28.6363232 | 7.85E-06 | 0.000192 | T | F | T | T |
| Trim25 | -0.618595 | 7.677507 | 28.5496043 | 8.04E-06 | 0.000196 | T | F | T | T |
| Asap2 | 1.14665861 | 4.52412406 | 28.5363474 | 8.06E-06 | 0.000196 | F | F | F | F |
| Ephb3 | 0.84975428 | 5.91163706 | 28.5120076 | 8.12E-06 | 0.000198 | F | F | F | F |
| H3f3a | 0.58670695 | 7.67745633 | 28.5057513 | 8.13E-06 | 0.000198 | T | F | F | F |
| Cln6 | -0.9295308 | 5.76951295 | 28.4414119 | 8.27E-06 | 0.000201 | F | F | T | F |
| Tnpo2 | -0.6544471 | 7.24350055 | 28.4132658 | 8.33E-06 | 0.000202 | T | F | T | F |
| Eda | 1.57809092 | 3.45649182 | 28.4027048 | 8.36E-06 | 0.000203 | T | F | T | F |
| Frem1 | 1.54561092 | 3.59301136 | 28.3959151 | 8.37E-06 | 0.000203 | T | F | F | F |
| Gna14 | 2.92995829 | 1.77561154 | 28.3803654 | 8.41E-06 | 0.000203 | T | F | T | T |
| Ndrg2 | 2.46017344 | 2.11939192 | 28.3432504 | 8.49E-06 | 0.000205 | T | F | T | F |
| Foxo3 | 0.84909449 | 5.78855938 | 28.2673907 | 8.67E-06 | 0.000209 | F | F | T | T |
| Parp4 | -1.1227395 | 4.94191602 | 28.2588868 | 8.69E-06 | 0.000209 | F | F | F | F |
| Limd2 | 0.95922527 | 5.25395403 | 28.2538548 | 8.70E-06 | 0.000209 | T | F | T | F |
| Sept9 | -0.5377883 | 8.68574801 | 28.2279683 | 8.76E-06 | 0.000211 | F | F | T | F |
| Nell2 | 3.82461184 | 1.21346103 | 28.2174847 | 8.79E-06 | 0.000211 | F | F | F | T |
| Apba2 | 3.63926506 | 1.37654179 | 28.2116894 | 8.80E-06 | 0.000211 | F | F | F | F |
| Capn1 | -0.7259049 | 6.78040894 | 28.2082112 | 8.81E-06 | 0.000211 | F | F | F | F |
| Lgals3 | -0.8949264 | 5.89026192 | 28.1741785 | 8.89E-06 | 0.000213 | T | F | F | F |
| Klhl23 | 1.83058605 | 2.95386459 | 28.1216978 | 9.02E-06 | 0.000216 | T | F | T | T |
| Rnf165 | 2.29589415 | 2.38641037 | 28.0543876 | 9.18E-06 | 0.000219 | F | F | T | T |
| Fam172a | -1.44664 | 4.32600763 | 28.0390616 | 9.22E-06 | 0.00022 | T | T | T | T |
| Hist1h2bq | -0.9102372 | 8.13730771 | 28.0096238 | 9.29E-06 | 0.000221 | F | F | T | F |
| Camkv | 2.64391657 | 1.99754671 | 27.9994597 | 9.32E-06 | 0.000222 | T | F | T | F |
| Rhox6 | -1.0293716 | 5.29567525 | 27.9769109 | 9.38E-06 | 0.000223 | F | F | T | F |
| Stxbp6 | 1.86692228 | 2.97839781 | 27.9644245 | 9.41E-06 | 0.000223 | T | T | T | T |
| mt-Nd3 | -0.7481403 | 6.78380102 | 27.8651901 | 9.67E-06 | 0.000229 | F | F | F | F |
| Plod2 | 0.78850802 | 6.29267173 | 27.7697435 | 9.92E-06 | 0.000235 | T | F | F | T |
| Plcd3 | -1.3404679 | 4.25406541 | 27.6543181 | 1.02E-05 | 0.000242 | F | F | F | F |
| Krt18 | -0.4956662 | 9.20316607 | 27.6297543 | 1.03E-05 | 0.000244 | F | F | F | F |
| Specc1 | -0.7708593 | 6.44873723 | 27.6042137 | 1.04E-05 | 0.000245 | T | T | F | T |
| Thsd7a | 2.60488418 | 1.95027965 | 27.5974619 | 1.04E-05 | 0.000245 | F | F | T | T |
| Zfp57 | 1.39710165 | 3.7987648 | 27.5784389 | 1.05E-05 | 0.000246 | F | F | F | F |
| Gjb2 | -1.2594645 | 4.51295598 | 27.5578911 | 1.05E-05 | 0.000247 | T | F | T | F |

| | | | | | | | | | |
|---|---|---|---|---|---|---|---|---|---|
| Psd4 | -1.1289527 | 4.81428423 | 27.5480072 | 1.05E-05 | 0.000248 | T | F | F | T |
| Sult4a1 | 1.90645193 | 3.05481551 | 27.5024704 | 1.07E-05 | 0.000251 | T | F | T | T |
| Hmga1 | 0.55015822 | 8.03656083 | 27.4936279 | 1.07E-05 | 0.000251 | F | F | F | F |
| Ikzf1 | -1.8848707 | 3.87461586 | 27.4849874 | 1.07E-05 | 0.000251 | T | F | F | T |
| Vill | -1.5883155 | 4.22397649 | 27.4845655 | 1.07E-05 | 0.000251 | F | F | F | F |
| Nrep | 2.22615708 | 2.76718094 | 27.4587965 | 1.08E-05 | 0.000253 | F | F | T | F |
| Anxa6 | -0.8222341 | 6.29261302 | 27.3874126 | 1.10E-05 | 0.000257 | F | F | F | F |
| Il2rg | -1.395286 | 4.12272417 | 27.3651268 | 1.11E-05 | 0.000259 | F | F | F | F |
| Rhox9 | -1.2672461 | 4.41022838 | 27.3441974 | 1.11E-05 | 0.00026 | F | F | F | F |
| Tmem198b | 1.97345473 | 2.69553963 | 27.3369581 | 1.12E-05 | 0.00026 | F | F | T | F |
| Ptbp3 | -0.6818151 | 7.24256178 | 27.3022231 | 1.13E-05 | 0.000262 | T | T | T | T |
| Ube2a | -0.9119278 | 5.93727192 | 27.2768229 | 1.14E-05 | 0.000264 | T | F | F | T |
| Prex2 | -1.0218049 | 5.31973324 | 27.256249 | 1.14E-05 | 0.000265 | F | F | T | T |
| Atxn10 | -0.5762947 | 7.99728396 | 27.2541787 | 1.14E-05 | 0.000265 | F | F | T | T |
| Kcnc3 | 1.78445428 | 3.01369806 | 27.2274539 | 1.15E-05 | 0.000267 | T | F | F | F |
| Zfp184 | 1.99288045 | 2.6419309 | 27.1551587 | 1.17E-05 | 0.000272 | F | F | F | F |
| Ccdc8 | 1.11876651 | 4.47989899 | 27.1466722 | 1.18E-05 | 0.000272 | F | F | F | F |
| Calml4 | -8.5491007 | 0.92284317 | 27.1266905 | 1.18E-05 | 0.000273 | F | F | T | T |
| Fbxl20 | 1.49603004 | 3.60466794 | 27.1219374 | 1.18E-05 | 0.000273 | T | F | T | T |
| Parva | -0.8945928 | 5.99505682 | 27.1125975 | 1.19E-05 | 0.000274 | T | F | T | T |
| Maged1 | 0.55804224 | 7.79917227 | 27.0568373 | 1.21E-05 | 0.000278 | T | F | F | F |
| Igf1r | 0.65576654 | 7.09725746 | 27.0479479 | 1.21E-05 | 0.000278 | T | F | T | T |
| Cul4b | -0.5892975 | 7.87631948 | 26.9936623 | 1.23E-05 | 0.000282 | F | F | T | T |
| Sesn3 | 0.79139049 | 6.09435888 | 26.9611176 | 1.24E-05 | 0.000284 | T | F | F | T |
| Rarg | 1.02076859 | 4.99492867 | 26.9610142 | 1.24E-05 | 0.000284 | F | F | T | F |
| Adgrf5 | -0.5607624 | 7.99441429 | 26.9550643 | 1.24E-05 | 0.000284 | T | F | T | T |
| Tmem132c | 2.79380374 | 2.06528459 | 26.9491401 | 1.24E-05 | 0.000284 | T | F | F | F |
| Ptprm | 1.81644916 | 2.91474728 | 26.9361285 | 1.25E-05 | 0.000285 | F | F | F | F |
| Jakmip2 | 3.45295706 | 1.29807724 | 26.9305926 | 1.25E-05 | 0.000285 | T | F | T | F |
| Clic6 | -0.5299116 | 8.49957254 | 26.9294491 | 1.25E-05 | 0.000285 | T | F | T | T |
| Mapk12 | 4.89218923 | 0.92945674 | 26.9274307 | 1.25E-05 | 0.000285 | F | F | F | F |
| Lonp2 | -0.8309392 | 6.12079459 | 26.9023887 | 1.26E-05 | 0.000286 | F | F | F | T |
| Gmip | -0.9955094 | 5.34862134 | 26.9010292 | 1.26E-05 | 0.000286 | T | F | F | F |
| Ergic1 | 0.78835165 | 6.06205904 | 26.882921 | 1.27E-05 | 0.000287 | F | F | T | T |
| Prxl2a | 1.10446854 | 4.62487695 | 26.8819982 | 1.27E-05 | 0.000287 | F | F | F | F |
| Epop | -0.5261598 | 8.66005914 | 26.8407354 | 1.28E-05 | 0.00029 | F | F | T | F |
| Plpp2 | -0.8437992 | 5.99293069 | 26.8099704 | 1.29E-05 | 0.000292 | T | F | T | F |
| Ptgis | 0.9546982 | 5.1194432 | 26.7401788 | 1.32E-05 | 0.000298 | F | F | F | F |
| Hat1 | -0.6380505 | 7.41970744 | 26.7101722 | 1.33E-05 | 0.0003 | T | F | F | F |
| Arxes1 | 1.42445453 | 3.67560219 | 26.6455264 | 1.35E-05 | 0.000305 | F | T | F | T |
| Stk10 | -0.7690948 | 6.42967023 | 26.6278268 | 1.36E-05 | 0.000306 | T | F | F | T |
| Esyt1 | -0.7799171 | 6.33862643 | 26.6155118 | 1.36E-05 | 0.000307 | T | F | F | T |

| | | | | | | | | | |
|---|---|---|---|---|---|---|---|---|---|
| Plch1 | 1.0949966 | 4.52396296 | 26.576537 | 1.38E-05 | 0.00031 | T | F | T | T |
| Ankrd13a | -0.5951223 | 7.70265448 | 26.5742392 | 1.38E-05 | 0.00031 | T | F | T | T |
| Esrp2 | -0.8862026 | 6.02732725 | 26.5489207 | 1.39E-05 | 0.000312 | T | F | F | T |
| Chst7 | 2.72706148 | 2.06510858 | 26.5042602 | 1.41E-05 | 0.000315 | T | F | T | T |
| Gata6 | 1.01024954 | 5.52653286 | 26.4582528 | 1.42E-05 | 0.000319 | T | F | T | F |
| Tpbg | 1.44816475 | 3.6713689 | 26.4436646 | 1.43E-05 | 0.00032 | T | F | F | F |
| Ptma | 0.41248463 | 10.4104326 | 26.4336723 | 1.43E-05 | 0.00032 | F | F | T | F |
| Fhl3 | -0.9788939 | 5.34197149 | 26.4006542 | 1.45E-05 | 0.000323 | F | F | T | F |
| H2aw | 0.58873241 | 7.3924322 | 26.3867952 | 1.45E-05 | 0.000324 | F | F | F | F |
| Satb2 | -0.7170295 | 6.72345098 | 26.3825231 | 1.45E-05 | 0.000324 | T | T | T | T |
| Id2 | -0.6338375 | 7.35819551 | 26.3752323 | 1.46E-05 | 0.000324 | F | F | T | F |
| Bcl11a | 4.13725929 | 0.91068836 | 26.3517392 | 1.47E-05 | 0.000326 | T | F | T | F |
| Apobec3 | 1.83392488 | 2.84293977 | 26.3447179 | 1.47E-05 | 0.000326 | T | F | T | T |
| Dnah5 | 4.9913786 | 0.64687121 | 26.340208 | 1.47E-05 | 0.000327 | T | F | T | T |
| Igfbp3 | 1.69516856 | 3.26793738 | 26.3256446 | 1.48E-05 | 0.000328 | T | F | T | T |
| Mical1 | 1.14732877 | 4.33964517 | 26.263164 | 1.50E-05 | 0.000333 | T | F | F | F |
| Rragc | -0.8424132 | 5.98239143 | 26.2316131 | 1.52E-05 | 0.000335 | T | F | T | T |
| Foxn2 | -0.8268201 | 6.05752779 | 26.2296569 | 1.52E-05 | 0.000335 | T | F | T | F |
| Nid1 | 0.47885249 | 8.96711289 | 26.2235907 | 1.52E-05 | 0.000336 | T | F | F | T |
| Lonrf3 | -2.508096 | 2.58842869 | 26.1930588 | 1.53E-05 | 0.000338 | T | F | F | T |
| Plce1 | 2.14487553 | 2.4427316 | 26.1877888 | 1.54E-05 | 0.000338 | T | F | F | F |
| Hsd17b2 | -1.324453 | 4.20280558 | 26.1289365 | 1.56E-05 | 0.000344 | F | F | F | F |
| Akna | 1.27886088 | 3.98501489 | 26.1171341 | 1.57E-05 | 0.000344 | T | F | F | F |
| Rapgef3 | -2.3161093 | 2.67405657 | 26.0612501 | 1.59E-05 | 0.000349 | F | F | T | T |
| Gata1 | -1.3348014 | 4.65575753 | 26.0464667 | 1.60E-05 | 0.000351 | F | F | F | F |
| Tfcp2l1 | -0.6487753 | 7.0400425 | 26.0356897 | 1.60E-05 | 0.000351 | T | F | T | T |
| Igsf8 | -0.7577016 | 6.39008483 | 26.0190035 | 1.61E-05 | 0.000353 | F | F | F | F |
| D430041D0 | 1.84991379 | 2.82756856 | 25.9901922 | 1.62E-05 | 0.000355 | T | F | T | F |
| Rreb1 | -0.5722919 | 7.75704831 | 25.9636868 | 1.64E-05 | 0.000357 | T | F | T | F |
| Eef2k | -0.8289266 | 6.15318716 | 25.8785132 | 1.67E-05 | 0.000366 | T | F | T | T |
| Grrp1 | 2.23565042 | 2.39564054 | 25.8461867 | 1.69E-05 | 0.000368 | T | F | F | T |
| Cxcr4 | 1.7030363 | 3.35695019 | 25.845545 | 1.69E-05 | 0.000368 | T | F | F | F |
| Cyp4f15 | 4.45124651 | 0.82510091 | 25.8051204 | 1.71E-05 | 0.000372 | T | F | F | F |
| Sorcs2 | 1.65235814 | 3.24906183 | 25.8027287 | 1.71E-05 | 0.000372 | T | F | T | T |
| Fermt3 | -2.421968 | 3.23811761 | 25.7871038 | 1.72E-05 | 0.000373 | F | F | T | F |
| Unc93b1 | -1.8239945 | 3.46160373 | 25.784691 | 1.72E-05 | 0.000373 | T | F | T | F |
| Espn | -0.8644654 | 5.83438026 | 25.7736178 | 1.73E-05 | 0.000374 | F | F | F | F |
| Rab34 | 1.26388581 | 4.00251187 | 25.7435574 | 1.74E-05 | 0.000377 | F | F | F | F |
| Arid1b | 0.67900432 | 6.59257495 | 25.7418221 | 1.74E-05 | 0.000377 | T | F | T | T |
| Zcchc18 | 3.28535431 | 1.3499284 | 25.7382246 | 1.74E-05 | 0.000377 | T | F | T | T |
| Tspan5 | 1.98576931 | 2.56179833 | 25.7267003 | 1.75E-05 | 0.000378 | T | F | T | F |
| Trap1a | -0.7331005 | 6.53769904 | 25.7117867 | 1.76E-05 | 0.000379 | F | F | F | F |

| | | | | | | | | | |
|---|---|---|---|---|---|---|---|---|---|
| Itpr1 | 0.757192 | 6.1320442 | 25.6940756 | 1.76E-05 | 0.00038 | T | F | F | F |
| Arid1a | 0.46627877 | 9.07145178 | 25.6608218 | 1.78E-05 | 0.000383 | T | F | T | T |
| Scn5a | 3.68486041 | 1.29244105 | 25.6046115 | 1.81E-05 | 0.000389 | F | F | T | T |
| Ntrk3 | 2.63319134 | 1.83534158 | 25.5918352 | 1.82E-05 | 0.00039 | T | F | T | T |
| Gk5 | -1.0633698 | 4.94692641 | 25.559836 | 1.83E-05 | 0.000393 | T | F | T | T |
| Fam107a | 4.16794155 | 0.94765054 | 25.5449796 | 1.84E-05 | 0.000395 | T | F | T | T |
| Arid5b | -0.7276727 | 7.03594373 | 25.5083734 | 1.86E-05 | 0.000398 | T | F | T | T |
| Anpep | 1.05746237 | 4.63396712 | 25.4828801 | 1.87E-05 | 0.000401 | F | F | F | F |
| Sulf2 | 0.61489475 | 6.97519716 | 25.4212819 | 1.91E-05 | 0.000408 | F | F | T | T |
| Nt5dc2 | 0.53231941 | 7.96876682 | 25.3884635 | 1.92E-05 | 0.000411 | F | F | F | F |
| H2-Q4 | 1.31252558 | 3.85958464 | 25.3608206 | 1.94E-05 | 0.000414 | F | F | T | T |
| Rbm15b | 0.59660717 | 7.13452648 | 25.3422937 | 1.95E-05 | 0.000416 | T | F | F | T |
| Il1rap | -1.6040896 | 3.75115207 | 25.3263345 | 1.96E-05 | 0.000417 | T | T | T | T |
| Pfkl | 0.48566131 | 8.69286537 | 25.3182957 | 1.96E-05 | 0.000418 | F | F | F | F |
| Msh6 | 0.54619806 | 7.6949378 | 25.3016974 | 1.97E-05 | 0.000419 | T | F | F | F |
| Ace | -1.6705286 | 3.61327705 | 25.2870415 | 1.98E-05 | 0.000421 | F | F | T | F |
| Trp73 | 4.96342111 | 0.57170493 | 25.2772501 | 1.99E-05 | 0.000421 | F | F | T | F |
| Kif5a | 2.32121112 | 2.1300684 | 25.245039 | 2.00E-05 | 0.000425 | T | F | T | F |
| Cts8 | 1.62467706 | 3.77669592 | 25.236523 | 2.01E-05 | 0.000425 | T | F | F | F |
| Adam11 | 3.89343984 | 1.0591016 | 25.1874666 | 2.04E-05 | 0.000431 | T | F | F | T |
| Foxa2 | 1.09759243 | 4.46643013 | 25.1850462 | 2.04E-05 | 0.000431 | T | T | F | T |
| Junb | -0.8320855 | 6.06219295 | 25.157793 | 2.05E-05 | 0.000434 | F | F | F | F |
| Slc16a6 | -0.7887506 | 6.2063407 | 25.156066 | 2.06E-05 | 0.000434 | F | F | T | F |
| Nodal | 4.33145324 | 0.78035389 | 25.1475898 | 2.06E-05 | 0.000434 | F | F | T | F |
| Plxnb2 | 0.49979774 | 8.34298687 | 25.1336796 | 2.07E-05 | 0.000436 | F | F | F | F |
| Rell1 | -0.7547154 | 6.33425069 | 25.0933956 | 2.09E-05 | 0.00044 | F | F | T | F |
| Fbxo6 | -1.1580258 | 4.55426327 | 25.0712359 | 2.11E-05 | 0.000443 | F | F | F | F |
| Heyl | -4.947135 | 1.35000947 | 25.0165848 | 2.14E-05 | 0.000449 | T | F | F | T |
| Itih5 | -1.2625451 | 4.54978133 | 25.0017873 | 2.15E-05 | 0.000451 | T | F | T | T |
| Tmem37 | -0.8190655 | 5.97374513 | 24.9835701 | 2.16E-05 | 0.000452 | F | F | F | F |
| Apobr | -0.9235034 | 5.51724144 | 24.9694906 | 2.17E-05 | 0.000454 | T | F | F | F |
| N4bp3 | -1.0068192 | 5.11825665 | 24.9541806 | 2.18E-05 | 0.000455 | F | F | F | T |
| Sdc4 | -0.6393818 | 7.00968541 | 24.9352998 | 2.19E-05 | 0.000457 | F | F | T | F |
| Kdm7a | 1.50388754 | 3.4450848 | 24.9111972 | 2.21E-05 | 0.00046 | T | F | T | T |
| Raver2 | 1.69047478 | 3.39595556 | 24.8585503 | 2.24E-05 | 0.000467 | T | F | T | T |
| Tead4 | -0.7076072 | 6.63913996 | 24.8538406 | 2.24E-05 | 0.000467 | T | F | T | F |
| Mob1a | -0.6268895 | 7.16281743 | 24.8239615 | 2.26E-05 | 0.00047 | T | F | T | T |
| Nynrin | 0.63713427 | 6.78013619 | 24.8083081 | 2.27E-05 | 0.000472 | F | F | T | F |
| Ckap4 | 0.53942204 | 7.71914491 | 24.8069721 | 2.27E-05 | 0.000472 | T | F | F | F |
| Lrp3 | 1.2039361 | 4.15754206 | 24.7595278 | 2.30E-05 | 0.000478 | F | F | F | F |
| Sept2 | -0.5371694 | 8.05829083 | 24.7387549 | 2.32E-05 | 0.00048 | T | F | T | T |
| Ank1 | -4.0498339 | 2.3894582 | 24.7176974 | 2.33E-05 | 0.000483 | T | F | T | T |

| | | | | | | | | | |
|---|---|---|---|---|---|---|---|---|---|
| Spred2 | 0.73528656 | 6.13496536 | 24.710637 | 2.34E-05 | 0.000483 | T | F | F | F |
| Pdgfra | 0.5315167 | 7.78589182 | 24.6208176 | 2.40E-05 | 0.000496 | T | F | T | T |
| Nwd1 | -2.958675 | 2.05465926 | 24.6137393 | 2.40E-05 | 0.000496 | T | F | T | T |
| Hip1r | -0.7981826 | 6.0932454 | 24.6092697 | 2.41E-05 | 0.000496 | F | F | F | F |
| Bmpr1b | 5.32111368 | 0.59402677 | 24.6070438 | 2.41E-05 | 0.000496 | T | F | T | T |
| Twsg1 | 0.82729483 | 5.60928249 | 24.5569601 | 2.44E-05 | 0.000503 | T | F | T | T |
| Tnrc18 | -0.5337216 | 8.01012094 | 24.5489873 | 2.45E-05 | 0.000504 | T | F | F | F |
| Gm14322 | -8.501812 | 0.90926562 | 24.4899854 | 2.49E-05 | 0.000512 | F | F | T | T |
| Xlr4a | -2.0949508 | 3.03729343 | 24.471806 | 2.50E-05 | 0.000514 | T | T | F | T |
| Slco5a1 | 1.04216578 | 4.57034912 | 24.3998221 | 2.56E-05 | 0.000524 | T | F | T | T |
| Gtf2h1 | -0.6315996 | 6.9954293 | 24.392894 | 2.56E-05 | 0.000525 | T | F | F | F |
| Tox3 | 3.03758595 | 1.4786862 | 24.3658025 | 2.58E-05 | 0.000528 | T | F | T | T |
| Smad7 | -0.6390688 | 7.31487129 | 24.3217744 | 2.61E-05 | 0.000535 | T | F | T | F |
| Tmem54 | -1.160998 | 4.58171275 | 24.318013 | 2.62E-05 | 0.000535 | F | F | F | F |
| Zxdc | -0.8625809 | 5.72851602 | 24.3013759 | 2.63E-05 | 0.000537 | T | T | T | T |
| Btaf1 | -0.5758716 | 7.55535191 | 24.289604 | 2.64E-05 | 0.000538 | T | F | F | T |
| Rcn1 | 0.66791973 | 6.63382949 | 24.287615 | 2.64E-05 | 0.000538 | F | F | F | F |
| Cadps2 | 3.54286364 | 1.12098636 | 24.2858637 | 2.64E-05 | 0.000538 | T | F | F | F |
| Map2 | 2.16163903 | 2.27423184 | 24.2840262 | 2.64E-05 | 0.000538 | F | F | T | T |
| Pcdhgc4 | 8.48278157 | -0.0127632 | 24.5215073 | 2.67E-05 | 0.000543 | F | F | F | F |
| Slc30a2 | -1.0566663 | 5.05754665 | 24.2239889 | 2.69E-05 | 0.000546 | F | F | F | F |
| Card9 | -2.4815003 | 2.44490627 | 24.1929474 | 2.71E-05 | 0.00055 | F | F | F | T |
| March9 | 1.50800334 | 3.34554752 | 24.1787355 | 2.73E-05 | 0.000552 | F | F | F | F |
| Baz2b | 1.01933945 | 4.61159595 | 24.170378 | 2.73E-05 | 0.000553 | T | F | F | F |
| Gkap1 | -1.1715393 | 4.4682923 | 24.1542932 | 2.74E-05 | 0.000555 | T | F | F | F |
| Acrbp | 1.68652306 | 2.98755033 | 24.1407343 | 2.76E-05 | 0.000557 | F | F | F | F |
| Med12l | -1.0200424 | 4.97467695 | 24.1198183 | 2.77E-05 | 0.00056 | T | F | T | F |
| Ccrl2 | -2.8201698 | 2.12420659 | 24.1131464 | 2.78E-05 | 0.00056 | F | F | T | T |
| Slc35a5 | -1.595743 | 3.55804217 | 24.0363241 | 2.84E-05 | 0.000572 | T | F | F | F |
| Pxylp1 | 2.76458883 | 1.68732867 | 23.9937156 | 2.88E-05 | 0.000579 | T | F | T | F |
| Chst1 | 3.06426677 | 1.56494846 | 23.9629962 | 2.90E-05 | 0.000584 | F | F | F | F |
| Tmc5 | -2.5500073 | 2.31318318 | 23.9432425 | 2.92E-05 | 0.000587 | F | F | T | F |
| Mycl | 1.11437072 | 4.26757552 | 23.8832938 | 2.97E-05 | 0.000596 | T | F | T | F |
| Sinhcaf | 0.5748023 | 7.51722103 | 23.8675129 | 2.98E-05 | 0.000599 | F | F | T | F |
| Strbp | 0.66649193 | 6.65346047 | 23.8385964 | 3.01E-05 | 0.000603 | T | F | T | T |
| Nbl1 | 1.66926487 | 3.20750655 | 23.8201423 | 3.03E-05 | 0.000606 | T | F | T | T |
| Hs3st1 | 0.98470626 | 4.7204223 | 23.7694047 | 3.07E-05 | 0.000614 | T | F | F | T |
| Evc2 | 1.02714846 | 4.57849052 | 23.732354 | 3.10E-05 | 0.00062 | T | F | F | F |
| Errfi1 | -0.6470408 | 6.87878683 | 23.6663007 | 3.17E-05 | 0.000632 | T | F | T | T |
| Mtpn | -0.5841139 | 7.43652865 | 23.6322589 | 3.20E-05 | 0.000638 | T | F | F | T |
| Nkd1 | 0.95682456 | 5.04721687 | 23.5825293 | 3.24E-05 | 0.000647 | F | F | T | F |
| Cdk5rap2 | 0.76073214 | 6.095622 | 23.5763702 | 3.25E-05 | 0.000647 | F | F | T | T |

| | | | | | | | | | |
|---|---|---|---|---|---|---|---|---|---|
| Kdm1b | -1.0344603 | 5.00246972 | 23.5744134 | 3.25E-05 | 0.000647 | T | F | T | T |
| Amer1 | 0.83925199 | 5.44146702 | 23.533507 | 3.29E-05 | 0.000654 | T | F | T | T |
| Tanc2 | -0.7101118 | 6.60470033 | 23.520485 | 3.30E-05 | 0.000656 | T | F | T | T |
| Msantd3 | 1.10263483 | 4.28372043 | 23.5183643 | 3.31E-05 | 0.000656 | F | F | T | T |
| Zfp105 | 1.37238703 | 3.63178096 | 23.4545822 | 3.37E-05 | 0.000668 | F | F | F | F |
| Fabp3 | -0.7366466 | 6.86284137 | 23.4436894 | 3.38E-05 | 0.000669 | F | F | T | F |
| Fam171a1 | -0.6634484 | 6.77344276 | 23.4327574 | 3.39E-05 | 0.000671 | F | F | T | F |
| Rab35 | -0.6447943 | 6.82543521 | 23.4289599 | 3.40E-05 | 0.000671 | F | F | T | F |
| Car7 | -0.6043138 | 7.08385153 | 23.4133332 | 3.41E-05 | 0.000674 | F | F | F | F |
| Cobll1 | -0.6493419 | 6.79455143 | 23.3893792 | 3.44E-05 | 0.000678 | T | F | F | T |
| Aldh1a2 | 2.76116623 | 1.61674192 | 23.3718922 | 3.45E-05 | 0.000681 | F | F | F | F |
| Vasp | -0.5738341 | 7.90336236 | 23.3169798 | 3.51E-05 | 0.000691 | F | F | T | T |
| Parp1 | 0.47477511 | 8.77503156 | 23.3051831 | 3.52E-05 | 0.000693 | F | F | F | T |
| Nppb | 1.40525034 | 3.61787386 | 23.3011372 | 3.53E-05 | 0.000693 | F | F | F | F |
| Pip5k1c | -0.5158046 | 8.08466343 | 23.257198 | 3.57E-05 | 0.000702 | F | F | F | F |
| Bend3 | 0.52654677 | 7.66901012 | 23.2224806 | 3.61E-05 | 0.000708 | T | F | T | T |
| Aprt | -0.5329008 | 7.85781118 | 23.1957967 | 3.64E-05 | 0.000713 | F | F | F | F |
| Dnaja2 | -0.5491651 | 7.89658513 | 23.1845588 | 3.65E-05 | 0.000715 | T | F | T | T |
| Nid2 | 0.48787369 | 8.2182314 | 23.1810721 | 3.65E-05 | 0.000715 | T | F | F | T |
| Mutyh | 1.9257228 | 2.52420393 | 23.1487073 | 3.69E-05 | 0.000721 | F | F | F | F |
| B3gnt3 | -1.0167713 | 4.89493157 | 23.1271616 | 3.71E-05 | 0.000725 | F | F | T | F |
| Anln | -0.5437838 | 7.79055573 | 23.0830816 | 3.76E-05 | 0.000734 | T | T | T | T |
| Fsd1 | 2.18268202 | 2.15116747 | 23.031936 | 3.82E-05 | 0.000745 | F | F | F | F |
| Litaf | -0.7061964 | 6.597944 | 23.0281526 | 3.82E-05 | 0.000745 | T | F | T | T |
| Dnajb5 | 2.27358608 | 2.06690739 | 23.0130145 | 3.84E-05 | 0.000748 | F | F | T | T |
| Pcsk5 | 1.89697035 | 2.64837481 | 22.9926408 | 3.86E-05 | 0.000751 | T | T | T | T |
| Tmem28 | 1.7799951 | 2.83792492 | 22.9483991 | 3.92E-05 | 0.000761 | F | F | T | F |
| Nlgn3 | 1.53486673 | 3.34437892 | 22.9324252 | 3.93E-05 | 0.000764 | F | F | T | T |
| Olfml2a | 3.1581111 | 1.31940665 | 22.9225744 | 3.95E-05 | 0.000765 | F | F | F | T |
| Fam163b | 6.30078802 | 0.17322015 | 22.844743 | 4.04E-05 | 0.000783 | F | F | T | F |
| Adamts7 | 1.25202535 | 3.94409875 | 22.7938919 | 4.10E-05 | 0.000794 | F | F | F | T |
| Il17rd | 1.04742407 | 5.43149993 | 22.7873548 | 4.11E-05 | 0.000795 | T | F | T | T |
| Mast4 | 1.01212485 | 4.60557659 | 22.7192012 | 4.19E-05 | 0.00081 | T | F | F | F |
| Slk | -0.5812462 | 7.23398343 | 22.7148744 | 4.20E-05 | 0.000811 | T | F | F | T |
| Ift172 | 1.10400567 | 4.2225849 | 22.7029238 | 4.21E-05 | 0.000813 | F | F | F | F |
| Clcn2 | 1.08374521 | 4.28895273 | 22.6823959 | 4.24E-05 | 0.000817 | F | F | F | F |
| Htra1 | 0.75408763 | 5.83564249 | 22.6813019 | 4.24E-05 | 0.000817 | F | F | T | F |
| Six4 | 1.04786212 | 4.64731183 | 22.669781 | 4.26E-05 | 0.000819 | T | F | T | T |
| Zswim4 | 0.94013822 | 4.92211404 | 22.6457875 | 4.29E-05 | 0.000824 | F | F | T | F |
| Khdrbs3 | 1.02241535 | 4.47856324 | 22.5774318 | 4.38E-05 | 0.00084 | T | T | F | T |
| Nr1h5 | -6.6104334 | 2.16141097 | 22.5450094 | 4.42E-05 | 0.000848 | F | F | T | F |
| Ptch2 | 1.59535497 | 3.04020165 | 22.5387925 | 4.43E-05 | 0.000849 | F | F | T | F |

| | | | | | | | | | |
|---|---|---|---|---|---|---|---|---|---|
| Fam171b | 3.15320977 | 1.26470722 | 22.5138441 | 4.46E-05 | 0.000854 | T | F | T | T |
| Hcls1 | -2.6527547 | 2.14949393 | 22.4870514 | 4.50E-05 | 0.00086 | T | F | F | F |
| Tle4 | 0.86978984 | 5.15523204 | 22.4748056 | 4.51E-05 | 0.000863 | T | T | F | T |
| Myadm | -0.5169285 | 8.16177571 | 22.4392972 | 4.56E-05 | 0.000871 | F | F | F | T |
| Shmt2 | -0.5155524 | 7.93444326 | 22.4215757 | 4.59E-05 | 0.000875 | F | F | T | T |
| Fam13c | 4.56947824 | 0.83557769 | 22.3827136 | 4.64E-05 | 0.000885 | T | F | F | F |
| Chst2 | -0.9541073 | 5.10368411 | 22.368278 | 4.66E-05 | 0.000888 | T | F | T | T |
| Thy1 | 2.74947754 | 1.60884917 | 22.3436526 | 4.69E-05 | 0.000894 | T | F | F | T |
| Mycn | 0.60052395 | 6.96914777 | 22.329475 | 4.72E-05 | 0.000897 | T | F | F | F |
| Myh3 | -3.1436802 | 1.85066429 | 22.2990583 | 4.76E-05 | 0.000904 | F | F | F | F |
| Rtl6 | 1.24153249 | 3.82756365 | 22.2950703 | 4.76E-05 | 0.000905 | T | F | T | F |
| Fignl2 | -0.5583047 | 7.64408061 | 22.2886551 | 4.77E-05 | 0.000906 | F | F | T | F |
| Epha2 | 0.83612042 | 5.45905387 | 22.2765948 | 4.79E-05 | 0.000908 | F | F | F | F |
| Epha7 | 2.60735281 | 1.72999199 | 22.2738857 | 4.79E-05 | 0.000908 | F | F | T | F |
| P4hb | 0.39369555 | 9.99392823 | 22.2634363 | 4.81E-05 | 0.00091 | T | T | T | T |
| Timp3 | 0.86842006 | 5.18709193 | 22.2611752 | 4.81E-05 | 0.00091 | T | F | T | T |
| Lrig1 | 0.98836824 | 4.64798338 | 22.2559573 | 4.82E-05 | 0.00091 | F | F | T | F |
| Arpc4 | -0.522136 | 7.83126046 | 22.2542983 | 4.82E-05 | 0.00091 | F | F | T | T |
| Mdfi | -0.9424171 | 5.12353201 | 22.2519068 | 4.83E-05 | 0.00091 | F | F | T | F |
| Kdm3b | 0.6073812 | 6.83809391 | 22.2378347 | 4.85E-05 | 0.000913 | T | F | T | F |
| Rhob | -0.6469822 | 6.78952704 | 22.1939723 | 4.91E-05 | 0.000925 | T | F | T | T |
| Syt7 | 1.60237884 | 3.01033408 | 22.1803706 | 4.93E-05 | 0.000928 | F | F | T | T |
| Enpp3 | -1.2908534 | 4.03758497 | 22.1751201 | 4.94E-05 | 0.000928 | T | F | F | F |
| Tmem131l | 0.9910997 | 4.65422538 | 22.1302993 | 5.01E-05 | 0.00094 | F | F | F | F |
| Rhox4b | -8.3653515 | 0.72614831 | 22.1222838 | 5.02E-05 | 0.000942 | F | F | F | F |
| Speg | 0.89227966 | 5.15226438 | 22.1182994 | 5.03E-05 | 0.000942 | F | F | T | F |
| Epb42 | -4.2894109 | 1.35942547 | 22.0907549 | 5.07E-05 | 0.000948 | T | F | F | F |
| Cs | -0.4787203 | 8.56440492 | 22.0906701 | 5.07E-05 | 0.000948 | F | F | F | F |
| Arhgap31 | -0.9964979 | 5.09319288 | 22.0562001 | 5.12E-05 | 0.000957 | T | T | T | T |
| Ube2q2 | -0.6919185 | 6.54587999 | 22.0306805 | 5.16E-05 | 0.000964 | T | F | T | T |
| Zmynd8 | 0.52936793 | 7.5344092 | 22.0282252 | 5.17E-05 | 0.000964 | F | F | T | F |
| Tcaf1 | 1.08886992 | 4.31962332 | 22.0154095 | 5.19E-05 | 0.000967 | F | F | T | F |
| S100a6 | -0.8522805 | 5.54770305 | 22.0099734 | 5.19E-05 | 0.000968 | F | F | F | F |
| Crb2 | 0.92709337 | 5.05892156 | 22.0070303 | 5.20E-05 | 0.000968 | F | F | T | F |
| Kdm2b | 0.6197651 | 6.71587817 | 21.9941693 | 5.22E-05 | 0.000971 | T | T | F | T |
| Irs1 | 0.9881974 | 4.65945893 | 21.9702443 | 5.26E-05 | 0.000977 | T | F | T | T |
| Mbd2 | -0.9284971 | 5.23714317 | 21.953829 | 5.28E-05 | 0.000981 | T | F | T | T |
| Ostf1 | -1.0060693 | 4.80375794 | 21.9200476 | 5.34E-05 | 0.00099 | F | F | F | F |
| Rims3 | 2.06979411 | 2.38448091 | 21.9000268 | 5.37E-05 | 0.000995 | T | F | T | T |
| Ephb1 | 1.19641762 | 3.91796675 | 21.8792016 | 5.41E-05 | 0.001001 | F | F | F | F |
| Itgb3 | -0.5370035 | 7.68570322 | 21.8612434 | 5.44E-05 | 0.001006 | F | F | T | T |
| Prl2c2 | 1.86230457 | 5.14459167 | 21.8467821 | 5.46E-05 | 0.001008 | F | F | F | F |

| | | | | | | | | | |
|---|---|---|---|---|---|---|---|---|---|
| Baz1a | -0.5800124 | 7.14634646 | 21.8462415 | 5.46E-05 | 0.001008 | T | F | T | F |
| Armcx1 | 1.61994637 | 3.07473165 | 21.833629 | 5.48E-05 | 0.001011 | T | F | F | F |
| Slc12a7 | -0.5279598 | 7.72587744 | 21.8331089 | 5.48E-05 | 0.001011 | T | F | F | F |
| Wiz | 0.77114543 | 5.79639615 | 21.815232 | 5.51E-05 | 0.001015 | F | F | F | F |
| Clstn1 | 0.55218675 | 7.13166903 | 21.7981792 | 5.54E-05 | 0.00102 | F | F | T | F |
| Pgghg | 1.63312549 | 2.91112005 | 21.7873065 | 5.56E-05 | 0.001022 | T | F | F | F |
| Tmem267 | 2.2478428 | 2.48579299 | 21.7728591 | 5.58E-05 | 0.001026 | F | F | F | F |
| Chchd4 | -0.7229416 | 6.19493823 | 21.768163 | 5.59E-05 | 0.001027 | F | F | T | F |
| Zfp799 | 0.97102388 | 4.59723111 | 21.7626446 | 5.60E-05 | 0.001027 | T | F | T | T |
| Ano8 | 1.6008357 | 3.1027253 | 21.7490109 | 5.62E-05 | 0.001031 | F | F | F | F |
| Frmd5 | 3.67794743 | 0.89969586 | 21.7394867 | 5.64E-05 | 0.001033 | T | F | F | F |
| Osbpl8 | -0.9260798 | 6.11368776 | 21.7240764 | 5.67E-05 | 0.001037 | T | T | F | T |
| Ttc21b | 1.24796007 | 4.08327491 | 21.7165486 | 5.68E-05 | 0.001039 | T | F | F | F |
| Ctnnb1 | 0.41311957 | 9.43132544 | 21.7012307 | 5.71E-05 | 0.001043 | F | F | F | F |
| S1pr5 | 5.56652437 | 0.31339129 | 21.6973559 | 5.71E-05 | 0.001043 | T | F | T | T |
| Nabp1 | -0.812318 | 5.70883252 | 21.6609637 | 5.78E-05 | 0.001054 | T | F | T | T |
| Hmgb3 | 0.75000085 | 5.81013225 | 21.6349133 | 5.82E-05 | 0.001061 | F | F | T | F |
| Cdh11 | 0.9657207 | 4.60619627 | 21.5732773 | 5.94E-05 | 0.001081 | F | F | F | F |
| Sms | 0.91654785 | 4.81278295 | 21.5399178 | 6.00E-05 | 0.001091 | T | F | T | T |
| Frmd4b | -1.0867039 | 4.530462 | 21.5313697 | 6.01E-05 | 0.001093 | F | F | T | F |
| Zeb2 | 1.02553128 | 4.78855211 | 21.5103202 | 6.05E-05 | 0.001099 | T | F | T | F |
| Rftn1 | 2.18661236 | 2.33084011 | 21.4954128 | 6.08E-05 | 0.001103 | F | F | F | T |
| Elavl2 | 2.26415467 | 2.13761339 | 21.4836088 | 6.10E-05 | 0.001106 | F | F | T | T |
| Fbxo16 | 5.64944348 | 0.23121877 | 21.4755273 | 6.12E-05 | 0.001108 | F | F | T | F |
| Sox2 | 0.98354772 | 4.7120724 | 21.4734298 | 6.12E-05 | 0.001108 | T | F | F | F |
| Cpxm1 | 1.45354033 | 3.38710237 | 21.4490305 | 6.17E-05 | 0.001115 | T | F | F | T |
| Csf2ra | -1.7238324 | 3.14913195 | 21.4335011 | 6.20E-05 | 0.00112 | F | F | F | F |
| Fmnl2 | 0.89900946 | 5.00657819 | 21.4286483 | 6.21E-05 | 0.00112 | T | F | T | T |
| Rcbtb1 | 0.7392354 | 5.86842106 | 21.3977385 | 6.26E-05 | 0.00113 | T | F | T | T |
| Helz2 | 0.94537813 | 4.73476793 | 21.3165396 | 6.42E-05 | 0.001158 | F | F | T | T |
| Anxa5 | -0.7283733 | 6.29665828 | 21.3066114 | 6.44E-05 | 0.00116 | F | F | F | F |
| Shb | 1.03404225 | 4.43161666 | 21.2571553 | 6.54E-05 | 0.001177 | T | F | T | F |
| Eif1a | -0.5457472 | 7.47111154 | 21.2536878 | 6.55E-05 | 0.001177 | T | T | T | T |
| Alcam | 1.88136542 | 2.45515755 | 21.2497177 | 6.56E-05 | 0.001177 | F | F | F | F |
| Cpeb2 | -0.924217 | 5.30108569 | 21.249651 | 6.56E-05 | 0.001177 | T | T | F | T |
| Rapgef2 | -0.6350419 | 6.94543529 | 21.2278803 | 6.60E-05 | 0.001184 | T | F | T | T |
| Dab1 | 2.18850786 | 2.48373302 | 21.1996662 | 6.66E-05 | 0.001193 | T | F | T | F |
| Dnmt3l | -0.8067394 | 6.00212858 | 21.1897851 | 6.68E-05 | 0.001196 | T | F | F | F |
| Rabggtb | -0.8558244 | 5.48913637 | 21.1705147 | 6.72E-05 | 0.001202 | F | F | F | F |
| Reep2 | 2.91078877 | 1.46835991 | 21.1598903 | 6.74E-05 | 0.001205 | F | F | T | F |
| Lrtm2 | 8.3212469 | -0.1542829 | 21.3635525 | 6.76E-05 | 0.001208 | F | F | T | F |
| Mfap2 | 2.75322124 | 1.61789791 | 21.1447915 | 6.77E-05 | 0.001209 | F | F | F | F |

| | | | | | | | | | |
|---|---|---|---|---|---|---|---|---|---|
| Tsku | 1.27274585 | 3.67124068 | 21.1336172 | 6.80E-05 | 0.001212 | F | F | F | T |
| Eea1 | -0.7245789 | 6.22497832 | 21.1259739 | 6.81E-05 | 0.001214 | T | F | F | T |
| Agap2 | 3.22416499 | 1.15369286 | 21.1127213 | 6.84E-05 | 0.001218 | F | F | T | F |
| Qser1 | 0.62433203 | 6.70961198 | 21.0865906 | 6.90E-05 | 0.001226 | F | F | T | T |
| Elavl3 | 1.7285107 | 2.81057395 | 21.085788 | 6.90E-05 | 0.001226 | F | F | T | F |
| Ptprz1 | 4.54948581 | 0.55910448 | 21.0799667 | 6.91E-05 | 0.001227 | T | F | T | T |
| Tnk2 | -0.6238163 | 6.8651991 | 21.078704 | 6.91E-05 | 0.001227 | F | F | T | F |
| Pear1 | 1.45694077 | 3.62614541 | 21.0546772 | 6.96E-05 | 0.001235 | F | F | F | F |
| Cemip | 1.16117057 | 4.39112788 | 21.0457128 | 6.98E-05 | 0.001237 | F | F | T | F |
| Ppp1r1a | 2.52761863 | 1.86958665 | 21.0329124 | 7.01E-05 | 0.001241 | F | F | F | F |
| Prom1 | 0.79195183 | 5.43293771 | 20.9736938 | 7.14E-05 | 0.001263 | T | F | T | T |
| Neb | 2.87406254 | 1.48128519 | 20.9700896 | 7.15E-05 | 0.001264 | F | F | F | F |
| Trp53bp1 | 0.64144786 | 6.3975747 | 20.9357535 | 7.23E-05 | 0.001276 | T | T | F | T |
| Cux2 | 3.3500576 | 1.05880591 | 20.9171512 | 7.27E-05 | 0.001282 | F | F | F | T |
| Bcat2 | 0.96807843 | 4.5911433 | 20.8886123 | 7.33E-05 | 0.001292 | T | F | F | T |
| Prpf40b | 1.34472548 | 3.56876278 | 20.887757 | 7.34E-05 | 0.001292 | T | F | F | F |
| Ssbp4 | -0.8340826 | 5.49472773 | 20.8862303 | 7.34E-05 | 0.001292 | T | F | F | F |
| Erlin1 | -0.7780628 | 5.78974397 | 20.8649837 | 7.39E-05 | 0.001299 | T | T | T | T |
| Myl7 | 2.34445135 | 1.9216589 | 20.8286991 | 7.47E-05 | 0.001313 | F | F | F | F |
| Prps1 | 0.84293774 | 5.20418388 | 20.7472393 | 7.66E-05 | 0.001346 | T | F | T | T |
| Ccdc151 | 3.69510839 | 0.82704217 | 20.722705 | 7.72E-05 | 0.001355 | F | F | F | F |
| Tpbgl | 2.18313573 | 2.09104162 | 20.7092759 | 7.76E-05 | 0.001359 | F | F | F | T |
| Epb41l1 | -0.7629436 | 5.8680386 | 20.6972433 | 7.78E-05 | 0.001363 | T | F | T | F |
| Rnf217 | 1.46265256 | 3.19675601 | 20.6716162 | 7.85E-05 | 0.001373 | F | F | T | T |
| Pold1 | 0.55579619 | 6.96846944 | 20.6578248 | 7.88E-05 | 0.001378 | F | F | F | F |
| Vim | 0.64366927 | 6.38789589 | 20.6425709 | 7.92E-05 | 0.001384 | F | F | F | F |
| Arhgef16 | -0.8833396 | 5.43380408 | 20.6207354 | 7.97E-05 | 0.001392 | T | T | F | T |
| Gm9112 | -8.219756 | 0.62189479 | 20.5813146 | 8.07E-05 | 0.001407 | F | F | F | F |
| Nfib | 2.89770514 | 1.42206478 | 20.5779465 | 8.08E-05 | 0.001407 | T | F | T | T |
| Mesp2 | 6.19831325 | 0.04403758 | 20.5754578 | 8.09E-05 | 0.001407 | F | F | T | F |
| Prkd3 | 0.79219673 | 5.39191902 | 20.5742144 | 8.09E-05 | 0.001407 | T | F | T | F |
| Cacna1d | 2.16779165 | 2.24504967 | 20.5723803 | 8.09E-05 | 0.001407 | F | F | F | T |
| Afp | 0.56649194 | 6.88544479 | 20.553168 | 8.14E-05 | 0.001414 | F | F | F | F |
| B3gnt2 | -0.8506956 | 5.38792111 | 20.5507279 | 8.15E-05 | 0.001414 | F | F | T | F |
| Add3 | 0.80308911 | 5.32469528 | 20.5506247 | 8.15E-05 | 0.001414 | T | T | T | T |
| Zmiz1 | -0.5287454 | 7.73132805 | 20.5483532 | 8.16E-05 | 0.001414 | F | T | T | T |
| Il6st | -0.5223644 | 7.65354432 | 20.5401489 | 8.18E-05 | 0.001415 | T | F | T | T |
| Alas1 | -1.2888274 | 3.90934724 | 20.5400615 | 8.18E-05 | 0.001415 | F | F | F | F |
| Sez6 | 8.28570155 | -0.1851711 | 20.7183326 | 8.24E-05 | 0.001424 | F | F | T | F |
| Cfap52 | 8.2856656 | -0.1852023 | 20.7176889 | 8.24E-05 | 0.001424 | F | F | F | F |
| Sec14l2 | -0.6059046 | 6.83264055 | 20.493276 | 8.30E-05 | 0.001433 | T | F | T | F |
| Slc25a4 | -0.4524044 | 8.65555305 | 20.489283 | 8.31E-05 | 0.001433 | T | F | F | T |

| | | | | | | | | | |
|---|---|---|---|---|---|---|---|---|---|
| Tmem170b | -1.3090141 | 4.01306506 | 20.4848116 | 8.32E-05 | 0.001434 | F | F | T | T |
| Iglon5 | 2.92650149 | 1.31209934 | 20.4726446 | 8.35E-05 | 0.001438 | F | F | T | F |
| Ret | 1.58974282 | 2.90284215 | 20.4703514 | 8.36E-05 | 0.001438 | T | F | T | T |
| Cdk6 | 0.8757452 | 5.13223863 | 20.4168357 | 8.50E-05 | 0.001461 | T | F | T | T |
| Cbx6 | 0.81569842 | 5.24750365 | 20.4096087 | 8.52E-05 | 0.001464 | T | F | T | T |
| Brsk1 | 1.28155824 | 3.60020923 | 20.3879913 | 8.58E-05 | 0.001472 | F | F | F | F |
| Cdh10 | 8.27780674 | -0.1920187 | 20.5774 | 8.61E-05 | 0.001476 | T | F | T | F |
| Sema6b | 1.10844867 | 4.09891639 | 20.3751252 | 8.61E-05 | 0.001476 | F | F | T | F |
| Adam22 | 2.04794045 | 2.16268659 | 20.3587283 | 8.66E-05 | 0.001483 | T | F | T | F |
| Slc35a3 | -0.7484393 | 6.03128707 | 20.3352863 | 8.72E-05 | 0.001492 | T | F | T | T |
| Pdhb | -0.6132983 | 6.80199808 | 20.2892293 | 8.85E-05 | 0.001513 | T | F | F | F |
| Rnf219 | 1.01264378 | 4.35040099 | 20.2469715 | 8.97E-05 | 0.001532 | T | F | F | F |
| Repin1 | 0.95274172 | 4.54727962 | 20.2403165 | 8.99E-05 | 0.001534 | F | F | F | F |
| Wnt2 | 3.12846762 | 1.15475533 | 20.2280045 | 9.02E-05 | 0.001539 | T | F | F | F |
| Tcf7 | 0.66021283 | 6.28347056 | 20.2055697 | 9.08E-05 | 0.001549 | F | F | F | F |
| Arl8a | -0.8228115 | 5.50522428 | 20.1972944 | 9.11E-05 | 0.001551 | F | F | T | F |
| Adar | 0.84835942 | 5.3636235 | 20.190773 | 9.13E-05 | 0.001553 | T | F | F | F |
| Neurl4 | 0.60232138 | 6.62830275 | 20.1889104 | 9.13E-05 | 0.001553 | F | F | T | F |
| Foxp1 | 1.1796399 | 3.84725345 | 20.1842415 | 9.15E-05 | 0.001554 | T | F | T | T |
| Atp2b2 | 8.26641133 | -0.2018942 | 20.3754881 | 9.16E-05 | 0.001555 | T | F | T | F |
| Frk | -1.1174536 | 4.37670916 | 20.1778909 | 9.16E-05 | 0.001555 | T | F | T | T |
| Ptprt | 3.99994467 | 0.65032306 | 20.1551188 | 9.23E-05 | 0.001565 | F | F | T | T |
| Chst15 | 0.84926629 | 5.03826783 | 20.1036011 | 9.38E-05 | 0.001589 | F | F | T | F |
| P3h1 | 0.75114323 | 5.5815647 | 20.0809156 | 9.45E-05 | 0.001599 | F | F | T | F |
| Bcl7a | 0.8846135 | 4.84910573 | 20.0529076 | 9.53E-05 | 0.001612 | T | F | T | T |
| Apbb2 | -0.7572754 | 6.26020892 | 20.0433956 | 9.56E-05 | 0.001616 | T | F | T | T |
| Gprin1 | 3.68147196 | 0.88197817 | 20.0390278 | 9.58E-05 | 0.001617 | F | F | T | F |
| Cep131 | 0.92032098 | 4.63527153 | 20.0143874 | 9.65E-05 | 0.001628 | F | F | F | F |
| Plod3 | 0.57252179 | 6.94996004 | 20.0013771 | 9.69E-05 | 0.001634 | T | F | F | T |
| Capn6 | 1.12263455 | 4.09785759 | 19.9775884 | 9.76E-05 | 0.001645 | T | F | F | T |
| Klf10 | -0.8061755 | 5.56014885 | 19.9512824 | 9.85E-05 | 0.001658 | F | F | T | F |
| Rpl35a | -0.6161208 | 6.73249394 | 19.9306148 | 9.91E-05 | 0.001667 | F | F | F | F |
| Mbnl2 | -0.6933879 | 6.2112403 | 19.8086806 | 0.0001 | 0.001732 | T | F | F | T |
| Bmp1 | 0.90506451 | 4.7666201 | 19.8041622 | 0.0001 | 0.001733 | F | F | T | F |
| Jak2 | -0.9111666 | 5.22787252 | 19.7978544 | 0.0001 | 0.001735 | F | F | T | F |
| Arhgap45 | 0.94414173 | 4.57457009 | 19.7771133 | 0.0001 | 0.001745 | F | F | F | F |
| 1110008P1[illegible] | -1.039411 | 4.50787053 | 19.7698722 | 0.0001 | 0.001747 | F | F | F | F |
| Tchh | 3.68373593 | 0.86241562 | 19.7697345 | 0.0001 | 0.001747 | F | F | F | F |
| Alx1 | 5.0716996 | 0.37936605 | 19.7639476 | 0.0001 | 0.001748 | T | F | T | T |
| Rassf8 | -0.7073162 | 6.11502434 | 19.7474167 | 0.00011 | 0.001755 | T | F | T | T |
| Phf13 | -0.5845951 | 7.05124839 | 19.7469296 | 0.00011 | 0.001755 | F | F | T | F |
| Fosl2 | -0.9260926 | 4.93253071 | 19.7234836 | 0.00011 | 0.001767 | T | F | T | F |

| | | | | | | | | | |
|---|---|---|---|---|---|---|---|---|---|
| Trps1 | 1.86195106 | 2.38917822 | 19.7175422 | 0.00011 | 0.001769 | T | F | T | T |
| mt-Nd4 | -0.4003378 | 9.66584489 | 19.7108519 | 0.00011 | 0.001771 | F | F | F | F |
| Fasl | -1.5174819 | 3.39903526 | 19.7044761 | 0.00011 | 0.001774 | F | F | T | F |
| Iffo2 | -0.9041461 | 5.62490452 | 19.692679 | 0.00011 | 0.001779 | F | F | T | F |
| Gucy1a2 | 3.15009455 | 1.15179 | 19.6570446 | 0.00011 | 0.001798 | T | T | T | T |
| Chst11 | 1.82170148 | 2.44888586 | 19.5984764 | 0.00011 | 0.001831 | T | F | T | T |
| Nr2f6 | -0.6083435 | 6.69054359 | 19.5915214 | 0.00011 | 0.001833 | T | F | T | F |
| Stc1 | 4.43618805 | 0.44010874 | 19.5886565 | 0.00011 | 0.001834 | T | F | F | F |
| Spock1 | 2.49340124 | 1.65681555 | 19.5592069 | 0.00011 | 0.001849 | T | F | T | F |
| Klhl42 | -0.8336487 | 5.47744299 | 19.5355329 | 0.00011 | 0.001862 | T | F | T | T |
| Gm50241 | 3.23747332 | 2.13469505 | 19.5311716 | 0.00011 | 0.001863 | F | F | F | F |
| Cast | -0.7319861 | 5.8942303 | 19.5255004 | 0.00011 | 0.001865 | T | F | T | T |
| Map3k5 | 0.96228003 | 4.50220519 | 19.5117392 | 0.00011 | 0.001872 | F | F | F | F |
| Nhsl1 | -0.6491394 | 6.59320793 | 19.5082216 | 0.00011 | 0.001873 | T | F | T | F |
| Cdc14b | 0.68907645 | 5.92096595 | 19.4849767 | 0.00011 | 0.001885 | T | F | T | T |
| Tmem151b | 2.8637506 | 1.47327812 | 19.4571562 | 0.00012 | 0.001901 | F | F | F | F |
| St6gal2 | 4.64679251 | 0.3351266 | 19.4521424 | 0.00012 | 0.001902 | T | F | T | T |
| Aldh3b2 | -1.6328359 | 3.30879535 | 19.4497217 | 0.00012 | 0.001902 | F | F | T | F |
| Arfgef3 | 2.00437191 | 2.16201835 | 19.4464013 | 0.00012 | 0.001903 | T | F | F | T |
| Bmp2 | 0.89510455 | 4.68128584 | 19.4359996 | 0.00012 | 0.001908 | T | F | F | T |
| Yars | -0.5185286 | 7.45244499 | 19.4181388 | 0.00012 | 0.001916 | F | F | T | F |
| Plcd1 | -0.9018466 | 4.98630883 | 19.4179654 | 0.00012 | 0.001916 | F | F | F | F |
| Wnt5b | 2.31685726 | 2.22568716 | 19.4145576 | 0.00012 | 0.001917 | T | T | T | T |
| Alad | -0.8048508 | 5.48970764 | 19.3777193 | 0.00012 | 0.001938 | F | F | T | T |
| Sash1 | 0.72379792 | 6.03962468 | 19.3597554 | 0.00012 | 0.001948 | T | F | T | T |
| Galnt9 | 5.04677923 | 0.35815166 | 19.3334851 | 0.00012 | 0.001962 | F | F | F | F |
| Sik1 | -0.5637076 | 7.00982941 | 19.3321395 | 0.00012 | 0.001962 | T | F | F | F |
| Atp1a3 | 3.55731191 | 0.98239864 | 19.291341 | 0.00012 | 0.001987 | F | F | F | F |
| Bcorl1 | 0.81977638 | 5.40375102 | 19.2729732 | 0.00012 | 0.001997 | F | F | F | F |
| Cystm1 | -1.0886613 | 4.42017751 | 19.2528002 | 0.00012 | 0.002009 | F | F | F | T |
| Ogdhl | 0.75133576 | 5.48390049 | 19.2343244 | 0.00012 | 0.002019 | F | F | F | F |
| Nmt2 | -0.5341142 | 7.22481952 | 19.2198901 | 0.00012 | 0.002027 | F | F | T | T |
| Cask | 0.94545616 | 4.48826842 | 19.1999499 | 0.00013 | 0.002038 | T | F | T | F |
| Tcf7l1 | 0.54551883 | 6.90519355 | 19.1847347 | 0.00013 | 0.002047 | F | F | T | F |
| Trip12 | -0.40338 | 9.64418916 | 19.1800478 | 0.00013 | 0.002049 | F | F | F | T |
| Zfp664 | 0.54795342 | 6.87874961 | 19.1558905 | 0.00013 | 0.002062 | T | F | T | T |
| Hbegf | -1.1313927 | 4.18712041 | 19.1556057 | 0.00013 | 0.002062 | F | F | T | F |
| Eif3j2 | -0.5852575 | 6.86539428 | 19.148601 | 0.00013 | 0.002065 | F | F | T | T |
| Nefl | 1.05242023 | 4.12904417 | 19.1454374 | 0.00013 | 0.002065 | F | F | T | T |
| Crispld1 | 2.06122741 | 2.05801293 | 19.1030742 | 0.00013 | 0.002092 | T | F | T | T |
| Irf2bpl | 0.87987523 | 4.9814952 | 19.0860524 | 0.00013 | 0.002102 | F | F | F | F |
| Spock2 | 1.12504791 | 3.89528676 | 19.063775 | 0.00013 | 0.002116 | T | T | T | T |

| | | | | | | | | | |
|---|---|---|---|---|---|---|---|---|---|
| Ephb6 | 1.87385947 | 2.29386138 | 19.0586345 | 0.00013 | 0.002118 | F | F | F | F |
| Prl5a1 | 1.06910045 | 4.0918783 | 19.0394133 | 0.00013 | 0.002129 | F | F | F | F |
| Fhod3 | 1.17318043 | 3.89692295 | 19.0161155 | 0.00013 | 0.002143 | T | T | F | T |
| Wnt5a | 1.18935384 | 3.97393241 | 19.0151873 | 0.00013 | 0.002143 | T | F | T | T |
| Bicd1 | 2.38648424 | 1.95569044 | 18.9608087 | 0.00014 | 0.00218 | T | F | T | F |
| Dnajc2 | -0.5407092 | 7.22609449 | 18.9571397 | 0.00014 | 0.002181 | F | F | F | F |
| Pkdcc | 0.69064688 | 6.035732 | 18.9458832 | 0.00014 | 0.002187 | F | F | F | F |
| Tulp3 | 0.71393601 | 5.67426266 | 18.9240845 | 0.00014 | 0.002201 | T | F | T | T |
| Ak3 | -0.702207 | 6.06386107 | 18.915683 | 0.00014 | 0.002205 | T | F | T | T |
| Dock5 | -0.5105147 | 7.59448055 | 18.8986151 | 0.00014 | 0.002216 | F | F | T | F |
| Arhgef12 | -0.5049718 | 7.52711119 | 18.8891645 | 0.00014 | 0.002221 | T | F | T | T |
| Dhcr24 | 0.43743333 | 8.391512 | 18.8832102 | 0.00014 | 0.002224 | F | F | T | F |
| Myof | -0.6705897 | 6.25779739 | 18.8733651 | 0.00014 | 0.002229 | T | F | T | F |
| Rcbtb2 | 0.70851365 | 5.71433592 | 18.8649974 | 0.00014 | 0.002234 | T | F | F | F |
| Trim24 | 0.85024198 | 5.44343772 | 18.8500139 | 0.00014 | 0.002243 | T | F | F | T |
| Pgm2 | -0.7477832 | 5.76103912 | 18.8143438 | 0.00014 | 0.002267 | T | F | F | F |
| Prl2c3 | 1.56626405 | 5.11810984 | 18.798975 | 0.00014 | 0.002277 | F | F | F | F |
| Cacna1g | 2.51767911 | 1.56523808 | 18.7686422 | 0.00014 | 0.002298 | F | F | F | F |
| Atf7ip | 0.52079453 | 7.20350164 | 18.7204108 | 0.00015 | 0.002333 | T | T | F | T |
| Tnrc6b | 0.63508779 | 6.17557707 | 18.7142623 | 0.00015 | 0.002336 | T | F | T | T |
| H4c8 | 0.40442239 | 8.929802 | 18.7091667 | 0.00015 | 0.002338 | F | F | F | F |
| Apoa1 | -0.4569031 | 8.92476147 | 18.7051512 | 0.00015 | 0.002339 | F | F | F | F |
| Gsto1 | -0.7965354 | 7.29609935 | 18.697452 | 0.00015 | 0.002343 | F | F | F | F |
| Ryr2 | 1.36708075 | 3.2404982 | 18.6860401 | 0.00015 | 0.00235 | F | F | F | F |
| Zfhx4 | 3.59950998 | 0.7649847 | 18.6531145 | 0.00015 | 0.002374 | F | F | F | F |
| Kmt5c | 0.79763247 | 5.14781017 | 18.64741 | 0.00015 | 0.002377 | F | F | F | F |
| Mcf2l | 1.54980753 | 2.94465208 | 18.628467 | 0.00015 | 0.00239 | T | F | T | F |
| Actb | -0.2856048 | 12.384293 | 18.6126477 | 0.00015 | 0.0024 | F | F | T | F |
| Mapk8ip1 | 0.85931405 | 4.77197669 | 18.6046953 | 0.00015 | 0.002405 | F | F | T | F |
| Syndig1l | 2.95211247 | 1.28970152 | 18.5984662 | 0.00015 | 0.002408 | F | F | T | F |
| Fkbp15 | -0.6630878 | 6.23531685 | 18.5897057 | 0.00015 | 0.002413 | T | F | F | T |
| Zfp644 | 0.66393983 | 5.96221451 | 18.5857671 | 0.00015 | 0.002414 | T | F | T | T |
| Rasgef1b | -1.132789 | 4.21589423 | 18.5790612 | 0.00015 | 0.002418 | T | F | F | T |
| Fyttd1 | -0.6969803 | 6.14748867 | 18.5623397 | 0.00015 | 0.002429 | T | F | T | T |
| Foxo1 | 0.80282011 | 5.22622271 | 18.5456753 | 0.00015 | 0.002441 | T | F | T | T |
| Pdia4 | 0.44501049 | 8.19831131 | 18.5103192 | 0.00016 | 0.002468 | F | F | F | F |
| Adamts5 | 3.11062918 | 1.06534905 | 18.496945 | 0.00016 | 0.002477 | T | F | T | T |
| Homez | 1.22083129 | 3.66755972 | 18.4863631 | 0.00016 | 0.002483 | T | F | F | T |
| Col6a2 | 2.13386264 | 1.95963241 | 18.4740404 | 0.00016 | 0.002491 | T | F | T | T |
| Fgd5 | 1.15881428 | 3.79133614 | 18.4732574 | 0.00016 | 0.002491 | F | F | F | F |
| Slc41a1 | 0.94832617 | 4.41024477 | 18.4447062 | 0.00016 | 0.002512 | T | F | T | F |
| Pgr | 2.83893481 | 1.31658341 | 18.4416497 | 0.00016 | 0.002513 | T | F | T | T |

| | | | | | | | | | |
|---|---|---|---|---|---|---|---|---|---|
| Slc1a3 | 1.21625371 | 3.59993039 | 18.4091874 | 0.00016 | 0.002538 | T | F | T | T |
| Esx1 | -1.2918376 | 3.96734641 | 18.3733845 | 0.00016 | 0.002563 | F | F | F | F |
| Tecr | 0.55927263 | 6.79296309 | 18.372753 | 0.00016 | 0.002563 | T | F | F | F |
| Tbc1d10b | -0.5317693 | 7.10679994 | 18.3718759 | 0.00016 | 0.002563 | F | F | F | F |
| Als2cl | -0.8812716 | 5.13092616 | 18.3710233 | 0.00016 | 0.002563 | T | F | F | F |
| Bbs1 | 1.51709745 | 2.87035682 | 18.3591374 | 0.00016 | 0.00257 | F | F | T | T |
| Rims2 | 1.13059887 | 3.84297433 | 18.3567023 | 0.00016 | 0.00257 | F | F | T | F |
| Inava | 0.82905883 | 4.97915503 | 18.3556926 | 0.00016 | 0.00257 | F | F | F | F |
| Stxbp5 | -0.6888596 | 6.0510686 | 18.3266322 | 0.00017 | 0.002593 | T | F | T | T |
| Prl2a1 | -1.6184441 | 3.71274683 | 18.3053912 | 0.00017 | 0.002609 | F | F | F | F |
| Adam8 | -0.9820889 | 4.53478913 | 18.302565 | 0.00017 | 0.00261 | F | F | F | F |
| Aacs | -0.8915704 | 5.05492624 | 18.29592 | 0.00017 | 0.002613 | F | F | T | T |
| Fam199x | 0.7696722 | 5.26916397 | 18.2895519 | 0.00017 | 0.002617 | T | F | T | T |
| Ahcy | 0.45052343 | 8.71610742 | 18.2867032 | 0.00017 | 0.002618 | F | F | F | F |
| Clstn3 | 1.048058 | 4.07505381 | 18.279461 | 0.00017 | 0.002622 | F | F | F | F |
| Batf3 | -1.7327153 | 2.94397384 | 18.2705198 | 0.00017 | 0.002628 | F | F | F | F |
| H2az2 | 0.86140652 | 4.72448918 | 18.2687798 | 0.00017 | 0.002628 | F | F | F | F |
| Lgi2 | 1.95518544 | 2.17145043 | 18.2600976 | 0.00017 | 0.002632 | F | F | T | F |
| Lrch2 | 2.10747764 | 2.30428984 | 18.2587804 | 0.00017 | 0.002632 | F | F | F | T |
| Sirt5 | -0.9055458 | 6.30315909 | 18.2554761 | 0.00017 | 0.002633 | T | F | T | F |
| Tmcc3 | -0.838726 | 5.25848027 | 18.2202475 | 0.00017 | 0.002662 | T | F | T | T |
| Prmt8 | 5.36945281 | 0.14592797 | 18.209228 | 0.00017 | 0.00267 | T | F | T | T |
| Slco2a1 | 1.04619293 | 4.0533719 | 18.1981179 | 0.00017 | 0.002678 | T | F | T | T |
| Pak2 | -0.48684 | 7.86447113 | 18.1366944 | 0.00018 | 0.002731 | T | F | T | F |
| Ccn1 | 0.81562192 | 5.28250693 | 18.1330856 | 0.00018 | 0.002732 | F | F | F | F |
| Elobl | 3.6499513 | 0.69419175 | 18.116705 | 0.00018 | 0.002745 | F | F | F | F |
| Nsd3 | -0.5816781 | 6.71268845 | 18.098739 | 0.00018 | 0.00276 | T | F | T | F |
| Sorbs2 | 3.13117279 | 1.00845536 | 18.0800278 | 0.00018 | 0.002775 | T | F | T | T |
| Sqstm1 | -0.4607903 | 7.98647119 | 18.070362 | 0.00018 | 0.002782 | F | F | F | T |
| Actr3 | -0.4837594 | 7.9994807 | 18.0671509 | 0.00018 | 0.002783 | T | T | T | T |
| Tsr1 | -0.4920632 | 7.78535463 | 18.0578831 | 0.00018 | 0.002789 | T | F | F | F |
| Dmrta1 | 4.3791441 | 0.40953405 | 18.0545353 | 0.00018 | 0.00279 | T | T | F | T |
| Mixl1 | 2.27729802 | 2.53500027 | 18.0448887 | 0.00018 | 0.002796 | F | F | F | F |
| H4c6 | 0.43923677 | 8.31532518 | 18.0444718 | 0.00018 | 0.002796 | F | F | F | F |
| Mob3c | -1.1349469 | 4.07799792 | 18.0369397 | 0.00018 | 0.002801 | T | F | T | T |
| Sec14l4 | 1.77259537 | 2.36248172 | 18.0084687 | 0.00018 | 0.002824 | F | F | F | F |
| Magi3 | -0.6358452 | 6.32277003 | 18.0079803 | 0.00018 | 0.002824 | F | F | F | F |
| Itga7 | -0.6067281 | 6.76781255 | 18.0008462 | 0.00019 | 0.002828 | T | F | F | T |
| Arhgef3 | -0.861369 | 5.01540512 | 17.9725634 | 0.00019 | 0.002853 | T | F | T | T |
| Mfsd9 | 8.13009726 | -0.3192452 | 18.0955935 | 0.00019 | 0.002885 | F | F | T | T |
| Myo5c | 2.08321751 | 1.94782119 | 17.9347094 | 0.00019 | 0.002885 | T | F | F | F |
| Wls | 0.74743136 | 5.544149 | 17.905575 | 0.00019 | 0.002909 | T | F | F | F |

| | | | | | | | | | |
|---|---|---|---|---|---|---|---|---|---|
| Armcx3 | 1.75018911 | 2.44462539 | 17.9032941 | 0.00019 | 0.002909 | T | F | T | T |
| Apela | 2.04619479 | 2.09599975 | 17.9005464 | 0.00019 | 0.002909 | F | F | F | F |
| Rtn4rl2 | 2.88279588 | 1.17299258 | 17.8998764 | 0.00019 | 0.002909 | F | F | F | T |
| Phf19 | -1.1233492 | 4.97822076 | 17.8992626 | 0.00019 | 0.002909 | T | F | T | F |
| Flrt3 | 0.67940773 | 5.81439276 | 17.8849893 | 0.00019 | 0.002921 | T | F | F | F |
| Syndig1 | 4.98733755 | 0.2179006 | 17.8785424 | 0.00019 | 0.002925 | T | F | F | F |
| S100a16 | -1.4262068 | 3.48288087 | 17.8738093 | 0.00019 | 0.002928 | F | F | T | F |
| Ssc5d | 3.62963771 | 0.83366669 | 17.8554981 | 0.00019 | 0.002943 | F | F | F | F |
| Insm1 | 2.64598026 | 1.37473837 | 17.835627 | 0.0002 | 0.002961 | T | F | F | T |
| Sptb | -1.6199538 | 4.87131121 | 17.8291227 | 0.0002 | 0.002963 | T | F | T | F |
| Arhgap27 | -0.9704336 | 5.20594048 | 17.8287605 | 0.0002 | 0.002963 | T | F | T | T |
| Crlf3 | -0.7202916 | 5.80837546 | 17.8268082 | 0.0002 | 0.002963 | F | F | F | F |
| Uchl5 | -0.783726 | 5.39994179 | 17.7794081 | 0.0002 | 0.003008 | F | F | F | T |
| Usp49 | -0.8525797 | 5.03011004 | 17.7551474 | 0.0002 | 0.003031 | T | F | T | T |
| Shc2 | 1.3277394 | 3.25815517 | 17.725151 | 0.0002 | 0.003059 | T | F | T | T |
| Pdia6 | 0.41816506 | 8.70166412 | 17.7185128 | 0.0002 | 0.003064 | F | F | F | F |
| Nefh | 4.57376527 | 0.29198935 | 17.7156662 | 0.0002 | 0.003064 | F | F | F | F |
| Chmp4b | -0.5387174 | 7.1170926 | 17.6853961 | 0.00021 | 0.003093 | F | F | F | T |
| Slc44a4 | 0.84582642 | 4.83600526 | 17.6573158 | 0.00021 | 0.003119 | F | F | F | F |
| Klkb1 | -1.5849286 | 3.10867399 | 17.6545744 | 0.00021 | 0.003119 | F | F | T | F |
| Efhc2 | 2.182207 | 1.88558542 | 17.6541903 | 0.00021 | 0.003119 | F | F | F | F |
| Fign | 1.66044647 | 2.57049558 | 17.6506009 | 0.00021 | 0.003121 | F | F | T | T |
| Pianp | 1.57672803 | 2.69470182 | 17.6462631 | 0.00021 | 0.003123 | F | F | T | F |
| Eno1 | 0.30782497 | 11.2779471 | 17.6404683 | 0.00021 | 0.003127 | T | T | F | T |
| Oas1c | -3.1503457 | 1.54912263 | 17.6211698 | 0.00021 | 0.003145 | F | F | F | F |
| Mknk1 | -1.0750186 | 4.19236203 | 17.5891645 | 0.00021 | 0.003177 | F | F | F | F |
| Fam120a | -0.4272319 | 8.51906359 | 17.561727 | 0.00021 | 0.003203 | F | F | T | F |
| Prss12 | 0.8388818 | 4.7343071 | 17.558691 | 0.00021 | 0.003203 | F | F | F | T |
| Dipk2a | 0.88171174 | 4.56062064 | 17.5582404 | 0.00021 | 0.003203 | T | F | T | T |
| Gpbp1 | -0.6455167 | 6.78349084 | 17.5131154 | 0.00022 | 0.00325 | T | F | F | T |
| Tfdp2 | 0.73729713 | 5.35487141 | 17.5059934 | 0.00022 | 0.003255 | T | F | T | T |
| Tcf7l2 | 0.71660449 | 5.51914963 | 17.4990397 | 0.00022 | 0.003261 | T | F | T | T |
| Ankrd13b | 1.62977365 | 2.73099229 | 17.4758257 | 0.00022 | 0.003282 | T | F | T | T |
| Lars2 | 0.51486195 | 6.98306057 | 17.4757999 | 0.00022 | 0.003282 | F | F | F | F |
| Itga5 | -0.4626492 | 7.81563462 | 17.4504528 | 0.00022 | 0.003307 | T | F | F | F |
| mt-Nd1 | -0.3606343 | 10.1853623 | 17.4486565 | 0.00022 | 0.003307 | F | F | F | F |
| Prkd1 | 1.23908809 | 3.565829 | 17.439405 | 0.00022 | 0.003315 | T | F | F | F |
| Fgf8 | 2.6627795 | 1.90390524 | 17.4126054 | 0.00023 | 0.003343 | F | F | T | F |
| Ldha | 0.31762948 | 10.7952916 | 17.3900514 | 0.00023 | 0.003366 | T | F | F | F |
| Sorbs3 | 1.1177496 | 3.88949535 | 17.3802739 | 0.00023 | 0.003375 | F | F | F | F |
| Xkr7 | 8.09368083 | -0.3503433 | 17.5276603 | 0.00023 | 0.003376 | F | F | F | T |
| Ptprf | 0.40257451 | 9.24755112 | 17.3708274 | 0.00023 | 0.003381 | F | F | T | T |

| | | | | | | | | | |
|---|---|---|---|---|---|---|---|---|---|
| C1galt1 | -0.7231066 | 5.71065199 | 17.3245717 | 0.00023 | 0.003432 | T | F | F | T |
| Hcn4 | 3.07261976 | 0.98861498 | 17.3142011 | 0.00023 | 0.003441 | F | F | T | F |
| Gprasp1 | 0.75631565 | 5.34195211 | 17.2828167 | 0.00024 | 0.003474 | T | F | F | T |
| Calcr | 5.99096961 | -0.0930636 | 17.2818449 | 0.00024 | 0.003474 | T | F | F | F |
| Map4 | -0.4427752 | 8.20524319 | 17.2448381 | 0.00024 | 0.003516 | T | F | T | T |
| Hebp1 | -0.9411124 | 4.61163322 | 17.2340433 | 0.00024 | 0.003526 | F | F | T | F |
| Fxyd5 | -1.1577426 | 3.94105641 | 17.2038485 | 0.00024 | 0.00356 | T | F | F | T |
| Ypel1 | 2.66706176 | 1.31477792 | 17.1963932 | 0.00024 | 0.003566 | T | F | T | F |
| Dlx5 | 4.11985638 | 0.3961268 | 17.1795057 | 0.00024 | 0.003584 | F | F | F | F |
| Plekha7 | -0.5338626 | 6.94574023 | 17.1635899 | 0.00025 | 0.003601 | T | F | T | F |
| Mfsd14b | -0.6041807 | 6.51976827 | 17.1494493 | 0.00025 | 0.003616 | T | F | T | F |
| Hipk3 | -0.5854663 | 6.5703868 | 17.1294186 | 0.00025 | 0.003638 | F | F | T | T |
| Anxa2 | -0.5516203 | 6.87032604 | 17.1108561 | 0.00025 | 0.003659 | F | F | F | F |
| Mrtfb | -0.6175088 | 6.37861391 | 17.0727575 | 0.00025 | 0.003704 | F | F | T | F |
| Tacstd2 | -0.9452357 | 4.59351085 | 17.0575711 | 0.00025 | 0.00372 | F | F | F | F |
| Cacng8 | 2.61575539 | 1.34401106 | 17.0412116 | 0.00026 | 0.003738 | F | F | T | F |
| Crtc3 | -0.8283646 | 5.09725903 | 17.0069109 | 0.00026 | 0.003779 | T | T | T | T |
| Gm4724 | 8.06640115 | -0.3735669 | 17.1132186 | 0.00026 | 0.003825 | T | F | T | T |
| Ybx2 | 2.22464355 | 1.78254172 | 16.9624607 | 0.00026 | 0.003832 | T | F | F | F |
| 4921524J17 | -0.9715426 | 4.64429084 | 16.9497158 | 0.00026 | 0.003846 | T | F | T | F |
| Zfp365 | 4.06906768 | 0.65093696 | 16.9391079 | 0.00026 | 0.003857 | F | F | T | F |
| Slc38a7 | -0.5439198 | 6.85678399 | 16.9208565 | 0.00027 | 0.003878 | F | F | F | F |
| Ralb | -0.5986223 | 6.47386349 | 16.9096885 | 0.00027 | 0.003891 | F | F | T | F |
| Sox13 | 0.86729259 | 4.72560409 | 16.875947 | 0.00027 | 0.003933 | T | F | F | T |
| Eloa | -0.5457963 | 6.8581866 | 16.8444191 | 0.00027 | 0.003973 | T | T | T | T |
| Ptprn | 1.15694083 | 3.62824565 | 16.8405601 | 0.00027 | 0.003973 | F | F | F | F |
| Ecm1 | 0.8914886 | 4.88169618 | 16.8399983 | 0.00027 | 0.003973 | F | F | F | F |
| Ddx39b | 0.42977067 | 8.26153494 | 16.8360774 | 0.00027 | 0.003976 | T | F | F | F |
| Hmox1 | -0.4856019 | 7.43473006 | 16.8200677 | 0.00028 | 0.003995 | F | F | T | F |
| Atp2a1 | 4.51901824 | 0.24651103 | 16.7992251 | 0.00028 | 0.00402 | F | F | F | F |
| Cnnm2 | 1.01386192 | 4.03325672 | 16.7977665 | 0.00028 | 0.00402 | F | F | F | T |
| Pdcd6ip | -0.4562398 | 7.80803109 | 16.7907638 | 0.00028 | 0.004027 | F | F | T | F |
| Otop3 | 8.05201597 | -0.385788 | 16.8984168 | 0.00028 | 0.00407 | F | F | F | F |
| Ocstamp | -1.9659428 | 2.44671566 | 16.7556056 | 0.00028 | 0.00407 | T | T | F | T |
| Parm1 | -0.8731444 | 4.77574737 | 16.7461902 | 0.00028 | 0.004081 | T | F | T | T |
| Gak | -0.4558353 | 7.78912774 | 16.7384689 | 0.00028 | 0.004089 | F | F | T | T |
| Lrguk | 1.94451849 | 2.05055473 | 16.7320883 | 0.00028 | 0.004095 | T | F | T | T |
| Syt14 | 0.97399608 | 4.15401261 | 16.7159627 | 0.00029 | 0.004115 | T | F | T | T |
| Sars | -0.4886528 | 7.56656301 | 16.7033342 | 0.00029 | 0.004129 | T | F | T | F |
| Pkn2 | -0.7460578 | 6.66117694 | 16.701675 | 0.00029 | 0.004129 | T | F | T | F |
| Nrp2 | 0.53341949 | 6.70986644 | 16.6913521 | 0.00029 | 0.004141 | T | F | T | F |
| Szt2 | 0.55861717 | 6.55357009 | 16.6854869 | 0.00029 | 0.004147 | F | F | F | F |

| | | | | | | | | | |
|---|---|---|---|---|---|---|---|---|---|
| Dync2h1 | 0.69233494 | 5.52504787 | 16.6225642 | 0.00029 | 0.004234 | F | F | T | F |
| Gm11627 | 8.04034399 | -0.3956912 | 16.7561618 | 0.0003 | 0.004238 | F | F | F | F |
| 1700017B0: | -0.518398 | 7.22670759 | 16.6159891 | 0.0003 | 0.004238 | T | F | F | F |
| Ttc9 | 1.44084127 | 2.96805975 | 16.6043301 | 0.0003 | 0.004252 | T | F | T | F |
| Mogat2 | 0.55134267 | 6.55430546 | 16.599962 | 0.0003 | 0.004256 | T | F | F | T |
| Fam129b | 0.51734643 | 6.82839284 | 16.5926357 | 0.0003 | 0.004264 | T | F | T | T |
| Ptgfrn | 0.51242206 | 6.90112189 | 16.5898595 | 0.0003 | 0.004265 | F | F | T | F |
| Fhod1 | -0.6002279 | 6.46991776 | 16.5608432 | 0.0003 | 0.004304 | F | F | F | F |
| Pcdh1 | 0.52288944 | 7.73587816 | 16.5596341 | 0.0003 | 0.004304 | F | F | T | T |
| Arhgef19 | 1.70428388 | 2.63482009 | 16.5397036 | 0.0003 | 0.00433 | F | F | T | T |
| Larp4b | -0.4414873 | 7.98456097 | 16.5234299 | 0.00031 | 0.004352 | T | F | T | T |
| Ift122 | 1.01903887 | 3.9954361 | 16.5117372 | 0.00031 | 0.004366 | T | F | T | F |
| Mrfap1 | -0.398172 | 9.24985635 | 16.4985659 | 0.00031 | 0.004383 | T | F | F | T |
| Art4 | -4.0936731 | 1.07934295 | 16.4735144 | 0.00031 | 0.004416 | T | F | T | T |
| Kcp | 3.84631929 | 0.64311972 | 16.47296 | 0.00031 | 0.004416 | F | F | F | F |
| Rps26 | -0.3962652 | 8.92638408 | 16.4669762 | 0.00031 | 0.004422 | F | F | F | F |
| Elovl1 | -0.5919037 | 6.58905738 | 16.4505984 | 0.00031 | 0.004444 | T | F | F | F |
| Arhgef25 | 0.87421588 | 4.50492746 | 16.4484446 | 0.00031 | 0.004445 | F | F | F | F |
| Rgs16 | 1.57785733 | 2.66496415 | 16.4465787 | 0.00031 | 0.004445 | T | F | F | T |
| Tfeb | -0.6565875 | 5.99744265 | 16.4412002 | 0.00031 | 0.00445 | F | F | F | F |
| Bex1 | -0.4228959 | 8.33727647 | 16.4312081 | 0.00031 | 0.004462 | F | F | F | F |
| Ncapd2 | 0.42321353 | 7.95170502 | 16.418217 | 0.00032 | 0.004479 | F | F | F | F |
| Pgbd5 | 2.5806448 | 1.33945117 | 16.3894543 | 0.00032 | 0.004521 | T | F | T | T |
| Usp12 | -0.5608151 | 6.73315471 | 16.3275803 | 0.00033 | 0.004615 | T | F | F | T |
| Sh3bp4 | 0.64497778 | 5.83906821 | 16.3237133 | 0.00033 | 0.004618 | F | F | T | F |
| Mafk | -0.7646508 | 5.37042772 | 16.2995691 | 0.00033 | 0.004654 | F | F | T | F |
| Tmem45a | 3.45150505 | 0.71112933 | 16.2966698 | 0.00033 | 0.004656 | T | F | T | T |
| Tia1 | 0.89973506 | 4.51015368 | 16.2945622 | 0.00033 | 0.004656 | T | F | T | T |
| Tet2 | -0.871736 | 4.7148965 | 16.2793955 | 0.00033 | 0.004675 | T | T | T | T |
| Zfp292 | 0.59396005 | 6.30992948 | 16.2791942 | 0.00033 | 0.004675 | T | F | F | T |
| Pygb | -0.492721 | 7.24304877 | 16.2673136 | 0.00033 | 0.004689 | F | F | F | T |
| Mageb16 | -0.8926228 | 4.66041528 | 16.2664661 | 0.00033 | 0.004689 | F | F | T | F |
| Psme3 | -0.4231964 | 8.25574881 | 16.2637646 | 0.00033 | 0.004689 | F | F | F | F |
| Lrrc61 | 1.54492135 | 2.68114558 | 16.2627792 | 0.00033 | 0.004689 | F | F | T | F |
| Dmc1 | 8.00911197 | -0.4221328 | 16.3851962 | 0.00033 | 0.004703 | T | F | T | T |
| Ralgps2 | 0.75787253 | 5.0488144 | 16.2403787 | 0.00034 | 0.004717 | T | F | T | T |
| Agrn | 0.40740913 | 8.48246542 | 16.2400681 | 0.00034 | 0.004717 | F | F | F | T |
| Tmem88 | 0.98660753 | 4.35047764 | 16.2211227 | 0.00034 | 0.004745 | T | F | T | T |
| Cdc34 | -0.4297851 | 8.23885913 | 16.2075844 | 0.00034 | 0.004764 | F | F | F | F |
| Soga3 | 1.87416784 | 2.26113134 | 16.204082 | 0.00034 | 0.004767 | F | F | F | F |
| Ahctf1 | -0.3889832 | 8.91094693 | 16.1898673 | 0.00034 | 0.004787 | F | F | T | T |
| Zfp467 | 1.88270033 | 2.15273844 | 16.1877531 | 0.00034 | 0.004787 | T | F | T | T |

| | | | | | | | | | |
|---|---|---|---|---|---|---|---|---|---|
| Sh3glb1 | -0.5580906 | 6.69114427 | 16.1426138 | 0.00035 | 0.00486 | T | F | T | T |
| mt-Nd2 | -0.446114 | 7.95269829 | 16.1190085 | 0.00035 | 0.004896 | F | F | F | F |
| Klf3 | 1.10233135 | 3.81589788 | 16.1062532 | 0.00035 | 0.004915 | T | F | T | T |
| Rab25 | 1.58357949 | 2.56011396 | 16.0919567 | 0.00035 | 0.004936 | T | F | T | T |
| Nectin4 | -1.9998423 | 2.32197 | 16.0872621 | 0.00035 | 0.004941 | F | F | T | F |
| Fgd3 | -1.2127017 | 3.70783987 | 16.0845307 | 0.00035 | 0.004942 | F | F | T | T |
| Nipa2 | -0.6186661 | 6.19850508 | 16.0734726 | 0.00036 | 0.004958 | F | F | F | F |
| Otx1 | 3.25475671 | 0.7786481 | 16.0689072 | 0.00036 | 0.004963 | F | F | F | F |
| Smarca1 | 0.81786945 | 4.6726885 | 16.0619399 | 0.00036 | 0.004972 | T | T | F | T |
| Lamb3 | 3.02230271 | 0.94429515 | 16.0107088 | 0.00036 | 0.005055 | F | F | F | T |
| Fzd1 | 1.07808238 | 3.932601 | 16.0106102 | 0.00036 | 0.005055 | T | F | T | T |
| Cfdp1 | -0.4897699 | 7.22981702 | 15.9997851 | 0.00037 | 0.00507 | F | F | T | F |
| Shank2 | 1.75339364 | 2.28924703 | 15.9924403 | 0.00037 | 0.005079 | F | F | T | T |
| Trim44 | 0.49960848 | 6.96599089 | 15.9911134 | 0.00037 | 0.005079 | T | T | T | T |
| Boc | 1.51747334 | 2.66788851 | 15.9875823 | 0.00037 | 0.005082 | T | T | F | T |
| Pfkfb2 | -1.2076567 | 3.93664479 | 15.9817686 | 0.00037 | 0.005089 | T | T | T | T |
| Pld1 | -1.0339427 | 4.14273511 | 15.9644935 | 0.00037 | 0.005117 | T | F | T | T |
| Nsg1 | 3.89658835 | 0.42057928 | 15.9407671 | 0.00037 | 0.005156 | T | F | T | F |
| Cep97 | 0.81361683 | 4.70835452 | 15.9020435 | 0.00038 | 0.005222 | T | F | T | T |
| Jade3 | -0.5328282 | 7.01497705 | 15.88987 | 0.00038 | 0.005239 | F | F | T | F |
| Dgki | 3.75637768 | 0.56757069 | 15.8891243 | 0.00038 | 0.005239 | T | F | T | T |
| Pdk3 | 0.90498654 | 4.3108975 | 15.8865914 | 0.00038 | 0.005241 | F | F | F | T |
| Zwint | -0.4873725 | 7.26884873 | 15.8765459 | 0.00038 | 0.00525 | T | F | T | T |
| Abi1 | -0.7084591 | 5.69701047 | 15.8753742 | 0.00038 | 0.00525 | T | T | F | T |
| Ugt8a | 5.22232734 | 0.02092472 | 15.8751148 | 0.00038 | 0.00525 | T | F | T | T |
| Hba-x | -5.4947371 | 3.12244705 | 15.8738868 | 0.00038 | 0.00525 | F | F | F | F |
| Smad5 | 0.69208549 | 5.52611079 | 15.8722863 | 0.00038 | 0.00525 | T | F | T | T |
| Stag3 | 1.9164082 | 2.28305676 | 15.8499422 | 0.00039 | 0.005287 | F | F | F | F |
| Adgrg6 | -0.9789108 | 4.29245301 | 15.8486895 | 0.00039 | 0.005287 | T | F | T | T |
| Ctdspl | -0.5561272 | 6.63849943 | 15.8329725 | 0.00039 | 0.005312 | T | F | T | T |
| Map2k3 | -0.597112 | 6.33372975 | 15.7919572 | 0.00039 | 0.005386 | F | F | T | T |
| Ppp4r4 | 3.45920456 | 0.64768426 | 15.7888989 | 0.00039 | 0.005388 | T | F | T | T |
| Spaar | -2.7906512 | 1.63717961 | 15.7803254 | 0.00039 | 0.0054 | T | F | T | F |
| Pphln1 | 0.50695483 | 6.80476881 | 15.7787365 | 0.00039 | 0.0054 | T | F | T | T |
| Jade1 | -0.4404891 | 8.04717454 | 15.7771833 | 0.00039 | 0.0054 | T | F | T | T |
| Gm8113 | 0.94213357 | 4.15051407 | 15.7664466 | 0.0004 | 0.005417 | F | F | F | F |
| Ppp1r26 | 1.12008225 | 3.72919819 | 15.7637838 | 0.0004 | 0.005417 | T | F | F | F |
| Btrc | 0.81140847 | 4.66415032 | 15.7629488 | 0.0004 | 0.005417 | T | F | T | F |
| Pde8b | 7.96489284 | -0.4594246 | 15.8716284 | 0.0004 | 0.005438 | T | F | T | T |
| Srd5a2 | 7.96489284 | -0.4594246 | 15.8716284 | 0.0004 | 0.005438 | F | F | F | F |
| Gm45623 | 7.96489284 | -0.4594246 | 15.8716284 | 0.0004 | 0.005438 | F | F | F | F |
| Smad4 | 0.52150273 | 6.76930149 | 15.7432904 | 0.0004 | 0.005438 | T | F | T | T |

| | | | | | | | | | |
|---|---|---|---|---|---|---|---|---|---|
| Gria4 | 2.52772441 | 1.32254151 | 15.7426884 | 0.0004 | 0.005438 | T | T | T | F |
| Mark4 | 0.69858924 | 5.41757759 | 15.738681 | 0.0004 | 0.005442 | F | F | T | F |
| Fkbp4 | 0.35765923 | 9.28614444 | 15.7266919 | 0.0004 | 0.005462 | T | F | F | T |
| Iqsec2 | 1.56259737 | 2.53742435 | 15.7181368 | 0.0004 | 0.005475 | T | F | T | T |
| Dnah9 | 7.96179115 | -0.4620339 | 15.8361137 | 0.0004 | 0.005485 | F | F | F | F |
| Wdr6 | 0.49914821 | 7.17432932 | 15.705136 | 0.00041 | 0.005493 | F | F | F | F |
| Pde1b | 1.66302596 | 2.35570757 | 15.6765558 | 0.00041 | 0.005544 | T | F | T | T |
| Fgfr2 | 0.55683376 | 6.47001501 | 15.6642853 | 0.00041 | 0.005565 | T | F | F | T |
| Mthfd2 | -0.6648636 | 6.38703625 | 15.6586109 | 0.00041 | 0.005571 | F | F | F | F |
| Tnfrsf9 | -1.5415293 | 3.30762677 | 15.6575676 | 0.00041 | 0.005571 | T | F | F | F |
| Nab1 | -0.9447279 | 4.39055765 | 15.6518178 | 0.00041 | 0.005579 | T | F | T | T |
| Eif2s2 | -0.4397865 | 8.37425243 | 15.6478698 | 0.00041 | 0.005583 | T | F | F | F |
| Arhgef15 | 1.4384804 | 2.78967171 | 15.6396707 | 0.00041 | 0.005595 | T | F | F | T |
| Nin | -0.5614419 | 6.59848112 | 15.6209042 | 0.00042 | 0.005629 | T | T | T | T |
| Pdlim1 | -0.5936169 | 6.34447522 | 15.6117407 | 0.00042 | 0.005644 | T | F | F | T |
| Klf13 | -0.4882749 | 7.47089992 | 15.6054961 | 0.00042 | 0.005652 | F | F | T | F |
| Plcxd2 | -1.0383253 | 4.11234365 | 15.5941445 | 0.00042 | 0.005671 | F | F | T | T |
| Armcx6 | 3.81365055 | 0.56840534 | 15.5903242 | 0.00042 | 0.005675 | T | F | F | T |
| Snx2 | -0.5184769 | 6.85737088 | 15.5752577 | 0.00042 | 0.005702 | T | T | F | T |
| Sertad2 | -0.6541708 | 5.89832344 | 15.5724319 | 0.00042 | 0.005704 | T | F | T | T |
| Lrrc56 | 2.08714745 | 1.78732728 | 15.5647159 | 0.00043 | 0.005716 | F | F | T | F |
| Dcun1d5 | -0.5927787 | 6.31348849 | 15.5532204 | 0.00043 | 0.005736 | F | F | T | T |
| Gata5 | 2.65002369 | 1.23764698 | 15.5473446 | 0.00043 | 0.005744 | F | F | T | F |
| Tmem121 | 2.25147718 | 1.63787388 | 15.5195004 | 0.00043 | 0.005797 | T | F | T | F |
| Zfp42 | -0.9311107 | 4.42183776 | 15.5101785 | 0.00043 | 0.005812 | T | T | F | T |
| Crocc | 0.64540201 | 5.84954827 | 15.4939005 | 0.00044 | 0.005842 | F | F | F | F |
| Slco4a1 | -1.6287174 | 2.80107496 | 15.4906906 | 0.00044 | 0.005845 | T | F | F | T |
| Epha4 | 1.38106737 | 3.14112974 | 15.4858188 | 0.00044 | 0.005851 | T | F | T | T |
| Actg1 | -0.285485 | 11.8659861 | 15.4672087 | 0.00044 | 0.005886 | F | F | F | F |
| Klf1 | -2.8138806 | 2.35359444 | 15.4618178 | 0.00044 | 0.005894 | F | F | F | F |
| Mxra8 | 1.14517112 | 3.51688229 | 15.4599558 | 0.00044 | 0.005894 | F | F | F | T |
| Tagln | -1.6048992 | 2.95772961 | 15.4543771 | 0.00044 | 0.005902 | F | F | F | F |
| Syt3 | 4.80083697 | 0.15154783 | 15.4459734 | 0.00044 | 0.005916 | F | F | F | T |
| Nfil3 | 1.2943839 | 3.15211424 | 15.4273765 | 0.00045 | 0.005951 | F | F | T | F |
| Coro1a | 2.76316691 | 1.20397994 | 15.3663507 | 0.00046 | 0.006077 | F | F | F | F |
| Itga2b | -1.8159427 | 3.63420608 | 15.3615312 | 0.00046 | 0.006083 | F | F | F | F |
| Ifi35 | -2.3641312 | 1.97909076 | 15.3496762 | 0.00046 | 0.006105 | F | F | F | F |
| Traf3ip1 | 0.96336171 | 4.16360791 | 15.3329358 | 0.00046 | 0.006138 | F | F | F | T |
| Zmat3 | 0.62671567 | 5.82887912 | 15.3263385 | 0.00046 | 0.006148 | T | F | T | T |
| Gabra3 | 4.23674122 | 0.22585097 | 15.3160206 | 0.00046 | 0.006167 | T | F | T | T |
| Msgn1 | 2.5989275 | 1.57461196 | 15.3078684 | 0.00047 | 0.006181 | F | F | F | F |
| Ttc7b | -0.5960991 | 6.24622617 | 15.2901522 | 0.00047 | 0.006215 | F | F | T | F |

| | | | | | | | | | |
|---|---|---|---|---|---|---|---|---|---|
| Nisch | 0.35960761 | 9.09894781 | 15.2889433 | 0.00047 | 0.006215 | T | F | T | T |
| H2bc9 | 0.46561425 | 7.19881277 | 15.274923 | 0.00047 | 0.006242 | F | F | F | F |
| Cux1 | 0.64454896 | 5.74077382 | 15.2612076 | 0.00047 | 0.006268 | T | F | T | T |
| Ttc6 | 7.91927541 | -0.4977139 | 15.3559684 | 0.00048 | 0.006313 | F | F | F | F |
| Gfra1 | 7.91927541 | -0.4977139 | 15.3559684 | 0.00048 | 0.006313 | T | F | T | T |
| Vwa5a | 1.23269986 | 3.2503569 | 15.2194844 | 0.00048 | 0.00635 | T | F | F | T |
| Wdr62 | -0.7171921 | 5.50762031 | 15.2069379 | 0.00048 | 0.006374 | F | F | F | F |
| Ppt2 | 0.66023177 | 5.62334778 | 15.2000136 | 0.00048 | 0.006383 | F | F | F | F |
| Arhgap21 | -0.476513 | 7.27311611 | 15.1996697 | 0.00048 | 0.006383 | F | F | T | F |
| Mif4gd | -0.8400728 | 4.71565954 | 15.1688866 | 0.00049 | 0.006449 | F | F | F | F |
| Psors1c2 | 2.01983625 | 1.8924813 | 15.1409783 | 0.00049 | 0.006509 | F | F | F | F |
| Eppk1 | 1.0882662 | 3.81937995 | 15.1338341 | 0.0005 | 0.006522 | F | F | F | F |
| Rbm38 | -0.5155246 | 7.1242433 | 15.1270358 | 0.0005 | 0.006533 | F | F | T | F |
| Ccnb1ip1 | 0.30710895 | 12.3830608 | 15.1255334 | 0.0005 | 0.006533 | F | F | F | F |
| Mef2c | 2.78176216 | 1.06019115 | 15.117955 | 0.0005 | 0.006547 | T | F | T | T |
| Igfbpl1 | 2.68208859 | 1.5187608 | 15.1092889 | 0.0005 | 0.006559 | F | F | T | F |
| Resf1 | 0.6949094 | 5.66106286 | 15.1092716 | 0.0005 | 0.006559 | F | F | T | F |
| Gm49909 | -0.494421 | 6.98468005 | 15.0928185 | 0.0005 | 0.006593 | F | F | F | F |
| Tuba1b | 0.33796976 | 10.6075018 | 15.088988 | 0.0005 | 0.006598 | F | F | F | T |
| C130026I21 | -7.834273 | 0.27341389 | 15.0855605 | 0.0005 | 0.006601 | F | F | F | F |
| Klhl24 | 0.71511467 | 5.18589535 | 15.0845786 | 0.0005 | 0.006601 | T | F | T | T |
| Baiap2 | -0.7391987 | 5.30082782 | 15.0818381 | 0.0005 | 0.006601 | F | F | T | F |
| Msh2 | 0.46222675 | 7.14390149 | 15.0809103 | 0.0005 | 0.006601 | F | F | T | F |
| Cox5a | -0.5261562 | 6.80099037 | 15.0531667 | 0.00051 | 0.006663 | F | F | F | F |
| Srl | 0.97785018 | 4.0289723 | 15.0444265 | 0.00051 | 0.00668 | T | T | T | T |
| Dst | 0.37145516 | 9.13919381 | 15.0393689 | 0.00051 | 0.006688 | T | F | F | F |
| Bicd2 | -0.576994 | 6.37113341 | 15.0228775 | 0.00052 | 0.006721 | F | F | T | F |
| Fam83h | -0.6017699 | 6.14550788 | 15.0220005 | 0.00052 | 0.006721 | F | F | T | F |
| Stum | 4.33853088 | 0.28564391 | 15.0040981 | 0.00052 | 0.00676 | T | F | T | T |
| Hrct1 | -5.0932337 | 0.66128196 | 14.9878366 | 0.00052 | 0.006795 | T | F | T | T |
| Slc29a4 | 1.96668208 | 1.92675243 | 14.9732706 | 0.00052 | 0.006826 | F | F | F | F |
| Srgap2 | -0.5242158 | 6.79141355 | 14.9716279 | 0.00052 | 0.006826 | T | F | F | T |
| Cacna1b | 1.9414704 | 2.04019105 | 14.9567848 | 0.00053 | 0.006858 | T | F | T | T |
| Spata21 | -4.356818 | 0.81032357 | 14.9517098 | 0.00053 | 0.006867 | F | F | F | F |
| Ctsh | -0.6294663 | 6.2529482 | 14.9493602 | 0.00053 | 0.006868 | F | F | T | F |
| Il17rb | -4.3486074 | 0.82659564 | 14.9318499 | 0.00053 | 0.006907 | F | F | F | F |
| Adgre5 | 0.97202032 | 4.05839236 | 14.9073796 | 0.00054 | 0.006963 | F | F | F | F |
| Enah | 0.43988984 | 7.47173326 | 14.9014156 | 0.00054 | 0.006972 | T | F | T | T |
| Prrg4 | 0.96781578 | 4.00741972 | 14.9007403 | 0.00054 | 0.006972 | T | F | T | T |
| Erich2 | 7.8879317 | -0.5239133 | 15.009885 | 0.00054 | 0.006978 | F | F | F | F |
| Adprh | -0.5801123 | 6.30070675 | 14.8854182 | 0.00054 | 0.007001 | F | F | F | F |
| Macf1 | 0.35344712 | 9.40959871 | 14.8833914 | 0.00054 | 0.007001 | T | T | F | T |

| | | | | | | | | | |
|---|---|---|---|---|---|---|---|---|---|
| Zfp282 | 0.76942699 | 4.77850175 | 14.8823148 | 0.00054 | 0.007001 | T | F | T | T |
| Acan | 3.32183647 | 0.66718496 | 14.8779699 | 0.00054 | 0.007008 | F | F | F | F |
| Etv4 | 0.85044987 | 4.55073712 | 14.8751925 | 0.00054 | 0.00701 | F | F | F | F |
| Ms4a4d | 2.99658932 | 0.85960349 | 14.866646 | 0.00054 | 0.007028 | T | T | F | T |
| Stk36 | 1.51804515 | 2.55245094 | 14.8606832 | 0.00055 | 0.007038 | F | F | T | F |
| Rpl22l1 | -0.7164311 | 5.80711712 | 14.8541464 | 0.00055 | 0.007051 | T | F | F | F |
| Pcolce2 | 2.43583222 | 1.33709307 | 14.839017 | 0.00055 | 0.007082 | T | F | F | T |
| Suz12 | -0.4903449 | 7.39177992 | 14.838605 | 0.00055 | 0.007082 | T | F | T | F |
| Fli1 | -1.1202107 | 3.97071249 | 14.8088676 | 0.00056 | 0.007153 | T | F | F | T |
| Slc27a3 | 0.76472053 | 4.87748024 | 14.8004205 | 0.00056 | 0.007171 | F | F | F | F |
| Fosb | -1.4530278 | 3.11980893 | 14.7946802 | 0.00056 | 0.007181 | F | F | F | T |
| Slain1 | 1.27345734 | 3.11083858 | 14.7816957 | 0.00056 | 0.00721 | T | F | F | F |
| Nefm | 0.87240242 | 4.58316156 | 14.7737283 | 0.00056 | 0.007227 | F | F | F | F |
| Dmrtc1b | -7.8166514 | 0.26672861 | 14.7606451 | 0.00057 | 0.007256 | T | F | F | F |
| Nphp4 | 1.8479712 | 2.10429819 | 14.7514185 | 0.00057 | 0.007276 | F | F | F | F |
| Dnajc13 | -0.3976181 | 8.35782608 | 14.7462402 | 0.00057 | 0.007285 | T | F | F | F |
| Rtn2 | 1.65042428 | 2.30545719 | 14.7189671 | 0.00057 | 0.007352 | F | F | T | F |
| Cdk18 | 0.97245637 | 3.94961322 | 14.7121444 | 0.00058 | 0.007366 | F | F | T | F |
| Limk1 | 0.51845129 | 6.59980066 | 14.6994842 | 0.00058 | 0.007395 | F | F | T | F |
| Ar | 3.3816055 | 0.75594383 | 14.6958327 | 0.00058 | 0.0074 | F | T | T | T |
| Bmp2k | -0.7399598 | 5.21610251 | 14.6941483 | 0.00058 | 0.007401 | T | F | F | F |
| Gm2381 | -1.1679734 | 3.6744695 | 14.6916864 | 0.00058 | 0.007403 | F | F | F | T |
| Dgkh | 0.89700189 | 4.22860596 | 14.6874977 | 0.00058 | 0.00741 | F | F | T | F |
| Rhox1 | -2.0048116 | 2.28246952 | 14.6747863 | 0.00058 | 0.007439 | T | F | T | T |
| Ptch1 | 0.45637494 | 7.18147483 | 14.6712228 | 0.00058 | 0.007444 | F | F | T | T |
| Actn3 | 2.26746312 | 1.53039956 | 14.6557407 | 0.00059 | 0.007481 | F | F | T | F |
| Vopp1 | 1.56679646 | 2.44654621 | 14.653964 | 0.00059 | 0.007482 | T | F | T | T |
| Itm2c | 0.62198855 | 5.72670149 | 14.6397553 | 0.00059 | 0.007515 | F | F | T | F |
| Creb3 | -0.689904 | 5.51164821 | 14.6383058 | 0.00059 | 0.007515 | F | F | F | F |
| Taf1 | -0.4752108 | 7.13493684 | 14.6249196 | 0.00059 | 0.007547 | T | F | T | T |
| Ppargc1b | -0.9296959 | 4.41011298 | 14.6108647 | 0.0006 | 0.007581 | F | F | F | T |
| Six5 | 1.45372455 | 2.66451426 | 14.6062631 | 0.0006 | 0.007587 | T | F | T | F |
| Cap1 | -0.402378 | 8.18186846 | 14.6054475 | 0.0006 | 0.007587 | F | F | T | F |
| Phf21b | 1.69446034 | 2.53097861 | 14.6025814 | 0.0006 | 0.00759 | T | F | T | F |
| P3h4 | 1.23081676 | 3.20190639 | 14.5928049 | 0.0006 | 0.007612 | F | F | F | F |
| Zfp703 | 0.74394524 | 5.28972115 | 14.5738848 | 0.00061 | 0.00766 | F | F | T | F |
| Gdf11 | 1.59036096 | 2.38728326 | 14.5319992 | 0.00061 | 0.007772 | F | F | T | T |
| Zfp839 | 1.2291091 | 3.19156807 | 14.5235116 | 0.00062 | 0.007791 | F | F | F | F |
| Cpeb4 | -0.6714595 | 5.60343745 | 14.5175331 | 0.00062 | 0.007803 | F | F | T | F |
| Gypc | -1.4528434 | 3.14373357 | 14.5107947 | 0.00062 | 0.007818 | F | F | T | F |
| Lactb | -0.8418561 | 4.74182262 | 14.5064404 | 0.00062 | 0.007825 | F | F | F | F |
| Itsn2 | -0.584252 | 6.23717909 | 14.4957069 | 0.00062 | 0.007851 | T | F | F | T |

| | | | | | | | | | |
|---|---|---|---|---|---|---|---|---|---|
| Kcnn2 | 7.84997155 | -0.555522 | 14.5996078 | 0.00062 | 0.007858 | F | F | F | F |
| Adrb3 | 2.58284136 | 1.20303591 | 14.4761407 | 0.00063 | 0.007898 | T | F | F | T |
| Dnajc10 | 0.47062569 | 6.969062 | 14.4449034 | 0.00063 | 0.007982 | T | F | T | T |
| Sun2 | -0.8501008 | 4.61472122 | 14.4433287 | 0.00063 | 0.007982 | T | F | T | F |
| Akr1b10 | 1.82319243 | 2.0269139 | 14.4322575 | 0.00064 | 0.00801 | F | F | F | F |
| Zik1 | 1.05359295 | 3.67321697 | 14.4294202 | 0.00064 | 0.008013 | F | F | T | T |
| Phf8 | 0.65957217 | 5.44175272 | 14.4030498 | 0.00064 | 0.008083 | T | T | F | T |
| Akap1 | -0.477051 | 7.0266854 | 14.4023117 | 0.00064 | 0.008083 | F | F | T | F |
| 1700031F05 | -5.0375083 | 0.60047695 | 14.3963768 | 0.00065 | 0.008095 | F | F | F | F |
| Sh3bgrl3 | -0.6607095 | 5.67657225 | 14.3843297 | 0.00065 | 0.008126 | F | F | T | F |
| Rinl | -0.8978065 | 4.6816531 | 14.3658059 | 0.00065 | 0.008176 | T | F | T | T |
| Igsf1 | 1.78999566 | 2.22358236 | 14.3505975 | 0.00066 | 0.008216 | T | F | T | F |
| Ppm1e | 3.49364433 | 0.51598232 | 14.3446639 | 0.00066 | 0.008225 | T | F | T | T |
| Map7d1 | -0.4685548 | 7.68496426 | 14.3445055 | 0.00066 | 0.008225 | F | F | F | T |
| Nhs | 1.15810313 | 3.46270154 | 14.3404356 | 0.00066 | 0.008232 | T | T | F | T |
| Zbtb5 | 1.05287313 | 3.68612388 | 14.3347018 | 0.00066 | 0.008245 | T | F | T | T |
| Top1mt | -0.8473504 | 4.61833674 | 14.3317106 | 0.00066 | 0.008249 | F | F | F | T |
| Prox1 | 1.20115361 | 3.27949258 | 14.3272211 | 0.00066 | 0.008258 | T | F | T | T |
| Cdh6 | 1.03550131 | 3.79696112 | 14.3201054 | 0.00066 | 0.008273 | F | F | F | T |
| Tmem98 | 1.09900375 | 3.56148797 | 14.3190688 | 0.00066 | 0.008273 | F | F | F | F |
| Arcn1 | -0.4073382 | 8.16548694 | 14.3032215 | 0.00067 | 0.008315 | T | F | T | T |
| Rbp4 | -0.404416 | 8.09436631 | 14.2921768 | 0.00067 | 0.008344 | F | F | T | F |
| Ift81 | 1.42679731 | 2.67142033 | 14.2817611 | 0.00067 | 0.00837 | F | F | F | F |
| 2410004P03 | 2.37727968 | 1.36956847 | 14.280504 | 0.00067 | 0.00837 | T | F | F | F |
| Zfp800 | -0.8598184 | 4.54775507 | 14.2754475 | 0.00067 | 0.008379 | T | F | T | T |
| Jmjd4 | 1.06093396 | 4.02290294 | 14.2742951 | 0.00067 | 0.008379 | T | F | T | T |
| Ate1 | -0.5354638 | 6.55214696 | 14.2682309 | 0.00068 | 0.008393 | T | F | T | T |
| Aldh7a1 | -0.4278956 | 7.72055337 | 14.2645958 | 0.00068 | 0.008399 | T | F | T | T |
| Trib3 | -1.9734643 | 3.49294635 | 14.2618945 | 0.00068 | 0.008402 | F | F | F | F |
| Dnah8 | -8.631823 | 1.06262428 | 14.2493009 | 0.00068 | 0.008436 | T | F | F | T |
| Itm2a | 2.1557756 | 1.81181122 | 14.2400203 | 0.00068 | 0.00846 | T | F | F | F |
| Kcnh8 | 7.82480771 | -0.5764012 | 14.3329296 | 0.00069 | 0.00849 | T | F | F | F |
| Foxd3 | 7.82347106 | -0.5775086 | 14.3188815 | 0.00069 | 0.008523 | T | F | T | T |
| Ankrd35 | 7.82347106 | -0.5775086 | 14.3188815 | 0.00069 | 0.008523 | F | F | F | F |
| Gap43 | 3.48079643 | 0.66317417 | 14.2117694 | 0.00069 | 0.008527 | F | F | T | F |
| Pofut1 | 0.63395039 | 5.5921233 | 14.2015543 | 0.00069 | 0.008554 | T | F | F | T |
| Pou2f3 | -2.0498061 | 2.14765037 | 14.1997129 | 0.00069 | 0.008555 | T | T | F | T |
| Bcl2 | -1.1939522 | 3.61139193 | 14.1919099 | 0.00069 | 0.008574 | T | F | T | F |
| Mapk13 | -1.406842 | 3.26774007 | 14.1525121 | 0.0007 | 0.008693 | F | F | F | T |
| Rap1b | -0.5190018 | 6.66460667 | 14.1443083 | 0.00071 | 0.008714 | T | F | T | F |
| Tmem254a | 2.24287914 | 1.52505961 | 14.1313957 | 0.00071 | 0.00875 | F | F | T | F |
| Scyl3 | -0.8310104 | 4.67681922 | 14.1234094 | 0.00071 | 0.008771 | T | F | F | F |

| | | | | | | | | | |
|---|---|---|---|---|---|---|---|---|---|
| Pim1 | -0.5738851 | 6.22905519 | 14.1192909 | 0.00071 | 0.008776 | F | F | T | F |
| Dcun1d4 | -0.6196933 | 5.88658834 | 14.1178535 | 0.00071 | 0.008776 | T | T | T | T |
| Megf8 | 0.48487797 | 6.84636343 | 14.115702 | 0.00071 | 0.008776 | T | F | F | F |
| Dcun1d2 | -0.8066341 | 4.72356038 | 14.1155518 | 0.00071 | 0.008776 | T | F | T | F |
| Gm20431 | 2.51178131 | 2.89961099 | 14.110934 | 0.00072 | 0.008786 | F | F | F | F |
| Vps37b | -0.640975 | 5.75454303 | 14.0924541 | 0.00072 | 0.00884 | T | F | T | F |
| Oscp1 | 1.72797759 | 2.12441553 | 14.0838069 | 0.00072 | 0.008863 | F | F | T | F |
| Gfi1b | -1.520172 | 2.911327 | 14.0785695 | 0.00072 | 0.008875 | T | F | F | T |
| Ftl1 | -0.2990931 | 10.7781898 | 14.0762942 | 0.00072 | 0.008877 | F | F | F | F |
| Tax1bp3 | -0.6021476 | 6.02192278 | 14.0561925 | 0.00073 | 0.008938 | F | F | T | F |
| Gpat3 | -0.918257 | 4.30492388 | 14.033785 | 0.00074 | 0.009006 | T | F | F | F |
| Igsf9b | 0.59201976 | 5.87166267 | 14.023375 | 0.00074 | 0.009035 | T | F | F | F |
| Fzd6 | 1.08874279 | 3.55705915 | 14.0169406 | 0.00074 | 0.009051 | T | F | T | T |
| Prkcsh | 0.43180373 | 7.76371219 | 14.0128902 | 0.00074 | 0.009059 | F | F | F | F |
| Dixdc1 | 1.34748594 | 2.84591785 | 14.0069078 | 0.00074 | 0.009074 | T | F | T | T |
| Zfp212 | 0.91263362 | 4.12946812 | 13.9807714 | 0.00075 | 0.009156 | F | F | F | F |
| Kcnma1 | 5.76159088 | -0.2831052 | 13.9623046 | 0.00076 | 0.009213 | T | F | T | T |
| Lgals2 | -0.9172898 | 4.33004372 | 13.9602624 | 0.00076 | 0.009215 | F | F | F | F |
| Rasa4 | 2.41698125 | 1.31562693 | 13.9562058 | 0.00076 | 0.009223 | F | F | F | F |
| Calca | 3.13996457 | 0.7518755 | 13.9533746 | 0.00076 | 0.009227 | F | F | F | F |
| Ankrd33b | -0.6171534 | 5.87796048 | 13.9467955 | 0.00076 | 0.009244 | T | F | T | T |
| Fgf4 | 2.9662025 | 0.92405013 | 13.9387938 | 0.00076 | 0.009266 | F | F | T | F |
| Anp32b | -0.3439353 | 9.41860678 | 13.9365575 | 0.00076 | 0.009269 | F | F | F | F |
| Epas1 | 0.43500992 | 7.32734186 | 13.9302036 | 0.00076 | 0.009285 | F | F | F | F |
| Irf2bp1 | 0.64734746 | 5.5046986 | 13.9227412 | 0.00077 | 0.009305 | F | F | F | F |
| Acox1 | -0.5427984 | 6.44188558 | 13.9209581 | 0.00077 | 0.009306 | T | F | T | T |
| Iqce | 1.00657033 | 3.77812054 | 13.9075906 | 0.00077 | 0.009346 | T | F | T | T |
| Egfr | -0.7377501 | 5.13378339 | 13.9053202 | 0.00077 | 0.009346 | T | F | T | T |
| Smg5 | -0.4194054 | 7.71727405 | 13.904722 | 0.00077 | 0.009346 | F | F | T | F |
| Cpne1 | 0.57626 | 6.04048522 | 13.8958737 | 0.00077 | 0.009371 | T | F | T | F |
| Faf1 | -0.4466226 | 7.30667823 | 13.893392 | 0.00077 | 0.009374 | T | F | T | T |
| Ppfia4 | 1.19165866 | 3.26414502 | 13.8888662 | 0.00078 | 0.009385 | T | F | T | F |
| Peli3 | 1.35815242 | 2.79239903 | 13.880989 | 0.00078 | 0.009407 | F | F | T | F |
| Hpcal1 | -0.8432253 | 4.55154878 | 13.8577972 | 0.00078 | 0.009482 | T | F | F | F |
| Acvr2b | 0.4730794 | 6.84568598 | 13.8550318 | 0.00079 | 0.009486 | F | F | T | F |
| Wnt7b | -0.5718361 | 6.50883086 | 13.848909 | 0.00079 | 0.009502 | T | F | T | T |
| Zfp13 | 0.88892024 | 4.14768098 | 13.8456167 | 0.00079 | 0.009508 | F | F | F | F |
| St8sia2 | 2.53840655 | 1.14844847 | 13.8427755 | 0.00079 | 0.009513 | T | F | F | T |
| Cfap65 | 7.78685449 | -0.6077783 | 13.9386071 | 0.00079 | 0.009519 | F | F | F | F |
| Kmt5a | -0.4195514 | 7.77967913 | 13.835449 | 0.00079 | 0.009528 | F | F | T | F |
| Abcb6 | 0.64642343 | 5.42952283 | 13.8120374 | 0.0008 | 0.009605 | F | F | F | F |
| Aebp1 | 0.92203086 | 4.38393516 | 13.7927278 | 0.0008 | 0.009668 | F | F | F | F |

| | | | | | | | | | |
|---|---|---|---|---|---|---|---|---|---|
| Pagr1a | 1.37783545 | 4.79073116 | 13.7802085 | 0.00081 | 0.009707 | T | F | F | F |
| Ptprk | 0.49408642 | 6.66029196 | 13.7775735 | 0.00081 | 0.009711 | T | T | F | T |
| Sgk3 | -0.6436632 | 5.75212003 | 13.7714495 | 0.00081 | 0.009727 | T | F | F | F |
| Kpna1 | -0.533413 | 6.56534571 | 13.7581489 | 0.00081 | 0.00977 | T | F | T | T |
| Kcnb1 | 2.56540572 | 1.11708563 | 13.7385239 | 0.00082 | 0.009835 | F | F | T | T |
| Plxdc2 | 1.34388402 | 2.87722761 | 13.7329527 | 0.00082 | 0.009848 | F | T | T | T |
| Zfp358 | 1.13119017 | 3.38261365 | 13.7320216 | 0.00082 | 0.009848 | F | F | F | F |
| Cfc1 | 1.69959668 | 2.18533621 | 13.6995911 | 0.00083 | 0.00996 | F | F | F | F |
| Atf7 | -0.6100357 | 5.88233142 | 13.6687494 | 0.00084 | 0.010068 | T | F | T | T |
| Reln | -0.7380412 | 5.82976289 | 13.6617676 | 0.00084 | 0.010089 | T | F | T | F |
| Uhrf1bp1 | -0.4651692 | 7.01982118 | 13.6576778 | 0.00084 | 0.010098 | T | F | T | F |
| Cul7 | -0.3874866 | 8.60011998 | 13.6487247 | 0.00085 | 0.010126 | F | F | F | F |
| Sh3kbp1 | 1.15319889 | 3.29668262 | 13.647293 | 0.00085 | 0.010126 | T | F | T | T |
| Sel1l3 | 0.8101657 | 4.456283 | 13.6368359 | 0.00085 | 0.01016 | T | F | T | F |
| Fam89a | -1.0345293 | 3.96739024 | 13.6308433 | 0.00085 | 0.010176 | F | F | F | F |
| Fermt2 | -0.4439365 | 7.39406683 | 13.6293276 | 0.00085 | 0.010177 | T | F | F | F |
| Dnajb13 | 5.73480954 | -0.3072461 | 13.5862676 | 0.00087 | 0.010334 | F | F | F | T |
| Zfp646 | 0.46431821 | 6.86047749 | 13.5659727 | 0.00087 | 0.010406 | T | F | T | F |
| Bmp7 | 0.73649066 | 4.82973793 | 13.550408 | 0.00088 | 0.01046 | F | F | T | F |
| Eno2 | 7.75734378 | -0.6320796 | 13.6384878 | 0.00088 | 0.010483 | F | F | F | F |
| Lmx1b | 1.42566698 | 2.65820591 | 13.5398337 | 0.00088 | 0.01049 | F | F | T | F |
| Slc35g1 | -0.5300988 | 6.47106918 | 13.5357053 | 0.00088 | 0.0105 | F | F | T | F |
| Cyp20a1 | -0.6204696 | 5.90583891 | 13.506524 | 0.00089 | 0.010608 | F | T | F | T |
| Phf21a | 0.79139975 | 4.63066739 | 13.5012357 | 0.0009 | 0.010623 | F | F | T | T |
| Epb41l5 | 0.60684871 | 5.65310021 | 13.4909995 | 0.0009 | 0.010657 | F | F | T | F |
| Sema5b | 2.99156624 | 0.7563159 | 13.4820809 | 0.0009 | 0.010687 | T | F | T | F |
| Abcb10 | -0.6149298 | 6.25780838 | 13.4753844 | 0.0009 | 0.010708 | T | F | T | T |
| Mapre2 | 0.57609733 | 5.93887548 | 13.46965 | 0.00091 | 0.010722 | T | F | T | F |
| Me2 | -0.4412841 | 7.31890762 | 13.4681734 | 0.00091 | 0.010722 | T | F | T | T |
| Ep300 | 0.39014028 | 7.913772 | 13.4673617 | 0.00091 | 0.010722 | T | F | F | F |
| Pacsin2 | -0.4346218 | 7.37640953 | 13.4559459 | 0.00091 | 0.010761 | F | F | T | F |
| Nans | -0.495408 | 6.75961822 | 13.4282825 | 0.00092 | 0.010866 | F | F | F | F |
| Plin2 | -0.5754025 | 6.13129745 | 13.4252216 | 0.00092 | 0.010873 | F | F | T | F |
| H2ax | 0.33262323 | 9.23953813 | 13.4082518 | 0.00093 | 0.010936 | F | F | F | F |
| Fbxo44 | 2.1204949 | 1.6161018 | 13.3988601 | 0.00093 | 0.010968 | T | F | T | F |
| Camta2 | -0.7057175 | 5.41957588 | 13.3839979 | 0.00093 | 0.011017 | F | F | F | F |
| Gusb | -0.630247 | 5.69717489 | 13.3837405 | 0.00093 | 0.011017 | T | F | F | T |
| Hipk2 | 0.51893128 | 6.46980931 | 13.3796699 | 0.00094 | 0.011028 | T | F | F | T |
| Gas2l1 | -0.5677665 | 6.2842742 | 13.3505877 | 0.00095 | 0.011142 | F | F | T | T |
| Plekhg2 | 0.54284831 | 6.13712031 | 13.3475843 | 0.00095 | 0.011148 | F | F | F | F |
| Lad1 | -0.4771408 | 6.86849433 | 13.335313 | 0.00095 | 0.011186 | T | F | T | T |
| Ahnak2 | 2.37807321 | 1.34157767 | 13.3345886 | 0.00095 | 0.011186 | F | F | F | F |

| | | | | | | | | | |
|---|---|---|---|---|---|---|---|---|---|
| Ccni | 0.46929355 | 6.9649988 | 13.3342184 | 0.00095 | 0.011186 | T | F | T | T |
| Gm15107 | 7.73566331 | -0.6498787 | 13.4215787 | 0.00095 | 0.011202 | F | F | F | F |
| Eif2s3x | -0.341209 | 9.26921894 | 13.3252207 | 0.00096 | 0.011211 | T | F | T | T |
| Ccdc88a | 0.71985363 | 5.01644475 | 13.3167787 | 0.00096 | 0.01124 | T | T | T | T |
| Tbkbp1 | 1.05362228 | 3.54701931 | 13.3114794 | 0.00096 | 0.011256 | F | F | T | F |
| Gm4631 | 0.71633432 | 5.41118875 | 13.3061637 | 0.00096 | 0.011272 | F | F | F | F |
| Eif1 | -0.3698348 | 8.61791318 | 13.3005954 | 0.00096 | 0.01129 | F | F | T | F |
| Trim10 | -4.5023908 | 0.90316246 | 13.286882 | 0.00097 | 0.011337 | F | F | F | F |
| Arhgap8 | -1.0826893 | 4.02331606 | 13.2851874 | 0.00097 | 0.011337 | F | F | F | T |
| Cpne8 | -1.101644 | 3.69695939 | 13.2837639 | 0.00097 | 0.011337 | T | T | T | T |
| Col14a1 | 3.08994444 | 0.70369982 | 13.2834254 | 0.00097 | 0.011337 | T | F | T | T |
| Hist1h2an | 0.47129864 | 7.08962833 | 13.2749036 | 0.00097 | 0.011367 | F | F | F | F |
| Zfp608 | 0.95126697 | 4.45133811 | 13.2643713 | 0.00098 | 0.011406 | T | F | T | F |
| Ccp110 | 0.62483734 | 5.48795181 | 13.2464486 | 0.00098 | 0.011476 | T | F | T | F |
| Fam76a | -0.7012349 | 5.26161543 | 13.2433701 | 0.00099 | 0.011483 | F | F | T | F |
| Capzb | -0.4171578 | 7.65806923 | 13.2363791 | 0.00099 | 0.011507 | F | F | T | F |
| Coro1c | -0.3571764 | 8.79881698 | 13.2211228 | 0.00099 | 0.011566 | T | F | T | F |
| Castor2 | 0.54230197 | 6.1629883 | 13.2159122 | 0.001 | 0.011583 | T | F | T | T |
| Slc25a12 | -1.2607414 | 3.31272967 | 13.2141922 | 0.001 | 0.011584 | T | F | T | T |
| Ube2d1 | 0.89477191 | 4.17350167 | 13.2044997 | 0.001 | 0.01162 | T | T | F | T |
| Pfn2 | 1.44288072 | 2.51400356 | 13.1950197 | 0.001 | 0.011654 | F | F | T | F |
| Grip1 | 1.05116269 | 3.58059961 | 13.1821849 | 0.00101 | 0.011704 | T | F | F | F |
| E2f7 | -0.5023896 | 6.65839709 | 13.177512 | 0.00101 | 0.011718 | T | F | T | T |
| Kctd15 | -0.6642442 | 5.47469879 | 13.1759691 | 0.00101 | 0.011719 | F | F | T | T |
| Nmrk1 | -0.8429698 | 4.49000344 | 13.1703689 | 0.00101 | 0.011737 | F | F | T | F |
| Hdhd2 | -0.7690863 | 4.86038709 | 13.1595656 | 0.00102 | 0.011769 | T | F | T | T |
| Sec24a | -0.5048085 | 6.74205112 | 13.1591549 | 0.00102 | 0.011769 | T | F | T | F |
| Pcgf5 | -0.60681 | 5.82168612 | 13.1588348 | 0.00102 | 0.011769 | F | F | T | T |
| Zswim5 | 1.20308664 | 3.08403169 | 13.1369321 | 0.00102 | 0.011856 | T | F | F | F |
| Rsf1 | 0.76345651 | 4.5777613 | 13.1362134 | 0.00103 | 0.011856 | F | T | T | T |
| Fam114a1 | 1.29839959 | 2.84395557 | 13.1196509 | 0.00103 | 0.011923 | T | F | T | T |
| Pml | 0.69972134 | 4.9347252 | 13.116537 | 0.00103 | 0.011931 | T | F | T | F |
| Fhl2 | -0.8361143 | 4.48594278 | 13.0994573 | 0.00104 | 0.012001 | F | F | F | F |
| Zcchc3 | 0.51645766 | 6.3239057 | 13.0948226 | 0.00104 | 0.012015 | F | F | F | F |
| Pak3 | 3.70738284 | 0.26968111 | 13.086347 | 0.00104 | 0.012044 | F | T | T | T |
| Xlr4b | -1.0221572 | 3.87539864 | 13.0854997 | 0.00104 | 0.012044 | T | F | F | T |
| Nsun2 | -0.3527214 | 8.82760329 | 13.0820953 | 0.00105 | 0.01205 | T | F | T | F |
| Cxxc5 | 0.57047898 | 5.86875945 | 13.0815104 | 0.00105 | 0.01205 | T | F | F | T |
| Tnip1 | -0.56353 | 6.1237398 | 13.0798733 | 0.00105 | 0.012051 | T | F | T | F |
| Nt5c2 | 0.99335183 | 3.7079179 | 13.0771598 | 0.00105 | 0.012056 | F | F | T | T |
| Ier3 | -1.3941207 | 3.00904472 | 13.0568954 | 0.00106 | 0.012142 | F | F | F | F |
| Cplx2 | 1.44053122 | 2.61421681 | 13.0526418 | 0.00106 | 0.012155 | F | F | T | T |

| | | | | | | | | | |
|---|---|---|---|---|---|---|---|---|---|
| Nphp1 | 1.45879773 | 2.47543397 | 13.0482878 | 0.00106 | 0.012155 | F | F | F | T |
| S100a11 | -0.6596676 | 5.45561272 | 13.0476854 | 0.00106 | 0.012155 | F | F | F | T |
| Cbx1 | 0.58031503 | 5.78069831 | 13.0474839 | 0.00106 | 0.012155 | T | F | T | F |
| Hnf4a | -0.4288194 | 7.46040389 | 13.0469103 | 0.00106 | 0.012155 | T | F | F | T |
| Cadm3 | 1.85049324 | 2.16594945 | 13.0406414 | 0.00106 | 0.012177 | T | F | F | T |
| Ilf3 | 0.39530212 | 7.80532275 | 13.0210475 | 0.00107 | 0.012261 | T | F | T | F |
| Ggta1 | -0.5683513 | 6.07338875 | 13.01602 | 0.00107 | 0.012277 | F | F | T | F |
| Cnksr2 | 1.15793879 | 3.19924305 | 13.006977 | 0.00108 | 0.012312 | F | F | T | T |
| Pfkp | -0.638751 | 5.9727963 | 12.9919635 | 0.00108 | 0.012375 | F | F | F | F |
| Prelid3b | -0.562552 | 6.12598842 | 12.9892283 | 0.00108 | 0.01238 | T | F | T | F |
| Rpl26 | 0.35762831 | 8.45061275 | 12.988172 | 0.00108 | 0.01238 | F | F | F | T |
| Hrh3 | 7.6999586 | -0.6790897 | 13.0708922 | 0.00109 | 0.012399 | F | F | T | F |
| Kcnmb4 | 4.0509677 | 0.1491187 | 12.9807361 | 0.00109 | 0.012401 | T | F | T | F |
| Zc4h2 | 1.5309112 | 2.33996344 | 12.9752693 | 0.00109 | 0.012416 | T | F | T | T |
| Pitpna | -0.4680314 | 6.90650781 | 12.974887 | 0.00109 | 0.012416 | T | F | T | T |
| Rcor2 | 0.48791749 | 6.57576661 | 12.9676555 | 0.00109 | 0.012443 | F | F | F | F |
| Egflam | 1.16449183 | 3.55122626 | 12.9650833 | 0.00109 | 0.012448 | T | F | T | T |
| Hirip3 | 0.66396663 | 5.19393852 | 12.9546098 | 0.0011 | 0.012487 | F | F | F | F |
| Dld | -0.528764 | 6.56978436 | 12.9539523 | 0.0011 | 0.012487 | F | F | F | T |
| Dpp9 | 0.50533287 | 6.44810992 | 12.9512604 | 0.0011 | 0.01249 | T | F | F | F |
| Notch4 | 1.3601974 | 2.66964135 | 12.9506173 | 0.0011 | 0.01249 | F | F | F | F |
| Golim4 | 0.91223105 | 4.00396049 | 12.9469773 | 0.0011 | 0.0125 | T | F | T | T |
| Chmp2b | -0.6432402 | 5.60706845 | 12.9433143 | 0.0011 | 0.012511 | T | T | T | T |
| Armh4 | 0.9311726 | 3.91312514 | 12.9348888 | 0.00111 | 0.012544 | T | F | T | F |
| Ago1 | 0.58188775 | 5.77058065 | 12.9282702 | 0.00111 | 0.012569 | F | F | T | T |
| Ninl | 0.64364051 | 5.29723953 | 12.9233656 | 0.00111 | 0.012585 | F | F | F | F |
| Timm44 | -0.52389 | 6.44253331 | 12.916318 | 0.00111 | 0.012612 | F | F | T | F |
| Gnao1 | 1.04303304 | 3.52950116 | 12.9000983 | 0.00112 | 0.012683 | T | F | T | F |
| Trappc8 | -0.5106326 | 6.52988269 | 12.8942667 | 0.00112 | 0.012704 | F | F | T | T |
| Esyt3 | -1.0331185 | 3.97173507 | 12.8886539 | 0.00112 | 0.012722 | T | F | T | T |
| Kras | -0.5716793 | 6.18377078 | 12.8876433 | 0.00113 | 0.012722 | T | F | T | T |
| Tll2 | 4.02044662 | 0.10009463 | 12.8748478 | 0.00113 | 0.012777 | F | F | F | F |
| Lsr | 0.42742493 | 7.37713241 | 12.8730592 | 0.00113 | 0.012779 | T | F | F | T |
| mt-Atp6 | -0.5345701 | 9.07994494 | 12.8684051 | 0.00113 | 0.01279 | F | F | F | F |
| Ubxn2a | -0.7113779 | 5.10176146 | 12.8679697 | 0.00113 | 0.01279 | T | F | T | T |
| Rab39b | 3.24961114 | 0.55357554 | 12.8665536 | 0.00113 | 0.01279 | T | F | T | T |
| Slc25a30 | -0.701543 | 5.22145498 | 12.8585958 | 0.00114 | 0.012822 | F | F | F | F |
| Gpc2 | 1.24306647 | 2.955742 | 12.8340066 | 0.00115 | 0.012935 | T | F | F | F |
| Zbtb39 | 0.53457775 | 6.17941376 | 12.8232442 | 0.00115 | 0.012981 | T | F | T | T |
| Rtl8a | 0.85557695 | 4.1350799 | 12.8212157 | 0.00115 | 0.012984 | T | F | T | F |
| Creld1 | 1.13406875 | 3.25858678 | 12.8113169 | 0.00116 | 0.013025 | F | F | F | F |
| H2-K1 | 1.40191027 | 2.58674071 | 12.8017368 | 0.00116 | 0.013066 | F | F | F | F |

| | | | | | | | | | |
|---|---|---|---|---|---|---|---|---|---|
| Ctsf | 1.40922255 | 2.76216899 | 12.7981438 | 0.00116 | 0.013073 | F | F | F | F |
| Gpx8 | 1.36207207 | 2.72627771 | 12.797496 | 0.00116 | 0.013073 | F | F | F | F |
| Ppp1r13l | -0.6733596 | 5.35004444 | 12.7934323 | 0.00117 | 0.013086 | F | F | F | F |
| Gm49337 | 7.67991035 | -0.6954357 | 12.8775098 | 0.00117 | 0.013089 | F | F | F | F |
| Sept7 | -0.472325 | 6.96849913 | 12.7895525 | 0.00117 | 0.013092 | T | F | T | T |
| Zfp445 | 0.47038262 | 6.69160116 | 12.7780188 | 0.00117 | 0.013142 | T | F | T | T |
| AI467606 | -2.0188499 | 2.01577732 | 12.7744173 | 0.00117 | 0.013153 | F | F | F | F |
| C2cd3 | 0.55650359 | 5.96308225 | 12.7709933 | 0.00118 | 0.013164 | T | F | F | F |
| Vstm2l | 5.6936544 | -0.3864476 | 12.7684854 | 0.00118 | 0.013169 | T | F | T | T |
| Rapgef5 | 0.87720825 | 4.11085342 | 12.7634754 | 0.00118 | 0.013187 | F | F | T | T |
| Bcar3 | -0.7124642 | 5.07213477 | 12.7603106 | 0.00118 | 0.013196 | F | F | F | F |
| Cdhr5 | -1.6668704 | 2.67845068 | 12.7510369 | 0.00118 | 0.013236 | F | F | F | F |
| mt-Atp8 | -0.6500905 | 5.57598986 | 12.7480189 | 0.00119 | 0.013244 | F | F | F | F |
| Gypa | -5.107439 | 0.60024516 | 12.7247561 | 0.0012 | 0.013348 | T | F | F | F |
| Tmed2 | -0.3835589 | 8.07905133 | 12.724527 | 0.0012 | 0.013348 | T | F | T | F |
| Lars | -0.3933268 | 7.82790971 | 12.7231499 | 0.0012 | 0.013349 | F | F | F | F |
| Stxbp1 | 1.45255006 | 2.44247876 | 12.7158359 | 0.0012 | 0.013379 | T | F | F | T |
| Arhgap20 | -1.8022872 | 2.24181232 | 12.706038 | 0.00121 | 0.013421 | T | F | T | F |
| Pmpca | -0.4016195 | 7.78155088 | 12.7043785 | 0.00121 | 0.013423 | T | F | F | T |
| Rasal1 | -0.8871966 | 4.2475697 | 12.697868 | 0.00121 | 0.013449 | F | F | F | F |
| S100g | -0.7179159 | 5.01560881 | 12.6944023 | 0.00121 | 0.013453 | F | F | F | F |
| Sipa1l2 | -0.4296315 | 7.26861845 | 12.6943512 | 0.00121 | 0.013453 | F | F | F | F |
| Cacnb2 | 4.13436307 | 0.02025715 | 12.6744439 | 0.00122 | 0.013548 | T | F | F | F |
| Marcksl1 | 0.35100088 | 8.81222546 | 12.6718834 | 0.00122 | 0.013554 | F | F | F | F |
| Sox17 | 0.57486192 | 5.90602801 | 12.6641307 | 0.00122 | 0.013587 | T | F | F | F |
| Etv5 | 0.42939851 | 7.04766949 | 12.6529078 | 0.00123 | 0.013636 | T | F | T | T |
| Plekhn1 | 1.03182424 | 3.53971988 | 12.651995 | 0.00123 | 0.013636 | F | F | F | F |
| Rbbp7 | -0.3548532 | 8.91290817 | 12.6439786 | 0.00123 | 0.01367 | F | F | F | F |
| Tspyl3 | 2.08415802 | 1.55280211 | 12.642084 | 0.00123 | 0.013673 | T | F | F | F |
| Gpsm1 | 0.719271 | 4.75900019 | 12.6396219 | 0.00124 | 0.013679 | F | F | T | F |
| Eid2 | 1.13106146 | 3.22613192 | 12.632835 | 0.00124 | 0.013707 | F | F | F | F |
| Mtmr1 | -0.8285573 | 4.52263955 | 12.625844 | 0.00124 | 0.013736 | T | F | F | F |
| Adgrg1 | -1.0432536 | 4.80002567 | 12.6041886 | 0.00125 | 0.013842 | F | F | T | F |
| Serping1 | 1.2360911 | 3.00757812 | 12.6007087 | 0.00125 | 0.013854 | F | F | F | F |
| Dusp4 | 0.56894182 | 5.84407866 | 12.5927527 | 0.00126 | 0.013888 | F | F | T | F |
| Fbxo27 | -1.1096481 | 3.5892685 | 12.5901057 | 0.00126 | 0.013895 | F | F | T | F |
| Zdhhc15 | 0.77734956 | 4.43300526 | 12.5886259 | 0.00126 | 0.013896 | T | F | F | T |
| Caln1 | 3.70569793 | 0.21868955 | 12.5811792 | 0.00126 | 0.013928 | F | F | T | T |
| Fbln2 | 1.01449246 | 3.57829798 | 12.5767592 | 0.00127 | 0.013944 | F | F | F | F |
| Acadvl | -0.6486529 | 5.46966239 | 12.5754741 | 0.00127 | 0.013944 | F | F | F | F |
| Dars | -0.4334196 | 7.25532997 | 12.572417 | 0.00127 | 0.013953 | T | T | T | T |
| Gm45837 | 3.26185671 | 0.87382914 | 12.5455941 | 0.00128 | 0.014088 | F | F | F | F |

| | | | | | | | | | |
|---|---|---|---|---|---|---|---|---|---|
| Cdc42bpb | -0.3656928 | 8.36016832 | 12.5444282 | 0.00128 | 0.014088 | F | F | T | F |
| Akt2 | -0.4546173 | 6.97257851 | 12.5364656 | 0.00129 | 0.014118 | F | F | T | F |
| Sap18b | -0.4793191 | 6.7797653 | 12.5361622 | 0.00129 | 0.014118 | F | F | F | F |
| A2m | 1.13988353 | 4.48264811 | 12.533489 | 0.00129 | 0.014125 | F | F | F | F |
| Lxn | -0.676553 | 5.2643443 | 12.52991 | 0.00129 | 0.014137 | F | F | F | F |
| Spsb4 | 1.56293304 | 2.33268031 | 12.511684 | 0.0013 | 0.014228 | F | F | F | F |
| Cfap44 | 5.6484327 | -0.3781254 | 12.5084047 | 0.0013 | 0.014239 | F | F | F | F |
| Adam10 | -0.5317169 | 6.28371268 | 12.5041702 | 0.0013 | 0.014255 | T | F | F | T |
| Tmem229a | 2.37610553 | 1.58192214 | 12.5019104 | 0.0013 | 0.01426 | T | F | T | T |
| Tbc1d10a | -0.6223199 | 5.62106416 | 12.4929144 | 0.00131 | 0.014301 | F | F | F | T |
| Ganab | 0.34901051 | 8.60991179 | 12.4902157 | 0.00131 | 0.014309 | T | F | T | T |
| Dazap2 | -0.6433704 | 5.54954786 | 12.4842868 | 0.00131 | 0.014325 | T | F | F | T |
| Camk1g | 3.22295687 | 0.53316905 | 12.4836458 | 0.00131 | 0.014325 | F | F | T | F |
| Lrrk1 | -0.9293237 | 4.30030396 | 12.4831841 | 0.00131 | 0.014325 | F | F | T | T |
| Hsph1 | -0.3596225 | 8.76847038 | 12.4764162 | 0.00131 | 0.014355 | T | F | F | T |
| Tmem132d | 5.64554198 | -0.3804852 | 12.4737126 | 0.00132 | 0.014363 | F | F | F | F |
| Tnc | 2.47542477 | 1.08093949 | 12.47084 | 0.00132 | 0.014371 | T | F | F | F |
| Sestd1 | 0.72333641 | 4.6895331 | 12.4663266 | 0.00132 | 0.014389 | T | T | T | T |
| Dmrtc1c1 | -3.9763932 | 1.04040077 | 12.4641417 | 0.00132 | 0.014393 | F | F | F | F |
| Ppp1r12c | -0.542607 | 6.17278204 | 12.4293178 | 0.00134 | 0.014578 | F | F | F | F |
| Hmmr | -0.4656174 | 6.92804329 | 12.4219006 | 0.00134 | 0.014608 | T | F | T | T |
| Plpp4 | 7.64030842 | -0.7276042 | 12.5028788 | 0.00134 | 0.014608 | F | F | F | F |
| Nbeal2 | -0.6309059 | 5.62963485 | 12.4114447 | 0.00135 | 0.014656 | F | F | F | F |
| H1f3 | 0.30738491 | 9.61140271 | 12.3971893 | 0.00136 | 0.014728 | T | F | F | T |
| Gdap1l1 | 2.66304547 | 0.90707812 | 12.3857562 | 0.00136 | 0.014785 | F | F | F | F |
| Inpp5b | -0.6033375 | 5.77968471 | 12.3829602 | 0.00136 | 0.014793 | F | F | F | F |
| Vav2 | -0.4722127 | 6.75824159 | 12.3723286 | 0.00137 | 0.014846 | F | F | F | T |
| Plk3 | -0.6919536 | 5.1154712 | 12.3623515 | 0.00137 | 0.014895 | F | F | F | F |
| Arfgef1 | -0.4174256 | 7.70305559 | 12.344589 | 0.00138 | 0.014988 | T | T | T | T |
| Rapgef6 | -0.4751391 | 6.74731757 | 12.3434646 | 0.00138 | 0.014988 | T | F | F | T |
| Atp1a2 | 2.52970556 | 1.20119048 | 12.3384401 | 0.00139 | 0.015009 | T | F | T | T |
| Elovl6 | 0.57707604 | 5.80649925 | 12.326855 | 0.00139 | 0.015068 | T | F | T | T |
| Jarid2 | 0.41061343 | 7.25412966 | 12.3136007 | 0.0014 | 0.015137 | T | F | T | F |
| Lbp | 1.97770218 | 1.84469015 | 12.2975659 | 0.00141 | 0.015223 | F | F | T | F |
| Mindy1 | -0.7221202 | 4.90492327 | 12.2826016 | 0.00142 | 0.015303 | F | F | F | F |
| Akap8 | -0.5693391 | 6.23337129 | 12.2804864 | 0.00142 | 0.015307 | T | F | F | F |
| Hectd1 | -0.349892 | 8.61740614 | 12.2737329 | 0.00142 | 0.015339 | F | F | T | F |
| Nme6 | -0.8198481 | 4.46322103 | 12.2677001 | 0.00142 | 0.015363 | F | F | F | F |
| Jph1 | 1.05095692 | 3.44555986 | 12.2671309 | 0.00142 | 0.015363 | F | F | F | F |
| Gm3264 | -7.5920926 | 0.06690608 | 12.2622691 | 0.00143 | 0.015384 | F | F | F | F |
| H2bc14 | 0.41742348 | 7.78684588 | 12.2605223 | 0.00143 | 0.015386 | F | F | F | F |
| Tab2 | -0.4413233 | 7.08165719 | 12.2573253 | 0.00143 | 0.015398 | F | F | T | F |

| | | | | | | | | | |
|---|---|---|---|---|---|---|---|---|---|
| Cluh | -0.3845128 | 8.07041842 | 12.2516119 | 0.00143 | 0.015424 | T | F | F | F |
| Dcx | 4.11265713 | 0.193102 | 12.2499522 | 0.00143 | 0.015426 | T | F | T | T |
| Mmp17 | 1.18611974 | 3.245869 | 12.2476095 | 0.00143 | 0.015432 | F | F | T | T |
| Asb4 | 1.31562413 | 3.25950708 | 12.2307578 | 0.00144 | 0.015522 | T | F | F | F |
| Tmem250-p | 0.75658213 | 4.78139209 | 12.2297752 | 0.00144 | 0.015522 | F | F | T | F |
| Maml3 | 0.70169448 | 4.8612097 | 12.2284468 | 0.00145 | 0.015523 | F | F | T | T |
| Gpr146 | -1.5080239 | 2.64190848 | 12.2265365 | 0.00145 | 0.015526 | T | F | T | T |
| Max | -0.5361994 | 6.28754112 | 12.220025 | 0.00145 | 0.015557 | T | F | T | F |
| Slc2a2 | -0.5988806 | 6.00611907 | 12.200625 | 0.00146 | 0.01566 | F | F | T | F |
| Cad | 0.32148667 | 9.53515467 | 12.2002187 | 0.00146 | 0.01566 | F | F | F | F |
| Col4a6 | 1.44752038 | 2.42002156 | 12.1930018 | 0.00147 | 0.015696 | F | F | F | F |
| Pfkm | -0.5163165 | 6.37585357 | 12.1866183 | 0.00147 | 0.015727 | F | F | F | T |
| Zfp655 | -0.5592735 | 6.00365839 | 12.1839201 | 0.00147 | 0.015735 | T | F | T | T |
| Abhd8 | 1.03950771 | 3.4980977 | 12.1822279 | 0.00147 | 0.015738 | F | F | T | F |
| Derl3 | -1.4090543 | 3.04485147 | 12.1741767 | 0.00148 | 0.015779 | F | F | F | F |
| F3 | 0.78905353 | 4.38693644 | 12.1691947 | 0.00148 | 0.015801 | F | F | T | F |
| Cdc42 | -0.3785114 | 8.18024577 | 12.1526964 | 0.00149 | 0.015893 | T | F | T | F |
| Tor1aip2 | -0.3865262 | 7.84820533 | 12.1514415 | 0.00149 | 0.015893 | T | F | T | T |
| Hspbap1 | -0.5735618 | 5.92211988 | 12.1341463 | 0.0015 | 0.015984 | T | F | T | T |
| Nucb1 | 0.45361443 | 6.73526284 | 12.1331563 | 0.0015 | 0.015984 | T | F | T | F |
| Pga5 | 0.99217062 | 3.69166402 | 12.1327908 | 0.0015 | 0.015984 | F | F | F | F |
| H4c2 | 0.36572639 | 8.39355793 | 12.1285807 | 0.0015 | 0.016001 | F | F | F | F |
| Ncoa6 | -0.449724 | 7.02332362 | 12.1274635 | 0.0015 | 0.016001 | T | F | F | F |
| Tmem200a | 3.08979374 | 0.57201806 | 12.1239614 | 0.0015 | 0.016015 | T | F | T | T |
| Zfp827 | 1.88489317 | 1.77228131 | 12.1161316 | 0.00151 | 0.016055 | T | F | T | T |
| Miip | 0.94291424 | 3.74847529 | 12.0997324 | 0.00152 | 0.016149 | F | F | F | F |
| Rnf17 | -0.8549829 | 4.28034989 | 12.0984073 | 0.00152 | 0.016149 | T | F | F | F |
| Hba-a2 | -1.4259612 | 3.83051226 | 12.0821385 | 0.00153 | 0.016242 | F | F | F | F |
| Txndc9 | -0.5646124 | 6.01143192 | 12.0766886 | 0.00153 | 0.016268 | T | F | T | T |
| Plekhm2 | -0.6569589 | 5.29888339 | 12.0727117 | 0.00153 | 0.016285 | F | F | T | F |
| Tbc1d4 | 1.37871443 | 2.54886905 | 12.0662353 | 0.00154 | 0.016318 | F | F | T | F |
| Hps1 | 0.88983106 | 4.08068915 | 12.0567794 | 0.00154 | 0.01637 | F | F | T | T |
| Mfap3 | 0.50341759 | 6.25314775 | 12.0526266 | 0.00155 | 0.016388 | T | F | T | T |
| Fut10 | 0.80455488 | 4.39820408 | 12.0349626 | 0.00156 | 0.016492 | T | F | F | F |
| mt-Cytb | -0.334267 | 9.84607385 | 12.0321705 | 0.00156 | 0.016501 | F | F | F | F |
| Tmem246 | 1.89452523 | 1.69343198 | 12.0275481 | 0.00156 | 0.016519 | F | F | T | F |
| Map3k11 | -0.463078 | 6.83062446 | 12.0267789 | 0.00156 | 0.016519 | F | F | F | F |
| Slc25a24 | -0.6608264 | 5.25593873 | 12.0239167 | 0.00156 | 0.01653 | T | F | T | F |
| Mycbp2 | -0.3589468 | 8.29464687 | 12.0083378 | 0.00157 | 0.016621 | T | F | T | F |
| Cntnap2 | 1.84813037 | 1.76970016 | 11.9986223 | 0.00158 | 0.016675 | T | F | F | T |
| Tril | 2.08775719 | 1.45664125 | 11.9961575 | 0.00158 | 0.016683 | T | F | T | T |
| Zfp658 | 1.75418116 | 1.87013256 | 11.994125 | 0.00158 | 0.016688 | T | F | T | F |

| | | | | | | | | | |
|---|---|---|---|---|---|---|---|---|---|
| Lpcat1 | 0.48876815 | 6.41272819 | 11.9781568 | 0.00159 | 0.016781 | F | F | T | F |
| Vps9d1 | -8.4111483 | 0.71844917 | 11.9772478 | 0.00159 | 0.016781 | F | F | F | F |
| Stx2 | -0.9514496 | 3.94761909 | 11.9732462 | 0.00159 | 0.016798 | T | F | T | T |
| Cryab | 1.09458927 | 3.25117973 | 11.9716993 | 0.0016 | 0.0168 | F | F | F | F |
| Egln3 | 0.79924988 | 4.36936589 | 11.9565958 | 0.0016 | 0.01689 | T | F | T | F |
| Cnpy2 | 0.5620829 | 5.78001293 | 11.9360501 | 0.00162 | 0.017016 | F | F | F | T |
| Mapk10 | 5.43693505 | -0.3772983 | 11.9329719 | 0.00162 | 0.017028 | T | F | T | T |
| Cgnl1 | -0.3915671 | 7.69472945 | 11.9313817 | 0.00162 | 0.017031 | T | F | T | T |
| Clca3a1 | 7.58476279 | -0.7724501 | 11.9936388 | 0.00163 | 0.017106 | F | F | T | F |
| Hs6st1 | 0.5138984 | 6.15815487 | 11.9174824 | 0.00163 | 0.017106 | F | F | T | F |
| Kiz | 1.09153966 | 3.25562904 | 11.906501 | 0.00164 | 0.01717 | F | F | F | F |
| Myrfl | 4.1710919 | -0.063835 | 11.8968074 | 0.00164 | 0.017226 | F | F | F | T |
| Slitrk2 | 2.73303303 | 0.83856476 | 11.891869 | 0.00165 | 0.017247 | F | T | F | T |
| Plekho1 | 0.97663133 | 3.62217817 | 11.891246 | 0.00165 | 0.017247 | F | F | F | F |
| Shank1 | 1.5862667 | 2.17619904 | 11.879615 | 0.00165 | 0.017316 | F | F | T | F |
| Stau1 | -0.4970233 | 6.56092986 | 11.8763733 | 0.00166 | 0.017328 | F | F | T | F |
| Jakmip1 | 2.32862123 | 1.17039464 | 11.8741756 | 0.00166 | 0.017328 | T | F | T | F |
| Top3b | 0.71170235 | 4.85118986 | 11.8740723 | 0.00166 | 0.017328 | T | F | F | T |
| Eef2kmt | -0.7761695 | 4.81219573 | 11.8680147 | 0.00166 | 0.017361 | F | F | T | F |
| Stxbp4 | 0.64172329 | 5.10744761 | 11.8637313 | 0.00166 | 0.017381 | T | F | T | T |
| Desi2 | -0.521328 | 6.35416582 | 11.8402632 | 0.00168 | 0.017531 | F | F | T | F |
| Cfl2 | -0.7969274 | 4.57431476 | 11.8370316 | 0.00168 | 0.017545 | F | F | T | F |
| Atp6v0e2 | 1.79998411 | 1.81505315 | 11.8340047 | 0.00168 | 0.017557 | F | F | T | T |
| Xlr3b | -1.3663937 | 3.35994419 | 11.8306881 | 0.00168 | 0.017571 | F | F | F | F |
| Cyp26a1 | 0.55838241 | 5.77401924 | 11.8181916 | 0.00169 | 0.017648 | T | T | F | T |
| Aak1 | -0.4029224 | 7.43577935 | 11.8124185 | 0.0017 | 0.017679 | T | F | T | T |
| Ifrd1 | -0.6526003 | 5.29063969 | 11.8061409 | 0.0017 | 0.017713 | F | F | F | F |
| Vcl | -0.3683009 | 7.98358228 | 11.7893541 | 0.00171 | 0.01782 | T | F | T | T |
| Fgd6 | -0.5383704 | 6.14787319 | 11.7706589 | 0.00172 | 0.017941 | F | F | T | T |
| Unc119b | 0.54262068 | 5.90152911 | 11.7650906 | 0.00173 | 0.017972 | T | F | T | F |
| Esf1 | -0.5021404 | 6.47626628 | 11.76311 | 0.00173 | 0.017977 | F | F | F | F |
| Atp1b1 | -0.3710081 | 8.0373588 | 11.7594851 | 0.00173 | 0.01799 | F | F | F | F |
| Wnt6 | -0.7596638 | 5.28107473 | 11.7588037 | 0.00173 | 0.01799 | T | F | T | F |
| Cecr2 | 0.41237332 | 7.30656439 | 11.7555766 | 0.00173 | 0.017992 | T | F | T | T |
| Mapkapk2 | -0.4731405 | 6.67517955 | 11.7551491 | 0.00173 | 0.017992 | T | F | T | T |
| Stat1 | -0.6390975 | 5.62657529 | 11.7548074 | 0.00174 | 0.017992 | T | F | T | T |
| Stk39 | 0.85513798 | 4.01303939 | 11.7465085 | 0.00174 | 0.018041 | F | F | F | F |
| Olfm1 | 0.94424036 | 3.73012916 | 11.7442201 | 0.00174 | 0.018049 | T | F | T | F |
| Cdr2l | 0.71135486 | 4.9618491 | 11.73401 | 0.00175 | 0.018112 | F | F | T | F |
| Hspb8 | -0.6899688 | 5.07728763 | 11.7317989 | 0.00175 | 0.018119 | F | F | T | F |
| Tead3 | -0.6951071 | 4.97153021 | 11.725668 | 0.00175 | 0.018153 | T | F | T | F |
| Dip2c | 0.62594985 | 5.26631212 | 11.7229982 | 0.00176 | 0.018163 | T | F | T | F |

| | | | | | | | | | |
|---|---|---|---|---|---|---|---|---|---|
| Rcn2 | 0.59048214 | 5.48977784 | 11.7186655 | 0.00176 | 0.018185 | T | F | F | T |
| Eras | 1.22718394 | 2.93511069 | 11.7165812 | 0.00176 | 0.018191 | F | F | F | F |
| Pno1 | -0.4647592 | 6.75933411 | 11.7147182 | 0.00176 | 0.018195 | F | F | F | F |
| Rax | 4.94342924 | -0.2835741 | 11.7136123 | 0.00176 | 0.018195 | F | F | F | F |
| Cep78 | 0.73837222 | 4.48591073 | 11.6987474 | 0.00177 | 0.018292 | F | F | F | F |
| Pebp1 | 0.36744378 | 7.81018917 | 11.6854838 | 0.00178 | 0.018376 | F | F | T | F |
| Plpp1 | -0.7423163 | 4.71728281 | 11.6844866 | 0.00178 | 0.018376 | F | F | T | F |
| Mfsd10 | 0.65382408 | 5.49580194 | 11.6674617 | 0.00179 | 0.018489 | F | F | F | F |
| Gramd2 | -1.1589803 | 3.33410499 | 11.6645196 | 0.0018 | 0.018502 | F | F | T | F |
| Ccdc88c | 0.50029116 | 6.21787893 | 11.6579135 | 0.0018 | 0.018541 | T | F | F | T |
| Slc8a1 | 2.07322085 | 1.43366523 | 11.6490757 | 0.00181 | 0.018596 | T | F | T | T |
| Lnx2 | -0.5582799 | 5.92249973 | 11.6390648 | 0.00181 | 0.018651 | T | F | F | T |
| Sv2a | 1.5077811 | 2.26489864 | 11.6390626 | 0.00181 | 0.018651 | T | F | T | T |
| Atcay | 7.55177884 | -0.7989264 | 11.7000432 | 0.00182 | 0.018724 | T | F | T | T |
| Itpr3 | -0.3734584 | 7.88479299 | 11.6223777 | 0.00183 | 0.018755 | F | F | F | F |
| Cfap54 | 7.54980436 | -0.8005076 | 11.6826733 | 0.00184 | 0.018832 | F | F | F | F |
| Stag2 | 0.45600676 | 6.6163614 | 11.6059212 | 0.00184 | 0.018857 | T | T | F | T |
| Pcgf2 | 0.57073644 | 5.61221486 | 11.5863736 | 0.00185 | 0.018989 | F | F | T | F |
| Sla2 | -2.7343008 | 1.46062255 | 11.585535 | 0.00185 | 0.018989 | F | F | F | T |
| Ckap5 | 0.35643341 | 7.95925285 | 11.5838166 | 0.00185 | 0.018989 | T | F | F | T |
| Tcf20 | -0.3398573 | 8.78875503 | 11.5832241 | 0.00185 | 0.018989 | F | F | F | T |
| Cstf2t | 0.51940425 | 6.0166648 | 11.5793357 | 0.00186 | 0.019009 | T | F | F | F |
| Olfr1152 | -0.9938864 | 4.22962012 | 11.5754376 | 0.00186 | 0.019029 | F | F | F | F |
| Necap1 | -0.574684 | 5.77083816 | 11.5607863 | 0.00187 | 0.019129 | F | F | T | F |
| Uhmk1 | -0.4422061 | 6.96361525 | 11.5490824 | 0.00188 | 0.019208 | T | F | T | T |
| Id3 | 0.49754055 | 6.22289594 | 11.545938 | 0.00188 | 0.019222 | T | F | T | T |
| Hk1 | -0.709375 | 5.71785594 | 11.5417809 | 0.00189 | 0.019244 | F | F | F | T |
| Exoc3l4 | -0.662524 | 5.47044291 | 11.532947 | 0.00189 | 0.019302 | F | F | F | T |
| Susd4 | 1.07600373 | 3.24112886 | 11.5173603 | 0.0019 | 0.019411 | F | F | T | F |
| Sh3gl3 | 2.53187889 | 1.03238439 | 11.5130126 | 0.00191 | 0.019434 | T | F | F | F |
| Itga8 | 2.08499343 | 1.64384529 | 11.5048243 | 0.00191 | 0.019488 | T | F | T | T |
| Twist1 | 1.26541207 | 2.84619805 | 11.5023399 | 0.00191 | 0.019497 | T | F | F | T |
| Olfr46 | -0.5484578 | 5.99367946 | 11.4945143 | 0.00192 | 0.019543 | F | F | F | F |
| Scn1a | 7.53632989 | -0.8112872 | 11.5647507 | 0.00192 | 0.019543 | T | T | T | T |
| Pex5 | -0.4697904 | 6.65691984 | 11.4924106 | 0.00192 | 0.019546 | F | F | T | F |
| Slc45a1 | 2.52163858 | 0.93255036 | 11.4854965 | 0.00193 | 0.019589 | T | F | F | F |
| Efemp2 | 0.81373963 | 4.14560986 | 11.48432 | 0.00193 | 0.019589 | F | F | F | T |
| Cited1 | -0.3819415 | 7.76390405 | 11.4827994 | 0.00193 | 0.019592 | F | F | F | F |
| Calcoco1 | 0.71700002 | 4.62448339 | 11.4767428 | 0.00193 | 0.019629 | T | F | F | F |
| Cdc42ep1 | -0.5557933 | 5.89430623 | 11.4652769 | 0.00194 | 0.019708 | T | F | T | F |
| Kirrel | 0.53089351 | 5.8982078 | 11.4544784 | 0.00195 | 0.019782 | T | F | T | T |
| Uvrag | -0.6130605 | 5.46912743 | 11.4387723 | 0.00196 | 0.019895 | T | F | F | F |

| | | | | | | | | | |
|---|---|---|---|---|---|---|---|---|---|
| Zfp91 | -0.4162995 | 7.24332701 | 11.4321228 | 0.00197 | 0.019933 | T | T | T | T |
| Hapln1 | -0.8398984 | 4.25931212 | 11.4314764 | 0.00197 | 0.019933 | T | T | T | T |
| Ehhadh | -0.6839309 | 4.9824504 | 11.4301614 | 0.00197 | 0.019934 | T | F | T | F |
| Esam | -0.5900483 | 5.71279896 | 11.4245664 | 0.00197 | 0.019969 | T | F | F | F |
| Mcm2 | 0.34169702 | 8.68531154 | 11.4159194 | 0.00198 | 0.020027 | T | F | F | T |
| Crabp2 | 1.09449465 | 3.16854498 | 11.4099781 | 0.00198 | 0.020065 | F | F | F | F |
| Npl | -0.4714355 | 6.60399518 | 11.405429 | 0.00199 | 0.020091 | F | F | F | F |
| Adcy5 | 3.92582374 | 0.05001848 | 11.3955601 | 0.002 | 0.02016 | F | F | T | T |
| Chp1 | -0.5048071 | 6.40037948 | 11.3914601 | 0.002 | 0.020183 | T | F | F | F |
| Abcc8 | 1.86921294 | 1.71134649 | 11.3849423 | 0.002 | 0.020225 | F | F | F | F |
| Foxn3 | 0.44575627 | 6.68594547 | 11.3764863 | 0.00201 | 0.020283 | T | F | T | T |
| Gli1 | 1.13803302 | 3.03367458 | 11.3686492 | 0.00202 | 0.020336 | T | F | F | F |
| Npcd | -7.4954296 | -0.0273974 | 11.3653462 | 0.00202 | 0.020353 | T | F | T | T |
| Cebpb | -0.6864618 | 5.50991231 | 11.361218 | 0.00202 | 0.020376 | F | F | T | T |
| Zscan18 | 1.26513881 | 2.69413263 | 11.3524741 | 0.00203 | 0.020437 | F | F | T | F |
| Zscan2 | 1.35853635 | 2.46936768 | 11.3489999 | 0.00203 | 0.020455 | F | F | F | T |
| App | 0.46144177 | 6.57404484 | 11.3427875 | 0.00204 | 0.020496 | T | F | T | T |
| Rsu1 | -0.6807157 | 4.98235844 | 11.3350682 | 0.00204 | 0.020548 | T | F | F | F |
| Zfp87 | 1.00052362 | 3.46102816 | 11.3291242 | 0.00205 | 0.020587 | T | F | T | F |
| Drp2 | 1.40923146 | 2.42309754 | 11.3182478 | 0.00206 | 0.020666 | T | F | T | F |
| Rubcn | -0.5954997 | 5.58299207 | 11.3063296 | 0.00207 | 0.020753 | T | F | T | F |
| Pthlh | -1.526123 | 2.57133477 | 11.2957235 | 0.00208 | 0.020828 | F | F | F | F |
| Tab3 | -0.5364152 | 6.01142405 | 11.2947535 | 0.00208 | 0.020828 | F | F | T | F |
| Ccdc69 | -2.8867737 | 1.14913188 | 11.2870816 | 0.00208 | 0.020874 | F | F | F | F |
| Fam171a2 | 1.23261944 | 2.7957763 | 11.2864739 | 0.00208 | 0.020874 | F | F | T | F |
| Brpf1 | -0.4237656 | 7.04257797 | 11.2855922 | 0.00208 | 0.020874 | F | F | T | F |
| Cds2 | 0.42027047 | 6.89841381 | 11.2845313 | 0.00209 | 0.020874 | T | F | F | T |
| G2e3 | 0.58319018 | 5.70192844 | 11.2733936 | 0.00209 | 0.020956 | T | F | T | T |
| Usp1 | -0.3877714 | 7.55323787 | 11.2624275 | 0.0021 | 0.021037 | T | F | F | T |
| Pimreg | 0.7044449 | 4.63016135 | 11.2575313 | 0.00211 | 0.021068 | F | F | T | F |
| Tef | 0.82015527 | 4.15443643 | 11.2533374 | 0.00211 | 0.021093 | T | F | T | F |
| Timp2 | 0.80195009 | 4.19423257 | 11.2443957 | 0.00212 | 0.021149 | T | F | T | F |
| Gdap2 | -0.6847542 | 4.95332708 | 11.2442902 | 0.00212 | 0.021149 | T | F | T | T |
| Tmem121b | 3.71672519 | 0.07367674 | 11.243048 | 0.00212 | 0.021149 | T | F | T | F |
| Mak16 | -0.4241766 | 6.99147442 | 11.23471 | 0.00213 | 0.021209 | T | T | T | T |
| Hunk | 0.46588484 | 6.44785572 | 11.2261016 | 0.00213 | 0.021271 | F | F | T | F |
| Gdpd2 | 7.50474135 | -0.8364809 | 11.2924764 | 0.00213 | 0.021273 | F | F | F | F |
| Eid1 | 0.72751989 | 4.61758321 | 11.2213331 | 0.00214 | 0.02129 | T | F | F | F |
| Pdpn | 0.41667314 | 6.91780488 | 11.2204209 | 0.00214 | 0.02129 | T | F | T | F |
| Prkra | 1.01793301 | 3.45374396 | 11.2191483 | 0.00214 | 0.021291 | T | F | F | F |
| Rassf1 | -0.7271721 | 4.67358386 | 11.206995 | 0.00215 | 0.021383 | T | F | T | F |
| 4931428F04 | 0.83224618 | 4.04461472 | 11.1942424 | 0.00216 | 0.021474 | F | F | F | F |

| | | | | | | | | | |
|---|---|---|---|---|---|---|---|---|---|
| Jmjd1c | 0.43450767 | 7.02765878 | 11.1939534 | 0.00216 | 0.021474 | T | T | T | T |
| Zfp647 | 1.30958938 | 2.69023405 | 11.1915059 | 0.00216 | 0.021483 | T | F | T | T |
| Aoc3 | -2.3050464 | 1.57644816 | 11.1905174 | 0.00216 | 0.021483 | T | F | T | T |
| Plac8 | -0.5420084 | 6.29625529 | 11.1776441 | 0.00217 | 0.021583 | F | F | F | F |
| Ttll12 | -0.3848116 | 7.71659929 | 11.174139 | 0.00218 | 0.021597 | F | F | F | F |
| Slc30a4 | -0.6541262 | 5.1737101 | 11.17204 | 0.00218 | 0.021597 | T | F | F | T |
| Dnm1l | -0.4281076 | 7.07106209 | 11.1714496 | 0.00218 | 0.021597 | T | F | F | T |
| Cdx1 | 1.86305874 | 1.85610025 | 11.1713788 | 0.00218 | 0.021597 | F | F | T | F |
| Brpf3 | -0.4505089 | 6.7640084 | 11.1655177 | 0.00219 | 0.021627 | T | F | T | F |
| Rilpl1 | 1.01190683 | 3.4201211 | 11.1654576 | 0.00219 | 0.021627 | F | F | F | F |
| 1700003E16 | 5.50353624 | -0.4961068 | 11.1325311 | 0.00221 | 0.021892 | F | F | F | F |
| Cpox | -0.5168898 | 6.32054374 | 11.1323278 | 0.00221 | 0.021892 | T | T | F | T |
| Kif21a | 0.48978457 | 6.1986462 | 11.1258502 | 0.00222 | 0.021939 | F | F | F | T |
| AW551984 | 3.22937623 | 0.34527991 | 11.1209294 | 0.00222 | 0.021971 | T | F | T | T |
| Myom3 | -2.2745575 | 1.60635499 | 11.1135158 | 0.00223 | 0.022026 | T | F | F | F |
| Jcad | -0.533466 | 6.17123726 | 11.1051063 | 0.00224 | 0.022089 | T | F | T | T |
| Tmed4 | 0.63464737 | 5.00114574 | 11.0962697 | 0.00225 | 0.022156 | T | F | T | T |
| Fam3c | -0.527535 | 6.13949269 | 11.0912961 | 0.00225 | 0.02219 | F | F | T | T |
| Trmt9b | 1.96710974 | 1.5090813 | 11.081848 | 0.00226 | 0.022263 | F | F | T | F |
| Ptprd | 1.25222682 | 2.68825939 | 11.073059 | 0.00227 | 0.02233 | T | F | T | T |
| Psmd7 | -0.386025 | 7.51828084 | 11.0479571 | 0.00229 | 0.022543 | T | F | F | T |
| Cyyr1 | 1.7428935 | 1.78324857 | 11.0439615 | 0.00229 | 0.022569 | T | F | T | T |
| B3galt1 | 5.51148692 | -0.5353437 | 11.0418211 | 0.00229 | 0.022578 | T | F | T | T |
| Col5a2 | 1.0861874 | 3.9427284 | 11.0383991 | 0.0023 | 0.022598 | F | F | F | T |
| Yeats2 | 0.4379682 | 6.74308585 | 11.0278991 | 0.00231 | 0.022676 | T | F | T | F |
| Gm9780 | -7.7100851 | 0.07819005 | 11.0274374 | 0.00231 | 0.022676 | F | F | F | F |
| Wasf3 | 0.76042899 | 4.34282496 | 11.0205681 | 0.00231 | 0.022727 | T | F | F | T |
| Zbtb11 | -0.525226 | 6.05160155 | 11.0194911 | 0.00232 | 0.022727 | T | F | F | F |
| Rac3 | -1.0584991 | 3.52195419 | 11.0125581 | 0.00232 | 0.022779 | F | F | F | F |
| Osbp | -0.4363168 | 6.83760093 | 11.0007526 | 0.00233 | 0.022876 | T | F | T | F |
| Wdr36 | -0.4043046 | 7.25782746 | 10.9944487 | 0.00234 | 0.022923 | F | F | T | F |
| Mpnd | 0.63326776 | 5.00752103 | 10.9869697 | 0.00235 | 0.02298 | F | F | F | F |
| Ddah1 | -0.5134247 | 6.1486753 | 10.9857998 | 0.00235 | 0.022981 | F | F | F | T |
| Zkscan3 | 0.63627627 | 4.96151549 | 10.9699359 | 0.00236 | 0.023115 | T | F | F | F |
| Zgrf1 | 0.71176408 | 4.50425185 | 10.9577482 | 0.00237 | 0.023217 | F | T | F | T |
| Drc1 | 3.82303639 | 0.22383356 | 10.9531536 | 0.00238 | 0.023249 | T | F | F | F |
| Taf1c | -0.6907521 | 4.83292861 | 10.9502733 | 0.00238 | 0.023265 | T | F | F | F |
| Csgalnact1 | -1.0917536 | 3.67958248 | 10.9484461 | 0.00238 | 0.023272 | T | T | T | T |
| Mogs | 0.39026095 | 7.32316465 | 10.9453938 | 0.00238 | 0.023289 | T | T | F | T |
| 2610008E11 | 0.89915035 | 3.79898745 | 10.9422748 | 0.00239 | 0.023308 | F | F | T | F |
| Grk3 | 1.95774013 | 2.2005051 | 10.9254735 | 0.0024 | 0.023453 | T | F | T | F |
| Dtnb | 0.63193312 | 4.98555399 | 10.9230186 | 0.00241 | 0.023454 | T | F | T | T |

| | | | | | | | | | |
|---|---|---|---|---|---|---|---|---|---|
| Pou2f1 | 0.45682523 | 6.69232225 | 10.9221314 | 0.00241 | 0.023454 | F | T | T | T |
| Sox18 | -3.2334585 | 0.96814962 | 10.9212092 | 0.00241 | 0.023454 | F | F | F | F |
| Wwtr1 | -0.5319739 | 5.9750153 | 10.9208835 | 0.00241 | 0.023454 | T | F | T | T |
| Selenot | -0.3895678 | 7.40613671 | 10.9149614 | 0.00241 | 0.023499 | T | F | T | T |
| Zfp202 | -2.3828922 | 1.53819401 | 10.9025419 | 0.00243 | 0.023605 | T | T | T | T |
| Epcam | 0.39649495 | 7.06724065 | 10.89712 | 0.00243 | 0.023645 | T | F | T | T |
| Arhgap1 | -0.4096295 | 7.1444286 | 10.8844033 | 0.00244 | 0.023754 | T | F | T | T |
| Oxct2b | -1.4709672 | 2.52439909 | 10.8710536 | 0.00246 | 0.02387 | F | F | F | F |
| Fndc4 | 0.81341554 | 4.05794403 | 10.8597614 | 0.00247 | 0.023967 | F | F | T | F |
| Dlg3 | 1.29488925 | 2.59729321 | 10.8585999 | 0.00247 | 0.023968 | T | F | T | F |
| Senp5 | -0.4644456 | 6.5817438 | 10.8550741 | 0.00247 | 0.023991 | T | F | T | T |
| Gm9833 | 7.45918194 | -0.8726242 | 10.9099486 | 0.00248 | 0.024064 | F | F | F | F |
| Jund | -0.5004847 | 6.2557797 | 10.8447459 | 0.00248 | 0.024068 | F | F | F | T |
| Abi2 | 0.42395559 | 6.77891335 | 10.8377524 | 0.00249 | 0.024124 | T | F | T | T |
| Golm1 | 0.41484203 | 6.86699396 | 10.8303571 | 0.0025 | 0.024185 | T | F | T | T |
| Impa2 | -0.6893921 | 5.21184378 | 10.8280849 | 0.0025 | 0.024194 | T | F | T | T |
| H4c3 | 0.32666225 | 8.5057397 | 10.82721 | 0.0025 | 0.024194 | F | F | F | F |
| Gm28042 | -8.19368 | 0.51094171 | 10.8160851 | 0.00251 | 0.02429 | F | F | F | F |
| Znrf1 | 0.61692333 | 5.09058798 | 10.8134295 | 0.00251 | 0.024305 | T | F | T | T |
| Gmfb | -0.4093064 | 7.1847686 | 10.8093985 | 0.00252 | 0.024334 | T | F | T | F |
| Cyp39a1 | 2.06351246 | 1.34287613 | 10.8002054 | 0.00253 | 0.024412 | T | F | F | F |
| Clmn | 0.76211008 | 4.23153166 | 10.7977828 | 0.00253 | 0.024425 | T | F | T | T |
| Mxi1 | 1.01838349 | 3.36072665 | 10.795373 | 0.00253 | 0.024438 | T | F | T | F |
| Rbpms | 0.47562913 | 6.35035537 | 10.7837606 | 0.00254 | 0.02454 | F | F | T | F |
| Tollip | -0.4648092 | 6.55375205 | 10.7819939 | 0.00254 | 0.024547 | F | F | T | T |
| Spink1 | 0.46695881 | 6.39586165 | 10.7776138 | 0.00255 | 0.024579 | F | F | F | T |
| Rnf24 | 1.06442552 | 3.16646781 | 10.7740786 | 0.00255 | 0.024602 | T | T | F | T |
| Nxn | 0.59151391 | 5.28118612 | 10.7668335 | 0.00256 | 0.024663 | F | F | T | F |
| Triml2 | -1.2626871 | 3.13995358 | 10.7631063 | 0.00256 | 0.024678 | F | F | F | F |
| Phactr1 | -0.6761305 | 5.25734407 | 10.7630135 | 0.00256 | 0.024678 | T | F | T | T |
| Col3a1 | -1.5080156 | 2.54581281 | 10.7617429 | 0.00257 | 0.024678 | T | F | T | F |
| Rabepk | -0.8794527 | 4.2148803 | 10.7608549 | 0.00257 | 0.024678 | F | F | T | T |
| Vwa2 | 2.20891731 | 1.37031359 | 10.7597735 | 0.00257 | 0.024678 | F | F | T | F |
| Tmed1 | 0.62058659 | 5.27208188 | 10.7570123 | 0.00257 | 0.024694 | F | F | T | F |
| Pm20d1 | 1.73010596 | 1.80789171 | 10.7550715 | 0.00257 | 0.024702 | T | F | T | T |
| Perp | -0.5630288 | 5.74683862 | 10.7514315 | 0.00258 | 0.024727 | T | F | T | F |
| Rspo3 | 0.98200674 | 3.89444435 | 10.7439389 | 0.00258 | 0.024787 | T | F | T | F |
| Pot1b | 0.82829682 | 3.97151425 | 10.743182 | 0.00258 | 0.024787 | T | F | T | T |
| Eya1 | 2.39608966 | 1.0598683 | 10.7344675 | 0.00259 | 0.024862 | T | F | T | T |
| Csrnp3 | 1.89053068 | 1.56307098 | 10.7185161 | 0.00261 | 0.025011 | T | T | T | T |
| Dcaf12 | -0.4203338 | 6.95134331 | 10.7112382 | 0.00262 | 0.025072 | F | F | T | F |
| Hectd3 | -0.6159657 | 5.32663601 | 10.7088737 | 0.00262 | 0.025085 | T | F | F | T |

| | | | | | | | | | |
|---|---|---|---|---|---|---|---|---|---|
| Ncor2 | 0.34516614 | 7.91312367 | 10.6861531 | 0.00264 | 0.025303 | F | F | F | F |
| Arl6 | 1.32553726 | 2.54543918 | 10.6781081 | 0.00265 | 0.025374 | T | F | T | F |
| Syvn1 | 0.4659567 | 6.40982964 | 10.6754985 | 0.00266 | 0.025389 | T | F | F | F |
| Ywhab | -0.3347909 | 8.431511 | 10.6727219 | 0.00266 | 0.025406 | T | F | T | F |
| Iars | -0.3337376 | 8.61943297 | 10.6624724 | 0.00267 | 0.025491 | F | F | F | T |
| Exoc5 | -0.5280977 | 6.14013777 | 10.6621559 | 0.00267 | 0.025491 | T | F | T | T |
| Fer | -0.6583689 | 5.00541981 | 10.6553541 | 0.00268 | 0.025538 | F | F | F | T |
| Cuedc1 | -0.9888521 | 3.67620124 | 10.6549558 | 0.00268 | 0.025538 | F | F | T | T |
| Slc35f2 | 0.73033869 | 4.59322902 | 10.6533025 | 0.00268 | 0.025538 | F | F | T | F |
| Eya4 | 3.69733283 | 0.10355073 | 10.6532093 | 0.00268 | 0.025538 | F | F | T | F |
| Cercam | 1.3862596 | 2.5080532 | 10.6471172 | 0.00269 | 0.025589 | F | F | F | F |
| Clec1a | 1.95886281 | 1.76562412 | 10.6456759 | 0.00269 | 0.025593 | T | F | T | T |
| Amotl1 | 0.47638946 | 6.31916687 | 10.634229 | 0.0027 | 0.025699 | T | F | T | T |
| Etv6 | 0.62482784 | 4.98340963 | 10.6298142 | 0.0027 | 0.025734 | T | F | T | T |
| Hdx | 1.04094044 | 3.33459862 | 10.619678 | 0.00272 | 0.025818 | T | F | T | T |
| Nfe2 | -2.4605537 | 1.49687815 | 10.6194235 | 0.00272 | 0.025818 | F | F | F | T |
| Cnbp | -0.3172909 | 9.00024029 | 10.6155571 | 0.00272 | 0.025847 | F | F | F | T |
| Rpl10l | -1.3239806 | 2.97839125 | 10.608011 | 0.00273 | 0.025914 | F | F | F | F |
| Gucy1a1 | -1.4362483 | 2.58392791 | 10.5985719 | 0.00274 | 0.025993 | T | T | F | F |
| Amhr2 | 2.34033895 | 1.02643284 | 10.598324 | 0.00274 | 0.025993 | F | F | F | F |
| Strada | 0.81267893 | 4.20973318 | 10.5919768 | 0.00275 | 0.026047 | F | F | T | F |
| Prr12 | 0.34121513 | 8.04126252 | 10.5891863 | 0.00275 | 0.026065 | F | F | F | F |
| Lpo | 7.42694678 | -0.8980573 | 10.6464268 | 0.00275 | 0.026081 | F | F | F | F |
| Shh | 7.42694678 | -0.8980573 | 10.6464268 | 0.00275 | 0.026081 | F | F | F | F |
| Eif1b | 0.69773111 | 4.54746747 | 10.5600433 | 0.00278 | 0.026338 | F | F | T | F |
| Rbms3 | 2.33031269 | 1.04284002 | 10.5573582 | 0.00278 | 0.026355 | T | T | T | T |
| Morf4l2 | -0.292744 | 9.53557277 | 10.5558538 | 0.00279 | 0.026359 | T | T | T | T |
| Efemp1 | 2.83529917 | 0.59855265 | 10.5494941 | 0.00279 | 0.026415 | F | F | F | T |
| Tspan32 | -7.7525474 | 0.10881066 | 10.5349799 | 0.00281 | 0.026558 | F | F | F | T |
| Zcchc14 | 0.48098125 | 6.26108587 | 10.5246577 | 0.00282 | 0.026657 | F | F | F | F |
| Fbxl7 | 2.15256449 | 1.33955771 | 10.5097299 | 0.00284 | 0.026806 | T | F | T | F |
| Lpar5 | -1.4451464 | 2.52793997 | 10.5080117 | 0.00284 | 0.026813 | F | F | F | T |
| Ddx19b | 0.57104957 | 5.38729744 | 10.5063853 | 0.00284 | 0.026819 | T | F | T | F |
| Ermn | 5.32343484 | -0.5206725 | 10.4895814 | 0.00286 | 0.026989 | T | F | T | T |
| Ifngr2 | -0.7948251 | 4.41853049 | 10.4827125 | 0.00287 | 0.027053 | T | F | T | T |
| Ulk4 | 1.25198078 | 3.11594407 | 10.4755468 | 0.00288 | 0.027114 | T | F | T | F |
| Mapk11 | 2.05695151 | 1.32049685 | 10.4749092 | 0.00288 | 0.027114 | F | F | F | F |
| Ctif | 0.68642637 | 4.57394153 | 10.458025 | 0.0029 | 0.027288 | T | F | T | T |
| Sfpq | 0.2907924 | 9.56938948 | 10.436153 | 0.00292 | 0.027517 | T | F | F | T |
| Cdc42ep3 | -1.1658195 | 3.12503292 | 10.4338289 | 0.00293 | 0.027531 | F | F | F | T |
| Ass1 | -0.4156512 | 7.19364672 | 10.4084574 | 0.00296 | 0.027803 | F | F | F | F |
| Trim32 | 0.57079081 | 5.35620486 | 10.3904663 | 0.00298 | 0.027987 | T | F | F | F |

| | | | | | | | | | |
|---|---|---|---|---|---|---|---|---|---|
| H3c10 | 0.35431005 | 7.65064522 | 10.3899706 | 0.00298 | 0.027987 | F | F | F | T |
| Mfn2 | -0.4428002 | 6.68487124 | 10.3782488 | 0.00299 | 0.028108 | F | F | T | T |
| Sec24b | -0.4098878 | 6.96352838 | 10.3713919 | 0.003 | 0.028174 | F | F | F | T |
| Pla2g4f | -7.4109476 | -0.1355532 | 10.3681854 | 0.00301 | 0.028194 | F | F | F | F |
| Clta | -0.3412562 | 8.10909248 | 10.3674543 | 0.00301 | 0.028194 | F | F | F | F |
| Sephs1 | 0.42452624 | 6.68448542 | 10.3523041 | 0.00302 | 0.028355 | F | F | T | F |
| Ptprs | -0.3594961 | 7.74420257 | 10.3487765 | 0.00303 | 0.028383 | T | F | T | F |
| Tnfaip2 | -0.7498876 | 4.65691601 | 10.3456554 | 0.00303 | 0.02839 | T | F | F | T |
| Thg1l | -0.64049 | 5.04589465 | 10.3455685 | 0.00303 | 0.02839 | F | F | T | T |
| Esrra | -0.5699467 | 5.56445883 | 10.3449843 | 0.00303 | 0.02839 | F | F | T | F |
| Gm49345 | 7.3961744 | -0.9222266 | 10.4002835 | 0.00304 | 0.028413 | F | F | F | F |
| Rrp7a | -0.3800036 | 7.3945108 | 10.33977 | 0.00304 | 0.028426 | F | F | F | T |
| Prkcb | 2.05711748 | 1.29170646 | 10.3367637 | 0.00304 | 0.028448 | T | F | T | T |
| Per2 | 1.05768036 | 3.13846116 | 10.3222946 | 0.00306 | 0.028603 | F | F | T | F |
| Mlec | 0.321678 | 8.58314536 | 10.3164525 | 0.00307 | 0.028658 | T | F | T | T |
| Hdac11 | 1.78869923 | 1.66276081 | 10.3154012 | 0.00307 | 0.028658 | F | F | T | F |
| Ighmbp2 | -0.6934133 | 4.70934945 | 10.3100984 | 0.00308 | 0.028707 | F | F | T | T |
| Adamts19 | 2.34123883 | 1.04134127 | 10.2970447 | 0.00309 | 0.028847 | T | F | F | F |
| Snx3 | -0.3648268 | 7.61620463 | 10.2882976 | 0.0031 | 0.028937 | T | F | F | F |
| Akap13 | 0.38384473 | 7.08467555 | 10.2778233 | 0.00312 | 0.029043 | T | F | T | F |
| Smarcc1 | 0.31345879 | 8.82682759 | 10.276831 | 0.00312 | 0.029043 | T | F | T | F |
| Arhgap6 | 1.02504999 | 3.30221831 | 10.2760864 | 0.00312 | 0.029043 | T | F | T | F |
| Ptgds | 5.40408047 | -0.5758805 | 10.2742646 | 0.00312 | 0.029045 | F | F | F | F |
| Vdac1 | -0.3242555 | 8.73386499 | 10.2738272 | 0.00312 | 0.029045 | F | F | T | F |
| Dlec1 | 3.52967548 | 0.36493921 | 10.2725893 | 0.00312 | 0.029047 | F | F | F | T |
| Fgfr3 | 0.57485376 | 5.37703103 | 10.2598525 | 0.00314 | 0.029185 | F | F | T | F |
| Snx8 | -0.4703624 | 6.3758065 | 10.2559129 | 0.00315 | 0.029219 | F | F | T | F |
| Cenpb | 0.47617703 | 6.23706299 | 10.2529509 | 0.00315 | 0.029242 | F | F | F | T |
| Ubn1 | -0.4418794 | 6.64458006 | 10.2441588 | 0.00316 | 0.029334 | F | F | F | F |
| Arsi | 1.14716967 | 3.12845924 | 10.2398771 | 0.00317 | 0.029372 | F | F | F | F |
| Bdp1 | -0.3829118 | 7.29421938 | 10.2348918 | 0.00317 | 0.029419 | T | F | T | T |
| Sacm1l | -0.5283819 | 5.94154952 | 10.2302955 | 0.00318 | 0.029456 | T | F | F | F |
| Garem2 | 2.41365708 | 0.88940562 | 10.229678 | 0.00318 | 0.029456 | F | F | F | F |
| Gm49496 | 2.64322332 | 1.60961547 | 10.2246072 | 0.00319 | 0.029505 | F | F | F | F |
| Gm3298 | 7.37989877 | -0.934966 | 10.2722108 | 0.0032 | 0.029604 | F | F | F | F |
| Itga4 | -0.8352098 | 4.28611064 | 10.2141992 | 0.0032 | 0.029604 | T | F | T | T |
| Slc25a37 | -1.0636219 | 3.82086144 | 10.2074713 | 0.00321 | 0.029673 | F | F | T | F |
| Zfp966 | 1.82648231 | 2.64332121 | 10.2056859 | 0.00321 | 0.029682 | F | F | T | F |
| Rabif | -0.6129883 | 5.20966565 | 10.1990013 | 0.00322 | 0.02975 | F | F | F | F |
| Mertk | 0.92201686 | 3.55247704 | 10.1737118 | 0.00325 | 0.030037 | F | F | F | T |
| Ptprg | 0.75748761 | 4.88279445 | 10.1732458 | 0.00325 | 0.030037 | T | T | T | T |
| Rsl1d1 | -0.3330821 | 8.21177069 | 10.171326 | 0.00326 | 0.030048 | F | F | F | T |

| | | | | | | | | | |
|---|---|---|---|---|---|---|---|---|---|
| Paqr4 | -0.7456303 | 4.4271659 | 10.1696783 | 0.00326 | 0.030055 | F | F | T | F |
| Dnah11 | 1.18085075 | 2.73757265 | 10.1615128 | 0.00327 | 0.030143 | F | F | F | F |
| Shisa7 | 5.40620231 | -0.6198284 | 10.1341645 | 0.0033 | 0.030467 | F | F | T | T |
| Epb41 | -0.3767392 | 7.40340133 | 10.1301082 | 0.00331 | 0.030505 | T | F | T | F |
| Ticam2 | 2.6027919 | 0.71276871 | 10.1157298 | 0.00333 | 0.03067 | T | F | F | T |
| Mtss2 | 0.52012033 | 5.71476156 | 10.1147659 | 0.00333 | 0.03067 | T | F | F | F |
| Vmn2r53 | -0.5357064 | 6.13642909 | 10.1038746 | 0.00335 | 0.030793 | F | F | F | F |
| Ldlrap1 | -0.5266239 | 5.87190825 | 10.0989757 | 0.00335 | 0.030833 | F | F | T | T |
| D630003M2 | 4.01935088 | -0.1834278 | 10.0986648 | 0.00335 | 0.030833 | F | F | T | F |
| Gprasp2 | 1.61696766 | 1.87748058 | 10.0936521 | 0.00336 | 0.030883 | F | F | F | T |
| Med15 | -0.5224617 | 5.92536318 | 10.084827 | 0.00337 | 0.030975 | T | F | T | F |
| Fmn2 | 1.37689558 | 2.3073553 | 10.0837873 | 0.00337 | 0.030975 | F | F | F | T |
| Zdhhc23 | -0.8242669 | 4.24411933 | 10.0832446 | 0.00337 | 0.030975 | F | F | T | F |
| Pkd2 | 0.56951612 | 5.30125131 | 10.0778291 | 0.00338 | 0.03103 | T | T | T | T |
| Zmynd10 | 7.36070965 | -0.9499465 | 10.1230698 | 0.0034 | 0.031148 | F | F | F | F |
| Gal3st4 | 7.36023175 | -0.9503191 | 10.1193811 | 0.0034 | 0.031181 | F | F | F | F |
| Dnttip2 | -0.3912759 | 7.13607729 | 10.0604688 | 0.00341 | 0.031211 | F | F | F | F |
| Pla2g2d | -2.2177917 | 1.52015819 | 10.0548722 | 0.00341 | 0.031269 | F | F | T | F |
| Per3 | 1.0678628 | 3.04610175 | 10.0534677 | 0.00342 | 0.031274 | T | F | F | F |
| Fam162a | 0.64072984 | 4.78217097 | 10.0467078 | 0.00342 | 0.031347 | F | F | F | F |
| Dpyd | 2.7306577 | 0.85271697 | 10.0416507 | 0.00343 | 0.031399 | T | T | F | T |
| Nbeal1 | 0.58737843 | 5.48823371 | 10.0361661 | 0.00344 | 0.031447 | T | F | T | T |
| Gm14124 | -1.2864018 | 2.77597872 | 10.0358464 | 0.00344 | 0.031447 | F | F | F | F |
| Eif2b1 | -0.4280325 | 6.72308752 | 10.0295062 | 0.00345 | 0.031515 | F | F | F | F |
| D16Ertd472 | 0.91086846 | 3.7068678 | 10.0260903 | 0.00345 | 0.031546 | T | F | T | T |
| Lig1 | 0.36636409 | 7.44924644 | 10.02155 | 0.00346 | 0.031591 | F | F | F | F |
| Polr1d | -0.3921891 | 7.08939146 | 10.0205058 | 0.00346 | 0.031591 | F | F | F | F |
| Metrn | 1.07684267 | 3.21106613 | 10.0051813 | 0.00348 | 0.031776 | F | F | F | F |
| Dzip1 | 1.26996153 | 2.52987399 | 9.98998761 | 0.00351 | 0.031961 | F | F | T | T |
| Klk8 | 1.76265235 | 1.63846903 | 9.98765378 | 0.00351 | 0.031978 | F | F | F | F |
| Rnf145 | 0.39408477 | 6.99017281 | 9.97521437 | 0.00353 | 0.032128 | T | F | T | T |
| Prss16 | -1.3399072 | 2.66003403 | 9.97372133 | 0.00353 | 0.032134 | F | F | T | F |
| Sh3bp5 | -0.7549517 | 4.60723352 | 9.96762081 | 0.00354 | 0.032201 | F | F | T | F |
| Scamp1 | -0.577032 | 5.44875699 | 9.96473109 | 0.00354 | 0.032225 | T | T | T | T |
| Rab12 | -0.6351428 | 4.99379848 | 9.96053779 | 0.00355 | 0.032267 | F | F | T | T |
| Gnai1 | 0.97432352 | 3.37195322 | 9.95386784 | 0.00356 | 0.032342 | T | F | T | T |
| Vps26a | -0.4353327 | 6.63587778 | 9.9482226 | 0.00357 | 0.032399 | T | F | F | T |
| Slc43a1 | 0.83823777 | 3.82614958 | 9.94756041 | 0.00357 | 0.032399 | T | F | F | T |
| Zfp354c | 0.89266808 | 3.64337719 | 9.93997593 | 0.00358 | 0.032486 | T | F | T | T |
| Ttc39b | 1.13609806 | 2.82848882 | 9.92649097 | 0.0036 | 0.032643 | F | F | T | F |
| Tmem65 | -0.6475782 | 5.05389829 | 9.92618289 | 0.0036 | 0.032643 | T | F | T | T |
| Metap1 | -0.387868 | 7.10594691 | 9.92440553 | 0.0036 | 0.032653 | T | F | T | F |

| | | | | | | | | | |
|---|---|---|---|---|---|---|---|---|---|
| Tpst2 | -0.5591392 | 5.6206761 | 9.92311582 | 0.0036 | 0.032656 | F | F | F | F |
| Zmym4 | 0.40744582 | 6.76541837 | 9.91577002 | 0.00361 | 0.032741 | F | F | T | F |
| Atrnl1 | 0.53130641 | 5.59417842 | 9.91307936 | 0.00362 | 0.032764 | F | F | T | F |
| Eif3j1 | -0.6086195 | 5.17657672 | 9.90956107 | 0.00362 | 0.032797 | T | F | F | T |
| Crct1 | 3.62969819 | 0.05146858 | 9.9068894 | 0.00363 | 0.03282 | T | F | T | F |
| Tbc1d32 | 1.5764459 | 2.08193653 | 9.9056314 | 0.00363 | 0.032823 | T | F | F | T |
| Rbm19 | -0.3645735 | 7.52007135 | 9.88322768 | 0.00366 | 0.033112 | F | F | T | T |
| Rcor1 | -0.3992582 | 6.96823722 | 9.86936725 | 0.00368 | 0.033287 | T | F | T | T |
| Il1r2 | 2.00974994 | 1.29275951 | 9.86675708 | 0.00369 | 0.033309 | F | F | F | F |
| Gprc5c | 0.47951748 | 6.69371183 | 9.86252329 | 0.00369 | 0.033349 | T | F | F | F |
| Evl | 0.52051567 | 5.64522976 | 9.86179906 | 0.00369 | 0.033349 | F | F | T | F |
| Tekt4 | 7.33279644 | -0.9716613 | 9.90967586 | 0.0037 | 0.033412 | F | F | F | F |
| Blzf1 | -0.6022861 | 5.21528448 | 9.84755336 | 0.00372 | 0.033517 | T | F | T | T |
| Lysmd2 | -1.3167326 | 2.68486824 | 9.83997265 | 0.00373 | 0.033608 | T | F | F | F |
| Clasp1 | 0.37413329 | 7.1213244 | 9.83563719 | 0.00373 | 0.033654 | T | F | T | T |
| Gm11992 | -2.6700261 | 1.11464872 | 9.8301549 | 0.00374 | 0.033716 | F | F | F | F |
| Bag3 | -0.4573569 | 6.4299558 | 9.82886433 | 0.00374 | 0.03372 | F | F | F | F |
| Elovl4 | 1.85896409 | 1.54680107 | 9.79731727 | 0.00379 | 0.034146 | T | F | F | T |
| Rab5c | -0.3805577 | 7.18767873 | 9.79335859 | 0.0038 | 0.034187 | F | F | T | T |
| Zfp12 | 0.63263196 | 4.72152327 | 9.79037988 | 0.0038 | 0.034215 | T | F | T | T |
| Idh3b | -0.4079646 | 6.86304131 | 9.7886245 | 0.00381 | 0.034226 | T | F | F | F |
| E2f2 | -0.7562542 | 4.30489342 | 9.78712273 | 0.00381 | 0.034233 | F | F | T | F |
| Net1 | -0.3745272 | 7.32137683 | 9.77888331 | 0.00382 | 0.034335 | T | F | T | F |
| Bnip3 | 0.54251609 | 5.47500027 | 9.77327782 | 0.00383 | 0.0344 | T | F | F | T |
| Cpxm2 | 3.40710554 | 0.13827126 | 9.7663464 | 0.00384 | 0.034484 | T | F | T | T |
| Ltv1 | -0.4577559 | 6.38554346 | 9.75293323 | 0.00386 | 0.034661 | T | F | F | F |
| Stat4 | 5.33965634 | -0.6269899 | 9.74791024 | 0.00387 | 0.034718 | F | F | T | F |
| Pde4d | 1.11173669 | 2.87374522 | 9.739325 | 0.00389 | 0.034827 | F | F | T | T |
| Maf | -0.535831 | 5.69442914 | 9.73806201 | 0.00389 | 0.034831 | T | F | T | T |
| Frat1 | 1.24696381 | 2.52453812 | 9.73599155 | 0.00389 | 0.034846 | T | F | T | F |
| Pacsin3 | 1.09011369 | 2.92914037 | 9.73152339 | 0.0039 | 0.034896 | F | F | T | F |
| Lmtk3 | 0.62677553 | 4.84965756 | 9.72274471 | 0.00391 | 0.035008 | T | F | F | T |
| Tspan3 | 0.43630883 | 6.43107114 | 9.71386392 | 0.00393 | 0.035122 | T | F | T | F |
| Gale | -0.6054583 | 5.16059469 | 9.71030157 | 0.00393 | 0.03515 | F | F | F | F |
| Tekt1 | 3.60319374 | -0.0366566 | 9.70871154 | 0.00393 | 0.03515 | F | F | F | F |
| Ccno | 3.7033905 | 0.13048136 | 9.70823118 | 0.00394 | 0.03515 | T | F | F | T |
| Kcnt2 | 2.41448362 | 0.82798224 | 9.70794487 | 0.00394 | 0.03515 | F | F | F | F |
| Metap2 | -0.3494383 | 7.75491772 | 9.70649956 | 0.00394 | 0.03515 | T | F | T | T |
| Ncoa4 | -0.4099709 | 7.07850731 | 9.70594323 | 0.00394 | 0.03515 | T | F | F | F |
| Hmox2 | -0.5465638 | 5.59541907 | 9.67574502 | 0.00399 | 0.035577 | F | F | F | T |
| Slc25a23 | 0.87401386 | 3.72504056 | 9.67327943 | 0.00399 | 0.035598 | T | F | F | F |
| Rims1 | 2.97707393 | 0.35294994 | 9.66999249 | 0.004 | 0.035621 | T | F | T | T |

| | | | | | | | | | |
|---|---|---|---|---|---|---|---|---|---|
| Mast1 | 3.21257489 | 0.26616195 | 9.66976979 | 0.004 | 0.035621 | T | F | F | F |
| Gspt1 | -0.3157383 | 8.53383337 | 9.66736593 | 0.004 | 0.035642 | T | F | T | T |
| Brinp2 | 7.30723972 | -0.9914625 | 9.71794378 | 0.004 | 0.035642 | T | F | T | F |
| Nfatc2ip | 0.70173074 | 4.33023578 | 9.66518581 | 0.00401 | 0.035644 | T | F | T | F |
| Ddx21 | -0.2779269 | 9.59987453 | 9.66064062 | 0.00401 | 0.035697 | F | F | T | F |
| Dennd1a | -0.5933293 | 5.23886344 | 9.65530424 | 0.00402 | 0.035761 | F | F | F | F |
| Avpr1b | 3.90285386 | -0.0686406 | 9.65277051 | 0.00403 | 0.035762 | T | F | F | T |
| Mospd1 | -0.8641494 | 3.95422325 | 9.65241118 | 0.00403 | 0.035762 | T | F | F | T |
| Aqp8 | 0.48377031 | 5.93332993 | 9.6522872 | 0.00403 | 0.035762 | F | F | F | F |
| Smarca4 | 0.29066873 | 9.06394261 | 9.6469128 | 0.00404 | 0.035827 | F | F | T | T |
| Col6a1 | 2.11547388 | 1.13951885 | 9.64256001 | 0.00404 | 0.035876 | F | F | F | F |
| Cnnm1 | 4.2630909 | -0.3132915 | 9.63263999 | 0.00406 | 0.036007 | F | F | F | F |
| Ap1s1 | -0.4871141 | 6.08046736 | 9.63180989 | 0.00406 | 0.036007 | F | F | F | F |
| Slc39a4 | 0.5068231 | 5.72824713 | 9.62854601 | 0.00407 | 0.036041 | F | F | T | F |
| Stx1b | 1.26054351 | 2.47440708 | 9.61827106 | 0.00408 | 0.03618 | F | F | F | F |
| Scn4a | 3.79800771 | -0.1251271 | 9.61037678 | 0.0041 | 0.036283 | T | F | F | F |
| Crtc2 | -0.6397485 | 4.87406876 | 9.60860906 | 0.0041 | 0.036295 | F | F | F | F |
| Adora1 | 0.82155957 | 3.83570804 | 9.60544262 | 0.00411 | 0.036328 | T | F | T | T |
| Slc32a1 | -1.3918595 | 2.6615133 | 9.6009128 | 0.00411 | 0.036381 | T | F | T | T |
| Serpine1 | 0.59254642 | 5.40181896 | 9.59652222 | 0.00412 | 0.036433 | T | F | T | T |
| Slc39a7 | 0.35267245 | 7.42594339 | 9.59506902 | 0.00412 | 0.03644 | T | F | F | F |
| Cpt1c | 1.92381787 | 1.37332491 | 9.57551521 | 0.00416 | 0.036721 | T | F | F | T |
| Gm10053 | -0.4208383 | 6.69338303 | 9.57072146 | 0.00417 | 0.036779 | F | F | F | F |
| Gm21992 | -1.802277 | 1.91467876 | 9.56729481 | 0.00417 | 0.036817 | F | F | F | F |
| Abce1 | -0.3229439 | 8.2097866 | 9.5641294 | 0.00418 | 0.03685 | T | F | T | T |
| Wdr35 | 0.8396068 | 3.75819701 | 9.56218678 | 0.00418 | 0.036857 | T | F | T | F |
| Gm14139 | -1.9145277 | 1.7899098 | 9.56122436 | 0.00418 | 0.036857 | F | F | F | F |
| Eogt | -0.6443548 | 4.82656952 | 9.56075727 | 0.00418 | 0.036857 | T | F | T | F |
| Ildr2 | 1.50414878 | 2.05483071 | 9.54854477 | 0.0042 | 0.037029 | F | T | T | T |
| H2bc6 | 0.39171067 | 7.50658357 | 9.5462686 | 0.00421 | 0.037049 | T | F | F | T |
| Cacna1h | 1.31820532 | 2.31008863 | 9.54107443 | 0.00422 | 0.037114 | F | F | T | F |
| Postn | -2.8309376 | 1.23057894 | 9.53908807 | 0.00422 | 0.037129 | T | F | F | F |
| Alkbh1 | 2.51107741 | 0.74670336 | 9.53768533 | 0.00422 | 0.037136 | F | F | T | F |
| Myo6 | -0.3596802 | 7.46218748 | 9.53103309 | 0.00424 | 0.037193 | T | F | T | T |
| Rbfox3 | 4.72408467 | -0.4607635 | 9.53101411 | 0.00424 | 0.037193 | T | F | T | F |
| Phyhip | 4.72408467 | -0.4607635 | 9.53101411 | 0.00424 | 0.037193 | T | F | F | T |
| Poglut2 | 0.66813229 | 4.88731576 | 9.52941342 | 0.00424 | 0.037203 | F | F | T | F |
| Camk2b | 4.30966029 | -0.2443757 | 9.52723931 | 0.00424 | 0.037222 | T | F | F | F |
| Smim1 | -0.8921939 | 3.94764054 | 9.51797512 | 0.00426 | 0.03735 | F | F | T | F |
| Rbp1 | 0.60485168 | 4.96125005 | 9.51634578 | 0.00426 | 0.03736 | F | F | F | F |
| Rspo1 | 2.62377669 | 0.6471417 | 9.51119064 | 0.00427 | 0.037425 | F | F | F | F |
| Man2b1 | 0.48317646 | 5.98392291 | 9.50929841 | 0.00427 | 0.037439 | F | F | T | F |

| | | | | | | | | | |
|---|---|---|---|---|---|---|---|---|---|
| Plagl2 | 0.54033897 | 5.44200022 | 9.50739886 | 0.00428 | 0.037451 | T | F | F | F |
| Pcdhga11 | 1.55569267 | 1.88587 | 9.50660346 | 0.00428 | 0.037451 | F | F | T | F |
| Reep6 | -0.3739898 | 7.31434584 | 9.50449041 | 0.00428 | 0.037456 | F | F | T | F |
| H2bc3 | 0.30569595 | 8.51757788 | 9.50433451 | 0.00428 | 0.037456 | F | F | F | F |
| Naa50 | -0.3624479 | 7.4864872 | 9.50131839 | 0.00429 | 0.037488 | T | F | T | T |
| Car3 | 2.72880461 | 0.79485784 | 9.49898866 | 0.00429 | 0.037509 | T | F | F | F |
| Stx6 | 0.70774828 | 4.29495285 | 9.49668688 | 0.0043 | 0.03753 | F | F | T | F |
| Nphp3 | 0.69243727 | 4.36750211 | 9.48495751 | 0.00432 | 0.037698 | T | F | T | F |
| Dmxl2 | -0.5689499 | 5.37574111 | 9.47531045 | 0.00433 | 0.037834 | T | F | F | T |
| Slfn9 | 0.76500576 | 4.06866777 | 9.47440465 | 0.00434 | 0.037834 | T | F | F | T |
| Astn2 | 2.69219875 | 0.59074099 | 9.45994434 | 0.00436 | 0.038046 | T | F | F | F |
| Tfpi | 0.55998974 | 5.22827239 | 9.45835155 | 0.00436 | 0.038056 | T | F | T | T |
| Dimt1 | -0.7640987 | 4.2307912 | 9.45218364 | 0.00438 | 0.038139 | T | F | T | T |
| Ube2o | -0.3529991 | 7.55126002 | 9.44561018 | 0.00439 | 0.038228 | F | F | F | F |
| Exoc6b | 0.60127886 | 4.97409269 | 9.44098085 | 0.0044 | 0.038286 | T | F | T | T |
| Dda1 | -0.47448 | 6.15151098 | 9.43846531 | 0.0044 | 0.038301 | F | F | T | F |
| Pla2g12a | -0.60302 | 5.21226059 | 9.43812493 | 0.0044 | 0.038301 | F | F | T | F |
| Filip1l | -1.3153037 | 2.61835493 | 9.43045833 | 0.00442 | 0.038396 | F | F | F | T |
| Capg | -0.5597418 | 5.73166237 | 9.43025231 | 0.00442 | 0.038396 | F | F | T | F |
| Hook3 | -0.5483503 | 5.7680427 | 9.427788 | 0.00442 | 0.03842 | F | F | T | T |
| March8 | -0.5509295 | 5.52780206 | 9.42622546 | 0.00442 | 0.03843 | T | T | T | T |
| Fam89b | 0.94968052 | 3.54504895 | 9.42495305 | 0.00443 | 0.038435 | F | F | F | F |
| Lrrc3b | 4.62970832 | -0.4967818 | 9.4111384 | 0.00445 | 0.038641 | F | F | F | T |
| Ywhaz | -0.2864932 | 9.61863319 | 9.40227234 | 0.00447 | 0.038755 | T | F | T | T |
| Nmd3 | -0.5567024 | 5.45332408 | 9.40217377 | 0.00447 | 0.038755 | T | F | F | T |
| Adamts1 | 0.58067408 | 5.04288813 | 9.40014956 | 0.00447 | 0.038772 | T | F | F | F |
| Enpp2 | 0.80894558 | 3.90052477 | 9.39916266 | 0.00447 | 0.038772 | T | F | F | F |
| Spag9 | -0.4245904 | 6.69428387 | 9.39132364 | 0.00449 | 0.038884 | T | F | T | T |
| Cyb561d2 | -0.8242803 | 4.03881761 | 9.38988916 | 0.00449 | 0.038892 | F | F | T | F |
| Sprtn | -0.5258844 | 5.69676007 | 9.3883254 | 0.00449 | 0.038901 | F | F | F | F |
| Snap29 | -0.5906009 | 5.19496382 | 9.38477078 | 0.0045 | 0.038944 | T | F | T | F |
| Pde3a | 0.77161243 | 3.98028874 | 9.36570251 | 0.00454 | 0.039239 | T | F | T | T |
| Galnt2 | 0.37854272 | 6.94615361 | 9.36406925 | 0.00454 | 0.03925 | T | F | T | T |
| Ikbkb | 0.74725567 | 4.18698606 | 9.35603795 | 0.00456 | 0.039359 | T | F | T | T |
| Zfp512 | 0.53153933 | 5.46904154 | 9.35549412 | 0.00456 | 0.039359 | T | T | T | T |
| Lin7c | -0.4364113 | 6.93439942 | 9.34882972 | 0.00457 | 0.039453 | T | F | F | F |
| Nfkb2 | 0.63137075 | 4.68615075 | 9.34380146 | 0.00458 | 0.039521 | T | F | F | F |
| Ttc23 | 1.01355805 | 3.29016217 | 9.33445384 | 0.0046 | 0.039659 | F | F | F | F |
| Zfp606 | 0.89569436 | 3.57387312 | 9.33193086 | 0.0046 | 0.039678 | F | F | T | F |
| Flrt1 | 3.35935412 | 0.0580593 | 9.33141543 | 0.0046 | 0.039678 | F | F | T | T |
| Blvra | -0.7988875 | 4.06832287 | 9.32740392 | 0.00461 | 0.039722 | F | F | F | F |
| Psg16 | 0.64504584 | 5.31541947 | 9.32684977 | 0.00461 | 0.039722 | T | T | F | T |

| | | | | | | | | | |
|---|---|---|---|---|---|---|---|---|---|
| Lims1 | -0.4329882 | 6.54863479 | 9.31972319 | 0.00462 | 0.039825 | T | F | T | T |
| Gfra2 | 1.36032165 | 2.21003194 | 9.31393671 | 0.00464 | 0.039895 | F | F | F | F |
| Ndrg4 | 1.76704171 | 1.57372269 | 9.31363294 | 0.00464 | 0.039895 | F | F | T | F |
| Xrcc1 | 0.47050336 | 5.97907057 | 9.31020328 | 0.00464 | 0.039936 | F | F | F | F |
| Zfp553 | 0.47022709 | 5.99225152 | 9.30711201 | 0.00465 | 0.039972 | F | F | F | F |
| Ptprj | 0.6920806 | 4.35194914 | 9.30145584 | 0.00466 | 0.040051 | T | F | T | F |
| Arid3b | 0.40407907 | 6.7894569 | 9.30018907 | 0.00466 | 0.040056 | T | F | T | T |
| Fubp1 | 0.32313273 | 8.03370109 | 9.2899872 | 0.00468 | 0.040212 | T | F | T | T |
| Calm1 | -0.3490997 | 8.25481546 | 9.28674147 | 0.00469 | 0.04025 | F | F | T | F |
| Mmp1b | 3.02637567 | 0.73109294 | 9.28554676 | 0.00469 | 0.040255 | T | F | T | F |
| Caskin1 | -0.7570363 | 4.2785811 | 9.28295711 | 0.0047 | 0.040282 | F | F | T | F |
| Rnf150 | 2.57926585 | 0.8333881 | 9.27728831 | 0.00471 | 0.040362 | T | T | T | T |
| Zfyve9 | 0.55421177 | 5.31312688 | 9.26794468 | 0.00473 | 0.040504 | F | F | T | T |
| Ptms | 0.47355345 | 6.03318739 | 9.26627705 | 0.00473 | 0.040506 | F | F | T | F |
| Agrp | -3.5294609 | 0.62053434 | 9.26592906 | 0.00473 | 0.040506 | F | F | F | F |
| Cnot6 | -0.4031848 | 6.80356949 | 9.26250054 | 0.00474 | 0.040542 | T | F | T | T |
| Agtrap | 0.55308314 | 5.2768786 | 9.26110588 | 0.00474 | 0.040542 | T | F | T | F |
| mt-Co2 | 1.10915124 | 4.75126337 | 9.26099321 | 0.00474 | 0.040542 | F | F | F | F |
| Slc25a26 | -1.1326331 | 3.06537157 | 9.25986077 | 0.00474 | 0.040545 | T | F | T | F |
| Smarce1 | 0.36436634 | 7.13281678 | 9.25860713 | 0.00474 | 0.040545 | F | F | T | F |
| Pdzk1 | -0.4383253 | 6.45704614 | 9.25801326 | 0.00475 | 0.040545 | T | F | F | F |
| Fam57a | -0.8021706 | 4.04530757 | 9.24587014 | 0.00477 | 0.040736 | F | F | F | F |
| Shd | 3.23124465 | 0.1134498 | 9.24298471 | 0.00478 | 0.040769 | F | F | F | F |
| Map3k9 | 0.73521184 | 4.10567776 | 9.23478754 | 0.00479 | 0.040893 | T | F | T | T |
| Ctnnd1 | -0.311945 | 8.68624663 | 9.23357743 | 0.00479 | 0.040898 | T | F | T | T |
| D430019H1 | 1.83906522 | 1.44452744 | 9.23172873 | 0.0048 | 0.040913 | T | F | T | F |
| Plekhh2 | 0.90818317 | 3.48604289 | 9.22783881 | 0.00481 | 0.040961 | T | F | T | T |
| Lap3 | -0.3472086 | 7.58071636 | 9.22709662 | 0.00481 | 0.040961 | T | F | F | T |
| Rab11fip1 | -0.3983373 | 6.83880209 | 9.22282904 | 0.00482 | 0.041018 | T | F | T | F |
| Moxd1 | 2.41732462 | 0.76245158 | 9.21770876 | 0.00483 | 0.04109 | T | F | T | F |
| Zfp746 | 0.65089576 | 4.54112262 | 9.21531418 | 0.00483 | 0.041115 | F | F | T | T |
| Cdkn1a | 0.90906545 | 3.44731768 | 9.21172979 | 0.00484 | 0.041161 | T | F | T | F |
| Zfp7 | 0.68320986 | 4.34800769 | 9.19379452 | 0.00488 | 0.041446 | T | F | F | F |
| Susd2 | 0.71117403 | 4.2182368 | 9.19339088 | 0.00488 | 0.041446 | T | F | F | T |
| Cdk19 | -0.8702526 | 3.82096658 | 9.19046206 | 0.00488 | 0.041481 | F | F | T | T |
| Ubc | -0.2612202 | 9.91213536 | 9.17920385 | 0.00491 | 0.041661 | F | F | F | F |
| Sec14l1 | -0.4019146 | 6.83904199 | 9.17209544 | 0.00492 | 0.041769 | T | F | T | T |
| Duox2 | 0.69525834 | 4.4978754 | 9.14949348 | 0.00497 | 0.042151 | T | F | F | T |
| Zmat4 | 3.92542492 | -0.220412 | 9.14718031 | 0.00497 | 0.042176 | T | F | T | T |
| Mmp16 | 3.21077282 | 0.14735912 | 9.14046884 | 0.00499 | 0.042278 | T | F | T | T |
| Plag1 | 1.26784488 | 2.36592552 | 9.1365177 | 0.00499 | 0.042332 | T | F | F | T |
| Isoc2a | 1.15243391 | 2.66876421 | 9.13415693 | 0.005 | 0.042358 | F | F | T | F |

| | | | | | | | | | |
|---|---|---|---|---|---|---|---|---|---|
| Fbxw4 | 1.10492686 | 2.80444093 | 9.13232022 | 0.005 | 0.042374 | F | F | T | F |
| Klf2 | -0.8323732 | 3.92624455 | 9.13130041 | 0.005 | 0.042375 | F | F | T | F |
| Rnf126 | -0.3670486 | 7.27571207 | 9.12384601 | 0.00502 | 0.042486 | F | F | F | F |
| Agap1 | 0.41906497 | 6.49360717 | 9.12327199 | 0.00502 | 0.042486 | T | T | T | T |
| Phb | -0.3343185 | 7.77446774 | 9.11670701 | 0.00504 | 0.042586 | T | F | F | F |
| Car13 | 1.74280011 | 1.55502758 | 9.1150095 | 0.00504 | 0.0426 | T | T | F | T |
| Avpi1 | -0.6182529 | 4.91488207 | 9.10817287 | 0.00505 | 0.042706 | F | F | T | T |
| Ptges2 | -0.6406422 | 4.75759422 | 9.10590773 | 0.00506 | 0.04273 | F | F | F | T |
| Gm3468 | -7.3298473 | -0.1309157 | 9.09692232 | 0.00508 | 0.042869 | F | F | F | F |
| Pou3f1 | 0.49347422 | 5.78608747 | 9.09569605 | 0.00508 | 0.042869 | F | F | F | F |
| Hkdc1 | 0.56263307 | 5.11346738 | 9.09472689 | 0.00508 | 0.042869 | F | F | F | F |
| Qk | 0.44621389 | 6.33476342 | 9.09451697 | 0.00508 | 0.042869 | T | F | T | T |
| Fbxl12 | -0.595754 | 5.07455428 | 9.08705595 | 0.0051 | 0.042987 | F | F | F | F |
| P4ha1 | 0.35475369 | 7.26381662 | 9.08163754 | 0.00511 | 0.043069 | F | T | T | T |
| Pcsk7 | -0.6698632 | 4.71807983 | 9.07701023 | 0.00512 | 0.043136 | T | F | T | T |
| Bcam | 0.54602986 | 5.64944936 | 9.07429071 | 0.00513 | 0.043169 | T | F | F | T |
| Vstm5 | -1.3030861 | 2.58263657 | 9.0722207 | 0.00513 | 0.04319 | F | F | T | F |
| Gsdme | 1.84493443 | 1.43049589 | 9.06886163 | 0.00514 | 0.043234 | F | F | F | F |
| Shoc2 | -0.4210715 | 6.58076081 | 9.06120481 | 0.00515 | 0.043357 | T | F | F | F |
| Plxna4 | 1.36870631 | 2.22212758 | 9.05704855 | 0.00516 | 0.043416 | F | F | T | T |
| Fdxr | 0.65000091 | 4.50886927 | 9.05556474 | 0.00517 | 0.043426 | F | F | F | F |
| Tuba1a | 0.35090945 | 7.28763739 | 9.0544112 | 0.00517 | 0.043431 | F | F | F | T |
| Stk32a | 2.5843152 | 0.64092754 | 9.0441413 | 0.00519 | 0.043602 | T | F | T | T |
| Ttbk1 | 0.67109439 | 4.37199759 | 9.03688297 | 0.00521 | 0.043719 | T | F | T | F |
| Ddhd2 | -0.5228475 | 5.63656488 | 9.03497389 | 0.00521 | 0.043737 | F | F | T | F |
| Tubb2b | 0.64178081 | 4.54887632 | 9.03299564 | 0.00522 | 0.043757 | F | F | F | T |
| Acp2 | 0.59589674 | 4.82782083 | 9.03138259 | 0.00522 | 0.04377 | T | F | T | T |
| Dna2 | -0.4504037 | 6.3204942 | 9.02313425 | 0.00524 | 0.043906 | T | F | F | T |
| Efcab6 | 7.21676582 | -1.0609322 | 9.06651971 | 0.00524 | 0.043927 | F | F | F | F |
| Rpl29 | -0.3053015 | 8.96365708 | 9.01960573 | 0.00525 | 0.043937 | F | F | F | F |
| Pik3r2 | 0.48529023 | 5.79266659 | 9.01401211 | 0.00526 | 0.044024 | F | F | T | T |
| Eno3 | 1.0232203 | 3.03770262 | 9.00932954 | 0.00527 | 0.044094 | F | F | F | F |
| Cd209c | -0.607852 | 4.99926414 | 9.0003562 | 0.00529 | 0.044244 | F | F | F | T |
| Polr2a | 0.27842135 | 9.03567459 | 8.97995683 | 0.00533 | 0.04461 | F | F | F | T |
| Zfp948 | -0.6227704 | 4.8319918 | 8.96663792 | 0.00536 | 0.04484 | F | F | F | F |
| Cep68 | 0.5046479 | 5.57994075 | 8.96599272 | 0.00537 | 0.04484 | T | F | T | F |
| Gpx4 | -0.3419598 | 7.58163329 | 8.94983212 | 0.0054 | 0.04513 | F | F | F | F |
| Tomm5 | -0.4453673 | 6.30451343 | 8.94662237 | 0.00541 | 0.045165 | F | F | F | F |
| Cacfd1 | 0.58842061 | 4.94444209 | 8.94529937 | 0.00541 | 0.045165 | F | F | T | F |
| Arhgap24 | 2.18649036 | 1.15508083 | 8.94525213 | 0.00541 | 0.045165 | T | F | T | T |
| Slc35f1 | 2.99910388 | 0.23346137 | 8.94268663 | 0.00542 | 0.045197 | T | F | T | F |
| Tmsb10 | -0.3032683 | 8.95973768 | 8.92598179 | 0.00546 | 0.0455 | F | F | F | F |

| | | | | | | | | | |
|---|---|---|---|---|---|---|---|---|---|
| Chrdl1 | 2.64881875 | 0.5201016 | 8.92170849 | 0.00547 | 0.045555 | T | F | T | T |
| Klc2 | -0.5299488 | 5.5680612 | 8.92130536 | 0.00547 | 0.045555 | F | F | F | F |
| Hspa9 | -0.2645082 | 9.67543405 | 8.91814904 | 0.00548 | 0.045599 | T | F | F | F |
| Ghitm | -0.3888899 | 6.91119686 | 8.91204121 | 0.00549 | 0.045699 | T | T | F | T |
| Fbxo21 | -0.4190222 | 8.54991557 | 8.91023428 | 0.00549 | 0.045717 | T | F | F | T |
| Cdk2ap2 | -0.7198316 | 4.34915162 | 8.90549945 | 0.00551 | 0.045791 | T | T | F | T |
| Dvl2 | 0.47449978 | 5.87116385 | 8.89572152 | 0.00553 | 0.045959 | T | F | F | F |
| Sbsn | 0.58408841 | 4.88527459 | 8.89505351 | 0.00553 | 0.045959 | F | F | F | F |
| Actr3b | 1.46569197 | 1.99007535 | 8.88940126 | 0.00554 | 0.046051 | T | F | F | F |
| Samd4 | 1.003408 | 3.16139122 | 8.8866726 | 0.00555 | 0.046087 | F | F | T | T |
| Magea5 | 5.24697077 | -0.7452689 | 8.87995887 | 0.00557 | 0.046201 | F | F | T | F |
| Pde10a | 0.94769404 | 3.2845854 | 8.8774293 | 0.00557 | 0.046233 | T | F | F | F |
| Dok2 | 1.88725963 | 1.43958012 | 8.87360414 | 0.00558 | 0.04629 | F | F | T | F |
| Smndc1 | -0.5200063 | 5.68902068 | 8.86877115 | 0.00559 | 0.046367 | T | F | F | F |
| Cltc | -0.2458003 | 10.2964922 | 8.86079336 | 0.00561 | 0.046507 | T | T | F | T |
| Kdm3a | 0.55828243 | 5.10884837 | 8.85203386 | 0.00563 | 0.046662 | T | F | T | T |
| Birc5 | -0.3828011 | 6.91938257 | 8.84020239 | 0.00566 | 0.046879 | T | F | T | F |
| Pald1 | 0.45710273 | 5.99254012 | 8.83590015 | 0.00567 | 0.046947 | F | F | T | T |
| Pdia3 | 0.27400172 | 9.28312471 | 8.83165073 | 0.00568 | 0.047014 | T | F | F | F |
| Adcy1 | 0.9522808 | 3.27286199 | 8.82462358 | 0.0057 | 0.047133 | T | F | T | T |
| Dennd2c | 1.20351404 | 2.53919111 | 8.82388101 | 0.0057 | 0.047133 | T | T | T | T |
| Agl | -0.5175011 | 5.62490877 | 8.82026144 | 0.00571 | 0.047188 | T | F | T | T |
| Stx12 | -0.4898006 | 5.85590176 | 8.8176145 | 0.00571 | 0.047223 | F | F | F | F |
| Lmnb2 | 0.42066148 | 6.4759492 | 8.81611338 | 0.00572 | 0.047235 | F | F | F | F |
| Slc27a2 | 2.34401417 | 0.78008529 | 8.81402023 | 0.00572 | 0.047259 | F | F | F | T |
| Edaradd | -3.9024519 | 0.37504131 | 8.80385545 | 0.00575 | 0.047439 | F | F | F | F |
| Ppp1r3b | -1.197871 | 2.81615943 | 8.80330522 | 0.00575 | 0.047439 | T | F | T | T |
| Fndc3b | -0.4012267 | 6.87626263 | 8.79169613 | 0.00578 | 0.047655 | T | F | T | F |
| Plekhg3 | 0.41588011 | 6.43709307 | 8.78566262 | 0.00579 | 0.047745 | T | F | T | T |
| Zyg11b | 0.43687938 | 6.33785987 | 8.78548911 | 0.00579 | 0.047745 | T | F | T | T |
| Hopx | 0.88692294 | 3.44764423 | 8.78418974 | 0.0058 | 0.047753 | T | T | F | T |
| Fam83e | -2.2551284 | 1.29790183 | 8.7784826 | 0.00581 | 0.047838 | T | F | F | F |
| Cdk20 | -2.2404198 | 1.36152574 | 8.77818605 | 0.00581 | 0.047838 | F | F | F | F |
| Amer2 | 3.28922885 | 0.02136035 | 8.77736369 | 0.00581 | 0.047838 | F | F | T | T |
| Aass | -0.6224788 | 5.07758226 | 8.77519377 | 0.00582 | 0.047864 | T | F | F | T |
| Slc25a32 | -0.7942815 | 4.19438373 | 8.76996925 | 0.00583 | 0.047942 | F | F | T | F |
| Cnot6l | -0.5409021 | 5.41873534 | 8.76957718 | 0.00583 | 0.047942 | F | F | T | T |
| Trp53bp2 | 0.48234907 | 5.82136949 | 8.76569913 | 0.00584 | 0.048003 | T | F | T | F |
| Rbm47 | -0.3520411 | 7.32694675 | 8.76477345 | 0.00584 | 0.048004 | T | F | T | T |
| Ipo7 | -0.3072536 | 8.39036251 | 8.76131802 | 0.00585 | 0.048056 | T | F | F | T |
| Nfya | -0.4194749 | 6.57244984 | 8.75860885 | 0.00586 | 0.048093 | T | F | T | T |
| Plekha2 | -0.4644235 | 6.33443932 | 8.75329437 | 0.00587 | 0.048184 | T | F | T | F |

| | | | | | | | | | |
|---|---|---|---|---|---|---|---|---|---|
| Ube2d3 | -0.3590292 | 7.71487557 | 8.74446106 | 0.0059 | 0.048348 | T | F | T | T |
| Cep350 | -0.3676301 | 7.07532096 | 8.7434658 | 0.0059 | 0.04835 | T | F | F | T |
| Paics | 0.29317404 | 8.47247134 | 8.74136285 | 0.0059 | 0.048375 | T | F | F | F |
| Scrib | 0.35297059 | 7.32545511 | 8.71648564 | 0.00597 | 0.04886 | F | F | F | F |
| Lrp6 | 0.32368429 | 7.67781045 | 8.71619248 | 0.00597 | 0.04886 | T | T | T | T |
| Ankrd37 | -1.2731386 | 2.61149823 | 8.71190021 | 0.00598 | 0.048931 | F | F | F | F |
| Syde2 | -0.444214 | 6.25215375 | 8.69634746 | 0.00602 | 0.049238 | F | T | F | T |
| Tmbim6 | 0.29838912 | 8.2545171 | 8.67379814 | 0.00608 | 0.049695 | F | F | T | F |

**Table S2F. DEGs between degenerating and non-degenerating female embryos.**

| | logFC | logCPM | F | PValue | FDR < 0.05 | microrna .org | mirdb | mir walk | target scan |
|---|---|---|---|---|---|---|---|---|---|
| Entpd1 | 8.96289958 | 8.14571567 | 1351.09044 | 2.58E-44 | 5.59E-40 | T | T | T | T |
| Plet1 | 7.30772613 | 8.78699773 | 1272.61257 | 1.56E-43 | 1.70E-39 | T | F | F | T |
| Serpinb9e | 9.66828363 | 7.00352076 | 1145.31661 | 3.69E-42 | 2.67E-38 | T | T | F | T |
| Adgrf5 | 7.02744109 | 8.51053469 | 1118.82468 | 7.43E-42 | 3.25E-38 | T | F | T | T |
| Fstl3 | 7.73288748 | 7.5653684 | 1118.45416 | 7.51E-42 | 3.25E-38 | F | F | F | F |
| Lgals9 | 7.99003424 | 7.23945131 | 1078.77422 | 2.21E-41 | 7.97E-38 | F | F | F | T |
| Tfap2c | 6.37905688 | 8.76872328 | 1072.31059 | 2.64E-41 | 8.18E-38 | T | F | F | F |
| Lipg | 8.10096634 | 7.65274836 | 1064.83629 | 3.25E-41 | 8.81E-38 | T | F | T | F |
| Gjb3 | 6.52584627 | 8.5756645 | 1017.51221 | 1.26E-40 | 3.03E-37 | F | F | T | F |
| Cdh5 | 7.76202384 | 9.63190292 | 993.439061 | 2.56E-40 | 5.17E-37 | T | T | F | T |
| AU018091 | 9.63048328 | 7.81579287 | 991.413083 | 2.72E-40 | 5.17E-37 | F | F | F | F |
| Prune2 | 6.28654777 | 8.27621682 | 989.726924 | 2.86E-40 | 5.17E-37 | T | F | T | F |
| Gjb5 | 7.8738867 | 7.59446571 | 982.784796 | 3.53E-40 | 5.89E-37 | T | F | F | T |
| Krt19 | 5.860071 | 8.89554684 | 979.116647 | 3.94E-40 | 6.11E-37 | F | F | T | F |
| Ldoc1 | 7.22028655 | 7.17167285 | 953.929314 | 8.54E-40 | 1.23E-36 | T | F | F | F |
| Itgb3 | 8.01641405 | 8.1184352 | 928.623532 | 1.89E-39 | 2.56E-36 | F | F | T | T |
| Plac1 | 6.81622267 | 7.85028034 | 926.812012 | 2.00E-39 | 2.56E-36 | T | F | F | F |
| Prl4a1 | 8.75843871 | 6.51350345 | 905.096912 | 4.04E-39 | 4.86E-36 | F | F | F | F |
| Sfmbt2 | 5.42841245 | 9.04270397 | 876.884557 | 1.03E-38 | 1.17E-35 | T | F | T | T |
| Apob | -6.5460474 | 10.8874784 | 867.531764 | 1.41E-38 | 1.53E-35 | F | F | F | F |
| Ascl2 | 7.50669501 | 6.74866673 | 848.69501 | 2.69E-38 | 2.77E-35 | F | F | T | F |
| Adcy7 | 6.37325999 | 7.16188111 | 816.354099 | 8.40E-38 | 8.11E-35 | T | F | T | F |
| Nav3 | 7.85119057 | 7.08225271 | 815.677432 | 8.61E-38 | 8.11E-35 | T | F | T | T |
| Spn | 6.8740821 | 6.8031936 | 802.48071 | 1.39E-37 | 1.25E-34 | T | F | T | F |
| Serpinb9g | 9.28403688 | 6.44328225 | 786.652557 | 2.49E-37 | 2.16E-34 | T | T | F | T |
| Nid1 | 5.25550518 | 8.6748364 | 784.766192 | 2.67E-37 | 2.22E-34 | T | F | F | T |
| Ceacam9 | 8.63210391 | 6.43525735 | 781.640515 | 3.00E-37 | 2.41E-34 | T | F | F | T |
| Itgb4 | 6.60988676 | 6.68897123 | 724.378151 | 2.75E-36 | 2.13E-33 | F | F | F | F |
| Cubn | -6.0228818 | 10.3291592 | 712.375934 | 4.47E-36 | 3.34E-33 | F | F | F | F |
| Pla2g2f | 9.71898438 | 5.64418756 | 708.104599 | 5.32E-36 | 3.84E-33 | F | F | F | F |
| Itga7 | 7.57901045 | 6.10278722 | 703.600781 | 6.40E-36 | 4.48E-33 | T | F | F | T |
| Adm | 6.46087247 | 8.24262342 | 685.250867 | 1.38E-35 | 9.16E-33 | T | F | T | F |
| Mical2 | 7.04927772 | 6.92468916 | 684.920748 | 1.39E-35 | 9.16E-33 | T | F | T | T |
| Slc2a3 | -5.9715783 | 10.5888886 | 678.460667 | 1.83E-35 | 1.17E-32 | T | F | T | T |
| Pltp | 6.64271425 | 6.48796009 | 673.512642 | 2.27E-35 | 1.40E-32 | F | F | F | F |
| Irs3 | 8.0476319 | 5.7411902 | 657.292387 | 4.58E-35 | 2.76E-32 | F | F | F | T |
| Elf4 | 5.68723364 | 6.90617385 | 651.919133 | 5.80E-35 | 3.40E-32 | T | F | T | T |
| Ero1l | 4.64432732 | 8.6600711 | 643.19411 | 8.55E-35 | 4.88E-32 | F | F | T | F |
| Apobr | 7.72513474 | 5.72157618 | 626.823436 | 1.79E-34 | 9.97E-32 | T | F | F | F |

| | | | | | | | | | |
|---|---|---|---|---|---|---|---|---|---|
| Gata2 | 5.05848437 | 7.8573115 | 624.278777 | 2.02E-34 | 1.09E-31 | F | F | F | F |
| Pcdh12 | 8.23954148 | 5.5971829 | 623.836808 | 2.06E-34 | 1.09E-31 | F | F | F | T |
| Fthl17a | 9.4711624 | 5.39123139 | 616.199739 | 2.93E-34 | 1.51E-31 | F | F | F | F |
| Lin28a | -5.5431752 | 10.0552171 | 604.155381 | 5.15E-34 | 2.60E-31 | F | F | F | F |
| Serpinb9f | 8.87154275 | 5.52287534 | 600.472026 | 6.14E-34 | 3.02E-31 | T | T | F | T |
| Lgals1 | 5.03535135 | 8.10060081 | 595.990702 | 7.60E-34 | 3.66E-31 | F | F | T | F |
| Sfn | 5.9918174 | 6.36820763 | 589.710967 | 1.03E-33 | 4.85E-31 | F | F | F | F |
| Sox11 | -7.1074273 | 8.09327693 | 582.825796 | 1.44E-33 | 6.63E-31 | T | F | T | T |
| Nfkbiz | 5.24109218 | 6.94789348 | 573.905661 | 2.23E-33 | 1.01E-30 | T | F | T | T |
| Ccnd2 | -7.0778614 | 8.06386414 | 565.578476 | 3.38E-33 | 1.50E-30 | T | F | T | T |
| Adap1 | 7.10130021 | 5.75121843 | 555.366796 | 5.67E-33 | 2.46E-30 | T | F | F | T |
| Gpc3 | -6.2166427 | 8.54042803 | 552.543611 | 6.55E-33 | 2.79E-30 | F | F | F | F |
| Pou5f1 | -7.9804873 | 7.73489174 | 552.10689 | 6.70E-33 | 2.80E-30 | T | F | F | F |
| Thbd | 5.80322351 | 6.30215097 | 536.415119 | 1.52E-32 | 6.18E-30 | T | F | F | T |
| Sema7a | 5.1574878 | 6.85406633 | 536.137183 | 1.54E-32 | 6.18E-30 | F | F | T | F |
| Serpinb9b | 6.69238041 | 5.81001246 | 530.506963 | 2.07E-32 | 8.18E-30 | T | F | F | T |
| Elf5 | 7.60420997 | 5.47315256 | 527.411929 | 2.45E-32 | 9.42E-30 | T | F | F | T |
| Krt7 | 5.0591163 | 6.80828432 | 527.175781 | 2.48E-32 | 9.42E-30 | F | F | F | F |
| Ppard | 4.44455187 | 7.64406863 | 522.637454 | 3.16E-32 | 1.18E-29 | F | F | F | T |
| Axin2 | -7.6257242 | 7.25195711 | 514.139917 | 5.02E-32 | 1.84E-29 | F | F | F | F |
| Sept5 | 6.30433737 | 6.1112795 | 512.876389 | 5.38E-32 | 1.94E-29 | F | F | F | F |
| Was | 6.44128876 | 5.71499352 | 512.466323 | 5.50E-32 | 1.95E-29 | T | F | T | F |
| Slc7a1 | 4.57497978 | 7.68423785 | 505.622096 | 8.03E-32 | 2.81E-29 | T | F | F | F |
| Col18a1 | -6.0112212 | 8.2331292 | 505.087457 | 8.27E-32 | 2.85E-29 | T | F | F | F |
| Arpc1b | 4.37970605 | 7.72120054 | 504.552601 | 8.52E-32 | 2.88E-29 | T | F | F | T |
| Lgals3 | 6.32492287 | 6.37164858 | 504.304091 | 8.64E-32 | 2.88E-29 | T | F | F | F |
| 3830417A1 | 7.57248027 | 5.25587604 | 500.727777 | 1.05E-31 | 3.46E-29 | F | F | F | F |
| Smo | -6.9161613 | 7.45099879 | 498.6295 | 1.19E-31 | 3.84E-29 | F | F | F | F |
| Gtf2i | -5.5359784 | 8.59866773 | 490.993042 | 1.83E-31 | 5.83E-29 | F | F | F | F |
| Hic1 | 7.09369888 | 5.29401892 | 489.750551 | 1.96E-31 | 6.17E-29 | F | F | F | F |
| Myof | 4.92889079 | 6.70141072 | 489.187723 | 2.03E-31 | 6.28E-29 | T | F | T | F |
| Tenm4 | -6.9716369 | 7.39332614 | 487.978955 | 2.17E-31 | 6.64E-29 | T | F | T | T |
| Cdkn1c | 3.90000324 | 9.24027638 | 486.636202 | 2.35E-31 | 7.07E-29 | T | T | F | T |
| Clip2 | 4.12777192 | 7.96391872 | 483.643814 | 2.79E-31 | 8.27E-29 | F | F | T | F |
| Flnb | 3.37735076 | 10.7984595 | 483.438357 | 2.82E-31 | 8.27E-29 | T | F | T | T |
| Arhgef6 | 8.33186492 | 4.93608661 | 482.43976 | 2.99E-31 | 8.64E-29 | T | F | T | T |
| Sdc1 | 3.81727906 | 9.17464825 | 480.02532 | 3.44E-31 | 9.72E-29 | T | F | T | T |
| Mbnl3 | 5.28837502 | 6.32656827 | 479.963818 | 3.45E-31 | 9.72E-29 | T | T | T | T |
| Apoe | -5.4473331 | 9.02155164 | 476.388915 | 4.25E-31 | 1.18E-28 | F | F | F | F |
| Greb1 | -7.9260268 | 6.93253501 | 475.959468 | 4.36E-31 | 1.20E-28 | F | F | F | F |
| Hist1h2br | -14.181628 | 6.57668411 | 473.524214 | 5.03E-31 | 1.36E-28 | F | F | T | T |
| Zc3h7b | -6.5937407 | 7.3816057 | 470.853586 | 5.89E-31 | 1.58E-28 | T | F | T | T |

| | | | | | | | | | |
|---|---|---|---|---|---|---|---|---|---|
| Hspb1 | 4.55832888 | 6.91894025 | 467.550144 | 7.17E-31 | 1.90E-28 | T | F | F | T |
| Dock8 | 5.01279803 | 6.54998333 | 466.765298 | 7.51E-31 | 1.96E-28 | T | F | F | T |
| Tgm2 | 4.53258945 | 6.96506473 | 464.605395 | 8.55E-31 | 2.20E-28 | F | F | T | T |
| Cdh3 | 4.00092199 | 7.9954788 | 464.456211 | 8.62E-31 | 2.20E-28 | F | F | T | F |
| Zbtb10 | 3.9365063 | 8.30085902 | 463.3155 | 9.24E-31 | 2.33E-28 | T | F | T | T |
| Prl2a1 | 8.73912565 | 4.87053119 | 462.998273 | 9.41E-31 | 2.35E-28 | F | F | F | F |
| Dab2 | -5.503284 | 8.23016419 | 460.198565 | 1.11E-30 | 2.75E-28 | T | F | T | T |
| Gata3 | 4.58604936 | 7.0322199 | 459.324073 | 1.17E-30 | 2.86E-28 | F | F | F | F |
| Mex3a | -6.4723198 | 7.31818135 | 455.891086 | 1.45E-30 | 3.49E-28 | T | F | T | T |
| Plec | 3.42993456 | 9.81177403 | 448.113273 | 2.33E-30 | 5.56E-28 | T | F | F | T |
| Lrp2 | -5.5205899 | 8.82147431 | 442.225801 | 3.36E-30 | 7.93E-28 | T | F | F | F |
| Lef1 | -7.5257219 | 6.80741252 | 438.474699 | 4.26E-30 | 9.92E-28 | T | F | F | F |
| Tram2 | 3.89796732 | 7.91993885 | 431.489516 | 6.63E-30 | 1.53E-27 | T | F | F | F |
| Frem2 | -7.3578429 | 6.93480298 | 431.199367 | 6.75E-30 | 1.54E-27 | T | T | F | T |
| Plac8 | 4.69185868 | 6.72116806 | 430.528543 | 7.05E-30 | 1.58E-27 | F | F | F | F |
| Pcdh19 | -7.4200713 | 6.89864824 | 430.462344 | 7.08E-30 | 1.58E-27 | T | F | T | T |
| Gm2a | 4.74722081 | 6.87954739 | 425.850259 | 9.53E-30 | 2.11E-27 | F | F | T | F |
| Psg29 | 5.51445972 | 7.11777032 | 425.135072 | 9.98E-30 | 2.19E-27 | T | F | T | F |
| Armcx2 | -8.115433 | 6.60319807 | 424.420454 | 1.05E-29 | 2.27E-27 | F | F | T | F |
| Slc13a4 | -6.6135241 | 7.07291901 | 423.073243 | 1.14E-29 | 2.45E-27 | T | F | F | F |
| Nrk | 3.42740764 | 9.46714356 | 418.964972 | 1.49E-29 | 3.17E-27 | T | F | T | T |
| Tex19.1 | 5.41354864 | 5.72174916 | 418.108492 | 1.58E-29 | 3.32E-27 | T | F | F | F |
| Efhd2 | 4.16280298 | 7.35093763 | 416.080374 | 1.80E-29 | 3.76E-27 | T | F | T | T |
| Chdh | 5.92991198 | 5.79555122 | 415.444928 | 1.88E-29 | 3.88E-27 | T | T | F | T |
| Msi1 | -6.0752097 | 7.13741913 | 412.099943 | 2.35E-29 | 4.80E-27 | F | F | F | F |
| Clic6 | -6.0884441 | 7.15044597 | 410.531504 | 2.61E-29 | 5.28E-27 | T | F | T | T |
| Esam | 5.78069182 | 5.80748136 | 409.549642 | 2.78E-29 | 5.59E-27 | T | F | F | F |
| Eps8l2 | 4.41864037 | 6.65563749 | 407.824392 | 3.12E-29 | 6.21E-27 | F | F | F | F |
| Trh | -9.2711901 | 6.56369692 | 407.018735 | 3.30E-29 | 6.50E-27 | F | F | F | F |
| Rusc2 | 5.43047065 | 5.61248389 | 405.418293 | 3.67E-29 | 7.18E-27 | F | F | T | F |
| Nacc1 | -4.7951876 | 8.55551903 | 404.135592 | 4.01E-29 | 7.76E-27 | T | F | T | T |
| Tnfrsf19 | -7.8510713 | 6.52945029 | 403.537303 | 4.17E-29 | 8.00E-27 | T | F | F | T |
| Dnase1l3 | 7.31601443 | 4.75566471 | 402.270722 | 4.55E-29 | 8.65E-27 | T | F | T | T |
| Fzd2 | -6.6942385 | 6.82350587 | 402.072768 | 4.61E-29 | 8.69E-27 | F | F | F | F |
| Sulf1 | -7.420945 | 6.6606646 | 401.940794 | 4.65E-29 | 8.69E-27 | T | F | T | T |
| Afp | -6.6422756 | 6.83556069 | 398.693452 | 5.80E-29 | 1.08E-26 | F | F | F | F |
| Tmem132e | 5.25460304 | 5.95184778 | 398.490086 | 5.88E-29 | 1.08E-26 | T | F | F | F |
| Prl3d1 | 5.9923519 | 6.33653599 | 397.437035 | 6.32E-29 | 1.15E-26 | F | F | F | F |
| Anxa4 | 5.04718981 | 5.87401735 | 397.387688 | 6.35E-29 | 1.15E-26 | T | F | T | F |
| Prkch | 4.32438512 | 6.70874757 | 396.773863 | 6.62E-29 | 1.19E-26 | F | F | F | F |
| Armcx4 | -7.9554523 | 6.4441088 | 391.164351 | 9.76E-29 | 1.73E-26 | F | F | F | F |
| Apoa1 | -6.2489576 | 7.74855428 | 389.212133 | 1.12E-28 | 1.97E-26 | F | F | F | F |

| | | | | | | | | | |
|---|---|---|---|---|---|---|---|---|---|
| Capn1 | 4.24705848 | 6.71654343 | 387.820825 | 1.23E-28 | 2.16E-26 | F | F | F | F |
| Nid2 | -5.15745 | 7.73796648 | 385.801506 | 1.42E-28 | 2.46E-26 | T | F | F | T |
| Sall2 | -7.754383 | 6.43346223 | 385.498301 | 1.45E-28 | 2.50E-26 | F | F | T | F |
| Cd82 | 4.54651397 | 6.45398841 | 383.252906 | 1.70E-28 | 2.90E-26 | F | F | T | F |
| Palm | 3.73788715 | 8.23603708 | 382.704781 | 1.77E-28 | 2.99E-26 | F | F | F | F |
| Serpinb9c | 8.57097271 | 4.69519177 | 382.398107 | 1.81E-28 | 3.04E-26 | T | F | F | F |
| Ptpn14 | -5.8512187 | 7.05670397 | 381.179699 | 1.97E-28 | 3.29E-26 | T | F | T | T |
| Pbx1 | -5.6435869 | 7.30120937 | 380.816098 | 2.02E-28 | 3.35E-26 | T | F | T | T |
| Aplnr | -6.5075906 | 6.76315527 | 378.185747 | 2.44E-28 | 4.01E-26 | T | F | T | T |
| Rgcc | 6.97458485 | 4.68354836 | 376.454138 | 2.76E-28 | 4.50E-26 | T | F | F | F |
| Gja1 | -4.7322741 | 8.19677816 | 375.89427 | 2.88E-28 | 4.65E-26 | F | F | F | F |
| Plpp1 | 4.81530395 | 5.92389241 | 375.445202 | 2.97E-28 | 4.77E-26 | F | F | T | F |
| Ldlr | 3.73170482 | 7.63185041 | 375.209561 | 3.02E-28 | 4.82E-26 | T | F | T | T |
| S100a6 | 5.4141583 | 5.65204567 | 374.800213 | 3.11E-28 | 4.93E-26 | F | F | F | F |
| Sct | 7.11011116 | 5.53197755 | 374.584195 | 3.16E-28 | 4.97E-26 | F | F | F | F |
| Adam8 | 5.8371921 | 5.17314755 | 374.236704 | 3.24E-28 | 5.06E-26 | F | F | F | F |
| Liph | 6.11727715 | 5.19444014 | 374.024717 | 3.29E-28 | 5.10E-26 | T | F | T | T |
| Fam129a | 4.57395399 | 6.17123033 | 372.303289 | 3.73E-28 | 5.74E-26 | F | F | F | T |
| Kif26a | 3.83107268 | 7.88664994 | 371.652396 | 3.91E-28 | 5.97E-26 | F | F | F | F |
| Dlx3 | 4.75545315 | 6.20222489 | 369.667054 | 4.52E-28 | 6.85E-26 | T | F | T | T |
| Rlim | 3.19262731 | 9.73372713 | 369.484014 | 4.58E-28 | 6.90E-26 | T | F | T | F |
| Zic3 | -7.6535592 | 6.50085708 | 368.33059 | 4.98E-28 | 7.45E-26 | T | F | T | T |
| AI661453 | 3.74398562 | 7.4452791 | 367.923179 | 5.14E-28 | 7.63E-26 | T | F | T | F |
| Crispld2 | 6.99620737 | 4.78461974 | 367.521733 | 5.29E-28 | 7.80E-26 | F | F | T | F |
| Ccnd1 | -5.5037093 | 7.28914973 | 367.144828 | 5.44E-28 | 7.97E-26 | T | F | T | T |
| Dbn1 | -5.3339406 | 7.30387824 | 366.292541 | 5.79E-28 | 8.42E-26 | F | F | F | F |
| Myc | 3.78697825 | 8.18980304 | 366.176202 | 5.84E-28 | 8.44E-26 | F | F | F | F |
| Sema4d | 3.68337635 | 7.69907451 | 365.230032 | 6.26E-28 | 8.99E-26 | F | F | F | F |
| Sema3f | 3.87725681 | 7.00821891 | 364.926361 | 6.40E-28 | 9.13E-26 | F | F | F | F |
| Tnfrsf9 | 7.31889529 | 4.84690646 | 364.69611 | 6.51E-28 | 9.23E-26 | T | F | F | F |
| Tenm3 | -6.272558 | 6.70238887 | 364.571085 | 6.57E-28 | 9.25E-26 | T | F | T | T |
| Pde4dip | 3.52064412 | 7.90935323 | 362.90161 | 7.44E-28 | 1.04E-25 | T | F | T | F |
| Mpzl2 | 4.24056331 | 6.4805981 | 360.799123 | 8.70E-28 | 1.21E-25 | T | F | F | F |
| Afap1l2 | 5.51407664 | 5.40501913 | 360.135774 | 9.14E-28 | 1.26E-25 | T | F | T | T |
| Rassf4 | 5.2645707 | 5.85300876 | 359.045949 | 9.92E-28 | 1.36E-25 | T | F | T | T |
| Hmox1 | 3.75452911 | 7.10530722 | 357.062458 | 1.15E-27 | 1.57E-25 | F | F | T | F |
| Asap3 | 4.71282408 | 5.96650507 | 357.000735 | 1.16E-27 | 1.57E-25 | F | F | T | F |
| T | -7.7201884 | 7.47584229 | 356.314603 | 1.22E-27 | 1.64E-25 | F | F | F | F |
| Myadm | 3.43351558 | 8.44671669 | 356.202977 | 1.23E-27 | 1.64E-25 | F | F | F | T |
| Sall3 | -7.5994028 | 6.5971343 | 356.100625 | 1.24E-27 | 1.65E-25 | T | F | T | T |
| Mvp | 4.2945143 | 6.50213465 | 355.557658 | 1.29E-27 | 1.70E-25 | F | F | F | F |
| Olr1 | 6.46055629 | 4.94085137 | 354.824784 | 1.36E-27 | 1.79E-25 | T | F | T | T |

| | | | | | | | | | |
|---|---|---|---|---|---|---|---|---|---|
| Ttr | -6.3193619 | 6.98846513 | 354.279464 | 1.42E-27 | 1.86E-25 | T | F | F | T |
| Nkx1-2 | -7.2828805 | 6.95216578 | 353.910704 | 1.46E-27 | 1.90E-25 | T | F | T | F |
| Mest | -5.8895007 | 7.87990837 | 352.862075 | 1.58E-27 | 2.04E-25 | T | F | T | T |
| Satb1 | 3.51184663 | 8.17432627 | 352.334432 | 1.65E-27 | 2.11E-25 | F | T | T | T |
| Robo1 | -6.2862969 | 6.73961959 | 351.58894 | 1.74E-27 | 2.22E-25 | T | F | F | T |
| Kcnk6 | 4.5135314 | 6.21669601 | 351.524361 | 1.75E-27 | 2.22E-25 | T | F | T | F |
| Spred1 | -6.2008997 | 6.62770692 | 349.872946 | 1.99E-27 | 2.50E-25 | T | F | F | T |
| Fes | 4.91269562 | 5.60617593 | 349.630017 | 2.02E-27 | 2.54E-25 | F | F | F | F |
| Cd24a | -5.0853448 | 8.18850201 | 349.225601 | 2.09E-27 | 2.60E-25 | F | F | F | F |
| Ptk7 | -4.395179 | 8.54171129 | 348.317369 | 2.24E-27 | 2.77E-25 | F | F | F | F |
| Kctd12 | 4.29375106 | 6.43744015 | 347.447693 | 2.39E-27 | 2.95E-25 | T | F | T | T |
| Rgma | -6.2963191 | 6.61164154 | 347.032338 | 2.47E-27 | 3.03E-25 | T | F | T | T |
| Mdk | -5.5880405 | 7.75109435 | 344.776279 | 2.94E-27 | 3.58E-25 | T | T | T | T |
| Exoc3l4 | 6.32546393 | 5.01924115 | 344.240691 | 3.07E-27 | 3.70E-25 | F | F | F | T |
| Tpm2 | 4.0269004 | 6.70859239 | 344.228676 | 3.07E-27 | 3.70E-25 | F | F | T | F |
| Car2 | 3.52007874 | 8.08044244 | 343.383566 | 3.28E-27 | 3.93E-25 | T | F | F | F |
| Jak3 | 7.4628855 | 4.37202156 | 342.926139 | 3.40E-27 | 4.05E-25 | T | T | F | T |
| Peg10 | 2.58444064 | 11.8505299 | 342.834625 | 3.42E-27 | 4.05E-25 | T | F | F | T |
| Igfbp2 | -6.7695425 | 6.35022088 | 342.212003 | 3.59E-27 | 4.23E-25 | F | F | F | F |
| Rbp4 | -6.8686318 | 6.78623987 | 341.406405 | 3.83E-27 | 4.48E-25 | F | F | T | F |
| Arhgap35 | -4.6012436 | 7.87133605 | 341.233558 | 3.88E-27 | 4.52E-25 | T | F | T | F |
| Dll1 | -6.4836712 | 6.54814119 | 340.683191 | 4.05E-27 | 4.69E-25 | F | F | F | F |
| Fn1 | -3.5339202 | 10.5001483 | 338.303766 | 4.88E-27 | 5.63E-25 | T | F | F | F |
| Serpine1 | 5.51106284 | 5.19288756 | 334.526379 | 6.58E-27 | 7.53E-25 | T | F | T | T |
| Cald1 | 3.23781964 | 8.4804903 | 334.47609 | 6.61E-27 | 7.53E-25 | T | T | T | T |
| Fbn2 | -7.0348085 | 6.2946361 | 334.433833 | 6.63E-27 | 7.53E-25 | T | F | T | F |
| Vrtn | -7.6672098 | 6.34712159 | 330.358653 | 9.19E-27 | 1.04E-24 | F | F | F | F |
| Jup | 3.27028318 | 8.24184566 | 330.185305 | 9.31E-27 | 1.05E-24 | F | F | T | T |
| Sfrp1 | -6.245428 | 6.59087417 | 329.619864 | 9.75E-27 | 1.09E-24 | F | F | T | T |
| Foxh1 | -6.5472601 | 6.30849964 | 329.008031 | 1.02E-26 | 1.14E-24 | F | F | F | F |
| Plxna2 | -5.4408816 | 6.83369998 | 328.765185 | 1.04E-26 | 1.16E-24 | T | F | T | T |
| Ano6 | 3.45111865 | 7.48510189 | 326.960213 | 1.21E-26 | 1.33E-24 | F | F | T | T |
| Procr | 6.31822949 | 4.92500069 | 326.518339 | 1.25E-26 | 1.37E-24 | T | F | F | T |
| Ror2 | -6.251168 | 6.44732546 | 325.451206 | 1.37E-26 | 1.49E-24 | T | F | T | F |
| Anxa2 | 4.15067962 | 6.7019232 | 324.550961 | 1.47E-26 | 1.59E-24 | F | F | F | F |
| Serpinb9 | 5.91192724 | 4.78044174 | 324.335804 | 1.50E-26 | 1.61E-24 | T | F | T | T |
| Hand1 | 3.33027263 | 8.68780894 | 322.855874 | 1.69E-26 | 1.81E-24 | T | T | F | T |
| Sin3b | 3.23333649 | 8.17433685 | 322.581335 | 1.73E-26 | 1.85E-24 | F | F | F | F |
| Car14 | -7.3772193 | 6.05836975 | 320.839152 | 1.99E-26 | 2.12E-24 | T | F | F | F |
| Shtn1 | 4.73308691 | 5.55224435 | 320.137993 | 2.11E-26 | 2.23E-24 | F | F | T | F |
| Ajuba | -4.7584211 | 7.46609879 | 318.781563 | 2.36E-26 | 2.49E-24 | F | F | F | F |
| Epn3 | 4.74980214 | 5.48333355 | 318.179206 | 2.48E-26 | 2.60E-24 | F | F | F | T |

| | | | | | | | | | |
|---|---|---|---|---|---|---|---|---|---|
| Gli2 | -7.6432997 | 5.91518182 | 316.646267 | 2.82E-26 | 2.94E-24 | T | F | T | F |
| Etv5 | -5.231558 | 6.83993108 | 315.810217 | 3.03E-26 | 3.14E-24 | T | F | T | T |
| Ets2 | 3.32586111 | 7.89947809 | 315.751918 | 3.04E-26 | 3.14E-24 | F | F | T | F |
| Jag2 | 4.46076705 | 5.93641071 | 312.507525 | 3.99E-26 | 4.10E-24 | F | F | F | F |
| Crim1 | 3.86888002 | 6.64813058 | 310.551863 | 4.71E-26 | 4.81E-24 | F | F | T | F |
| Nynrin | -4.990984 | 6.95667849 | 310.262597 | 4.82E-26 | 4.91E-24 | F | F | T | F |
| Errfi1 | 3.43282245 | 7.27800672 | 310.108795 | 4.89E-26 | 4.95E-24 | T | F | T | T |
| Rhobtb3 | -6.7836803 | 6.04554199 | 309.660357 | 5.08E-26 | 5.12E-24 | T | F | T | F |
| Crmp1 | -8.7522193 | 5.68175652 | 309.582789 | 5.11E-26 | 5.13E-24 | F | F | F | F |
| Fras1 | -6.2404359 | 6.23665957 | 308.358272 | 5.67E-26 | 5.67E-24 | F | F | F | F |
| Zic2 | -6.8866462 | 6.46708597 | 308.061331 | 5.82E-26 | 5.79E-24 | F | F | F | F |
| Notch3 | -6.5626634 | 6.1013057 | 307.633009 | 6.03E-26 | 5.97E-24 | F | F | F | F |
| Pcdh18 | -9.7566924 | 5.65742998 | 306.856974 | 6.45E-26 | 6.35E-24 | T | F | T | T |
| Cxcl12 | -8.0476064 | 5.74648107 | 306.524355 | 6.63E-26 | 6.51E-24 | T | F | T | T |
| Spink1 | -6.944158 | 5.94515018 | 303.519476 | 8.59E-26 | 8.39E-24 | F | F | F | T |
| Nf1 | -5.7309673 | 6.46317668 | 303.291909 | 8.76E-26 | 8.52E-24 | T | F | F | F |
| Prag1 | -7.2661764 | 5.87470937 | 303.056072 | 8.94E-26 | 8.65E-24 | F | F | F | F |
| Hmga1b | -3.4300794 | 10.0083927 | 301.788372 | 9.97E-26 | 9.61E-24 | F | F | F | T |
| Rbpms2 | -5.3482803 | 6.64022425 | 300.687111 | 1.10E-25 | 1.05E-23 | T | F | T | F |
| Bcor | 2.97914217 | 8.78679433 | 300.643758 | 1.10E-25 | 1.05E-23 | T | F | F | F |
| Esyt1 | 3.91471057 | 6.28965293 | 298.390834 | 1.34E-25 | 1.28E-23 | T | F | F | T |
| Epb41l3 | -5.4440946 | 6.57977796 | 298.242119 | 1.36E-25 | 1.29E-23 | F | F | F | F |
| Zscan10 | -7.3619338 | 6.04343471 | 298.200405 | 1.36E-25 | 1.29E-23 | F | F | F | F |
| Nr2f2 | 4.61518669 | 5.91790929 | 297.452802 | 1.46E-25 | 1.37E-23 | T | F | T | F |
| Macroh2a2 | -6.721827 | 6.30213108 | 296.995026 | 1.52E-25 | 1.42E-23 | F | F | T | F |
| Ccdc184 | 5.72693896 | 4.646259 | 296.933616 | 1.52E-25 | 1.42E-23 | F | F | F | F |
| H1f10 | -6.043682 | 6.36110282 | 296.258411 | 1.62E-25 | 1.50E-23 | T | F | F | T |
| Notch1 | -4.5344742 | 7.21071106 | 295.647925 | 1.71E-25 | 1.57E-23 | T | F | F | F |
| Ror1 | -8.3941622 | 5.69002375 | 294.696266 | 1.86E-25 | 1.71E-23 | F | F | F | F |
| C530008M | -6.4450056 | 6.01891051 | 294.578386 | 1.88E-25 | 1.72E-23 | T | F | F | T |
| Frk | 5.22969827 | 5.3982055 | 293.253991 | 2.11E-25 | 1.92E-23 | T | F | T | T |
| Slc39a8 | -5.8746816 | 6.24064786 | 291.825792 | 2.40E-25 | 2.17E-23 | T | F | F | T |
| Tns1 | 3.58716817 | 6.9637412 | 291.154702 | 2.54E-25 | 2.30E-23 | T | T | T | T |
| Mbnl2 | 3.86934246 | 6.25608294 | 290.538124 | 2.69E-25 | 2.42E-23 | T | F | F | T |
| Slit3 | -8.2587264 | 5.55500243 | 289.622949 | 2.92E-25 | 2.61E-23 | F | F | T | T |
| Cachd1 | -4.8023073 | 7.14025791 | 289.119459 | 3.05E-25 | 2.72E-23 | T | F | F | F |
| Bin2 | 5.00331391 | 4.99513582 | 289.003534 | 3.08E-25 | 2.74E-23 | F | F | F | T |
| Tmprss2 | -5.8300825 | 6.23870679 | 288.478127 | 3.23E-25 | 2.86E-23 | T | F | T | F |
| Gli3 | -4.7404435 | 6.99427051 | 288.384066 | 3.26E-25 | 2.87E-23 | T | F | F | T |
| Myo1c | 3.19010276 | 8.08247698 | 287.998603 | 3.38E-25 | 2.96E-23 | T | F | T | T |
| Hmgb3 | -5.3712922 | 6.50799721 | 287.023594 | 3.69E-25 | 3.22E-23 | F | F | T | F |
| Tspan7 | -12.889403 | 5.28854265 | 286.978623 | 3.70E-25 | 3.22E-23 | T | T | T | T |

| | | | | | | | | | |
|---|---|---|---|---|---|---|---|---|---|
| Dgkk | -7.4174061 | 5.90915784 | 286.94391 | 3.71E-25 | 3.22E-23 | T | F | T | T |
| C1qtnf6 | 4.5792341 | 5.28891929 | 286.29001 | 3.94E-25 | 3.40E-23 | T | F | T | T |
| Stard8 | -5.148073 | 6.63906123 | 285.702669 | 4.15E-25 | 3.57E-23 | T | F | F | T |
| Rragd | -5.6487321 | 6.38978199 | 284.281087 | 4.73E-25 | 4.05E-23 | T | F | T | T |
| Pramef12 | 5.38149071 | 4.73945909 | 284.157049 | 4.78E-25 | 4.08E-23 | F | F | T | F |
| Fzd7 | -4.3132218 | 7.35409104 | 283.897456 | 4.90E-25 | 4.16E-23 | T | F | T | T |
| Fasl | 7.56810391 | 4.1914074 | 283.538274 | 5.06E-25 | 4.28E-23 | F | F | T | F |
| Psd4 | 4.73826522 | 5.47506239 | 283.12237 | 5.25E-25 | 4.43E-23 | T | F | F | T |
| Pard6g | 3.48154834 | 6.91265584 | 280.946394 | 6.42E-25 | 5.39E-23 | F | F | T | F |
| Pkp2 | 3.20305382 | 7.40844336 | 279.741956 | 7.17E-25 | 6.00E-23 | T | F | F | T |
| Mmp9 | 5.27524167 | 6.56186722 | 277.591525 | 8.75E-25 | 7.30E-23 | F | F | F | F |
| Plau | 5.8627892 | 4.48993755 | 276.972539 | 9.27E-25 | 7.70E-23 | T | F | F | F |
| Apln | -6.8644235 | 5.86637063 | 276.898204 | 9.33E-25 | 7.72E-23 | T | F | T | T |
| Gsn | 3.11738073 | 7.75588954 | 275.539737 | 1.06E-24 | 8.73E-23 | F | F | F | F |
| Upp1 | 3.85575693 | 6.08403624 | 274.724117 | 1.14E-24 | 9.39E-23 | F | F | F | T |
| Prl5a1 | 6.74454568 | 4.15427253 | 273.713593 | 1.26E-24 | 1.03E-22 | F | F | F | F |
| Kctd17 | 3.48593028 | 6.76976303 | 273.49227 | 1.28E-24 | 1.05E-22 | F | F | F | F |
| Psap | 2.82857616 | 8.90768902 | 273.126257 | 1.33E-24 | 1.08E-22 | F | F | T | F |
| Mogat2 | -5.2878858 | 6.42686425 | 271.919633 | 1.49E-24 | 1.20E-22 | T | F | F | T |
| Tinagl1 | 4.39991639 | 6.02653562 | 271.373326 | 1.57E-24 | 1.26E-22 | F | F | T | F |
| Nxf7 | 3.25097762 | 7.05909765 | 271.199019 | 1.59E-24 | 1.28E-22 | F | F | F | F |
| Phc1 | -4.5304763 | 7.10956585 | 271.028175 | 1.62E-24 | 1.30E-22 | F | F | F | F |
| Nnat | -8.6518995 | 5.38692883 | 270.596166 | 1.69E-24 | 1.34E-22 | F | F | F | F |
| H2bu2 | -7.2655077 | 5.54030832 | 270.42656 | 1.71E-24 | 1.36E-22 | F | F | F | F |
| Il1rap | 5.46559822 | 4.57476847 | 269.754769 | 1.83E-24 | 1.44E-22 | T | T | T | T |
| Lyn | 4.23244692 | 5.55919417 | 269.749935 | 1.83E-24 | 1.44E-22 | T | T | F | T |
| Stk10 | 3.46153318 | 6.70822146 | 269.71881 | 1.83E-24 | 1.44E-22 | T | F | F | T |
| Frzb | -12.830851 | 5.23061167 | 269.391004 | 1.89E-24 | 1.48E-22 | T | F | T | T |
| Dysf | 3.73447174 | 6.45712025 | 269.105867 | 1.94E-24 | 1.52E-22 | F | F | F | F |
| Zfp629 | -6.6850809 | 5.68813471 | 269.001978 | 1.96E-24 | 1.53E-22 | T | F | T | F |
| Gab2 | 4.91212698 | 5.16635151 | 268.700913 | 2.02E-24 | 1.56E-22 | F | F | T | T |
| Anp32a | -4.2569206 | 7.1524194 | 267.877271 | 2.19E-24 | 1.69E-22 | F | F | T | T |
| Nos2 | 7.12957412 | 4.0407021 | 267.478202 | 2.27E-24 | 1.75E-22 | T | F | F | F |
| Msx1 | -8.7985066 | 5.53240628 | 266.360375 | 2.53E-24 | 1.94E-22 | F | F | F | F |
| Trap1a | 3.85848201 | 6.02563278 | 264.661947 | 2.98E-24 | 2.27E-22 | F | F | F | F |
| Sez6l2 | 3.90661502 | 5.87926869 | 264.534766 | 3.01E-24 | 2.29E-22 | F | F | F | F |
| Mttp | -4.8413948 | 6.6539652 | 264.400219 | 3.05E-24 | 2.31E-22 | F | F | T | F |
| Sp8 | -8.8409071 | 5.57601002 | 264.02122 | 3.17E-24 | 2.39E-22 | F | F | T | F |
| Tmem132a | -4.2118099 | 7.17346496 | 263.61923 | 3.29E-24 | 2.48E-22 | F | F | F | F |
| Apoa4 | -6.0617771 | 6.12905881 | 262.793394 | 3.57E-24 | 2.68E-22 | F | F | F | F |
| Klhl22 | 3.31694542 | 7.08067114 | 262.392517 | 3.71E-24 | 2.77E-22 | T | F | F | T |
| Col17a1 | 7.55477742 | 3.93575109 | 261.853521 | 3.91E-24 | 2.91E-22 | T | F | F | F |

| | | | | | | | | | |
|---|---|---|---|---|---|---|---|---|---|
| Trpv2 | 5.17740013 | 4.66080809 | 261.513415 | 4.04E-24 | 2.99E-22 | F | F | F | T |
| Celsr2 | -7.0367471 | 5.53074512 | 261.511566 | 4.04E-24 | 2.99E-22 | F | F | F | F |
| Sox3 | -7.4695762 | 5.4848099 | 261.430241 | 4.07E-24 | 3.00E-22 | F | F | F | F |
| Prps1 | -6.9639648 | 5.52444994 | 260.960775 | 4.27E-24 | 3.13E-22 | T | F | T | T |
| Fam83b | 4.72991225 | 5.15817148 | 260.648991 | 4.40E-24 | 3.22E-22 | F | F | T | T |
| Pclaf | -4.5620961 | 6.828999 | 260.143048 | 4.62E-24 | 3.37E-22 | F | F | T | F |
| Amn | -5.8644123 | 5.93416225 | 259.852805 | 4.75E-24 | 3.46E-22 | F | F | T | F |
| Irx2 | 3.87107562 | 5.88864022 | 259.459659 | 4.94E-24 | 3.57E-22 | T | F | T | F |
| Stox2 | -6.3023773 | 5.72044149 | 259.456429 | 4.94E-24 | 3.57E-22 | T | F | T | T |
| Ly75 | -5.5770502 | 6.0624325 | 259.399964 | 4.97E-24 | 3.58E-22 | T | F | T | T |
| Sox4 | -4.084768 | 7.26677412 | 258.944501 | 5.20E-24 | 3.73E-22 | F | F | T | T |
| Hnf4a | -5.6230154 | 6.00163907 | 257.257701 | 6.14E-24 | 4.39E-22 | T | F | F | T |
| Hipk2 | -4.3207882 | 6.94428214 | 256.722368 | 6.47E-24 | 4.62E-22 | T | F | F | T |
| Shroom1 | 5.55957471 | 4.50566544 | 256.094916 | 6.89E-24 | 4.90E-22 | T | F | T | T |
| Pcdh7 | -7.9730671 | 5.27110263 | 255.783208 | 7.10E-24 | 5.03E-22 | T | F | T | T |
| Dsp | 2.60678465 | 9.56551005 | 255.762578 | 7.12E-24 | 5.03E-22 | F | F | F | T |
| Foxf1 | -8.512823 | 5.2486647 | 254.357465 | 8.18E-24 | 5.76E-22 | F | F | F | T |
| Slc30a2 | 4.81125414 | 5.12475475 | 254.159953 | 8.35E-24 | 5.86E-22 | F | F | F | F |
| Irx5 | -8.4747009 | 5.21132116 | 253.962164 | 8.51E-24 | 5.96E-22 | F | F | F | F |
| Vgll3 | 4.92869286 | 4.94864498 | 253.014157 | 9.36E-24 | 6.53E-22 | T | F | F | T |
| Arhgdib | 5.34280181 | 4.55246024 | 252.749846 | 9.61E-24 | 6.68E-22 | F | F | T | T |
| Adgrg6 | 5.95663184 | 4.23683891 | 252.436549 | 9.92E-24 | 6.87E-22 | T | F | T | T |
| Zc3hav1 | 3.29106892 | 6.81868514 | 252.031549 | 1.03E-23 | 7.13E-22 | T | F | T | F |
| Phf6 | -6.0169147 | 5.78323671 | 251.546369 | 1.08E-23 | 7.47E-22 | T | F | F | F |
| Bex2 | -7.1960458 | 5.42005401 | 251.38958 | 1.10E-23 | 7.56E-22 | F | F | F | F |
| Elf3 | 3.92725289 | 6.87668876 | 250.466731 | 1.21E-23 | 8.27E-22 | T | F | T | F |
| Tbx3 | -5.1582514 | 6.22383755 | 250.034457 | 1.26E-23 | 8.61E-22 | F | F | T | F |
| Lima1 | 3.02973428 | 7.58859674 | 249.47248 | 1.34E-23 | 9.07E-22 | F | F | T | F |
| Ctsh | -12.630615 | 5.0308068 | 249.461673 | 1.34E-23 | 9.07E-22 | F | F | T | F |
| Myo1d | 3.55306326 | 6.34464905 | 249.384762 | 1.35E-23 | 9.11E-22 | T | F | T | F |
| Sbk1 | -4.7212423 | 6.51581959 | 248.985916 | 1.40E-23 | 9.46E-22 | T | F | T | T |
| Lin28b | -5.1277805 | 6.19893375 | 248.553399 | 1.47E-23 | 9.86E-22 | T | T | T | T |
| Otx2 | -7.9822415 | 5.28049967 | 248.355755 | 1.50E-23 | 1.00E-21 | T | F | T | T |
| Sp5 | -5.751584 | 5.82277511 | 248.123159 | 1.53E-23 | 1.02E-21 | F | F | F | F |
| Pdzd4 | -6.4349384 | 5.81441619 | 248.115063 | 1.53E-23 | 1.02E-21 | F | F | F | F |
| Mgat4b | 3.12695353 | 7.15296073 | 247.889357 | 1.57E-23 | 1.04E-21 | F | F | F | F |
| Hoxb1 | -7.8364822 | 5.51163919 | 246.980312 | 1.72E-23 | 1.14E-21 | F | F | T | F |
| Igfbp4 | -4.3383722 | 7.02588393 | 246.822587 | 1.75E-23 | 1.15E-21 | F | F | T | T |
| Tro | -5.8784619 | 5.7277898 | 246.54948 | 1.80E-23 | 1.18E-21 | F | F | F | F |
| Ralbp1 | 2.86718977 | 7.88242255 | 246.020446 | 1.90E-23 | 1.25E-21 | F | F | T | F |
| Selenop | -4.7492449 | 6.45717667 | 245.973747 | 1.91E-23 | 1.25E-21 | F | F | F | F |
| Rarg | -5.3138404 | 6.03857341 | 245.90155 | 1.92E-23 | 1.25E-21 | F | F | T | F |

| | | | | | | | | | |
|---|---|---|---|---|---|---|---|---|---|
| Dusp4 | -5.7948021 | 6.17079097 | 244.734989 | 2.17E-23 | 1.41E-21 | F | F | T | F |
| Bcl9 | -4.0934638 | 7.32299385 | 244.642857 | 2.19E-23 | 1.42E-21 | F | F | T | F |
| Slc7a5 | 2.71867232 | 8.5407927 | 244.420241 | 2.24E-23 | 1.45E-21 | F | F | T | F |
| Irs1 | -5.8644432 | 5.71383071 | 244.212692 | 2.29E-23 | 1.47E-21 | T | F | T | T |
| Smarca1 | -7.879685 | 5.17821865 | 244.020948 | 2.33E-23 | 1.50E-21 | T | T | F | T |
| Nkd1 | -5.8393405 | 5.71500699 | 243.927161 | 2.36E-23 | 1.51E-21 | F | F | T | F |
| Smad3 | -5.2117237 | 6.06283403 | 242.41763 | 2.76E-23 | 1.76E-21 | T | F | T | T |
| Akt1 | 2.96817891 | 8.31109025 | 241.862357 | 2.92E-23 | 1.86E-21 | T | F | F | F |
| Bex4 | -4.5660462 | 6.56265128 | 241.690135 | 2.97E-23 | 1.88E-21 | F | F | F | F |
| Eml1 | -12.59847 | 4.99870306 | 239.515006 | 3.73E-23 | 2.36E-21 | F | F | F | T |
| Col2a1 | -8.437731 | 5.17395584 | 239.393467 | 3.78E-23 | 2.38E-21 | T | F | T | F |
| Litaf | 3.23326866 | 6.69922208 | 239.334271 | 3.80E-23 | 2.39E-21 | T | F | T | T |
| Pfkp | 3.68455899 | 5.99786245 | 237.761967 | 4.48E-23 | 2.81E-21 | F | F | F | F |
| Prex2 | 3.47134697 | 6.3361798 | 237.579017 | 4.57E-23 | 2.85E-21 | F | F | T | T |
| Slco4a1 | 12.0886064 | 3.4780888 | 240.745847 | 4.88E-23 | 3.04E-21 | T | F | F | T |
| Specc1 | 3.31720883 | 6.47881737 | 235.66185 | 5.59E-23 | 3.48E-21 | T | T | F | T |
| Tead2 | -3.3945919 | 8.33291911 | 235.378211 | 5.77E-23 | 3.57E-21 | F | F | T | F |
| Celsr1 | -4.3941686 | 6.70037159 | 235.018797 | 5.99E-23 | 3.70E-21 | T | F | T | F |
| Vim | -4.5983769 | 6.93294177 | 234.900472 | 6.07E-23 | 3.74E-21 | F | F | F | F |
| Nefm | -7.8300995 | 5.12879619 | 234.392369 | 6.40E-23 | 3.93E-21 | F | F | F | F |
| Prtg | -3.1283654 | 9.79263636 | 233.930151 | 6.73E-23 | 4.12E-21 | T | F | T | F |
| Foxc2 | -7.8062929 | 5.1053014 | 233.817279 | 6.81E-23 | 4.16E-21 | F | F | F | F |
| Mtcl1 | -3.9647582 | 7.22050241 | 233.743262 | 6.86E-23 | 4.18E-21 | F | F | F | T |
| Sall4 | -3.0251255 | 9.69157181 | 233.064228 | 7.38E-23 | 4.48E-21 | T | F | T | T |
| Mrc2 | -5.5845491 | 5.78790105 | 232.985278 | 7.44E-23 | 4.51E-21 | F | F | F | F |
| Fxyd5 | 5.46732025 | 4.3586321 | 232.460768 | 7.87E-23 | 4.75E-21 | T | F | F | T |
| Ifitm1 | -9.3374966 | 5.14250911 | 232.191553 | 8.10E-23 | 4.88E-21 | F | F | F | F |
| Nostrin | 4.52876863 | 5.11350419 | 231.43655 | 8.78E-23 | 5.27E-21 | F | F | F | F |
| Zfp266 | -3.8977402 | 7.43956197 | 231.419579 | 8.80E-23 | 5.27E-21 | F | F | T | T |
| Satb2 | 3.49929472 | 6.14080107 | 231.34981 | 8.87E-23 | 5.30E-21 | T | T | T | T |
| Tgif1 | -5.4664139 | 5.95245665 | 231.19815 | 9.01E-23 | 5.37E-21 | T | F | T | F |
| Ank3 | -5.9862265 | 5.50900863 | 230.982272 | 9.22E-23 | 5.48E-21 | T | T | T | T |
| Rgl3 | 12.040283 | 3.42972859 | 234.135155 | 9.72E-23 | 5.76E-21 | F | F | T | F |
| Npl | -8.3459852 | 5.08290454 | 230.458027 | 9.76E-23 | 5.77E-21 | F | F | F | F |
| Ltbp1 | -9.1457717 | 4.95253275 | 230.319496 | 9.91E-23 | 5.84E-21 | F | F | T | T |
| Anxa5 | 3.51992203 | 6.14162474 | 230.154174 | 1.01E-22 | 5.93E-21 | F | F | F | F |
| Foxc1 | -6.5220576 | 5.37634351 | 229.716043 | 1.06E-22 | 6.20E-21 | F | F | T | F |
| Snx9 | 2.90903597 | 7.38034965 | 229.099963 | 1.13E-22 | 6.61E-21 | F | F | T | F |
| Csgalnact1 | 6.39777567 | 4.08135637 | 228.942333 | 1.15E-22 | 6.70E-21 | T | T | T | T |
| Nt5dc2 | -3.4169128 | 8.07158037 | 228.887825 | 1.16E-22 | 6.72E-21 | F | F | F | F |
| Fgfbp1 | 4.66048384 | 4.85975088 | 228.08061 | 1.26E-22 | 7.32E-21 | F | F | F | F |
| Kif5c | -5.6427047 | 5.62771412 | 227.630103 | 1.33E-22 | 7.67E-21 | F | F | T | T |

| | | | | | | | | | |
|---|---|---|---|---|---|---|---|---|---|
| Igdcc3 | -5.5282893 | 5.9077455 | 227.069185 | 1.41E-22 | 8.13E-21 | F | F | F | F |
| Hmgn5 | 3.31136632 | 6.3521896 | 226.40716 | 1.52E-22 | 8.72E-21 | F | F | T | T |
| Fabp3 | 4.55936144 | 4.75093959 | 225.308677 | 1.71E-22 | 9.81E-21 | F | F | T | F |
| Fam199x | -5.1200677 | 5.93261651 | 225.220313 | 1.73E-22 | 9.88E-21 | T | F | T | T |
| Gm17067 | 4.3708526 | 5.04369872 | 224.724793 | 1.82E-22 | 1.04E-20 | F | F | F | T |
| Cldn6 | -3.7920592 | 7.09964548 | 224.438004 | 1.88E-22 | 1.07E-20 | T | F | F | T |
| Crybg3 | 2.89143196 | 7.18832731 | 224.102042 | 1.95E-22 | 1.11E-20 | F | F | F | F |
| Ptpn13 | -3.6473979 | 7.38822094 | 223.915828 | 1.99E-22 | 1.13E-20 | T | F | F | F |
| Krt8 | 2.48197656 | 9.93151482 | 223.789278 | 2.02E-22 | 1.14E-20 | F | F | F | F |
| Bcam | -5.5296626 | 5.83762032 | 223.659819 | 2.05E-22 | 1.15E-20 | T | F | F | T |
| Btbd2 | -4.4628813 | 6.40863764 | 223.563777 | 2.07E-22 | 1.16E-20 | T | F | F | F |
| Ggta1 | 3.4396536 | 6.12843143 | 223.321062 | 2.13E-22 | 1.19E-20 | F | F | T | F |
| Foxb1 | -8.1772226 | 5.03574126 | 223.118573 | 2.18E-22 | 1.22E-20 | F | T | T | T |
| Adamts4 | 3.75713463 | 5.52116359 | 222.910379 | 2.23E-22 | 1.24E-20 | F | F | F | F |
| Wasf1 | -6.6022273 | 5.28941677 | 222.243363 | 2.40E-22 | 1.33E-20 | T | F | F | T |
| Mt2 | 3.17314677 | 6.89231205 | 221.593004 | 2.58E-22 | 1.43E-20 | F | F | F | F |
| Tubb4a | -7.7028943 | 5.00279725 | 221.587434 | 2.58E-22 | 1.43E-20 | F | F | T | F |
| Sox2 | -8.3151006 | 5.05213784 | 221.573316 | 2.58E-22 | 1.43E-20 | T | F | F | F |
| Trp53i11 | -5.1218866 | 5.83386938 | 221.092406 | 2.73E-22 | 1.50E-20 | T | F | T | T |
| B3gnt2 | 3.73690355 | 5.57168779 | 220.497862 | 2.91E-22 | 1.60E-20 | F | F | T | F |
| Wipf3 | 3.53668201 | 5.97555245 | 219.517278 | 3.25E-22 | 1.78E-20 | F | F | T | F |
| Rnd3 | -5.6575197 | 5.50960395 | 219.304203 | 3.33E-22 | 1.82E-20 | F | F | T | F |
| Ralgds | 3.13037772 | 6.75042171 | 219.181007 | 3.38E-22 | 1.84E-20 | T | F | T | F |
| Myh9 | 2.10143792 | 11.4351241 | 219.010862 | 3.44E-22 | 1.87E-20 | T | F | T | F |
| Smad7 | 2.81795548 | 7.33173697 | 218.560411 | 3.62E-22 | 1.96E-20 | T | F | T | F |
| Efna5 | -7.6825322 | 4.98276461 | 217.985386 | 3.86E-22 | 2.09E-20 | T | F | T | T |
| Adcy6 | -6.1389553 | 5.30172148 | 217.341881 | 4.15E-22 | 2.24E-20 | F | F | T | F |
| Myrf | -4.3706611 | 6.38589873 | 217.304167 | 4.17E-22 | 2.24E-20 | F | F | T | T |
| Hmga1 | -3.377814 | 7.90592717 | 217.178941 | 4.23E-22 | 2.27E-20 | F | F | F | F |
| Rsad2 | 6.75446624 | 3.78978913 | 217.047528 | 4.29E-22 | 2.30E-20 | T | F | T | T |
| Trf | -5.3905525 | 5.68682163 | 216.772495 | 4.43E-22 | 2.36E-20 | F | F | F | T |
| Fosl1 | 6.67249148 | 4.32907504 | 216.736927 | 4.45E-22 | 2.37E-20 | F | F | F | F |
| Fam83f | 4.72014785 | 4.87958888 | 216.649924 | 4.49E-22 | 2.39E-20 | T | F | T | T |
| Smpdl3b | 4.67541437 | 4.60028558 | 216.582742 | 4.52E-22 | 2.40E-20 | F | F | F | F |
| Tdh | -6.8775657 | 5.15448726 | 216.124844 | 4.77E-22 | 2.52E-20 | T | F | F | F |
| Dpp4 | -5.9444745 | 5.43459493 | 215.967005 | 4.85E-22 | 2.56E-20 | T | T | T | T |
| Mpzl1 | 2.76296518 | 7.77333696 | 215.862733 | 4.91E-22 | 2.58E-20 | T | F | T | F |
| Nhsl2 | -6.8892593 | 5.10718533 | 215.726257 | 4.99E-22 | 2.62E-20 | F | F | T | T |
| Lpar1 | -5.3877447 | 5.65487187 | 215.38249 | 5.18E-22 | 2.71E-20 | F | F | T | F |
| Mfsd2a | 4.47377237 | 4.72120762 | 215.371747 | 5.19E-22 | 2.71E-20 | F | F | F | F |
| Syna | 11.9295301 | 3.31889858 | 218.622335 | 5.20E-22 | 2.71E-20 | F | F | F | T |
| Zcchc3 | -4.0492878 | 6.68055449 | 215.203137 | 5.29E-22 | 2.75E-20 | F | F | F | F |

| | | | | | | | | | |
|---|---|---|---|---|---|---|---|---|---|
| Gm2381 | 5.87731648 | 4.01887718 | 215.073348 | 5.37E-22 | 2.79E-20 | F | F | F | T |
| Mgat5 | 3.14236061 | 6.53039432 | 214.978873 | 5.43E-22 | 2.81E-20 | T | F | F | T |
| Car4 | -3.4534072 | 7.67022303 | 214.65473 | 5.63E-22 | 2.91E-20 | F | F | F | F |
| Adgra2 | -6.2813148 | 5.86486341 | 213.476138 | 6.45E-22 | 3.32E-20 | F | F | F | F |
| Fhl1 | -4.6710578 | 6.08548728 | 213.394601 | 6.51E-22 | 3.34E-20 | F | F | T | F |
| Hic2 | 2.58009672 | 8.24784261 | 212.926544 | 6.86E-22 | 3.52E-20 | T | F | T | T |
| Fzd10 | -6.7111616 | 5.39612038 | 212.635136 | 7.10E-22 | 3.62E-20 | T | F | T | T |
| A4galt | 3.24906826 | 6.35928865 | 212.623128 | 7.11E-22 | 3.62E-20 | F | F | F | F |
| Tmem47 | -6.1700042 | 5.31465265 | 212.612116 | 7.12E-22 | 3.62E-20 | F | F | T | F |
| Cd47 | 3.630547 | 5.74762006 | 212.355056 | 7.33E-22 | 3.72E-20 | T | F | T | T |
| Id3 | -4.5211707 | 6.35674694 | 212.068518 | 7.58E-22 | 3.84E-20 | T | F | T | T |
| Crb2 | -5.3804902 | 5.70598541 | 211.81703 | 7.80E-22 | 3.94E-20 | F | F | T | F |
| Col4a1 | -3.0264929 | 9.02144043 | 211.698575 | 7.91E-22 | 3.99E-20 | T | F | T | T |
| Mmp2 | -7.1919451 | 5.20895641 | 211.544742 | 8.05E-22 | 4.05E-20 | F | F | T | F |
| Bahcc1 | -3.4525965 | 7.62015929 | 211.380101 | 8.20E-22 | 4.12E-20 | T | F | F | T |
| Apom | -6.6363268 | 5.3220686 | 211.235765 | 8.34E-22 | 4.18E-20 | F | F | F | F |
| Kif7 | -7.6190235 | 4.91975942 | 211.138433 | 8.43E-22 | 4.21E-20 | F | F | F | F |
| Slc39a5 | -7.2831503 | 4.98698404 | 210.906564 | 8.66E-22 | 4.32E-20 | F | F | F | F |
| Cd200 | 6.23782096 | 3.90242142 | 209.785006 | 9.87E-22 | 4.91E-20 | T | F | T | T |
| Ldhb | -4.3683116 | 6.31152426 | 209.529755 | 1.02E-21 | 5.04E-20 | T | F | T | T |
| Zfp423 | -4.4662539 | 6.18074017 | 208.945663 | 1.09E-21 | 5.39E-20 | F | F | F | F |
| Itga5 | 2.65812233 | 8.00616506 | 208.907517 | 1.09E-21 | 5.40E-20 | T | F | F | F |
| Zdbf2 | -4.422346 | 6.45454871 | 208.243357 | 1.18E-21 | 5.82E-20 | F | T | F | T |
| Zfp521 | -7.2910127 | 4.99465116 | 207.973711 | 1.22E-21 | 5.99E-20 | T | F | T | T |
| Pkdcc | -4.2786569 | 6.33171413 | 207.573954 | 1.28E-21 | 6.26E-20 | F | F | F | F |
| Fmnl2 | -5.557107 | 5.41081762 | 207.512681 | 1.29E-21 | 6.29E-20 | T | F | T | T |
| Fgd3 | 5.84665533 | 4.46883835 | 206.962523 | 1.37E-21 | 6.70E-20 | F | F | T | T |
| Tmbim1 | 3.229801 | 6.2020988 | 206.902081 | 1.38E-21 | 6.73E-20 | F | F | T | F |
| Zfp667 | 3.87733325 | 5.17692661 | 206.355162 | 1.47E-21 | 7.16E-20 | T | F | F | T |
| Zdhhc15 | -8.0675366 | 4.80687734 | 205.847492 | 1.56E-21 | 7.59E-20 | T | F | F | T |
| Il2rg | 5.5022936 | 4.38670019 | 205.637086 | 1.60E-21 | 7.76E-20 | F | F | F | F |
| Zfp36l3 | 4.13548427 | 4.88020705 | 205.102496 | 1.71E-21 | 8.25E-20 | F | F | F | F |
| Arnt2 | -7.4910994 | 4.86671481 | 204.779137 | 1.77E-21 | 8.55E-20 | T | F | T | F |
| Prss8 | 2.75192286 | 7.18597968 | 204.040547 | 1.94E-21 | 9.31E-20 | T | F | F | F |
| Sri | 3.324934 | 6.06761581 | 203.057907 | 2.18E-21 | 1.04E-19 | F | F | F | F |
| Epb41l5 | -4.3534946 | 6.1854695 | 202.754771 | 2.26E-21 | 1.08E-19 | F | F | T | F |
| Dhrs9 | 7.20968592 | 3.69709293 | 202.741546 | 2.26E-21 | 1.08E-19 | T | F | F | F |
| Als2cl | 4.12640903 | 4.90040524 | 202.691653 | 2.27E-21 | 1.08E-19 | T | F | F | F |
| Trib3 | 4.65938645 | 4.43373867 | 202.388814 | 2.36E-21 | 1.12E-19 | F | F | F | F |
| Sept6 | -5.6088013 | 5.2934741 | 201.699616 | 2.56E-21 | 1.21E-19 | T | F | T | T |
| Dusp6 | -6.7329113 | 5.22857695 | 201.249091 | 2.70E-21 | 1.28E-19 | T | F | F | T |
| Atp2c2 | 4.53050119 | 4.63415948 | 200.962774 | 2.80E-21 | 1.32E-19 | F | F | F | F |

| | | | | | | | | | |
|---|---|---|---|---|---|---|---|---|---|
| Cd9 | 3.15693278 | 6.25089212 | 200.588637 | 2.92E-21 | 1.38E-19 | T | T | T | T |
| Rspo3 | -8.9310074 | 4.7391857 | 200.467468 | 2.97E-21 | 1.39E-19 | T | F | T | F |
| H2-D1 | 3.2712778 | 6.18170206 | 200.454281 | 2.97E-21 | 1.39E-19 | F | F | T | F |
| Nfasc | -8.9045966 | 4.71348386 | 200.277771 | 3.04E-21 | 1.42E-19 | F | F | T | F |
| Mmp28 | 6.22053394 | 3.81734034 | 200.227664 | 3.05E-21 | 1.43E-19 | F | F | T | F |
| Junb | 3.6995172 | 5.44234232 | 199.774413 | 3.22E-21 | 1.50E-19 | F | F | F | F |
| Casz1 | 3.52136115 | 5.52107256 | 198.948526 | 3.56E-21 | 1.66E-19 | F | F | T | F |
| Mapk8ip2 | -8.8948248 | 4.70379573 | 198.940935 | 3.57E-21 | 1.66E-19 | F | F | F | F |
| Pitpnc1 | -7.6146849 | 4.91580914 | 198.646507 | 3.70E-21 | 1.71E-19 | T | T | T | T |
| Car7 | -6.0212977 | 5.58615993 | 197.99034 | 4.00E-21 | 1.85E-19 | F | F | F | F |
| Nrxn2 | -8.9439295 | 4.75266709 | 197.604906 | 4.19E-21 | 1.93E-19 | F | F | T | T |
| Gpc1 | -3.7741401 | 6.73103314 | 197.394513 | 4.30E-21 | 1.98E-19 | F | F | T | F |
| Igfbp5 | -4.8157228 | 6.15790939 | 197.310949 | 4.35E-21 | 2.00E-19 | T | F | T | F |
| Zswim6 | -12.140267 | 4.5441119 | 197.243592 | 4.38E-21 | 2.01E-19 | T | F | F | F |
| Zfp462 | -3.3052554 | 7.56733374 | 197.205126 | 4.40E-21 | 2.01E-19 | T | T | F | T |
| Ago1 | -4.279711 | 6.20897941 | 197.112194 | 4.45E-21 | 2.03E-19 | F | F | T | T |
| Sgpp1 | 3.52650608 | 5.47641615 | 196.984736 | 4.52E-21 | 2.06E-19 | T | T | F | T |
| Zbtb12 | -6.0904163 | 5.09929379 | 196.938857 | 4.55E-21 | 2.07E-19 | F | F | T | F |
| Lpl | 3.26465262 | 6.0325403 | 196.869113 | 4.59E-21 | 2.08E-19 | T | F | F | F |
| Dpysl5 | -3.9571114 | 6.56772257 | 196.8574 | 4.59E-21 | 2.08E-19 | F | F | T | F |
| Ina | -7.0132322 | 5.03310267 | 196.542264 | 4.77E-21 | 2.16E-19 | T | F | T | F |
| Kif1a | -7.5133901 | 5.08990831 | 196.45469 | 4.82E-21 | 2.17E-19 | F | F | F | F |
| Extl1 | 4.4901219 | 4.68216224 | 196.333091 | 4.90E-21 | 2.20E-19 | F | F | F | T |
| Gata4 | -4.424733 | 6.04868563 | 196.152527 | 5.01E-21 | 2.25E-19 | F | F | T | F |
| Apoc2 | -4.7747724 | 6.44138227 | 196.023746 | 5.09E-21 | 2.28E-19 | T | F | F | F |
| Pthlh | 8.34067428 | 3.32851655 | 195.963214 | 5.12E-21 | 2.29E-19 | F | F | F | F |
| Zeb2 | -4.7052646 | 5.75252075 | 195.747966 | 5.26E-21 | 2.35E-19 | T | F | T | F |
| Hdac7 | -12.136591 | 4.54027573 | 195.639045 | 5.33E-21 | 2.37E-19 | F | F | T | F |
| Nek6 | 3.20963788 | 6.17493195 | 195.606448 | 5.35E-21 | 2.38E-19 | F | F | T | F |
| Zfp219 | -4.373779 | 6.04567796 | 195.435901 | 5.47E-21 | 2.42E-19 | F | F | F | F |
| Zic5 | -6.8363496 | 5.21149149 | 195.134866 | 5.67E-21 | 2.51E-19 | F | F | F | T |
| Ndst1 | -4.4995564 | 5.91993365 | 194.982059 | 5.78E-21 | 2.55E-19 | T | F | T | T |
| Ergic1 | -4.0142816 | 6.44242003 | 194.962129 | 5.79E-21 | 2.55E-19 | F | F | T | T |
| Snn | -5.6206944 | 5.21332392 | 194.439417 | 6.18E-21 | 2.72E-19 | T | F | T | T |
| Emilin1 | 3.19366143 | 7.08656533 | 194.398316 | 6.21E-21 | 2.72E-19 | F | F | F | F |
| Peli2 | -7.0147547 | 4.85618442 | 194.072802 | 6.46E-21 | 2.83E-19 | T | F | T | T |
| Arid3a | 2.48841939 | 8.3813445 | 193.959321 | 6.55E-21 | 2.86E-19 | F | F | T | F |
| Pld1 | 4.39058709 | 4.60576522 | 193.827453 | 6.66E-21 | 2.91E-19 | T | F | T | T |
| Limk1 | 2.74886184 | 7.01786435 | 193.400169 | 7.02E-21 | 3.06E-19 | F | F | T | F |
| Wnt5a | -6.2861957 | 5.24525531 | 193.308391 | 7.10E-21 | 3.09E-19 | T | F | T | T |
| Rab8b | 2.6773874 | 7.25108189 | 193.222682 | 7.18E-21 | 3.11E-19 | T | F | T | F |
| Slc38a4 | 2.48091546 | 8.41386325 | 193.142021 | 7.25E-21 | 3.14E-19 | F | F | T | T |

| | | | | | | | | | |
|---|---|---|---|---|---|---|---|---|---|
| Rinl | 4.58826446 | 4.60375743 | 193.033379 | 7.35E-21 | 3.17E-19 | T | F | T | T |
| Stat1 | 3.41719085 | 5.59612955 | 192.677432 | 7.68E-21 | 3.31E-19 | T | F | T | T |
| Lrig3 | -4.6060091 | 5.7923605 | 192.657597 | 7.70E-21 | 3.31E-19 | T | F | F | F |
| Pim2 | -4.0537891 | 7.16388229 | 192.50527 | 7.84E-21 | 3.37E-19 | F | F | F | F |
| Mapk8ip1 | -6.6146563 | 4.89461129 | 192.194866 | 8.15E-21 | 3.49E-19 | F | F | T | F |
| Chek2 | -12.173884 | 4.57766918 | 191.258943 | 9.16E-21 | 3.92E-19 | F | F | F | F |
| Usp44 | -7.2033336 | 4.90859813 | 190.922059 | 9.55E-21 | 4.08E-19 | T | F | T | T |
| Fstl1 | -4.2238067 | 6.29547234 | 190.888367 | 9.59E-21 | 4.09E-19 | T | F | T | T |
| Mid1ip1 | -5.9318197 | 5.07854016 | 190.854568 | 9.63E-21 | 4.10E-19 | T | F | F | F |
| Fam89a | 4.63254182 | 4.38764171 | 190.452494 | 1.01E-20 | 4.30E-19 | F | F | F | F |
| Stag3 | 6.69780745 | 3.59667553 | 190.111111 | 1.06E-20 | 4.48E-19 | F | F | F | F |
| Egfr | 3.90549514 | 4.93742199 | 189.976204 | 1.08E-20 | 4.54E-19 | T | F | T | T |
| Map1a | -4.6992683 | 5.74323435 | 189.592036 | 1.13E-20 | 4.76E-19 | T | F | F | F |
| Flrt3 | -5.2089923 | 5.47886489 | 189.459606 | 1.15E-20 | 4.83E-19 | T | F | F | F |
| Rhox6 | 4.77584334 | 4.2497128 | 189.336845 | 1.16E-20 | 4.89E-19 | F | F | T | F |
| Ripor1 | 3.01001942 | 6.53125291 | 189.265175 | 1.18E-20 | 4.93E-19 | T | F | F | F |
| Capg | 3.27037024 | 5.82801586 | 189.073431 | 1.20E-20 | 5.04E-19 | F | F | T | F |
| Ppp1r13b | 3.22869062 | 6.28999335 | 189.056237 | 1.21E-20 | 5.04E-19 | F | F | F | F |
| Unc5c | -8.1295748 | 4.91340527 | 189.020415 | 1.21E-20 | 5.05E-19 | T | F | T | T |
| Igsf8 | 3.07329887 | 6.51143435 | 188.943703 | 1.22E-20 | 5.09E-19 | F | F | F | F |
| Asns | 2.63775859 | 7.27926859 | 188.413371 | 1.31E-20 | 5.43E-19 | F | F | F | T |
| Inhbb | 6.78640852 | 3.56229918 | 187.958128 | 1.39E-20 | 5.75E-19 | F | F | F | F |
| Cdh11 | -6.7736569 | 5.02260497 | 186.759262 | 1.61E-20 | 6.67E-19 | F | F | F | F |
| Agrn | -2.9757425 | 8.18854614 | 186.50618 | 1.67E-20 | 6.88E-19 | F | F | F | T |
| Plch1 | -6.2530983 | 4.94393943 | 186.329973 | 1.70E-20 | 7.02E-19 | T | F | T | T |
| Gys1 | 2.7077624 | 6.93776485 | 184.764445 | 2.08E-20 | 8.56E-19 | T | F | F | T |
| Vcan | -5.1025078 | 6.77297908 | 184.028714 | 2.29E-20 | 9.38E-19 | T | F | T | F |
| Prdm6 | -6.9931767 | 4.78156508 | 183.816117 | 2.35E-20 | 9.63E-19 | T | F | F | F |
| Svil | 2.90537055 | 6.46482506 | 183.50222 | 2.44E-20 | 1.00E-18 | T | T | F | T |
| Rai1 | -3.432304 | 6.93897796 | 183.132358 | 2.56E-20 | 1.05E-18 | T | F | F | T |
| Arid5b | 2.66455212 | 7.00138074 | 183.044338 | 2.59E-20 | 1.06E-18 | T | F | T | T |
| Cited1 | -4.8187958 | 5.82758737 | 183.034519 | 2.60E-20 | 1.06E-18 | F | F | F | F |
| Fkbp10 | -6.5340005 | 4.81479516 | 182.771293 | 2.69E-20 | 1.09E-18 | F | F | T | F |
| Dll3 | -5.9964014 | 5.12356492 | 182.476699 | 2.79E-20 | 1.13E-18 | F | F | F | F |
| Sfrp2 | -11.992301 | 4.39738474 | 182.415321 | 2.81E-20 | 1.14E-18 | T | F | T | F |
| Osbpl8 | 2.77418707 | 6.78240914 | 182.361552 | 2.83E-20 | 1.14E-18 | T | T | F | T |
| Fgfr1 | -2.9959087 | 8.12069308 | 181.906966 | 3.00E-20 | 1.21E-18 | F | F | F | F |
| Adgre5 | 5.31702635 | 3.95417906 | 181.567655 | 3.14E-20 | 1.26E-18 | F | F | F | F |
| Gm4737 | -11.979316 | 4.38450909 | 181.330241 | 3.24E-20 | 1.30E-18 | F | F | T | F |
| Tceal8 | -3.9315218 | 6.32327225 | 181.258352 | 3.27E-20 | 1.31E-18 | F | F | F | F |
| Basp1 | 2.32905786 | 8.56024283 | 181.066334 | 3.35E-20 | 1.34E-18 | T | F | F | F |
| Erbb2 | -5.5896619 | 5.11339152 | 180.953365 | 3.40E-20 | 1.36E-18 | F | F | T | F |

| | | | | | | | | | |
|---|---|---|---|---|---|---|---|---|---|
| Scin | 4.25410475 | 4.60610371 | 180.50803 | 3.60E-20 | 1.44E-18 | F | F | F | F |
| Zfp282 | -5.5786884 | 5.05998519 | 180.316796 | 3.69E-20 | 1.47E-18 | T | F | T | T |
| Ccser1 | 7.0330793 | 3.40085702 | 179.12412 | 4.31E-20 | 1.71E-18 | F | F | F | T |
| Slc39a14 | 2.60134221 | 7.08399786 | 179.101076 | 4.33E-20 | 1.72E-18 | T | F | T | T |
| Gm9 | 8.77163047 | 3.25441585 | 179.004597 | 4.38E-20 | 1.73E-18 | T | F | F | F |
| Ugp2 | 2.97783152 | 6.23381562 | 178.639693 | 4.60E-20 | 1.82E-18 | F | F | T | T |
| Evc2 | -5.6257465 | 5.01426543 | 177.816321 | 5.12E-20 | 2.02E-18 | T | F | F | F |
| Erf | -3.8823861 | 6.28925274 | 176.435203 | 6.15E-20 | 2.42E-18 | F | F | F | F |
| Amotl2 | -3.1539265 | 7.28567172 | 176.425332 | 6.16E-20 | 2.42E-18 | F | F | T | F |
| Pkp3 | 3.73355649 | 4.98731066 | 176.378168 | 6.19E-20 | 2.43E-18 | F | F | F | F |
| Sox9 | -6.4898314 | 4.77108071 | 176.37488 | 6.20E-20 | 2.43E-18 | F | F | F | F |
| Rasgrp3 | -4.7978647 | 5.39763775 | 176.16693 | 6.37E-20 | 2.49E-18 | T | F | T | F |
| Ezr | 2.18611065 | 9.49295269 | 176.075062 | 6.45E-20 | 2.51E-18 | F | F | T | F |
| B3gnt3 | 4.24388914 | 4.57625548 | 176.043363 | 6.48E-20 | 2.52E-18 | F | F | T | F |
| Ankrd33b | 3.40063441 | 5.34073199 | 175.627509 | 6.84E-20 | 2.66E-18 | T | F | T | T |
| Hsd17b2 | 4.31249049 | 4.43218998 | 175.195271 | 7.25E-20 | 2.81E-18 | F | F | F | F |
| Plxna3 | -7.0756904 | 4.72185562 | 175.10935 | 7.33E-20 | 2.84E-18 | T | F | T | F |
| Tnfrsf1b | 3.68955848 | 4.98402277 | 174.953194 | 7.48E-20 | 2.89E-18 | T | F | T | F |
| Arrb1 | -3.500165 | 6.91998013 | 174.746297 | 7.69E-20 | 2.97E-18 | T | F | T | T |
| Evx1 | -7.2837034 | 4.58746609 | 174.634709 | 7.81E-20 | 3.01E-18 | T | F | T | F |
| Il13ra1 | 4.96210208 | 4.08554776 | 174.28288 | 8.18E-20 | 3.15E-18 | T | F | F | T |
| Golim4 | -6.5231896 | 4.71704622 | 174.237775 | 8.23E-20 | 3.16E-18 | T | F | T | T |
| Gpc4 | -5.5146974 | 5.36977796 | 173.684111 | 8.87E-20 | 3.40E-18 | F | F | F | T |
| Palm3 | -7.2759144 | 4.57974846 | 173.515714 | 9.07E-20 | 3.47E-18 | F | F | T | F |
| Psg22 | 5.03935544 | 4.2048677 | 172.856215 | 9.91E-20 | 3.78E-18 | T | F | F | T |
| Paqr4 | 4.10187417 | 4.73694895 | 172.6422 | 1.02E-19 | 3.89E-18 | F | F | T | F |
| Zfp12 | -7.3968382 | 4.69996631 | 172.611051 | 1.02E-19 | 3.90E-18 | T | F | T | T |
| Cntfr | -12.105471 | 4.50915801 | 172.578395 | 1.03E-19 | 3.91E-18 | F | F | T | F |
| Nacad | 5.47176651 | 3.82702903 | 172.443051 | 1.05E-19 | 3.97E-18 | F | F | F | F |
| E330009J07 | -4.8666606 | 5.31327908 | 172.308266 | 1.07E-19 | 4.04E-18 | T | F | T | F |
| Patz1 | -3.3426652 | 6.85609505 | 172.167848 | 1.09E-19 | 4.11E-18 | T | F | T | T |
| Fhod1 | 2.90200198 | 6.25061899 | 172.079513 | 1.10E-19 | 4.15E-18 | F | F | F | F |
| Rhox9 | 7.04333572 | 3.4237468 | 172.056396 | 1.10E-19 | 4.15E-18 | F | F | F | F |
| Smad6 | -5.366699 | 5.05507903 | 171.716097 | 1.16E-19 | 4.34E-18 | F | F | F | F |
| Klf6 | 2.48104812 | 7.38218519 | 171.370249 | 1.21E-19 | 4.54E-18 | T | F | F | T |
| Cd276 | -3.8404983 | 6.19257781 | 171.194062 | 1.24E-19 | 4.64E-18 | T | F | T | T |
| Ate1 | 2.55813712 | 7.03096401 | 170.79464 | 1.31E-19 | 4.89E-18 | T | F | T | T |
| Arhgap22 | 3.94036492 | 4.78695073 | 170.786245 | 1.31E-19 | 4.89E-18 | F | F | F | F |
| Jph4 | -7.4065732 | 4.70965359 | 170.662165 | 1.33E-19 | 4.97E-18 | T | F | F | F |
| Meis2 | -4.5724165 | 6.0696698 | 170.100432 | 1.44E-19 | 5.35E-18 | T | F | F | F |
| Parp4 | 3.46515354 | 5.27117624 | 169.430324 | 1.58E-19 | 5.85E-18 | F | F | F | F |
| Dnm2 | 2.35621579 | 7.92221884 | 169.058841 | 1.66E-19 | 6.14E-18 | F | F | T | F |

| | | | | | | | | | |
|---|---|---|---|---|---|---|---|---|---|
| Pdzk1 | -5.2116521 | 5.07085959 | 168.940318 | 1.68E-19 | 6.23E-18 | T | F | F | F |
| Baz2b | -4.3836441 | 5.5723094 | 168.813952 | 1.71E-19 | 6.33E-18 | T | F | F | F |
| Jak1 | 2.45367248 | 7.43420034 | 168.716331 | 1.74E-19 | 6.41E-18 | F | F | T | F |
| Parp1 | -2.6329771 | 8.86652514 | 168.347793 | 1.83E-19 | 6.73E-18 | F | F | F | T |
| Slc9a3r1 | -2.8860241 | 7.99722614 | 168.217144 | 1.86E-19 | 6.84E-18 | F | F | T | F |
| Col5a1 | -4.3018744 | 5.70006935 | 168.182024 | 1.87E-19 | 6.86E-18 | F | F | F | F |
| Slc39a4 | -6.8614345 | 4.56987592 | 168.146627 | 1.88E-19 | 6.88E-18 | F | F | T | F |
| Foxi3 | -8.7625513 | 4.57285128 | 167.786102 | 1.97E-19 | 7.22E-18 | F | F | F | F |
| Rnf144a | -5.8991079 | 4.78840256 | 167.348129 | 2.10E-19 | 7.65E-18 | T | F | T | F |
| Frem1 | -5.4142885 | 5.07114936 | 167.238843 | 2.13E-19 | 7.75E-18 | T | F | F | F |
| Pdpn | -4.2868876 | 5.62940285 | 167.228904 | 2.13E-19 | 7.75E-18 | T | F | T | F |
| Lhfpl2 | -4.3631782 | 5.58412726 | 166.887437 | 2.23E-19 | 8.11E-18 | T | T | T | T |
| Acp5 | 3.68513099 | 5.24831018 | 166.872078 | 2.24E-19 | 8.12E-18 | F | F | T | F |
| Cd164 | 2.26570432 | 8.37901398 | 166.803142 | 2.26E-19 | 8.18E-18 | T | F | T | F |
| Tspan13 | -4.869651 | 5.20514018 | 166.720282 | 2.29E-19 | 8.26E-18 | T | T | F | T |
| Tcf3 | -2.8051106 | 8.02236139 | 166.657427 | 2.31E-19 | 8.32E-18 | F | F | T | F |
| Has2 | -11.823655 | 4.23024609 | 165.699265 | 2.63E-19 | 9.48E-18 | T | T | T | T |
| Neo1 | -3.3503524 | 6.72242634 | 165.666942 | 2.64E-19 | 9.51E-18 | T | F | T | F |
| Abcc2 | -6.1659208 | 4.6958477 | 165.525789 | 2.70E-19 | 9.68E-18 | F | F | T | T |
| Dlk1 | -5.6525982 | 5.85409567 | 165.138368 | 2.85E-19 | 1.02E-17 | T | F | T | T |
| Nrp1 | -5.1887334 | 5.26618087 | 165.054572 | 2.88E-19 | 1.03E-17 | T | F | T | F |
| Fosl2 | 3.44444508 | 5.13481602 | 165.018539 | 2.89E-19 | 1.03E-17 | T | F | T | F |
| Rab34 | -6.3576441 | 4.6407118 | 164.773515 | 2.99E-19 | 1.07E-17 | F | F | F | F |
| Tmem88 | -5.6840799 | 4.8340854 | 164.021976 | 3.32E-19 | 1.18E-17 | T | F | T | T |
| Jmjd8 | -11.820147 | 4.22677715 | 163.862638 | 3.40E-19 | 1.21E-17 | F | F | T | F |
| Pmp22 | -6.3995046 | 5.08743069 | 163.671144 | 3.49E-19 | 1.24E-17 | F | F | T | F |
| Palb2 | -11.848448 | 4.25481621 | 163.410378 | 3.62E-19 | 1.28E-17 | T | F | F | F |
| Mfsd14b | 2.61888032 | 6.74237247 | 163.350283 | 3.65E-19 | 1.29E-17 | T | F | T | F |
| Fjx1 | -7.2777849 | 4.58138196 | 163.284056 | 3.69E-19 | 1.30E-17 | F | F | F | F |
| Fbln1 | -3.3827407 | 6.88131909 | 163.273357 | 3.69E-19 | 1.30E-17 | F | F | T | F |
| Vasp | 2.31429162 | 7.89927873 | 163.25122 | 3.70E-19 | 1.30E-17 | F | F | T | T |
| Cdcp1 | 2.94483687 | 5.96361343 | 163.236187 | 3.71E-19 | 1.30E-17 | T | F | T | F |
| Rbp1 | -5.0930016 | 5.35484463 | 162.81269 | 3.94E-19 | 1.38E-17 | F | F | F | F |
| Tbx20 | -4.7144275 | 5.23147588 | 162.116742 | 4.34E-19 | 1.52E-17 | T | F | T | T |
| Serpinb6b | 5.88517853 | 3.77167817 | 161.71822 | 4.59E-19 | 1.61E-17 | F | F | F | T |
| Phactr1 | 3.16728213 | 5.47955384 | 161.266519 | 4.90E-19 | 1.71E-17 | T | F | T | T |
| Dock9 | 2.74239681 | 6.51394218 | 161.116201 | 5.00E-19 | 1.74E-17 | F | F | T | F |
| Pcbp4 | -2.7528982 | 8.01860046 | 160.979287 | 5.10E-19 | 1.78E-17 | F | F | F | F |
| Ski | -3.1732623 | 6.89633412 | 160.834866 | 5.21E-19 | 1.81E-17 | T | F | T | F |
| Gsta4 | -5.6948647 | 4.84452351 | 160.462969 | 5.49E-19 | 1.90E-17 | F | F | F | F |
| Sbsn | 4.05816819 | 4.41614669 | 160.428391 | 5.51E-19 | 1.91E-17 | F | F | F | F |
| Tln1 | 2.13574202 | 9.29051146 | 160.193327 | 5.70E-19 | 1.97E-17 | F | F | F | F |

| | | | | | | | | | |
|---|---|---|---|---|---|---|---|---|---|
| Fam171a1 | 2.52119783 | 6.92108991 | 160.041547 | 5.83E-19 | 2.01E-17 | F | F | T | F |
| Dsc2 | 2.88133683 | 6.23284563 | 160.032455 | 5.83E-19 | 2.01E-17 | T | F | T | F |
| Cdh2 | -3.6177174 | 6.25188482 | 159.679396 | 6.13E-19 | 2.11E-17 | T | F | F | T |
| Antxr1 | -6.1564684 | 4.62279144 | 159.06053 | 6.70E-19 | 2.30E-17 | T | F | T | T |
| Repin1 | -4.837341 | 5.11517414 | 158.767328 | 6.99E-19 | 2.40E-17 | F | F | F | F |
| Atp7b | -4.1395906 | 5.69972865 | 158.651041 | 7.10E-19 | 2.43E-17 | F | F | F | F |
| Gbx2 | -5.5436612 | 5.47351565 | 158.579514 | 7.18E-19 | 2.45E-17 | T | T | T | T |
| Atp10a | 3.10119502 | 5.83541168 | 158.503405 | 7.26E-19 | 2.48E-17 | F | F | F | T |
| Foxj1 | -6.5855633 | 4.86665624 | 158.384102 | 7.38E-19 | 2.52E-17 | T | F | T | F |
| Grik3 | -4.3490115 | 5.91485484 | 158.35958 | 7.41E-19 | 2.52E-17 | F | F | T | T |
| Arhgap23 | 2.58480454 | 6.7342456 | 158.35012 | 7.42E-19 | 2.52E-17 | F | F | T | F |
| Zfpm1 | -3.8987617 | 5.94267554 | 158.292704 | 7.48E-19 | 2.54E-17 | F | F | T | F |
| Pknox2 | -4.8432198 | 5.17867448 | 157.850962 | 7.97E-19 | 2.70E-17 | T | F | T | F |
| Foxo4 | -3.1601689 | 6.94980884 | 157.804322 | 8.02E-19 | 2.71E-17 | F | F | T | F |
| Cdk6 | -3.8235926 | 5.91557029 | 157.037186 | 8.96E-19 | 3.03E-17 | T | F | T | T |
| Fhl3 | 2.98986893 | 5.72769689 | 156.8 | 9.27E-19 | 3.13E-17 | F | F | T | F |
| Pstpip1 | 5.9058467 | 3.55176388 | 156.663324 | 9.45E-19 | 3.18E-17 | F | F | F | F |
| Tbrg1 | 2.25717876 | 8.04676253 | 156.322484 | 9.93E-19 | 3.33E-17 | F | F | F | F |
| Rims2 | -5.6263306 | 4.82567104 | 156.320307 | 9.94E-19 | 3.33E-17 | F | F | T | F |
| Evc | -6.8073904 | 4.47236459 | 155.968304 | 1.05E-18 | 3.50E-17 | T | F | T | T |
| Ece1 | -4.2222801 | 5.96818235 | 155.921277 | 1.05E-18 | 3.52E-17 | F | F | T | F |
| Gpr161 | -11.738324 | 4.14573542 | 155.91986 | 1.05E-18 | 3.52E-17 | F | F | T | T |
| Wdr86 | -7.0744301 | 4.45460869 | 155.812446 | 1.07E-18 | 3.57E-17 | F | F | F | T |
| Nup62cl | 3.22471749 | 5.29179595 | 155.598078 | 1.10E-18 | 3.67E-17 | F | F | F | F |
| Arhgap27 | 3.10184887 | 5.51835457 | 155.487453 | 1.12E-18 | 3.73E-17 | T | F | T | T |
| Enpp3 | 3.52152688 | 5.04464908 | 155.169568 | 1.17E-18 | 3.90E-17 | T | F | F | F |
| Nedd4 | -2.2217364 | 10.4722494 | 155.133289 | 1.18E-18 | 3.91E-17 | T | F | T | T |
| Eif1ax | -3.3243066 | 6.56177998 | 154.991961 | 1.20E-18 | 3.99E-17 | F | F | T | T |
| Aldh1a2 | -8.3910047 | 6.40015737 | 154.527161 | 1.29E-18 | 4.26E-17 | F | F | F | F |
| Stmn1 | -2.8253899 | 7.5304067 | 154.4878 | 1.30E-18 | 4.28E-17 | T | F | T | T |
| Sox17 | -4.8290882 | 5.10731531 | 154.036306 | 1.38E-18 | 4.56E-17 | T | F | F | F |
| Fam83h | 2.79717662 | 6.14926462 | 153.868677 | 1.42E-18 | 4.67E-17 | F | F | T | F |
| Ddah2 | -4.5307829 | 5.22693088 | 153.638343 | 1.47E-18 | 4.82E-17 | F | F | F | T |
| Slc44a4 | 3.93496643 | 4.42028537 | 153.498699 | 1.50E-18 | 4.91E-17 | F | F | F | F |
| Myo18a | 2.21427909 | 8.14368172 | 153.474056 | 1.50E-18 | 4.93E-17 | T | F | F | T |
| Slit2 | -11.715853 | 4.12347943 | 153.280682 | 1.55E-18 | 5.06E-17 | T | F | F | T |
| Msi2 | -2.957493 | 7.08166126 | 153.185971 | 1.57E-18 | 5.12E-17 | T | T | T | T |
| Hs6st2 | -8.4040968 | 4.21756193 | 153.135529 | 1.58E-18 | 5.15E-17 | T | F | T | F |
| Crabp1 | -8.8446591 | 4.65311751 | 153.029276 | 1.61E-18 | 5.23E-17 | F | F | F | F |
| Fgd6 | 2.63770469 | 6.6151905 | 152.819737 | 1.66E-18 | 5.38E-17 | F | F | T | T |
| Dyrk2 | 2.32436104 | 7.46901416 | 152.312525 | 1.78E-18 | 5.79E-17 | T | F | F | T |
| Pparg | 6.53966938 | 3.41491961 | 152.116906 | 1.84E-18 | 5.95E-17 | F | F | F | T |

| | | | | | | | | | |
|---|---|---|---|---|---|---|---|---|---|
| Gtf2ird1 | -3.6608055 | 6.03862124 | 152.020213 | 1.86E-18 | 6.03E-17 | T | F | T | F |
| Hip1r | 2.68535692 | 6.32626287 | 151.817648 | 1.92E-18 | 6.19E-17 | F | F | F | F |
| Tgm1 | 3.19101131 | 5.35533261 | 151.815109 | 1.92E-18 | 6.19E-17 | F | F | F | T |
| Tagln2 | 2.27346122 | 7.75420496 | 151.792964 | 1.93E-18 | 6.20E-17 | T | F | F | T |
| Hdgfl3 | -4.2974511 | 5.41396476 | 151.522067 | 2.00E-18 | 6.44E-17 | T | F | T | F |
| Dchs1 | -6.097226 | 4.60099943 | 151.520537 | 2.00E-18 | 6.44E-17 | F | F | F | F |
| Zmynd11 | -3.6348066 | 6.03735949 | 151.485385 | 2.02E-18 | 6.46E-17 | T | T | T | T |
| Map3k9 | -7.649737 | 4.39345264 | 151.464624 | 2.02E-18 | 6.47E-17 | T | F | T | T |
| Myd88 | 3.61698388 | 4.71288709 | 151.463164 | 2.02E-18 | 6.47E-17 | F | F | T | F |
| Ppp1r3b | 4.37170605 | 4.11665846 | 151.398852 | 2.04E-18 | 6.52E-17 | T | F | T | T |
| Gm49394 | 2.50958645 | 7.19708077 | 151.116815 | 2.13E-18 | 6.79E-17 | F | F | F | F |
| Zfp64 | 2.4315671 | 6.95604267 | 151.034593 | 2.15E-18 | 6.86E-17 | F | F | T | F |
| Foxp1 | -4.4348696 | 5.4251454 | 151.012386 | 2.16E-18 | 6.87E-17 | T | F | T | T |
| Sort1 | -3.4149214 | 6.31894501 | 150.589035 | 2.30E-18 | 7.31E-17 | T | F | T | T |
| Rbm24 | -6.6643551 | 4.37529856 | 150.151172 | 2.46E-18 | 7.78E-17 | T | F | F | T |
| Gata6 | -4.9808826 | 5.07616618 | 150.146536 | 2.46E-18 | 7.78E-17 | T | F | T | F |
| Rasgef1b | 3.52611248 | 4.77920615 | 150.110899 | 2.47E-18 | 7.81E-17 | T | F | F | T |
| Crabp2 | -7.0489246 | 4.75448007 | 150.100367 | 2.48E-18 | 7.81E-17 | F | F | F | F |
| Myef2 | -2.9238878 | 7.09726223 | 150.030391 | 2.50E-18 | 7.88E-17 | T | F | F | T |
| Slc2a2 | -6.427704 | 4.52090433 | 149.686565 | 2.63E-18 | 8.28E-17 | F | F | T | F |
| Oaf | 3.02281111 | 6.07560255 | 149.636914 | 2.65E-18 | 8.33E-17 | T | F | T | T |
| Ndrg1 | 3.41620917 | 6.06476887 | 149.50286 | 2.71E-18 | 8.49E-17 | T | F | T | F |
| Clip3 | -4.226598 | 5.38728263 | 149.381019 | 2.76E-18 | 8.63E-17 | T | F | T | T |
| Sox12 | -3.315756 | 6.4620075 | 149.000854 | 2.92E-18 | 9.13E-17 | F | F | F | T |
| Smagp | 3.68758111 | 4.68494317 | 148.874864 | 2.97E-18 | 9.29E-17 | F | F | T | F |
| Dhx35 | -11.671417 | 4.07968214 | 148.732502 | 3.04E-18 | 9.47E-17 | F | F | F | T |
| Dio3 | 4.60779988 | 3.96414029 | 148.530366 | 3.13E-18 | 9.75E-17 | F | F | F | F |
| Tmem28 | -11.675733 | 4.0837359 | 148.365348 | 3.21E-18 | 9.98E-17 | F | F | T | F |
| Sh3bp1 | -4.5137907 | 5.15018924 | 147.898632 | 3.44E-18 | 1.07E-16 | T | F | F | F |
| Sirpa | -4.6636903 | 5.06002206 | 147.709354 | 3.54E-18 | 1.10E-16 | T | F | F | T |
| Slc16a6 | 2.80549499 | 5.98939816 | 147.593977 | 3.60E-18 | 1.12E-16 | F | F | T | F |
| Kif26b | -4.5364725 | 5.27616848 | 147.578804 | 3.61E-18 | 1.12E-16 | F | F | F | T |
| Baiap2 | 3.11396239 | 5.3351197 | 147.485668 | 3.66E-18 | 1.13E-16 | F | F | T | F |
| Arap2 | 2.95691802 | 5.58929843 | 147.415174 | 3.70E-18 | 1.14E-16 | F | F | T | F |
| Ttc21b | -5.4219162 | 4.70028497 | 147.324072 | 3.75E-18 | 1.16E-16 | T | F | F | F |
| Col4a2 | -2.6417156 | 7.94012239 | 147.179344 | 3.84E-18 | 1.18E-16 | F | F | T | F |
| Etv6 | -3.9126027 | 5.5985348 | 146.760332 | 4.09E-18 | 1.25E-16 | T | F | T | T |
| Tbx6 | -4.9909974 | 4.85468869 | 146.715251 | 4.11E-18 | 1.26E-16 | F | F | F | F |
| Lrp11 | -6.0367571 | 4.54189061 | 146.539644 | 4.22E-18 | 1.29E-16 | T | F | T | T |
| Twist1 | -7.0752265 | 4.38131436 | 146.144137 | 4.49E-18 | 1.37E-16 | T | F | F | T |
| Pacs2 | -3.6431106 | 5.91019041 | 145.925318 | 4.64E-18 | 1.42E-16 | T | F | F | F |
| Fam57b | -11.654806 | 4.06300445 | 145.827747 | 4.71E-18 | 1.44E-16 | F | F | T | F |

| | | | | | | | | | |
|---|---|---|---|---|---|---|---|---|---|
| Tpm4 | 2.0416582 | 9.06774978 | 145.767829 | 4.75E-18 | 1.45E-16 | T | F | T | T |
| Ppp1r9b | 2.38087077 | 7.02881708 | 145.709319 | 4.79E-18 | 1.46E-16 | T | F | T | T |
| Serpinh1 | -2.8981517 | 7.93652824 | 145.151076 | 5.22E-18 | 1.58E-16 | F | F | T | F |
| Dusp8 | 5.21665438 | 3.64461256 | 145.109813 | 5.25E-18 | 1.59E-16 | F | F | T | T |
| Ube2q2 | 2.44845764 | 6.77121092 | 144.739551 | 5.56E-18 | 1.68E-16 | T | F | T | T |
| Il17rd | -4.3086936 | 5.74172163 | 144.189654 | 6.04E-18 | 1.83E-16 | T | F | T | T |
| Mgat4a | 2.88760565 | 5.66239867 | 144.084765 | 6.14E-18 | 1.85E-16 | F | F | T | T |
| Cer1 | -5.5254578 | 5.63813547 | 143.724073 | 6.49E-18 | 1.96E-16 | T | F | T | T |
| Lasp1 | 2.08468971 | 8.48217205 | 143.666064 | 6.55E-18 | 1.97E-16 | T | F | T | T |
| Tafa5 | -8.319899 | 4.13428689 | 143.425914 | 6.79E-18 | 2.04E-16 | T | F | T | F |
| Sipa1l2 | 2.27578239 | 7.29995456 | 143.422719 | 6.80E-18 | 2.04E-16 | F | F | F | F |
| Nub1 | 2.37174007 | 6.95793622 | 143.316078 | 6.91E-18 | 2.07E-16 | T | F | T | T |
| Pou3f1 | -5.5935352 | 4.74568869 | 143.260476 | 6.97E-18 | 2.09E-16 | F | F | F | F |
| Fam168a | 2.32983355 | 7.18700978 | 143.066469 | 7.18E-18 | 2.15E-16 | T | F | T | T |
| Fgfr3 | -4.5623378 | 5.02359644 | 142.987608 | 7.27E-18 | 2.17E-16 | F | F | T | F |
| Socs2 | -8.3195018 | 4.13129719 | 142.876005 | 7.39E-18 | 2.21E-16 | T | F | T | T |
| Reln | -5.5376042 | 4.59918809 | 142.834398 | 7.44E-18 | 2.21E-16 | T | F | T | F |
| Zfp623 | -8.3059334 | 4.12053532 | 142.831682 | 7.45E-18 | 2.21E-16 | F | F | F | T |
| Msgn1 | -8.5903327 | 4.40149189 | 142.630239 | 7.68E-18 | 2.28E-16 | F | F | F | F |
| Tead4 | 2.76697578 | 6.25059462 | 142.46353 | 7.88E-18 | 2.34E-16 | T | F | T | F |
| Pecam1 | 4.00394129 | 5.25110313 | 142.038246 | 8.42E-18 | 2.49E-16 | F | F | F | F |
| Stk26 | -11.607618 | 4.01631511 | 141.801708 | 8.73E-18 | 2.58E-16 | T | F | F | F |
| Msx2 | -5.5469236 | 4.58825335 | 141.679836 | 8.90E-18 | 2.63E-16 | F | F | T | F |
| Cldn2 | -5.3647149 | 4.64386392 | 141.349985 | 9.36E-18 | 2.76E-16 | T | F | T | F |
| Bmerb1 | -11.628738 | 4.03713802 | 141.29357 | 9.45E-18 | 2.78E-16 | T | F | F | F |
| N4bp3 | 3.0195301 | 5.30975227 | 141.209851 | 9.57E-18 | 2.82E-16 | F | F | F | T |
| Gpat3 | 3.67911898 | 4.6380454 | 141.048595 | 9.81E-18 | 2.88E-16 | T | F | F | F |
| Vangl2 | -3.1683323 | 6.50150588 | 140.850304 | 1.01E-17 | 2.97E-16 | F | F | T | T |
| Smarcd3 | -8.0949332 | 4.2562361 | 140.762944 | 1.03E-17 | 3.01E-16 | F | F | F | F |
| Gab1 | 2.26073244 | 7.29001771 | 140.666228 | 1.04E-17 | 3.05E-16 | F | F | T | F |
| Chsy1 | 2.15813191 | 7.96048296 | 140.659144 | 1.04E-17 | 3.05E-16 | T | F | T | F |
| Pfpl | 4.47840456 | 6.31485934 | 140.291946 | 1.10E-17 | 3.22E-16 | F | F | F | F |
| Atp1b1 | 2.2131043 | 7.48753493 | 140.128651 | 1.13E-17 | 3.30E-16 | F | F | F | F |
| Fst | -3.6784829 | 5.94483376 | 140.086086 | 1.14E-17 | 3.32E-16 | T | F | T | T |
| Wnt3a | -8.3500427 | 4.16422562 | 139.996387 | 1.16E-17 | 3.36E-16 | T | F | T | F |
| Vil1 | -4.0547538 | 5.31555627 | 139.83247 | 1.19E-17 | 3.44E-16 | T | F | F | T |
| Tmem132c | -7.4181485 | 4.16445467 | 139.667312 | 1.22E-17 | 3.53E-16 | T | F | F | F |
| Cep164 | 2.19267539 | 7.67324627 | 139.363945 | 1.28E-17 | 3.70E-16 | F | F | T | F |
| Hap1 | 3.67893382 | 4.51887935 | 139.184486 | 1.31E-17 | 3.80E-16 | F | F | T | F |
| Zfp36l1 | -3.0181594 | 6.69789724 | 139.051705 | 1.34E-17 | 3.87E-16 | F | F | F | F |
| Plpp2 | 2.72757493 | 6.09079415 | 138.990575 | 1.35E-17 | 3.90E-16 | T | F | T | F |
| Tspan6 | -3.9161756 | 5.43297444 | 138.896376 | 1.37E-17 | 3.96E-16 | T | F | T | F |

| | | | | | | | | | |
|---|---|---|---|---|---|---|---|---|---|
| Etv4 | -5.0790213 | 4.85894107 | 138.862323 | 1.38E-17 | 3.97E-16 | F | F | F | F |
| Itga1 | 3.11780028 | 5.57663451 | 138.84131 | 1.39E-17 | 3.98E-16 | T | F | F | T |
| Skiv2l | -3.9842841 | 5.48563457 | 138.747727 | 1.41E-17 | 4.03E-16 | F | F | F | F |
| Trim6 | 2.43445014 | 6.70370076 | 138.351112 | 1.50E-17 | 4.29E-16 | F | F | T | F |
| Cdh6 | -11.545955 | 3.95554893 | 138.337213 | 1.50E-17 | 4.29E-16 | F | F | F | T |
| Rapgef2 | 2.21448295 | 7.41929021 | 138.268265 | 1.52E-17 | 4.33E-16 | T | F | T | T |
| Greb1l | -3.2470362 | 7.93363853 | 138.215658 | 1.53E-17 | 4.36E-16 | T | F | T | F |
| Dnmt3b | -2.8290721 | 9.98250542 | 138.182255 | 1.54E-17 | 4.38E-16 | F | F | F | F |
| Pxdn | -2.4808672 | 8.16030134 | 137.875634 | 1.61E-17 | 4.59E-16 | T | F | F | F |
| Il6st | 2.1159072 | 7.92995512 | 137.872805 | 1.61E-17 | 4.59E-16 | T | F | T | T |
| Noct | 1.91534924 | 10.6194608 | 137.641527 | 1.67E-17 | 4.75E-16 | F | T | F | F |
| Irs2 | -3.1428833 | 6.44212944 | 137.501059 | 1.71E-17 | 4.85E-16 | F | F | F | T |
| Wnk3 | 2.40070994 | 6.73944705 | 137.438919 | 1.73E-17 | 4.90E-16 | T | F | T | T |
| Tmod2 | -4.3145411 | 5.10920168 | 137.193824 | 1.80E-17 | 5.08E-16 | T | F | T | T |
| Prps2 | -3.764294 | 5.63866389 | 137.190613 | 1.80E-17 | 5.08E-16 | T | F | T | T |
| Ugcg | 2.2198655 | 7.31228183 | 137.057311 | 1.84E-17 | 5.18E-16 | T | F | T | T |
| Dtx3 | -4.6636226 | 4.94576361 | 136.99752 | 1.85E-17 | 5.22E-16 | F | F | T | F |
| Sh3pxd2b | -3.2939415 | 6.1998215 | 136.94732 | 1.87E-17 | 5.26E-16 | T | F | F | T |
| Bcl2 | 3.11901088 | 5.08863706 | 136.913906 | 1.88E-17 | 5.28E-16 | T | F | T | F |
| Pid1 | -5.739222 | 4.53599514 | 136.808491 | 1.91E-17 | 5.36E-16 | T | F | F | T |
| Ppp2r2c | 4.48069008 | 3.90068908 | 136.787599 | 1.92E-17 | 5.37E-16 | T | F | T | T |
| Pde3b | -4.6597967 | 4.88105862 | 136.774127 | 1.92E-17 | 5.38E-16 | T | F | T | T |
| Eef2k | 2.56667397 | 6.28216374 | 136.754587 | 1.93E-17 | 5.39E-16 | T | F | T | T |
| Grb2 | 2.18739291 | 7.52024223 | 136.700912 | 1.94E-17 | 5.43E-16 | T | F | T | F |
| Rab11fip4 | -2.913775 | 6.84797736 | 136.488652 | 2.01E-17 | 5.60E-16 | F | F | T | T |
| Tcaf1 | -4.1780915 | 5.16763107 | 136.444088 | 2.03E-17 | 5.64E-16 | F | F | T | F |
| Lpar4 | -8.2429779 | 4.05830205 | 136.434344 | 2.03E-17 | 5.64E-16 | T | F | T | T |
| Slc44a2 | 2.3698154 | 6.78188057 | 136.24371 | 2.09E-17 | 5.81E-16 | T | F | T | T |
| Nbea | -4.0233752 | 5.33228265 | 136.057236 | 2.15E-17 | 5.97E-16 | T | F | F | T |
| Wnt8a | -5.5013761 | 4.77754095 | 135.840091 | 2.23E-17 | 6.18E-16 | F | F | F | F |
| Gmip | 2.88846562 | 5.49792986 | 135.809199 | 2.24E-17 | 6.20E-16 | T | F | F | F |
| Slc38a7 | 2.36665233 | 6.80441635 | 135.66175 | 2.29E-17 | 6.34E-16 | F | F | F | F |
| Slc39a6 | 2.5645575 | 6.46546884 | 135.298711 | 2.43E-17 | 6.71E-16 | F | F | F | T |
| Hspb8 | 3.26201695 | 4.87616979 | 135.229509 | 2.46E-17 | 6.77E-16 | F | F | T | F |
| Trim25 | 2.21574421 | 7.32340378 | 135.196039 | 2.47E-17 | 6.80E-16 | T | F | T | T |
| Nrep | -5.4750107 | 4.49374179 | 135.141412 | 2.49E-17 | 6.85E-16 | F | F | T | F |
| Cotl1 | -3.0270377 | 6.68833962 | 134.998677 | 2.55E-17 | 7.00E-16 | F | F | F | F |
| Aff1 | -3.5552745 | 6.11893158 | 134.948231 | 2.57E-17 | 7.05E-16 | T | F | F | T |
| Mycl | -4.5213939 | 4.92110752 | 134.942826 | 2.57E-17 | 7.05E-16 | T | F | T | F |
| Bcl9l | -2.9883742 | 6.75361427 | 134.919826 | 2.58E-17 | 7.06E-16 | F | F | T | F |
| Pls3 | -3.028433 | 6.68427197 | 134.871743 | 2.60E-17 | 7.11E-16 | T | F | F | T |
| Col23a1 | -4.9312833 | 4.71478222 | 134.762082 | 2.65E-17 | 7.23E-16 | T | F | T | T |

| | | | | | | | | | |
|---|---|---|---|---|---|---|---|---|---|
| Iqgap2 | -3.4907396 | 5.87120183 | 134.729815 | 2.66E-17 | 7.26E-16 | F | F | F | F |
| Irx1 | 2.43902174 | 6.60625387 | 134.423194 | 2.80E-17 | 7.61E-16 | T | F | F | F |
| Gas1 | -3.9662464 | 5.31913257 | 134.396247 | 2.81E-17 | 7.64E-16 | T | F | T | T |
| Rhob | 2.44536532 | 6.98188461 | 134.387971 | 2.81E-17 | 7.64E-16 | T | F | T | T |
| Camsap3 | -3.1795492 | 6.42036148 | 134.297822 | 2.86E-17 | 7.74E-16 | F | F | F | F |
| Sesn3 | -3.1077072 | 6.61433297 | 134.184854 | 2.91E-17 | 7.86E-16 | T | F | F | T |
| Slc16a13 | -4.1114304 | 5.17469174 | 134.184484 | 2.91E-17 | 7.86E-16 | F | F | F | F |
| Celsr3 | -4.688941 | 5.0861827 | 133.966681 | 3.01E-17 | 8.13E-16 | F | F | T | F |
| Lhx1 | -5.8916402 | 4.39903901 | 133.845092 | 3.07E-17 | 8.28E-16 | F | F | T | T |
| S100a11 | 3.2586921 | 4.96701729 | 133.399432 | 3.30E-17 | 8.89E-16 | F | F | F | T |
| Laptm4b | -3.1185219 | 6.47166506 | 133.385363 | 3.31E-17 | 8.90E-16 | F | F | F | F |
| Aup1 | -11.49748 | 3.90784336 | 133.331715 | 3.34E-17 | 8.97E-16 | F | F | F | F |
| Fbxo6 | 3.50946726 | 4.56225525 | 133.282914 | 3.36E-17 | 9.03E-16 | F | F | F | F |
| Sos1 | 2.33592218 | 6.90117681 | 133.201875 | 3.41E-17 | 9.13E-16 | T | F | T | F |
| Pkd1l1 | -8.2698547 | 4.0846859 | 133.127951 | 3.45E-17 | 9.23E-16 | F | F | F | F |
| Pdzrn3 | -3.6503086 | 5.61543152 | 133.054671 | 3.49E-17 | 9.33E-16 | T | F | F | T |
| Zfyve9 | -3.5469205 | 5.82040659 | 132.873835 | 3.59E-17 | 9.60E-16 | F | F | T | T |
| Ncoa3 | -2.9179118 | 6.70609645 | 132.471486 | 3.84E-17 | 1.02E-15 | T | F | T | T |
| Irs4 | -7.3901746 | 4.13694557 | 132.434907 | 3.86E-17 | 1.03E-15 | T | F | T | T |
| Fndc3c1 | -3.197507 | 6.26744045 | 132.314249 | 3.94E-17 | 1.05E-15 | F | F | F | F |
| Pgm2 | 2.66310511 | 5.88521826 | 132.129506 | 4.06E-17 | 1.08E-15 | T | F | F | F |
| Csrp1 | 2.31065627 | 7.33662326 | 132.111061 | 4.07E-17 | 1.08E-15 | F | F | F | T |
| Dusp9 | 2.03066099 | 8.79886217 | 131.919658 | 4.20E-17 | 1.11E-15 | F | F | T | F |
| Sulf2 | -2.9015803 | 6.71773301 | 131.748877 | 4.32E-17 | 1.14E-15 | F | F | T | T |
| Mical1 | -3.9361955 | 5.24154359 | 130.548598 | 5.25E-17 | 1.39E-15 | T | F | F | F |
| Sqstm1 | 2.07244258 | 7.93152394 | 130.329467 | 5.44E-17 | 1.44E-15 | F | F | F | T |
| Pde5a | -3.9862157 | 5.19307797 | 130.171464 | 5.59E-17 | 1.47E-15 | F | F | T | T |
| Qser1 | -2.687834 | 7.17828095 | 130.140341 | 5.62E-17 | 1.48E-15 | F | F | T | T |
| Sh3tc1 | -6.448767 | 4.19827918 | 129.999453 | 5.75E-17 | 1.51E-15 | F | F | F | T |
| Rhox5 | 2.35909645 | 6.72681787 | 129.876079 | 5.86E-17 | 1.54E-15 | F | F | F | F |
| H3f3a | -2.3738303 | 8.24264834 | 129.840605 | 5.90E-17 | 1.55E-15 | T | F | F | F |
| Loxl2 | -2.9086503 | 6.63902356 | 129.547765 | 6.19E-17 | 1.62E-15 | F | F | T | T |
| Arhgap45 | -4.3000566 | 4.95751877 | 129.225728 | 6.53E-17 | 1.71E-15 | F | F | F | F |
| Plagl2 | -3.2188599 | 6.11261166 | 129.183397 | 6.57E-17 | 1.72E-15 | T | F | F | F |
| Cltb | 3.77596182 | 4.38226185 | 128.974271 | 6.81E-17 | 1.78E-15 | T | F | T | T |
| Arhgap10 | 2.37396836 | 6.62007845 | 128.866921 | 6.93E-17 | 1.81E-15 | F | F | T | F |
| Pcdh8 | -6.4128016 | 4.43946688 | 128.529425 | 7.33E-17 | 1.91E-15 | F | F | F | F |
| Pmaip1 | -6.1339718 | 4.26305038 | 128.424162 | 7.46E-17 | 1.94E-15 | F | F | T | F |
| Arap1 | 2.75738887 | 5.7666755 | 128.41693 | 7.46E-17 | 1.94E-15 | F | F | F | F |
| Card9 | 6.72081025 | 3.18225119 | 127.947426 | 8.07E-17 | 2.10E-15 | F | F | F | T |
| Crtc3 | 2.7808649 | 5.49102354 | 127.931673 | 8.09E-17 | 2.10E-15 | T | T | T | T |
| Plekhg3 | -3.2591941 | 6.0039155 | 127.781883 | 8.30E-17 | 2.15E-15 | T | F | T | T |

| | | | | | | | | | |
|---|---|---|---|---|---|---|---|---|---|
| Ccnjl | -4.3582042 | 4.90502709 | 127.776788 | 8.30E-17 | 2.15E-15 | F | F | F | F |
| Hs6st1 | -3.3198246 | 5.93652324 | 127.764129 | 8.32E-17 | 2.15E-15 | F | F | T | F |
| Cd209c | 3.05879672 | 5.14524369 | 127.333703 | 8.94E-17 | 2.31E-15 | F | F | F | T |
| Srl | -4.5544445 | 4.77845652 | 127.286327 | 9.01E-17 | 2.32E-15 | T | T | T | T |
| Inf2 | 2.37348301 | 6.62479241 | 127.092689 | 9.31E-17 | 2.40E-15 | F | F | F | F |
| Fndc4 | -7.3492217 | 4.09661901 | 127.001242 | 9.45E-17 | 2.43E-15 | F | F | T | F |
| Sema5a | -4.7465288 | 4.71343367 | 126.961001 | 9.51E-17 | 2.44E-15 | T | F | T | T |
| Add3 | -3.240346 | 6.239179 | 126.939124 | 9.55E-17 | 2.45E-15 | T | T | T | T |
| Sorl1 | -3.3015956 | 5.92873247 | 126.90616 | 9.60E-17 | 2.46E-15 | F | F | T | T |
| Hcls1 | 6.96611778 | 3.18390736 | 126.858274 | 9.68E-17 | 2.48E-15 | T | F | F | F |
| Trib2 | -6.5141073 | 4.22719446 | 126.830892 | 9.72E-17 | 2.49E-15 | T | T | T | T |
| Bicd1 | -6.7658472 | 4.15718762 | 126.754566 | 9.85E-17 | 2.52E-15 | T | F | T | F |
| Amz1 | 5.24534438 | 3.44597838 | 126.633662 | 1.00E-16 | 2.56E-15 | T | F | F | T |
| Zxdc | 2.54693551 | 6.03229094 | 126.395603 | 1.05E-16 | 2.66E-15 | T | T | T | T |
| Spin2c | -7.3371381 | 4.08471513 | 126.050909 | 1.11E-16 | 2.82E-15 | F | F | F | F |
| Ncs1 | -3.291971 | 5.90510832 | 125.778714 | 1.16E-16 | 2.95E-15 | T | F | T | T |
| Jade2 | -4.4432496 | 4.8995009 | 125.739322 | 1.17E-16 | 2.96E-15 | T | F | F | F |
| Cenpb | -2.9805108 | 6.41304887 | 125.66337 | 1.18E-16 | 3.00E-15 | F | F | F | T |
| Klhl13 | 2.55141024 | 6.05893677 | 125.527712 | 1.21E-16 | 3.07E-15 | T | F | T | T |
| Rtn1 | 3.1541749 | 4.83271232 | 125.391169 | 1.24E-16 | 3.13E-15 | T | F | T | F |
| Amer1 | -3.3737185 | 5.85529318 | 125.208037 | 1.28E-16 | 3.23E-15 | T | F | T | T |
| Lmx1b | -6.8703479 | 4.18218609 | 124.989254 | 1.33E-16 | 3.35E-15 | F | F | T | F |
| Tmcc2 | -11.385214 | 3.79670143 | 124.982332 | 1.33E-16 | 3.35E-15 | F | F | T | F |
| C1galt1 | 2.4669439 | 6.2408367 | 124.862483 | 1.35E-16 | 3.41E-15 | T | F | F | T |
| Atp2a3 | 3.25463304 | 4.7322958 | 124.756649 | 1.38E-16 | 3.47E-15 | T | F | F | T |
| Zfp42 | 5.5247842 | 3.45567237 | 124.696295 | 1.39E-16 | 3.50E-15 | T | T | F | T |
| Pnma2 | -6.2023406 | 4.23184174 | 124.627847 | 1.41E-16 | 3.54E-15 | T | F | T | T |
| Tnfaip2 | 3.8095383 | 4.74700214 | 124.571147 | 1.42E-16 | 3.57E-15 | T | F | F | T |
| Pip4k2c | 2.21981389 | 6.91751716 | 124.504446 | 1.44E-16 | 3.60E-15 | T | F | T | F |
| Ecm1 | 3.56485159 | 4.32177603 | 124.237815 | 1.51E-16 | 3.76E-15 | F | F | F | F |
| Zfp334 | -2.9894416 | 6.35399515 | 124.060043 | 1.55E-16 | 3.87E-15 | T | F | T | T |
| Wdr1 | 1.94904133 | 8.42764753 | 123.756447 | 1.63E-16 | 4.08E-15 | T | F | T | F |
| Fbxo27 | 3.81557091 | 4.14895011 | 123.669089 | 1.66E-16 | 4.13E-15 | F | F | T | F |
| Amph | -11.378452 | 3.79052287 | 123.025819 | 1.85E-16 | 4.61E-15 | T | F | T | T |
| Las1l | 2.04406115 | 7.6411805 | 122.670303 | 1.97E-16 | 4.89E-15 | T | F | F | T |
| Igsf9 | -3.2722761 | 5.84061624 | 122.642457 | 1.98E-16 | 4.91E-15 | T | F | F | F |
| Cxadr | -2.9473755 | 6.60206355 | 122.024395 | 2.20E-16 | 5.45E-15 | T | T | T | T |
| Ccnd3 | 2.10816388 | 7.95719886 | 121.926685 | 2.23E-16 | 5.53E-15 | T | F | T | F |
| Reg2 | 2.42969416 | 6.72033588 | 121.841561 | 2.27E-16 | 5.61E-15 | F | F | F | F |
| Cpeb2 | 2.9366804 | 5.09596065 | 121.479866 | 2.41E-16 | 5.96E-15 | T | T | F | T |
| Anxa11 | 2.33009848 | 6.63639077 | 121.263225 | 2.50E-16 | 6.18E-15 | T | F | T | F |
| Adam10 | 2.35726556 | 6.42068942 | 121.211235 | 2.53E-16 | 6.23E-15 | T | F | F | T |

| | | | | | | | | | |
|---|---|---|---|---|---|---|---|---|---|
| Dock5 | 2.09363009 | 7.45812979 | 121.028954 | 2.61E-16 | 6.42E-15 | F | F | T | F |
| Tgfb3 | 5.58229153 | 3.4462698 | 120.976516 | 2.63E-16 | 6.47E-15 | F | F | T | F |
| Galnt16 | -6.1097608 | 4.14085168 | 120.42109 | 2.90E-16 | 7.12E-15 | T | F | T | T |
| Tceal9 | -2.9168461 | 6.560847 | 120.256605 | 2.98E-16 | 7.32E-15 | T | F | F | F |
| Rragb | -6.7268503 | 4.03784429 | 120.076462 | 3.07E-16 | 7.54E-15 | F | F | T | F |
| Syngap1 | -3.8146894 | 5.35288414 | 120.056537 | 3.08E-16 | 7.56E-15 | F | F | F | T |
| Zfp280c | -3.5878952 | 5.47589349 | 120.040918 | 3.09E-16 | 7.57E-15 | T | F | F | F |
| Mthfd2 | 2.2246361 | 6.80692635 | 119.955144 | 3.14E-16 | 7.67E-15 | F | F | F | F |
| Castor1 | 3.2745615 | 4.61433787 | 119.942958 | 3.15E-16 | 7.68E-15 | F | F | F | F |
| Gldc | -2.3765219 | 7.73760709 | 119.883627 | 3.18E-16 | 7.75E-15 | T | F | F | T |
| Slc43a1 | -11.326025 | 3.738751 | 119.800673 | 3.22E-16 | 7.85E-15 | T | F | F | T |
| Gprc5b | -6.728223 | 4.03917733 | 119.797285 | 3.23E-16 | 7.85E-15 | T | F | T | T |
| Wnk2 | 2.15048743 | 7.04334081 | 119.619401 | 3.33E-16 | 8.09E-15 | T | F | F | F |
| Hmga2 | -2.3637537 | 7.92820205 | 119.40521 | 3.45E-16 | 8.39E-15 | T | F | T | F |
| Apela | -7.2595809 | 4.00835926 | 119.384005 | 3.47E-16 | 8.41E-15 | F | F | F | F |
| Cuedc2 | -4.5179319 | 4.65185631 | 119.17394 | 3.60E-16 | 8.71E-15 | F | F | F | F |
| Prl3d2 | 5.91615481 | 3.12474905 | 119.158652 | 3.61E-16 | 8.73E-15 | F | F | F | F |
| Grik5 | -3.8599588 | 5.1524327 | 119.130079 | 3.62E-16 | 8.76E-15 | F | F | T | F |
| Gas6 | -11.319233 | 3.73211092 | 119.00877 | 3.70E-16 | 8.94E-15 | F | F | F | F |
| Pwwp3b | -3.4563986 | 5.48235559 | 118.632415 | 3.95E-16 | 9.54E-15 | F | F | F | F |
| Ctdspl | 2.33889138 | 6.41083294 | 118.417676 | 4.10E-16 | 9.89E-15 | T | F | T | T |
| Shroom4 | 2.25918917 | 6.65254269 | 118.231842 | 4.24E-16 | 1.02E-14 | F | F | T | T |
| Ptbp3 | 2.01472155 | 7.65437598 | 118.185615 | 4.28E-16 | 1.03E-14 | T | T | T | T |
| Nphp1 | -11.290826 | 3.70371031 | 118.159451 | 4.29E-16 | 1.03E-14 | F | F | F | T |
| Fzd4 | -4.2016672 | 4.94471219 | 118.023506 | 4.40E-16 | 1.06E-14 | F | F | F | T |
| Rhpn2 | -4.374868 | 4.71957842 | 118.014249 | 4.41E-16 | 1.06E-14 | T | F | T | F |
| Fat3 | -6.0850114 | 4.11642465 | 117.923865 | 4.48E-16 | 1.07E-14 | T | F | T | F |
| Castor2 | -2.9520424 | 6.36909153 | 117.508512 | 4.82E-16 | 1.15E-14 | T | F | T | T |
| Fign | -11.33097 | 3.74263069 | 117.4121 | 4.90E-16 | 1.17E-14 | F | F | T | T |
| Nefl | -4.6676001 | 4.53913987 | 117.246741 | 5.04E-16 | 1.20E-14 | F | F | T | T |
| Prom2 | 4.81822859 | 3.44525488 | 117.231111 | 5.06E-16 | 1.20E-14 | T | F | F | T |
| Plin2 | 2.42743114 | 6.26454113 | 116.951287 | 5.31E-16 | 1.26E-14 | F | F | T | F |
| Gm50241 | -11.314752 | 3.72662169 | 116.397474 | 5.86E-16 | 1.39E-14 | F | F | F | F |
| Tubb6 | 2.27172716 | 6.80474273 | 116.331313 | 5.93E-16 | 1.41E-14 | F | F | T | T |
| Egflam | -11.269252 | 3.68215521 | 116.163931 | 6.11E-16 | 1.45E-14 | T | F | T | T |
| Sult4a1 | -6.6670027 | 3.97908348 | 116.162817 | 6.11E-16 | 1.45E-14 | T | F | T | T |
| Prxl2a | -3.5354532 | 5.34498961 | 116.147271 | 6.13E-16 | 1.45E-14 | F | F | F | F |
| Mt1 | 2.36907472 | 6.36253523 | 116.127397 | 6.15E-16 | 1.45E-14 | F | F | F | T |
| Lrrn1 | -8.0510747 | 3.86851137 | 115.914825 | 6.38E-16 | 1.51E-14 | F | F | F | F |
| Nfkb1 | -3.5302729 | 5.320073 | 115.868639 | 6.44E-16 | 1.52E-14 | T | F | F | T |
| Prkd1 | -4.2769375 | 4.73764526 | 115.796119 | 6.52E-16 | 1.54E-14 | T | F | F | F |
| Slc38a3 | 3.01033834 | 4.845494 | 115.773809 | 6.55E-16 | 1.54E-14 | T | F | T | T |

| | | | | | | | | | |
|---|---|---|---|---|---|---|---|---|---|
| Perp | 3.07517675 | 4.75972377 | 115.76046 | 6.56E-16 | 1.54E-14 | T | F | T | F |
| Gprasp1 | -3.1996967 | 5.75342569 | 115.645279 | 6.70E-16 | 1.57E-14 | T | F | F | T |
| Ubqln4 | -2.4476418 | 7.21197018 | 115.53289 | 6.83E-16 | 1.60E-14 | F | F | F | F |
| Tspan12 | -5.2686487 | 4.29143515 | 115.527875 | 6.84E-16 | 1.60E-14 | T | F | T | T |
| Fgb | -7.1739712 | 3.92398716 | 115.342339 | 7.07E-16 | 1.66E-14 | F | F | F | T |
| Adgrb2 | -4.1225314 | 4.83118942 | 115.310228 | 7.11E-16 | 1.66E-14 | F | F | F | F |
| Kif1c | 1.89322892 | 8.26058758 | 115.114206 | 7.36E-16 | 1.72E-14 | T | F | T | T |
| Lama5 | -2.2082959 | 8.27796924 | 115.079622 | 7.41E-16 | 1.73E-14 | F | F | F | F |
| Gjc1 | -4.4343486 | 4.63082582 | 115.001638 | 7.51E-16 | 1.75E-14 | T | F | F | T |
| Slc38a1 | 1.79568363 | 9.03290122 | 114.993529 | 7.52E-16 | 1.75E-14 | T | F | T | T |
| Ttc3 | -2.2925612 | 7.82620942 | 114.703059 | 7.92E-16 | 1.84E-14 | T | F | F | F |
| Trim8 | 1.93427582 | 7.8955228 | 114.58102 | 8.10E-16 | 1.88E-14 | F | F | F | T |
| Dkkl1 | 5.67648575 | 3.0440552 | 114.136181 | 8.77E-16 | 2.04E-14 | F | F | F | F |
| 4933427D1 | -11.234475 | 3.64795771 | 114.036326 | 8.93E-16 | 2.07E-14 | F | F | T | F |
| Diaph2 | 2.63608222 | 5.46012801 | 114.013534 | 8.96E-16 | 2.08E-14 | F | F | T | F |
| Lrrc8c | 2.53856221 | 5.75867972 | 113.933865 | 9.09E-16 | 2.10E-14 | F | F | F | T |
| Klf13 | 1.95529262 | 7.99466948 | 113.80407 | 9.31E-16 | 2.15E-14 | F | F | T | F |
| Ube2a | 2.45091433 | 5.93365575 | 113.70622 | 9.47E-16 | 2.19E-14 | T | F | F | T |
| Inava | -3.6250702 | 5.17897705 | 113.48157 | 9.86E-16 | 2.28E-14 | F | F | F | F |
| Rara | -3.5753663 | 5.33149459 | 113.430058 | 9.96E-16 | 2.29E-14 | F | F | F | F |
| Foxa1 | -7.1227979 | 3.87327029 | 113.166143 | 1.04E-15 | 2.40E-14 | T | F | F | T |
| Myo6 | -2.9546614 | 6.10139506 | 113.126622 | 1.05E-15 | 2.42E-14 | T | F | T | T |
| Rasgrp2 | -11.222563 | 3.63612156 | 113.010896 | 1.07E-15 | 2.47E-14 | F | F | F | F |
| B4galnt4 | -3.5706804 | 5.22640958 | 112.953692 | 1.09E-15 | 2.49E-14 | F | F | F | T |
| Pik3cb | 2.66129789 | 5.36715405 | 112.952648 | 1.09E-15 | 2.49E-14 | T | F | F | F |
| Sdk2 | 2.41757263 | 6.22453627 | 112.947742 | 1.09E-15 | 2.49E-14 | F | F | F | F |
| Mcf2 | -11.313169 | 3.72477462 | 112.881256 | 1.10E-15 | 2.51E-14 | T | F | F | F |
| Magel2 | -7.973711 | 3.79243358 | 112.853419 | 1.10E-15 | 2.52E-14 | T | F | F | F |
| Cap2 | -7.9271858 | 3.78658157 | 112.765021 | 1.12E-15 | 2.56E-14 | F | F | F | T |
| St6gal1 | -3.4968271 | 5.26651566 | 112.628412 | 1.15E-15 | 2.62E-14 | T | T | F | T |
| Zhx1 | -4.9850778 | 4.30378273 | 112.174682 | 1.25E-15 | 2.85E-14 | T | F | T | T |
| Fhl2 | 3.02786421 | 4.75834695 | 111.956205 | 1.30E-15 | 2.96E-14 | F | F | F | F |
| Zfp36 | 2.8722835 | 5.03086638 | 111.935802 | 1.30E-15 | 2.97E-14 | F | F | F | F |
| Lrp3 | -3.9196059 | 4.91797126 | 111.622899 | 1.38E-15 | 3.14E-14 | F | F | F | F |
| Casp3 | -2.8686889 | 6.27269586 | 111.583179 | 1.39E-15 | 3.16E-14 | T | F | F | F |
| Cln6 | 2.55603471 | 5.60391205 | 111.413138 | 1.44E-15 | 3.25E-14 | F | F | T | F |
| Fndc3b | 2.09498321 | 7.12586535 | 111.400825 | 1.44E-15 | 3.26E-14 | T | F | T | F |
| Tmem37 | 2.90213057 | 5.1230608 | 111.31045 | 1.46E-15 | 3.31E-14 | F | F | F | F |
| Fbxw5 | -4.5449116 | 4.48508953 | 110.801018 | 1.60E-15 | 3.62E-14 | F | F | F | F |
| Rimbp2 | -5.9125508 | 4.50296867 | 110.677957 | 1.64E-15 | 3.70E-14 | T | F | T | F |
| Aldob | -5.218096 | 4.24204097 | 110.486766 | 1.70E-15 | 3.83E-14 | T | F | F | T |
| Cdk2ap1 | -2.2683169 | 7.7196058 | 110.448068 | 1.71E-15 | 3.85E-14 | F | F | T | F |

| | | | | | | | | | |
|---|---|---|---|---|---|---|---|---|---|
| Nfatc4 | -4.6305406 | 4.43597144 | 110.284169 | 1.76E-15 | 3.97E-14 | T | F | F | T |
| Hip1 | -2.3856792 | 7.1861847 | 110.217053 | 1.79E-15 | 4.01E-14 | T | F | T | T |
| Nin | 2.08068188 | 6.91508562 | 109.816322 | 1.92E-15 | 4.32E-14 | T | T | T | T |
| Plekhg5 | -4.5907487 | 4.46449915 | 109.649531 | 1.98E-15 | 4.45E-14 | T | F | T | T |
| Fabp5 | -2.9526186 | 6.03219733 | 109.319303 | 2.11E-15 | 4.72E-14 | F | F | F | F |
| Ppp1r1a | -6.6426601 | 3.95553434 | 109.312594 | 2.11E-15 | 4.72E-14 | F | F | F | F |
| Dapk1 | -2.9295588 | 6.40919128 | 109.285907 | 2.12E-15 | 4.74E-14 | T | F | F | T |
| Pth1r | -3.2492963 | 5.52317232 | 109.158885 | 2.17E-15 | 4.85E-14 | F | F | F | F |
| Rell1 | 2.34921561 | 6.13889821 | 109.109756 | 2.19E-15 | 4.89E-14 | F | F | T | F |
| Hmcn1 | -7.9703518 | 3.79013564 | 109.096668 | 2.20E-15 | 4.89E-14 | T | T | T | T |
| Ctdsp2 | -2.2843688 | 7.73298149 | 109.096097 | 2.20E-15 | 4.89E-14 | T | F | T | T |
| Spaar | 10.8128213 | 2.2030658 | 110.044184 | 2.24E-15 | 4.98E-14 | T | F | T | F |
| Slc12a7 | 1.95235452 | 7.68910559 | 108.968508 | 2.25E-15 | 4.99E-14 | T | F | F | F |
| Ctsd | 1.92490477 | 7.98617434 | 108.88539 | 2.28E-15 | 5.07E-14 | F | F | T | F |
| Tef | -4.4317046 | 4.58769644 | 108.845207 | 2.30E-15 | 5.10E-14 | T | F | T | F |
| Gm14440 | 10.8080362 | 2.19829725 | 109.733719 | 2.37E-15 | 5.24E-14 | F | F | F | T |
| Dact1 | -3.8304759 | 4.97178519 | 108.470651 | 2.47E-15 | 5.45E-14 | T | F | T | F |
| Plcg2 | -3.74937 | 5.0219419 | 108.24597 | 2.57E-15 | 5.68E-14 | T | T | F | T |
| Sema4c | 1.98283779 | 7.2663717 | 108.116391 | 2.63E-15 | 5.81E-14 | F | F | F | F |
| Slco4c1 | -7.891664 | 3.71201621 | 108.031793 | 2.67E-15 | 5.90E-14 | F | F | F | T |
| Derl3 | 3.65838332 | 4.09732911 | 107.617993 | 2.89E-15 | 6.37E-14 | F | F | F | F |
| Phf21b | -11.159253 | 3.57335776 | 107.553591 | 2.92E-15 | 6.44E-14 | T | F | T | F |
| Ptch1 | -2.4330034 | 7.74769856 | 107.449923 | 2.98E-15 | 6.55E-14 | F | F | T | T |
| Arpc2 | 1.84800532 | 8.08452324 | 107.429762 | 2.99E-15 | 6.57E-14 | T | F | F | F |
| Blmh | -2.4952304 | 6.83645059 | 107.397389 | 3.01E-15 | 6.60E-14 | F | F | F | F |
| Igdcc4 | -7.8838865 | 3.70419055 | 107.396124 | 3.01E-15 | 6.60E-14 | T | F | T | T |
| Abcc1 | 2.13873071 | 6.66986697 | 107.299291 | 3.07E-15 | 6.71E-14 | F | F | T | F |
| Slc23a2 | 2.38798461 | 5.89875877 | 107.109782 | 3.18E-15 | 6.95E-14 | T | T | F | T |
| Wnt2b | -6.6291084 | 4.34088256 | 107.048261 | 3.21E-15 | 7.02E-14 | T | F | T | T |
| Vav3 | 3.2728855 | 4.32233511 | 106.80122 | 3.37E-15 | 7.35E-14 | T | F | T | F |
| Sept3 | -6.1526662 | 3.93569073 | 106.596701 | 3.50E-15 | 7.62E-14 | T | F | T | T |
| Map2k3 | 2.27940019 | 6.28686836 | 106.594869 | 3.50E-15 | 7.62E-14 | F | F | T | T |
| Rcbtb1 | -2.7513824 | 6.29005554 | 106.578045 | 3.51E-15 | 7.64E-14 | T | F | T | T |
| Cnn3 | -2.2451214 | 7.61982652 | 106.473517 | 3.58E-15 | 7.78E-14 | T | F | F | T |
| Boc | -4.9735383 | 4.2613173 | 106.271788 | 3.72E-15 | 8.07E-14 | T | T | F | T |
| Tmcc3 | 2.78287515 | 5.10323971 | 106.108865 | 3.83E-15 | 8.32E-14 | T | F | T | T |
| Syngr1 | 4.33966159 | 3.67068958 | 106.074594 | 3.86E-15 | 8.36E-14 | T | F | T | F |
| Wls | -3.1159197 | 5.9017034 | 105.983343 | 3.92E-15 | 8.50E-14 | T | F | F | F |
| Arhgef26 | -3.9799027 | 4.73750671 | 105.895593 | 3.99E-15 | 8.63E-14 | F | F | F | F |
| Spata13 | -2.2887229 | 7.52891729 | 105.687261 | 4.15E-15 | 8.97E-14 | T | F | T | F |
| Ccrl2 | 6.70347469 | 2.81609418 | 105.622625 | 4.20E-15 | 9.07E-14 | F | F | T | T |
| Ppt2 | -3.6253968 | 5.047359 | 105.560735 | 4.25E-15 | 9.17E-14 | F | F | F | F |

| | | | | | | | | | |
|---|---|---|---|---|---|---|---|---|---|
| Prkar2b | -4.9700607 | 4.25792513 | 105.473261 | 4.32E-15 | 9.31E-14 | F | F | T | F |
| Aass | -5.7480438 | 4.04136885 | 105.465471 | 4.33E-15 | 9.31E-14 | T | F | F | T |
| Gpc6 | -5.3020953 | 4.17535222 | 105.279725 | 4.48E-15 | 9.64E-14 | T | F | T | T |
| Ttyh3 | -2.2650206 | 7.43550131 | 105.229362 | 4.52E-15 | 9.72E-14 | T | F | F | F |
| Rimklb | -4.2637345 | 4.55708859 | 105.225427 | 4.53E-15 | 9.72E-14 | T | F | T | T |
| Cdhr2 | -5.6676533 | 4.17905108 | 105.201652 | 4.55E-15 | 9.75E-14 | F | F | F | F |
| Folr1 | -4.3867836 | 4.48572511 | 105.195729 | 4.55E-15 | 9.75E-14 | F | F | F | F |
| Mcc | -2.7389857 | 6.32432565 | 105.166039 | 4.58E-15 | 9.80E-14 | T | F | T | T |
| Ucp2 | -3.2708989 | 5.42225143 | 104.987601 | 4.74E-15 | 1.01E-13 | F | F | F | F |
| DXBay18 | 3.38434055 | 4.45093229 | 104.90347 | 4.81E-15 | 1.03E-13 | F | F | F | F |
| Sdc3 | 2.57712487 | 5.4816997 | 104.784115 | 4.92E-15 | 1.05E-13 | T | F | T | T |
| Rassf8 | 2.43224061 | 5.84492031 | 104.696884 | 5.00E-15 | 1.07E-13 | T | F | T | T |
| Nwd1 | 7.1634019 | 2.49385616 | 104.67675 | 5.02E-15 | 1.07E-13 | T | F | T | T |
| Itpripl2 | 2.64947233 | 5.16893966 | 104.663382 | 5.04E-15 | 1.07E-13 | T | F | T | F |
| Lonrf3 | 5.34944234 | 3.09552807 | 104.483375 | 5.21E-15 | 1.11E-13 | T | F | F | T |
| Stat3 | 2.32138668 | 6.10243408 | 104.270572 | 5.43E-15 | 1.15E-13 | F | F | T | T |
| 2610318N0 | -4.6012507 | 4.34611745 | 104.183059 | 5.52E-15 | 1.17E-13 | F | F | F | F |
| Fgf8 | -7.0054272 | 3.75809032 | 104.061468 | 5.65E-15 | 1.20E-13 | F | F | T | F |
| Gramd2 | 4.39705841 | 3.52809609 | 103.799207 | 5.93E-15 | 1.26E-13 | F | F | T | F |
| Fgfr2 | -2.7577904 | 6.21631972 | 103.739971 | 6.00E-15 | 1.27E-13 | T | F | F | T |
| Fignl2 | 1.80035068 | 8.19278951 | 103.731583 | 6.01E-15 | 1.27E-13 | F | F | T | F |
| Gk5 | 2.72495827 | 5.02964692 | 103.675456 | 6.08E-15 | 1.28E-13 | T | F | T | T |
| Apcdd1 | -3.8922405 | 4.74842359 | 103.580615 | 6.19E-15 | 1.31E-13 | F | F | F | F |
| Itsn2 | 2.35245966 | 5.97027287 | 103.37538 | 6.44E-15 | 1.36E-13 | T | F | F | T |
| Hoxb2 | -6.3401261 | 4.05639939 | 103.109945 | 6.77E-15 | 1.43E-13 | F | F | F | F |
| Zhx2 | -5.9055313 | 3.94023373 | 103.089448 | 6.80E-15 | 1.43E-13 | F | F | F | F |
| Ccdc88c | -2.6737872 | 6.32239209 | 102.920007 | 7.02E-15 | 1.48E-13 | T | F | F | T |
| Fgf15 | -8.04868 | 3.86766861 | 102.812343 | 7.17E-15 | 1.50E-13 | F | F | F | F |
| Ninj1 | 2.61646048 | 5.32498811 | 102.738611 | 7.27E-15 | 1.52E-13 | F | F | F | F |
| Zfp7 | -4.353721 | 4.46004035 | 102.531349 | 7.57E-15 | 1.58E-13 | T | F | F | F |
| Prl7a1 | 4.91990352 | 3.63010406 | 102.483946 | 7.63E-15 | 1.60E-13 | F | F | F | F |
| Cplx2 | -5.7528298 | 3.95770354 | 102.266943 | 7.96E-15 | 1.66E-13 | F | F | T | T |
| B4galt6 | -3.0792358 | 5.5601786 | 102.091419 | 8.23E-15 | 1.72E-13 | F | F | T | F |
| Trim28 | -1.792726 | 10.0987629 | 101.870673 | 8.59E-15 | 1.79E-13 | F | F | F | F |
| Tbc1d16 | -2.5601845 | 6.5342741 | 101.849085 | 8.63E-15 | 1.80E-13 | F | F | T | T |
| Anxa7 | 2.06204156 | 6.73159159 | 101.795126 | 8.72E-15 | 1.82E-13 | T | F | T | T |
| Kirrel | -2.8460343 | 6.05228041 | 101.399661 | 9.41E-15 | 1.96E-13 | T | F | T | T |
| Man2a2 | 2.38324373 | 6.31851596 | 101.391827 | 9.43E-15 | 1.96E-13 | T | F | T | T |
| Atxn1l | -2.5315811 | 6.57019056 | 101.308789 | 9.58E-15 | 1.99E-13 | T | F | T | T |
| Raver2 | -4.5652492 | 4.35335833 | 101.243988 | 9.70E-15 | 2.01E-13 | T | F | T | T |
| 4933402E1 | -6.1373527 | 3.85695218 | 101.184481 | 9.81E-15 | 2.03E-13 | F | F | F | F |
| Thbs3 | -7.696616 | 3.64992005 | 101.041188 | 1.01E-14 | 2.09E-13 | T | F | F | T |

| | | | | | | | | | |
|---|---|---|---|---|---|---|---|---|---|
| Cmklr1 | 2.75634186 | 4.92070348 | 101.015304 | 1.01E-14 | 2.10E-13 | F | F | F | T |
| Ppp1r26 | -4.2090308 | 4.48566543 | 100.804149 | 1.06E-14 | 2.18E-13 | T | F | F | F |
| Rcn1 | -2.7920334 | 6.2235819 | 100.775466 | 1.06E-14 | 2.19E-13 | F | F | F | F |
| Hs3st1 | 3.94570971 | 3.74343814 | 100.687342 | 1.08E-14 | 2.23E-13 | T | F | F | T |
| Fat4 | -5.6563495 | 3.95143381 | 100.277299 | 1.17E-14 | 2.41E-13 | T | F | F | F |
| Cadm1 | -2.6750402 | 6.22923433 | 100.207048 | 1.19E-14 | 2.44E-13 | F | F | T | F |
| Meis1 | -4.8271026 | 4.42171941 | 100.096327 | 1.21E-14 | 2.49E-13 | F | F | F | T |
| Ern1 | 2.3731794 | 5.85545009 | 99.9215686 | 1.25E-14 | 2.58E-13 | T | F | T | T |
| Cts7 | 4.18521358 | 4.26874815 | 99.5870349 | 1.34E-14 | 2.75E-13 | F | F | F | F |
| Slitrk5 | -10.997836 | 3.41503704 | 99.4287747 | 1.38E-14 | 2.83E-13 | T | F | T | T |
| Oas1a | 4.8812128 | 3.20516478 | 99.4043611 | 1.39E-14 | 2.84E-13 | F | F | F | T |
| Heph | -6.9329215 | 3.68646396 | 99.2750195 | 1.42E-14 | 2.92E-13 | T | F | F | T |
| Fbn1 | -4.424814 | 4.8710432 | 99.1348135 | 1.46E-14 | 2.99E-13 | T | F | F | F |
| Dach1 | -11.019714 | 3.43768805 | 98.9293776 | 1.52E-14 | 3.11E-13 | T | F | T | F |
| Pappa | -5.4162605 | 3.9913648 | 98.7951877 | 1.56E-14 | 3.19E-13 | T | T | T | T |
| Dipk2a | -3.2079273 | 5.28622842 | 98.7238543 | 1.59E-14 | 3.24E-13 | T | F | T | T |
| Slc44a5 | -7.7495022 | 3.57207595 | 98.6137753 | 1.62E-14 | 3.30E-13 | T | F | T | T |
| Ptp4a3 | 2.55554846 | 5.22527785 | 98.5181379 | 1.65E-14 | 3.36E-13 | F | F | T | T |
| Nrp2 | -2.6099988 | 6.37391132 | 98.3148937 | 1.72E-14 | 3.50E-13 | T | F | T | F |
| Gng12 | 2.13386476 | 6.37718953 | 98.274974 | 1.73E-14 | 3.52E-13 | T | F | T | T |
| Fgf3 | -4.8440946 | 4.24859932 | 98.2567805 | 1.74E-14 | 3.53E-13 | F | F | T | F |
| Myo7a | -3.5311051 | 4.92980885 | 98.223399 | 1.75E-14 | 3.55E-13 | T | F | F | F |
| Lrch2 | -7.7483397 | 3.57068565 | 98.218684 | 1.75E-14 | 3.55E-13 | F | F | F | T |
| Tanc2 | 2.07152243 | 6.65826063 | 98.1561951 | 1.77E-14 | 3.59E-13 | T | F | T | T |
| Myl7 | -10.977698 | 3.39510522 | 98.1546538 | 1.77E-14 | 3.59E-13 | F | F | F | F |
| Tcf20 | 1.6757333 | 8.92168816 | 97.977177 | 1.84E-14 | 3.71E-13 | F | F | F | T |
| Aldh3b2 | 6.80345533 | 2.47699932 | 97.9442444 | 1.85E-14 | 3.74E-13 | F | F | T | F |
| Ptma | -1.6685267 | 10.7530294 | 97.5689575 | 1.99E-14 | 4.02E-13 | F | F | T | F |
| Bmp2 | -3.8250038 | 4.67256372 | 97.5649539 | 1.99E-14 | 4.02E-13 | T | F | F | T |
| Mast4 | -2.9913385 | 5.54711431 | 97.5493627 | 2.00E-14 | 4.03E-13 | T | F | F | F |
| Foxa2 | -4.3165802 | 4.41805467 | 97.3711378 | 2.07E-14 | 4.17E-13 | T | T | F | T |
| Plekhh2 | -11.004768 | 3.42322796 | 97.2093644 | 2.14E-14 | 4.30E-13 | T | F | T | T |
| Ksr1 | 2.28434369 | 6.07481188 | 97.1313473 | 2.17E-14 | 4.36E-13 | F | F | T | F |
| BC067074 | -4.8431553 | 4.13434169 | 96.9623332 | 2.25E-14 | 4.51E-13 | F | F | F | F |
| Capn6 | -4.6961203 | 4.20988549 | 96.9366183 | 2.26E-14 | 4.53E-13 | T | F | F | T |
| Lonp2 | 2.12776662 | 6.35190246 | 96.8865693 | 2.28E-14 | 4.57E-13 | F | F | F | T |
| Hoxa1 | -5.2273177 | 5.37109986 | 96.7458869 | 2.35E-14 | 4.70E-13 | T | F | F | F |
| Lrp5 | -2.2223997 | 7.22350113 | 96.5891689 | 2.42E-14 | 4.84E-13 | T | F | F | F |
| Ric8b | -3.8900321 | 4.64347032 | 96.5577419 | 2.44E-14 | 4.87E-13 | F | F | T | F |
| Slc1a3 | -6.0796685 | 3.80006269 | 96.5207296 | 2.46E-14 | 4.90E-13 | T | F | T | T |
| Ncoa2 | -2.5841958 | 6.29317273 | 96.4326627 | 2.50E-14 | 4.98E-13 | T | T | T | T |
| Calu | 1.68932671 | 8.63401263 | 96.2501423 | 2.59E-14 | 5.16E-13 | T | F | T | T |

| | | | | | | | | | |
|---|---|---|---|---|---|---|---|---|---|
| Adgrv1 | -7.7614735 | 3.58548785 | 96.0967568 | 2.67E-14 | 5.32E-13 | F | F | F | F |
| Furin | 1.76275746 | 8.04239867 | 96.0105145 | 2.72E-14 | 5.40E-13 | F | F | F | F |
| Tmc7 | -4.7737901 | 4.18020372 | 95.967416 | 2.74E-14 | 5.45E-13 | T | F | T | T |
| Plaur | 6.10488984 | 2.92477466 | 95.938356 | 2.76E-14 | 5.47E-13 | T | F | F | F |
| Cpeb4 | 2.35488265 | 5.63814044 | 95.9031476 | 2.78E-14 | 5.51E-13 | F | F | T | F |
| Arl8a | 2.33617003 | 5.70637046 | 95.8182067 | 2.83E-14 | 5.60E-13 | F | F | T | F |
| Tmed5 | 2.05762113 | 6.63559076 | 95.7532321 | 2.86E-14 | 5.66E-13 | T | F | T | T |
| Tspan5 | -11.071581 | 3.4859174 | 95.7357652 | 2.87E-14 | 5.68E-13 | T | F | T | F |
| Zfp503 | -6.8969936 | 4.60401593 | 95.7195051 | 2.88E-14 | 5.69E-13 | F | F | T | F |
| Ssbp3 | -2.1862257 | 7.21493109 | 95.4602313 | 3.04E-14 | 5.99E-13 | T | F | F | F |
| Plk3 | 3.03276132 | 4.55525645 | 95.2215708 | 3.19E-14 | 6.28E-13 | F | F | F | F |
| Pcdh1 | -2.2511165 | 7.02278551 | 95.0399329 | 3.30E-14 | 6.51E-13 | F | F | T | T |
| Pou2f3 | 5.41072118 | 2.87202284 | 94.9853287 | 3.34E-14 | 6.57E-13 | T | T | F | T |
| B4galt5 | 1.791143 | 7.72481184 | 94.8973941 | 3.40E-14 | 6.68E-13 | F | F | F | F |
| Hmgn2 | -1.9406595 | 8.50727559 | 94.8490674 | 3.43E-14 | 6.74E-13 | T | T | T | T |
| Parvb | -2.6704609 | 6.05440049 | 94.8388574 | 3.44E-14 | 6.75E-13 | T | F | F | T |
| Sfi1 | 2.59373987 | 5.07384925 | 94.7918792 | 3.47E-14 | 6.81E-13 | T | F | F | F |
| Slc12a2 | -2.8037257 | 5.93774164 | 94.7825325 | 3.48E-14 | 6.82E-13 | T | F | T | F |
| Rgs5 | -7.9288106 | 3.74997564 | 94.7679979 | 3.49E-14 | 6.83E-13 | T | F | T | T |
| Fzd3 | -2.6089099 | 6.3134226 | 94.7128287 | 3.53E-14 | 6.90E-13 | F | F | T | T |
| Rgs19 | -4.2761101 | 4.30929155 | 94.6590686 | 3.57E-14 | 6.97E-13 | F | F | T | F |
| Rab11fip3 | -2.635178 | 6.11556647 | 94.6429682 | 3.58E-14 | 6.99E-13 | F | F | T | T |
| Pdk3 | -3.2882267 | 5.08061424 | 94.6299965 | 3.59E-14 | 7.00E-13 | F | F | F | T |
| Jakmip2 | -10.941152 | 3.35849042 | 94.5671281 | 3.63E-14 | 7.08E-13 | T | F | T | F |
| Fgd5 | -4.1457186 | 4.44275783 | 94.5425994 | 3.65E-14 | 7.11E-13 | F | F | F | F |
| Tedc1 | -10.94126 | 3.35854181 | 94.408092 | 3.75E-14 | 7.30E-13 | T | F | F | F |
| Foxo1 | -3.4380926 | 5.05176277 | 94.3241281 | 3.82E-14 | 7.42E-13 | T | F | T | T |
| Nipa2 | 2.10275434 | 6.41734153 | 94.2558041 | 3.87E-14 | 7.51E-13 | F | F | F | F |
| N4bp2 | 1.70925749 | 8.39666347 | 94.2341491 | 3.89E-14 | 7.53E-13 | F | F | F | T |
| Rtl8b | -2.5997808 | 6.18690307 | 94.2341293 | 3.89E-14 | 7.53E-13 | T | F | F | F |
| Bmpr2 | 1.86948692 | 7.17814488 | 94.1898708 | 3.92E-14 | 7.59E-13 | F | F | T | T |
| Dclk1 | -11.208954 | 3.62093704 | 94.185381 | 3.93E-14 | 7.59E-13 | T | F | T | T |
| Cdh1 | -1.9868068 | 8.62747448 | 94.08508 | 4.01E-14 | 7.74E-13 | T | F | F | F |
| Arhgap9 | 4.48549085 | 3.29394048 | 94.0748287 | 4.02E-14 | 7.75E-13 | F | F | F | F |
| Tmppe | 2.41878196 | 5.34932345 | 93.8538375 | 4.20E-14 | 8.10E-13 | F | F | T | T |
| Dkk1 | -6.7326052 | 3.6352363 | 93.8136636 | 4.23E-14 | 8.16E-13 | T | F | F | F |
| Pgap1 | -2.9351367 | 5.67583208 | 93.8078359 | 4.24E-14 | 8.16E-13 | F | T | T | T |
| Emb | -1.8148363 | 9.48981801 | 93.7797079 | 4.26E-14 | 8.20E-13 | F | F | F | F |
| C5ar2 | 4.503842 | 3.64445846 | 93.6682418 | 4.36E-14 | 8.38E-13 | T | T | F | T |
| Zfp212 | -3.8092063 | 4.61626008 | 93.5351267 | 4.48E-14 | 8.61E-13 | F | F | F | F |
| Rftn1 | -6.1100737 | 3.73678229 | 93.4511238 | 4.56E-14 | 8.75E-13 | F | F | F | T |
| Shmt2 | 1.7571086 | 7.87251863 | 93.4157288 | 4.59E-14 | 8.80E-13 | F | F | T | T |

| | | | | | | | | | |
|---|---|---|---|---|---|---|---|---|---|
| Mgat3 | -4.3245177 | 4.28320618 | 93.3266507 | 4.67E-14 | 8.95E-13 | F | F | T | F |
| Cdk18 | -3.9045349 | 4.52074834 | 93.2818571 | 4.72E-14 | 9.03E-13 | F | F | T | F |
| Cdon | -2.9550734 | 5.46106294 | 93.0975535 | 4.90E-14 | 9.37E-13 | T | F | T | T |
| Vegfb | -3.6155987 | 4.73260344 | 92.8576818 | 5.14E-14 | 9.83E-13 | F | F | F | T |
| Ippk | 2.71754251 | 4.85413947 | 92.7793061 | 5.23E-14 | 9.98E-13 | F | F | F | F |
| Stx2 | 2.74066797 | 4.75510339 | 92.724533 | 5.29E-14 | 1.01E-12 | T | F | T | T |
| Nhsl1 | 1.896432 | 6.99547073 | 92.5470412 | 5.48E-14 | 1.04E-12 | T | F | T | F |
| Fam76a | 2.3473612 | 5.52919139 | 92.497315 | 5.54E-14 | 1.05E-12 | F | F | T | F |
| Parva | 2.26266688 | 6.21840019 | 92.4323767 | 5.61E-14 | 1.07E-12 | T | F | T | T |
| Usp13 | -5.5588607 | 3.85573869 | 92.0863029 | 6.02E-14 | 1.14E-12 | T | F | F | T |
| Cyp26a1 | -4.4367223 | 4.53562285 | 91.9826888 | 6.15E-14 | 1.17E-12 | T | T | F | T |
| Tet2 | 2.40762255 | 5.44780577 | 91.8832865 | 6.28E-14 | 1.19E-12 | T | T | T | T |
| Fam171a2 | -5.5330793 | 3.83083202 | 91.2539905 | 7.15E-14 | 1.36E-12 | F | F | T | F |
| Fer | 2.35671863 | 5.44131076 | 91.2385412 | 7.17E-14 | 1.36E-12 | F | F | F | T |
| Top1mt | 2.82108356 | 4.58913196 | 91.1877821 | 7.25E-14 | 1.37E-12 | F | F | F | T |
| Fnbp1l | -2.3887288 | 6.66377835 | 90.8052305 | 7.85E-14 | 1.48E-12 | T | F | T | F |
| Slc31a2 | 3.15170738 | 4.18586004 | 90.7205779 | 7.99E-14 | 1.51E-12 | F | F | F | F |
| Nab1 | 2.64888385 | 4.83289373 | 90.6222739 | 8.15E-14 | 1.54E-12 | T | F | T | T |
| Bcar3 | 2.24994776 | 5.71005739 | 90.591615 | 8.20E-14 | 1.55E-12 | F | F | F | F |
| Nsd3 | 1.83176811 | 7.11992674 | 90.4804232 | 8.39E-14 | 1.58E-12 | T | F | T | F |
| Arhgef15 | -5.2899266 | 3.87997695 | 90.1086986 | 9.07E-14 | 1.71E-12 | T | F | F | T |
| Ildr2 | -10.874646 | 3.29555026 | 90.0769497 | 9.13E-14 | 1.72E-12 | F | T | T | T |
| Crispld1 | -10.839471 | 3.25951027 | 90.064759 | 9.15E-14 | 1.72E-12 | T | F | T | T |
| Elovl2 | -5.5274691 | 3.82563663 | 89.9215837 | 9.43E-14 | 1.77E-12 | F | F | F | F |
| Dll4 | 3.90553032 | 3.94762983 | 89.8800761 | 9.51E-14 | 1.78E-12 | F | F | F | F |
| Ninl | -3.1060624 | 5.17368079 | 89.7763211 | 9.72E-14 | 1.82E-12 | F | F | F | F |
| Cdx1 | -6.5344859 | 6.72680166 | 89.6973158 | 9.88E-14 | 1.85E-12 | F | F | T | F |
| Bcorl1 | -2.6030231 | 6.21173628 | 89.6576311 | 9.96E-14 | 1.86E-12 | F | F | F | F |
| AA467197 | 7.3047576 | 2.27450213 | 89.6223769 | 1.00E-13 | 1.88E-12 | F | F | F | F |
| Sgk1 | -2.7735007 | 5.65526154 | 89.617795 | 1.00E-13 | 1.88E-12 | T | F | F | T |
| Nptx2 | -3.3899122 | 5.05249876 | 89.6136246 | 1.01E-13 | 1.88E-12 | F | F | T | F |
| Ubqln2 | -1.9900672 | 7.80050301 | 89.4134376 | 1.05E-13 | 1.95E-12 | F | F | F | F |
| Llgl1 | -2.0900654 | 7.28094219 | 89.2783518 | 1.08E-13 | 2.01E-12 | F | F | F | F |
| Sept9 | 1.745801 | 8.91289745 | 89.2022212 | 1.10E-13 | 2.04E-12 | F | F | T | F |
| Osbpl6 | -10.834462 | 3.25562007 | 89.0981502 | 1.12E-13 | 2.08E-12 | T | F | T | T |
| Grip1 | -3.6734648 | 4.60694376 | 89.0730968 | 1.13E-13 | 2.09E-12 | T | F | F | F |
| Mesp1 | -6.9367063 | 3.68892741 | 89.0448852 | 1.13E-13 | 2.10E-12 | F | F | F | F |
| Lgals8 | 4.27052017 | 3.29200717 | 88.9543577 | 1.15E-13 | 2.14E-12 | F | F | T | F |
| Ada | 3.1833216 | 4.19326362 | 88.9102795 | 1.16E-13 | 2.16E-12 | T | F | T | F |
| Cacna1a | -10.866688 | 3.28476133 | 88.8447176 | 1.18E-13 | 2.19E-12 | F | F | F | T |
| Slc22a23 | -2.6071333 | 6.07896888 | 88.7091135 | 1.21E-13 | 2.25E-12 | T | F | T | T |
| B4galt1 | 1.85362136 | 6.97527892 | 88.6944132 | 1.22E-13 | 2.25E-12 | T | F | F | F |

| | | | | | | | | | |
|---|---|---|---|---|---|---|---|---|---|
| Myh10 | -1.8082002 | 8.93850807 | 88.6597261 | 1.23E-13 | 2.27E-12 | T | F | T | F |
| St8sia3 | -6.7382372 | 3.52924822 | 88.639529 | 1.23E-13 | 2.27E-12 | T | F | T | T |
| Tbc1d4 | -10.830606 | 3.24991428 | 88.5397626 | 1.26E-13 | 2.32E-12 | F | F | T | F |
| Rnf130 | -3.438313 | 4.77432336 | 88.4428361 | 1.28E-13 | 2.37E-12 | T | F | T | T |
| Hsd11b2 | -5.7833966 | 3.82148335 | 88.4109833 | 1.29E-13 | 2.38E-12 | F | F | F | F |
| Krt18 | 1.63610211 | 8.99996114 | 88.3690211 | 1.30E-13 | 2.40E-12 | F | F | F | F |
| Mapk4 | -6.3022584 | 3.62040535 | 88.3099632 | 1.32E-13 | 2.43E-12 | T | F | F | T |
| Gli1 | -5.075569 | 4.10315169 | 88.2666093 | 1.33E-13 | 2.45E-12 | T | F | F | F |
| Cyp11a1 | 3.88194553 | 3.61078667 | 88.1704285 | 1.36E-13 | 2.50E-12 | T | F | F | F |
| Zbtb16 | -3.7090229 | 4.56397921 | 87.9501159 | 1.43E-13 | 2.61E-12 | T | F | T | T |
| Flna | -1.6880364 | 9.85880297 | 87.9052818 | 1.44E-13 | 2.64E-12 | F | F | F | F |
| Prdm5 | -6.2813963 | 3.60013825 | 87.7227074 | 1.50E-13 | 2.74E-12 | T | F | F | T |
| Caskin1 | 3.00412124 | 4.35525653 | 87.7211925 | 1.50E-13 | 2.74E-12 | F | F | T | F |
| Slc30a1 | 1.92754299 | 6.69295349 | 87.6526132 | 1.52E-13 | 2.77E-12 | T | F | T | T |
| Lin9 | -10.85079 | 3.26902005 | 87.6396702 | 1.52E-13 | 2.78E-12 | T | T | T | T |
| Cts8 | 3.82308912 | 3.61202744 | 87.554967 | 1.55E-13 | 2.83E-12 | T | F | F | F |
| Tacc1 | 1.88006843 | 6.83442747 | 87.3953737 | 1.60E-13 | 2.92E-12 | T | F | T | T |
| Cxcr4 | -4.0100091 | 4.31218549 | 87.3348985 | 1.62E-13 | 2.96E-12 | T | F | F | F |
| Tnrc18 | 1.68543128 | 8.09267503 | 87.2895186 | 1.64E-13 | 2.98E-12 | T | F | F | F |
| Sema3e | 2.45463234 | 5.32653196 | 87.1219234 | 1.70E-13 | 3.09E-12 | F | F | T | F |
| Prl6a1 | 8.37581687 | 1.99124887 | 87.0423179 | 1.73E-13 | 3.14E-12 | F | F | F | F |
| Unc93b1 | 4.00981937 | 3.50110015 | 87.0397357 | 1.73E-13 | 3.14E-12 | T | F | T | F |
| Eps8 | 2.03246773 | 6.28553062 | 87.0057447 | 1.74E-13 | 3.16E-12 | F | F | F | T |
| Foxn3 | -2.2310424 | 6.88687968 | 86.8328291 | 1.81E-13 | 3.27E-12 | T | F | T | T |
| Bicd2 | 1.92371559 | 6.67047807 | 86.5639828 | 1.91E-13 | 3.46E-12 | F | F | T | F |
| Zfp105 | -3.9017556 | 4.37607114 | 86.4600426 | 1.96E-13 | 3.53E-12 | F | F | F | F |
| Prickle1 | -2.9320313 | 5.70489292 | 86.4594629 | 1.96E-13 | 3.53E-12 | T | F | T | F |
| Shc2 | -4.5112783 | 4.12134113 | 86.4291181 | 1.97E-13 | 3.55E-12 | T | F | T | T |
| Erlin1 | 2.34739838 | 5.5879632 | 86.3368637 | 2.01E-13 | 3.62E-12 | T | T | T | T |
| Bcl7a | -2.8711578 | 5.35724288 | 86.1960772 | 2.07E-13 | 3.73E-12 | T | F | T | T |
| Park7 | -2.3012986 | 6.57284134 | 86.1772769 | 2.08E-13 | 3.74E-12 | F | F | F | F |
| Gdpd5 | 2.31585098 | 5.35552686 | 86.0201769 | 2.15E-13 | 3.87E-12 | T | F | F | T |
| Cdca7 | -2.0432058 | 7.33943981 | 85.9957212 | 2.16E-13 | 3.88E-12 | F | F | F | F |
| Dcbld1 | -4.1350022 | 4.22981199 | 85.9429138 | 2.18E-13 | 3.92E-12 | T | F | F | F |
| Filip1l | 4.45859625 | 3.18255464 | 85.9359896 | 2.19E-13 | 3.93E-12 | F | F | F | T |
| Xkr5 | -10.766888 | 3.18806226 | 85.9174322 | 2.20E-13 | 3.94E-12 | F | F | F | F |
| Col9a3 | -10.833624 | 3.25188478 | 85.8358358 | 2.23E-13 | 4.00E-12 | T | F | F | F |
| Cast | 2.11086506 | 5.9612746 | 85.6761802 | 2.31E-13 | 4.14E-12 | T | F | T | T |
| Asb4 | -4.6012868 | 4.11757046 | 85.6447151 | 2.33E-13 | 4.16E-12 | T | F | F | F |
| Brsk1 | -3.82619 | 4.40574745 | 85.5033979 | 2.40E-13 | 4.29E-12 | F | F | F | F |
| Gm14698 | 10.4146302 | 1.8069982 | 86.1673304 | 2.41E-13 | 4.31E-12 | F | F | F | T |
| Actr2 | 1.6158419 | 8.4146201 | 85.4517859 | 2.43E-13 | 4.33E-12 | T | T | T | T |

| | | | | | | | | | |
|---|---|---|---|---|---|---|---|---|---|
| Tfap2a | 2.24455959 | 5.53227793 | 85.4466589 | 2.43E-13 | 4.33E-12 | F | F | T | F |
| Trip6 | -2.6703521 | 5.68658208 | 85.4274424 | 2.44E-13 | 4.35E-12 | F | F | F | F |
| Lrrn4 | -2.9396918 | 5.23188587 | 85.3954174 | 2.46E-13 | 4.37E-12 | T | F | F | T |
| Chst2 | 2.16288692 | 6.04492906 | 85.3347828 | 2.49E-13 | 4.43E-12 | T | F | T | T |
| Sap30 | -3.4084215 | 4.72636439 | 85.3266028 | 2.49E-13 | 4.43E-12 | F | F | T | F |
| Prss12 | -4.2462091 | 4.20750854 | 85.3211188 | 2.50E-13 | 4.43E-12 | F | F | F | T |
| Gm12166 | 10.4111487 | 1.80354322 | 85.976283 | 2.51E-13 | 4.46E-12 | F | F | F | F |
| Rcbtb2 | -2.4133331 | 6.25298251 | 85.2215977 | 2.55E-13 | 4.52E-12 | T | F | F | F |
| Scarb2 | 2.14241006 | 5.84928732 | 85.1675182 | 2.58E-13 | 4.57E-12 | F | F | T | F |
| Rxra | 1.82100071 | 6.9721844 | 85.1515549 | 2.59E-13 | 4.58E-12 | T | F | F | F |
| Spon1 | -5.3098071 | 3.82848608 | 85.1286972 | 2.60E-13 | 4.60E-12 | T | F | T | T |
| Rnf219 | -2.8636463 | 5.32698743 | 85.0670595 | 2.64E-13 | 4.65E-12 | T | F | F | F |
| Syngr2 | 1.86050556 | 6.81070606 | 85.0655906 | 2.64E-13 | 4.65E-12 | F | F | T | F |
| Sfmbt1 | -3.0289485 | 5.11758944 | 85.0268486 | 2.66E-13 | 4.69E-12 | F | F | T | T |
| Tox3 | -7.5559657 | 3.38090863 | 84.9798102 | 2.69E-13 | 4.73E-12 | T | F | T | T |
| Spry1 | -4.696912 | 3.99233646 | 84.9397198 | 2.71E-13 | 4.77E-12 | F | F | F | T |
| Fads2 | -2.3538573 | 6.38414473 | 84.7517307 | 2.82E-13 | 4.96E-12 | T | F | T | F |
| Nasp | -1.7626067 | 8.93284445 | 84.6831812 | 2.86E-13 | 5.03E-12 | T | F | F | T |
| Colgalt1 | 1.84113155 | 6.89634335 | 84.6576443 | 2.88E-13 | 5.05E-12 | F | F | T | T |
| Camkv | -6.229469 | 3.54927044 | 84.6568722 | 2.88E-13 | 5.05E-12 | T | F | T | F |
| Foxd4 | -10.791126 | 3.21408495 | 84.6253019 | 2.90E-13 | 5.09E-12 | F | F | F | F |
| Vcl | 1.63080249 | 8.15777248 | 84.6070823 | 2.91E-13 | 5.10E-12 | T | F | T | T |
| Dab1 | -5.0094297 | 3.88950507 | 84.4990417 | 2.98E-13 | 5.22E-12 | T | F | T | F |
| Met | -3.738807 | 4.50622738 | 84.3894154 | 3.05E-13 | 5.34E-12 | T | F | T | T |
| Gfod1 | 3.63759587 | 3.71056303 | 84.3835972 | 3.05E-13 | 5.34E-12 | T | F | F | F |
| Tmprss4 | 10.3928115 | 1.78534842 | 84.9751987 | 3.11E-13 | 5.43E-12 | F | F | T | F |
| Arhgef7 | 1.77736302 | 7.0885589 | 84.2922298 | 3.12E-13 | 5.44E-12 | T | F | T | F |
| Wdr62 | 2.33170207 | 5.93438937 | 84.2771218 | 3.13E-13 | 5.45E-12 | F | F | F | F |
| Tcn2 | -3.4588142 | 4.86036771 | 84.2049449 | 3.17E-13 | 5.53E-12 | T | F | T | T |
| Arl4c | -2.3395949 | 6.44562896 | 84.1761141 | 3.19E-13 | 5.56E-12 | F | F | T | F |
| Cnn2 | 1.82570436 | 7.26207809 | 84.077472 | 3.26E-13 | 5.68E-12 | T | F | T | T |
| Hnf1b | -3.110118 | 5.00476023 | 84.0497021 | 3.28E-13 | 5.71E-12 | T | F | T | F |
| Col5a2 | -3.4947621 | 4.68369967 | 83.7073134 | 3.54E-13 | 6.14E-12 | F | F | F | T |
| Zfp354c | -3.5748706 | 4.55091726 | 83.7009201 | 3.54E-13 | 6.15E-12 | T | F | T | T |
| Cdhr5 | 4.2450599 | 3.60463364 | 83.6666728 | 3.57E-13 | 6.19E-12 | F | F | F | F |
| Dsg2 | 1.66495433 | 7.80599351 | 83.5622603 | 3.65E-13 | 6.33E-12 | T | F | T | F |
| Rsf1 | -2.9327186 | 5.19481537 | 83.5123312 | 3.69E-13 | 6.39E-12 | F | T | T | T |
| Slain1 | -6.6712742 | 3.43036393 | 83.4354457 | 3.75E-13 | 6.49E-12 | T | F | F | F |
| Fgf4 | -10.718733 | 3.1409911 | 83.4124015 | 3.77E-13 | 6.52E-12 | F | F | T | F |
| Vash2 | -2.8252532 | 5.33972171 | 83.3896601 | 3.79E-13 | 6.55E-12 | F | F | T | F |
| Chst7 | -6.6880979 | 3.44733349 | 83.3814115 | 3.80E-13 | 6.55E-12 | T | F | T | T |
| Ralb | 1.97303563 | 6.45352376 | 83.2656538 | 3.89E-13 | 6.72E-12 | F | F | T | F |

| | | | | | | | | | |
|---|---|---|---|---|---|---|---|---|---|
| Slc4a4 | -3.2477848 | 4.83325374 | 83.2406293 | 3.91E-13 | 6.75E-12 | T | F | T | T |
| Cadm3 | -7.5320715 | 3.36015499 | 83.2221921 | 3.93E-13 | 6.77E-12 | T | F | F | T |
| Shisa2 | -2.8359074 | 5.33617693 | 83.178016 | 3.97E-13 | 6.83E-12 | T | F | F | T |
| Mn1 | -2.3369858 | 6.35829836 | 83.1652306 | 3.98E-13 | 6.84E-12 | T | F | T | F |
| Lrp10 | 1.95951532 | 6.86105451 | 83.0098706 | 4.12E-13 | 7.07E-12 | F | F | F | F |
| Mfap2 | -7.5155001 | 3.34375247 | 82.9537941 | 4.17E-13 | 7.15E-12 | F | F | F | F |
| Atf6 | 1.98309657 | 6.3204011 | 82.9510709 | 4.17E-13 | 7.15E-12 | T | F | T | T |
| Serinc2 | 2.9825662 | 4.21236346 | 82.8327653 | 4.28E-13 | 7.33E-12 | F | F | F | F |
| Tcf7 | -2.3539181 | 6.30068325 | 82.7906531 | 4.32E-13 | 7.40E-12 | F | F | F | F |
| Sel1l3 | -3.8699379 | 4.33754786 | 82.7115849 | 4.39E-13 | 7.52E-12 | T | F | T | F |
| Stxbp6 | -3.8481623 | 4.34174963 | 82.7100538 | 4.40E-13 | 7.52E-12 | T | T | T | T |
| Lrp4 | -2.6812843 | 5.56646665 | 82.557734 | 4.54E-13 | 7.77E-12 | T | T | T | T |
| Slc35a3 | 2.10040197 | 5.90644896 | 82.5287953 | 4.57E-13 | 7.81E-12 | T | F | T | T |
| Col3a1 | 3.77665516 | 3.51843042 | 82.258234 | 4.85E-13 | 8.28E-12 | T | F | T | F |
| Hexa | -3.05925 | 5.03536333 | 82.2194446 | 4.89E-13 | 8.34E-12 | F | F | F | F |
| Ncam1 | 1.98254752 | 6.40533886 | 82.1866309 | 4.93E-13 | 8.40E-12 | T | F | T | T |
| Snai1 | 2.00356721 | 6.20464903 | 82.1549773 | 4.96E-13 | 8.45E-12 | F | F | F | F |
| Tmem145 | -10.748714 | 3.16857456 | 82.1049155 | 5.02E-13 | 8.54E-12 | F | F | F | F |
| Gkap1 | 2.44886818 | 4.95742079 | 82.0154933 | 5.12E-13 | 8.70E-12 | T | F | F | F |
| Ocln | -4.9599361 | 3.84144121 | 81.9320354 | 5.21E-13 | 8.85E-12 | T | F | T | F |
| Trp53bp2 | -2.4430385 | 6.04194984 | 81.7241598 | 5.46E-13 | 9.26E-12 | T | F | T | F |
| Zfp618 | -3.3567366 | 4.71377343 | 81.6668185 | 5.53E-13 | 9.37E-12 | T | F | T | F |
| Serpinb9d | 8.27848407 | 1.89450697 | 81.631452 | 5.57E-13 | 9.44E-12 | F | F | F | F |
| Nbeal1 | -2.6081712 | 5.8112135 | 81.5578291 | 5.66E-13 | 9.58E-12 | T | F | T | T |
| Rufy4 | 7.57578848 | 2.02389369 | 81.5522349 | 5.67E-13 | 9.59E-12 | T | F | F | T |
| Creb3 | 2.4390013 | 5.29611625 | 81.4947479 | 5.74E-13 | 9.70E-12 | F | F | F | F |
| Arid1b | -1.9917567 | 7.23165478 | 81.4108742 | 5.85E-13 | 9.88E-12 | T | F | T | T |
| Grb10 | -1.7004838 | 9.33188869 | 81.3936433 | 5.87E-13 | 9.91E-12 | T | F | T | T |
| Atp11a | 1.96115297 | 6.30514546 | 81.3734507 | 5.90E-13 | 9.94E-12 | F | F | F | F |
| Epha1 | -2.3602441 | 6.23178975 | 81.2686547 | 6.04E-13 | 1.02E-11 | F | F | F | F |
| Shroom3 | -2.1454606 | 6.79883013 | 81.2412012 | 6.07E-13 | 1.02E-11 | T | F | T | T |
| Prr13 | 2.40337776 | 5.04982762 | 81.2091828 | 6.11E-13 | 1.03E-11 | T | F | T | T |
| Pgm1 | 1.80330058 | 6.86740875 | 81.0541816 | 6.33E-13 | 1.06E-11 | T | F | F | F |
| Sgk2 | 4.88984133 | 2.95879307 | 80.9077761 | 6.54E-13 | 1.10E-11 | F | F | F | T |
| Pygb | 1.71493338 | 7.25808739 | 80.8990499 | 6.55E-13 | 1.10E-11 | F | F | F | T |
| Fgd1 | -2.3148565 | 6.31718606 | 80.7917198 | 6.71E-13 | 1.12E-11 | F | F | F | F |
| Ctsb | 1.57831746 | 8.39430452 | 80.7876498 | 6.71E-13 | 1.12E-11 | T | F | T | T |
| Pgghg | -10.673033 | 3.09574598 | 80.7805466 | 6.72E-13 | 1.13E-11 | T | F | F | F |
| Rnf217 | -4.4131818 | 4.03320927 | 80.7309158 | 6.80E-13 | 1.14E-11 | F | F | T | T |
| Grk3 | -5.5600727 | 3.64155165 | 80.6896769 | 6.86E-13 | 1.15E-11 | T | F | T | F |
| Gdf1 | -10.739333 | 3.16389622 | 80.5850286 | 7.02E-13 | 1.17E-11 | T | F | F | F |
| B3gnt5 | -2.9541338 | 5.08350194 | 80.5821669 | 7.03E-13 | 1.17E-11 | T | F | T | T |

| | | | | | | | | | |
|---|---|---|---|---|---|---|---|---|---|
| Nnt | -2.6567965 | 5.53711921 | 80.5628871 | 7.06E-13 | 1.18E-11 | T | F | T | T |
| Kdm1b | 2.22956534 | 5.37282136 | 80.5092359 | 7.14E-13 | 1.19E-11 | T | F | T | T |
| Rin1 | 2.48470001 | 4.89817996 | 80.3863723 | 7.34E-13 | 1.22E-11 | T | F | F | T |
| Optn | 2.75598255 | 4.45217071 | 80.3698416 | 7.37E-13 | 1.23E-11 | F | F | T | T |
| Elavl2 | -6.6677402 | 3.42521204 | 80.2421353 | 7.58E-13 | 1.26E-11 | F | F | T | T |
| Irf7 | 6.89178002 | 2.18679687 | 80.1807218 | 7.68E-13 | 1.28E-11 | F | F | F | F |
| Zyx | 1.6020786 | 8.0202996 | 80.1693757 | 7.70E-13 | 1.28E-11 | T | F | T | T |
| Esyt3 | 2.96644969 | 4.17679471 | 80.1381706 | 7.76E-13 | 1.29E-11 | T | F | T | T |
| Gpr153 | -10.675117 | 3.10001606 | 80.0672809 | 7.88E-13 | 1.31E-11 | T | F | T | F |
| Apobec3 | -10.662574 | 3.08524138 | 80.0206351 | 7.96E-13 | 1.32E-11 | T | F | T | T |
| Oat | -1.9663392 | 7.34413042 | 79.8615552 | 8.25E-13 | 1.37E-11 | T | T | T | T |
| Sesn2 | 1.87509678 | 6.6067576 | 79.7404128 | 8.48E-13 | 1.40E-11 | F | F | F | T |
| Nlgn3 | -3.6687615 | 4.36484952 | 79.7224471 | 8.51E-13 | 1.41E-11 | F | F | T | T |
| Trp53inp1 | -10.685677 | 3.10688487 | 79.5492358 | 8.85E-13 | 1.46E-11 | T | F | F | F |
| Zfp647 | -5.0445808 | 3.75803553 | 79.5036507 | 8.94E-13 | 1.47E-11 | T | F | T | T |
| Armcx3 | -7.4222431 | 3.25150762 | 79.4965541 | 8.95E-13 | 1.48E-11 | T | F | T | T |
| Prlr | 2.7448328 | 4.40345618 | 79.3845144 | 9.18E-13 | 1.51E-11 | T | F | T | T |
| Homer2 | 2.34436139 | 5.1658218 | 79.2327717 | 9.50E-13 | 1.56E-11 | T | T | T | T |
| Hmgn1 | -1.724914 | 8.80660907 | 79.0499598 | 9.89E-13 | 1.63E-11 | T | F | T | F |
| Vstm4 | 4.88457083 | 2.81211205 | 79.0170221 | 9.97E-13 | 1.64E-11 | F | F | F | F |
| Soat2 | -2.8576151 | 5.14947392 | 78.9890467 | 1.00E-12 | 1.65E-11 | F | F | T | F |
| Tnpo2 | 1.72706363 | 7.11125761 | 78.9619958 | 1.01E-12 | 1.66E-11 | T | F | T | F |
| Pcsk6 | 2.58012175 | 4.66279027 | 78.8161527 | 1.04E-12 | 1.71E-11 | T | F | F | F |
| Atf7 | 2.00799753 | 6.01991289 | 78.7769254 | 1.05E-12 | 1.72E-11 | T | F | T | T |
| Rnf122 | -7.3794411 | 3.23861673 | 78.7004485 | 1.07E-12 | 1.75E-11 | T | F | T | T |
| Trim24 | -2.4600694 | 5.87702233 | 78.6457969 | 1.08E-12 | 1.77E-11 | T | F | F | T |
| Gprasp2 | -10.627028 | 3.05219113 | 78.5466199 | 1.11E-12 | 1.81E-11 | F | F | F | T |
| Med12 | -1.958759 | 7.18778406 | 78.5428332 | 1.11E-12 | 1.81E-11 | T | T | F | T |
| Pxylp1 | -6.4999974 | 3.37185856 | 78.4057058 | 1.14E-12 | 1.87E-11 | T | F | T | F |
| Hapln4 | 4.95189569 | 2.77295718 | 78.3542037 | 1.16E-12 | 1.89E-11 | F | F | F | F |
| Klf12 | -4.6625223 | 3.95845923 | 78.3443081 | 1.16E-12 | 1.89E-11 | T | F | T | T |
| Mob3c | 2.74286656 | 4.43787997 | 78.2780454 | 1.18E-12 | 1.92E-11 | T | F | T | T |
| Heyl | 6.06184184 | 2.52468139 | 78.2308813 | 1.19E-12 | 1.94E-11 | T | F | F | T |
| Tpbg | -3.277662 | 4.68819884 | 78.2128042 | 1.19E-12 | 1.94E-11 | T | F | F | F |
| Lrp1 | -1.7464768 | 8.58140988 | 78.172854 | 1.21E-12 | 1.96E-11 | T | T | F | T |
| Rtl8c | -10.625852 | 3.05143232 | 78.1719361 | 1.21E-12 | 1.96E-11 | F | F | F | F |
| Pbx3 | -2.8051212 | 5.2398336 | 78.1413896 | 1.21E-12 | 1.97E-11 | T | F | T | T |
| Jak2 | 2.2817195 | 5.1659442 | 77.9781056 | 1.26E-12 | 2.04E-11 | F | F | T | F |
| Ccdc8 | -2.6902986 | 5.37043002 | 77.9721975 | 1.26E-12 | 2.04E-11 | F | F | F | F |
| St3gal4 | 2.79561086 | 4.30751184 | 77.8805619 | 1.29E-12 | 2.09E-11 | T | F | T | F |
| Itgb1 | 1.54371079 | 8.38481627 | 77.8502525 | 1.30E-12 | 2.10E-11 | T | F | T | T |
| Lpcat4 | 2.01022534 | 5.99535612 | 77.7742441 | 1.32E-12 | 2.13E-11 | F | F | F | T |

| | | | | | | | | | |
|---|---|---|---|---|---|---|---|---|---|
| Pla2g4d | 8.20571829 | 1.82227253 | 77.7458766 | 1.33E-12 | 2.15E-11 | T | F | F | F |
| Ifitm2 | -2.4293598 | 5.89830479 | 77.6714346 | 1.35E-12 | 2.18E-11 | F | F | F | T |
| Camta2 | 2.11802772 | 5.57761584 | 77.4830831 | 1.41E-12 | 2.27E-11 | F | F | F | F |
| Ppp1r13l | 2.29934072 | 5.1087263 | 77.4320029 | 1.43E-12 | 2.30E-11 | F | F | F | F |
| Cabp7 | -7.4275826 | 3.25467728 | 77.364643 | 1.45E-12 | 2.33E-11 | F | F | T | T |
| Stxbp5 | 1.93062259 | 6.24245789 | 77.3354458 | 1.46E-12 | 2.35E-11 | T | F | T | T |
| Acadvl | 2.39473718 | 4.9831702 | 77.3114532 | 1.47E-12 | 2.36E-11 | F | F | F | F |
| Copz2 | 4.17436496 | 3.23366608 | 77.2565182 | 1.48E-12 | 2.39E-11 | F | F | F | F |
| Actr3 | 1.55402628 | 8.20751658 | 77.2094776 | 1.50E-12 | 2.41E-11 | T | T | T | T |
| Plk2 | 2.8820184 | 4.47821819 | 77.198873 | 1.50E-12 | 2.41E-11 | F | F | F | F |
| Tubb2b | -3.0754113 | 5.18831563 | 77.180078 | 1.51E-12 | 2.42E-11 | F | F | F | T |
| Rbm15b | -1.8636958 | 7.66477314 | 77.1085739 | 1.53E-12 | 2.46E-11 | T | F | F | T |
| Clptm1l | 1.84327593 | 6.5794975 | 77.017563 | 1.57E-12 | 2.51E-11 | F | F | F | F |
| Cdc42ep3 | 3.84693594 | 3.37871947 | 77.0076925 | 1.57E-12 | 2.51E-11 | F | F | F | T |
| Arhgap31 | 2.5166655 | 5.26850064 | 76.970293 | 1.58E-12 | 2.53E-11 | T | T | T | T |
| Fam219a | 3.03224662 | 4.05738665 | 76.9458013 | 1.59E-12 | 2.54E-11 | F | F | T | F |
| Aacs | 2.2579442 | 5.18267789 | 76.7567472 | 1.66E-12 | 2.66E-11 | F | F | T | T |
| Micall2 | 3.10585111 | 3.9830797 | 76.5648518 | 1.74E-12 | 2.77E-11 | F | F | F | F |
| S100a16 | 3.28131296 | 3.89195509 | 76.5644238 | 1.74E-12 | 2.77E-11 | F | F | T | F |
| H1f6 | -4.508517 | 3.92263704 | 76.5379305 | 1.75E-12 | 2.79E-11 | F | F | F | F |
| Col13a1 | -6.0818687 | 3.40485352 | 76.5253235 | 1.75E-12 | 2.79E-11 | T | F | F | T |
| Tnfrsf1a | 2.06594965 | 5.72887076 | 76.4995219 | 1.76E-12 | 2.81E-11 | F | F | T | F |
| Grrp1 | -7.3770434 | 3.20596957 | 76.4698807 | 1.78E-12 | 2.82E-11 | T | F | F | T |
| Zfp358 | -3.9589702 | 4.13940792 | 76.4321447 | 1.79E-12 | 2.85E-11 | F | F | F | F |
| Cpxm1 | -6.0903969 | 3.41470897 | 76.3869862 | 1.81E-12 | 2.87E-11 | T | F | F | T |
| Klhl42 | 2.13115146 | 5.52577593 | 76.2895581 | 1.85E-12 | 2.94E-11 | T | F | T | T |
| Bend4 | -2.2069024 | 6.47847985 | 76.2386112 | 1.87E-12 | 2.97E-11 | F | T | T | T |
| Gm14226 | -10.725197 | 3.15104382 | 76.2108704 | 1.88E-12 | 2.98E-11 | F | F | F | F |
| Gnas | -1.6886201 | 8.76580725 | 76.1997632 | 1.89E-12 | 2.99E-11 | T | F | T | F |
| Ankrd13b | -6.2934746 | 3.3566571 | 76.1331304 | 1.92E-12 | 3.03E-11 | T | F | T | T |
| Kdm5b | -1.8064281 | 8.37302633 | 76.0904882 | 1.94E-12 | 3.06E-11 | T | F | T | T |
| Cystm1 | 2.89741284 | 4.16783173 | 76.0692894 | 1.95E-12 | 3.07E-11 | F | F | F | T |
| Prkd3 | -2.3870648 | 5.92378895 | 76.0465875 | 1.96E-12 | 3.09E-11 | T | F | T | F |
| Dut | -1.8708929 | 7.48086434 | 75.8893097 | 2.03E-12 | 3.20E-11 | T | F | F | T |
| Atp13a3 | 1.54614423 | 8.31397145 | 75.7705517 | 2.08E-12 | 3.28E-11 | F | F | F | T |
| Reep3 | 1.7805861 | 6.7292989 | 75.7658348 | 2.09E-12 | 3.29E-11 | T | F | T | T |
| Pip5k1c | 1.63225433 | 7.73571956 | 75.7527709 | 2.09E-12 | 3.29E-11 | F | F | F | F |
| Fam120a | 1.51873476 | 8.42681343 | 75.7479587 | 2.10E-12 | 3.29E-11 | F | F | T | F |
| Ric3 | -10.588964 | 3.01601864 | 75.6601689 | 2.14E-12 | 3.36E-11 | T | F | T | F |
| Nfat5 | 2.68844839 | 7.00804707 | 75.6460365 | 2.15E-12 | 3.37E-11 | T | F | T | T |
| Lgr5 | -10.563169 | 2.98858167 | 75.6004788 | 2.17E-12 | 3.40E-11 | T | F | T | T |
| Cnot6l | 2.00429357 | 5.97387815 | 75.5711098 | 2.18E-12 | 3.42E-11 | F | F | T | T |

| | | | | | | | | | |
|---|---|---|---|---|---|---|---|---|---|
| Sptbn2 | 1.76113763 | 6.83099797 | 75.5692792 | 2.18E-12 | 3.42E-11 | F | F | T | F |
| Tmem144 | 2.43499592 | 4.77991026 | 75.4683204 | 2.23E-12 | 3.50E-11 | F | F | F | T |
| Tspan14 | 1.76974824 | 6.7582235 | 75.4626922 | 2.24E-12 | 3.50E-11 | T | T | T | T |
| Fam129b | -2.3132449 | 6.10922371 | 75.4614881 | 2.24E-12 | 3.50E-11 | T | F | T | T |
| Cnksr3 | -2.781561 | 5.17469272 | 75.4199179 | 2.26E-12 | 3.53E-11 | F | F | F | T |
| Car3 | -7.366982 | 3.19843561 | 75.4074931 | 2.27E-12 | 3.54E-11 | T | F | F | F |
| Armcx1 | -4.0584225 | 4.09939918 | 75.3731796 | 2.28E-12 | 3.56E-11 | T | F | F | F |
| Gabbr1 | -5.7928331 | 3.51846185 | 75.3406149 | 2.30E-12 | 3.59E-11 | T | F | T | T |
| Nedd4l | 2.21929978 | 5.20136588 | 75.3250344 | 2.31E-12 | 3.60E-11 | T | F | T | T |
| Arhgef40 | -4.0973695 | 4.04944613 | 75.2438873 | 2.35E-12 | 3.66E-11 | T | F | T | T |
| Trim2 | -2.6797396 | 5.28046975 | 75.1887547 | 2.38E-12 | 3.71E-11 | T | F | T | T |
| Fgf13 | -10.553421 | 2.9792338 | 75.1841525 | 2.39E-12 | 3.71E-11 | F | F | F | T |
| mt-Co1 | 1.18437943 | 12.203845 | 75.159387 | 2.40E-12 | 3.73E-11 | F | F | F | F |
| Cr1l | 1.92243595 | 6.20865383 | 75.116496 | 2.42E-12 | 3.76E-11 | T | F | F | T |
| Rab4a | -3.2354415 | 4.6209692 | 74.9777995 | 2.50E-12 | 3.88E-11 | F | F | T | F |
| Mmgt2 | 2.7320604 | 4.31597579 | 74.8479078 | 2.58E-12 | 4.00E-11 | F | F | F | F |
| Ostf1 | 2.3864972 | 4.87317164 | 74.7933109 | 2.61E-12 | 4.05E-11 | F | F | F | F |
| Fbxo21 | -1.9178607 | 7.12132058 | 74.7673121 | 2.63E-12 | 4.07E-11 | T | F | F | T |
| Ogt | 1.58291128 | 7.9324272 | 74.6591635 | 2.69E-12 | 4.17E-11 | T | F | T | F |
| Shh | -10.614597 | 3.0421798 | 74.5958498 | 2.73E-12 | 4.23E-11 | F | F | F | F |
| Plce1 | -5.4787247 | 3.52265982 | 74.5925206 | 2.74E-12 | 4.23E-11 | T | F | F | F |
| Gm14124 | 6.13291089 | 2.14098168 | 74.5054312 | 2.79E-12 | 4.31E-11 | F | F | F | F |
| Wiz | -2.2509612 | 6.20605961 | 74.4993353 | 2.80E-12 | 4.31E-11 | F | F | F | F |
| Arid3b | -1.954609 | 7.38760399 | 74.4122461 | 2.85E-12 | 4.40E-11 | T | F | T | T |
| Coch | -6.0651933 | 3.38803025 | 74.3362175 | 2.90E-12 | 4.47E-11 | F | F | F | F |
| Tbc1d2 | 2.69342885 | 4.36931502 | 74.2018632 | 3.00E-12 | 4.61E-11 | T | F | T | F |
| Arhgap32 | -2.412586 | 5.8003753 | 74.1403693 | 3.04E-12 | 4.67E-11 | F | F | T | T |
| Flrt2 | -6.5022206 | 3.26492061 | 74.1261805 | 3.05E-12 | 4.69E-11 | F | F | T | F |
| Crybg1 | 2.21403701 | 5.1766892 | 74.1039841 | 3.07E-12 | 4.71E-11 | F | F | T | F |
| Dnajc22 | -5.4517986 | 3.49716576 | 73.8952553 | 3.22E-12 | 4.94E-11 | F | F | T | T |
| Ints6l | -3.4486374 | 4.38174862 | 73.8500779 | 3.25E-12 | 4.99E-11 | F | F | F | F |
| Cdc14b | -2.3380684 | 5.99284601 | 73.8377191 | 3.26E-12 | 5.00E-11 | T | F | T | T |
| Neb | -10.579201 | 3.00204162 | 73.8190281 | 3.28E-12 | 5.02E-11 | F | F | F | F |
| Myo1f | 7.01470511 | 1.9863593 | 73.781306 | 3.31E-12 | 5.06E-11 | F | F | F | F |
| Rapgef3 | 4.10664251 | 3.09798399 | 73.7768604 | 3.31E-12 | 5.06E-11 | F | F | T | T |
| Zfp362 | -2.7872269 | 5.09480474 | 73.7121468 | 3.36E-12 | 5.13E-11 | F | F | T | F |
| Dmrtc1c2 | 10.1848457 | 1.57934933 | 74.2343737 | 3.37E-12 | 5.14E-11 | F | F | F | F |
| Flt1 | -3.4784807 | 4.43667822 | 73.6564895 | 3.40E-12 | 5.19E-11 | F | F | F | T |
| Psrc1 | -3.1328369 | 4.68867637 | 73.6415342 | 3.41E-12 | 5.21E-11 | F | F | T | F |
| Frmd4b | 2.38484106 | 5.40883909 | 73.5427445 | 3.49E-12 | 5.32E-11 | F | F | T | F |
| Cpe | -3.8940898 | 4.14043585 | 73.5391896 | 3.50E-12 | 5.33E-11 | F | F | T | F |
| Esrp2 | 2.69404768 | 4.38234642 | 73.3805125 | 3.63E-12 | 5.52E-11 | T | F | F | T |

| | | | | | | | | | |
|---|---|---|---|---|---|---|---|---|---|
| Abhd6 | 2.85050467 | 4.14604868 | 73.3037755 | 3.70E-12 | 5.62E-11 | T | F | F | T |
| Fam83g | 2.16667784 | 5.30848003 | 73.2781301 | 3.72E-12 | 5.65E-11 | T | F | F | F |
| Dpysl2 | -1.8142322 | 7.57255838 | 73.0946221 | 3.88E-12 | 5.89E-11 | T | F | T | T |
| Fscn1 | -1.6008757 | 9.28965101 | 73.0854379 | 3.89E-12 | 5.90E-11 | F | F | T | F |
| Hes1 | -3.4617574 | 4.44300058 | 73.0633961 | 3.91E-12 | 5.93E-11 | T | F | F | F |
| Hapln1 | -6.5730566 | 3.75551129 | 72.9470549 | 4.02E-12 | 6.09E-11 | T | T | T | T |
| Trim56 | -3.873541 | 4.12769701 | 72.8809859 | 4.08E-12 | 6.18E-11 | T | F | F | F |
| Pagr1a | -10.629814 | 3.05050142 | 72.8454217 | 4.12E-12 | 6.23E-11 | T | F | F | F |
| Dzip1l | -2.895271 | 4.91745108 | 72.7621008 | 4.20E-12 | 6.35E-11 | F | F | T | F |
| Kif21a | -2.3968043 | 6.01298678 | 72.5895922 | 4.37E-12 | 6.60E-11 | F | F | F | T |
| Rcor1 | 1.64586858 | 7.09171255 | 72.4931367 | 4.47E-12 | 6.75E-11 | T | F | T | T |
| Efemp2 | -3.4699031 | 4.33731603 | 72.3988931 | 4.57E-12 | 6.90E-11 | F | F | F | T |
| Rhbdf1 | 2.59337039 | 4.47489302 | 72.2796979 | 4.70E-12 | 7.09E-11 | F | F | F | F |
| Slc30a4 | 2.13207819 | 5.31227803 | 72.2733343 | 4.71E-12 | 7.10E-11 | T | F | F | T |
| Klf3 | -3.1262685 | 4.64156836 | 72.2178687 | 4.77E-12 | 7.18E-11 | T | F | T | T |
| Plcd3 | 3.08820798 | 3.8951514 | 72.1786023 | 4.82E-12 | 7.25E-11 | F | F | F | F |
| Arpc5 | 1.65795798 | 7.04703788 | 72.1665314 | 4.83E-12 | 7.26E-11 | F | F | F | F |
| Arfgef1 | 1.54122762 | 7.87030822 | 72.1295124 | 4.87E-12 | 7.32E-11 | T | T | T | T |
| Map7d3 | -5.2706313 | 3.57389935 | 72.0537763 | 4.96E-12 | 7.45E-11 | F | F | T | F |
| Nlgn2 | -2.2741622 | 6.21873094 | 72.0388111 | 4.98E-12 | 7.47E-11 | T | F | T | F |
| Ssbp4 | 2.18535818 | 5.18146843 | 71.9678762 | 5.06E-12 | 7.59E-11 | T | F | F | F |
| Hbb-bs | 5.01475719 | 2.55540027 | 71.9476406 | 5.09E-12 | 7.62E-11 | F | F | F | F |
| Fam131b | -4.9461717 | 3.6619208 | 71.9109429 | 5.13E-12 | 7.68E-11 | F | F | T | F |
| 1700019D0 | -4.6452446 | 3.81856042 | 71.8490419 | 5.21E-12 | 7.79E-11 | F | F | T | T |
| Plcxd2 | 2.66595875 | 4.2987291 | 71.8437963 | 5.21E-12 | 7.79E-11 | F | F | T | T |
| Prl8a9 | 5.0330466 | 2.5514221 | 71.7252187 | 5.36E-12 | 8.01E-11 | T | F | F | F |
| Nectin4 | 3.9679536 | 3.21131199 | 71.6949175 | 5.40E-12 | 8.06E-11 | F | F | T | F |
| Ube2d1 | -2.9048365 | 4.86809449 | 71.6560271 | 5.45E-12 | 8.13E-11 | T | T | F | T |
| Eml2 | 2.24879341 | 5.04979336 | 71.4900504 | 5.67E-12 | 8.45E-11 | F | F | F | F |
| Asxl1 | -1.7326295 | 7.93923249 | 71.4741918 | 5.69E-12 | 8.48E-11 | T | F | F | F |
| Eogt | 2.32365429 | 4.89694159 | 71.4416674 | 5.73E-12 | 8.54E-11 | T | F | T | F |
| Lfng | -5.1915585 | 3.52584806 | 71.3954278 | 5.80E-12 | 8.63E-11 | F | F | T | F |
| Vsir | 3.20261964 | 3.76633601 | 71.3647242 | 5.84E-12 | 8.68E-11 | F | F | T | F |
| Sept8 | 1.58917273 | 7.5095634 | 71.3551626 | 5.85E-12 | 8.70E-11 | T | T | T | T |
| Gale | 2.30504253 | 4.94884772 | 71.3458183 | 5.87E-12 | 8.71E-11 | F | F | F | F |
| Arpc4 | 1.56522338 | 7.70675958 | 71.2739189 | 5.97E-12 | 8.86E-11 | F | F | T | T |
| Asap2 | -2.4705202 | 5.55287406 | 71.1744804 | 6.11E-12 | 9.06E-11 | F | F | F | F |
| Setbp1 | -2.5598191 | 5.34967277 | 71.1202982 | 6.19E-12 | 9.17E-11 | T | F | T | T |
| Ttc7b | 1.86035534 | 6.23626432 | 71.1187078 | 6.19E-12 | 9.17E-11 | F | F | T | F |
| Tgfbr3 | 1.7316066 | 6.68978229 | 71.0045787 | 6.36E-12 | 9.42E-11 | T | F | T | T |
| Slc7a3 | -3.3223975 | 4.78738622 | 70.9408013 | 6.46E-12 | 9.55E-11 | F | F | F | F |
| Rbak | -4.8955556 | 3.69067915 | 70.8873842 | 6.54E-12 | 9.67E-11 | F | F | F | F |

| | | | | | | | | | |
|---|---|---|---|---|---|---|---|---|---|
| Baiap3 | 5.7525781 | 2.49813561 | 70.8242289 | 6.64E-12 | 9.81E-11 | F | F | T | F |
| Rhox1 | 8.06900274 | 1.68679301 | 70.819202 | 6.65E-12 | 9.82E-11 | T | F | T | T |
| Cdc42ep5 | -4.8922821 | 3.61040025 | 70.7611584 | 6.74E-12 | 9.95E-11 | F | F | F | F |
| Slco5a1 | 2.2781785 | 4.96808386 | 70.6182689 | 6.98E-12 | 1.03E-10 | T | F | T | T |
| Lpar5 | 6.54573885 | 1.98015465 | 70.6167103 | 6.98E-12 | 1.03E-10 | F | F | F | T |
| Muc13 | -5.1498704 | 3.51963637 | 70.5682404 | 7.06E-12 | 1.04E-10 | F | F | T | F |
| Csf2ra | 4.12600939 | 3.02612886 | 70.4875898 | 7.20E-12 | 1.06E-10 | F | F | F | F |
| Tmod3 | 1.61792603 | 7.11502224 | 70.4569237 | 7.25E-12 | 1.07E-10 | T | F | T | T |
| Trp53 | -1.7861569 | 7.52284153 | 70.4341137 | 7.29E-12 | 1.07E-10 | F | F | F | F |
| Eloa | 1.65620075 | 6.96656058 | 70.3767389 | 7.39E-12 | 1.09E-10 | T | T | T | T |
| Fgg | -7.2635898 | 3.09750335 | 70.3732187 | 7.40E-12 | 1.09E-10 | T | F | F | T |
| Cep89 | -3.2730749 | 4.43965729 | 70.2736553 | 7.58E-12 | 1.11E-10 | F | F | T | F |
| Cdip1 | -2.566135 | 5.28094722 | 70.2552786 | 7.61E-12 | 1.11E-10 | T | F | T | T |
| Ihh | -6.0326666 | 3.28193895 | 70.1551733 | 7.80E-12 | 1.14E-10 | F | F | F | T |
| Ppp2r5a | -2.2021389 | 6.11317893 | 70.1224845 | 7.86E-12 | 1.15E-10 | F | F | T | F |
| Lrpap1 | -2.0399887 | 6.65095654 | 70.110026 | 7.88E-12 | 1.15E-10 | T | F | T | T |
| Cyp2s1 | -10.450573 | 2.87766563 | 70.0725021 | 7.95E-12 | 1.16E-10 | T | F | F | T |
| Rgl2 | -2.7437998 | 5.03751036 | 70.0212694 | 8.05E-12 | 1.18E-10 | F | F | F | F |
| Phf8 | -2.5039883 | 5.39534805 | 69.9954712 | 8.10E-12 | 1.18E-10 | T | T | F | T |
| Tal1 | -5.7675783 | 3.4970968 | 69.9758498 | 8.14E-12 | 1.19E-10 | T | T | T | T |
| Bmp2k | 2.07076835 | 5.38458045 | 69.8778644 | 8.33E-12 | 1.21E-10 | T | F | F | F |
| Ccdc69 | 7.34955686 | 1.79950129 | 69.8436574 | 8.40E-12 | 1.22E-10 | F | F | F | F |
| Stmn2 | -5.1775882 | 3.48444458 | 69.8228695 | 8.44E-12 | 1.23E-10 | T | F | T | T |
| Nfe2l1 | 1.47353487 | 8.30619368 | 69.61708 | 8.87E-12 | 1.29E-10 | T | F | T | F |
| Bambi | -3.4741118 | 4.33745388 | 69.6003112 | 8.91E-12 | 1.29E-10 | T | T | T | T |
| Prr5l | -4.7449139 | 3.63955113 | 69.5668483 | 8.98E-12 | 1.30E-10 | F | F | T | F |
| Sowahb | -3.3486949 | 4.34828865 | 69.52411 | 9.07E-12 | 1.32E-10 | F | F | F | F |
| Pik3c2b | -2.4015228 | 5.73182586 | 69.3479733 | 9.47E-12 | 1.37E-10 | T | F | T | F |
| Brpf3 | 1.58587456 | 7.25352532 | 69.2278242 | 9.75E-12 | 1.41E-10 | T | F | T | F |
| Nceh1 | 2.39122618 | 4.68686244 | 69.1820408 | 9.85E-12 | 1.43E-10 | T | F | F | T |
| Spata21 | 7.32369447 | 1.83651947 | 69.1192448 | 1.00E-11 | 1.45E-10 | F | F | F | F |
| Noto | -10.419638 | 2.84952357 | 69.0653042 | 1.01E-11 | 1.47E-10 | F | F | T | F |
| Uchl1 | -5.3720047 | 3.41865287 | 68.9584726 | 1.04E-11 | 1.50E-10 | F | F | F | F |
| Lsr | -1.8727595 | 6.99393174 | 68.8959068 | 1.06E-11 | 1.53E-10 | T | F | F | T |
| Svep1 | -4.5945362 | 3.89377641 | 68.8473053 | 1.07E-11 | 1.54E-10 | T | F | T | T |
| Plg | -7.2428942 | 3.07772168 | 68.8190137 | 1.08E-11 | 1.55E-10 | T | F | F | F |
| Aldh5a1 | -3.3185859 | 4.34475579 | 68.7965486 | 1.08E-11 | 1.56E-10 | F | F | T | T |
| Selenon | -2.4251966 | 5.53527795 | 68.7234101 | 1.10E-11 | 1.59E-10 | T | T | T | T |
| Mllt3 | -3.5056243 | 4.22255009 | 68.6403651 | 1.12E-11 | 1.62E-10 | T | F | T | T |
| Maf | -3.3276871 | 4.35099934 | 68.6198097 | 1.13E-11 | 1.62E-10 | T | F | T | T |
| Fermt2 | 1.55742305 | 7.53971669 | 68.5835358 | 1.14E-11 | 1.64E-10 | T | F | F | F |
| Epb41l4a | -3.7702117 | 4.0783693 | 68.4835638 | 1.17E-11 | 1.68E-10 | F | F | T | F |

| | | | | | | | | | |
|---|---|---|---|---|---|---|---|---|---|
| Smad1 | -2.4772259 | 5.37340415 | 68.4449996 | 1.18E-11 | 1.69E-10 | T | F | F | T |
| Plekha7 | 1.67530443 | 6.78488632 | 68.2939662 | 1.22E-11 | 1.75E-10 | T | F | T | F |
| Espn | -4.3542351 | 3.87927466 | 68.2687384 | 1.23E-11 | 1.76E-10 | F | F | F | F |
| Mboat2 | -3.1352599 | 4.55526909 | 68.2230094 | 1.24E-11 | 1.78E-10 | F | F | T | T |
| Gaa | 1.82604632 | 6.50939252 | 68.1965918 | 1.25E-11 | 1.79E-10 | F | F | F | F |
| Gdf11 | -5.1457538 | 3.4537162 | 68.172754 | 1.26E-11 | 1.80E-10 | F | F | T | T |
| Camta1 | -3.5211726 | 4.25469302 | 68.122174 | 1.27E-11 | 1.82E-10 | T | F | T | T |
| Gng3 | -6.4103193 | 3.17362733 | 68.0838052 | 1.29E-11 | 1.84E-10 | T | F | F | T |
| Rims3 | -5.0516135 | 3.57649691 | 68.0746613 | 1.29E-11 | 1.84E-10 | T | F | T | T |
| Arhgef17 | -1.8217113 | 7.1033931 | 68.0735484 | 1.29E-11 | 1.84E-10 | T | F | T | T |
| Epha7 | -4.7883972 | 3.58299253 | 68.0148577 | 1.31E-11 | 1.87E-10 | F | F | T | F |
| Chst11 | -4.517398 | 3.69549227 | 68.0143695 | 1.31E-11 | 1.87E-10 | T | F | T | T |
| Zxdb | 2.63670805 | 4.25124185 | 67.9763798 | 1.32E-11 | 1.88E-10 | T | F | T | T |
| Lzts1 | 3.034685 | 3.8575492 | 67.8242494 | 1.37E-11 | 1.95E-10 | F | F | T | T |
| Dnajb5 | -7.1889915 | 3.0233053 | 67.7476679 | 1.40E-11 | 1.99E-10 | F | F | T | T |
| H4c8 | -1.5737272 | 8.81211833 | 67.7311116 | 1.40E-11 | 1.99E-10 | F | F | F | F |
| Peli3 | -3.8356445 | 4.0213247 | 67.6928218 | 1.42E-11 | 2.01E-10 | F | F | T | F |
| Nadk2 | -2.2251825 | 5.93414441 | 67.6758994 | 1.42E-11 | 2.02E-10 | T | F | T | F |
| Bmp4 | -3.8682671 | 4.02946038 | 67.565795 | 1.46E-11 | 2.07E-10 | F | F | F | F |
| Dennd1a | 2.04233802 | 5.37927836 | 67.5629307 | 1.46E-11 | 2.07E-10 | F | F | F | F |
| Gata5 | -10.742594 | 3.16960567 | 67.5544223 | 1.46E-11 | 2.07E-10 | F | F | T | F |
| Atmin | 1.73481671 | 6.52284101 | 67.4846881 | 1.49E-11 | 2.11E-10 | F | F | T | F |
| Kcnq4 | 4.1915538 | 2.96712775 | 67.4342532 | 1.51E-11 | 2.13E-10 | F | F | T | F |
| Rpl10l | 6.26415567 | 1.94710374 | 67.4328012 | 1.51E-11 | 2.13E-10 | F | F | F | F |
| Asic1 | -10.381393 | 2.81073308 | 67.423686 | 1.51E-11 | 2.14E-10 | F | F | T | F |
| Gas2l1 | 1.89198579 | 5.87496442 | 67.4105638 | 1.52E-11 | 2.14E-10 | F | F | T | T |
| Megf6 | -2.7196581 | 4.96443358 | 67.3255963 | 1.55E-11 | 2.19E-10 | F | F | T | T |
| Esyt2 | 1.68799195 | 6.74477248 | 67.2265792 | 1.59E-11 | 2.24E-10 | T | F | F | T |
| Mapk12 | -10.681967 | 3.10006674 | 67.2082406 | 1.59E-11 | 2.25E-10 | F | F | F | F |
| Vstm5 | 4.46780572 | 2.88958737 | 67.1759979 | 1.61E-11 | 2.26E-10 | F | F | T | F |
| Myl12a | 1.57412107 | 7.22317415 | 67.0820798 | 1.64E-11 | 2.31E-10 | F | F | F | T |
| Braf | 1.4873189 | 8.02952631 | 67.0137334 | 1.67E-11 | 2.35E-10 | T | F | T | T |
| Baz1a | 1.77930559 | 6.48344789 | 66.9969002 | 1.68E-11 | 2.36E-10 | T | F | T | F |
| Ndrg2 | -10.469854 | 2.89370158 | 66.9349806 | 1.70E-11 | 2.40E-10 | T | F | T | F |
| Glcci1 | -2.9846718 | 4.73919153 | 66.8156629 | 1.76E-11 | 2.46E-10 | T | T | T | T |
| Apba2 | -5.8891088 | 3.21770849 | 66.7470635 | 1.79E-11 | 2.51E-10 | F | F | F | F |
| Lpar6 | -3.6660418 | 4.08684918 | 66.6097487 | 1.85E-11 | 2.59E-10 | F | F | F | T |
| Zfp956 | -10.361073 | 2.7908977 | 66.5766686 | 1.86E-11 | 2.61E-10 | F | F | F | F |
| Slc25a23 | -3.2401215 | 4.34980283 | 66.5208308 | 1.89E-11 | 2.64E-10 | T | F | F | F |
| Cobll1 | 1.71751298 | 6.69449513 | 66.4692931 | 1.91E-11 | 2.68E-10 | T | F | F | T |
| Aprt | 1.57965429 | 7.4596373 | 66.4660871 | 1.91E-11 | 2.68E-10 | F | F | F | F |
| Foxn2 | 1.88963312 | 5.87677639 | 66.3988544 | 1.95E-11 | 2.72E-10 | T | F | T | F |

| | | | | | | | | | |
|---|---|---|---|---|---|---|---|---|---|
| Hba-a2 | 3.81495277 | 3.18425826 | 66.3986468 | 1.95E-11 | 2.72E-10 | F | F | F | F |
| Ccdc40 | -3.7854201 | 4.03631325 | 66.3915931 | 1.95E-11 | 2.72E-10 | F | F | T | F |
| Nipsnap1 | -2.3787858 | 5.48433949 | 66.1944746 | 2.05E-11 | 2.85E-10 | F | F | T | F |
| Arhgef25 | -2.8838102 | 4.71629149 | 66.1939782 | 2.05E-11 | 2.85E-10 | F | F | F | F |
| Klhl23 | -3.949096 | 3.92298979 | 66.1755381 | 2.06E-11 | 2.86E-10 | T | F | T | T |
| Rps6ka4 | 2.34135147 | 4.82265963 | 66.1739663 | 2.06E-11 | 2.86E-10 | F | F | T | F |
| Phf13 | 1.63942815 | 6.85188452 | 66.0453559 | 2.12E-11 | 2.95E-10 | F | F | T | F |
| Epha4 | -4.3778444 | 3.71580476 | 65.9692265 | 2.16E-11 | 3.01E-10 | T | F | T | T |
| Gnao1 | -3.1806638 | 4.38685832 | 65.9309814 | 2.18E-11 | 3.04E-10 | T | F | T | F |
| Itga8 | -10.342621 | 2.77451654 | 65.8326142 | 2.24E-11 | 3.11E-10 | T | F | T | T |
| Pex14 | 1.73205345 | 6.5624064 | 65.7860033 | 2.26E-11 | 3.14E-10 | F | F | T | F |
| Cacna1g | -4.983435 | 3.51268627 | 65.7539352 | 2.28E-11 | 3.17E-10 | F | F | F | F |
| Cdca7l | -2.2087314 | 5.87329208 | 65.6970956 | 2.31E-11 | 3.21E-10 | T | F | F | T |
| Dnah5 | -10.380341 | 2.80741824 | 65.6485103 | 2.34E-11 | 3.25E-10 | T | F | T | T |
| Piezo1 | 1.47601942 | 7.84599166 | 65.6406022 | 2.35E-11 | 3.25E-10 | F | F | F | T |
| Dusp1 | 2.74259173 | 4.10032034 | 65.5949885 | 2.37E-11 | 3.29E-10 | F | F | F | T |
| Rab25 | -5.1070204 | 3.42711728 | 65.5283322 | 2.41E-11 | 3.34E-10 | T | F | T | T |
| Atm | -1.8457202 | 7.00544088 | 65.4691324 | 2.45E-11 | 3.39E-10 | T | F | F | T |
| Tmed1 | -2.7560214 | 4.84528209 | 65.4109868 | 2.49E-11 | 3.43E-10 | F | F | T | F |
| Plekhm2 | 1.96521441 | 5.50990894 | 65.401093 | 2.49E-11 | 3.44E-10 | F | F | T | F |
| Fga | -5.93165 | 3.26115306 | 65.3874714 | 2.50E-11 | 3.45E-10 | F | F | T | T |
| Gprin1 | -10.329179 | 2.76106203 | 65.3629056 | 2.52E-11 | 3.47E-10 | F | F | T | F |
| Gfra3 | -10.331463 | 2.76185613 | 65.3156684 | 2.55E-11 | 3.51E-10 | F | F | F | F |
| Tmem229a | -10.338061 | 2.7676312 | 65.2817531 | 2.57E-11 | 3.53E-10 | T | F | T | T |
| Ptch2 | -10.913993 | 3.33748749 | 65.1831098 | 2.63E-11 | 3.62E-10 | F | F | T | F |
| Fkbp4 | -1.4892028 | 9.32083273 | 65.1547378 | 2.65E-11 | 3.64E-10 | T | F | F | T |
| Itpr1 | -2.0341041 | 6.36158898 | 65.1283193 | 2.67E-11 | 3.66E-10 | T | F | F | F |
| Elovl1 | 1.82880562 | 6.07106915 | 65.0736913 | 2.70E-11 | 3.71E-10 | T | F | F | F |
| Atp1b2 | -4.8760668 | 3.59327761 | 64.9876949 | 2.76E-11 | 3.79E-10 | T | F | T | T |
| Rbms1 | 1.43222237 | 8.21250597 | 64.9558855 | 2.78E-11 | 3.82E-10 | T | F | T | T |
| Eya1 | -7.0316134 | 3.01170107 | 64.931257 | 2.80E-11 | 3.84E-10 | T | F | T | T |
| Zfp800 | 2.1738045 | 4.94976376 | 64.8730134 | 2.84E-11 | 3.89E-10 | T | F | T | T |
| Insl3 | 9.99315517 | 1.39015166 | 65.2775883 | 2.86E-11 | 3.91E-10 | T | T | T | T |
| Creb3l3 | -5.1204729 | 3.43028867 | 64.8212681 | 2.88E-11 | 3.94E-10 | F | F | F | F |
| E2f7 | 1.5425351 | 7.18103277 | 64.7783919 | 2.91E-11 | 3.98E-10 | T | F | T | T |
| Cpn1 | -3.770135 | 4.06011367 | 64.7511913 | 2.93E-11 | 4.00E-10 | T | F | T | F |
| Fam222a | -4.9292491 | 3.45973758 | 64.7228644 | 2.95E-11 | 4.03E-10 | F | F | F | F |
| Itpr3 | 1.55168565 | 7.63221063 | 64.6759594 | 2.99E-11 | 4.07E-10 | F | F | F | F |
| Enpp2 | -5.1640728 | 3.46970335 | 64.6117294 | 3.03E-11 | 4.14E-10 | T | F | F | F |
| Khdrbs3 | -2.4678292 | 5.22004271 | 64.5393858 | 3.09E-11 | 4.21E-10 | T | T | F | T |
| Snx22 | 5.3825137 | 2.32588734 | 64.5153906 | 3.11E-11 | 4.23E-10 | F | F | F | F |
| Oas1c | 4.77443895 | 2.54542632 | 64.5009471 | 3.12E-11 | 4.25E-10 | F | F | F | F |

| | | | | | | | | | |
|---|---|---|---|---|---|---|---|---|---|
| Alpk3 | -4.7740996 | 3.49608637 | 64.4927659 | 3.13E-11 | 4.25E-10 | F | F | F | T |
| Capns1 | 1.86873424 | 6.24856914 | 64.4216208 | 3.18E-11 | 4.33E-10 | T | F | T | T |
| Col4a5 | -5.1075744 | 3.63384636 | 64.3793548 | 3.22E-11 | 4.37E-10 | F | F | F | F |
| Hspa4l | 2.33930641 | 4.72939041 | 64.3225335 | 3.26E-11 | 4.43E-10 | T | F | T | T |
| Tpbpa | 7.15608272 | 1.64156572 | 64.2956257 | 3.29E-11 | 4.46E-10 | T | F | F | F |
| Elk3 | 2.53177807 | 4.29303998 | 64.1993826 | 3.37E-11 | 4.56E-10 | F | F | T | T |
| Gusb | 1.80082826 | 6.12875979 | 64.1661107 | 3.39E-11 | 4.60E-10 | T | F | F | T |
| Slc43a3 | 2.33660479 | 4.60579484 | 64.1522634 | 3.41E-11 | 4.61E-10 | F | F | F | F |
| Myo5c | -5.4544349 | 3.26067436 | 64.1276286 | 3.43E-11 | 4.64E-10 | T | F | F | F |
| Lnpep | 1.67084256 | 6.75389671 | 64.1120235 | 3.44E-11 | 4.65E-10 | T | F | T | T |
| Cd63 | 1.73665004 | 6.4781521 | 64.1064261 | 3.45E-11 | 4.66E-10 | F | F | F | F |
| Gpr20 | 5.66017655 | 2.03183744 | 64.0264825 | 3.52E-11 | 4.75E-10 | F | F | F | F |
| March9 | -3.75459 | 4.00682408 | 63.9883072 | 3.55E-11 | 4.79E-10 | F | F | F | F |
| Gria4 | -10.313602 | 2.74319051 | 63.9428607 | 3.59E-11 | 4.84E-10 | T | T | T | F |
| Ier3 | 4.09292062 | 3.00661469 | 63.8321969 | 3.69E-11 | 4.98E-10 | F | F | F | F |
| Slc35g1 | 1.77450741 | 6.15070014 | 63.8275641 | 3.70E-11 | 4.98E-10 | F | F | T | F |
| Kif21b | -2.1400495 | 5.94764003 | 63.750983 | 3.77E-11 | 5.07E-10 | T | F | T | T |
| Syt14 | -2.7809242 | 4.77615499 | 63.7142576 | 3.80E-11 | 5.12E-10 | T | F | T | T |
| Man2b2 | -2.5508497 | 5.07289679 | 63.7116284 | 3.81E-11 | 5.12E-10 | F | F | F | T |
| Cxxc4 | -10.323824 | 2.75839177 | 63.6772422 | 3.84E-11 | 5.16E-10 | T | F | T | F |
| Sh3gl3 | -10.290086 | 2.72405036 | 63.4938166 | 4.02E-11 | 5.40E-10 | T | F | F | F |
| Pdlim4 | -3.4491844 | 4.1862921 | 63.4802785 | 4.03E-11 | 5.42E-10 | F | F | F | F |
| Nedd9 | -4.3824513 | 3.68878597 | 63.4675038 | 4.05E-11 | 5.43E-10 | F | F | T | F |
| Ttll3 | 2.94956289 | 3.83705802 | 63.3830396 | 4.14E-11 | 5.55E-10 | F | F | T | F |
| Mixl1 | -10.292838 | 2.72718234 | 63.3616907 | 4.16E-11 | 5.57E-10 | F | F | F | F |
| Tmem54 | 2.8889137 | 3.88683452 | 63.3204079 | 4.20E-11 | 5.63E-10 | F | F | F | F |
| Igfbpl1 | -10.321334 | 2.75623455 | 63.302844 | 4.22E-11 | 5.65E-10 | F | F | T | F |
| Ptk2b | 2.16343138 | 5.00621151 | 63.214596 | 4.32E-11 | 5.77E-10 | T | F | F | T |
| Creb3l1 | -4.0991432 | 3.82264126 | 63.2142652 | 4.32E-11 | 5.77E-10 | F | F | T | F |
| Zcchc18 | -10.274278 | 2.70679168 | 63.1861961 | 4.35E-11 | 5.81E-10 | T | F | T | T |
| Snph | -5.4969772 | 3.23091323 | 62.9673911 | 4.59E-11 | 6.13E-10 | F | F | F | F |
| Il2rb | 5.79425429 | 1.96202151 | 62.9047108 | 4.67E-11 | 6.23E-10 | T | F | T | T |
| Mapk13 | 3.25342811 | 3.50316565 | 62.9004494 | 4.67E-11 | 6.23E-10 | F | F | F | T |
| Chst1 | -10.290401 | 2.72027776 | 62.8439871 | 4.74E-11 | 6.31E-10 | F | F | F | F |
| Apba1 | 5.61884756 | 2.11003968 | 62.8439439 | 4.74E-11 | 6.31E-10 | T | T | T | T |
| Ptprt | -5.7930677 | 3.14328163 | 62.8254489 | 4.76E-11 | 6.34E-10 | F | F | T | T |
| Pdgfa | -2.479914 | 5.14828919 | 62.7450436 | 4.86E-11 | 6.47E-10 | F | F | T | F |
| Pde3a | -4.0743411 | 3.79101571 | 62.7428146 | 4.86E-11 | 6.47E-10 | T | F | T | T |
| Fam160a1 | 3.96104728 | 2.91818082 | 62.6382301 | 5.00E-11 | 6.64E-10 | F | F | F | F |
| Ankrd13a | 1.47761492 | 7.55425464 | 62.6241217 | 5.01E-11 | 6.66E-10 | T | F | T | T |
| Mmp14 | -2.0717102 | 6.08610408 | 62.5894487 | 5.06E-11 | 6.71E-10 | F | F | T | T |
| Tril | -7.0867006 | 2.92517289 | 62.4949402 | 5.18E-11 | 6.87E-10 | T | F | T | T |

| | | | | | | | | | |
|---|---|---|---|---|---|---|---|---|---|
| Sft2d2 | -2.5226048 | 5.06455377 | 62.368242 | 5.35E-11 | 7.09E-10 | T | F | T | T |
| Cacna2d1 | -10.270432 | 2.70563971 | 62.3575379 | 5.37E-11 | 7.11E-10 | T | F | T | F |
| Mpnd | -2.407848 | 5.30757606 | 62.3284768 | 5.41E-11 | 7.16E-10 | F | F | F | F |
| Smarcd1 | -1.6187753 | 7.85573427 | 62.2491596 | 5.52E-11 | 7.30E-10 | F | F | F | F |
| Nol4l | -2.2500077 | 5.60838821 | 62.2053902 | 5.58E-11 | 7.38E-10 | T | F | F | F |
| Gcnt1 | 1.81292378 | 5.93946457 | 62.1378454 | 5.68E-11 | 7.50E-10 | T | F | F | T |
| Tcf15 | -10.325524 | 2.76133682 | 62.1039486 | 5.73E-11 | 7.56E-10 | T | F | F | F |
| Zzef1 | 1.67066643 | 6.48368306 | 62.090273 | 5.75E-11 | 7.58E-10 | T | F | T | F |
| Rnf165 | -4.7716151 | 3.61002025 | 62.0787 | 5.76E-11 | 7.60E-10 | F | F | T | T |
| Fhod3 | -4.6069419 | 3.4999023 | 62.0693611 | 5.78E-11 | 7.61E-10 | T | T | F | T |
| Scarb1 | 1.66730081 | 6.69603676 | 61.9652264 | 5.93E-11 | 7.82E-10 | T | F | T | T |
| Itprid2 | 1.76901408 | 6.11534275 | 61.9158874 | 6.01E-11 | 7.91E-10 | F | F | F | F |
| Hipk3 | 1.6454978 | 6.57318727 | 61.8353955 | 6.13E-11 | 8.07E-10 | F | F | T | T |
| Msmo1 | 2.29044305 | 4.70801121 | 61.6752171 | 6.39E-11 | 8.40E-10 | F | F | T | F |
| Ccn1 | -3.0235879 | 4.42196584 | 61.5782671 | 6.55E-11 | 8.61E-10 | F | F | F | F |
| St3gal1 | 2.00829008 | 5.57527885 | 61.5516828 | 6.60E-11 | 8.66E-10 | T | F | T | T |
| Sorbs1 | 2.24896937 | 4.67352502 | 61.5282279 | 6.64E-11 | 8.71E-10 | T | F | T | T |
| Smad4 | -1.7832697 | 7.05531823 | 61.5178079 | 6.65E-11 | 8.73E-10 | T | F | T | T |
| Nucb2 | 2.76763987 | 3.99601497 | 61.4903814 | 6.70E-11 | 8.79E-10 | T | F | F | F |
| Etl4 | 1.45804662 | 7.78393379 | 61.4847186 | 6.71E-11 | 8.79E-10 | T | T | F | T |
| Frmd4a | 1.63062085 | 6.63347284 | 61.4245839 | 6.82E-11 | 8.92E-10 | T | T | T | T |
| Mageb16 | 2.93185022 | 3.82561065 | 61.4192352 | 6.83E-11 | 8.93E-10 | F | F | T | F |
| Rab43 | 2.35732679 | 4.4724802 | 61.3276104 | 6.99E-11 | 9.14E-10 | F | F | T | F |
| Porcn | -6.2417213 | 3.01127119 | 61.3198547 | 7.00E-11 | 9.15E-10 | F | F | F | T |
| Nr2f6 | 1.60970221 | 6.72770545 | 61.3087033 | 7.02E-11 | 9.17E-10 | T | F | T | F |
| Sema4a | -10.233713 | 2.66972407 | 61.1174666 | 7.38E-11 | 9.63E-10 | F | F | T | F |
| Tmsb4x | 1.88593171 | 6.32452416 | 61.0503108 | 7.51E-11 | 9.79E-10 | F | F | T | F |
| Git2 | 1.76810217 | 6.0511999 | 61.0269724 | 7.55E-11 | 9.85E-10 | T | F | T | F |
| Rap1gap2 | -10.423255 | 2.8582681 | 60.9136939 | 7.78E-11 | 1.01E-09 | T | F | T | T |
| Pcgf2 | -2.0949713 | 5.92253751 | 60.8610275 | 7.88E-11 | 1.03E-09 | F | F | T | F |
| Zfhx2 | -2.3859349 | 5.36146708 | 60.8576724 | 7.89E-11 | 1.03E-09 | F | F | F | T |
| Cdc42bpb | 1.37025017 | 8.35757255 | 60.841865 | 7.92E-11 | 1.03E-09 | F | F | T | F |
| Arhgap24 | -10.212165 | 2.64663452 | 60.8235943 | 7.96E-11 | 1.03E-09 | T | F | T | T |
| Ddc | -10.220493 | 2.65671722 | 60.7078857 | 8.20E-11 | 1.07E-09 | T | F | F | T |
| Ephb6 | -6.2292397 | 2.99821336 | 60.6817832 | 8.26E-11 | 1.07E-09 | F | F | F | F |
| Slc35e1 | 1.46698899 | 7.41581705 | 60.6698656 | 8.28E-11 | 1.07E-09 | T | F | T | T |
| D630045J1: | 1.64614962 | 6.52807355 | 60.6624879 | 8.30E-11 | 1.08E-09 | T | F | T | F |
| Neurl1b | 3.0545297 | 3.7050793 | 60.6470726 | 8.33E-11 | 1.08E-09 | T | F | T | T |
| Skida1 | -3.2903489 | 4.20398664 | 60.6124064 | 8.41E-11 | 1.09E-09 | T | F | F | T |
| Aqp11 | -4.3016868 | 3.61077108 | 60.6040528 | 8.42E-11 | 1.09E-09 | F | F | T | T |
| Ptprs | 1.38773235 | 8.09548295 | 60.5273014 | 8.59E-11 | 1.11E-09 | T | F | T | F |
| Nlrp4c | 4.72081832 | 2.3825602 | 60.496427 | 8.66E-11 | 1.12E-09 | F | F | T | F |

| | | | | | | | | | |
|---|---|---|---|---|---|---|---|---|---|
| Prl2c5 | 7.83414656 | 1.4927946 | 60.4868169 | 8.68E-11 | 1.12E-09 | F | F | F | F |
| Pakap | -1.9437776 | 6.65873326 | 60.4398133 | 8.79E-11 | 1.14E-09 | T | F | T | T |
| Col4a6 | -4.7471354 | 3.47117281 | 60.4296909 | 8.81E-11 | 1.14E-09 | F | F | F | F |
| Slc20a1 | 1.50022151 | 7.10793914 | 60.4050408 | 8.87E-11 | 1.14E-09 | T | T | F | T |
| Pura | 2.01573021 | 5.24918408 | 60.3828086 | 8.92E-11 | 1.15E-09 | F | F | T | T |
| Nuak1 | -2.8036618 | 4.61628418 | 60.3765859 | 8.94E-11 | 1.15E-09 | T | F | F | F |
| Pdlim1 | 1.8914448 | 5.58994934 | 60.3645317 | 8.96E-11 | 1.15E-09 | T | F | F | T |
| Lrrc4b | -10.206574 | 2.64288221 | 60.3248649 | 9.06E-11 | 1.17E-09 | F | F | F | F |
| Pfn1 | 1.25659381 | 9.71005699 | 60.2118383 | 9.33E-11 | 1.20E-09 | T | F | F | F |
| Gm8797 | -1.928952 | 6.37158264 | 60.1417789 | 9.50E-11 | 1.22E-09 | F | F | F | F |
| Mbd2 | 1.99168276 | 5.17517897 | 60.0144646 | 9.82E-11 | 1.26E-09 | T | F | T | T |
| Myorg | -2.8695828 | 4.52390776 | 59.9766981 | 9.91E-11 | 1.27E-09 | F | F | T | T |
| Zbtb7b | -2.9530183 | 4.54373856 | 59.9521245 | 9.98E-11 | 1.28E-09 | T | F | T | F |
| Cebpb | 2.43159809 | 4.29630278 | 59.9401341 | 1.00E-10 | 1.28E-09 | F | F | T | T |
| Trim46 | -10.195444 | 2.62883484 | 59.8907645 | 1.01E-10 | 1.30E-09 | F | F | T | F |
| Tfap4 | -2.441711 | 5.10021614 | 59.8833009 | 1.02E-10 | 1.30E-09 | F | F | F | F |
| Fam78b | 3.03226386 | 3.83995321 | 59.8582054 | 1.02E-10 | 1.31E-09 | T | F | T | T |
| Alx1 | -10.206764 | 2.6386413 | 59.6779532 | 1.07E-10 | 1.37E-09 | T | F | T | T |
| Jph3 | -10.189359 | 2.62289734 | 59.6619181 | 1.08E-10 | 1.38E-09 | T | F | T | T |
| Nars | 1.45130576 | 7.46386018 | 59.6363101 | 1.08E-10 | 1.38E-09 | F | F | F | F |
| Mylip | -3.4728525 | 4.04875999 | 59.6360651 | 1.08E-10 | 1.38E-09 | F | F | F | F |
| Amt | -2.1457815 | 5.6923346 | 59.5844998 | 1.10E-10 | 1.40E-09 | F | F | F | F |
| Nell2 | -5.4215678 | 3.15774179 | 59.4510534 | 1.14E-10 | 1.45E-09 | F | F | F | T |
| Yap1 | -1.578048 | 7.83501227 | 59.399388 | 1.15E-10 | 1.47E-09 | T | F | T | F |
| Fbxl7 | -6.9083937 | 2.85899394 | 59.3471299 | 1.17E-10 | 1.49E-09 | T | F | T | F |
| Tspoap1 | -10.240429 | 2.66975977 | 59.3347928 | 1.17E-10 | 1.49E-09 | T | F | T | F |
| Cnpy2 | -2.0904312 | 5.8904325 | 59.3336734 | 1.17E-10 | 1.49E-09 | F | F | F | T |
| Mdfi | 2.15685723 | 4.77390114 | 59.292578 | 1.19E-10 | 1.51E-09 | F | F | T | F |
| Entpd7 | 1.87260165 | 5.53054994 | 59.2699091 | 1.19E-10 | 1.52E-09 | T | F | T | T |
| Cth | -4.1946419 | 3.62003833 | 59.2515366 | 1.20E-10 | 1.52E-09 | T | F | F | T |
| Mgat2 | 1.60235363 | 6.62822962 | 59.209699 | 1.21E-10 | 1.54E-09 | F | F | F | T |
| Ciz1 | 1.73292217 | 6.04413592 | 59.1323749 | 1.24E-10 | 1.57E-09 | F | F | F | F |
| Hps1 | -3.1981201 | 4.19889924 | 59.1181402 | 1.24E-10 | 1.57E-09 | F | F | T | T |
| Lbh | -3.4986005 | 4.17176905 | 59.0854668 | 1.25E-10 | 1.59E-09 | F | F | T | T |
| Crb3 | -4.4397611 | 3.48676166 | 59.0434329 | 1.27E-10 | 1.60E-09 | F | F | F | F |
| Srgap2 | 1.47451117 | 7.16981641 | 59.0077764 | 1.28E-10 | 1.62E-09 | T | F | F | T |
| Mta1 | -1.5821565 | 7.77298479 | 58.9400415 | 1.30E-10 | 1.64E-09 | F | F | T | F |
| Wnt6 | -5.7993211 | 3.13300093 | 58.9267335 | 1.30E-10 | 1.65E-09 | T | F | T | F |
| Efnb3 | -2.9640992 | 4.47614919 | 58.8656955 | 1.33E-10 | 1.68E-09 | F | F | F | F |
| Rtl8a | -2.5319767 | 4.92711329 | 58.8087686 | 1.35E-10 | 1.70E-09 | T | F | T | F |
| Slc8b1 | 2.05082003 | 5.05577666 | 58.7784865 | 1.36E-10 | 1.71E-09 | T | F | T | T |
| Mcm2 | -1.4762268 | 8.61027699 | 58.728054 | 1.37E-10 | 1.73E-09 | T | F | F | T |

| | | | | | | | | | |
|---|---|---|---|---|---|---|---|---|---|
| Plcg1 | -1.8504535 | 6.55781291 | 58.7063798 | 1.38E-10 | 1.74E-09 | T | F | F | F |
| Chrdl1 | -10.159597 | 2.5970947 | 58.6708472 | 1.40E-10 | 1.76E-09 | T | F | T | T |
| Tspan18 | 2.03277489 | 5.28352339 | 58.6596952 | 1.40E-10 | 1.76E-09 | T | F | T | T |
| Col9a1 | -10.200002 | 2.63880566 | 58.5884364 | 1.43E-10 | 1.79E-09 | F | F | F | F |
| Ripor2 | 3.618622 | 3.14335593 | 58.5400245 | 1.44E-10 | 1.82E-09 | F | F | T | F |
| Nme4 | -3.2809514 | 4.13839562 | 58.5303008 | 1.45E-10 | 1.82E-09 | F | F | F | F |
| Arhgap20 | 3.56465409 | 3.54382486 | 58.43021 | 1.49E-10 | 1.87E-09 | T | F | T | F |
| Brpf1 | 1.46790742 | 7.16854206 | 58.4215722 | 1.49E-10 | 1.87E-09 | F | F | T | F |
| Pla2g12b | -4.6518045 | 3.37825743 | 58.3954346 | 1.50E-10 | 1.88E-09 | F | F | F | F |
| Msantd4 | -2.312332 | 5.32143419 | 58.3909304 | 1.50E-10 | 1.88E-09 | T | F | T | T |
| Lmo2 | -7.21021 | 3.04904005 | 58.3723598 | 1.51E-10 | 1.89E-09 | T | F | F | T |
| Fbln2 | -3.977573 | 3.70585101 | 58.3582778 | 1.52E-10 | 1.90E-09 | F | F | F | F |
| Tbx2 | -10.209103 | 2.64826068 | 58.2455485 | 1.56E-10 | 1.95E-09 | F | F | T | F |
| Gabbr2 | -6.8567651 | 3.01061773 | 58.0488649 | 1.64E-10 | 2.06E-09 | F | F | F | F |
| Ccdc27 | 9.83320278 | 1.23289882 | 58.390339 | 1.65E-10 | 2.06E-09 | F | F | F | F |
| Fam177a | 2.72609107 | 3.91784501 | 58.0327364 | 1.65E-10 | 2.06E-09 | F | F | T | T |
| Astn1 | -4.3497772 | 3.56238533 | 58.0195407 | 1.66E-10 | 2.07E-09 | T | F | T | T |
| Eno3 | -3.7201767 | 3.85708268 | 58.0053507 | 1.66E-10 | 2.08E-09 | F | F | F | F |
| Zfp532 | -1.8323912 | 6.56770322 | 57.8963653 | 1.71E-10 | 2.14E-09 | T | F | T | F |
| Tdrd5 | -6.999115 | 2.83546104 | 57.8600247 | 1.73E-10 | 2.16E-09 | F | F | F | T |
| Pcsk7 | 2.08137326 | 4.97909186 | 57.7884624 | 1.76E-10 | 2.20E-09 | T | F | T | T |
| Rhbg | -10.22209 | 2.66157574 | 57.7631326 | 1.77E-10 | 2.21E-09 | F | F | T | F |
| Pros1 | 2.05299698 | 4.93905558 | 57.6735526 | 1.82E-10 | 2.26E-09 | T | F | F | F |
| Cfc1 | -5.7112085 | 3.04329496 | 57.6429663 | 1.83E-10 | 2.28E-09 | F | F | F | F |
| Vwa5b1 | -6.9717123 | 2.8461957 | 57.6421783 | 1.83E-10 | 2.28E-09 | F | F | F | T |
| Dock11 | -1.8103621 | 6.89284299 | 57.6393071 | 1.83E-10 | 2.28E-09 | F | F | F | T |
| Nfil3 | -3.3612586 | 4.1030835 | 57.6079223 | 1.85E-10 | 2.30E-09 | F | F | T | F |
| Fzd6 | -4.0300721 | 3.66423707 | 57.5374355 | 1.88E-10 | 2.34E-09 | T | F | T | T |
| Cacna2d2 | -10.136153 | 2.57052899 | 57.5076782 | 1.90E-10 | 2.35E-09 | F | F | T | F |
| Ccdc177 | -10.137163 | 2.57632809 | 57.4871051 | 1.91E-10 | 2.37E-09 | F | F | T | F |
| Necap2 | 1.84249387 | 5.50986779 | 57.454024 | 1.93E-10 | 2.39E-09 | F | F | F | F |
| Rtkn | -4.5263737 | 3.42284309 | 57.4167581 | 1.94E-10 | 2.41E-09 | F | F | F | F |
| Iars | 1.30259227 | 8.70802508 | 57.3758385 | 1.97E-10 | 2.43E-09 | F | F | F | T |
| Adam19 | 1.4348566 | 7.39852193 | 57.3489933 | 1.98E-10 | 2.45E-09 | F | F | T | F |
| Map3k20 | -2.4861746 | 4.92052486 | 57.2377259 | 2.04E-10 | 2.52E-09 | T | F | T | F |
| Etv2 | -7.0330732 | 2.87549467 | 57.1717064 | 2.08E-10 | 2.56E-09 | F | F | F | F |
| Gm28048 | 9.81054441 | 1.21067511 | 57.4585554 | 2.10E-10 | 2.60E-09 | F | F | F | F |
| Med12l | 2.14185025 | 4.71775191 | 57.0644214 | 2.14E-10 | 2.64E-09 | T | F | T | F |
| F3 | 3.74965235 | 2.91346541 | 56.992861 | 2.18E-10 | 2.69E-09 | F | F | T | F |
| Arhgap21 | 1.42118126 | 7.42325631 | 56.9645052 | 2.19E-10 | 2.70E-09 | F | F | T | F |
| Tcfl5 | -2.7370695 | 4.58496303 | 56.9565017 | 2.20E-10 | 2.71E-09 | T | F | T | F |
| Gipc2 | -5.7024481 | 3.07156476 | 56.8897876 | 2.24E-10 | 2.75E-09 | F | F | F | F |

| | | | | | | | | | |
|---|---|---|---|---|---|---|---|---|---|
| Arfgef2 | 1.4996798 | 6.91923336 | 56.8883267 | 2.24E-10 | 2.75E-09 | T | F | T | F |
| Fmnl3 | -2.6924997 | 4.64144664 | 56.7232444 | 2.34E-10 | 2.88E-09 | F | F | F | F |
| Msh6 | -1.541494 | 7.95016523 | 56.7229521 | 2.34E-10 | 2.88E-09 | T | F | F | F |
| Pank1 | -3.7538218 | 3.77957167 | 56.6311346 | 2.40E-10 | 2.95E-09 | T | F | T | T |
| Mtmr1 | 2.31704301 | 4.45020911 | 56.6138941 | 2.41E-10 | 2.96E-09 | T | F | F | F |
| Mns1 | -2.2391849 | 5.45375643 | 56.6053749 | 2.41E-10 | 2.96E-09 | F | F | T | F |
| Necab2 | -10.106004 | 2.54565351 | 56.6026849 | 2.42E-10 | 2.96E-09 | F | F | F | F |
| Sypl | 1.55699812 | 6.65285639 | 56.6010606 | 2.42E-10 | 2.96E-09 | T | T | T | T |
| Mcoln3 | -10.096244 | 2.53344016 | 56.5800697 | 2.43E-10 | 2.98E-09 | F | F | F | F |
| Glt1d1 | -6.075045 | 2.91225364 | 56.4912985 | 2.49E-10 | 3.05E-09 | F | F | T | T |
| Eras | -5.6673112 | 3.00212007 | 56.4317983 | 2.53E-10 | 3.10E-09 | F | F | F | F |
| Sgpl1 | -1.7105079 | 6.86380931 | 56.3965763 | 2.55E-10 | 3.12E-09 | T | F | T | F |
| 2510009E0′ | -4.5027711 | 3.40016673 | 56.3153729 | 2.61E-10 | 3.19E-09 | T | F | T | T |
| Bclaf3 | 2.2099778 | 4.69383821 | 56.3018114 | 2.62E-10 | 3.20E-09 | F | F | F | F |
| Osbp | 1.53622545 | 6.73046259 | 56.2360992 | 2.67E-10 | 3.25E-09 | T | F | T | F |
| Prpf40b | -3.370179 | 4.03505805 | 56.234688 | 2.67E-10 | 3.25E-09 | T | F | F | F |
| Slc25a24 | 1.74472212 | 5.80505098 | 56.1869266 | 2.70E-10 | 3.29E-09 | T | F | T | F |
| Pidd1 | -10.213697 | 2.65436357 | 56.1199744 | 2.75E-10 | 3.35E-09 | F | F | T | F |
| Alad | 1.86104176 | 5.60522433 | 56.1011507 | 2.76E-10 | 3.37E-09 | F | F | T | T |
| Anln | 1.3806689 | 7.70475179 | 56.071374 | 2.79E-10 | 3.39E-09 | T | T | T | T |
| Rrbp1 | 1.28161267 | 8.78382202 | 55.96831 | 2.87E-10 | 3.49E-09 | F | F | F | F |
| Nudt11 | -3.6524465 | 3.82465212 | 55.9388347 | 2.89E-10 | 3.51E-09 | T | F | T | F |
| Inpp5b | 1.78887737 | 5.67738418 | 55.9088171 | 2.91E-10 | 3.54E-09 | F | F | F | F |
| Tcirg1 | 2.4997099 | 4.62754959 | 55.8976798 | 2.92E-10 | 3.55E-09 | F | F | F | F |
| Ptbp2 | -2.021273 | 5.94268129 | 55.8818872 | 2.93E-10 | 3.56E-09 | F | F | T | F |
| Gabrb3 | -5.7735126 | 3.10160542 | 55.7983959 | 3.00E-10 | 3.64E-09 | T | T | T | T |
| Blvra | 3.04302567 | 3.57325018 | 55.6789612 | 3.10E-10 | 3.76E-09 | F | F | F | F |
| Scara3 | -6.9920849 | 2.83581714 | 55.6382832 | 3.13E-10 | 3.80E-09 | T | F | T | T |
| Mpp2 | -6.9051296 | 2.74792028 | 55.628681 | 3.14E-10 | 3.80E-09 | F | F | T | F |
| Six1 | -10.101821 | 2.54324608 | 55.6178557 | 3.15E-10 | 3.81E-09 | T | T | F | T |
| Trib1 | -2.460905 | 4.89376875 | 55.5999601 | 3.16E-10 | 3.83E-09 | T | F | F | F |
| Gp1bb | 9.7721928 | 1.17309118 | 55.9064315 | 3.18E-10 | 3.84E-09 | F | F | T | F |
| Dzip1 | -4.0243696 | 3.65835836 | 55.5778101 | 3.18E-10 | 3.85E-09 | F | F | T | T |
| Hcn4 | -5.335292 | 3.0761443 | 55.4460638 | 3.30E-10 | 3.99E-09 | F | F | T | F |
| Klhl3 | -10.117362 | 2.55001227 | 55.4420046 | 3.30E-10 | 3.99E-09 | F | F | T | F |
| Tmed2 | 1.33834558 | 8.07310486 | 55.4342076 | 3.31E-10 | 3.99E-09 | T | F | T | F |
| Sacs | -2.3348351 | 5.12147523 | 55.3835369 | 3.36E-10 | 4.05E-09 | T | F | F | T |
| Tbc1d10b | 1.47813126 | 6.92285161 | 55.3488457 | 3.39E-10 | 4.08E-09 | F | F | F | F |
| Ubxn2a | 2.03328699 | 4.89468104 | 55.3332192 | 3.40E-10 | 4.10E-09 | T | F | T | T |
| Pla2g2d | 4.58734853 | 2.32239333 | 55.2709893 | 3.46E-10 | 4.17E-09 | F | F | T | F |
| Kyat3 | 3.22193178 | 3.31103976 | 55.2688445 | 3.46E-10 | 4.17E-09 | F | F | F | F |
| Trabd2b | -4.2380452 | 3.50374255 | 55.2547747 | 3.47E-10 | 4.18E-09 | T | F | T | F |

| | | | | | | | | | |
|---|---|---|---|---|---|---|---|---|---|
| Notch4 | -10.094825 | 2.53675111 | 55.1973433 | 3.53E-10 | 4.24E-09 | F | F | F | F |
| Egln1 | 1.62999783 | 6.23846183 | 55.1708783 | 3.55E-10 | 4.27E-09 | T | F | T | T |
| Zfp428 | -2.3489815 | 5.05336569 | 55.164386 | 3.56E-10 | 4.28E-09 | F | F | F | F |
| Prickle2 | -4.5870947 | 3.31615764 | 55.0607141 | 3.66E-10 | 4.40E-09 | F | T | T | T |
| Runx1t1 | -5.2764376 | 3.06989938 | 55.0454047 | 3.68E-10 | 4.41E-09 | T | F | T | T |
| Adamts9 | 1.69370992 | 6.25870782 | 55.020626 | 3.70E-10 | 4.44E-09 | F | F | F | F |
| Apoa2 | -10.150486 | 2.59262006 | 55.0115563 | 3.71E-10 | 4.45E-09 | F | F | F | T |
| Lhx5 | -10.091006 | 2.53322744 | 54.9490233 | 3.78E-10 | 4.52E-09 | F | F | F | F |
| Metrn | -3.1640776 | 4.38475719 | 54.7096115 | 4.03E-10 | 4.82E-09 | F | F | F | F |
| Klf2 | 2.91427419 | 3.61368078 | 54.7011908 | 4.04E-10 | 4.83E-09 | F | F | T | F |
| Cc2d2a | -3.6872337 | 3.76738844 | 54.6685872 | 4.07E-10 | 4.87E-09 | T | F | F | F |
| Bcat2 | -2.736507 | 4.57332601 | 54.6666417 | 4.08E-10 | 4.87E-09 | T | F | F | T |
| Plcd1 | 2.20841786 | 4.5304901 | 54.6499647 | 4.10E-10 | 4.89E-09 | F | F | F | F |
| Pkd1 | 1.49565021 | 6.84294707 | 54.5949388 | 4.16E-10 | 4.96E-09 | F | F | F | F |
| Cacng7 | -2.7235938 | 4.56179208 | 54.5889745 | 4.16E-10 | 4.97E-09 | F | F | F | F |
| Spred2 | -1.7275403 | 6.70805773 | 54.4818945 | 4.29E-10 | 5.11E-09 | T | F | F | F |
| Vat1l | -4.7035357 | 3.24113269 | 54.4474134 | 4.33E-10 | 5.16E-09 | T | F | T | T |
| S100a13 | 4.18962017 | 2.57830374 | 54.429266 | 4.35E-10 | 5.18E-09 | F | F | F | F |
| Syne1 | 1.85102058 | 5.32258471 | 54.4200422 | 4.36E-10 | 5.19E-09 | T | T | T | T |
| Agap2 | -10.034068 | 2.47264205 | 54.3854255 | 4.40E-10 | 5.24E-09 | F | F | T | F |
| Fgfr4 | -4.3416127 | 3.39265957 | 54.3486863 | 4.45E-10 | 5.29E-09 | T | F | F | F |
| Rspo1 | -10.034816 | 2.47648833 | 54.2574363 | 4.56E-10 | 5.42E-09 | F | F | F | F |
| Gins2 | -2.2574011 | 5.18951011 | 54.2425814 | 4.58E-10 | 5.44E-09 | T | F | F | F |
| Abhd5 | 2.22544817 | 4.47453929 | 54.1341405 | 4.72E-10 | 5.60E-09 | F | F | T | F |
| Secisbp2l | 1.55517064 | 6.50292588 | 54.0890401 | 4.77E-10 | 5.66E-09 | T | T | F | T |
| Prpsap2 | -2.7037582 | 4.50677295 | 54.0094869 | 4.88E-10 | 5.78E-09 | T | F | T | F |
| Hadhb | 1.72009539 | 5.80990248 | 53.9379148 | 4.98E-10 | 5.90E-09 | T | F | F | F |
| Depdc7 | -10.019508 | 2.46096127 | 53.9313293 | 4.98E-10 | 5.90E-09 | T | T | T | T |
| Pofut1 | -2.0708044 | 5.57351942 | 53.8762607 | 5.06E-10 | 5.99E-09 | T | F | F | T |
| Arhgef12 | 1.36759581 | 7.56052367 | 53.858884 | 5.08E-10 | 6.01E-09 | T | F | T | T |
| Snx10 | 1.75722038 | 5.74125034 | 53.8046208 | 5.16E-10 | 6.10E-09 | F | F | F | T |
| Daam2 | -10.026052 | 2.46854095 | 53.8036265 | 5.16E-10 | 6.10E-09 | T | F | T | T |
| Whrn | -4.0093771 | 3.54774884 | 53.7909 | 5.18E-10 | 6.12E-09 | F | F | T | F |
| Slc3a2 | 1.31793478 | 8.00543237 | 53.7885634 | 5.18E-10 | 6.12E-09 | T | F | T | T |
| Itgb5 | -1.8585051 | 6.2258398 | 53.779634 | 5.20E-10 | 6.13E-09 | F | F | F | T |
| Tpst2 | 1.91083985 | 5.11713341 | 53.7267478 | 5.27E-10 | 6.22E-09 | F | F | F | F |
| Map7d1 | 1.32218216 | 7.9747372 | 53.7015409 | 5.31E-10 | 6.26E-09 | F | F | F | T |
| Hdac2 | -1.6340265 | 6.96680975 | 53.6520457 | 5.38E-10 | 6.34E-09 | T | F | F | F |
| Plekha2 | 1.69457809 | 5.90980756 | 53.6461912 | 5.39E-10 | 6.35E-09 | T | F | T | F |
| Rbpms | -1.7359532 | 6.64460151 | 53.6053197 | 5.45E-10 | 6.41E-09 | F | F | T | F |
| Shisa6 | -5.0865129 | 3.1185269 | 53.5605478 | 5.52E-10 | 6.49E-09 | T | F | T | T |
| Tnc | -10.065301 | 2.50912266 | 53.5386307 | 5.55E-10 | 6.53E-09 | T | F | F | F |

| | | | | | | | | | |
|---|---|---|---|---|---|---|---|---|---|
| Maged1 | -1.545661 | 7.785361 | 53.503235 | 5.61E-10 | 6.59E-09 | T | F | F | F |
| Epas1 | 1.75703322 | 5.59011552 | 53.5011601 | 5.61E-10 | 6.59E-09 | F | F | F | F |
| Cxxc5 | -1.8988167 | 6.06910265 | 53.4602472 | 5.67E-10 | 6.66E-09 | T | F | F | T |
| Sema5b | -3.7749507 | 3.6679081 | 53.4535269 | 5.68E-10 | 6.67E-09 | T | F | T | F |
| L1td1 | -2.2231845 | 5.87498461 | 53.4058074 | 5.76E-10 | 6.75E-09 | F | F | F | F |
| Fzd5 | -1.938028 | 5.92822242 | 53.3949288 | 5.78E-10 | 6.77E-09 | T | F | T | T |
| Ptov1 | -1.574669 | 7.21017029 | 53.3604264 | 5.83E-10 | 6.83E-09 | F | F | T | F |
| Fam3c | 1.61509701 | 6.19337855 | 53.3553604 | 5.84E-10 | 6.83E-09 | F | F | T | T |
| Ccn2 | -6.0499306 | 2.82411925 | 53.2786477 | 5.96E-10 | 6.98E-09 | T | F | T | F |
| Cntnap2 | -5.1522727 | 3.0577649 | 53.2238354 | 6.06E-10 | 7.08E-09 | T | F | F | T |
| Pfkm | 1.53847139 | 6.53827552 | 53.2123079 | 6.08E-10 | 7.10E-09 | F | F | F | T |
| Usp28 | -2.1218311 | 5.68526075 | 53.1707328 | 6.15E-10 | 7.18E-09 | F | F | F | F |
| Fkbp15 | 1.61004956 | 6.19293701 | 53.1006538 | 6.27E-10 | 7.31E-09 | T | F | F | T |
| Arhgef19 | -3.8567061 | 3.67510349 | 53.0596341 | 6.34E-10 | 7.39E-09 | F | F | T | T |
| Tiam1 | 1.39277143 | 7.22904053 | 53.0228467 | 6.40E-10 | 7.46E-09 | T | F | T | T |
| Fam13c | -9.9859002 | 2.42759391 | 53.0097864 | 6.42E-10 | 7.49E-09 | T | F | F | F |
| Ndn | -2.8460113 | 4.49262563 | 52.9956211 | 6.45E-10 | 7.51E-09 | F | F | F | F |
| Tmem9 | -2.2316494 | 5.16822266 | 52.9862227 | 6.47E-10 | 7.53E-09 | F | F | T | F |
| Actg1 | 1.00343363 | 11.9792536 | 52.9582503 | 6.52E-10 | 7.58E-09 | F | F | F | F |
| Sars | 1.35396131 | 7.66936435 | 52.9378492 | 6.55E-10 | 7.62E-09 | T | F | T | F |
| Fads1 | -1.9839249 | 5.84459355 | 52.9260921 | 6.58E-10 | 7.64E-09 | T | F | T | T |
| Snrpn | -1.6451128 | 7.03751259 | 52.9225855 | 6.58E-10 | 7.64E-09 | T | F | F | F |
| Gucy1a2 | -9.990401 | 2.43354057 | 52.8986105 | 6.63E-10 | 7.69E-09 | T | T | T | T |
| Cgnl1 | 1.33025579 | 8.1537803 | 52.8957484 | 6.63E-10 | 7.69E-09 | T | F | T | T |
| Tmem40 | 6.55278653 | 1.53115352 | 52.8815368 | 6.66E-10 | 7.72E-09 | F | F | F | F |
| Nhlrc3 | 1.87900206 | 5.20685542 | 52.8744759 | 6.67E-10 | 7.73E-09 | F | F | T | F |
| Mvb12b | -2.3410816 | 4.97270193 | 52.8722597 | 6.67E-10 | 7.73E-09 | T | F | T | F |
| Bex3 | -1.6246676 | 7.14558566 | 52.86369 | 6.69E-10 | 7.74E-09 | T | F | F | F |
| Hist1h2bp | -2.1464525 | 6.24674472 | 52.8499934 | 6.72E-10 | 7.77E-09 | F | F | F | F |
| Atrn | 1.73012766 | 5.65834713 | 52.7960377 | 6.82E-10 | 7.88E-09 | T | F | F | T |
| Myh3 | 3.53514259 | 3.00069602 | 52.7724901 | 6.86E-10 | 7.93E-09 | F | F | F | F |
| Ddr1 | 1.44074204 | 7.02125742 | 52.7681616 | 6.87E-10 | 7.94E-09 | T | F | F | F |
| Sox13 | -2.3492774 | 5.57754764 | 52.7464823 | 6.91E-10 | 7.98E-09 | T | F | F | T |
| Ccdc92b | -6.8750932 | 2.71411851 | 52.7458958 | 6.91E-10 | 7.98E-09 | T | F | T | T |
| Cpsf7 | -1.8221871 | 6.26335653 | 52.7364683 | 6.93E-10 | 7.99E-09 | T | F | F | T |
| Klhl7 | -2.726647 | 4.56911179 | 52.7073613 | 6.99E-10 | 8.05E-09 | F | F | T | T |
| Armcx6 | -9.9754829 | 2.4167985 | 52.7023659 | 7.00E-10 | 8.06E-09 | T | F | F | T |
| Fam189b | -3.2624812 | 3.98287624 | 52.7004346 | 7.00E-10 | 8.06E-09 | F | F | F | F |
| Pced1a | -9.9769016 | 2.41939619 | 52.6982659 | 7.00E-10 | 8.06E-09 | F | F | F | T |
| Ift81 | -3.4794832 | 3.83074698 | 52.668039 | 7.06E-10 | 8.12E-09 | F | F | F | F |
| Cyb5b | 1.30720857 | 8.00480891 | 52.6556822 | 7.09E-10 | 8.15E-09 | T | F | T | F |
| Igsf11 | -4.5834565 | 3.36187547 | 52.6464059 | 7.10E-10 | 8.16E-09 | T | F | T | F |

| | | | | | | | | | |
|---|---|---|---|---|---|---|---|---|---|
| Tub | -5.5992555 | 2.93422184 | 52.6416712 | 7.11E-10 | 8.17E-09 | T | F | T | F |
| Prl3d3 | 5.91611973 | 1.76313501 | 52.6323584 | 7.13E-10 | 8.19E-09 | F | F | F | F |
| Agl | 1.70759272 | 5.74623144 | 52.6245373 | 7.15E-10 | 8.20E-09 | T | F | T | T |
| B4galt2 | -2.515985 | 4.68418829 | 52.5302536 | 7.34E-10 | 8.41E-09 | F | F | T | F |
| Tm9sf3 | 1.31861993 | 7.85816373 | 52.5268397 | 7.34E-10 | 8.42E-09 | T | F | T | F |
| Tbc1d1 | -1.7651095 | 6.4517498 | 52.5228232 | 7.35E-10 | 8.42E-09 | T | F | T | T |
| Mcf2l | -4.3004747 | 3.35224043 | 52.4810156 | 7.44E-10 | 8.52E-09 | T | F | T | F |
| Phf21a | -2.1171009 | 5.37798194 | 52.4526761 | 7.50E-10 | 8.58E-09 | F | F | T | T |
| Zfp365 | -9.9843797 | 2.42281085 | 52.4481323 | 7.51E-10 | 8.59E-09 | F | F | T | F |
| Inppl1 | -1.5617473 | 7.1819013 | 52.4287608 | 7.55E-10 | 8.63E-09 | F | F | F | F |
| Prl8a2 | 2.86928744 | 3.80070577 | 52.4160344 | 7.57E-10 | 8.65E-09 | F | F | F | F |
| Ikbkb | -2.7169509 | 4.41441357 | 52.3888218 | 7.63E-10 | 8.71E-09 | T | F | T | T |
| Psg19 | 7.64826879 | 1.27235289 | 52.3797796 | 7.65E-10 | 8.73E-09 | T | F | F | F |
| Large2 | -4.4575317 | 3.27671957 | 52.355717 | 7.70E-10 | 8.79E-09 | F | F | F | F |
| Thyn1 | -2.3595087 | 4.91932751 | 52.3485048 | 7.72E-10 | 8.80E-09 | F | F | F | F |
| Tln2 | -1.504246 | 7.58551837 | 52.2877152 | 7.85E-10 | 8.94E-09 | T | F | T | T |
| Stum | -10.0879 | 2.51904949 | 52.2528984 | 7.93E-10 | 9.03E-09 | T | F | T | T |
| Pcx | 1.73521454 | 5.65433443 | 52.2458873 | 7.94E-10 | 9.04E-09 | F | F | F | F |
| Tbck | 1.61740613 | 6.09226911 | 52.1449063 | 8.17E-10 | 9.29E-09 | T | F | T | T |
| Cep131 | -2.2559952 | 5.07298891 | 52.0806199 | 8.32E-10 | 9.45E-09 | F | F | F | F |
| Chga | -6.0289918 | 2.8027026 | 52.0292978 | 8.44E-10 | 9.59E-09 | F | F | F | F |
| Rbms3 | -6.8142488 | 2.66073318 | 52.0171688 | 8.46E-10 | 9.61E-09 | T | T | T | T |
| Marveld2 | 2.01720084 | 4.83973601 | 51.9901381 | 8.53E-10 | 9.68E-09 | F | F | T | F |
| Dhfr | -1.8976066 | 5.98549893 | 51.9698632 | 8.58E-10 | 9.73E-09 | T | F | T | T |
| Ptpn12 | 1.56431065 | 6.31990498 | 51.959704 | 8.60E-10 | 9.75E-09 | T | F | T | F |
| Alpl | -2.4861842 | 5.24701491 | 51.9382314 | 8.65E-10 | 9.81E-09 | F | F | F | F |
| Snd1 | 1.23444218 | 8.75811735 | 51.8891746 | 8.77E-10 | 9.94E-09 | F | F | T | F |
| Zc3h7a | 1.907568 | 5.07365545 | 51.8332042 | 8.91E-10 | 1.01E-08 | F | F | F | F |
| Iffo2 | 1.93738837 | 4.97013399 | 51.7989433 | 9.00E-10 | 1.02E-08 | F | F | T | F |
| Ppp1r14a | -9.9448618 | 2.38750655 | 51.7976159 | 9.00E-10 | 1.02E-08 | F | F | F | F |
| Igf2 | 1.32333885 | 10.0235156 | 51.7884041 | 9.02E-10 | 1.02E-08 | T | F | T | F |
| Pcsk2 | -10.063616 | 2.49557528 | 51.7721812 | 9.06E-10 | 1.02E-08 | T | F | F | F |
| Sox21 | -5.6577381 | 2.99680849 | 51.6806087 | 9.30E-10 | 1.05E-08 | F | F | T | F |
| Nppb | 4.88204699 | 2.00920621 | 51.5722185 | 9.59E-10 | 1.08E-08 | F | F | F | F |
| H2az2 | -2.1089568 | 5.41880459 | 51.5431366 | 9.66E-10 | 1.09E-08 | F | F | F | F |
| Il22ra1 | -6.021015 | 2.79901332 | 51.5213184 | 9.72E-10 | 1.10E-08 | F | F | F | F |
| Gramd1b | -1.9630552 | 5.714088 | 51.4961074 | 9.79E-10 | 1.10E-08 | T | F | T | T |
| Pdhb | 1.48707454 | 6.66602596 | 51.420986 | 1.00E-09 | 1.13E-08 | T | F | F | F |
| Ldb2 | -6.8017124 | 2.64458305 | 51.4072665 | 1.00E-09 | 1.13E-08 | T | F | T | F |
| Sox1 | -10.16258 | 2.60673687 | 51.3838948 | 1.01E-09 | 1.14E-08 | F | F | T | T |
| Zfp711 | -9.9792342 | 2.42547673 | 51.3729418 | 1.01E-09 | 1.14E-08 | T | T | F | T |
| Nphp3 | -2.5396149 | 4.60887982 | 51.336421 | 1.02E-09 | 1.15E-08 | T | F | T | F |

| | | | | | | | | | |
|---|---|---|---|---|---|---|---|---|---|
| Dab2ip | 1.35173179 | 7.69108855 | 51.3338871 | 1.02E-09 | 1.15E-08 | T | F | T | F |
| Dlg4 | -2.0669833 | 5.73598868 | 51.3242709 | 1.03E-09 | 1.15E-08 | F | F | T | F |
| Atxn10 | 1.29803239 | 7.91819385 | 51.2874764 | 1.04E-09 | 1.17E-08 | F | F | T | T |
| Suco | 1.49118009 | 6.63216566 | 51.2719231 | 1.04E-09 | 1.17E-08 | F | F | F | T |
| Ms4a4d | -9.9903932 | 2.42603134 | 51.2321349 | 1.05E-09 | 1.18E-08 | T | T | F | T |
| F5 | -3.9898476 | 3.71602115 | 51.2182131 | 1.06E-09 | 1.19E-08 | F | F | T | F |
| Ttyh2 | 2.0949954 | 4.57752738 | 51.2081065 | 1.06E-09 | 1.19E-08 | T | F | T | F |
| Parp10 | 3.33449027 | 3.07978812 | 51.2036342 | 1.06E-09 | 1.19E-08 | F | F | F | F |
| Lmtk3 | -2.394998 | 4.84826053 | 51.1926532 | 1.07E-09 | 1.19E-08 | T | F | F | T |
| Cep97 | -2.0095564 | 5.62555345 | 51.1786994 | 1.07E-09 | 1.20E-08 | T | F | T | T |
| Zfp827 | -3.7226106 | 3.62630888 | 51.1705559 | 1.07E-09 | 1.20E-08 | T | F | T | T |
| Lrmp | -9.9472865 | 2.38622577 | 51.137088 | 1.08E-09 | 1.21E-08 | F | F | F | F |
| Gstm1 | -2.9447655 | 4.19812804 | 51.0447118 | 1.11E-09 | 1.24E-08 | F | F | T | F |
| Habp2 | -9.9601719 | 2.40678017 | 51.0158289 | 1.12E-09 | 1.25E-08 | F | F | T | F |
| Ak1 | -5.0168566 | 3.07758325 | 50.9770952 | 1.13E-09 | 1.26E-08 | F | F | T | T |
| Trim27 | 1.28114411 | 8.02832569 | 50.9666666 | 1.14E-09 | 1.27E-08 | T | F | T | F |
| Zdhhc8 | 1.57831455 | 6.23205268 | 50.9490695 | 1.14E-09 | 1.27E-08 | F | F | T | T |
| Gak | 1.33164569 | 7.55056615 | 50.9347832 | 1.15E-09 | 1.28E-08 | F | F | T | T |
| H2aw | -1.4729243 | 7.65735698 | 50.8856826 | 1.16E-09 | 1.29E-08 | F | F | F | F |
| Serpinf2 | -5.0522416 | 3.11244079 | 50.8844695 | 1.16E-09 | 1.29E-08 | F | F | T | F |
| Wwtr1 | 1.57876363 | 6.23780977 | 50.8590696 | 1.17E-09 | 1.30E-08 | T | F | T | T |
| Il6ra | 3.52470139 | 2.91697229 | 50.7934953 | 1.19E-09 | 1.33E-08 | T | F | T | T |
| Scd1 | 1.45625914 | 6.75544934 | 50.7621963 | 1.20E-09 | 1.34E-08 | T | F | T | T |
| Alx3 | -9.9513218 | 2.39826735 | 50.7525895 | 1.21E-09 | 1.34E-08 | F | F | F | T |
| Lrfn4 | -1.8505168 | 6.03102517 | 50.7432878 | 1.21E-09 | 1.34E-08 | F | F | F | F |
| Ptpre | 2.77461948 | 3.63586707 | 50.6772824 | 1.23E-09 | 1.37E-08 | T | F | T | T |
| Cers2 | 1.46679614 | 6.77438109 | 50.6535219 | 1.24E-09 | 1.38E-08 | T | F | F | T |
| Bri3 | -2.1241142 | 5.28085555 | 50.6489332 | 1.24E-09 | 1.38E-08 | T | F | F | F |
| Slc7a9 | -5.2451774 | 2.94201871 | 50.6129114 | 1.26E-09 | 1.39E-08 | F | F | F | F |
| Slc43a2 | -2.1159384 | 5.28533468 | 50.601711 | 1.26E-09 | 1.39E-08 | T | F | T | T |
| Adamts19 | -9.9036573 | 2.34734039 | 50.5841967 | 1.27E-09 | 1.40E-08 | T | F | F | F |
| Bst2 | 2.1172191 | 4.52507274 | 50.5833613 | 1.27E-09 | 1.40E-08 | F | F | T | F |
| Stard9 | -9.9186514 | 2.36503944 | 50.5655251 | 1.27E-09 | 1.41E-08 | T | F | F | F |
| Pde9a | -9.9262851 | 2.36583987 | 50.5271399 | 1.29E-09 | 1.42E-08 | T | F | F | T |
| Hectd3 | 1.78689108 | 5.35262751 | 50.4809303 | 1.30E-09 | 1.44E-08 | T | F | F | T |
| Pls1 | -4.4880772 | 3.2192292 | 50.4760191 | 1.30E-09 | 1.44E-08 | F | F | T | T |
| Tmem121 | -9.9036945 | 2.34611563 | 50.4668734 | 1.31E-09 | 1.44E-08 | T | F | T | F |
| Frat1 | -3.8683092 | 3.51009412 | 50.4507568 | 1.31E-09 | 1.45E-08 | T | F | T | F |
| Evl | -1.7901609 | 6.23822019 | 50.4398736 | 1.32E-09 | 1.45E-08 | F | F | T | F |
| Adamts15 | -2.7900924 | 4.36696769 | 50.4035048 | 1.33E-09 | 1.47E-08 | T | F | F | T |
| Spry4 | -2.639372 | 4.82351409 | 50.3926643 | 1.34E-09 | 1.47E-08 | T | F | T | T |
| Wscd2 | 2.83946458 | 3.68062744 | 50.300097 | 1.37E-09 | 1.51E-08 | T | F | F | T |

| | | | | | | | | | |
|---|---|---|---|---|---|---|---|---|---|
| Atp11b | 1.62968852 | 6.09983967 | 50.2441655 | 1.39E-09 | 1.53E-08 | T | F | T | T |
| Ydjc | 2.36328352 | 4.21325667 | 50.2399152 | 1.40E-09 | 1.53E-08 | T | F | T | F |
| Ell3 | -9.938052 | 2.3758045 | 50.1643416 | 1.43E-09 | 1.57E-08 | F | F | F | F |
| Chpf | 1.63146135 | 6.15355936 | 50.1387496 | 1.44E-09 | 1.58E-08 | F | F | F | F |
| Cryzl1 | 1.94704684 | 4.85495971 | 50.0838216 | 1.46E-09 | 1.60E-08 | T | F | F | F |
| Crip2 | -2.3604935 | 4.94435409 | 50.0459226 | 1.47E-09 | 1.62E-08 | T | T | F | T |
| Abi2 | -1.5511613 | 7.05574506 | 49.9606257 | 1.51E-09 | 1.66E-08 | T | F | T | T |
| Nckap1l | 4.55003573 | 2.72292792 | 49.9486434 | 1.52E-09 | 1.66E-08 | F | F | T | F |
| Spsb4 | -3.9190997 | 3.55976973 | 49.9211606 | 1.53E-09 | 1.67E-08 | F | F | F | F |
| Chd6 | -1.5701892 | 7.09075397 | 49.9166589 | 1.53E-09 | 1.67E-08 | T | F | F | F |
| Tcf7l2 | -2.1295628 | 5.59767556 | 49.8086092 | 1.58E-09 | 1.73E-08 | T | F | T | T |
| Cbx6 | -2.2463247 | 5.09528786 | 49.764387 | 1.60E-09 | 1.75E-08 | T | F | T | T |
| Serpine2 | 1.71797027 | 5.50186421 | 49.7123518 | 1.62E-09 | 1.77E-08 | F | F | F | T |
| Eif1a | 1.31581975 | 7.54016867 | 49.6060747 | 1.67E-09 | 1.83E-08 | T | T | T | T |
| Atp6v0e2 | -6.7319801 | 2.5796173 | 49.4756682 | 1.73E-09 | 1.89E-08 | F | F | T | T |
| Dlx5 | -6.7315833 | 2.57851702 | 49.4555239 | 1.74E-09 | 1.90E-08 | F | F | F | F |
| Wnk4 | -4.5409247 | 3.15188733 | 49.334111 | 1.81E-09 | 1.97E-08 | F | F | F | F |
| Unc80 | -9.8646222 | 2.30823838 | 49.3295998 | 1.81E-09 | 1.97E-08 | T | F | F | T |
| Fgfrl1 | -2.9009902 | 4.24934575 | 49.3203823 | 1.81E-09 | 1.98E-08 | F | F | T | F |
| Fat1 | 1.11146029 | 10.08331 | 49.2899044 | 1.83E-09 | 1.99E-08 | F | F | F | F |
| Mllt6 | 1.4438399 | 6.70894862 | 49.2807516 | 1.83E-09 | 2.00E-08 | F | F | T | T |
| Parm1 | 1.95974361 | 5.01901418 | 49.2799007 | 1.83E-09 | 2.00E-08 | T | F | T | T |
| Mgat5b | -4.9702647 | 3.0282387 | 49.2454724 | 1.85E-09 | 2.01E-08 | F | F | F | F |
| Dmxl2 | 1.69518848 | 5.66749009 | 49.2354706 | 1.86E-09 | 2.02E-08 | T | F | F | T |
| Ext1 | 1.41529073 | 6.82447912 | 49.1711936 | 1.89E-09 | 2.05E-08 | F | F | F | F |
| Hes3 | -6.8537689 | 2.70326876 | 49.1706359 | 1.89E-09 | 2.05E-08 | F | F | T | F |
| Kdelr3 | 2.26020104 | 4.42910249 | 49.156592 | 1.90E-09 | 2.06E-08 | T | F | F | T |
| Ncan | -5.2727069 | 2.90097423 | 49.1486007 | 1.90E-09 | 2.07E-08 | T | F | T | T |
| B4galnt2 | -4.4517622 | 3.18608076 | 49.1362798 | 1.91E-09 | 2.07E-08 | T | F | T | T |
| Dag1 | 1.24198496 | 8.33124393 | 49.1341096 | 1.91E-09 | 2.07E-08 | F | T | T | T |
| Six3 | -4.6889848 | 3.08485458 | 49.0495478 | 1.96E-09 | 2.12E-08 | F | F | T | F |
| Tmem246 | -6.7211957 | 2.56794164 | 49.0460201 | 1.96E-09 | 2.12E-08 | F | F | T | F |
| Sema6a | -1.5139023 | 7.17081093 | 49.0140452 | 1.98E-09 | 2.14E-08 | F | F | T | F |
| Ppargc1b | 2.53433348 | 3.86965447 | 48.9955252 | 1.99E-09 | 2.15E-08 | F | F | F | T |
| Sorcs2 | -3.3628282 | 3.80972169 | 48.9414297 | 2.02E-09 | 2.18E-08 | T | F | T | T |
| Tax1bp3 | 1.67662054 | 5.65774182 | 48.9142283 | 2.04E-09 | 2.20E-08 | F | F | T | F |
| L3mbtl3 | -1.9991842 | 5.60628293 | 48.9111491 | 2.04E-09 | 2.20E-08 | T | F | F | F |
| Iqgap1 | 1.19528149 | 8.99037111 | 48.8860694 | 2.05E-09 | 2.22E-08 | T | F | T | F |
| Kcnc3 | -3.2120201 | 3.97153772 | 48.8655913 | 2.07E-09 | 2.23E-08 | T | F | F | F |
| Cenpk | -9.848151 | 2.29583065 | 48.8401131 | 2.08E-09 | 2.24E-08 | T | F | F | F |
| Mtss2 | -1.7138722 | 6.37943382 | 48.7885427 | 2.11E-09 | 2.28E-08 | T | F | F | F |
| Crtap | 1.59514857 | 6.03314101 | 48.761627 | 2.13E-09 | 2.29E-08 | T | F | F | F |

| | | | | | | | | | |
|---|---|---|---|---|---|---|---|---|---|
| Bcat1 | 1.25599463 | 8.02056762 | 48.6828348 | 2.18E-09 | 2.34E-08 | T | F | T | T |
| 2810459M1 | -3.7967636 | 3.53395512 | 48.6492754 | 2.20E-09 | 2.36E-08 | T | T | T | T |
| Apbb2 | 1.5257043 | 6.28258874 | 48.6197026 | 2.22E-09 | 2.38E-08 | T | F | T | T |
| Vav2 | 1.36387552 | 7.0272326 | 48.5770466 | 2.24E-09 | 2.41E-08 | F | F | F | T |
| Rap1gap | 3.22502734 | 3.23193426 | 48.5732447 | 2.25E-09 | 2.41E-08 | T | F | F | F |
| Slc9a6 | 1.56645313 | 6.06558566 | 48.5408303 | 2.27E-09 | 2.43E-08 | F | F | F | F |
| Tmem245 | 1.31498636 | 7.74487033 | 48.5360282 | 2.27E-09 | 2.44E-08 | T | F | T | T |
| Cdk8 | -1.8849162 | 5.75459316 | 48.5329613 | 2.27E-09 | 2.44E-08 | T | F | F | T |
| Ift27 | -2.9501343 | 4.09289556 | 48.5185686 | 2.28E-09 | 2.45E-08 | F | F | F | F |
| Tab2 | 1.31606788 | 7.3954445 | 48.5002744 | 2.29E-09 | 2.46E-08 | F | F | T | F |
| Bdh1 | 2.45327166 | 3.95648632 | 48.4693341 | 2.31E-09 | 2.48E-08 | F | F | T | F |
| Cchcr1 | -4.724622 | 3.04669459 | 48.4624585 | 2.32E-09 | 2.48E-08 | F | F | F | F |
| Sidt2 | 1.3386138 | 7.16440916 | 48.4532437 | 2.33E-09 | 2.49E-08 | T | F | F | F |
| Dgkd | 1.30228899 | 7.53685179 | 48.4221013 | 2.35E-09 | 2.51E-08 | T | F | T | F |
| Itga6 | -2.0086525 | 5.39533658 | 48.4170917 | 2.35E-09 | 2.51E-08 | T | F | T | T |
| Gse1 | 1.41150057 | 7.00911359 | 48.3958366 | 2.36E-09 | 2.53E-08 | T | F | T | F |
| Acsl5 | 1.63326641 | 5.73681299 | 48.3663471 | 2.38E-09 | 2.55E-08 | F | F | F | T |
| Ccdc88b | 4.24415167 | 2.36839119 | 48.3659087 | 2.38E-09 | 2.55E-08 | F | F | F | F |
| Hbegf | 2.62985705 | 3.74846489 | 48.3364915 | 2.40E-09 | 2.57E-08 | F | F | T | F |
| Zfhx4 | -5.4941123 | 2.83730805 | 48.3206451 | 2.42E-09 | 2.58E-08 | F | F | F | F |
| 1700031F0: | 7.54159351 | 1.16801362 | 48.3071218 | 2.43E-09 | 2.59E-08 | F | F | F | F |
| Prrt3 | -5.912875 | 2.69216465 | 48.3002448 | 2.43E-09 | 2.59E-08 | F | F | F | F |
| Tmem150a | -2.6228404 | 4.47434198 | 48.2966912 | 2.43E-09 | 2.59E-08 | F | F | F | F |
| Lhpp | -2.0849987 | 5.30251729 | 48.2805025 | 2.44E-09 | 2.60E-08 | F | F | F | T |
| Tmpo | -1.3587923 | 8.26292187 | 48.2342724 | 2.48E-09 | 2.64E-08 | T | F | T | F |
| Rbm47 | 1.40331074 | 6.82027269 | 48.1835099 | 2.51E-09 | 2.67E-08 | T | F | T | T |
| Efna4 | -2.8566457 | 4.14763845 | 48.1777586 | 2.52E-09 | 2.68E-08 | F | F | T | F |
| Nmrk1 | 2.12423072 | 4.4091802 | 48.1711284 | 2.52E-09 | 2.68E-08 | F | F | T | F |
| Cobl | -9.8257354 | 2.27454623 | 48.1519013 | 2.54E-09 | 2.69E-08 | F | F | T | F |
| Dlgap3 | -3.1312865 | 3.93062443 | 48.1042437 | 2.57E-09 | 2.73E-08 | T | F | F | T |
| Adcy1 | -3.4391345 | 3.70787516 | 48.0641316 | 2.60E-09 | 2.76E-08 | T | F | T | T |
| Ceacam15 | 3.81908682 | 2.9873231 | 48.0279716 | 2.63E-09 | 2.79E-08 | F | F | T | F |
| Adgra3 | -1.4737379 | 7.33112577 | 48.0026075 | 2.65E-09 | 2.81E-08 | F | F | T | F |
| Lrrn2 | -9.8169298 | 2.26225826 | 47.9952164 | 2.65E-09 | 2.81E-08 | F | F | T | F |
| Sephs1 | -1.5047408 | 7.10596155 | 47.9835486 | 2.66E-09 | 2.82E-08 | F | F | T | F |
| Proser2 | 2.83919184 | 3.45132714 | 47.9476307 | 2.69E-09 | 2.85E-08 | T | F | F | T |
| Ehd1 | -1.5384438 | 6.96457482 | 47.9369537 | 2.70E-09 | 2.86E-08 | F | F | T | T |
| Dlc1 | -1.8348414 | 5.88011689 | 47.9219685 | 2.71E-09 | 2.87E-08 | T | F | T | F |
| Lmo4 | -2.1554853 | 5.05852288 | 47.9172894 | 2.71E-09 | 2.87E-08 | T | F | F | T |
| Fam20c | -9.8100037 | 2.25750417 | 47.9052099 | 2.72E-09 | 2.88E-08 | F | F | F | F |
| Apba3 | 1.56073274 | 6.07699295 | 47.8866155 | 2.74E-09 | 2.89E-08 | F | F | F | F |
| Stx6 | -2.2952211 | 4.81576395 | 47.8751562 | 2.75E-09 | 2.90E-08 | F | F | T | F |

| | | | | | | | | | |
|---|---|---|---|---|---|---|---|---|---|
| Ets1 | -2.9451926 | 4.06588187 | 47.8238993 | 2.79E-09 | 2.94E-08 | T | F | F | T |
| Bid | -2.5902598 | 4.3733491 | 47.7986329 | 2.81E-09 | 2.96E-08 | F | F | F | T |
| Dusp5 | 1.83361081 | 5.31495561 | 47.776457 | 2.83E-09 | 2.98E-08 | F | F | F | F |
| Mafk | 1.89109235 | 4.87715895 | 47.7695743 | 2.83E-09 | 2.98E-08 | F | F | T | F |
| Akr7a5 | -2.2629405 | 4.95094007 | 47.6836707 | 2.90E-09 | 3.06E-08 | F | F | F | F |
| 2900026A0 | 1.42003776 | 6.74423329 | 47.6572318 | 2.93E-09 | 3.08E-08 | T | F | T | T |
| 2700081O1 | -2.0107483 | 5.33540427 | 47.6436893 | 2.94E-09 | 3.09E-08 | T | F | T | T |
| Astn2 | -9.8075421 | 2.25258059 | 47.6374194 | 2.94E-09 | 3.09E-08 | T | F | F | F |
| Ocstamp | 3.25529249 | 3.02551761 | 47.5948224 | 2.98E-09 | 3.13E-08 | T | T | F | T |
| Auts2 | 1.37484916 | 6.99113878 | 47.5608424 | 3.01E-09 | 3.16E-08 | T | F | T | F |
| Gpr162 | -9.8031099 | 2.24842031 | 47.5351084 | 3.03E-09 | 3.18E-08 | F | F | F | F |
| Msh2 | -1.4723336 | 7.3124918 | 47.5267858 | 3.04E-09 | 3.19E-08 | F | F | T | F |
| Inpp5f | -2.1647918 | 5.02971351 | 47.4887676 | 3.07E-09 | 3.22E-08 | T | F | T | T |
| Itgb8 | -9.8278853 | 2.27914917 | 47.4452316 | 3.11E-09 | 3.26E-08 | T | F | T | T |
| Lyst | 1.63295181 | 5.73516816 | 47.4390376 | 3.12E-09 | 3.27E-08 | T | F | F | F |
| Hhex | -6.6685306 | 2.51902554 | 47.4034384 | 3.15E-09 | 3.30E-08 | T | F | T | F |
| Nuak2 | -3.2594553 | 3.81720154 | 47.4006754 | 3.15E-09 | 3.30E-08 | F | F | F | F |
| Scube3 | -3.6896956 | 3.76489288 | 47.3539514 | 3.20E-09 | 3.34E-08 | F | F | T | T |
| Thnsl1 | -5.8863185 | 2.66711778 | 47.3514557 | 3.20E-09 | 3.35E-08 | T | F | T | T |
| Gm12569 | 3.97617939 | 2.4960477 | 47.3217244 | 3.23E-09 | 3.37E-08 | F | F | T | F |
| Stc2 | 2.2296381 | 4.30464006 | 47.3115155 | 3.24E-09 | 3.38E-08 | T | T | T | T |
| Afap1 | 1.50937231 | 6.26403606 | 47.3092829 | 3.24E-09 | 3.38E-08 | T | F | T | F |
| Pon2 | 1.99335501 | 4.62694231 | 47.2907459 | 3.26E-09 | 3.40E-08 | T | F | F | T |
| Fxyd6 | -2.4215421 | 4.64212002 | 47.2718875 | 3.27E-09 | 3.42E-08 | F | F | F | T |
| Pbld1 | -4.8515744 | 2.95921965 | 47.2312496 | 3.31E-09 | 3.45E-08 | T | F | T | F |
| Wnt5b | -3.6107791 | 3.67295092 | 47.2270826 | 3.32E-09 | 3.46E-08 | T | T | T | T |
| Gfra2 | -3.5394236 | 3.6079407 | 47.1629987 | 3.38E-09 | 3.52E-08 | F | F | F | F |
| Trim32 | -1.917748 | 5.57029687 | 47.144842 | 3.40E-09 | 3.54E-08 | T | F | F | F |
| Fam172a | 2.07814531 | 4.46788628 | 47.1388374 | 3.40E-09 | 3.54E-08 | T | T | T | T |
| Sqle | 1.50328893 | 6.30930005 | 47.1375491 | 3.41E-09 | 3.54E-08 | F | F | F | F |
| Map3k10 | -2.5277789 | 4.43163248 | 47.1373642 | 3.41E-09 | 3.54E-08 | F | F | T | F |
| Ptms | -1.6554431 | 6.48024994 | 47.1300232 | 3.41E-09 | 3.55E-08 | F | F | T | F |
| Fzd1 | -2.6227998 | 4.31105936 | 47.1035908 | 3.44E-09 | 3.57E-08 | T | F | T | T |
| Ctsf | -9.7817708 | 2.23145371 | 47.0371823 | 3.51E-09 | 3.64E-08 | F | F | F | F |
| Lrrc55 | -9.7903217 | 2.23521447 | 47.011489 | 3.53E-09 | 3.67E-08 | T | F | F | T |
| Atf4 | 1.27966845 | 7.60632997 | 46.9612068 | 3.59E-09 | 3.72E-08 | F | F | F | F |
| Zswim5 | -3.3769989 | 3.7092421 | 46.9458162 | 3.60E-09 | 3.73E-08 | T | F | F | F |
| Ccni | -1.4382202 | 7.49181159 | 46.9114675 | 3.64E-09 | 3.77E-08 | T | F | T | T |
| Smco4 | -5.1225414 | 2.87115656 | 46.8759498 | 3.68E-09 | 3.81E-08 | F | F | T | F |
| Rcor2 | -1.4791926 | 7.16696881 | 46.7983459 | 3.76E-09 | 3.89E-08 | F | F | F | F |
| Mycbp2 | 1.22164645 | 8.13115281 | 46.7834705 | 3.78E-09 | 3.91E-08 | T | F | T | F |
| Eml4 | -1.4517586 | 7.40187257 | 46.7455877 | 3.82E-09 | 3.95E-08 | T | F | T | F |

| | | | | | | | | | |
|---|---|---|---|---|---|---|---|---|---|
| S1pr2 | -2.0835612 | 5.15219722 | 46.7084124 | 3.86E-09 | 3.99E-08 | T | F | T | F |
| Itih5 | 2.20059171 | 4.21474646 | 46.7025808 | 3.87E-09 | 4.00E-08 | T | F | T | T |
| B4galt3 | 1.58979688 | 5.85750365 | 46.6867289 | 3.89E-09 | 4.01E-08 | F | F | F | F |
| Spns2 | -3.4719375 | 3.67614012 | 46.6107088 | 3.97E-09 | 4.10E-08 | T | F | T | F |
| Pla2g5 | 3.95749162 | 2.55666944 | 46.5609236 | 4.03E-09 | 4.16E-08 | T | T | T | T |
| Zmym2 | -1.6515843 | 6.52344577 | 46.5606068 | 4.03E-09 | 4.16E-08 | F | F | T | T |
| Zbtb43 | -2.2793893 | 4.81283428 | 46.5120355 | 4.09E-09 | 4.21E-08 | T | F | T | F |
| Nav1 | 1.29005203 | 7.57499295 | 46.491557 | 4.11E-09 | 4.24E-08 | F | F | T | T |
| Tmem237 | -3.5369259 | 3.60148541 | 46.4825099 | 4.13E-09 | 4.25E-08 | T | F | F | F |
| Ldlrad3 | -1.8732172 | 5.66810586 | 46.4620327 | 4.15E-09 | 4.27E-08 | T | F | T | T |
| Hspg2 | 1.09954269 | 9.72642096 | 46.4300189 | 4.19E-09 | 4.31E-08 | F | F | F | T |
| 2210418O1 | 1.83534432 | 4.98321168 | 46.4271612 | 4.19E-09 | 4.31E-08 | F | F | T | F |
| Stra6 | -2.156682 | 4.99409136 | 46.3629287 | 4.27E-09 | 4.39E-08 | T | F | F | F |
| Syt7 | -3.0829964 | 3.90771711 | 46.328431 | 4.32E-09 | 4.43E-08 | F | F | T | T |
| Tspyl3 | -9.7523993 | 2.20182054 | 46.3076327 | 4.34E-09 | 4.46E-08 | T | F | F | F |
| Ubc | 1.10429291 | 9.70747296 | 46.3024517 | 4.35E-09 | 4.46E-08 | F | F | F | F |
| Arhgap30 | 5.0011513 | 2.06198497 | 46.2275272 | 4.45E-09 | 4.56E-08 | T | F | F | T |
| Usp12 | 1.40292776 | 6.67461443 | 46.1611188 | 4.53E-09 | 4.65E-08 | T | F | F | T |
| Rock1 | 1.30882687 | 7.2938148 | 46.1567844 | 4.54E-09 | 4.65E-08 | T | F | F | T |
| Irf6 | -2.9331187 | 4.01949274 | 46.1336603 | 4.57E-09 | 4.68E-08 | T | T | T | T |
| Rtp4 | 5.8364686 | 1.64101064 | 46.1288651 | 4.58E-09 | 4.69E-08 | F | F | T | F |
| Gm20671 | 9.51555094 | 0.92274376 | 46.3303724 | 4.62E-09 | 4.72E-08 | F | F | F | F |
| Dok4 | -2.4157503 | 4.85173566 | 46.0625568 | 4.67E-09 | 4.77E-08 | F | F | T | F |
| Aoc3 | 4.49877634 | 2.053625 | 45.9858207 | 4.77E-09 | 4.88E-08 | T | F | T | T |
| Pdgfrb | -2.6125165 | 4.63873263 | 45.9504622 | 4.82E-09 | 4.93E-08 | T | T | T | T |
| Nradd | -4.8577969 | 2.92343369 | 45.9429556 | 4.84E-09 | 4.94E-08 | F | F | F | F |
| Camk1g | -9.7389951 | 2.18910992 | 45.9371688 | 4.84E-09 | 4.94E-08 | F | F | T | F |
| Ssh1 | 1.30250652 | 7.22602136 | 45.9150108 | 4.88E-09 | 4.98E-08 | F | F | T | T |
| Scml2 | -6.6388418 | 2.48638858 | 45.8672444 | 4.94E-09 | 5.04E-08 | T | F | T | T |
| Nectin1 | -1.428997 | 7.41084767 | 45.8403026 | 4.98E-09 | 5.08E-08 | T | F | T | T |
| Pcsk9 | -3.7607207 | 3.40747132 | 45.8140809 | 5.02E-09 | 5.12E-08 | T | F | T | T |
| Myl12b | -1.951029 | 5.37824091 | 45.7248404 | 5.16E-09 | 5.25E-08 | F | F | F | F |
| Magi3 | 1.49468483 | 6.20566338 | 45.6500011 | 5.27E-09 | 5.37E-08 | F | F | F | F |
| Fut9 | -9.8228623 | 2.26169218 | 45.6494507 | 5.27E-09 | 5.37E-08 | T | T | T | T |
| Aebp1 | -2.9166005 | 4.03908918 | 45.6179153 | 5.32E-09 | 5.41E-08 | F | F | F | F |
| Rnf128 | 1.63607138 | 5.54737077 | 45.6126879 | 5.33E-09 | 5.42E-08 | T | F | T | F |
| Meox1 | -6.961091 | 4.26862319 | 45.6056361 | 5.34E-09 | 5.43E-08 | F | F | F | F |
| Dop1a | 1.85515814 | 4.8625606 | 45.5279712 | 5.47E-09 | 5.55E-08 | F | F | F | F |
| Fggy | -9.7232135 | 2.17400714 | 45.5051998 | 5.50E-09 | 5.59E-08 | T | F | T | F |
| Klf7 | 1.70094214 | 5.27940703 | 45.5015534 | 5.51E-09 | 5.59E-08 | F | F | F | T |
| Yars | 1.28676181 | 7.44214059 | 45.4634199 | 5.57E-09 | 5.65E-08 | F | F | T | F |
| H2ac19 | -13.221537 | 5.61657916 | 45.4586805 | 5.58E-09 | 5.66E-08 | F | F | F | F |

| | | | | | | | | | |
|---|---|---|---|---|---|---|---|---|---|
| Gm29695 | -5.8325091 | 2.61360966 | 45.4529402 | 5.59E-09 | 5.66E-08 | F | F | F | F |
| Tex2 | 1.4967992 | 6.15744212 | 45.4166709 | 5.65E-09 | 5.72E-08 | F | F | T | F |
| Gm14288 | -9.7391196 | 2.18485681 | 45.3981919 | 5.68E-09 | 5.75E-08 | T | F | F | F |
| Pla2g4e | 6.1569704 | 1.39875331 | 45.3433358 | 5.77E-09 | 5.84E-08 | T | F | F | T |
| Actb | 0.88898561 | 12.6061891 | 45.2598053 | 5.92E-09 | 5.99E-08 | F | F | T | F |
| Ubxn8 | 1.68298279 | 5.36994677 | 45.251765 | 5.93E-09 | 6.00E-08 | T | F | T | T |
| Lrfn3 | -4.2340661 | 3.1412304 | 45.2429687 | 5.95E-09 | 6.01E-08 | F | F | F | F |
| Lhx6 | 3.98753074 | 2.34316924 | 45.2302725 | 5.97E-09 | 6.03E-08 | T | F | T | T |
| Mfn2 | 1.38063515 | 6.71596477 | 45.1908833 | 6.04E-09 | 6.10E-08 | F | F | T | T |
| Hoxd1 | -6.6741281 | 2.52903842 | 45.1832441 | 6.06E-09 | 6.11E-08 | T | F | T | T |
| Prrg4 | 2.31835415 | 4.26308381 | 45.1541442 | 6.11E-09 | 6.16E-08 | T | F | T | T |
| Igsf10 | -2.9591035 | 3.97057037 | 45.1472793 | 6.12E-09 | 6.17E-08 | F | F | T | F |
| Sipa1 | 1.8030864 | 5.16676256 | 45.1470573 | 6.12E-09 | 6.17E-08 | F | F | T | F |
| Elovl5 | 1.32729597 | 6.96709648 | 45.0832244 | 6.24E-09 | 6.28E-08 | T | F | T | F |
| Galnt10 | -1.7998604 | 5.78127664 | 45.0821347 | 6.24E-09 | 6.28E-08 | F | F | F | F |
| Anp32e | -1.3824371 | 7.64428188 | 45.0795738 | 6.25E-09 | 6.28E-08 | F | F | T | T |
| Trip12 | 1.09239645 | 9.62125597 | 45.0687767 | 6.27E-09 | 6.30E-08 | F | F | F | T |
| Slc25a30 | 1.69723482 | 5.26726762 | 45.027752 | 6.34E-09 | 6.38E-08 | F | F | F | F |
| Zbed3 | -2.3417672 | 4.60237406 | 44.9900433 | 6.41E-09 | 6.45E-08 | T | F | T | F |
| Rnf149 | -1.9393374 | 5.36037639 | 44.9842794 | 6.43E-09 | 6.45E-08 | T | F | F | T |
| Ppp1r15b | 1.20480343 | 8.04590413 | 44.9814414 | 6.43E-09 | 6.46E-08 | F | F | T | F |
| Ddt | -2.1038879 | 5.08391192 | 44.9797876 | 6.43E-09 | 6.46E-08 | F | F | T | F |
| Abcc5 | -1.740281 | 6.00190605 | 44.9296076 | 6.53E-09 | 6.55E-08 | T | F | T | T |
| Lpcat1 | -1.6508248 | 6.31599365 | 44.9218419 | 6.55E-09 | 6.56E-08 | F | F | T | F |
| Dact3 | -5.0836276 | 2.83473795 | 44.8385553 | 6.71E-09 | 6.72E-08 | F | F | F | F |
| Spp2 | -9.7050907 | 2.15866378 | 44.8178364 | 6.75E-09 | 6.76E-08 | F | F | F | F |
| Reep5 | 1.58562928 | 5.72196378 | 44.7743914 | 6.84E-09 | 6.85E-08 | T | F | T | F |
| Zfp2 | -6.4359939 | 2.46873851 | 44.7422466 | 6.91E-09 | 6.91E-08 | F | F | T | F |
| Mef2c | -3.9970761 | 3.3203986 | 44.7281859 | 6.94E-09 | 6.94E-08 | T | F | T | T |
| Vopp1 | -3.5292695 | 3.52254683 | 44.7147915 | 6.96E-09 | 6.96E-08 | T | F | T | T |
| Angptl2 | -4.6799465 | 3.00657972 | 44.6954157 | 7.00E-09 | 7.00E-08 | T | F | F | T |
| Maz | -1.3272929 | 8.00930364 | 44.6788725 | 7.04E-09 | 7.03E-08 | T | F | T | F |
| Ptn | -3.7871713 | 3.61667818 | 44.6489872 | 7.10E-09 | 7.09E-08 | F | F | F | T |
| Jund | 1.4765734 | 6.21127041 | 44.6407576 | 7.12E-09 | 7.10E-08 | F | F | F | T |
| Lmnb2 | -1.6090838 | 6.45286386 | 44.6370219 | 7.13E-09 | 7.11E-08 | F | F | F | F |
| Slc38a2 | 1.19095871 | 8.74562603 | 44.626372 | 7.15E-09 | 7.13E-08 | T | F | T | T |
| Npepl1 | -2.4420422 | 4.49373131 | 44.6241454 | 7.15E-09 | 7.13E-08 | F | F | F | F |
| Tmem170b | 2.38019851 | 3.98525857 | 44.6101778 | 7.18E-09 | 7.16E-08 | F | F | T | T |
| Traf3ip1 | -2.2960115 | 4.65549752 | 44.6057959 | 7.19E-09 | 7.16E-08 | F | F | F | T |
| Limd2 | -1.7640592 | 5.9267696 | 44.5754962 | 7.26E-09 | 7.22E-08 | T | F | T | F |
| Gm9780 | 9.47038458 | 0.87891882 | 44.7899863 | 7.26E-09 | 7.22E-08 | F | F | F | F |
| Tor1aip1 | 1.50237009 | 6.07526557 | 44.5269665 | 7.37E-09 | 7.32E-08 | T | F | F | T |

| | | | | | | | | | |
|---|---|---|---|---|---|---|---|---|---|
| Tuba1a | -1.3087122 | 8.20557097 | 44.4971964 | 7.43E-09 | 7.38E-08 | F | F | F | T |
| Trip11 | 1.38384793 | 6.67844578 | 44.4772935 | 7.48E-09 | 7.43E-08 | F | F | T | T |
| Ttll5 | 1.50832614 | 6.06944355 | 44.4550267 | 7.53E-09 | 7.47E-08 | T | F | F | F |
| Chrd | -3.8874514 | 3.52633155 | 44.4412412 | 7.56E-09 | 7.50E-08 | T | F | T | F |
| Hoxb3 | -6.0661729 | 2.84762856 | 44.432969 | 7.58E-09 | 7.51E-08 | T | F | F | F |
| Col1a1 | -2.3809453 | 5.0448399 | 44.3686958 | 7.72E-09 | 7.66E-08 | F | F | F | F |
| Eda | -3.3741146 | 3.78201705 | 44.2961888 | 7.89E-09 | 7.82E-08 | T | F | T | F |
| Chn2 | -5.8074608 | 2.5931243 | 44.2055047 | 8.11E-09 | 8.03E-08 | T | F | T | F |
| Htatip2 | 3.04716062 | 3.13495949 | 44.1672506 | 8.20E-09 | 8.12E-08 | F | F | T | F |
| Lmf2 | 1.50971476 | 5.99014611 | 44.1629967 | 8.21E-09 | 8.13E-08 | T | F | F | F |
| Vgll1 | 5.91805196 | 1.38167778 | 44.1627739 | 8.22E-09 | 8.13E-08 | F | F | F | T |
| Cyp4f15 | -9.7142689 | 2.15902859 | 44.154667 | 8.24E-09 | 8.14E-08 | T | F | F | F |
| Sept2 | 1.21368724 | 7.88743583 | 44.1501294 | 8.25E-09 | 8.15E-08 | T | F | T | T |
| Arap3 | -3.1319983 | 3.80136121 | 44.1052081 | 8.36E-09 | 8.26E-08 | T | F | F | T |
| Msantd3 | -2.0802683 | 5.00507942 | 44.0417486 | 8.52E-09 | 8.41E-08 | F | F | T | T |
| Pak3 | -9.6682802 | 2.12179588 | 44.0040657 | 8.62E-09 | 8.50E-08 | F | T | T | T |
| Adcy5 | -6.5673001 | 2.42275442 | 43.9855994 | 8.66E-09 | 8.55E-08 | F | F | T | T |
| Tle4 | -1.8297364 | 5.69007583 | 43.9349921 | 8.80E-09 | 8.67E-08 | T | T | F | T |
| Itm2b | -2.0226151 | 5.12515535 | 43.9172879 | 8.84E-09 | 8.72E-08 | T | F | F | T |
| Gal | -9.6857849 | 2.14216773 | 43.8495837 | 9.03E-09 | 8.89E-08 | F | F | F | F |
| Tmem200a | -9.7349922 | 2.17718555 | 43.8323789 | 9.07E-09 | 8.93E-08 | T | F | T | T |
| Nrm | -2.4881058 | 4.32393773 | 43.812775 | 9.13E-09 | 8.98E-08 | F | F | T | F |
| Sun2 | 1.95532803 | 4.53373169 | 43.774314 | 9.23E-09 | 9.08E-08 | T | F | T | F |
| Spred3 | -2.9630367 | 3.93617286 | 43.7403938 | 9.33E-09 | 9.17E-08 | T | F | T | T |
| Hdac11 | -4.0001674 | 3.19948388 | 43.7104481 | 9.41E-09 | 9.25E-08 | F | F | T | F |
| Pear1 | 4.10723437 | 2.939357 | 43.6965495 | 9.45E-09 | 9.29E-08 | F | F | F | F |
| Ryr2 | -4.5331441 | 2.9592498 | 43.645366 | 9.60E-09 | 9.43E-08 | F | F | F | F |
| Dnajc13 | 1.1827667 | 8.09193675 | 43.6264865 | 9.66E-09 | 9.48E-08 | T | F | F | F |
| Xxylt1 | -1.9050585 | 5.37465697 | 43.6113817 | 9.70E-09 | 9.52E-08 | T | F | T | F |
| Chrna7 | -9.65012 | 2.10378212 | 43.5495549 | 9.88E-09 | 9.69E-08 | F | F | T | T |
| Bmf | -2.1621953 | 4.87334017 | 43.5461353 | 9.89E-09 | 9.70E-08 | T | F | F | T |
| Pwwp2b | -2.5754754 | 4.23567389 | 43.5342191 | 9.93E-09 | 9.73E-08 | F | F | F | T |
| H3c14 | -9.6543357 | 2.10492475 | 43.5105182 | 1.00E-08 | 9.79E-08 | F | F | F | T |
| Ift80 | -2.7948652 | 4.05863712 | 43.4970474 | 1.00E-08 | 9.83E-08 | T | F | T | T |
| Slc25a1 | -1.8716218 | 5.46414429 | 43.4963931 | 1.00E-08 | 9.83E-08 | F | F | T | T |
| Il17rb | 6.29924928 | 1.28403002 | 43.4911356 | 1.01E-08 | 9.84E-08 | F | F | F | F |
| Saraf | 1.46360542 | 6.24124649 | 43.4889427 | 1.01E-08 | 9.84E-08 | F | F | T | F |
| Gm19426 | -9.663162 | 2.11182059 | 43.4585668 | 1.02E-08 | 9.93E-08 | F | F | F | F |
| Plxnd1 | 1.60509803 | 5.53328368 | 43.4525752 | 1.02E-08 | 9.94E-08 | F | F | F | F |
| Ldlrap1 | 1.77918068 | 4.94243928 | 43.424476 | 1.03E-08 | 1.00E-07 | F | F | T | T |
| Olfr46 | 1.46125899 | 6.23313602 | 43.3714059 | 1.04E-08 | 1.02E-07 | F | F | F | F |
| Desi2 | 1.41858883 | 6.40521227 | 43.3679854 | 1.04E-08 | 1.02E-07 | F | F | T | F |

| | | | | | | | | | |
|---|---|---|---|---|---|---|---|---|---|
| Klb | -3.7992407 | 3.34857507 | 43.3384398 | 1.05E-08 | 1.03E-07 | F | F | F | T |
| Zbtb34 | -2.0369291 | 5.1197643 | 43.3323616 | 1.06E-08 | 1.03E-07 | T | F | T | T |
| Tpm1 | 1.22162284 | 8.66307797 | 43.3316459 | 1.06E-08 | 1.03E-07 | T | F | T | F |
| Rasl10b | -9.6729578 | 2.13057415 | 43.3016833 | 1.07E-08 | 1.04E-07 | F | F | T | T |
| Fsd1 | -3.9441451 | 3.20803833 | 43.1559722 | 1.11E-08 | 1.08E-07 | F | F | F | F |
| Slc16a2 | -2.9513998 | 3.92533385 | 43.1220653 | 1.12E-08 | 1.09E-07 | T | F | F | T |
| Ilf3 | -1.3016688 | 8.01456657 | 43.1126123 | 1.13E-08 | 1.10E-07 | T | F | T | F |
| Adcy8 | -6.5815975 | 2.47029569 | 43.0906963 | 1.14E-08 | 1.10E-07 | F | F | F | F |
| Mrfap1 | 1.08448706 | 9.36810745 | 43.0734974 | 1.14E-08 | 1.11E-07 | T | F | F | T |
| Sdk1 | -4.9918041 | 2.76095668 | 43.0679651 | 1.14E-08 | 1.11E-07 | F | F | F | F |
| Gdap1l1 | -9.6337203 | 2.08587512 | 43.0560122 | 1.15E-08 | 1.11E-07 | F | F | F | F |
| Zfp385a | -2.1862913 | 4.75549802 | 43.0333517 | 1.16E-08 | 1.12E-07 | F | F | F | F |
| Iqsec2 | -3.3036669 | 3.63973639 | 43.0320766 | 1.16E-08 | 1.12E-07 | T | F | T | T |
| Arhgef10 | -3.1171742 | 3.76000483 | 42.9978412 | 1.17E-08 | 1.13E-07 | T | T | F | T |
| Id2 | -1.9285206 | 5.38822042 | 42.9807348 | 1.17E-08 | 1.14E-07 | F | F | T | F |
| Cars | 1.33046671 | 6.79879455 | 42.9633732 | 1.18E-08 | 1.14E-07 | T | F | T | T |
| Slc37a2 | 3.09747917 | 3.0399968 | 42.9264143 | 1.19E-08 | 1.16E-07 | T | F | T | T |
| Sh3kbp1 | -2.2701883 | 4.61116642 | 42.8836568 | 1.21E-08 | 1.17E-07 | T | F | T | T |
| Tiam2 | -2.8349651 | 3.99409831 | 42.856361 | 1.22E-08 | 1.18E-07 | T | F | F | F |
| Pard6b | -1.7742137 | 5.71211371 | 42.8412497 | 1.22E-08 | 1.18E-07 | F | F | T | T |
| Kctd9 | 1.80559293 | 4.85119923 | 42.8257364 | 1.23E-08 | 1.19E-07 | T | F | T | T |
| Misp3 | -5.312061 | 2.66092349 | 42.8186684 | 1.23E-08 | 1.19E-07 | F | F | F | F |
| Tulp3 | -1.700626 | 5.95981716 | 42.8129328 | 1.24E-08 | 1.19E-07 | T | F | T | T |
| Card14 | -9.7004051 | 2.15954386 | 42.8113465 | 1.24E-08 | 1.19E-07 | F | F | F | F |
| Tbx1 | -9.8159202 | 2.27351959 | 42.8067591 | 1.24E-08 | 1.19E-07 | F | F | T | F |
| Impa2 | 1.72605207 | 5.13161442 | 42.7803679 | 1.25E-08 | 1.20E-07 | T | F | T | T |
| Msl3 | 2.01170294 | 4.36764987 | 42.7523854 | 1.26E-08 | 1.21E-07 | T | T | T | T |
| Psip1 | -1.2944906 | 8.1193128 | 42.7515231 | 1.26E-08 | 1.21E-07 | T | T | T | F |
| Prl7c1 | 7.36614536 | 1.04133595 | 42.7308347 | 1.27E-08 | 1.22E-07 | F | F | F | F |
| Ccdc102a | -2.7035453 | 4.09625547 | 42.7297683 | 1.27E-08 | 1.22E-07 | F | F | F | F |
| Pmpca | 1.21852769 | 7.59690939 | 42.7142385 | 1.27E-08 | 1.22E-07 | T | F | F | T |
| Tfrc | 1.25144296 | 7.55083387 | 42.6841439 | 1.28E-08 | 1.24E-07 | T | F | F | F |
| Mob3b | -2.5268403 | 4.26403545 | 42.651207 | 1.30E-08 | 1.25E-07 | F | F | T | T |
| Piezo2 | -3.3295503 | 3.61086341 | 42.6431997 | 1.30E-08 | 1.25E-07 | F | F | T | F |
| Rtn4rl2 | -9.6178528 | 2.07027705 | 42.6123551 | 1.31E-08 | 1.26E-07 | F | F | F | T |
| Sun1 | 1.38314386 | 6.50412488 | 42.5736888 | 1.33E-08 | 1.28E-07 | T | F | T | F |
| Grin2c | -5.0921991 | 2.83551887 | 42.5609401 | 1.33E-08 | 1.28E-07 | F | F | F | F |
| Rpia | 2.00709415 | 4.36361306 | 42.558091 | 1.34E-08 | 1.28E-07 | T | F | F | F |
| Ajap1 | -9.6209654 | 2.07202337 | 42.5291798 | 1.35E-08 | 1.29E-07 | T | F | F | F |
| Tmem119 | -9.7168231 | 2.17638266 | 42.5130683 | 1.35E-08 | 1.30E-07 | F | F | F | F |
| Pklr | -9.6523936 | 2.11174332 | 42.4774227 | 1.37E-08 | 1.31E-07 | T | F | T | T |
| Grem2 | 3.8834009 | 2.34700378 | 42.4135514 | 1.40E-08 | 1.34E-07 | T | F | T | T |

| | | | | | | | | | |
|---|---|---|---|---|---|---|---|---|---|
| Gm37240 | 3.94724926 | 2.23286196 | 42.4020854 | 1.40E-08 | 1.34E-07 | F | F | F | F |
| Pcnx2 | -6.6188791 | 2.46256322 | 42.4017932 | 1.40E-08 | 1.34E-07 | F | F | T | F |
| Tmem30a | 1.50427591 | 5.89166959 | 42.3737901 | 1.41E-08 | 1.35E-07 | T | F | T | F |
| Rapgef6 | 1.31513171 | 6.86745788 | 42.3143835 | 1.44E-08 | 1.37E-07 | T | F | F | T |
| Samd3 | -9.6663396 | 2.11100437 | 42.2441767 | 1.47E-08 | 1.40E-07 | T | F | T | F |
| Gne | 1.91966727 | 4.77377317 | 42.2388835 | 1.47E-08 | 1.41E-07 | F | F | T | F |
| Mfsd13a | -3.1767316 | 3.81555536 | 42.2347189 | 1.47E-08 | 1.41E-07 | F | F | F | F |
| Pcsk5 | -3.6189228 | 3.42421892 | 42.2338604 | 1.47E-08 | 1.41E-07 | T | T | T | T |
| Fos | 2.6539672 | 3.46579542 | 42.215662 | 1.48E-08 | 1.41E-07 | T | F | F | T |
| Diaph3 | 1.53999475 | 5.67715529 | 42.1990077 | 1.49E-08 | 1.42E-07 | F | F | F | F |
| Clstn1 | -1.479559 | 6.78400735 | 42.1834711 | 1.50E-08 | 1.43E-07 | F | F | T | F |
| Ace | 3.42671167 | 2.67157303 | 42.1733677 | 1.50E-08 | 1.43E-07 | F | F | T | F |
| Ro60 | -1.7684571 | 5.75188352 | 42.1589015 | 1.51E-08 | 1.44E-07 | T | F | T | T |
| Plekhn1 | -2.7604875 | 4.02735043 | 42.1500823 | 1.51E-08 | 1.44E-07 | F | F | F | F |
| Fktn | 1.59190013 | 5.52836997 | 42.1365977 | 1.52E-08 | 1.44E-07 | T | F | T | T |
| Plekha1 | -1.8631171 | 5.4054031 | 42.1187073 | 1.53E-08 | 1.45E-07 | T | F | T | T |
| Ppfia4 | -3.4743183 | 3.46087699 | 42.1163522 | 1.53E-08 | 1.45E-07 | T | F | T | F |
| Rsph3a | -4.1492271 | 3.06873507 | 42.1115519 | 1.53E-08 | 1.45E-07 | T | F | F | F |
| Prl2c3 | 3.44243318 | 2.92564104 | 42.1112855 | 1.53E-08 | 1.45E-07 | F | F | F | F |
| 2610008E1 | -2.108838 | 4.84940192 | 41.9971349 | 1.59E-08 | 1.50E-07 | F | F | T | F |
| Usp9x | 1.08267648 | 9.31977871 | 41.954657 | 1.61E-08 | 1.52E-07 | T | F | T | T |
| Ppm1e | -9.5877332 | 2.04427263 | 41.915842 | 1.63E-08 | 1.54E-07 | T | F | T | T |
| Spag9 | 1.26283408 | 7.07420082 | 41.9100448 | 1.63E-08 | 1.54E-07 | T | F | T | T |
| Rrp1 | -1.3810339 | 7.23551406 | 41.8775513 | 1.64E-08 | 1.56E-07 | T | F | T | T |
| Soga3 | -3.9531778 | 3.15415193 | 41.782041 | 1.69E-08 | 1.60E-07 | F | F | F | F |
| Dgke | 1.76797994 | 4.89186946 | 41.7757039 | 1.70E-08 | 1.61E-07 | F | F | T | T |
| Pax2 | -5.2897228 | 3.03706153 | 41.7645901 | 1.70E-08 | 1.61E-07 | F | F | T | F |
| Fbxl20 | -2.0490582 | 4.97882955 | 41.7194204 | 1.73E-08 | 1.63E-07 | T | F | T | T |
| E2f2 | 1.89855491 | 4.55917245 | 41.7178917 | 1.73E-08 | 1.63E-07 | F | F | T | F |
| Dock4 | -2.4978382 | 4.27856855 | 41.695943 | 1.74E-08 | 1.64E-07 | T | F | F | T |
| Zfp292 | -1.4587729 | 6.92389336 | 41.6708148 | 1.75E-08 | 1.66E-07 | T | F | F | T |
| Bcap29 | 2.42035523 | 3.76713058 | 41.6488918 | 1.76E-08 | 1.67E-07 | F | F | T | T |
| Prkaa2 | -3.7911317 | 3.23491939 | 41.6411223 | 1.77E-08 | 1.67E-07 | T | T | T | T |
| Mmp15 | -1.5664254 | 6.37695437 | 41.56277 | 1.81E-08 | 1.71E-07 | T | F | T | T |
| Map4 | 1.1400538 | 8.3163313 | 41.5499607 | 1.82E-08 | 1.71E-07 | T | F | T | T |
| Fam57a | 1.90001661 | 4.56590568 | 41.5475582 | 1.82E-08 | 1.72E-07 | F | F | F | F |
| Lratd2 | -1.7632392 | 5.67045578 | 41.5467208 | 1.82E-08 | 1.72E-07 | F | F | T | T |
| Itm2c | -1.7751114 | 5.69654512 | 41.5436587 | 1.82E-08 | 1.72E-07 | F | F | T | F |
| H4c9 | -1.9130441 | 6.39227443 | 41.5402334 | 1.82E-08 | 1.72E-07 | F | F | F | T |
| Flnc | 1.23601384 | 7.90779733 | 41.5370753 | 1.83E-08 | 1.72E-07 | F | F | T | F |
| Epdr1 | -5.7140796 | 2.50274882 | 41.5193134 | 1.84E-08 | 1.73E-07 | T | F | T | F |
| Ctsl | 1.12185468 | 8.81261279 | 41.4864659 | 1.85E-08 | 1.74E-07 | T | F | T | T |

| | | | | | | | | | |
|---|---|---|---|---|---|---|---|---|---|
| Cd151 | 1.4878656 | 5.90350525 | 41.4794938 | 1.86E-08 | 1.75E-07 | F | F | F | F |
| Tgfb1i1 | -6.4717517 | 2.33064678 | 41.4459439 | 1.88E-08 | 1.76E-07 | F | F | T | T |
| Adgrl2 | 1.15476225 | 8.19395655 | 41.4381545 | 1.88E-08 | 1.77E-07 | T | F | F | F |
| Nudt10 | -4.4864512 | 2.89599003 | 41.3794195 | 1.92E-08 | 1.80E-07 | T | F | T | F |
| Nxn | -1.7641126 | 5.71643739 | 41.3199028 | 1.95E-08 | 1.83E-07 | F | F | T | F |
| Zfp62 | -2.1803956 | 4.68255068 | 41.2658723 | 1.99E-08 | 1.86E-07 | T | F | F | T |
| Dcp1b | -9.5671102 | 2.02102747 | 41.2637844 | 1.99E-08 | 1.86E-07 | F | F | F | F |
| Gramd4 | -2.9020246 | 3.87984519 | 41.2559457 | 1.99E-08 | 1.87E-07 | F | F | T | F |
| Sema6c | -6.4282907 | 2.49562916 | 41.2208388 | 2.01E-08 | 1.89E-07 | F | F | T | F |
| Rnf213 | -1.4625451 | 7.11638752 | 41.1924982 | 2.03E-08 | 1.90E-07 | F | F | F | F |
| Ubn1 | 1.32120104 | 6.71513981 | 41.1806104 | 2.04E-08 | 1.91E-07 | F | F | F | F |
| Snhg4 | -10.083548 | 2.53390041 | 41.175284 | 2.04E-08 | 1.91E-07 | F | F | F | F |
| Cdk19 | 2.41880554 | 4.4363 | 41.1463392 | 2.06E-08 | 1.93E-07 | F | F | T | T |
| Adap2 | 3.86854261 | 2.29271697 | 41.1393391 | 2.06E-08 | 1.93E-07 | F | F | T | F |
| Amer2 | -9.5565004 | 2.01352719 | 41.1389787 | 2.07E-08 | 1.93E-07 | F | F | T | T |
| Tmem168 | 1.56556163 | 5.51530533 | 41.1293974 | 2.07E-08 | 1.93E-07 | F | F | T | T |
| Hoxb4 | -6.0512923 | 2.59168694 | 41.0855745 | 2.10E-08 | 1.96E-07 | F | F | F | F |
| Gabra3 | -9.5597143 | 2.01381719 | 41.065232 | 2.11E-08 | 1.97E-07 | T | F | T | T |
| Stat5b | -2.3503922 | 4.38603343 | 41.055226 | 2.12E-08 | 1.98E-07 | T | F | T | T |
| Pkm | -0.9806186 | 11.5791388 | 41.0476669 | 2.12E-08 | 1.98E-07 | F | F | F | F |
| Plxna4 | -4.2792425 | 3.01765548 | 41.0216359 | 2.14E-08 | 1.99E-07 | F | F | T | T |
| Trp73 | -6.4789055 | 2.33067683 | 41.0144377 | 2.15E-08 | 2.00E-07 | F | F | T | F |
| Glud1 | 1.14558208 | 8.16450674 | 40.9569482 | 2.18E-08 | 2.03E-07 | F | F | T | T |
| Hoxa3 | -6.6457247 | 2.5048057 | 40.947266 | 2.19E-08 | 2.04E-07 | T | F | T | T |
| Galnt9 | -9.5870786 | 2.03614445 | 40.9135426 | 2.21E-08 | 2.06E-07 | F | F | F | F |
| Cdh20 | -9.5821489 | 2.03175534 | 40.8995549 | 2.22E-08 | 2.07E-07 | T | F | T | T |
| Hlx | -10.010046 | 2.46298918 | 40.8723933 | 2.24E-08 | 2.08E-07 | F | F | F | F |
| Trim71 | -1.1232934 | 9.81287725 | 40.8364939 | 2.27E-08 | 2.11E-07 | F | F | T | T |
| Ptges2 | 1.81670039 | 4.71995494 | 40.8266539 | 2.28E-08 | 2.11E-07 | F | F | F | T |
| Tagln | -6.5111811 | 2.37260409 | 40.7961417 | 2.30E-08 | 2.13E-07 | F | F | F | F |
| Adam22 | -3.3099105 | 3.52254713 | 40.7900426 | 2.30E-08 | 2.13E-07 | T | F | T | F |
| Tarsl2 | -9.595449 | 2.05828781 | 40.6984675 | 2.37E-08 | 2.19E-07 | T | F | F | T |
| Lrguk | -3.6338503 | 3.28742017 | 40.6745072 | 2.39E-08 | 2.21E-07 | T | F | T | T |
| Rffl | 1.46062237 | 5.93237819 | 40.6584451 | 2.40E-08 | 2.22E-07 | T | F | T | F |
| Cdk4 | -1.2467732 | 8.31582975 | 40.6422009 | 2.41E-08 | 2.23E-07 | T | T | F | T |
| Cluh | 1.18553578 | 7.62920761 | 40.6344346 | 2.42E-08 | 2.24E-07 | T | F | F | F |
| Rubcn | 1.5512817 | 5.51283712 | 40.5965109 | 2.44E-08 | 2.26E-07 | T | F | T | F |
| Tmem198b | -3.3768569 | 3.45897924 | 40.5927519 | 2.45E-08 | 2.26E-07 | F | F | T | F |
| Arsi | -4.3962585 | 2.94337245 | 40.5804031 | 2.46E-08 | 2.27E-07 | F | F | F | F |
| Kalrn | -2.5289281 | 4.41041229 | 40.5397435 | 2.49E-08 | 2.30E-07 | T | F | T | F |
| B4galt7 | 1.67676383 | 5.07419194 | 40.5260362 | 2.50E-08 | 2.31E-07 | F | F | F | F |
| Mpdz | -1.4285599 | 6.93374499 | 40.5105604 | 2.51E-08 | 2.32E-07 | T | F | F | T |

| | | | | | | | | | |
|---|---|---|---|---|---|---|---|---|---|
| Vmn2r53 | 1.26662754 | 7.2737488 | 40.4818026 | 2.53E-08 | 2.34E-07 | F | F | F | F |
| Eid2 | -3.3928367 | 3.46685878 | 40.4477664 | 2.56E-08 | 2.36E-07 | F | F | F | F |
| Cacnb2 | -9.5331259 | 1.98837433 | 40.4094367 | 2.59E-08 | 2.39E-07 | T | F | F | F |
| Epop | 1.21676494 | 8.10360835 | 40.4061922 | 2.59E-08 | 2.39E-07 | F | F | T | F |
| Cdx4 | -7.2220106 | 3.0632851 | 40.401821 | 2.60E-08 | 2.39E-07 | F | F | F | F |
| Rasl11a | -3.46832 | 3.38781774 | 40.3862854 | 2.61E-08 | 2.40E-07 | F | F | F | F |
| Pxn | 1.29897684 | 6.82481598 | 40.3680078 | 2.62E-08 | 2.41E-07 | T | F | T | F |
| Rab32 | -4.9858043 | 2.66042339 | 40.3668243 | 2.63E-08 | 2.41E-07 | F | F | T | F |
| Ikzf4 | -4.6033995 | 2.84812565 | 40.3284842 | 2.66E-08 | 2.44E-07 | T | F | T | T |
| Robo3 | -9.5557609 | 2.00658127 | 40.3192387 | 2.66E-08 | 2.45E-07 | F | F | T | F |
| Ildr1 | 2.70884924 | 3.46506842 | 40.270397 | 2.71E-08 | 2.49E-07 | T | F | F | T |
| Gcnt4 | 2.81508113 | 3.22921745 | 40.2559717 | 2.72E-08 | 2.50E-07 | F | F | T | F |
| Bbs10 | -9.5245756 | 1.9804899 | 40.2287701 | 2.74E-08 | 2.51E-07 | F | F | F | T |
| Rarb | -5.3732372 | 2.72456011 | 40.2272777 | 2.74E-08 | 2.51E-07 | T | F | T | T |
| Wdr6 | -1.3205728 | 7.44474696 | 40.2272675 | 2.74E-08 | 2.51E-07 | F | F | F | F |
| Akr1c19 | -9.5328253 | 1.98668699 | 40.2136411 | 2.75E-08 | 2.52E-07 | F | F | F | F |
| Six4 | -1.6734171 | 5.91046995 | 40.2029121 | 2.76E-08 | 2.53E-07 | T | F | T | T |
| Kmt2c | 1.19856498 | 7.45438262 | 40.1959412 | 2.77E-08 | 2.54E-07 | T | F | F | T |
| Ddx39b | -1.2215944 | 8.46050491 | 40.195039 | 2.77E-08 | 2.54E-07 | T | F | F | F |
| Zfp24 | -1.65829 | 5.91231261 | 40.1699708 | 2.79E-08 | 2.55E-07 | T | F | T | F |
| Bcl2l2 | 1.62830384 | 5.17440128 | 40.1383659 | 2.82E-08 | 2.58E-07 | T | F | T | T |
| Zfp644 | -1.5356965 | 6.41904165 | 40.1322875 | 2.82E-08 | 2.58E-07 | T | F | T | T |
| Zscan25 | -3.7160447 | 3.27611574 | 40.1204031 | 2.84E-08 | 2.59E-07 | F | F | F | T |
| Cfl2 | 1.71980138 | 4.93422127 | 40.1099884 | 2.84E-08 | 2.60E-07 | F | F | T | F |
| Arid1a | -1.1455017 | 9.44656099 | 40.0802624 | 2.87E-08 | 2.62E-07 | T | F | T | T |
| Abhd17c | -1.5989838 | 6.1455994 | 40.0706376 | 2.88E-08 | 2.63E-07 | F | F | T | T |
| Tmem106a | 2.76641911 | 3.24065023 | 40.0580692 | 2.89E-08 | 2.64E-07 | F | F | T | T |
| Ptprd | -3.0571385 | 3.70275845 | 40.0576924 | 2.89E-08 | 2.64E-07 | T | F | T | T |
| Ntrk2 | -9.5317906 | 1.98489115 | 40.0449668 | 2.90E-08 | 2.65E-07 | F | F | T | F |
| 0610040J01 | -4.9174016 | 2.67129806 | 40.03792 | 2.91E-08 | 2.65E-07 | F | F | T | T |
| BC016579 | 4.76933938 | 1.66800264 | 39.9680112 | 2.97E-08 | 2.71E-07 | F | F | F | F |
| Birc6 | 1.03338877 | 9.67001585 | 39.9441441 | 3.00E-08 | 2.73E-07 | F | F | F | F |
| Kcnt2 | -9.532319 | 1.98480756 | 39.935517 | 3.00E-08 | 2.73E-07 | F | F | F | F |
| Shoc2 | 1.27862372 | 6.83566994 | 39.9208561 | 3.02E-08 | 2.75E-07 | T | F | F | F |
| Arl3 | -2.6371241 | 4.09714791 | 39.8557951 | 3.08E-08 | 2.80E-07 | F | F | F | T |
| Pla2g4f | 4.34147329 | 1.84984299 | 39.851341 | 3.08E-08 | 2.80E-07 | F | F | F | F |
| Tspan2 | -3.9005031 | 3.10505973 | 39.8293543 | 3.11E-08 | 2.82E-07 | F | F | F | F |
| Ssbp2 | -4.4888351 | 2.82055157 | 39.7996935 | 3.13E-08 | 2.85E-07 | T | F | T | T |
| Cyp39a1 | -4.9125885 | 2.66608686 | 39.7724096 | 3.16E-08 | 2.87E-07 | T | F | F | F |
| 2310030G0 | 2.56366965 | 3.47029191 | 39.7631621 | 3.17E-08 | 2.88E-07 | T | F | T | F |
| Apeh | -1.8315092 | 5.27027307 | 39.7600583 | 3.17E-08 | 2.88E-07 | F | F | F | F |
| Fam53c | -1.6367902 | 5.98361554 | 39.7584957 | 3.18E-08 | 2.88E-07 | T | F | T | F |

| | | | | | | | | | |
|---|---|---|---|---|---|---|---|---|---|
| Zfp811 | -9.4977868 | 1.95584867 | 39.6397572 | 3.30E-08 | 2.99E-07 | F | T | T | T |
| Prr36 | -4.5566358 | 2.8815768 | 39.6356174 | 3.30E-08 | 2.99E-07 | F | F | T | F |
| Slc7a6 | -1.6495857 | 5.87552108 | 39.6337059 | 3.30E-08 | 2.99E-07 | T | F | T | F |
| Mbd1 | 1.39911382 | 6.15443998 | 39.6008037 | 3.34E-08 | 3.02E-07 | T | F | F | F |
| mt-Co2 | 1.8559073 | 5.53870882 | 39.5875158 | 3.35E-08 | 3.03E-07 | F | F | F | F |
| Lamb3 | -9.4927528 | 1.95253658 | 39.5480307 | 3.39E-08 | 3.07E-07 | F | F | F | T |
| Tacstd2 | 2.97401898 | 3.13757422 | 39.5194794 | 3.42E-08 | 3.10E-07 | F | F | F | F |
| Cry2 | -1.8847789 | 5.14136886 | 39.4998863 | 3.44E-08 | 3.11E-07 | T | F | T | T |
| Igip | 4.22603909 | 1.98722892 | 39.4876102 | 3.46E-08 | 3.12E-07 | F | F | F | F |
| Csrnp3 | -4.0215113 | 3.02894016 | 39.4805911 | 3.47E-08 | 3.13E-07 | T | T | T | T |
| Vps13c | 1.27487822 | 6.9757073 | 39.4476518 | 3.50E-08 | 3.16E-07 | T | F | F | F |
| Ecpas | 1.16496456 | 7.72644411 | 39.3841701 | 3.57E-08 | 3.22E-07 | F | F | F | F |
| H2bc13 | -1.492131 | 6.50340458 | 39.3417403 | 3.62E-08 | 3.27E-07 | F | F | F | F |
| Pex5 | 1.36646049 | 6.32475811 | 39.311387 | 3.65E-08 | 3.30E-07 | F | F | T | F |
| Arg1 | -6.7992162 | 3.55954789 | 39.3047669 | 3.66E-08 | 3.30E-07 | F | F | F | F |
| Srpr | 1.21985796 | 7.18028596 | 39.3000217 | 3.67E-08 | 3.30E-07 | F | F | T | F |
| Cdkn1a | -2.4780935 | 4.18936802 | 39.2986667 | 3.67E-08 | 3.30E-07 | T | F | T | F |
| Galns | 1.91975915 | 4.36746321 | 39.2861053 | 3.68E-08 | 3.32E-07 | F | F | T | F |
| Gna12 | -1.3946661 | 6.88662642 | 39.2412239 | 3.74E-08 | 3.36E-07 | F | F | F | F |
| Sinhcaf | -1.228027 | 8.1879518 | 39.2351793 | 3.74E-08 | 3.37E-07 | F | F | T | F |
| Tex13b | 5.32192204 | 1.3516095 | 39.2344839 | 3.74E-08 | 3.37E-07 | F | F | T | F |
| Kdm2b | -1.3085224 | 7.39552319 | 39.2037197 | 3.78E-08 | 3.40E-07 | T | T | F | T |
| Ptprz1 | -5.7033839 | 2.49651118 | 39.1578609 | 3.84E-08 | 3.45E-07 | T | F | T | T |
| Cdx2 | -1.652976 | 6.52722264 | 39.1524025 | 3.84E-08 | 3.45E-07 | F | F | F | F |
| Bcas2 | -1.7266754 | 5.5884579 | 39.1372245 | 3.86E-08 | 3.47E-07 | F | F | F | F |
| Nabp1 | 1.67471056 | 5.03638281 | 39.1340336 | 3.87E-08 | 3.47E-07 | T | F | T | T |
| Prr12 | -1.2334935 | 8.18311481 | 39.1304319 | 3.87E-08 | 3.47E-07 | F | F | F | F |
| Nog | -4.9088415 | 2.66122525 | 39.1244527 | 3.88E-08 | 3.48E-07 | F | F | T | F |
| Wasf3 | -1.8347099 | 5.22550534 | 39.1231651 | 3.88E-08 | 3.48E-07 | T | F | F | T |
| Mtf1 | 1.56954553 | 5.29286489 | 39.0722144 | 3.94E-08 | 3.53E-07 | T | F | T | T |
| Traf6 | -1.8835808 | 5.11845543 | 39.0582672 | 3.96E-08 | 3.54E-07 | T | F | T | T |
| Ambp | -9.4722337 | 1.9340576 | 39.0147184 | 4.01E-08 | 3.59E-07 | F | F | F | F |
| Pcyt1b | -3.5345896 | 3.28897233 | 39.0019896 | 4.03E-08 | 3.61E-07 | T | F | T | T |
| Mex3d | -1.5721423 | 6.1261914 | 38.9971542 | 4.04E-08 | 3.61E-07 | T | F | T | F |
| Knop1 | -1.2888958 | 7.52069803 | 38.9305304 | 4.12E-08 | 3.68E-07 | F | F | T | T |
| Scamp1 | 1.48517644 | 5.66767741 | 38.9087501 | 4.15E-08 | 3.71E-07 | T | T | T | T |
| Lrrc20 | 2.05299371 | 4.15130806 | 38.8768692 | 4.19E-08 | 3.74E-07 | T | F | T | T |
| Otud5 | 1.18230907 | 7.37842866 | 38.8767097 | 4.19E-08 | 3.74E-07 | T | F | F | F |
| Cyp20a1 | 1.39348409 | 6.14652256 | 38.8653762 | 4.21E-08 | 3.76E-07 | F | T | F | T |
| Chchd7 | -2.8659315 | 3.80032235 | 38.8423895 | 4.24E-08 | 3.78E-07 | T | F | T | F |
| Tmco4 | 2.5042803 | 3.50193684 | 38.8355435 | 4.25E-08 | 3.79E-07 | T | F | F | F |
| Lcor | 1.26220097 | 6.8159739 | 38.8315709 | 4.25E-08 | 3.79E-07 | T | F | T | T |

| | | | | | | | | | |
|---|---|---|---|---|---|---|---|---|---|
| Vma21 | 1.61962045 | 5.13868757 | 38.8273135 | 4.26E-08 | 3.79E-07 | T | F | T | T |
| Plac9a | 5.56791651 | 1.36700225 | 38.81321 | 4.28E-08 | 3.81E-07 | F | F | F | F |
| Megf8 | -1.4088987 | 6.8840871 | 38.8018031 | 4.29E-08 | 3.82E-07 | T | F | F | F |
| Tbc1d32 | -3.8515279 | 3.08927848 | 38.7786183 | 4.33E-08 | 3.85E-07 | T | F | F | T |
| Paics | -1.17446 | 8.69202022 | 38.7766712 | 4.33E-08 | 3.85E-07 | T | F | F | F |
| Ankib1 | 1.44777871 | 5.85034216 | 38.7652396 | 4.34E-08 | 3.86E-07 | T | F | T | F |
| Iqce | -2.4971695 | 4.147158 | 38.744979 | 4.37E-08 | 3.89E-07 | T | F | T | T |
| Pcolce2 | -4.8825771 | 2.64136487 | 38.7369891 | 4.38E-08 | 3.89E-07 | T | F | F | T |
| Sclt1 | -9.4633293 | 1.92659312 | 38.7303302 | 4.39E-08 | 3.90E-07 | T | F | F | F |
| Ifi35 | 5.33675347 | 1.47620592 | 38.7232786 | 4.40E-08 | 3.91E-07 | F | F | F | F |
| Zfp395 | -1.7065517 | 5.59943751 | 38.7141375 | 4.41E-08 | 3.92E-07 | T | F | T | F |
| Usp11 | -2.6208558 | 4.05634909 | 38.7049627 | 4.43E-08 | 3.93E-07 | T | F | F | F |
| Blzf1 | 1.6308077 | 5.08950996 | 38.695963 | 4.44E-08 | 3.94E-07 | T | F | T | T |
| Zfp13 | -2.3552806 | 4.29569368 | 38.6874913 | 4.45E-08 | 3.95E-07 | F | F | F | F |
| Dpyd | -9.4765541 | 1.93169909 | 38.6840085 | 4.46E-08 | 3.95E-07 | T | T | F | T |
| Slc37a1 | 2.4470361 | 3.62905556 | 38.6674624 | 4.48E-08 | 3.97E-07 | F | F | F | F |
| Hnrnpab | -1.0603942 | 10.112908 | 38.6281015 | 4.54E-08 | 4.02E-07 | T | F | F | F |
| Ablim1 | -1.9986063 | 4.90044842 | 38.5797446 | 4.61E-08 | 4.08E-07 | T | F | T | T |
| Masp1 | -9.554559 | 2.02183599 | 38.5614548 | 4.63E-08 | 4.10E-07 | T | F | T | T |
| Dcaf12 | 1.21127505 | 7.07326661 | 38.5232301 | 4.69E-08 | 4.15E-07 | F | F | T | F |
| Tmem127 | 1.30939235 | 6.53566577 | 38.5145319 | 4.70E-08 | 4.16E-07 | T | F | F | F |
| Trps1 | -3.4356681 | 3.3565449 | 38.4952717 | 4.73E-08 | 4.18E-07 | T | F | T | T |
| Cask | -1.8134894 | 5.29721107 | 38.4914831 | 4.74E-08 | 4.18E-07 | T | F | T | F |
| Thra | -2.9032179 | 3.7608251 | 38.4655828 | 4.78E-08 | 4.22E-07 | T | F | T | F |
| Elovl4 | -4.3082457 | 2.86021007 | 38.4218721 | 4.84E-08 | 4.27E-07 | T | F | F | T |
| Edem1 | 1.20741928 | 7.14438125 | 38.3665546 | 4.93E-08 | 4.35E-07 | T | F | T | T |
| Golga2 | 1.41320904 | 5.96757199 | 38.3435379 | 4.97E-08 | 4.38E-07 | F | F | T | F |
| Syt11 | -1.8919288 | 5.16365584 | 38.340585 | 4.97E-08 | 4.38E-07 | T | F | T | T |
| Unkl | 1.6396379 | 5.08899881 | 38.3322325 | 4.98E-08 | 4.39E-07 | T | F | F | F |
| Shank1 | -2.8074419 | 3.85066344 | 38.3138427 | 5.01E-08 | 4.41E-07 | F | F | T | F |
| Tle3 | 1.0453959 | 9.02342963 | 38.2944773 | 5.05E-08 | 4.44E-07 | F | F | T | F |
| Zbtb11 | 1.42375493 | 5.90397986 | 38.2745139 | 5.08E-08 | 4.47E-07 | T | F | F | F |
| Arglu1 | -1.7009674 | 5.57670268 | 38.2538628 | 5.11E-08 | 4.49E-07 | F | F | F | T |
| Hist1h2an | -1.320183 | 7.1299838 | 38.2142687 | 5.18E-08 | 4.55E-07 | F | F | F | F |
| Hey1 | -4.6430565 | 2.71892209 | 38.1764077 | 5.24E-08 | 4.60E-07 | T | F | T | T |
| Entpd4 | 1.85405971 | 4.46380173 | 38.1383141 | 5.30E-08 | 4.66E-07 | F | F | T | F |
| Capn15 | 1.41406322 | 5.95333561 | 38.1268434 | 5.32E-08 | 4.67E-07 | F | F | F | F |
| Rapgef1 | 1.09175337 | 8.31779539 | 38.0856047 | 5.39E-08 | 4.73E-07 | T | F | F | F |
| Schip1 | -2.0437732 | 4.74301932 | 38.0609999 | 5.44E-08 | 4.77E-07 | T | F | T | F |
| Cdc20 | -1.2551103 | 7.67664467 | 38.0398676 | 5.47E-08 | 4.80E-07 | T | F | F | F |
| Ccnc | 1.80332653 | 4.60154145 | 38.0371594 | 5.48E-08 | 4.80E-07 | T | F | T | T |
| Casp8 | -3.3579775 | 3.36138446 | 37.9787685 | 5.58E-08 | 4.89E-07 | T | T | F | T |

| | | | | | | | | | |
|---|---|---|---|---|---|---|---|---|---|
| Hdx | -2.331338 | 4.40395239 | 37.9773666 | 5.58E-08 | 4.89E-07 | T | F | T | T |
| Dgki | -6.4860277 | 2.33235642 | 37.9743696 | 5.59E-08 | 4.89E-07 | T | F | T | T |
| Plekhb2 | 1.75297288 | 4.74211525 | 37.9587133 | 5.62E-08 | 4.91E-07 | T | F | T | T |
| Fzd8 | -2.2467993 | 4.43330159 | 37.9573845 | 5.62E-08 | 4.91E-07 | F | F | F | F |
| F2 | -6.3301351 | 2.19216738 | 37.9558787 | 5.62E-08 | 4.91E-07 | F | F | F | T |
| Gpc2 | -3.059256 | 3.59990999 | 37.947088 | 5.64E-08 | 4.93E-07 | T | F | F | F |
| Lrrc42 | -1.7245728 | 5.46997917 | 37.9340148 | 5.66E-08 | 4.95E-07 | F | F | T | F |
| mt-Atp6 | 1.02892689 | 9.55206649 | 37.8945798 | 5.73E-08 | 5.01E-07 | F | F | F | F |
| Xlr4b | 2.69203335 | 3.24885854 | 37.8895998 | 5.74E-08 | 5.01E-07 | T | F | F | T |
| Frmd5 | -5.6359456 | 2.42615218 | 37.8886766 | 5.74E-08 | 5.01E-07 | T | F | F | F |
| Tdrd7 | 1.8972051 | 4.33542731 | 37.8477001 | 5.82E-08 | 5.08E-07 | F | F | F | F |
| Ift172 | -2.1208306 | 4.60643255 | 37.8406785 | 5.83E-08 | 5.08E-07 | F | F | F | F |
| Cul7 | 1.09735509 | 8.3473868 | 37.8319743 | 5.85E-08 | 5.10E-07 | F | F | F | F |
| Zfp281 | -1.5624381 | 6.15305271 | 37.8231053 | 5.86E-08 | 5.11E-07 | T | F | T | T |
| Gm32687 | -9.452417 | 1.90699455 | 37.8034404 | 5.90E-08 | 5.14E-07 | F | F | T | F |
| Tdgf1 | -10.528977 | 2.94574429 | 37.7911498 | 5.93E-08 | 5.16E-07 | T | F | T | F |
| Abraxas1 | 3.759143 | 3.07097881 | 37.7483682 | 6.01E-08 | 5.23E-07 | T | F | F | F |
| Sufu | -1.3738354 | 6.84829021 | 37.7193627 | 6.06E-08 | 5.27E-07 | T | F | T | F |
| Avpr1b | -9.4414186 | 1.89708484 | 37.6848377 | 6.13E-08 | 5.33E-07 | T | F | F | T |
| Abhd8 | -2.4328427 | 4.15922974 | 37.6777576 | 6.14E-08 | 5.34E-07 | F | F | T | F |
| Prrx2 | -4.2979671 | 3.35436996 | 37.6275 | 6.24E-08 | 5.42E-07 | F | F | F | T |
| Cadps | -9.4195657 | 1.88555571 | 37.6190161 | 6.26E-08 | 5.44E-07 | T | F | T | F |
| Pim1 | 1.56424643 | 6.56892569 | 37.5941667 | 6.31E-08 | 5.48E-07 | F | F | T | F |
| Lzts3 | 1.88513953 | 4.36746016 | 37.5408093 | 6.42E-08 | 5.57E-07 | F | F | F | F |
| Oxr1 | 1.41093626 | 5.89625228 | 37.5296556 | 6.44E-08 | 5.59E-07 | T | T | T | T |
| Chd7 | 1.01618444 | 9.32091284 | 37.4576206 | 6.59E-08 | 5.72E-07 | T | F | F | T |
| Zfp580 | -6.3156176 | 2.17620701 | 37.4461465 | 6.62E-08 | 5.74E-07 | F | F | F | F |
| Psen1 | 1.23627816 | 6.85330559 | 37.4424113 | 6.63E-08 | 5.74E-07 | F | F | T | F |
| Elk4 | 1.53080954 | 5.39199392 | 37.4012546 | 6.72E-08 | 5.82E-07 | F | F | T | F |
| Fbh1 | -1.6768141 | 5.59198077 | 37.3973421 | 6.72E-08 | 5.82E-07 | F | F | F | F |
| Coro2b | -3.0547021 | 3.57451938 | 37.3925707 | 6.73E-08 | 5.83E-07 | F | F | F | F |
| Gpr146 | 3.85375664 | 2.07488928 | 37.3815713 | 6.76E-08 | 5.84E-07 | T | F | T | T |
| Lif | 3.58134282 | 2.31267532 | 37.3780962 | 6.77E-08 | 5.85E-07 | F | F | T | F |
| Lama3 | 3.90564443 | 2.04873856 | 37.3752773 | 6.77E-08 | 5.85E-07 | F | F | F | T |
| Eml3 | 1.28175043 | 6.58254776 | 37.3679846 | 6.79E-08 | 5.86E-07 | F | F | F | F |
| Lgals2 | -5.6619129 | 2.45775551 | 37.3607483 | 6.80E-08 | 5.87E-07 | F | F | F | F |
| Fbxo16 | -9.4003642 | 1.86410977 | 37.3227851 | 6.89E-08 | 5.94E-07 | F | F | T | F |
| Sp4 | -2.1455808 | 4.59492483 | 37.3200948 | 6.89E-08 | 5.95E-07 | T | T | T | T |
| Vamp2 | 1.71213459 | 4.77493532 | 37.3072984 | 6.92E-08 | 5.97E-07 | T | F | T | T |
| Efna2 | -2.5710743 | 4.01340328 | 37.2952775 | 6.95E-08 | 5.99E-07 | T | F | F | F |
| Eif4ebp1 | 1.26698872 | 6.73065835 | 37.2944489 | 6.95E-08 | 5.99E-07 | F | F | T | T |
| Gm14434 | 4.50445708 | 2.00890177 | 37.2693598 | 7.01E-08 | 6.04E-07 | F | F | T | T |

| | | | | | | | | | |
|---|---|---|---|---|---|---|---|---|---|
| Tlr5 | 5.0666138 | 1.56031507 | 37.2564441 | 7.04E-08 | 6.06E-07 | F | F | F | F |
| M1ap | -9.3979849 | 1.86287487 | 37.2490935 | 7.05E-08 | 6.07E-07 | F | F | F | F |
| Polk | 2.10540595 | 4.01956737 | 37.247159 | 7.06E-08 | 6.07E-07 | T | F | T | T |
| Resf1 | -1.4876389 | 6.35533483 | 37.2460818 | 7.06E-08 | 6.07E-07 | F | F | T | F |
| Acpp | 3.60381907 | 2.29051578 | 37.2339319 | 7.09E-08 | 6.09E-07 | T | F | T | F |
| Arid3c | -9.3974636 | 1.86267718 | 37.2257735 | 7.11E-08 | 6.11E-07 | F | F | T | F |
| Plscr1 | -2.1991806 | 4.42425531 | 37.2017402 | 7.16E-08 | 6.15E-07 | F | F | F | F |
| Tnk2 | 1.20534552 | 7.02515813 | 37.1921359 | 7.18E-08 | 6.17E-07 | F | F | T | F |
| Arhgef16 | 1.60652013 | 5.06188921 | 37.1773234 | 7.22E-08 | 6.20E-07 | T | T | F | T |
| Adrb3 | -9.6276246 | 2.06704267 | 37.1586952 | 7.26E-08 | 6.23E-07 | T | F | F | T |
| Kdelr2 | 1.17978226 | 7.31414626 | 37.1508162 | 7.28E-08 | 6.24E-07 | F | F | F | T |
| Cln3 | 2.10236333 | 4.02697971 | 37.1457298 | 7.29E-08 | 6.25E-07 | T | F | T | T |
| Dpysl3 | -2.1628114 | 4.47353664 | 37.1390169 | 7.31E-08 | 6.26E-07 | F | F | T | T |
| Zfp46 | 1.60830909 | 5.05801522 | 37.1232257 | 7.34E-08 | 6.29E-07 | T | F | T | F |
| Elf2 | 1.21082992 | 7.02490936 | 37.0918717 | 7.42E-08 | 6.35E-07 | F | F | T | F |
| Isl1 | -4.7015308 | 2.6888926 | 37.0816759 | 7.44E-08 | 6.37E-07 | T | F | F | F |
| Zfp467 | -3.7906462 | 3.13196854 | 37.0714047 | 7.47E-08 | 6.39E-07 | T | F | T | T |
| Dpy19l1 | 1.16940098 | 7.25525885 | 37.0272245 | 7.58E-08 | 6.48E-07 | T | F | T | T |
| Slc29a2 | 1.51208542 | 5.40727474 | 36.9705043 | 7.72E-08 | 6.60E-07 | F | F | T | F |
| Rpl35a | -1.4471052 | 6.50510456 | 36.9563347 | 7.75E-08 | 6.62E-07 | F | F | F | F |
| Sept4 | 4.16191927 | 1.83490958 | 36.8792671 | 7.95E-08 | 6.79E-07 | F | F | F | F |
| Slit1 | -3.5634285 | 3.41165181 | 36.824892 | 8.09E-08 | 6.91E-07 | F | F | F | T |
| Mpped2 | -3.3698386 | 3.29586738 | 36.8131371 | 8.12E-08 | 6.93E-07 | T | F | F | T |
| Crot | -1.8418364 | 5.12120706 | 36.7970223 | 8.16E-08 | 6.96E-07 | F | F | T | F |
| Tuba4a | -2.4378978 | 4.12541338 | 36.7739674 | 8.22E-08 | 7.01E-07 | F | F | T | F |
| Zfp523 | -2.9716555 | 3.6426784 | 36.7579057 | 8.27E-08 | 7.05E-07 | F | F | F | F |
| Zcchc24 | -2.8573289 | 3.7597588 | 36.7476014 | 8.29E-08 | 7.07E-07 | F | F | T | T |
| Prmt8 | -9.3796771 | 1.8417398 | 36.745129 | 8.30E-08 | 7.07E-07 | T | F | T | T |
| Lims1 | 1.22116735 | 6.91751569 | 36.7399288 | 8.31E-08 | 7.08E-07 | T | F | T | T |
| Fam13a | -5.5309695 | 2.32358982 | 36.7356921 | 8.32E-08 | 7.09E-07 | T | F | T | T |
| Gucy1b1 | -9.3769693 | 1.84345178 | 36.7322573 | 8.33E-08 | 7.09E-07 | F | F | F | F |
| Smpd3 | -9.3814399 | 1.84929748 | 36.721101 | 8.36E-08 | 7.12E-07 | T | F | T | F |
| Mdfic | -5.1091199 | 2.4625884 | 36.6911249 | 8.45E-08 | 7.18E-07 | T | F | T | F |
| Ube2ql1 | -9.4177558 | 1.87233234 | 36.6774483 | 8.48E-08 | 7.21E-07 | T | F | F | F |
| Scaper | 1.75483049 | 4.62055932 | 36.6694993 | 8.51E-08 | 7.23E-07 | T | F | F | F |
| Sh2d5 | 3.01313697 | 2.8234791 | 36.6404341 | 8.59E-08 | 7.29E-07 | F | F | F | F |
| Zmiz2 | -1.4254964 | 6.57624576 | 36.5845015 | 8.74E-08 | 7.42E-07 | T | F | T | T |
| Ptprm | -2.9233813 | 3.64670035 | 36.5791809 | 8.76E-08 | 7.43E-07 | F | F | F | F |
| Mrtfb | 1.25367869 | 6.72093185 | 36.5607007 | 8.81E-08 | 7.47E-07 | F | F | T | F |
| Gm35315 | -9.3789413 | 1.83922223 | 36.5533725 | 8.83E-08 | 7.49E-07 | F | F | T | F |
| Zdhhc17 | 1.85600596 | 4.35652423 | 36.5433877 | 8.86E-08 | 7.51E-07 | T | F | F | T |
| Stxbp4 | -1.6237123 | 5.7382799 | 36.4870636 | 9.02E-08 | 7.65E-07 | T | F | T | T |

| | | | | | | | | | |
|---|---|---|---|---|---|---|---|---|---|
| Tfdp2 | -1.5197744 | 6.17733148 | 36.4817766 | 9.04E-08 | 7.66E-07 | T | F | T | T |
| Sdsl | -9.3709856 | 1.83250218 | 36.4661607 | 9.09E-08 | 7.69E-07 | F | F | F | F |
| Piwil2 | 3.23460694 | 2.60276135 | 36.4501692 | 9.13E-08 | 7.73E-07 | T | T | F | T |
| Zfp703 | -1.7757001 | 6.48057641 | 36.4187135 | 9.23E-08 | 7.81E-07 | F | F | T | F |
| Sprtn | 1.44752703 | 5.60262913 | 36.3623416 | 9.40E-08 | 7.95E-07 | F | F | F | F |
| Car13 | -4.5629653 | 2.64038404 | 36.3324924 | 9.49E-08 | 8.02E-07 | T | T | F | T |
| Vstm2b | -6.2707015 | 2.13306183 | 36.321152 | 9.52E-08 | 8.05E-07 | T | F | T | F |
| Lad1 | 1.35998967 | 6.28436721 | 36.3095732 | 9.56E-08 | 8.08E-07 | T | F | T | T |
| Mfap4 | -9.3569442 | 1.82300855 | 36.3074098 | 9.57E-08 | 8.08E-07 | T | F | F | T |
| Tmem250-p | -1.807432 | 5.1527159 | 36.2848034 | 9.64E-08 | 8.13E-07 | F | F | T | F |
| Dipk1a | -4.7875794 | 2.54881308 | 36.2814168 | 9.65E-08 | 8.14E-07 | T | F | T | F |
| Sema4b | 1.31836294 | 6.27853641 | 36.2790783 | 9.66E-08 | 8.14E-07 | F | F | T | F |
| Hs1bp3 | 1.42874312 | 5.73851478 | 36.264359 | 9.70E-08 | 8.18E-07 | F | F | T | F |
| Tle5 | -1.8481724 | 5.02504968 | 36.2369564 | 9.79E-08 | 8.25E-07 | F | F | F | F |
| Slc23a1 | -2.2987992 | 4.2401076 | 36.2346624 | 9.80E-08 | 8.25E-07 | F | F | T | F |
| Map3k2 | 1.52474744 | 5.24262203 | 36.232759 | 9.80E-08 | 8.25E-07 | F | T | T | T |
| Hpn | -3.1317185 | 3.49887029 | 36.2288929 | 9.81E-08 | 8.26E-07 | F | F | F | F |
| Ncapd2 | -1.1657695 | 8.2644529 | 36.2277058 | 9.82E-08 | 8.26E-07 | F | F | F | F |
| Src | 1.20340262 | 6.92843472 | 36.2132857 | 9.86E-08 | 8.30E-07 | T | F | F | T |
| Marveld1 | -3.7184726 | 3.05475784 | 36.2048634 | 9.89E-08 | 8.32E-07 | T | F | F | F |
| Chp1 | 1.24512973 | 6.69841674 | 36.1862477 | 9.95E-08 | 8.36E-07 | T | F | F | F |
| Phlpp2 | 1.49927756 | 5.38110117 | 36.1783987 | 9.98E-08 | 8.38E-07 | F | F | T | F |
| Ide | -1.292904 | 7.07269733 | 36.1568811 | 1.00E-07 | 8.44E-07 | F | F | T | F |
| Daam1 | 1.24438755 | 6.72961853 | 36.1333938 | 1.01E-07 | 8.50E-07 | T | F | T | T |
| Zfp422 | -1.7685828 | 5.18845347 | 36.0960805 | 1.02E-07 | 8.60E-07 | T | F | T | T |
| Doxl1 | 3.87644125 | 2.05156529 | 36.0822556 | 1.03E-07 | 8.64E-07 | T | F | F | F |
| Zfp799 | -1.6789426 | 5.59891877 | 36.0687341 | 1.03E-07 | 8.67E-07 | T | F | T | T |
| Map2k1 | 1.18810721 | 6.99834899 | 36.0291059 | 1.05E-07 | 8.78E-07 | T | T | T | T |
| Myo1b | 1.15436098 | 7.23258061 | 36.0022498 | 1.06E-07 | 8.85E-07 | T | F | T | T |
| Myh7b | 3.18964203 | 2.7304925 | 35.985846 | 1.06E-07 | 8.90E-07 | F | F | F | F |
| Faf1 | 1.14402798 | 7.32964229 | 35.9852307 | 1.06E-07 | 8.90E-07 | T | F | T | T |
| Tfr2 | 3.28098715 | 2.53369174 | 35.9644199 | 1.07E-07 | 8.95E-07 | F | F | T | T |
| Ppm1h | -2.4410194 | 4.08127512 | 35.9476378 | 1.08E-07 | 9.00E-07 | T | F | T | F |
| Zfp936 | 3.48786765 | 2.34008732 | 35.9453937 | 1.08E-07 | 9.00E-07 | T | F | T | T |
| Fam219b | 1.62474297 | 4.95060265 | 35.9271898 | 1.08E-07 | 9.05E-07 | F | F | T | F |
| Gjb2 | 1.86001672 | 4.35049545 | 35.8988504 | 1.09E-07 | 9.13E-07 | T | F | T | F |
| Fyco1 | 1.49431208 | 5.33984412 | 35.8581828 | 1.11E-07 | 9.25E-07 | T | F | T | T |
| Ncor2 | -1.1560369 | 8.29424674 | 35.8345343 | 1.12E-07 | 9.32E-07 | F | F | F | F |
| Mycn | -1.265307 | 7.21938455 | 35.7537532 | 1.15E-07 | 9.57E-07 | T | F | F | F |
| Trim44 | -1.2405425 | 7.42315946 | 35.6966196 | 1.17E-07 | 9.74E-07 | T | T | T | T |
| Kctd12b | -6.1897772 | 2.12604059 | 35.6749698 | 1.18E-07 | 9.81E-07 | T | F | T | T |
| 2810408A1 | -9.3635826 | 1.82119272 | 35.6574003 | 1.18E-07 | 9.86E-07 | F | F | F | F |

| | | | | | | | | | |
|---|---|---|---|---|---|---|---|---|---|
| Mtmr11 | -9.3286466 | 1.7973016 | 35.6264241 | 1.19E-07 | 9.96E-07 | F | F | T | F |
| Clock | 1.4508671 | 5.52122221 | 35.6117496 | 1.20E-07 | 1.00E-06 | T | F | T | T |
| Osr1 | -9.3628591 | 1.83602271 | 35.5814602 | 1.21E-07 | 1.01E-06 | F | F | T | F |
| Pnrc1 | -1.5799899 | 5.97313806 | 35.5778121 | 1.21E-07 | 1.01E-06 | T | F | F | T |
| Dscam | -5.0580397 | 2.41547623 | 35.5700508 | 1.22E-07 | 1.01E-06 | T | F | T | T |
| Bmp5 | -3.1560958 | 3.39852712 | 35.5680399 | 1.22E-07 | 1.01E-06 | F | F | T | F |
| Aldh9a1 | 1.15550045 | 7.11392669 | 35.5658323 | 1.22E-07 | 1.01E-06 | F | F | T | F |
| Nfib | -4.8697158 | 2.50453043 | 35.5612529 | 1.22E-07 | 1.01E-06 | T | F | T | T |
| Mab21l3 | 4.14100766 | 1.80307923 | 35.5415051 | 1.23E-07 | 1.02E-06 | F | F | T | F |
| Hpcal1 | 1.67673371 | 4.77346956 | 35.5248628 | 1.24E-07 | 1.03E-06 | T | F | F | F |
| Stx7 | -1.7236352 | 5.2819396 | 35.5237395 | 1.24E-07 | 1.03E-06 | T | F | F | F |
| Mta3 | 1.32943549 | 6.13615455 | 35.4923193 | 1.25E-07 | 1.04E-06 | T | F | F | F |
| Prdx6 | -1.5626485 | 5.82861837 | 35.4913693 | 1.25E-07 | 1.04E-06 | T | F | T | T |
| Pir | -5.054979 | 2.41207183 | 35.4700762 | 1.26E-07 | 1.04E-06 | F | F | F | T |
| Tnrc6b | -1.3458512 | 7.05441529 | 35.4115378 | 1.28E-07 | 1.06E-06 | T | F | T | T |
| Arhgap18 | 1.69327515 | 4.72813372 | 35.338279 | 1.31E-07 | 1.09E-06 | T | F | F | T |
| Pipox | -9.3302918 | 1.79180655 | 35.3352159 | 1.31E-07 | 1.09E-06 | T | F | F | F |
| Cpne3 | 1.34787589 | 6.1407052 | 35.3343829 | 1.32E-07 | 1.09E-06 | T | F | T | T |
| Sycp2 | 5.96468718 | 0.9987173 | 35.3293327 | 1.32E-07 | 1.09E-06 | T | F | F | F |
| Pdzd3 | -6.284759 | 2.15633225 | 35.3259355 | 1.32E-07 | 1.09E-06 | F | F | F | F |
| Gm8113 | -2.0089336 | 4.65211178 | 35.3134287 | 1.32E-07 | 1.10E-06 | F | F | F | F |
| Dpf1 | -4.0684051 | 2.81986845 | 35.3113451 | 1.33E-07 | 1.10E-06 | T | F | F | F |
| Ubxn7 | -1.2684837 | 7.34432773 | 35.2690937 | 1.34E-07 | 1.11E-06 | T | F | T | T |
| Ptger3 | 4.90760947 | 1.36698617 | 35.2671682 | 1.34E-07 | 1.11E-06 | T | T | F | T |
| Meis3 | -3.8917684 | 3.00895032 | 35.2667752 | 1.34E-07 | 1.11E-06 | T | F | F | F |
| Klhl28 | 1.65307897 | 4.85353168 | 35.245874 | 1.35E-07 | 1.12E-06 | T | F | T | T |
| Dnajc3 | 1.13315401 | 7.31979168 | 35.2345431 | 1.36E-07 | 1.12E-06 | F | F | T | F |
| Mif4gd | 1.85043947 | 4.3011509 | 35.2302438 | 1.36E-07 | 1.12E-06 | F | F | F | F |
| Maneal | -9.3094558 | 1.77731306 | 35.2246675 | 1.36E-07 | 1.12E-06 | F | F | T | F |
| Ogdhl | -1.6361251 | 5.81549153 | 35.2085632 | 1.37E-07 | 1.13E-06 | F | F | F | F |
| Tsc22d3 | -3.0640437 | 3.45701061 | 35.2056342 | 1.37E-07 | 1.13E-06 | T | F | T | T |
| Prex1 | -1.3634747 | 6.73954807 | 35.1900955 | 1.38E-07 | 1.14E-06 | T | F | F | T |
| Cracr2b | 2.78617118 | 3.03059772 | 35.1799156 | 1.38E-07 | 1.14E-06 | F | F | F | F |
| Sdc2 | -2.9954545 | 3.51886565 | 35.1667155 | 1.39E-07 | 1.14E-06 | T | F | T | T |
| Ghitm | 1.1904424 | 6.88103282 | 35.1432591 | 1.40E-07 | 1.15E-06 | T | T | F | T |
| Gnai1 | -2.9665844 | 3.54424149 | 35.1244083 | 1.41E-07 | 1.16E-06 | T | F | T | T |
| Ehd2 | 1.90504367 | 4.22004542 | 35.099055 | 1.42E-07 | 1.17E-06 | T | F | T | F |
| Gnpda1 | -1.6352893 | 5.56434564 | 35.0969555 | 1.42E-07 | 1.17E-06 | F | F | T | F |
| Tnfaip8 | 1.71080127 | 4.63908018 | 34.9704655 | 1.48E-07 | 1.22E-06 | F | F | T | F |
| Syne2 | -1.2727802 | 7.14722052 | 34.9557228 | 1.49E-07 | 1.22E-06 | T | F | F | F |
| Slc6a17 | -9.2983875 | 1.76840261 | 34.9476363 | 1.49E-07 | 1.23E-06 | T | F | T | F |
| Vsig10 | -9.3607491 | 1.81574915 | 34.9225492 | 1.51E-07 | 1.24E-06 | T | F | T | T |

| | | | | | | | | | |
|---|---|---|---|---|---|---|---|---|---|
| Tnfaip3 | -9.3060858 | 1.76962362 | 34.9072491 | 1.51E-07 | 1.24E-06 | F | F | T | F |
| Mxd4 | 1.58241369 | 5.0270191 | 34.9054588 | 1.51E-07 | 1.24E-06 | F | F | T | F |
| Irak4 | 2.71006629 | 3.08494314 | 34.9004329 | 1.52E-07 | 1.24E-06 | T | F | F | F |
| Col27a1 | -3.0541744 | 3.47738185 | 34.8916371 | 1.52E-07 | 1.25E-06 | T | F | F | T |
| Arfip2 | -2.3668768 | 4.13095768 | 34.8870996 | 1.52E-07 | 1.25E-06 | F | F | T | F |
| Trpc2 | 9.14695492 | 0.56759076 | 35.0089154 | 1.53E-07 | 1.25E-06 | T | F | F | F |
| Ptprg | -1.6291674 | 5.53958134 | 34.8632747 | 1.54E-07 | 1.26E-06 | T | T | T | T |
| Samhd1 | -1.2892943 | 6.97558473 | 34.8630538 | 1.54E-07 | 1.26E-06 | T | F | F | F |
| Zfp169 | -3.7552947 | 2.96716549 | 34.8557959 | 1.54E-07 | 1.26E-06 | T | F | T | T |
| Smg5 | 1.08499122 | 7.81598501 | 34.8339278 | 1.55E-07 | 1.27E-06 | F | F | T | F |
| Rhpn1 | -9.29175 | 1.7612134 | 34.8215926 | 1.56E-07 | 1.27E-06 | F | F | T | T |
| Shank2 | -2.5281045 | 3.95215498 | 34.8062493 | 1.57E-07 | 1.28E-06 | F | F | T | T |
| Susd5 | -9.2933566 | 1.75955733 | 34.791376 | 1.57E-07 | 1.28E-06 | T | F | F | T |
| Pnma5 | 2.18335428 | 3.82903388 | 34.7702656 | 1.58E-07 | 1.29E-06 | F | F | T | F |
| Map4k4 | 0.99873227 | 8.99454168 | 34.7701485 | 1.58E-07 | 1.29E-06 | T | F | F | T |
| Agap1 | -1.3107457 | 6.92746243 | 34.7632486 | 1.59E-07 | 1.29E-06 | T | T | T | T |
| Txnrd2 | 1.82474054 | 4.45403157 | 34.7631571 | 1.59E-07 | 1.29E-06 | F | F | F | F |
| Klc1 | 1.27628863 | 6.36157888 | 34.7578967 | 1.59E-07 | 1.30E-06 | F | F | T | F |
| Klf4 | 2.04807374 | 3.98212874 | 34.7419335 | 1.60E-07 | 1.30E-06 | T | F | T | F |
| Unc13b | -2.2758448 | 4.23771642 | 34.7383681 | 1.60E-07 | 1.30E-06 | T | F | T | T |
| Dnah11 | -3.1183461 | 3.40584383 | 34.7375571 | 1.60E-07 | 1.30E-06 | F | F | F | F |
| Bmpr1b | -9.2950702 | 1.75979855 | 34.7288415 | 1.61E-07 | 1.31E-06 | T | F | T | T |
| Abcb1b | -9.2950745 | 1.7676799 | 34.714956 | 1.61E-07 | 1.31E-06 | T | F | F | F |
| Oaz2 | -1.7291788 | 5.19705869 | 34.6836435 | 1.63E-07 | 1.33E-06 | T | F | T | F |
| Cnrip1 | -9.3189256 | 1.79418649 | 34.6557685 | 1.65E-07 | 1.34E-06 | F | F | F | T |
| Fam171b | -3.5976053 | 3.05202905 | 34.6490448 | 1.65E-07 | 1.34E-06 | T | F | T | T |
| Ranbp17 | 1.63073547 | 4.84571982 | 34.6302292 | 1.66E-07 | 1.35E-06 | T | F | T | T |
| Pcnp | -1.4566752 | 6.25157295 | 34.620539 | 1.66E-07 | 1.35E-06 | T | F | T | F |
| Usp49 | 1.50561212 | 5.25332553 | 34.5994773 | 1.68E-07 | 1.36E-06 | T | F | T | T |
| Tmem98 | -2.0803877 | 4.46239513 | 34.5874193 | 1.68E-07 | 1.37E-06 | F | F | F | F |
| Col24a1 | -9.2802895 | 1.74756419 | 34.5238557 | 1.72E-07 | 1.39E-06 | T | F | T | F |
| Syde1 | 1.42545446 | 5.53413098 | 34.5118477 | 1.73E-07 | 1.40E-06 | T | F | T | T |
| Emc8 | -1.4623394 | 6.15372238 | 34.4782107 | 1.75E-07 | 1.41E-06 | T | F | T | T |
| Scd2 | -1.0841711 | 9.07032802 | 34.4723194 | 1.75E-07 | 1.42E-06 | T | F | T | T |
| Tmem151b | -3.5544306 | 3.07089843 | 34.470237 | 1.75E-07 | 1.42E-06 | F | F | F | F |
| Ep300 | -1.1516496 | 8.1467132 | 34.4636525 | 1.75E-07 | 1.42E-06 | T | F | F | F |
| Slc6a9 | 2.54284754 | 3.31322766 | 34.461209 | 1.76E-07 | 1.42E-06 | F | F | T | F |
| Alms1 | -1.3287271 | 6.78117245 | 34.4416355 | 1.77E-07 | 1.43E-06 | F | F | F | F |
| Bbs7 | -3.5937538 | 3.04937449 | 34.4372357 | 1.77E-07 | 1.43E-06 | T | F | F | T |
| Irf2bp1 | -1.551457 | 5.76787365 | 34.4155875 | 1.78E-07 | 1.44E-06 | F | F | F | F |
| Mob1a | 1.14429813 | 7.07413834 | 34.3856122 | 1.80E-07 | 1.45E-06 | T | F | T | T |
| Begain | -9.2736088 | 1.74106742 | 34.3696018 | 1.81E-07 | 1.46E-06 | T | F | T | T |

| | | | | | | | | | |
|---|---|---|---|---|---|---|---|---|---|
| Frs3 | -9.2766134 | 1.74941372 | 34.3692541 | 1.81E-07 | 1.46E-06 | T | F | F | F |
| Pou2f1 | -1.2611638 | 7.20001992 | 34.3477134 | 1.82E-07 | 1.47E-06 | F | T | T | T |
| AI837181 | -1.8964916 | 4.79847752 | 34.3425505 | 1.83E-07 | 1.47E-06 | F | F | F | F |
| Hook3 | 1.25872047 | 6.45826594 | 34.3263625 | 1.84E-07 | 1.48E-06 | F | F | T | T |
| Faah | 2.72200258 | 3.13662285 | 34.3192696 | 1.84E-07 | 1.48E-06 | F | F | F | F |
| Magt1 | 1.21511285 | 6.66488337 | 34.3109993 | 1.84E-07 | 1.49E-06 | F | F | T | F |
| Xndc1 | -2.141677 | 4.36021664 | 34.302288 | 1.85E-07 | 1.49E-06 | F | F | T | T |
| Dmrta1 | -9.2707219 | 1.73744309 | 34.2561608 | 1.88E-07 | 1.51E-06 | T | T | F | T |
| Rab10 | 1.09889578 | 7.53359509 | 34.2525723 | 1.88E-07 | 1.51E-06 | T | F | T | T |
| Rsu1 | 1.52493145 | 5.13263703 | 34.2382234 | 1.89E-07 | 1.52E-06 | T | F | F | F |
| P3h4 | -3.929598 | 2.85326514 | 34.2169038 | 1.90E-07 | 1.53E-06 | F | F | F | F |
| Zfp57 | -2.215489 | 4.2409441 | 34.1591332 | 1.94E-07 | 1.56E-06 | F | F | F | F |
| Zfp704 | -1.3815063 | 6.48511433 | 34.1518143 | 1.95E-07 | 1.56E-06 | T | F | T | T |
| D16Ertd472 | -2.0786009 | 4.43704204 | 34.1442286 | 1.95E-07 | 1.57E-06 | T | F | T | T |
| Zdhhc23 | 2.30194858 | 3.56487562 | 34.1273384 | 1.96E-07 | 1.58E-06 | F | F | T | F |
| Pttg1ip | 1.30891032 | 6.16620178 | 34.1154196 | 1.97E-07 | 1.58E-06 | T | F | T | T |
| Ahnak | 1.12902875 | 8.97516432 | 34.0863758 | 1.99E-07 | 1.60E-06 | T | F | T | T |
| Taf1 | 1.13567613 | 7.16318481 | 34.0725973 | 2.00E-07 | 1.60E-06 | T | F | T | T |
| Nova2 | -3.7299258 | 2.9454937 | 34.0619784 | 2.00E-07 | 1.61E-06 | F | F | T | F |
| Lpp | 1.10740583 | 7.41429694 | 34.0426146 | 2.02E-07 | 1.62E-06 | T | F | T | T |
| St8sia2 | -4.037327 | 2.78818063 | 34.0281422 | 2.03E-07 | 1.62E-06 | T | F | F | T |
| Tbl1xr1 | -1.42418 | 6.40236384 | 34.0268447 | 2.03E-07 | 1.62E-06 | T | T | T | T |
| Sco1 | 1.39449199 | 5.64581749 | 33.9545955 | 2.08E-07 | 1.66E-06 | T | T | T | T |
| Pygo1 | -9.2726717 | 1.74881934 | 33.9304518 | 2.09E-07 | 1.68E-06 | F | F | T | T |
| Cux2 | -4.5151832 | 2.59034138 | 33.9116689 | 2.11E-07 | 1.69E-06 | F | F | F | T |
| B3gnt8 | 5.71478426 | 1.00822155 | 33.8925056 | 2.12E-07 | 1.70E-06 | T | F | F | F |
| Hmgcr | 1.109846 | 7.32102162 | 33.8436236 | 2.16E-07 | 1.72E-06 | T | F | T | T |
| Aff3 | -1.8527652 | 4.85255098 | 33.822835 | 2.17E-07 | 1.74E-06 | F | F | F | T |
| Ckmt1 | 3.34522988 | 2.35636175 | 33.7834238 | 2.20E-07 | 1.76E-06 | F | F | F | F |
| Marf1 | 1.18690365 | 6.86823535 | 33.7647495 | 2.21E-07 | 1.77E-06 | F | F | F | T |
| Vps37b | 1.34738034 | 5.86559015 | 33.6967326 | 2.26E-07 | 1.81E-06 | T | F | T | F |
| Tmem33 | 1.33010675 | 5.93769912 | 33.6663328 | 2.29E-07 | 1.83E-06 | T | F | T | F |
| Man1a2 | 1.11753494 | 7.19136299 | 33.6264906 | 2.32E-07 | 1.85E-06 | T | T | T | T |
| Xbp1 | 1.25424643 | 6.61050362 | 33.6212125 | 2.32E-07 | 1.85E-06 | T | F | F | T |
| Cfap52 | -9.240941 | 1.70929475 | 33.6180645 | 2.33E-07 | 1.85E-06 | F | F | F | F |
| Tulp4 | -1.3921423 | 6.38993337 | 33.6151514 | 2.33E-07 | 1.85E-06 | T | F | T | T |
| Echs1 | -1.614638 | 5.5139388 | 33.6090078 | 2.33E-07 | 1.86E-06 | T | F | F | F |
| Clcn5 | -1.1772068 | 7.65187222 | 33.5981836 | 2.34E-07 | 1.86E-06 | F | F | T | T |
| Ccdc80 | -5.0234722 | 2.3879822 | 33.5879048 | 2.35E-07 | 1.87E-06 | T | F | F | T |
| Alcam | -2.7306922 | 3.68437824 | 33.5735304 | 2.36E-07 | 1.88E-06 | F | F | F | F |
| Col26a1 | -4.4323598 | 2.77296168 | 33.5667867 | 2.37E-07 | 1.88E-06 | F | F | F | F |
| Prox1 | -2.6153313 | 3.82901144 | 33.5624677 | 2.37E-07 | 1.88E-06 | T | F | T | T |

| | | | | | | | | | |
|---|---|---|---|---|---|---|---|---|---|
| Csf1r | 3.63317804 | 2.0751483 | 33.5577424 | 2.37E-07 | 1.89E-06 | T | F | F | F |
| Mrtfa | 1.29754713 | 6.13636374 | 33.5429142 | 2.38E-07 | 1.89E-06 | F | F | T | F |
| Fhdc1 | 1.41243662 | 5.81402758 | 33.5350135 | 2.39E-07 | 1.90E-06 | F | F | T | F |
| Gria3 | -9.2360682 | 1.70499387 | 33.5307733 | 2.39E-07 | 1.90E-06 | T | F | F | T |
| Sgsm2 | -2.8720481 | 3.54549702 | 33.527802 | 2.40E-07 | 1.90E-06 | T | F | T | F |
| Phlda3 | -2.3662675 | 4.25562329 | 33.5106367 | 2.41E-07 | 1.91E-06 | F | F | F | T |
| Phlda2 | 1.04499844 | 8.08707808 | 33.4773662 | 2.44E-07 | 1.93E-06 | T | F | F | F |
| Kcnmb4 | -9.2315541 | 1.70166103 | 33.4730475 | 2.44E-07 | 1.94E-06 | T | F | T | F |
| Phlda1 | -1.9784863 | 4.72110083 | 33.4614351 | 2.45E-07 | 1.94E-06 | T | F | T | T |
| Tbkbp1 | -2.2405429 | 4.18406777 | 33.4430365 | 2.47E-07 | 1.95E-06 | F | F | T | F |
| R3hcc1 | -1.8497963 | 4.82944293 | 33.4405569 | 2.47E-07 | 1.95E-06 | F | F | F | F |
| B3gat3 | 1.41644744 | 5.5191276 | 33.4375284 | 2.47E-07 | 1.96E-06 | F | F | T | F |
| Clic5 | -2.4422115 | 3.97894016 | 33.3571841 | 2.54E-07 | 2.01E-06 | T | F | T | T |
| Fam89b | -2.5504462 | 3.92700874 | 33.35432 | 2.54E-07 | 2.01E-06 | F | F | F | F |
| Rps6ka6 | -1.2632158 | 6.96317855 | 33.3485299 | 2.55E-07 | 2.01E-06 | T | F | F | T |
| Vash1 | -2.8228495 | 3.64464848 | 33.3374884 | 2.55E-07 | 2.02E-06 | T | F | T | T |
| Prl7d1 | 7.07607591 | 0.7176704 | 33.3174639 | 2.57E-07 | 2.03E-06 | F | F | F | F |
| Uhmk1 | 1.13803091 | 7.00728745 | 33.307569 | 2.58E-07 | 2.04E-06 | T | F | T | T |
| Capn5 | 1.86439467 | 4.20872561 | 33.267237 | 2.62E-07 | 2.07E-06 | F | F | T | F |
| Phf19 | 3.26468245 | 2.41966539 | 33.261324 | 2.62E-07 | 2.07E-06 | T | F | T | F |
| Psat1 | 1.04300993 | 8.01399213 | 33.2570329 | 2.62E-07 | 2.07E-06 | T | F | T | T |
| Ppp1r14b | -1.2127664 | 7.61570467 | 33.2553571 | 2.63E-07 | 2.07E-06 | F | F | F | F |
| Lancl3 | -9.2693846 | 1.72915303 | 33.2383505 | 2.64E-07 | 2.08E-06 | T | F | T | T |
| Frmpd1 | 2.2493626 | 3.6438717 | 33.2369451 | 2.64E-07 | 2.08E-06 | F | F | F | F |
| Mdh1 | -1.3264238 | 6.72551832 | 33.2099909 | 2.67E-07 | 2.10E-06 | T | T | T | T |
| B3glct | 1.43473346 | 5.35779049 | 33.1890428 | 2.69E-07 | 2.12E-06 | T | F | F | T |
| Chmp4b | 1.14263152 | 7.01110517 | 33.1839725 | 2.69E-07 | 2.12E-06 | F | F | F | T |
| Irf3 | -2.7533491 | 3.64147606 | 33.1690474 | 2.70E-07 | 2.13E-06 | F | F | F | F |
| Dennd3 | -5.3388623 | 2.22064347 | 33.163822 | 2.71E-07 | 2.13E-06 | T | F | F | F |
| Ccp110 | -1.497326 | 5.91819595 | 33.1600235 | 2.71E-07 | 2.13E-06 | T | F | T | F |
| Plag1 | -2.6372049 | 3.80137701 | 33.1443825 | 2.73E-07 | 2.14E-06 | T | F | F | T |
| Lactb | 1.74337409 | 4.46419069 | 33.1041007 | 2.76E-07 | 2.17E-06 | F | F | F | F |
| Iglon5 | -3.5454804 | 3.01962308 | 33.0706995 | 2.79E-07 | 2.20E-06 | F | F | T | F |
| Arhgef9 | -6.128592 | 2.00043642 | 33.0640253 | 2.80E-07 | 2.20E-06 | T | F | T | T |
| Sephs2 | -1.284273 | 6.82425544 | 33.0570103 | 2.81E-07 | 2.20E-06 | T | T | T | T |
| Syde2 | 1.28383438 | 6.13955505 | 33.020861 | 2.84E-07 | 2.23E-06 | F | T | F | T |
| Capzb | 1.07021005 | 7.73701796 | 32.9826632 | 2.88E-07 | 2.26E-06 | F | F | T | F |
| Ubxn10 | -6.1386244 | 2.00510593 | 32.9779006 | 2.88E-07 | 2.26E-06 | F | F | T | F |
| Irak1 | -1.5481677 | 5.69881397 | 32.9463449 | 2.91E-07 | 2.28E-06 | T | F | T | T |
| Bcar1 | -1.3163166 | 6.66603978 | 32.936278 | 2.92E-07 | 2.29E-06 | T | F | F | F |
| Ubash3b | 1.88265361 | 4.15629329 | 32.9320098 | 2.93E-07 | 2.29E-06 | T | F | T | F |
| Grsf1 | -1.3299982 | 7.94207384 | 32.9318461 | 2.93E-07 | 2.29E-06 | T | F | T | T |

| | | | | | | | | | |
|---|---|---|---|---|---|---|---|---|---|
| Gm9112 | 9.07180312 | 0.49596715 | 33.0516902 | 2.93E-07 | 2.30E-06 | F | F | F | F |
| Acnat1 | -9.2042736 | 1.67800231 | 32.9086577 | 2.95E-07 | 2.31E-06 | F | F | T | T |
| Trpa1 | -9.2468949 | 1.70819497 | 32.8860712 | 2.97E-07 | 2.33E-06 | F | F | T | F |
| BC034090 | -2.6915122 | 3.68560177 | 32.8858552 | 2.97E-07 | 2.33E-06 | T | F | T | F |
| Dek | -1.2001302 | 7.31827857 | 32.8661543 | 2.99E-07 | 2.34E-06 | T | F | F | T |
| Cadps2 | -5.0024106 | 2.3588895 | 32.8527626 | 3.01E-07 | 2.35E-06 | T | F | F | F |
| Cfap44 | -9.2017022 | 1.67333541 | 32.8321592 | 3.03E-07 | 2.37E-06 | F | F | F | F |
| Synm | -4.0134493 | 2.77036146 | 32.7607946 | 3.10E-07 | 2.42E-06 | T | F | T | F |
| Actn2 | -9.2023403 | 1.6720506 | 32.7570262 | 3.11E-07 | 2.43E-06 | T | F | F | F |
| Fabp7 | -9.3322633 | 1.81330872 | 32.7558633 | 3.11E-07 | 2.43E-06 | F | F | F | F |
| S1pr3 | -6.3200493 | 2.1949807 | 32.7514168 | 3.11E-07 | 2.43E-06 | T | F | T | T |
| E2f6 | -1.8340039 | 4.8545952 | 32.7126864 | 3.15E-07 | 2.46E-06 | T | F | T | T |
| Ttc7 | 1.47546507 | 5.35625711 | 32.6863132 | 3.18E-07 | 2.48E-06 | F | F | F | T |
| Thpo | -9.2076738 | 1.67491504 | 32.6837926 | 3.19E-07 | 2.48E-06 | F | F | T | F |
| Lmna | 1.66068599 | 4.71273303 | 32.6649903 | 3.21E-07 | 2.50E-06 | F | F | F | F |
| Igf1r | -1.1753666 | 7.56794932 | 32.6526413 | 3.22E-07 | 2.51E-06 | T | F | T | T |
| Jdp2 | 2.40441639 | 3.50269818 | 32.6492665 | 3.22E-07 | 2.51E-06 | F | F | F | T |
| Zfp948 | 1.7126345 | 4.47872825 | 32.6309173 | 3.24E-07 | 2.52E-06 | F | F | F | F |
| Ism1 | -9.250278 | 1.73172359 | 32.625198 | 3.25E-07 | 2.53E-06 | F | F | F | F |
| 1110008P14 | 1.81031816 | 4.25408519 | 32.6217108 | 3.25E-07 | 2.53E-06 | F | F | F | F |
| Ctdsp1 | -1.3395755 | 6.5224478 | 32.6199808 | 3.25E-07 | 2.53E-06 | F | F | T | F |
| Rbbp7 | 0.96582219 | 9.04848286 | 32.6144988 | 3.26E-07 | 2.53E-06 | F | F | F | F |
| Tsku | -2.0923091 | 4.31208717 | 32.6114573 | 3.26E-07 | 2.54E-06 | F | F | F | T |
| Fkbp8 | -1.2452445 | 6.99257225 | 32.6078513 | 3.27E-07 | 2.54E-06 | F | F | T | F |
| Zfp821 | -2.3203033 | 4.08261416 | 32.5900223 | 3.29E-07 | 2.55E-06 | F | F | T | F |
| Olfr839-ps1 | 1.29255994 | 6.76903246 | 32.5852431 | 3.29E-07 | 2.55E-06 | F | F | F | F |
| Rwdd4a | -1.7814115 | 4.93385914 | 32.560324 | 3.32E-07 | 2.58E-06 | T | F | T | T |
| Zfp28 | 2.44085249 | 3.29772053 | 32.5506759 | 3.33E-07 | 2.58E-06 | T | F | T | F |
| Cntn2 | -3.2315382 | 3.20490193 | 32.5373516 | 3.35E-07 | 2.59E-06 | T | F | T | F |
| Gba2 | 1.3150863 | 5.90422832 | 32.534488 | 3.35E-07 | 2.60E-06 | F | F | F | F |
| Igfbp3 | -2.142239 | 4.39061802 | 32.5183327 | 3.37E-07 | 2.61E-06 | T | F | T | T |
| Wbp1l | 1.09191614 | 7.27576391 | 32.507008 | 3.38E-07 | 2.62E-06 | T | F | T | T |
| Lrch1 | 1.8540173 | 4.19044935 | 32.4667721 | 3.43E-07 | 2.65E-06 | T | F | T | F |
| Wrnip1 | -1.8886278 | 4.68945514 | 32.4219997 | 3.48E-07 | 2.69E-06 | T | F | F | F |
| Cpa2 | -4.6466099 | 2.41244289 | 32.407323 | 3.50E-07 | 2.71E-06 | F | F | F | F |
| Cbx5 | -1.0126315 | 9.64632355 | 32.4039329 | 3.50E-07 | 2.71E-06 | T | T | T | T |
| Efs | 1.31471206 | 5.99093423 | 32.3994441 | 3.51E-07 | 2.71E-06 | F | F | T | F |
| Tcf7l1 | -1.314013 | 7.01909846 | 32.3615857 | 3.55E-07 | 2.74E-06 | F | F | T | F |
| Fam122b | 1.73797228 | 4.38792049 | 32.3331032 | 3.59E-07 | 2.77E-06 | T | F | T | F |
| Wdr90 | -1.4831659 | 5.83077633 | 32.3287531 | 3.59E-07 | 2.77E-06 | F | F | F | F |
| Jph2 | -9.1807162 | 1.65847322 | 32.3074555 | 3.62E-07 | 2.79E-06 | F | F | T | T |
| Slc2a1 | 0.96223018 | 10.1909111 | 32.2845206 | 3.65E-07 | 2.81E-06 | F | F | F | F |

| | | | | | | | | | |
|---|---|---|---|---|---|---|---|---|---|
| Mpc1 | 2.90931015 | 2.73074017 | 32.2769283 | 3.66E-07 | 2.82E-06 | F | F | T | F |
| Cuta | 1.55719942 | 4.87730895 | 32.2727537 | 3.66E-07 | 2.82E-06 | F | F | F | F |
| Tmc5 | 3.20788227 | 2.40378242 | 32.2725242 | 3.66E-07 | 2.82E-06 | F | F | T | F |
| Ak3 | 1.27652213 | 6.13546118 | 32.2713486 | 3.66E-07 | 2.82E-06 | T | F | T | T |
| Tuba1c | 0.9726991 | 8.87889626 | 32.2699156 | 3.67E-07 | 2.82E-06 | F | F | T | F |
| Mfge8 | 1.36555864 | 5.80265755 | 32.2525177 | 3.69E-07 | 2.84E-06 | F | F | T | T |
| Rreb1 | 1.0893437 | 7.24472442 | 32.236492 | 3.71E-07 | 2.85E-06 | T | F | T | F |
| Lxn | 1.70313329 | 4.61810904 | 32.2142367 | 3.74E-07 | 2.87E-06 | F | F | F | F |
| Wdr31 | -4.25498 | 2.59944249 | 32.2044318 | 3.75E-07 | 2.88E-06 | F | F | T | F |
| Galk1 | 1.08858598 | 7.27654996 | 32.2039776 | 3.75E-07 | 2.88E-06 | F | F | F | F |
| Tmtc3 | 1.23768648 | 6.3093086 | 32.1991413 | 3.75E-07 | 2.89E-06 | T | T | F | T |
| Nfya | 1.16967243 | 6.74116944 | 32.1710412 | 3.79E-07 | 2.91E-06 | T | F | T | T |
| Rnf19a | 1.35493024 | 5.64459539 | 32.169489 | 3.79E-07 | 2.91E-06 | F | F | F | F |
| Slitrk2 | -9.1742572 | 1.64522249 | 32.1562776 | 3.81E-07 | 2.92E-06 | F | T | F | T |
| Arfip1 | 1.33638208 | 5.74115491 | 32.1454596 | 3.82E-07 | 2.93E-06 | T | F | T | F |
| Dnah9 | -9.1674045 | 1.64171579 | 32.1255364 | 3.85E-07 | 2.95E-06 | F | F | F | F |
| Nomo1 | 1.03505082 | 7.93888308 | 32.1140646 | 3.87E-07 | 2.96E-06 | F | F | T | F |
| Hspa1b | 2.41253114 | 3.29763463 | 32.1005284 | 3.88E-07 | 2.98E-06 | T | F | F | T |
| 4931406C0 | 2.01105454 | 3.91543592 | 32.0899778 | 3.90E-07 | 2.99E-06 | T | F | T | F |
| Zc3hav1l | -1.4353251 | 6.19194377 | 32.0818346 | 3.91E-07 | 2.99E-06 | F | F | T | T |
| Myo5b | -1.965534 | 4.51222223 | 32.0696456 | 3.92E-07 | 3.00E-06 | T | F | F | F |
| Nisch | -1.0205974 | 9.23662506 | 32.0505131 | 3.95E-07 | 3.02E-06 | T | F | T | T |
| Eva1a | 4.70839843 | 1.29505217 | 32.046423 | 3.96E-07 | 3.03E-06 | T | F | T | F |
| Sik3 | -1.40735 | 6.13669032 | 31.9720743 | 4.06E-07 | 3.10E-06 | T | F | F | F |
| Rgs12 | -1.5726933 | 5.46775762 | 31.9476036 | 4.09E-07 | 3.13E-06 | T | F | F | T |
| Aldh16a1 | 1.2868336 | 6.02451416 | 31.9439493 | 4.10E-07 | 3.13E-06 | F | F | F | T |
| Ticam2 | -6.0878075 | 1.95746272 | 31.9419006 | 4.10E-07 | 3.13E-06 | T | F | F | T |
| Eva1b | -4.1278629 | 2.6946281 | 31.9326538 | 4.11E-07 | 3.14E-06 | F | F | F | F |
| Crtc2 | 1.53089617 | 4.92973299 | 31.9000826 | 4.16E-07 | 3.17E-06 | F | F | F | F |
| Mapk7 | -1.5254126 | 5.61013439 | 31.8579183 | 4.22E-07 | 3.22E-06 | T | F | F | F |
| Tmem186 | 1.43809687 | 5.21877442 | 31.8353677 | 4.25E-07 | 3.24E-06 | F | F | T | F |
| Kitl | 1.27559648 | 6.09210219 | 31.8228663 | 4.27E-07 | 3.26E-06 | T | T | T | T |
| Sdcbp | 1.17918933 | 6.6110532 | 31.8134772 | 4.28E-07 | 3.27E-06 | T | F | T | T |
| Adam17 | 1.24310953 | 6.23129563 | 31.8089086 | 4.29E-07 | 3.27E-06 | T | F | F | F |
| Hydin | -9.1526151 | 1.62964895 | 31.8088146 | 4.29E-07 | 3.27E-06 | T | F | F | F |
| Kctd1 | -2.2593618 | 4.08836182 | 31.8008915 | 4.30E-07 | 3.28E-06 | T | F | F | F |
| Stk32a | -5.9715388 | 2.06519154 | 31.7969437 | 4.31E-07 | 3.28E-06 | T | F | T | T |
| F2r | -1.4024041 | 6.14236898 | 31.795308 | 4.31E-07 | 3.28E-06 | F | F | F | T |
| Serinc1 | 1.21617087 | 6.39128761 | 31.7739008 | 4.34E-07 | 3.30E-06 | T | F | T | F |
| Lars | 1.02221408 | 7.96190403 | 31.7183117 | 4.42E-07 | 3.37E-06 | F | F | F | F |
| Tnni1 | -9.3070203 | 1.79063102 | 31.6961749 | 4.46E-07 | 3.39E-06 | F | F | F | F |
| Rnf150 | -9.1464013 | 1.62274094 | 31.6913063 | 4.47E-07 | 3.39E-06 | T | T | T | T |

| | | | | | | | | | |
|---|---|---|---|---|---|---|---|---|---|
| Fiz1 | -1.41175 | 6.08287088 | 31.681622 | 4.48E-07 | 3.40E-06 | F | F | F | F |
| Fam163b | -9.1995608 | 1.66167565 | 31.6629468 | 4.51E-07 | 3.42E-06 | F | F | T | F |
| Tbc1d14 | -1.4677501 | 5.82865726 | 31.6626793 | 4.51E-07 | 3.42E-06 | T | F | T | T |
| C2cd5 | -1.9387449 | 4.53344268 | 31.6525755 | 4.53E-07 | 3.43E-06 | F | F | F | F |
| Btla | -9.1446742 | 1.62289984 | 31.6238979 | 4.57E-07 | 3.47E-06 | T | F | T | T |
| Zfp966 | 2.26882252 | 3.57839668 | 31.6155535 | 4.58E-07 | 3.48E-06 | F | F | T | F |
| Rhox2a | 9.01738031 | 0.44429081 | 31.6858306 | 4.65E-07 | 3.53E-06 | F | F | F | F |
| Frmpd3 | -9.1457581 | 1.61822981 | 31.5685577 | 4.66E-07 | 3.53E-06 | F | F | T | F |
| Klhdc8b | -9.1765736 | 1.64171452 | 31.565819 | 4.66E-07 | 3.53E-06 | T | F | F | F |
| Arhgef4 | 3.25962852 | 2.3087067 | 31.5575768 | 4.68E-07 | 3.54E-06 | T | F | F | T |
| Dlg5 | -1.160237 | 7.46421732 | 31.5535123 | 4.68E-07 | 3.54E-06 | T | F | T | T |
| Man1a | 1.69808892 | 4.74742094 | 31.5375193 | 4.71E-07 | 3.56E-06 | T | F | T | T |
| 1810046K0 | 7.00849055 | 0.65311947 | 31.5035446 | 4.76E-07 | 3.60E-06 | T | T | F | T |
| Hba-a1 | 4.72552618 | 1.46228776 | 31.5017787 | 4.77E-07 | 3.60E-06 | F | F | F | F |
| Ulk1 | -1.4007696 | 6.12874075 | 31.4967633 | 4.77E-07 | 3.61E-06 | F | F | F | T |
| Fam160b2 | 1.38916775 | 5.40113716 | 31.4740678 | 4.81E-07 | 3.64E-06 | F | F | T | F |
| Edem2 | 1.40061429 | 5.38675727 | 31.4679561 | 4.82E-07 | 3.64E-06 | F | F | F | F |
| Kif20a | -1.1410087 | 7.60308167 | 31.4600665 | 4.83E-07 | 3.65E-06 | F | F | T | F |
| Tmem266 | 3.20626565 | 2.35741099 | 31.4393866 | 4.87E-07 | 3.68E-06 | T | F | T | F |
| Ifngr2 | 1.77487614 | 4.30725976 | 31.4340357 | 4.88E-07 | 3.68E-06 | T | F | T | T |
| Cd109 | -6.0563798 | 1.93002794 | 31.4145652 | 4.91E-07 | 3.70E-06 | T | F | F | T |
| Cdkn2c | 3.19308918 | 2.36419967 | 31.4137124 | 4.91E-07 | 3.70E-06 | T | F | F | F |
| Tmem68 | 1.76579694 | 4.27493339 | 31.389976 | 4.95E-07 | 3.73E-06 | T | F | T | F |
| Slc25a37 | -3.2937624 | 3.12004867 | 31.3772012 | 4.97E-07 | 3.75E-06 | F | F | T | F |
| Stx4a | 1.64957381 | 4.6340006 | 31.3392328 | 5.04E-07 | 3.80E-06 | F | F | F | T |
| Zfp655 | 1.25195547 | 6.1579508 | 31.3330691 | 5.05E-07 | 3.80E-06 | T | F | T | T |
| Gtf2h1 | 1.18777023 | 6.61499507 | 31.3231708 | 5.07E-07 | 3.82E-06 | T | F | F | F |
| Ubr4 | 0.91462348 | 9.71692152 | 31.3220053 | 5.07E-07 | 3.82E-06 | F | F | F | F |
| Vmn2r26 | 9.00663244 | 0.43410508 | 31.4212141 | 5.09E-07 | 3.83E-06 | F | F | F | F |
| Herpud1 | 1.44929457 | 5.23983954 | 31.2886322 | 5.13E-07 | 3.86E-06 | F | F | T | T |
| Xlr5a | 2.63384524 | 2.98059386 | 31.2881339 | 5.13E-07 | 3.86E-06 | F | F | T | F |
| Abhd17a | -1.4908108 | 5.69829759 | 31.2876303 | 5.13E-07 | 3.86E-06 | F | F | F | F |
| Fam181b | -6.049959 | 1.92663032 | 31.2536264 | 5.19E-07 | 3.90E-06 | F | F | F | F |
| Wnt3 | -6.1186964 | 1.98069891 | 31.2529102 | 5.19E-07 | 3.90E-06 | T | F | T | F |
| Ube3b | -1.2385702 | 6.86794136 | 31.2306755 | 5.23E-07 | 3.93E-06 | T | F | T | T |
| Rmdn3 | 1.32643461 | 5.70283267 | 31.2216737 | 5.25E-07 | 3.94E-06 | F | F | F | F |
| Btbd6 | -2.3366873 | 3.97351192 | 31.204938 | 5.28E-07 | 3.96E-06 | F | F | F | F |
| Fmr1nb | 1.89712241 | 4.09422293 | 31.2011241 | 5.28E-07 | 3.96E-06 | F | F | F | F |
| Thsd7a | -2.9746325 | 3.49980161 | 31.1973299 | 5.29E-07 | 3.97E-06 | F | F | T | T |
| Map3k11 | 1.13623742 | 6.83881656 | 31.1858978 | 5.31E-07 | 3.98E-06 | F | F | F | F |
| Adk | -1.4097852 | 6.10505964 | 31.1830031 | 5.32E-07 | 3.98E-06 | T | T | T | T |
| Ddit4l | -9.1276246 | 1.60066427 | 31.1750456 | 5.33E-07 | 3.99E-06 | T | F | T | T |

| | | | | | | | | | |
|---|---|---|---|---|---|---|---|---|---|
| Ube2r2 | -1.195399 | 7.06199615 | 31.1637133 | 5.35E-07 | 4.01E-06 | T | F | T | T |
| Ephb4 | -1.1419472 | 7.53482832 | 31.1379505 | 5.40E-07 | 4.04E-06 | T | F | F | F |
| Crk | 1.026468 | 7.84756604 | 31.1143105 | 5.45E-07 | 4.07E-06 | T | F | T | T |
| Wdr35 | -2.0146127 | 4.34181923 | 31.0923584 | 5.49E-07 | 4.10E-06 | T | F | T | F |
| Kcnh2 | -3.6504891 | 2.87034341 | 31.0857377 | 5.50E-07 | 4.11E-06 | F | F | F | F |
| Adamts6 | -9.1203714 | 1.59454404 | 31.0698342 | 5.53E-07 | 4.13E-06 | T | F | T | T |
| Hhip | -6.0401982 | 1.91461056 | 31.0552099 | 5.56E-07 | 4.15E-06 | F | F | T | T |
| Ppp1r12c | 1.25781506 | 6.07618889 | 31.0474056 | 5.57E-07 | 4.16E-06 | F | F | F | F |
| Adcyap1r1 | -3.0765655 | 3.3226153 | 31.0390738 | 5.59E-07 | 4.17E-06 | T | F | T | T |
| Shisa7 | -9.1142095 | 1.59140096 | 31.0253631 | 5.61E-07 | 4.19E-06 | F | F | T | T |
| Rap1a | 1.37654793 | 5.41485811 | 31.0034609 | 5.66E-07 | 4.22E-06 | T | F | T | T |
| Zfp965 | 3.52462353 | 2.04158479 | 30.9816409 | 5.70E-07 | 4.25E-06 | F | F | T | T |
| Enpep | -9.1151067 | 1.59615155 | 30.9679417 | 5.73E-07 | 4.27E-06 | T | F | T | F |
| Scyl3 | 1.63492943 | 4.59878946 | 30.948044 | 5.77E-07 | 4.30E-06 | T | F | F | F |
| Naa11 | 6.29766206 | 0.77489445 | 30.9364387 | 5.79E-07 | 4.31E-06 | T | F | T | F |
| Tmx4 | 1.55914607 | 4.7674412 | 30.9363165 | 5.79E-07 | 4.31E-06 | T | F | F | T |
| Mcoln2 | -9.1357425 | 1.62054186 | 30.9238539 | 5.82E-07 | 4.33E-06 | F | F | T | T |
| Zranb3 | -1.7808215 | 4.79827821 | 30.9080821 | 5.85E-07 | 4.35E-06 | F | F | F | F |
| Acta2 | -4.9469253 | 2.71033191 | 30.8810204 | 5.90E-07 | 4.39E-06 | F | F | T | F |
| Tspyl2 | -2.3494862 | 3.96391632 | 30.8801363 | 5.90E-07 | 4.39E-06 | F | F | F | F |
| Uprt | -4.0480611 | 2.61508896 | 30.8540775 | 5.96E-07 | 4.43E-06 | F | F | T | F |
| Zbtb37 | 1.21632139 | 6.30878504 | 30.842882 | 5.98E-07 | 4.45E-06 | F | F | T | T |
| Egfl7 | -2.4267126 | 3.9733376 | 30.7980154 | 6.07E-07 | 4.51E-06 | F | F | F | F |
| Hsd17b7 | 1.38386228 | 5.35132585 | 30.7894236 | 6.09E-07 | 4.53E-06 | T | F | T | T |
| Tead1 | -1.1002787 | 7.87933034 | 30.7846197 | 6.10E-07 | 4.53E-06 | T | F | F | F |
| Pitrm1 | 1.01301514 | 7.88818606 | 30.7755259 | 6.12E-07 | 4.54E-06 | T | F | F | T |
| Cdo1 | -4.2580228 | 2.49624854 | 30.7653674 | 6.14E-07 | 4.56E-06 | T | F | T | F |
| Pfkl | -1.0425118 | 8.63426978 | 30.7613413 | 6.15E-07 | 4.56E-06 | F | F | F | F |
| Mecom | -9.101577 | 1.58184672 | 30.7503815 | 6.17E-07 | 4.58E-06 | T | F | T | T |
| Gask1b | -2.6917436 | 3.70532585 | 30.7407562 | 6.20E-07 | 4.59E-06 | T | F | T | F |
| Prdx1 | -1.0563995 | 8.41562113 | 30.7297085 | 6.22E-07 | 4.61E-06 | F | F | F | F |
| Rbmxl1 | -1.1367234 | 7.52641213 | 30.7252606 | 6.23E-07 | 4.61E-06 | F | F | F | F |
| Bmper | -2.9895937 | 3.28688808 | 30.7150903 | 6.25E-07 | 4.63E-06 | T | F | T | F |
| Nectin2 | -1.1642715 | 7.37540013 | 30.6672539 | 6.36E-07 | 4.70E-06 | T | F | F | F |
| 4921524J17 | 1.68511737 | 4.39841551 | 30.6602223 | 6.37E-07 | 4.71E-06 | T | F | T | F |
| Grk6 | -1.2652356 | 6.66895973 | 30.6599385 | 6.37E-07 | 4.71E-06 | T | F | T | F |
| Ttc21a | -9.119745 | 1.58914412 | 30.6454852 | 6.40E-07 | 4.74E-06 | F | F | F | F |
| Smarcc1 | -0.9995247 | 9.28277152 | 30.6140873 | 6.47E-07 | 4.79E-06 | T | F | T | F |
| Camkk2 | -1.9007708 | 4.53009823 | 30.5775028 | 6.56E-07 | 4.84E-06 | F | F | T | F |
| Rab38 | -9.127554 | 1.5948631 | 30.5673625 | 6.58E-07 | 4.86E-06 | F | F | T | T |
| Akap1 | 1.20017231 | 6.35221177 | 30.5330294 | 6.66E-07 | 4.92E-06 | F | F | T | F |
| Cenpf | -1.0625406 | 8.42366095 | 30.5252428 | 6.68E-07 | 4.93E-06 | T | F | F | T |

| | | | | | | | | | |
|---|---|---|---|---|---|---|---|---|---|
| Gpr155 | -3.8034109 | 2.73431902 | 30.5181764 | 6.69E-07 | 4.94E-06 | T | F | T | T |
| Emc3 | 1.33476603 | 5.59254464 | 30.4712001 | 6.80E-07 | 5.02E-06 | F | F | F | F |
| Kctd18 | -9.0962702 | 1.58035804 | 30.4603487 | 6.83E-07 | 5.04E-06 | T | F | T | F |
| Clic4 | 0.93129278 | 9.07027527 | 30.423897 | 6.91E-07 | 5.10E-06 | T | F | T | T |
| Tap2 | 4.2474065 | 1.50338044 | 30.3885294 | 7.00E-07 | 5.16E-06 | F | F | T | F |
| Efemp1 | -9.0883578 | 1.56348576 | 30.3855238 | 7.01E-07 | 5.16E-06 | F | F | F | T |
| Gm8108 | 5.74329016 | 0.96276892 | 30.3803043 | 7.02E-07 | 5.17E-06 | F | F | F | F |
| Nodal | -9.1060669 | 1.5761107 | 30.3712814 | 7.04E-07 | 5.18E-06 | F | F | T | F |
| Cybc1 | 1.45600134 | 5.05404224 | 30.3709517 | 7.04E-07 | 5.18E-06 | F | F | T | F |
| Slc9a1 | 1.18893532 | 6.39875324 | 30.3696482 | 7.05E-07 | 5.18E-06 | F | F | T | F |
| Stat4 | -9.0811859 | 1.56138506 | 30.358852 | 7.07E-07 | 5.20E-06 | F | F | T | F |
| Fth1 | -1.0801631 | 8.39904806 | 30.3421564 | 7.11E-07 | 5.23E-06 | F | F | F | F |
| Prdm1 | 1.50377116 | 5.0585985 | 30.3327707 | 7.14E-07 | 5.25E-06 | F | F | T | F |
| Mlxip | -1.2366614 | 6.7744381 | 30.3297157 | 7.15E-07 | 5.25E-06 | F | F | T | F |
| Enah | -1.0886378 | 7.97156564 | 30.3160095 | 7.18E-07 | 5.27E-06 | T | F | T | T |
| Uggt2 | -2.1935973 | 4.1101864 | 30.2940336 | 7.23E-07 | 5.31E-06 | T | F | T | T |
| Exoc6 | -1.8853772 | 4.54094813 | 30.293916 | 7.23E-07 | 5.31E-06 | F | F | F | F |
| Dennd1b | 1.37888391 | 5.32645227 | 30.2820638 | 7.26E-07 | 5.33E-06 | T | F | T | T |
| Pebp1 | -1.103382 | 7.83761347 | 30.2794346 | 7.27E-07 | 5.33E-06 | F | F | T | F |
| Jade1 | 1.06310315 | 7.29648688 | 30.2662578 | 7.30E-07 | 5.36E-06 | T | F | T | T |
| Plac9b | 2.88837691 | 2.63362907 | 30.2535874 | 7.34E-07 | 5.38E-06 | F | F | F | F |
| Unc5a | 2.48538957 | 3.15098814 | 30.1997834 | 7.48E-07 | 5.48E-06 | T | F | T | T |
| Vsig2 | 4.14383159 | 1.59203928 | 30.1957853 | 7.49E-07 | 5.48E-06 | F | F | F | F |
| Emilin2 | -4.0566476 | 2.62721768 | 30.1950402 | 7.49E-07 | 5.48E-06 | T | F | T | F |
| Ephx2 | -3.0521969 | 3.21739003 | 30.1619261 | 7.58E-07 | 5.54E-06 | T | F | T | T |
| Sh3bgrl3 | 1.33301829 | 5.60170769 | 30.1614613 | 7.58E-07 | 5.54E-06 | F | F | T | F |
| Zfp91 | 1.03567189 | 7.48497693 | 30.156486 | 7.59E-07 | 5.55E-06 | T | T | T | T |
| Agps | 1.01715682 | 7.70765245 | 30.1391157 | 7.64E-07 | 5.58E-06 | T | F | T | T |
| Adam15 | 1.97337014 | 3.91255034 | 30.131376 | 7.66E-07 | 5.60E-06 | F | F | F | F |
| Cdk5rap2 | -1.2684119 | 6.59430939 | 30.110623 | 7.71E-07 | 5.64E-06 | F | F | T | T |
| Dpp6 | -9.06848 | 1.55000575 | 30.0979452 | 7.75E-07 | 5.66E-06 | T | F | F | T |
| Mybl2 | 1.04135564 | 7.51178009 | 30.0768156 | 7.80E-07 | 5.70E-06 | F | F | F | F |
| Pank2 | -1.7326945 | 4.84810496 | 30.0672932 | 7.83E-07 | 5.71E-06 | F | F | F | T |
| Rcc2 | -0.9795298 | 9.36904014 | 30.0672179 | 7.83E-07 | 5.71E-06 | F | F | T | F |
| Qprt | -1.9910137 | 4.31696593 | 30.0663655 | 7.83E-07 | 5.71E-06 | F | F | F | F |
| Stx12 | 1.25881688 | 5.93206328 | 30.0536076 | 7.87E-07 | 5.74E-06 | F | F | F | F |
| Hoxa2 | -9.0817725 | 1.56820412 | 30.0365614 | 7.91E-07 | 5.77E-06 | F | F | F | F |
| Cap1 | 0.98583713 | 8.1311962 | 30.029267 | 7.93E-07 | 5.78E-06 | F | F | T | F |
| Slfn9 | -1.6015746 | 5.1744866 | 30.016698 | 7.97E-07 | 5.81E-06 | T | F | F | T |
| Rdh11 | 1.42931867 | 5.11410677 | 30.0114131 | 7.98E-07 | 5.81E-06 | T | F | T | F |
| Traf4 | -1.3640305 | 6.27383817 | 30.0108119 | 7.99E-07 | 5.81E-06 | T | F | T | F |
| Pam16 | 2.1296985 | 3.79500303 | 30.0072751 | 7.99E-07 | 5.82E-06 | F | F | F | F |

| | | | | | | | | | |
|---|---|---|---|---|---|---|---|---|---|
| Ccdc160 | -9.06657 | 1.55085169 | 29.9698525 | 8.10E-07 | 5.89E-06 | F | F | F | F |
| Vwa2 | -3.6925408 | 2.77287923 | 29.937326 | 8.19E-07 | 5.96E-06 | F | F | T | F |
| Nagpa | 1.46603301 | 5.05934905 | 29.9148084 | 8.26E-07 | 6.00E-06 | F | F | F | F |
| Stat2 | 1.88214379 | 4.02996926 | 29.9070577 | 8.28E-07 | 6.02E-06 | F | F | T | F |
| St6galnac6 | -3.890918 | 2.6491405 | 29.9039493 | 8.29E-07 | 6.02E-06 | F | F | T | F |
| Sptb | -3.6806352 | 2.76393641 | 29.8881114 | 8.33E-07 | 6.05E-06 | T | F | T | F |
| Agpat4 | 1.28786527 | 5.74412732 | 29.8873357 | 8.34E-07 | 6.05E-06 | T | F | F | F |
| Asb1 | -2.3017857 | 3.94609605 | 29.8869908 | 8.34E-07 | 6.05E-06 | F | F | T | F |
| Hid1 | 2.98207355 | 2.51137073 | 29.8691276 | 8.39E-07 | 6.09E-06 | F | F | T | F |
| Rb1cc1 | 1.27373602 | 6.01147531 | 29.8629089 | 8.41E-07 | 6.10E-06 | T | F | F | F |
| Tmem263 | 1.35445321 | 5.39795983 | 29.8609012 | 8.41E-07 | 6.10E-06 | T | T | F | T |
| Actc1 | -9.0606833 | 1.54519706 | 29.8592009 | 8.42E-07 | 6.10E-06 | F | F | F | F |
| Gucy1a1 | -5.2456885 | 2.05604804 | 29.8525328 | 8.44E-07 | 6.12E-06 | T | T | F | F |
| Atp2a1 | -4.82444 | 2.1912541 | 29.8422454 | 8.47E-07 | 6.14E-06 | F | F | F | F |
| Slc35d1 | 1.31389139 | 5.60021543 | 29.8334483 | 8.50E-07 | 6.15E-06 | T | F | T | F |
| Yipf6 | 1.42539622 | 5.13174531 | 29.8294 | 8.51E-07 | 6.16E-06 | F | F | T | F |
| Calca | -9.1243176 | 1.58794549 | 29.8073058 | 8.57E-07 | 6.20E-06 | F | F | F | F |
| Rnf208 | -5.3039256 | 2.09990751 | 29.8072668 | 8.57E-07 | 6.20E-06 | F | F | F | T |
| Limch1 | -2.4269483 | 3.92861673 | 29.7983596 | 8.60E-07 | 6.22E-06 | T | T | F | T |
| Ahdc1 | 0.9979803 | 7.90383911 | 29.7777832 | 8.66E-07 | 6.26E-06 | F | F | F | F |
| Dcun1d5 | 1.19488151 | 6.28281788 | 29.7640032 | 8.70E-07 | 6.29E-06 | F | F | T | T |
| Clasp1 | -1.1116509 | 7.66095242 | 29.7631787 | 8.71E-07 | 6.29E-06 | T | F | T | T |
| Gpr85 | -9.0934234 | 1.56140335 | 29.7565982 | 8.73E-07 | 6.30E-06 | T | T | T | T |
| Plxna1 | -1.0697025 | 8.11626441 | 29.741458 | 8.77E-07 | 6.34E-06 | F | F | T | T |
| Klf15 | -4.5289971 | 2.30182194 | 29.7236677 | 8.83E-07 | 6.37E-06 | F | F | F | F |
| Clcn4 | 1.48246762 | 4.97451216 | 29.7198602 | 8.84E-07 | 6.38E-06 | T | F | T | T |
| Derl1 | 1.09598215 | 6.87973613 | 29.7141694 | 8.86E-07 | 6.39E-06 | T | F | T | T |
| Ackr2 | 6.93755043 | 0.5856506 | 29.7102129 | 8.87E-07 | 6.40E-06 | F | F | T | T |
| Pdpk1 | 1.11276768 | 6.77332735 | 29.6690151 | 9.00E-07 | 6.49E-06 | T | F | T | T |
| Lrrc45 | -2.1075562 | 4.16177433 | 29.6675155 | 9.00E-07 | 6.49E-06 | T | F | F | F |
| Strn | 1.09767142 | 6.87581624 | 29.6661766 | 9.01E-07 | 6.49E-06 | T | F | T | T |
| Eif1b | -1.6857552 | 4.93135022 | 29.6507009 | 9.06E-07 | 6.52E-06 | F | F | T | F |
| Olfm1 | -1.785971 | 4.70028694 | 29.6278385 | 9.13E-07 | 6.57E-06 | T | F | T | F |
| S1pr5 | -5.9731448 | 1.85067678 | 29.5938554 | 9.24E-07 | 6.65E-06 | T | F | T | T |
| Prl2c2 | 3.07124833 | 2.56687185 | 29.5885658 | 9.26E-07 | 6.66E-06 | F | F | F | F |
| Hacd4 | -9.0425186 | 1.52532107 | 29.5879246 | 9.26E-07 | 6.66E-06 | T | F | T | F |
| Tjap1 | -1.3884881 | 5.94780559 | 29.5720847 | 9.31E-07 | 6.69E-06 | F | F | F | F |
| Gucd1 | -1.6828543 | 4.9320767 | 29.5607825 | 9.35E-07 | 6.72E-06 | T | F | T | T |
| Phox2a | -9.0555385 | 1.52986529 | 29.5581242 | 9.36E-07 | 6.72E-06 | F | F | F | F |
| Tsacc | -9.0574082 | 1.53114685 | 29.55043 | 9.38E-07 | 6.74E-06 | F | F | F | F |
| Gpr156 | 5.16035958 | 0.9962493 | 29.5470934 | 9.39E-07 | 6.74E-06 | T | F | T | F |
| Tor1aip2 | 1.0087859 | 7.68175767 | 29.5436261 | 9.40E-07 | 6.75E-06 | T | F | T | T |

| | | | | | | | | | |
|---|---|---|---|---|---|---|---|---|---|
| Cpne4 | -9.0874158 | 1.55503558 | 29.5380174 | 9.42E-07 | 6.76E-06 | T | F | T | T |
| Ube2o | 1.02388735 | 7.4949099 | 29.503426 | 9.54E-07 | 6.84E-06 | F | F | F | F |
| Rasip1 | -3.3037936 | 3.14768839 | 29.5009912 | 9.55E-07 | 6.84E-06 | F | F | F | F |
| Leng8 | 1.0824393 | 7.12837195 | 29.4905402 | 9.58E-07 | 6.87E-06 | F | F | T | F |
| Slc30a7 | 1.20950499 | 6.15381892 | 29.4347724 | 9.77E-07 | 7.00E-06 | F | F | T | T |
| Stoml1 | -5.9666658 | 1.84840229 | 29.4133102 | 9.84E-07 | 7.05E-06 | F | F | T | T |
| Camk1 | -9.0893126 | 1.5804132 | 29.4129058 | 9.85E-07 | 7.05E-06 | F | F | F | F |
| Ptprf | -0.9777117 | 9.26295994 | 29.4113038 | 9.85E-07 | 7.05E-06 | F | F | T | T |
| Tbx4 | -6.2947336 | 3.01797462 | 29.3766015 | 9.97E-07 | 7.14E-06 | F | F | F | F |
| Slc8a3 | -9.0507567 | 1.52401759 | 29.3421556 | 1.01E-06 | 7.22E-06 | F | F | F | F |
| Mmp17 | -2.4280425 | 3.77549909 | 29.3081048 | 1.02E-06 | 7.31E-06 | F | F | T | T |
| Ghr | -2.8794518 | 3.30284421 | 29.2987605 | 1.02E-06 | 7.33E-06 | T | F | T | T |
| Agpat3 | 1.11541882 | 6.74280028 | 29.2910519 | 1.03E-06 | 7.34E-06 | T | F | T | T |
| Ppm1l | -2.0509856 | 4.20573164 | 29.273619 | 1.03E-06 | 7.39E-06 | T | T | T | T |
| Ptgis | 1.28386906 | 5.68981172 | 29.223107 | 1.05E-06 | 7.52E-06 | F | F | F | F |
| Smad9 | -2.9247602 | 3.2992661 | 29.2218838 | 1.05E-06 | 7.52E-06 | F | F | T | F |
| Naa25 | 1.03907615 | 7.29128072 | 29.2119779 | 1.06E-06 | 7.54E-06 | T | F | T | T |
| Cklf | 1.95474762 | 3.90101991 | 29.2044038 | 1.06E-06 | 7.56E-06 | T | F | T | F |
| Acat1 | 1.66371024 | 4.39681826 | 29.1933258 | 1.06E-06 | 7.59E-06 | T | F | F | T |
| Senp1 | 1.03209963 | 7.31724041 | 29.1848992 | 1.07E-06 | 7.61E-06 | T | F | T | T |
| Col1a2 | -2.1962442 | 4.10191801 | 29.1618382 | 1.08E-06 | 7.67E-06 | T | F | F | F |
| Map2k6 | -1.7484511 | 4.74128613 | 29.1462755 | 1.08E-06 | 7.71E-06 | F | F | T | T |
| Epb41l1 | 1.48805622 | 5.19522 | 29.1386451 | 1.08E-06 | 7.72E-06 | T | F | T | F |
| Plpp3 | 1.29505021 | 5.72576122 | 29.1237137 | 1.09E-06 | 7.76E-06 | T | F | F | F |
| Rufy1 | -2.4023747 | 3.79224104 | 29.0949944 | 1.10E-06 | 7.84E-06 | F | F | T | F |
| Prkra | -2.0444279 | 4.21264137 | 29.0774796 | 1.11E-06 | 7.89E-06 | T | F | F | F |
| Slc30a5 | 1.17423939 | 6.33893232 | 29.0723779 | 1.11E-06 | 7.90E-06 | F | F | F | T |
| Xlr3c | 3.43623398 | 2.17727723 | 29.0714821 | 1.11E-06 | 7.90E-06 | F | F | F | F |
| Cyb561a3 | 1.94031533 | 3.89961709 | 29.0712157 | 1.11E-06 | 7.90E-06 | T | F | T | F |
| Cpxm2 | -9.0199555 | 1.49894993 | 29.0536113 | 1.12E-06 | 7.94E-06 | T | F | T | T |
| Bicdl1 | 1.44925649 | 5.02861205 | 29.0474826 | 1.12E-06 | 7.96E-06 | F | F | F | F |
| Cbx1 | -1.3474544 | 6.08128125 | 29.0422226 | 1.12E-06 | 7.97E-06 | T | F | T | F |
| Nfyb | -1.4458618 | 5.62474936 | 29.0325738 | 1.13E-06 | 7.99E-06 | T | T | F | T |
| Mutyh | -2.876043 | 3.29231486 | 29.0085834 | 1.14E-06 | 8.06E-06 | F | F | F | F |
| Syndig1l | -6.1505574 | 2.00532209 | 28.9629358 | 1.15E-06 | 8.19E-06 | F | F | T | F |
| Adcy3 | -1.9252195 | 4.36465553 | 28.9553489 | 1.16E-06 | 8.21E-06 | F | F | F | T |
| Pfkfb2 | 1.9529624 | 3.8670645 | 28.9506804 | 1.16E-06 | 8.22E-06 | T | T | T | T |
| Sp100 | 3.638292 | 1.89891977 | 28.9449109 | 1.16E-06 | 8.23E-06 | T | F | T | F |
| Adamts20 | -3.7824739 | 2.64384824 | 28.9398503 | 1.16E-06 | 8.24E-06 | T | F | T | T |
| B2m | 1.84476804 | 4.06745457 | 28.9021745 | 1.18E-06 | 8.35E-06 | F | F | F | F |
| Kmt5a | 0.99515078 | 7.72406884 | 28.8742312 | 1.19E-06 | 8.43E-06 | F | F | T | F |
| Bag4 | -1.3890688 | 5.91182657 | 28.8693747 | 1.19E-06 | 8.44E-06 | T | F | T | T |

| | | | | | | | | | |
|---|---|---|---|---|---|---|---|---|---|
| Atp1a3 | -5.1884211 | 2.0515417 | 28.8590242 | 1.20E-06 | 8.47E-06 | F | F | F | F |
| Eea1 | 1.39336981 | 5.53085797 | 28.8559128 | 1.20E-06 | 8.48E-06 | T | F | F | T |
| Kcnj3 | -4.6684504 | 2.24859284 | 28.8457761 | 1.20E-06 | 8.51E-06 | T | F | T | F |
| Eomes | -3.5750009 | 3.08575535 | 28.8337882 | 1.21E-06 | 8.54E-06 | T | F | T | F |
| Nexn | -9.0028568 | 1.48811651 | 28.8144348 | 1.22E-06 | 8.59E-06 | F | F | T | F |
| Stag2 | -1.148027 | 7.16196412 | 28.8123945 | 1.22E-06 | 8.60E-06 | T | T | F | T |
| Espnl | 3.83519889 | 1.66783047 | 28.8109648 | 1.22E-06 | 8.60E-06 | T | F | F | T |
| Neu1 | 1.32927571 | 5.53858955 | 28.8061781 | 1.22E-06 | 8.61E-06 | F | F | F | F |
| Sumo1 | -1.2842143 | 6.36850608 | 28.8010343 | 1.22E-06 | 8.62E-06 | F | F | T | F |
| Dera | 1.26782402 | 5.71633489 | 28.7777874 | 1.23E-06 | 8.69E-06 | T | F | F | T |
| Hpse | 2.66082655 | 2.80536629 | 28.7326303 | 1.25E-06 | 8.83E-06 | T | F | F | T |
| Irf4 | 6.20996849 | 0.691633 | 28.7006859 | 1.27E-06 | 8.93E-06 | T | F | F | T |
| Tollip | 1.19426147 | 6.23020342 | 28.7001003 | 1.27E-06 | 8.93E-06 | F | F | T | T |
| Zbtb42 | -4.7840228 | 2.15104758 | 28.6924988 | 1.27E-06 | 8.95E-06 | F | F | F | T |
| Nme3 | -2.4232921 | 3.73693793 | 28.6701688 | 1.28E-06 | 9.02E-06 | F | F | F | F |
| Gjb1 | -9.0285768 | 1.52119271 | 28.6544244 | 1.29E-06 | 9.06E-06 | T | F | T | T |
| Myh7 | -4.2827126 | 2.48747522 | 28.6472987 | 1.29E-06 | 9.08E-06 | F | F | F | F |
| Ahcy | -1.0004125 | 8.6595401 | 28.6380553 | 1.29E-06 | 9.11E-06 | F | F | F | F |
| Arpin | 1.45246858 | 4.94175018 | 28.6193949 | 1.30E-06 | 9.17E-06 | T | F | T | T |
| Reep2 | -3.2788699 | 2.95384307 | 28.6182086 | 1.30E-06 | 9.17E-06 | F | F | T | F |
| Cthrc1 | -3.1399796 | 3.08257481 | 28.6104787 | 1.31E-06 | 9.19E-06 | F | F | F | F |
| Eef2kmt | 1.66470401 | 4.30607726 | 28.5903065 | 1.32E-06 | 9.25E-06 | F | F | T | F |
| Arxes1 | -2.1358835 | 4.09279705 | 28.5856911 | 1.32E-06 | 9.27E-06 | F | T | F | T |
| Rgs3 | -2.566865 | 3.56918256 | 28.5566661 | 1.33E-06 | 9.36E-06 | T | F | T | T |
| Cfap54 | -8.9976257 | 1.47647516 | 28.5500743 | 1.34E-06 | 9.38E-06 | F | F | F | F |
| Cux1 | -1.2401972 | 6.55323515 | 28.5081044 | 1.36E-06 | 9.52E-06 | T | F | T | T |
| Itgav | 1.11735085 | 6.63379156 | 28.4815997 | 1.37E-06 | 9.60E-06 | T | F | T | F |
| Lrrc8d | 1.25762043 | 5.7386716 | 28.4786698 | 1.37E-06 | 9.61E-06 | T | F | F | T |
| Fermt1 | -3.389478 | 2.88107467 | 28.4786366 | 1.37E-06 | 9.61E-06 | T | F | F | T |
| Zdhhc12 | 1.35237126 | 5.25908784 | 28.4572152 | 1.38E-06 | 9.68E-06 | F | F | F | F |
| G3bp2 | -1.0568653 | 8.08758909 | 28.4541699 | 1.38E-06 | 9.69E-06 | F | F | F | F |
| Atp13a2 | -1.6745211 | 4.87147273 | 28.4465841 | 1.39E-06 | 9.71E-06 | F | F | F | F |
| Trp53bp1 | -1.1488522 | 7.05985809 | 28.440634 | 1.39E-06 | 9.73E-06 | T | T | F | T |
| Doc2a | -5.9257084 | 1.80288445 | 28.428356 | 1.39E-06 | 9.76E-06 | T | F | F | F |
| Itga2 | 3.30381611 | 2.07059031 | 28.4024862 | 1.41E-06 | 9.85E-06 | F | F | T | T |
| Chrnb2 | -8.9808411 | 1.46586729 | 28.3994122 | 1.41E-06 | 9.86E-06 | T | F | T | T |
| Tubb2a | 1.56204828 | 4.57315369 | 28.3958653 | 1.41E-06 | 9.87E-06 | F | F | F | F |
| Rmnd5b | 1.59773423 | 4.50594134 | 28.3942586 | 1.41E-06 | 9.87E-06 | F | F | T | F |
| Npc1l1 | -8.9824409 | 1.47055602 | 28.3854433 | 1.42E-06 | 9.90E-06 | F | F | F | F |
| Zcchc12 | -8.9780058 | 1.46366277 | 28.3458889 | 1.44E-06 | 1.00E-05 | T | T | T | T |
| Rbm12b1 | 1.94230056 | 3.87386046 | 28.3456893 | 1.44E-06 | 1.00E-05 | T | F | T | T |
| Tmem130 | -4.090226 | 2.44089695 | 28.3090954 | 1.46E-06 | 1.02E-05 | F | F | T | F |

| | | | | | | | | | |
|---|---|---|---|---|---|---|---|---|---|
| Prkcz | 1.36269488 | 5.20414665 | 28.3085765 | 1.46E-06 | 1.02E-05 | T | F | T | T |
| Yipf4 | 1.60531962 | 4.43956086 | 28.3039078 | 1.46E-06 | 1.02E-05 | T | F | F | T |
| Pacsin3 | -2.5936177 | 3.55644486 | 28.2631492 | 1.48E-06 | 1.03E-05 | F | F | T | F |
| Cacna1d | -3.8304369 | 2.62592103 | 28.2374067 | 1.49E-06 | 1.04E-05 | F | F | F | T |
| Nr1h5 | -5.0572618 | 3.35226782 | 28.2292852 | 1.50E-06 | 1.04E-05 | F | F | T | F |
| Ikzf5 | 1.47331389 | 4.82001105 | 28.2008875 | 1.51E-06 | 1.05E-05 | T | F | F | T |
| Mgll | 3.17232894 | 2.31313931 | 28.157483 | 1.54E-06 | 1.07E-05 | T | F | T | T |
| Rnft1 | 2.02191585 | 3.67450548 | 28.1521225 | 1.54E-06 | 1.07E-05 | T | F | F | T |
| Zfp286 | -8.9675006 | 1.45374352 | 28.1461664 | 1.54E-06 | 1.07E-05 | T | F | T | F |
| Mcm7 | -0.996659 | 8.62549437 | 28.1375429 | 1.55E-06 | 1.08E-05 | F | F | F | F |
| Hykk | -2.1993785 | 3.96788259 | 28.0923906 | 1.57E-06 | 1.09E-05 | T | F | T | T |
| 0610030E2( | 1.3152771 | 5.3903427 | 28.0814505 | 1.58E-06 | 1.10E-05 | T | F | T | T |
| Nrarp | -2.1132893 | 4.07040204 | 28.0490785 | 1.60E-06 | 1.11E-05 | T | F | T | F |
| Syt3 | -5.9037453 | 1.78286427 | 28.0428515 | 1.60E-06 | 1.11E-05 | F | F | F | T |
| Ppcdc | 1.27740175 | 5.58774948 | 28.0249511 | 1.61E-06 | 1.12E-05 | F | F | T | F |
| Tctn3 | 1.71102807 | 4.32277243 | 28.023186 | 1.61E-06 | 1.12E-05 | T | F | F | T |
| Paip1 | -1.3374981 | 5.99769505 | 28.0225859 | 1.61E-06 | 1.12E-05 | T | F | T | T |
| Agrp | 4.4866702 | 1.35943685 | 28.0195635 | 1.61E-06 | 1.12E-05 | F | F | F | F |
| Rundc3a | -8.9623641 | 1.44585914 | 28.003403 | 1.62E-06 | 1.13E-05 | F | F | T | F |
| Gxylt1 | 1.09419763 | 6.72175781 | 27.9945893 | 1.63E-06 | 1.13E-05 | T | T | T | T |
| Slc4a3 | -8.9729447 | 1.45171703 | 27.9872466 | 1.63E-06 | 1.13E-05 | F | F | T | T |
| Col4a3bp | 1.15519808 | 6.32500273 | 27.9772848 | 1.64E-06 | 1.14E-05 | T | F | T | T |
| Sv2a | -3.376906 | 2.84521195 | 27.967978 | 1.64E-06 | 1.14E-05 | T | F | T | T |
| Stard10 | -1.672622 | 4.8563616 | 27.9654224 | 1.65E-06 | 1.14E-05 | F | F | F | F |
| Gm6871 | 3.09373095 | 2.2468785 | 27.9585035 | 1.65E-06 | 1.14E-05 | F | F | F | T |
| Trim41 | 1.17225808 | 6.21026526 | 27.9459605 | 1.66E-06 | 1.15E-05 | T | F | F | T |
| Aff2 | -8.9916788 | 1.48696193 | 27.9226904 | 1.67E-06 | 1.16E-05 | T | F | F | F |
| Cd55 | -3.8805108 | 2.55027614 | 27.893881 | 1.69E-06 | 1.17E-05 | T | F | T | F |
| Itga9 | 1.36601668 | 5.19646916 | 27.8803252 | 1.70E-06 | 1.17E-05 | F | F | T | T |
| Oasl2 | 4.1276765 | 1.71990882 | 27.8785201 | 1.70E-06 | 1.17E-05 | F | F | F | T |
| Pick1 | -2.2129869 | 3.94700337 | 27.8753908 | 1.70E-06 | 1.17E-05 | F | F | F | F |
| Cs | 0.94901198 | 8.25141073 | 27.8678837 | 1.70E-06 | 1.18E-05 | F | F | F | F |
| Gap43 | -3.6989402 | 2.63561416 | 27.8575151 | 1.71E-06 | 1.18E-05 | F | F | T | F |
| Eln | -8.9763276 | 1.47095365 | 27.8424666 | 1.72E-06 | 1.19E-05 | F | F | F | F |
| Hoxb6 | -4.5135279 | 2.48855289 | 27.8342868 | 1.72E-06 | 1.19E-05 | F | F | F | F |
| Jade3 | 1.11434824 | 6.59635401 | 27.8266771 | 1.73E-06 | 1.19E-05 | F | F | T | F |
| St7l | 1.64850051 | 4.29609213 | 27.821835 | 1.73E-06 | 1.20E-05 | F | F | T | T |
| Dpy19l3 | 1.30859096 | 5.39137909 | 27.7995349 | 1.75E-06 | 1.20E-05 | T | F | T | T |
| Ncoa6 | 1.04219632 | 7.01051512 | 27.7894879 | 1.75E-06 | 1.21E-05 | T | F | F | F |
| Tns3 | -1.2534022 | 6.4083518 | 27.7651323 | 1.77E-06 | 1.22E-05 | T | F | T | F |
| Scn4a | -8.973184 | 1.44932058 | 27.7609641 | 1.77E-06 | 1.22E-05 | T | F | F | F |
| Mmgt1 | 1.21847314 | 5.88206765 | 27.7503461 | 1.78E-06 | 1.22E-05 | T | T | T | T |

| | | | | | | | | | |
|---|---|---|---|---|---|---|---|---|---|
| She | -5.9007078 | 1.78949766 | 27.7436849 | 1.78E-06 | 1.23E-05 | T | F | T | T |
| Tex264 | 1.47121199 | 4.82600082 | 27.7339962 | 1.79E-06 | 1.23E-05 | F | F | F | F |
| Plcb3 | 1.08725818 | 6.71204389 | 27.7304481 | 1.79E-06 | 1.23E-05 | F | F | F | T |
| Aifm1 | 1.1623562 | 6.23433429 | 27.7235293 | 1.79E-06 | 1.23E-05 | F | F | F | F |
| Lefty2 | -4.2004509 | 4.43618264 | 27.718797 | 1.80E-06 | 1.24E-05 | T | F | T | F |
| Gtpbp1 | -1.2660549 | 6.31918422 | 27.7037093 | 1.81E-06 | 1.24E-05 | T | F | F | T |
| Slc39a10 | 1.02079354 | 7.46174984 | 27.7029486 | 1.81E-06 | 1.24E-05 | T | F | F | T |
| Emp1 | -8.9437988 | 1.43125348 | 27.6990443 | 1.81E-06 | 1.24E-05 | T | F | T | T |
| Kif13a | 1.11316549 | 6.54078456 | 27.6973594 | 1.81E-06 | 1.24E-05 | F | F | T | F |
| Otud3 | 1.59795045 | 4.4400041 | 27.696393 | 1.81E-06 | 1.24E-05 | F | F | F | T |
| Rsph4a | -8.9531425 | 1.43417309 | 27.6950884 | 1.81E-06 | 1.24E-05 | F | F | T | F |
| Rdh13 | 1.26270149 | 5.61880484 | 27.6709457 | 1.83E-06 | 1.25E-05 | F | F | F | F |
| Ralgapa2 | -1.3500823 | 5.96770959 | 27.6692731 | 1.83E-06 | 1.26E-05 | T | F | F | T |
| Pea15a | 1.12980873 | 6.43739736 | 27.6574825 | 1.84E-06 | 1.26E-05 | T | T | T | T |
| Arhgap8 | 2.29680768 | 3.21251708 | 27.6366636 | 1.85E-06 | 1.27E-05 | F | F | F | T |
| Sparcl1 | -5.4129153 | 2.22586841 | 27.6345125 | 1.85E-06 | 1.27E-05 | F | F | F | F |
| Mfng | -8.9851849 | 1.45780063 | 27.634098 | 1.85E-06 | 1.27E-05 | F | F | T | T |
| Maob | -8.941427 | 1.431178 | 27.633432 | 1.85E-06 | 1.27E-05 | T | F | F | T |
| Marc2 | 2.0326665 | 3.71813573 | 27.6049429 | 1.87E-06 | 1.28E-05 | F | F | T | F |
| Xkr4 | -8.9470496 | 1.42863851 | 27.5942873 | 1.88E-06 | 1.29E-05 | F | F | T | F |
| Dmd | -2.9875031 | 3.12018444 | 27.5886921 | 1.88E-06 | 1.29E-05 | T | F | F | T |
| Zdhhc20 | 1.20905163 | 5.95417741 | 27.5799591 | 1.89E-06 | 1.29E-05 | T | F | T | T |
| Got2 | 0.90757803 | 8.78689138 | 27.5797501 | 1.89E-06 | 1.29E-05 | F | F | T | F |
| 1700037H0 | -1.8260945 | 4.46573281 | 27.5735648 | 1.89E-06 | 1.29E-05 | F | F | F | F |
| Pdcd6ip | 0.97608754 | 7.66252221 | 27.540782 | 1.92E-06 | 1.31E-05 | F | F | T | F |
| Phactr2 | -2.0999429 | 4.07928159 | 27.5325559 | 1.92E-06 | 1.31E-05 | F | F | T | T |
| Ptprk | -1.179991 | 6.74032671 | 27.5237939 | 1.93E-06 | 1.32E-05 | T | T | F | T |
| Nova1 | -3.3619738 | 2.83075942 | 27.5145351 | 1.93E-06 | 1.32E-05 | T | T | T | T |
| Ap1s1 | 1.30153535 | 5.54571317 | 27.5114557 | 1.94E-06 | 1.32E-05 | F | F | F | F |
| Cebpa | 3.10218101 | 2.33159609 | 27.4989925 | 1.94E-06 | 1.33E-05 | F | F | T | F |
| Pard3b | -4.1177985 | 2.37437374 | 27.4779045 | 1.96E-06 | 1.34E-05 | T | F | T | T |
| Rab6b | 1.30528426 | 5.48366792 | 27.4738221 | 1.96E-06 | 1.34E-05 | T | F | T | F |
| Epb41l4b | -2.1460391 | 3.99384704 | 27.4486415 | 1.98E-06 | 1.35E-05 | F | F | T | F |
| Axl | -4.2233616 | 2.31513142 | 27.446073 | 1.98E-06 | 1.35E-05 | T | F | T | T |
| Hmgcll1 | -8.9375466 | 1.42019238 | 27.4458324 | 1.98E-06 | 1.35E-05 | T | F | T | T |
| Ppp2r5d | -1.0824153 | 7.41146066 | 27.4191523 | 2.00E-06 | 1.36E-05 | F | F | T | F |
| Stt3a | 0.97209545 | 7.7162367 | 27.4087943 | 2.01E-06 | 1.37E-05 | F | F | T | T |
| Klhl6 | -4.4693651 | 2.23975329 | 27.4040981 | 2.01E-06 | 1.37E-05 | F | F | F | F |
| Dclk3 | 4.08182076 | 1.57010353 | 27.3792671 | 2.03E-06 | 1.38E-05 | F | F | F | F |
| Ctnnd2 | -3.9063345 | 2.482227 | 27.3761791 | 2.03E-06 | 1.38E-05 | T | T | T | T |
| Aplp2 | -1.0195702 | 8.1169626 | 27.3754332 | 2.03E-06 | 1.38E-05 | F | F | F | F |
| Vps50 | 1.25572849 | 5.63489244 | 27.3588557 | 2.05E-06 | 1.39E-05 | T | F | F | F |

| | | | | | | | | | |
|---|---|---|---|---|---|---|---|---|---|
| Gpr108 | 1.5196648 | 4.63267692 | 27.3553787 | 2.05E-06 | 1.39E-05 | F | F | F | T |
| Map3k15 | -4.753353 | 2.11883501 | 27.333672 | 2.06E-06 | 1.40E-05 | T | F | F | T |
| Yes1 | 1.19267543 | 5.98941663 | 27.3265154 | 2.07E-06 | 1.41E-05 | T | F | F | T |
| Actr3b | -4.0463308 | 2.39904104 | 27.3259064 | 2.07E-06 | 1.41E-05 | T | F | F | F |
| Tpcn2 | 2.03623027 | 3.68291687 | 27.3212212 | 2.07E-06 | 1.41E-05 | F | F | F | T |
| Nek4 | -1.62362 | 4.8924628 | 27.2915555 | 2.10E-06 | 1.42E-05 | T | F | T | F |
| Itpkb | -2.7290104 | 3.3252634 | 27.2915517 | 2.10E-06 | 1.42E-05 | T | F | T | T |
| Gadd45g | 1.94735744 | 3.84251627 | 27.2738502 | 2.11E-06 | 1.43E-05 | F | F | T | F |
| Tfcp2l1 | -1.3802617 | 5.71581353 | 27.2650739 | 2.12E-06 | 1.43E-05 | T | F | T | T |
| Cib1 | 1.46793996 | 4.81215308 | 27.2114126 | 2.16E-06 | 1.46E-05 | F | F | F | F |
| Anxa1 | 4.22542004 | 1.57045911 | 27.2001141 | 2.17E-06 | 1.47E-05 | F | F | T | F |
| Hnrnpa0 | -0.9497321 | 9.01384126 | 27.1946447 | 2.17E-06 | 1.47E-05 | T | F | T | F |
| Bmp7 | -1.4986218 | 5.30763812 | 27.1588093 | 2.20E-06 | 1.49E-05 | F | F | T | F |
| Tex261 | 1.2279925 | 5.84151472 | 27.1569721 | 2.20E-06 | 1.49E-05 | T | F | F | F |
| Col11a1 | -8.9230791 | 1.41748702 | 27.1562174 | 2.20E-06 | 1.49E-05 | F | F | F | F |
| Slk | 1.04549733 | 6.97221768 | 27.1248052 | 2.23E-06 | 1.51E-05 | T | F | F | T |
| Slc25a32 | 1.6421802 | 4.28355848 | 27.1195432 | 2.23E-06 | 1.51E-05 | F | F | T | F |
| Med25 | -1.0952037 | 7.16087083 | 27.1131168 | 2.24E-06 | 1.51E-05 | T | F | T | T |
| Gnb2 | 0.97050595 | 7.78609473 | 27.1093741 | 2.24E-06 | 1.51E-05 | F | F | F | F |
| Ldb1 | -1.0937893 | 7.17456609 | 27.0995932 | 2.25E-06 | 1.52E-05 | T | F | F | F |
| Ap1s2 | -2.2228564 | 3.94813193 | 27.0871073 | 2.26E-06 | 1.52E-05 | T | T | T | T |
| Ube2l6 | -2.608234 | 3.43830298 | 27.0814225 | 2.26E-06 | 1.53E-05 | F | F | T | F |
| Ccdc28b | -4.2279382 | 2.30802207 | 27.0785407 | 2.26E-06 | 1.53E-05 | F | F | F | F |
| Erlin2 | 1.07398243 | 6.72254546 | 27.0432186 | 2.29E-06 | 1.55E-05 | T | T | T | T |
| Fbxl12 | 1.37764945 | 5.04725823 | 27.0404128 | 2.29E-06 | 1.55E-05 | F | F | F | F |
| Rab39b | -4.6989318 | 2.07134905 | 27.0126912 | 2.32E-06 | 1.56E-05 | T | F | T | T |
| Ccr1l1 | 8.81955033 | 0.25792456 | 27.0853857 | 2.33E-06 | 1.57E-05 | F | F | T | F |
| Il24 | 8.81955033 | 0.25792456 | 27.0853857 | 2.33E-06 | 1.57E-05 | F | F | F | F |
| Tll2 | -4.1950285 | 2.29221026 | 26.99757 | 2.33E-06 | 1.57E-05 | F | F | F | F |
| Mars2 | 1.42235525 | 4.90264649 | 26.977591 | 2.35E-06 | 1.58E-05 | T | F | T | T |
| Ecm2 | 8.81743547 | 0.25594564 | 27.0392781 | 2.37E-06 | 1.59E-05 | T | F | T | T |
| Penk | -8.9468356 | 1.4213255 | 26.9307797 | 2.39E-06 | 1.61E-05 | F | F | F | F |
| C130074G1 | -3.4816878 | 2.71109236 | 26.9159162 | 2.40E-06 | 1.62E-05 | F | F | T | F |
| Tcf21 | -8.9302445 | 1.42873211 | 26.9032675 | 2.41E-06 | 1.62E-05 | T | F | F | T |
| Tmem8 | -2.7698273 | 3.26038296 | 26.8746091 | 2.44E-06 | 1.64E-05 | F | F | F | T |
| Gm28635 | -10.080099 | 2.53298594 | 26.8638031 | 2.45E-06 | 1.64E-05 | F | F | F | F |
| Ckap5 | -0.9948826 | 8.29861165 | 26.8331339 | 2.47E-06 | 1.66E-05 | T | F | F | T |
| Slc25a40 | 2.34961159 | 3.11367577 | 26.816879 | 2.49E-06 | 1.67E-05 | F | F | F | F |
| Arhgap42 | -1.8868733 | 4.30823948 | 26.8117022 | 2.49E-06 | 1.67E-05 | F | F | T | T |
| Ugt8a | -9.0164121 | 1.48035994 | 26.8047491 | 2.50E-06 | 1.68E-05 | T | F | T | T |
| Zdhhc13 | 1.33988747 | 5.15500304 | 26.8021512 | 2.50E-06 | 1.68E-05 | F | F | F | F |
| Ky | -8.899112 | 1.38547253 | 26.8013383 | 2.50E-06 | 1.68E-05 | T | F | T | T |

| | | | | | | | | | |
|---|---|---|---|---|---|---|---|---|---|
| Sv2c | -8.924332 | 1.40230179 | 26.7915924 | 2.51E-06 | 1.68E-05 | F | F | T | T |
| Mif | -0.9630015 | 8.89891056 | 26.7895248 | 2.51E-06 | 1.69E-05 | F | F | F | F |
| Glyctk | -3.10872 | 2.97204915 | 26.758205 | 2.54E-06 | 1.70E-05 | T | F | T | T |
| Ncstn | 1.06801193 | 6.72289721 | 26.7575369 | 2.54E-06 | 1.70E-05 | F | F | F | F |
| Zfp260 | -1.4599539 | 5.36394937 | 26.7557531 | 2.54E-06 | 1.70E-05 | F | F | T | F |
| Rnf17 | 1.87330475 | 3.86309331 | 26.7477341 | 2.55E-06 | 1.71E-05 | T | F | F | F |
| Gpr27 | -4.3940944 | 2.17610429 | 26.7319105 | 2.57E-06 | 1.72E-05 | F | F | T | F |
| Lnpk | 1.72028231 | 4.24735242 | 26.71914 | 2.58E-06 | 1.73E-05 | T | F | T | F |
| Zfp968 | 5.08155173 | 1.0471172 | 26.715311 | 2.58E-06 | 1.73E-05 | F | F | F | F |
| Ccnf | 0.95406652 | 7.79216637 | 26.7140465 | 2.58E-06 | 1.73E-05 | T | F | T | F |
| Parp8 | -8.9064776 | 1.40408472 | 26.7086004 | 2.59E-06 | 1.73E-05 | T | T | T | T |
| Ankrd29 | 4.10548532 | 1.32997631 | 26.6864557 | 2.61E-06 | 1.74E-05 | T | F | T | F |
| Cby1 | -2.1904132 | 3.89542056 | 26.6808734 | 2.61E-06 | 1.75E-05 | F | F | T | F |
| Dcaf10 | 1.22797979 | 5.69840928 | 26.6525489 | 2.64E-06 | 1.76E-05 | T | F | T | T |
| Ar | -4.1793632 | 2.27509544 | 26.6495091 | 2.64E-06 | 1.77E-05 | F | T | T | T |
| Bcl2l11 | -1.5306852 | 5.1254138 | 26.6368518 | 2.66E-06 | 1.77E-05 | T | F | F | T |
| Bmp8b | 3.20214239 | 2.16918564 | 26.6040785 | 2.69E-06 | 1.79E-05 | T | F | T | T |
| Fundc2 | -2.5752397 | 3.43293891 | 26.5859645 | 2.71E-06 | 1.81E-05 | T | T | F | T |
| Kcnk4 | 2.98827074 | 2.27178484 | 26.5688227 | 2.72E-06 | 1.82E-05 | F | F | F | F |
| Fkbp3 | -1.1332361 | 6.86839351 | 26.5526591 | 2.74E-06 | 1.83E-05 | F | F | F | F |
| H3c8 | -0.9724665 | 8.76323213 | 26.5419686 | 2.75E-06 | 1.83E-05 | F | F | F | F |
| Degs1 | 1.04273515 | 6.90616075 | 26.5335976 | 2.76E-06 | 1.84E-05 | F | F | F | F |
| Gls | 1.13567085 | 6.28623086 | 26.5279467 | 2.76E-06 | 1.84E-05 | T | F | T | T |
| Lamp1 | 0.89434747 | 8.67496742 | 26.5110441 | 2.78E-06 | 1.85E-05 | T | F | T | T |
| Pink1 | 1.45759836 | 4.79873999 | 26.5010673 | 2.79E-06 | 1.86E-05 | T | F | F | F |
| Slc10a7 | 1.57156859 | 4.38548507 | 26.4769244 | 2.82E-06 | 1.87E-05 | F | F | T | F |
| Ccl27a | -8.9172439 | 1.39382692 | 26.4714545 | 2.82E-06 | 1.88E-05 | F | F | F | F |
| Chst15 | -1.7032459 | 4.6851331 | 26.4608815 | 2.83E-06 | 1.88E-05 | F | F | T | F |
| Gapdh | -0.8473398 | 10.7624086 | 26.4529357 | 2.84E-06 | 1.89E-05 | F | F | T | F |
| Abca9 | -5.8615097 | 1.75595103 | 26.4494112 | 2.84E-06 | 1.89E-05 | T | F | F | T |
| Sec14l1 | 1.05226448 | 6.87944421 | 26.4454449 | 2.85E-06 | 1.89E-05 | T | F | T | T |
| Dock6 | 1.10554239 | 6.51605872 | 26.4421642 | 2.85E-06 | 1.89E-05 | T | T | F | T |
| Ccdc194 | -8.8752758 | 1.36585072 | 26.4287664 | 2.87E-06 | 1.90E-05 | F | F | T | F |
| Glis1 | -2.4223241 | 3.61990459 | 26.4207497 | 2.87E-06 | 1.91E-05 | F | F | F | F |
| Gprc5a | 1.96064419 | 3.80638757 | 26.420596 | 2.87E-06 | 1.91E-05 | F | F | T | F |
| Cend1 | -8.8856842 | 1.36988565 | 26.419297 | 2.88E-06 | 1.91E-05 | T | F | F | F |
| Mppe1 | 2.21919395 | 3.2494018 | 26.4178135 | 2.88E-06 | 1.91E-05 | T | F | F | F |
| Glipr2 | 1.35374732 | 5.08651155 | 26.413816 | 2.88E-06 | 1.91E-05 | F | F | T | F |
| Rnf216 | -1.3223046 | 5.87147296 | 26.3974113 | 2.90E-06 | 1.92E-05 | F | F | T | F |
| Igf2bp3 | -1.0275408 | 7.73688585 | 26.3801463 | 2.92E-06 | 1.93E-05 | T | F | T | T |
| Id4 | -5.8217214 | 1.70455705 | 26.3661326 | 2.93E-06 | 1.94E-05 | F | F | T | F |
| Rhbdd1 | 1.75763476 | 4.02601057 | 26.3634059 | 2.93E-06 | 1.94E-05 | T | F | T | F |

| | | | | | | | | | |
|---|---|---|---|---|---|---|---|---|---|
| Arsa | -3.4866958 | 2.70424377 | 26.3440913 | 2.96E-06 | 1.96E-05 | F | F | T | T |
| Fgf5 | -3.9004394 | 2.78787558 | 26.3062247 | 3.00E-06 | 1.98E-05 | T | F | T | F |
| Aoc2 | 1.82536639 | 3.95037876 | 26.3009725 | 3.00E-06 | 1.99E-05 | F | F | F | F |
| mt-Nd4 | 0.83168656 | 10.043426 | 26.2898473 | 3.01E-06 | 1.99E-05 | F | F | F | F |
| Tceal5 | -8.870212 | 1.35856646 | 26.2852621 | 3.02E-06 | 2.00E-05 | F | F | T | F |
| Rtn4r | 2.46017279 | 2.92161209 | 26.2732474 | 3.03E-06 | 2.01E-05 | F | F | F | F |
| Nagk | 2.442042 | 2.95742821 | 26.2684448 | 3.04E-06 | 2.01E-05 | F | F | F | F |
| Fdps | 1.13533794 | 6.20657038 | 26.2582519 | 3.05E-06 | 2.02E-05 | F | F | F | F |
| Aldh3a2 | 1.30931618 | 5.22068396 | 26.2517308 | 3.06E-06 | 2.02E-05 | T | F | F | F |
| Scap | 1.00690416 | 7.07459119 | 26.2211261 | 3.09E-06 | 2.04E-05 | F | F | F | F |
| Tbc1d9b | 1.12561087 | 6.38416608 | 26.2157112 | 3.10E-06 | 2.04E-05 | T | F | F | T |
| Slc16a10 | -1.8272885 | 4.3353073 | 26.2101958 | 3.10E-06 | 2.05E-05 | F | F | T | T |
| Zfc3h1 | 1.04835758 | 6.98446164 | 26.1988465 | 3.12E-06 | 2.06E-05 | F | F | F | F |
| Asah1 | 1.32383751 | 5.20828227 | 26.184578 | 3.13E-06 | 2.07E-05 | F | F | T | F |
| Tmem109 | 1.301666 | 5.29572833 | 26.1622268 | 3.16E-06 | 2.08E-05 | T | F | T | T |
| Zkscan3 | -1.3855081 | 5.62638119 | 26.1543137 | 3.17E-06 | 2.09E-05 | T | F | F | F |
| Lgmn | -1.0149851 | 7.84031348 | 26.1230363 | 3.20E-06 | 2.11E-05 | F | F | F | F |
| Add2 | -8.893026 | 1.37159764 | 26.1210828 | 3.21E-06 | 2.11E-05 | T | F | T | T |
| Kdm3a | -1.3651012 | 5.70162749 | 26.1134407 | 3.22E-06 | 2.12E-05 | T | F | T | T |
| Zmpste24 | 1.12843062 | 6.23759543 | 26.1115111 | 3.22E-06 | 2.12E-05 | F | F | T | F |
| Klk13 | -8.8634149 | 1.35074596 | 26.1108724 | 3.22E-06 | 2.12E-05 | F | F | F | T |
| Pm20d2 | -4.3742297 | 2.1529044 | 26.1102395 | 3.22E-06 | 2.12E-05 | F | F | T | T |
| Pnpt1 | 1.18404402 | 5.89895805 | 26.0925062 | 3.24E-06 | 2.13E-05 | F | F | F | F |
| Kctd15 | 1.2233914 | 5.70779402 | 26.0921377 | 3.24E-06 | 2.13E-05 | F | F | T | T |
| Arhgap17 | 1.13374279 | 6.19364126 | 26.0907348 | 3.24E-06 | 2.13E-05 | T | F | F | T |
| Pbxip1 | 1.38118783 | 4.97374855 | 26.0758955 | 3.26E-06 | 2.14E-05 | F | F | T | T |
| Alas1 | 1.54981326 | 4.42225815 | 26.0313671 | 3.31E-06 | 2.18E-05 | F | F | F | F |
| Mesp2 | -5.7977052 | 1.68447161 | 26.0272355 | 3.32E-06 | 2.18E-05 | F | F | T | F |
| Aamdc | -8.8586586 | 1.3571652 | 25.9941285 | 3.36E-06 | 2.21E-05 | F | F | F | T |
| Vps13b | 0.97857258 | 7.31162835 | 25.9866166 | 3.37E-06 | 2.21E-05 | T | F | T | F |
| Xlr5b | 2.39178052 | 3.02666546 | 25.9701534 | 3.39E-06 | 2.22E-05 | F | F | F | F |
| Steap2 | -2.2663767 | 3.76402279 | 25.9425198 | 3.42E-06 | 2.25E-05 | T | F | T | T |
| Dcaf11 | -1.5799404 | 4.93378779 | 25.9407647 | 3.43E-06 | 2.25E-05 | F | F | F | F |
| Car9 | 8.76718756 | 0.20901643 | 25.9627148 | 3.50E-06 | 2.29E-05 | F | F | F | F |
| 2510039O1 | 1.11264917 | 6.34029839 | 25.8828995 | 3.50E-06 | 2.29E-05 | T | F | T | F |
| Poglut2 | -1.5477856 | 5.00415112 | 25.8801916 | 3.50E-06 | 2.29E-05 | F | F | T | F |
| Cacna1c | -4.1849219 | 2.28378867 | 25.8245568 | 3.57E-06 | 2.34E-05 | F | F | T | T |
| Bnc1 | -8.8417231 | 1.33417071 | 25.8229482 | 3.58E-06 | 2.34E-05 | T | F | F | F |
| Sipa1l3 | -1.236482 | 6.22263603 | 25.8190644 | 3.58E-06 | 2.34E-05 | T | F | T | T |
| Chmp2a | 1.27973128 | 5.3541516 | 25.8122489 | 3.59E-06 | 2.35E-05 | F | F | F | F |
| Il11ra1 | 1.82347458 | 3.92617568 | 25.7922529 | 3.62E-06 | 2.37E-05 | F | F | F | F |
| Zfp335 | 1.12782583 | 6.19352022 | 25.7751374 | 3.64E-06 | 2.38E-05 | F | F | F | F |

| | | | | | | | | | |
|---|---|---|---|---|---|---|---|---|---|
| Hk2 | 0.88655106 | 8.62090621 | 25.745026 | 3.68E-06 | 2.41E-05 | T | F | T | T |
| Dok1 | -8.8386681 | 1.32933625 | 25.7316215 | 3.70E-06 | 2.42E-05 | F | F | T | F |
| Thbs4 | -8.8354699 | 1.32941683 | 25.7191167 | 3.72E-06 | 2.43E-05 | F | F | F | F |
| Creg1 | 1.89147389 | 3.78825664 | 25.7021156 | 3.74E-06 | 2.44E-05 | F | F | F | F |
| Rabggta | 1.27612923 | 5.31378652 | 25.6890586 | 3.76E-06 | 2.45E-05 | F | F | T | F |
| Vtn | -8.9368202 | 1.44249992 | 25.6639484 | 3.79E-06 | 2.48E-05 | F | F | F | F |
| Tmem238 | -3.7899851 | 2.43783701 | 25.6611404 | 3.80E-06 | 2.48E-05 | T | F | F | T |
| H2ac7 | -0.9907379 | 8.00652518 | 25.6610283 | 3.80E-06 | 2.48E-05 | F | F | F | F |
| Myo15b | -3.1922852 | 2.83680041 | 25.6577397 | 3.80E-06 | 2.48E-05 | T | F | F | F |
| Zscan12 | -2.1068627 | 3.93688193 | 25.654827 | 3.80E-06 | 2.48E-05 | T | F | T | F |
| Pik3r3 | -1.6791778 | 4.62956295 | 25.6348897 | 3.83E-06 | 2.50E-05 | F | F | T | F |
| Usp40 | -1.6015695 | 4.80509694 | 25.6333387 | 3.83E-06 | 2.50E-05 | T | F | F | F |
| Cdh13 | -8.8310849 | 1.32390947 | 25.630171 | 3.84E-06 | 2.50E-05 | T | F | T | F |
| Pcdhga2 | -8.8359383 | 1.32478153 | 25.6160997 | 3.86E-06 | 2.51E-05 | F | F | F | F |
| Tagln3 | -8.8421071 | 1.32821758 | 25.6058921 | 3.87E-06 | 2.52E-05 | F | F | F | F |
| Plcl2 | -1.8806495 | 4.27073579 | 25.575543 | 3.92E-06 | 2.55E-05 | F | F | F | T |
| Insm1 | -5.9233425 | 1.79307663 | 25.5506153 | 3.95E-06 | 2.57E-05 | T | F | F | T |
| Ube3c | 0.96379989 | 7.35845507 | 25.5442864 | 3.96E-06 | 2.58E-05 | T | F | T | F |
| 3110001I22 | -8.8340717 | 1.32199457 | 25.5367551 | 3.97E-06 | 2.58E-05 | F | F | F | F |
| Myocd | -4.036081 | 2.39814814 | 25.52411 | 3.99E-06 | 2.60E-05 | T | F | T | T |
| Tgfbr2 | 1.72320079 | 4.04394411 | 25.4979804 | 4.03E-06 | 2.62E-05 | F | F | T | F |
| Tunar | -8.8553386 | 1.33622509 | 25.4878842 | 4.05E-06 | 2.63E-05 | F | F | F | F |
| Ifih1 | 3.00295035 | 2.20078165 | 25.4873849 | 4.05E-06 | 2.63E-05 | T | T | T | T |
| Tal2 | -8.8389967 | 1.34148877 | 25.4811305 | 4.06E-06 | 2.63E-05 | F | F | T | T |
| Ephb1 | -1.8160482 | 4.30787565 | 25.4764137 | 4.06E-06 | 2.64E-05 | F | F | F | F |
| Slc27a3 | -1.5674275 | 4.87810764 | 25.4753617 | 4.06E-06 | 2.64E-05 | F | F | F | F |
| Decr2 | 1.64671805 | 4.23705596 | 25.4701477 | 4.07E-06 | 2.64E-05 | F | F | F | F |
| Msh3 | -1.4592065 | 5.18864175 | 25.3909052 | 4.19E-06 | 2.72E-05 | T | F | F | F |
| Scn8a | -2.5408255 | 3.38586867 | 25.3631508 | 4.24E-06 | 2.75E-05 | T | F | T | F |
| Gpsm1 | -1.3249349 | 5.74548997 | 25.3464529 | 4.26E-06 | 2.76E-05 | F | F | T | F |
| Edn3 | -8.8199454 | 1.32028939 | 25.3435181 | 4.27E-06 | 2.77E-05 | F | F | T | F |
| Nt5dc3 | 1.3675132 | 4.92451036 | 25.3133266 | 4.32E-06 | 2.80E-05 | T | F | T | T |
| Xkr8 | 1.95999667 | 3.65570304 | 25.2791995 | 4.37E-06 | 2.83E-05 | T | F | T | F |
| Rhbdl3 | -3.4177784 | 2.733969 | 25.2654968 | 4.39E-06 | 2.84E-05 | F | F | T | F |
| Cenpt | 1.39251077 | 4.86811397 | 25.2476093 | 4.42E-06 | 2.86E-05 | F | F | F | F |
| Kcnj15 | -4.9535879 | 1.96399475 | 25.2258082 | 4.46E-06 | 2.88E-05 | F | F | F | T |
| Rcor3 | -3.9248519 | 2.33459025 | 25.2229608 | 4.46E-06 | 2.89E-05 | T | F | T | T |
| Slc29a4 | -2.779744 | 3.15318194 | 25.218204 | 4.47E-06 | 2.89E-05 | F | F | F | F |
| Fads6 | 2.64886332 | 2.63394616 | 25.2016837 | 4.50E-06 | 2.91E-05 | F | F | F | T |
| Dld | 1.07651694 | 6.44882622 | 25.1943506 | 4.51E-06 | 2.91E-05 | F | F | F | T |
| Elavl3 | -2.2399531 | 4.07807 | 25.1884342 | 4.52E-06 | 2.92E-05 | F | F | T | F |
| Gfra1 | -8.8051219 | 1.30222078 | 25.1810602 | 4.53E-06 | 2.93E-05 | T | F | T | T |

| | | | | | | | | | |
|---|---|---|---|---|---|---|---|---|---|
| Cic | -1.0794037 | 7.01705294 | 25.1588255 | 4.57E-06 | 2.95E-05 | F | F | F | F |
| Socs3 | 1.48326572 | 4.60275862 | 25.1557516 | 4.57E-06 | 2.95E-05 | F | F | T | F |
| Unc119b | -1.2282336 | 6.23629055 | 25.1444616 | 4.59E-06 | 2.96E-05 | T | F | T | F |
| Haus7 | -1.6497906 | 4.64591623 | 25.1324582 | 4.61E-06 | 2.98E-05 | T | F | T | T |
| Ddx58 | 1.96353382 | 3.59673097 | 25.1266037 | 4.62E-06 | 2.98E-05 | F | F | F | F |
| Tmem121b | -8.8019222 | 1.29980877 | 25.1201549 | 4.64E-06 | 2.99E-05 | T | F | T | F |
| Surf4 | 0.90405707 | 8.24824962 | 25.0838327 | 4.70E-06 | 3.03E-05 | T | F | F | T |
| Gm11992 | 3.87392582 | 1.43915588 | 25.0708579 | 4.72E-06 | 3.04E-05 | F | F | F | F |
| Tns2 | 1.36668271 | 4.93066615 | 25.041375 | 4.77E-06 | 3.07E-05 | F | F | F | F |
| Sumo2 | -0.9957617 | 7.90582466 | 25.022491 | 4.81E-06 | 3.09E-05 | T | F | F | F |
| Lnx2 | 1.17206713 | 5.79395308 | 25.016821 | 4.82E-06 | 3.10E-05 | T | F | F | T |
| Fam71f1 | 8.7246108 | 0.16938808 | 25.0787785 | 4.84E-06 | 3.11E-05 | F | F | F | F |
| Cyp51 | 1.22690189 | 5.48703717 | 25.0034112 | 4.84E-06 | 3.11E-05 | T | F | T | T |
| Ralgps2 | -1.2742838 | 6.28619904 | 24.9856684 | 4.87E-06 | 3.13E-05 | T | F | T | T |
| Elmo1 | -1.4504875 | 5.33808581 | 24.9796069 | 4.88E-06 | 3.14E-05 | T | F | T | F |
| Htatsf1 | 1.0231436 | 6.79801209 | 24.970997 | 4.90E-06 | 3.15E-05 | T | F | F | F |
| Tmem123 | 1.15726429 | 5.88957794 | 24.9632022 | 4.91E-06 | 3.16E-05 | T | F | F | T |
| Spef1 | -8.7937486 | 1.28676183 | 24.9304817 | 4.97E-06 | 3.19E-05 | T | F | F | F |
| Tmem185b | 1.5130371 | 4.45816729 | 24.9267145 | 4.98E-06 | 3.20E-05 | F | F | T | F |
| Ankrd37 | 2.77706765 | 2.49950419 | 24.9261353 | 4.98E-06 | 3.20E-05 | F | F | F | F |
| Zfp664 | -1.0304732 | 7.42262924 | 24.9245181 | 4.98E-06 | 3.20E-05 | T | F | T | T |
| Agap3 | -1.1844923 | 6.37761078 | 24.9138182 | 5.00E-06 | 3.21E-05 | F | F | F | F |
| Klkb1 | 1.60532041 | 4.32291346 | 24.9017737 | 5.03E-06 | 3.22E-05 | F | F | T | F |
| Hoxb8 | -6.5150884 | 2.38572606 | 24.8985316 | 5.03E-06 | 3.23E-05 | F | F | F | F |
| Kdm7a | -1.5190939 | 4.96196948 | 24.8852004 | 5.06E-06 | 3.24E-05 | T | F | T | T |
| Rasl11b | -3.0471752 | 2.9163838 | 24.8466025 | 5.13E-06 | 3.29E-05 | F | F | T | F |
| Rbm44 | 2.90387594 | 2.24386449 | 24.8442381 | 5.13E-06 | 3.29E-05 | F | F | F | F |
| Mynn | 1.68226207 | 4.22496634 | 24.8134846 | 5.19E-06 | 3.33E-05 | T | F | F | T |
| Adamts17 | -3.0116483 | 2.97205076 | 24.8059338 | 5.21E-06 | 3.34E-05 | T | F | T | T |
| Bcl11a | -4.0088119 | 2.57174581 | 24.8035192 | 5.21E-06 | 3.34E-05 | T | F | T | F |
| Eid1 | -1.386472 | 5.36484188 | 24.7939645 | 5.23E-06 | 3.35E-05 | T | F | F | F |
| Gm14295 | 3.35551261 | 2.55821638 | 24.780271 | 5.26E-06 | 3.36E-05 | F | F | T | T |
| Nsg1 | -4.9945288 | 1.81520274 | 24.7773994 | 5.26E-06 | 3.37E-05 | T | F | T | F |
| Aldh18a1 | 0.94984282 | 7.43847578 | 24.764362 | 5.29E-06 | 3.38E-05 | T | F | T | T |
| Lrrc1 | -1.6063089 | 4.73545363 | 24.7604047 | 5.30E-06 | 3.39E-05 | F | F | F | F |
| Dnajb2 | 1.78695968 | 3.88801181 | 24.7444582 | 5.33E-06 | 3.41E-05 | F | F | T | F |
| Rtl6 | -1.6652064 | 4.5743519 | 24.743603 | 5.33E-06 | 3.41E-05 | T | F | T | F |
| Nkx2-5 | -8.9546553 | 1.46290089 | 24.7399519 | 5.34E-06 | 3.41E-05 | F | F | T | T |
| Klhl18 | 1.35138597 | 4.93132706 | 24.7370367 | 5.34E-06 | 3.41E-05 | T | F | T | T |
| Zmat3 | -1.1599055 | 6.5201812 | 24.7343901 | 5.35E-06 | 3.41E-05 | T | F | T | T |
| Nos1 | 2.08498104 | 3.33795996 | 24.733667 | 5.35E-06 | 3.41E-05 | T | F | F | F |
| Atp9a | 1.25051644 | 5.30552972 | 24.7257083 | 5.37E-06 | 3.42E-05 | F | F | F | T |

| | | | | | | | | | |
|---|---|---|---|---|---|---|---|---|---|
| Pawr | 1.15728299 | 5.85312798 | 24.7133494 | 5.39E-06 | 3.44E-05 | T | F | F | F |
| Casp7 | 1.65824619 | 4.10851056 | 24.713309 | 5.39E-06 | 3.44E-05 | F | F | F | F |
| Capza1 | 0.98684761 | 7.38750348 | 24.6856347 | 5.45E-06 | 3.47E-05 | T | F | T | F |
| Robo2 | -3.1178842 | 2.98415754 | 24.6841753 | 5.45E-06 | 3.47E-05 | T | F | T | F |
| Smim3 | -5.7380825 | 1.6368943 | 24.6134092 | 5.60E-06 | 3.56E-05 | T | F | F | F |
| Actn1 | 0.86978792 | 8.68095932 | 24.6125484 | 5.60E-06 | 3.56E-05 | T | F | F | T |
| Clk2 | -1.4325903 | 5.17482194 | 24.6123482 | 5.60E-06 | 3.56E-05 | F | F | F | F |
| Pdzd8 | 0.95550144 | 7.3486372 | 24.6104228 | 5.60E-06 | 3.57E-05 | T | F | T | F |
| Ddx19b | -1.3181554 | 5.63455441 | 24.5967377 | 5.63E-06 | 3.58E-05 | T | F | T | F |
| Zfp119a | -8.7738275 | 1.26794059 | 24.5820976 | 5.66E-06 | 3.60E-05 | F | F | T | F |
| Gpr39 | -8.7744697 | 1.2680651 | 24.5791873 | 5.67E-06 | 3.60E-05 | T | F | F | T |
| Foxj2 | 1.41858015 | 4.68482205 | 24.5755031 | 5.68E-06 | 3.61E-05 | T | F | F | T |
| F8 | -8.7698855 | 1.26912867 | 24.57068 | 5.69E-06 | 3.61E-05 | F | F | F | T |
| Enox2 | -3.1744153 | 2.7874503 | 24.5554327 | 5.72E-06 | 3.63E-05 | F | F | T | F |
| Dmgdh | -8.7689046 | 1.26613823 | 24.5491263 | 5.73E-06 | 3.64E-05 | F | F | T | T |
| Sfr1 | -1.1339482 | 6.64819496 | 24.5447202 | 5.74E-06 | 3.65E-05 | T | F | F | T |
| Tpra1 | -2.2520982 | 3.67722236 | 24.5422669 | 5.75E-06 | 3.65E-05 | T | F | F | T |
| Sord | -1.5349613 | 5.01048884 | 24.5355754 | 5.76E-06 | 3.66E-05 | T | F | F | F |
| Scn5a | -4.4844768 | 2.2479389 | 24.5244371 | 5.78E-06 | 3.67E-05 | F | F | T | T |
| Def8 | -1.3072118 | 5.68606105 | 24.5178786 | 5.80E-06 | 3.68E-05 | F | F | T | F |
| Shf | -1.8103049 | 4.36901372 | 24.5106708 | 5.81E-06 | 3.69E-05 | F | F | F | F |
| Bdp1 | 0.94736728 | 7.45996189 | 24.5100439 | 5.82E-06 | 3.69E-05 | T | F | T | T |
| Hoxb9 | -9.2663689 | 1.75966735 | 24.5076797 | 5.82E-06 | 3.69E-05 | T | F | T | T |
| Arl8b | 1.01329535 | 6.79554239 | 24.5004382 | 5.84E-06 | 3.70E-05 | F | F | T | F |
| Trim37 | -1.0878632 | 6.8711711 | 24.4955935 | 5.85E-06 | 3.70E-05 | T | F | T | F |
| Sema6d | 1.42103964 | 5.43355844 | 24.494215 | 5.85E-06 | 3.70E-05 | T | F | F | T |
| Cacna1i | -8.7675937 | 1.26965709 | 24.4941473 | 5.85E-06 | 3.70E-05 | T | F | T | T |
| Plppr1 | -8.7729472 | 1.27745294 | 24.4902799 | 5.86E-06 | 3.71E-05 | T | F | T | T |
| Rcn3 | -1.3662088 | 5.54718536 | 24.4872556 | 5.87E-06 | 3.71E-05 | T | F | F | F |
| Specc1l | -1.1417734 | 6.55405428 | 24.4867933 | 5.87E-06 | 3.71E-05 | F | F | F | F |
| Apex1 | -1.0476493 | 7.10021882 | 24.4863759 | 5.87E-06 | 3.71E-05 | F | F | F | F |
| Prmt2 | 1.38465397 | 4.7982364 | 24.4799642 | 5.88E-06 | 3.72E-05 | F | F | F | F |
| Tfec | -8.7648077 | 1.26190467 | 24.4745776 | 5.89E-06 | 3.72E-05 | T | F | F | T |
| Tspan9 | -1.3383534 | 5.77748734 | 24.4725383 | 5.90E-06 | 3.73E-05 | T | F | F | T |
| Adra2b | 1.68677554 | 4.05408702 | 24.4664148 | 5.91E-06 | 3.73E-05 | T | F | F | T |
| Tgfbi | -4.2890774 | 2.07917223 | 24.4490446 | 5.95E-06 | 3.76E-05 | F | F | F | F |
| Tnfsf12 | -8.7629733 | 1.26319858 | 24.4487159 | 5.95E-06 | 3.76E-05 | F | F | F | F |
| Dnajc1 | 1.39793404 | 4.79272523 | 24.4392912 | 5.97E-06 | 3.77E-05 | T | F | T | T |
| Pak2 | 0.91137651 | 7.78522011 | 24.4347486 | 5.98E-06 | 3.77E-05 | T | F | T | F |
| Nlrx1 | 2.65461527 | 2.54205178 | 24.4277129 | 6.00E-06 | 3.78E-05 | T | F | F | F |
| Zfp354a | -3.3117373 | 2.70565046 | 24.4268137 | 6.00E-06 | 3.78E-05 | T | F | T | T |
| Lmo1 | -4.9897873 | 1.81687186 | 24.4144292 | 6.03E-06 | 3.80E-05 | F | F | F | F |

| | | | | | | | | | |
|---|---|---|---|---|---|---|---|---|---|
| Pask | -1.2973745 | 5.71583335 | 24.3966549 | 6.07E-06 | 3.82E-05 | F | F | F | F |
| Timp3 | -1.9970203 | 4.0685322 | 24.3886995 | 6.09E-06 | 3.83E-05 | T | F | T | T |
| Cipc | -1.435973 | 5.14471803 | 24.3866424 | 6.09E-06 | 3.83E-05 | T | F | T | T |
| Vwce | -3.8549071 | 2.29911473 | 24.3673662 | 6.13E-06 | 3.86E-05 | F | F | T | F |
| Adam9 | 1.18770227 | 5.61512412 | 24.3603974 | 6.15E-06 | 3.87E-05 | T | F | T | F |
| Nans | 1.04762073 | 6.58400335 | 24.3598221 | 6.15E-06 | 3.87E-05 | F | F | F | F |
| Ptcd1 | 1.19058864 | 5.59441306 | 24.359045 | 6.15E-06 | 3.87E-05 | F | F | T | F |
| Clec2d | 8.69152324 | 0.13868085 | 24.4096077 | 6.20E-06 | 3.90E-05 | F | F | T | T |
| Dennd6b | -9.0239468 | 1.48025685 | 24.3379582 | 6.20E-06 | 3.90E-05 | T | F | T | F |
| Gm29776 | -5.7215025 | 1.60796438 | 24.3364046 | 6.21E-06 | 3.90E-05 | F | F | T | F |
| Srms | 5.55638444 | 0.69714943 | 24.3184166 | 6.25E-06 | 3.92E-05 | F | F | F | F |
| Calb2 | -8.7896987 | 1.29924135 | 24.3180304 | 6.25E-06 | 3.92E-05 | T | F | F | T |
| Sapcd2 | -2.1979185 | 3.73541979 | 24.3042316 | 6.28E-06 | 3.94E-05 | T | F | T | T |
| Kif12 | -5.70936 | 1.60696789 | 24.3019852 | 6.29E-06 | 3.95E-05 | F | F | F | F |
| Slc2a5 | -5.6044555 | 1.62002363 | 24.2926678 | 6.31E-06 | 3.96E-05 | F | F | F | F |
| Cited2 | 0.92115572 | 7.62053025 | 24.2831107 | 6.33E-06 | 3.97E-05 | F | F | T | F |
| Spcs3 | 1.01524428 | 6.73266435 | 24.2816924 | 6.33E-06 | 3.97E-05 | T | F | T | F |
| Mknk1 | 1.77567559 | 3.86410688 | 24.2758659 | 6.35E-06 | 3.98E-05 | F | F | F | F |
| Senp5 | 1.05621604 | 6.45745779 | 24.2706668 | 6.36E-06 | 3.99E-05 | T | F | T | T |
| Akap12 | 0.91067859 | 7.75394094 | 24.2615307 | 6.38E-06 | 4.00E-05 | F | F | F | F |
| Dctpp1 | -1.1997407 | 6.27736637 | 24.2346899 | 6.45E-06 | 4.04E-05 | F | F | F | F |
| Galnt14 | -8.8711455 | 1.34104142 | 24.2296535 | 6.46E-06 | 4.04E-05 | T | F | F | T |
| Unc45a | 1.03738388 | 6.57904277 | 24.227798 | 6.46E-06 | 4.05E-05 | F | F | T | F |
| Syndig1 | -5.6995297 | 1.59340726 | 24.2072206 | 6.51E-06 | 4.08E-05 | T | F | F | F |
| Acp6 | -2.3111383 | 3.60909005 | 24.1950872 | 6.54E-06 | 4.09E-05 | T | F | F | F |
| Rin2 | -2.2772835 | 3.70943142 | 24.1923232 | 6.55E-06 | 4.10E-05 | F | F | F | F |
| Zfand2a | 1.46366485 | 4.50812547 | 24.1883263 | 6.56E-06 | 4.10E-05 | F | F | T | F |
| Slc16a9 | -2.3912992 | 3.47652373 | 24.1592302 | 6.63E-06 | 4.15E-05 | F | F | F | F |
| Nkd2 | -4.9811246 | 1.80986612 | 24.1222356 | 6.72E-06 | 4.20E-05 | F | F | T | F |
| Rida | -2.0958793 | 3.85020664 | 24.0910671 | 6.80E-06 | 4.25E-05 | F | F | F | F |
| Arhgap26 | -3.31237 | 2.65729438 | 24.0563672 | 6.89E-06 | 4.30E-05 | T | F | T | T |
| Tppp3 | -3.2914691 | 2.65417697 | 24.0517677 | 6.90E-06 | 4.31E-05 | F | F | T | F |
| Pdlim5 | 1.07195049 | 6.34620599 | 24.0425568 | 6.93E-06 | 4.32E-05 | F | T | T | T |
| Ddhd2 | 1.17869676 | 5.70914486 | 24.0094369 | 7.02E-06 | 4.38E-05 | F | F | T | F |
| Ap4e1 | 1.27522184 | 5.18296498 | 23.9944242 | 7.06E-06 | 4.40E-05 | T | F | F | T |
| Dtx4 | -1.587304 | 4.75743575 | 23.9765951 | 7.10E-06 | 4.43E-05 | T | F | T | T |
| Dync1h1 | 0.76335913 | 10.3395448 | 23.9599969 | 7.15E-06 | 4.46E-05 | F | F | F | F |
| Btbd17 | -5.6864826 | 1.58368401 | 23.9508366 | 7.17E-06 | 4.47E-05 | F | F | T | F |
| Nmt2 | 0.95899458 | 7.08134812 | 23.9484905 | 7.18E-06 | 4.47E-05 | F | F | T | T |
| Amot | -0.860264 | 9.47992692 | 23.9022821 | 7.30E-06 | 4.55E-05 | T | F | T | T |
| Dsc3 | 5.95163018 | 0.54115667 | 23.8922525 | 7.33E-06 | 4.57E-05 | F | F | T | F |
| Fam83d | -1.5988658 | 4.68247184 | 23.8916189 | 7.33E-06 | 4.57E-05 | F | F | F | F |

| | | | | | | | | | |
|---|---|---|---|---|---|---|---|---|---|
| Oprd1 | -8.7317351 | 1.2289981 | 23.8778711 | 7.37E-06 | 4.59E-05 | T | F | T | F |
| Dstn | 0.89278653 | 7.95478754 | 23.8765197 | 7.38E-06 | 4.59E-05 | T | F | F | F |
| Xrcc1 | -1.1773268 | 6.27807861 | 23.8712536 | 7.39E-06 | 4.60E-05 | F | F | F | F |
| Dnajb12 | 1.04707811 | 6.49456461 | 23.8588124 | 7.42E-06 | 4.62E-05 | F | F | T | T |
| Morn4 | -2.3382787 | 3.53143263 | 23.8526182 | 7.44E-06 | 4.63E-05 | F | F | T | F |
| Apoc1 | -2.8751135 | 2.98570708 | 23.8268702 | 7.51E-06 | 4.67E-05 | F | F | F | F |
| Vcpip1 | 1.0184068 | 6.70805812 | 23.7912158 | 7.62E-06 | 4.73E-05 | T | F | T | T |
| Prkab1 | -1.4197721 | 5.18302959 | 23.7899825 | 7.62E-06 | 4.73E-05 | F | F | T | F |
| Igsf3 | 1.02713845 | 6.77770136 | 23.7881191 | 7.62E-06 | 4.73E-05 | T | F | T | T |
| Pias2 | -1.4919025 | 5.06918597 | 23.7679404 | 7.68E-06 | 4.77E-05 | T | F | T | T |
| S100g | -4.1061887 | 2.21539137 | 23.7677068 | 7.68E-06 | 4.77E-05 | F | F | F | F |
| Gm10324 | 6.67168814 | 0.33556578 | 23.7533255 | 7.73E-06 | 4.79E-05 | F | F | F | F |
| Tbc1d13 | -1.1314913 | 6.50395109 | 23.7236211 | 7.81E-06 | 4.84E-05 | F | F | F | F |
| Klhdc2 | -1.1569684 | 6.3681438 | 23.7232544 | 7.81E-06 | 4.84E-05 | T | F | T | F |
| 1600014C1( | 1.80813376 | 3.77044967 | 23.7007076 | 7.88E-06 | 4.88E-05 | T | F | F | F |
| Maml2 | -3.5222231 | 2.46969801 | 23.6983062 | 7.89E-06 | 4.89E-05 | T | F | T | T |
| Mtpn | 0.91100708 | 7.61302568 | 23.68591 | 7.92E-06 | 4.91E-05 | T | F | F | T |
| Btrc | -1.4058723 | 5.19652161 | 23.6817692 | 7.94E-06 | 4.92E-05 | T | F | T | F |
| Fbxl14 | -1.0952704 | 6.70157947 | 23.6730546 | 7.96E-06 | 4.93E-05 | T | F | F | T |
| Dync2h1 | -1.1649384 | 6.31347642 | 23.6625986 | 7.99E-06 | 4.95E-05 | F | F | T | F |
| Brinp1 | -8.838239 | 1.30945713 | 23.6539805 | 8.02E-06 | 4.96E-05 | F | F | F | F |
| Nefh | -5.7213082 | 1.60202504 | 23.6438644 | 8.05E-06 | 4.98E-05 | F | F | F | F |
| Cyfip2 | -1.3221228 | 5.53380328 | 23.61738 | 8.13E-06 | 5.03E-05 | T | F | T | F |
| Ephb3 | -1.2620836 | 5.76260298 | 23.5741814 | 8.27E-06 | 5.11E-05 | F | F | F | F |
| Maco1 | 1.08046678 | 6.18239647 | 23.5695218 | 8.28E-06 | 5.12E-05 | F | F | F | F |
| Vstm2l | -8.7135182 | 1.22022422 | 23.559738 | 8.31E-06 | 5.14E-05 | T | F | T | T |
| Cds1 | 2.12698855 | 3.27183304 | 23.5357944 | 8.39E-06 | 5.18E-05 | T | F | T | F |
| Zfp184 | -2.1408818 | 3.80248629 | 23.5355053 | 8.39E-06 | 5.18E-05 | F | F | F | F |
| Blcap | 1.2840402 | 5.08910679 | 23.5265899 | 8.42E-06 | 5.20E-05 | T | F | T | F |
| Gnpat | -1.05322 | 6.92022052 | 23.5241331 | 8.42E-06 | 5.20E-05 | F | F | T | T |
| Csrnp2 | -1.6157403 | 4.58975095 | 23.4830896 | 8.56E-06 | 5.28E-05 | T | F | T | T |
| Mitf | 2.50798855 | 2.65197383 | 23.4619553 | 8.62E-06 | 5.32E-05 | T | F | T | T |
| Cdk2ap2 | 1.6392746 | 4.0789852 | 23.4379492 | 8.70E-06 | 5.37E-05 | T | T | F | T |
| Cldn9 | -8.7083332 | 1.20423679 | 23.3986527 | 8.83E-06 | 5.45E-05 | F | F | F | F |
| Kif5a | -2.2508234 | 3.6254031 | 23.3971572 | 8.84E-06 | 5.45E-05 | T | F | T | F |
| Myl9 | -3.5153529 | 2.46500782 | 23.3931194 | 8.85E-06 | 5.45E-05 | F | F | F | T |
| Plpp5 | 1.93892318 | 3.52274398 | 23.387397 | 8.87E-06 | 5.46E-05 | F | F | T | F |
| Man1b1 | 0.96970743 | 6.95532022 | 23.3737285 | 8.92E-06 | 5.49E-05 | T | F | T | T |
| Ngfr | -3.8743798 | 2.23028564 | 23.3602377 | 8.96E-06 | 5.52E-05 | F | F | F | F |
| Rcsd1 | -3.0570488 | 2.86141592 | 23.3580733 | 8.97E-06 | 5.52E-05 | F | F | F | T |
| Rnf126 | 0.94079632 | 7.10597926 | 23.353953 | 8.98E-06 | 5.53E-05 | F | F | F | F |
| Siva1 | -1.3753573 | 5.23481656 | 23.3457465 | 9.01E-06 | 5.54E-05 | F | F | T | F |

| | | | | | | | | | |
|---|---|---|---|---|---|---|---|---|---|
| Hkdc1 | -2.3906844 | 3.41205839 | 23.3453727 | 9.01E-06 | 5.54E-05 | F | F | F | F |
| Chid1 | 1.47259466 | 4.48710809 | 23.2990603 | 9.17E-06 | 5.64E-05 | T | F | T | T |
| Ccdc151 | -4.5314481 | 1.91021905 | 23.2981582 | 9.18E-06 | 5.64E-05 | F | F | F | F |
| Pitx1 | -3.2328632 | 3.0925231 | 23.2960858 | 9.18E-06 | 5.64E-05 | F | F | T | F |
| Cep78 | -1.4332473 | 5.0939459 | 23.2932889 | 9.19E-06 | 5.65E-05 | F | F | F | F |
| Slc2a4 | -8.6925695 | 1.19626349 | 23.2697164 | 9.28E-06 | 5.70E-05 | F | F | T | F |
| Npr3 | -4.5799297 | 1.97303428 | 23.2693568 | 9.28E-06 | 5.70E-05 | F | F | T | F |
| Frrs1l | -4.4527927 | 1.9103442 | 23.2678111 | 9.28E-06 | 5.70E-05 | F | F | T | T |
| Tnks1bp1 | -1.1569733 | 6.29072525 | 23.2670402 | 9.29E-06 | 5.70E-05 | T | F | T | F |
| Idh3b | 1.00099091 | 6.70102914 | 23.252125 | 9.34E-06 | 5.73E-05 | T | F | F | F |
| Dnah8 | -8.6914102 | 1.19604272 | 23.2490786 | 9.35E-06 | 5.73E-05 | T | F | F | T |
| Rpp14 | 1.20293448 | 5.36592409 | 23.2380039 | 9.39E-06 | 5.76E-05 | F | F | F | T |
| Lpgat1 | 0.95417874 | 7.02426795 | 23.2335041 | 9.40E-06 | 5.76E-05 | T | F | T | T |
| Rusc1 | -2.9729566 | 2.84589117 | 23.2068891 | 9.50E-06 | 5.82E-05 | F | F | F | F |
| H4c6 | -0.9612096 | 8.11236721 | 23.2063554 | 9.50E-06 | 5.82E-05 | F | F | F | F |
| Kcnk3 | -8.8613669 | 1.37674216 | 23.2041238 | 9.51E-06 | 5.82E-05 | T | F | T | T |
| Uba7 | 8.63315836 | 0.08471068 | 23.2666934 | 9.52E-06 | 5.83E-05 | F | F | F | F |
| Tmem259 | 0.97414257 | 6.87318711 | 23.1887142 | 9.57E-06 | 5.85E-05 | F | F | F | F |
| Anp32b | 0.79834964 | 9.46987931 | 23.1858079 | 9.58E-06 | 5.86E-05 | F | F | F | F |
| Pmf1 | -1.2110954 | 6.05201068 | 23.1800221 | 9.60E-06 | 5.87E-05 | F | F | F | T |
| Exosc9 | -1.294069 | 5.55950474 | 23.1644884 | 9.65E-06 | 5.90E-05 | T | F | T | F |
| Pitpnm2 | -1.2548887 | 5.76648212 | 23.1515718 | 9.70E-06 | 5.93E-05 | T | F | T | F |
| Lurap1l | -5.6638837 | 1.55167968 | 23.1404234 | 9.74E-06 | 5.95E-05 | T | F | F | T |
| Igf2bp1 | -0.7913271 | 10.4218143 | 23.1403325 | 9.74E-06 | 5.95E-05 | T | F | T | T |
| Zfp180 | 1.26833123 | 5.07327915 | 23.1261907 | 9.80E-06 | 5.98E-05 | F | F | T | T |
| Tfeb | 1.41047269 | 4.5781276 | 23.1249936 | 9.80E-06 | 5.98E-05 | F | F | F | F |
| Srgap3 | 1.30273172 | 5.07667702 | 23.124715 | 9.80E-06 | 5.98E-05 | T | F | T | T |
| Atl3 | 1.1198135 | 5.88255826 | 23.1188514 | 9.82E-06 | 5.99E-05 | F | F | T | T |
| Arhgdig | -4.01935 | 2.12208612 | 23.1123196 | 9.85E-06 | 6.01E-05 | F | F | F | F |
| Atg4c | 1.61570521 | 4.1632934 | 23.1122343 | 9.85E-06 | 6.01E-05 | T | F | F | T |
| Psme2 | -1.7209141 | 4.31500893 | 23.1112152 | 9.85E-06 | 6.01E-05 | F | F | F | F |
| Ncmap | 3.95299399 | 1.49035428 | 23.1104191 | 9.85E-06 | 6.01E-05 | F | F | T | F |
| Glis2 | -1.4878058 | 4.8706514 | 23.1022593 | 9.89E-06 | 6.02E-05 | T | F | T | T |
| Gjb4 | 5.5542549 | 0.57549055 | 23.0926841 | 9.92E-06 | 6.04E-05 | F | F | F | T |
| Dnajb9 | 1.54555928 | 4.21579901 | 23.0916538 | 9.93E-06 | 6.05E-05 | T | F | T | T |
| Sbf2 | 1.01087463 | 6.61376139 | 23.0484065 | 1.01E-05 | 6.14E-05 | T | F | F | T |
| Ly6g2 | -4.9091212 | 1.73988295 | 23.0423157 | 1.01E-05 | 6.16E-05 | F | F | F | F |
| Kbtbd11 | 4.00394977 | 1.23573327 | 23.0414171 | 1.01E-05 | 6.16E-05 | T | F | T | F |
| Fcho2 | 1.65883805 | 4.03525335 | 23.0339879 | 1.01E-05 | 6.17E-05 | T | F | T | T |
| Bloc1s4 | 1.61068509 | 4.13946533 | 23.0177846 | 1.02E-05 | 6.21E-05 | F | F | F | F |
| mt-Nd6 | 1.20744244 | 5.33156444 | 23.0167088 | 1.02E-05 | 6.21E-05 | F | F | F | F |
| Slc24a3 | -8.6772932 | 1.18119181 | 23.0161474 | 1.02E-05 | 6.21E-05 | T | F | F | T |

| | | | | | | | | | |
|---|---|---|---|---|---|---|---|---|---|
| Rfx4 | -8.7242454 | 1.24079069 | 23.0141296 | 1.02E-05 | 6.21E-05 | F | F | F | F |
| Psmb10 | -8.6824773 | 1.19313694 | 22.9948058 | 1.03E-05 | 6.26E-05 | F | F | F | F |
| Fkbp14 | -8.6782364 | 1.18725586 | 22.9844554 | 1.03E-05 | 6.28E-05 | F | F | T | T |
| Oard1 | -1.4742333 | 4.89870695 | 22.9805574 | 1.04E-05 | 6.29E-05 | F | F | F | F |
| A1cf | -5.6393887 | 1.54329859 | 22.9587144 | 1.04E-05 | 6.34E-05 | T | F | T | F |
| Nfatc2ip | -1.5674286 | 4.65891002 | 22.9561767 | 1.05E-05 | 6.34E-05 | T | F | T | F |
| Bmp6 | -2.6627003 | 3.10440929 | 22.9471332 | 1.05E-05 | 6.36E-05 | F | F | T | F |
| Amigo2 | -8.6759572 | 1.185236 | 22.9450281 | 1.05E-05 | 6.37E-05 | F | F | T | F |
| mt-Nd3 | 0.97446832 | 7.00552829 | 22.9424288 | 1.05E-05 | 6.37E-05 | F | F | F | F |
| H4c12 | -0.9776051 | 7.76439376 | 22.9414082 | 1.05E-05 | 6.37E-05 | F | F | F | F |
| Pi4kb | 1.07326626 | 6.17251561 | 22.9346346 | 1.05E-05 | 6.39E-05 | T | F | T | T |
| Slc35f5 | 1.30638522 | 4.90870019 | 22.9329163 | 1.05E-05 | 6.39E-05 | F | F | T | F |
| Pcna | -0.9484045 | 7.81605809 | 22.9282394 | 1.06E-05 | 6.40E-05 | T | F | F | F |
| H2bc15 | -1.0050698 | 7.18290299 | 22.9252246 | 1.06E-05 | 6.40E-05 | F | F | F | F |
| Ift74 | -1.7203401 | 4.29152223 | 22.9177199 | 1.06E-05 | 6.42E-05 | F | F | T | F |
| Ank2 | -1.0921829 | 6.61203807 | 22.9172949 | 1.06E-05 | 6.42E-05 | T | F | T | T |
| Ndc1 | -1.1219437 | 6.43903498 | 22.9104172 | 1.06E-05 | 6.43E-05 | F | F | F | T |
| Aldh2 | -1.19413 | 6.12816138 | 22.9059248 | 1.07E-05 | 6.44E-05 | T | F | T | T |
| Trpm6 | 1.77922131 | 3.75944292 | 22.869288 | 1.08E-05 | 6.53E-05 | T | F | F | T |
| Glul | -0.9039486 | 8.47809916 | 22.860762 | 1.08E-05 | 6.55E-05 | T | F | T | F |
| Ctnnd1 | 0.83018792 | 8.80416121 | 22.8472417 | 1.09E-05 | 6.58E-05 | T | F | T | T |
| Baiap2l1 | -1.6675301 | 4.41432266 | 22.807714 | 1.11E-05 | 6.68E-05 | T | F | T | T |
| Dusp27 | -8.6735505 | 1.18596826 | 22.8064466 | 1.11E-05 | 6.68E-05 | F | F | T | T |
| Zfp119b | -8.7370547 | 1.21787549 | 22.7852047 | 1.12E-05 | 6.74E-05 | F | F | T | F |
| Cmpk1 | 1.16696929 | 5.51890981 | 22.7813915 | 1.12E-05 | 6.74E-05 | T | F | T | T |
| Ctla2b | 5.02957089 | 0.68485696 | 22.776949 | 1.12E-05 | 6.75E-05 | T | F | T | F |
| Phldb2 | 0.94158607 | 7.28885643 | 22.7698071 | 1.12E-05 | 6.77E-05 | T | F | T | T |
| Zc4h2 | -2.4223521 | 3.36099598 | 22.7553493 | 1.13E-05 | 6.80E-05 | T | F | T | T |
| Ino80 | 0.9742193 | 6.83582842 | 22.7547072 | 1.13E-05 | 6.80E-05 | T | F | F | F |
| Slc34a2 | -2.2762406 | 3.52035251 | 22.749866 | 1.13E-05 | 6.82E-05 | T | F | T | F |
| Ndnf | -8.6800637 | 1.19516597 | 22.7391404 | 1.14E-05 | 6.84E-05 | T | T | T | T |
| Lemd3 | 1.13505544 | 5.80905591 | 22.7141858 | 1.15E-05 | 6.91E-05 | F | F | F | T |
| Coro1a | -4.0820476 | 2.04325911 | 22.7116777 | 1.15E-05 | 6.91E-05 | F | F | F | F |
| Zik1 | -1.5795603 | 4.59628483 | 22.7110439 | 1.15E-05 | 6.91E-05 | F | F | T | T |
| Map2 | -2.2820261 | 3.55936447 | 22.6892797 | 1.16E-05 | 6.97E-05 | F | F | T | T |
| Mthfd2l | -3.4019563 | 2.50777393 | 22.6727869 | 1.16E-05 | 7.01E-05 | F | F | F | F |
| Bend3 | -0.9675725 | 7.96175181 | 22.672506 | 1.16E-05 | 7.01E-05 | T | F | T | T |
| Dmbx1 | -3.525465 | 2.51916299 | 22.6719416 | 1.16E-05 | 7.01E-05 | T | F | T | T |
| Slc13a3 | -3.243147 | 2.61166991 | 22.6667119 | 1.17E-05 | 7.02E-05 | F | F | T | T |
| Ehbp1 | 1.19697037 | 5.31466598 | 22.6581632 | 1.17E-05 | 7.04E-05 | T | F | T | F |
| Crocc2 | -8.65516 | 1.16235161 | 22.6573913 | 1.17E-05 | 7.04E-05 | F | F | T | T |
| Cenpi | 1.41282317 | 4.52429371 | 22.6565246 | 1.17E-05 | 7.04E-05 | T | F | F | T |

| | | | | | | | | | |
|---|---|---|---|---|---|---|---|---|---|
| Mapre2 | -1.0977897 | 6.54460425 | 22.648035 | 1.18E-05 | 7.06E-05 | T | F | T | F |
| Rab3il1 | -1.6762724 | 4.36778074 | 22.6364188 | 1.18E-05 | 7.09E-05 | T | F | T | T |
| Nbas | 0.9942038 | 6.63902361 | 22.635806 | 1.18E-05 | 7.09E-05 | F | F | F | F |
| Zfp454 | -4.6815993 | 1.77552033 | 22.6264182 | 1.19E-05 | 7.11E-05 | F | F | F | F |
| Sobp | -3.3832066 | 2.48649202 | 22.6262966 | 1.19E-05 | 7.11E-05 | T | F | F | T |
| Ctnnb1 | -0.8275597 | 9.64354191 | 22.6255169 | 1.19E-05 | 7.11E-05 | F | F | F | F |
| Clmp | -3.4315871 | 2.53592312 | 22.6184924 | 1.19E-05 | 7.13E-05 | F | F | T | F |
| St6gal2 | -3.8141922 | 2.18362111 | 22.6144989 | 1.19E-05 | 7.14E-05 | T | F | T | T |
| Ralgps1 | -2.3777992 | 3.41008775 | 22.6062213 | 1.19E-05 | 7.16E-05 | F | F | T | T |
| Gpr19 | -3.3048073 | 2.54399856 | 22.5948167 | 1.20E-05 | 7.19E-05 | F | F | F | F |
| Pim3 | 1.06449291 | 6.3180066 | 22.5899458 | 1.20E-05 | 7.20E-05 | F | F | T | F |
| Gdf6 | -4.4672274 | 1.85836445 | 22.5898967 | 1.20E-05 | 7.20E-05 | T | F | F | F |
| Vegfc | -8.6642661 | 1.17881096 | 22.5893792 | 1.20E-05 | 7.20E-05 | F | F | F | F |
| Ccnb1ip1 | 0.6567928 | 12.0825274 | 22.5816475 | 1.21E-05 | 7.22E-05 | F | F | F | F |
| Gabre | -5.1951626 | 1.63602533 | 22.5548037 | 1.22E-05 | 7.29E-05 | F | F | F | T |
| Tbc1d23 | 1.44493152 | 4.40641677 | 22.5508111 | 1.22E-05 | 7.30E-05 | T | T | T | T |
| Gm2897 | -8.6771951 | 1.16854866 | 22.5344738 | 1.23E-05 | 7.34E-05 | T | F | F | F |
| Pxk | 1.31409398 | 4.84240972 | 22.5205167 | 1.23E-05 | 7.38E-05 | T | F | T | F |
| Golga4 | -1.0029052 | 7.08730232 | 22.5096856 | 1.24E-05 | 7.41E-05 | F | F | F | T |
| Pkn2 | 0.95110687 | 7.01078128 | 22.5079935 | 1.24E-05 | 7.41E-05 | T | F | T | F |
| 1600012H0 | 1.36273176 | 4.67000034 | 22.5067963 | 1.24E-05 | 7.41E-05 | T | F | F | T |
| Erg28 | 1.19883807 | 5.27717239 | 22.5067051 | 1.24E-05 | 7.41E-05 | F | F | T | F |
| Kat6b | 1.07142971 | 6.07539278 | 22.5022828 | 1.24E-05 | 7.42E-05 | T | F | T | F |
| Coro1c | 0.82478146 | 8.7046587 | 22.4906463 | 1.25E-05 | 7.45E-05 | T | F | T | F |
| Tmem41b | 1.31346863 | 4.86692103 | 22.4840526 | 1.25E-05 | 7.47E-05 | T | F | T | T |
| H2ac10 | -1.0468311 | 6.98750572 | 22.4808791 | 1.25E-05 | 7.48E-05 | F | F | F | F |
| Tmtc2 | 2.29173688 | 2.87738886 | 22.4777114 | 1.25E-05 | 7.48E-05 | T | F | T | F |
| Tdg | -0.9453728 | 7.71751535 | 22.4752428 | 1.26E-05 | 7.49E-05 | F | F | T | F |
| Insyn1 | -5.6008089 | 1.50102415 | 22.4605466 | 1.26E-05 | 7.53E-05 | F | F | F | T |
| Uhrf2 | -1.0274881 | 6.99102307 | 22.4549183 | 1.27E-05 | 7.54E-05 | T | F | T | F |
| Znrf3 | -1.2349061 | 5.79134304 | 22.4408329 | 1.27E-05 | 7.58E-05 | T | F | T | T |
| Sez6 | -8.818308 | 1.33733172 | 22.4391563 | 1.27E-05 | 7.58E-05 | F | F | T | F |
| Exoc5 | 1.05806037 | 6.16931699 | 22.4381869 | 1.27E-05 | 7.58E-05 | T | F | T | T |
| Jcad | 1.04856428 | 6.23653519 | 22.4215267 | 1.28E-05 | 7.63E-05 | T | F | T | T |
| Flywch2 | -4.4568429 | 1.84611899 | 22.4037757 | 1.29E-05 | 7.68E-05 | F | F | F | F |
| Cbfb | 0.97828781 | 6.7226002 | 22.3892356 | 1.30E-05 | 7.72E-05 | T | F | T | T |
| D630003M | -5.6117328 | 1.50412719 | 22.3749994 | 1.31E-05 | 7.76E-05 | F | F | T | F |
| Rbfox3 | -8.6376266 | 1.14160317 | 22.3428138 | 1.32E-05 | 7.86E-05 | T | F | T | F |
| Usp38 | 1.01476075 | 6.45114686 | 22.341091 | 1.32E-05 | 7.86E-05 | F | F | F | F |
| Zfp598 | 0.95740807 | 6.85983677 | 22.3326338 | 1.33E-05 | 7.88E-05 | F | F | F | F |
| Hectd1 | 0.83445379 | 8.43346194 | 22.3320186 | 1.33E-05 | 7.88E-05 | F | F | T | F |
| Ppfibp2 | 1.69854689 | 3.888862 | 22.3270837 | 1.33E-05 | 7.90E-05 | T | F | T | F |

| | | | | | | | | | |
|---|---|---|---|---|---|---|---|---|---|
| Yrdc | 1.14832614 | 5.57361151 | 22.3066471 | 1.34E-05 | 7.96E-05 | F | F | F | F |
| Rbbp4 | -0.8870532 | 8.5014979 | 22.3045773 | 1.34E-05 | 7.96E-05 | T | F | T | T |
| Amer3 | -8.6400945 | 1.14074427 | 22.3011499 | 1.34E-05 | 7.97E-05 | T | F | T | T |
| Fbxl3 | 1.15941391 | 5.49204965 | 22.2955406 | 1.35E-05 | 7.98E-05 | T | F | F | T |
| Ogdh | 0.85242914 | 8.06304755 | 22.2950986 | 1.35E-05 | 7.98E-05 | T | F | T | T |
| Mbp | -1.8818793 | 4.04526625 | 22.2926905 | 1.35E-05 | 7.99E-05 | T | F | T | T |
| Lrrc61 | -2.4993207 | 3.23734829 | 22.2881851 | 1.35E-05 | 8.00E-05 | F | F | T | F |
| Hunk | -1.0915644 | 6.52051998 | 22.2752925 | 1.36E-05 | 8.04E-05 | F | F | T | F |
| Lipa | -1.6833247 | 4.30882945 | 22.2745654 | 1.36E-05 | 8.04E-05 | T | F | F | T |
| Trnau1ap | -1.8117778 | 4.11835593 | 22.2622233 | 1.36E-05 | 8.07E-05 | F | F | F | F |
| Mrpl34 | -1.4111953 | 5.01440956 | 22.2463668 | 1.37E-05 | 8.12E-05 | F | F | F | F |
| Eid2b | -4.8580168 | 1.6894085 | 22.2336707 | 1.38E-05 | 8.16E-05 | F | F | T | T |
| Ttc14 | 1.17275283 | 5.60248149 | 22.2323174 | 1.38E-05 | 8.16E-05 | T | F | T | T |
| Pou3f3 | -8.6946581 | 1.17855157 | 22.2140657 | 1.39E-05 | 8.21E-05 | T | F | F | F |
| Aldh1a3 | 2.34343679 | 3.07870814 | 22.211357 | 1.39E-05 | 8.22E-05 | T | F | T | F |
| Pacsin2 | 0.97976262 | 6.8149614 | 22.201555 | 1.40E-05 | 8.25E-05 | F | F | T | F |
| Gga3 | -1.1968378 | 5.92286972 | 22.2011298 | 1.40E-05 | 8.25E-05 | F | F | T | F |
| Arl10 | 1.35238733 | 4.67371465 | 22.2004607 | 1.40E-05 | 8.25E-05 | F | F | T | T |
| Gm20431 | -9.4707382 | 1.95323038 | 22.1738681 | 1.41E-05 | 8.33E-05 | F | F | F | F |
| Hapln3 | -4.3244959 | 1.99558122 | 22.1501686 | 1.42E-05 | 8.41E-05 | F | F | F | F |
| Slmap | 0.92161401 | 7.09546126 | 22.1409198 | 1.43E-05 | 8.43E-05 | T | F | T | T |
| Jazf1 | 4.4665594 | 1.00546931 | 22.1227409 | 1.44E-05 | 8.49E-05 | T | F | F | T |
| Emc7 | 1.19285704 | 5.25643385 | 22.1162252 | 1.44E-05 | 8.51E-05 | F | F | F | T |
| H1f3 | -0.823338 | 9.79620283 | 22.11616 | 1.44E-05 | 8.51E-05 | T | F | F | T |
| Tesk1 | -1.8751115 | 4.0192396 | 22.0901167 | 1.46E-05 | 8.59E-05 | F | F | F | F |
| Cers5 | 1.05218153 | 6.14291276 | 22.0812082 | 1.46E-05 | 8.62E-05 | T | F | F | T |
| Odc1 | -0.8600477 | 8.95516902 | 22.0745888 | 1.47E-05 | 8.64E-05 | F | F | T | T |
| Usp21 | -1.2615741 | 5.54446058 | 22.0475511 | 1.48E-05 | 8.72E-05 | F | F | F | F |
| Pigm | 0.93676081 | 6.96700921 | 22.0472769 | 1.48E-05 | 8.72E-05 | T | F | T | T |
| Tgoln1 | 0.94089567 | 6.9212001 | 22.0466825 | 1.48E-05 | 8.72E-05 | F | F | T | T |
| Mgst1 | -5.5780474 | 1.47783742 | 22.0462385 | 1.48E-05 | 8.72E-05 | F | F | F | T |
| Cdh10 | -8.6588021 | 1.1484904 | 22.0277846 | 1.49E-05 | 8.78E-05 | T | F | T | F |
| Apoa5 | -8.6230314 | 1.13893556 | 22.0142869 | 1.50E-05 | 8.83E-05 | F | F | F | F |
| Asxl3 | -2.8378031 | 2.87756718 | 22.0115863 | 1.50E-05 | 8.83E-05 | F | F | T | T |
| Hells | -0.9872388 | 7.42308498 | 22.0103627 | 1.50E-05 | 8.83E-05 | T | F | F | F |
| Prkaca | -1.0948516 | 6.50148582 | 21.9923134 | 1.51E-05 | 8.89E-05 | T | F | F | F |
| AU022252 | -1.1793552 | 5.97323828 | 21.9921718 | 1.51E-05 | 8.89E-05 | F | F | T | F |
| Kank2 | -1.1782645 | 5.98962643 | 21.9524439 | 1.54E-05 | 9.03E-05 | T | F | F | F |
| Gk | -2.9740254 | 2.76097287 | 21.9371999 | 1.54E-05 | 9.08E-05 | T | F | T | F |
| Prrx1 | -2.8521459 | 2.95859646 | 21.8790111 | 1.58E-05 | 9.28E-05 | T | F | T | T |
| Zc3h12c | -2.1445498 | 3.6190066 | 21.8729788 | 1.58E-05 | 9.30E-05 | F | F | T | T |
| Srsf9 | -1.0676121 | 6.60401023 | 21.8729496 | 1.58E-05 | 9.30E-05 | F | F | T | F |

| | | | | | | | | | |
|---|---|---|---|---|---|---|---|---|---|
| Fbxo44 | -3.1455071 | 2.61105227 | 21.8699322 | 1.59E-05 | 9.31E-05 | T | F | T | F |
| Ptgfr | -8.6870131 | 1.21061257 | 21.8656357 | 1.59E-05 | 9.32E-05 | T | F | T | T |
| Chpf2 | 1.24371158 | 5.06912176 | 21.8493703 | 1.60E-05 | 9.38E-05 | T | F | F | T |
| Gls2 | 2.04689036 | 3.20482688 | 21.8306938 | 1.61E-05 | 9.44E-05 | F | F | F | F |
| Fbrs | 1.02511238 | 6.29117053 | 21.8273206 | 1.61E-05 | 9.45E-05 | T | F | F | T |
| Xkr7 | -8.6223626 | 1.11949188 | 21.8170276 | 1.62E-05 | 9.49E-05 | F | F | F | T |
| Cx3cl1 | -3.0374659 | 2.70383119 | 21.8150787 | 1.62E-05 | 9.49E-05 | F | F | T | T |
| Ttc28 | -1.6111083 | 4.55987193 | 21.813137 | 1.62E-05 | 9.50E-05 | T | F | T | T |
| Arl4d | -2.4244766 | 3.48204666 | 21.805624 | 1.63E-05 | 9.52E-05 | F | F | F | F |
| Hebp1 | 1.74504183 | 3.74823965 | 21.8021408 | 1.63E-05 | 9.53E-05 | F | F | T | F |
| Trim59 | -1.0823001 | 6.68239887 | 21.8020982 | 1.63E-05 | 9.53E-05 | F | F | T | F |
| Usp51 | -2.5087352 | 3.17396811 | 21.792226 | 1.63E-05 | 9.56E-05 | F | F | F | T |
| Pcsk1 | -8.6375533 | 1.15967477 | 21.7814226 | 1.64E-05 | 9.60E-05 | T | F | F | F |
| Hcn3 | -8.6002437 | 1.10842276 | 21.7733277 | 1.65E-05 | 9.63E-05 | F | F | F | F |
| Grip2 | -8.6665395 | 1.15187935 | 21.7664761 | 1.65E-05 | 9.65E-05 | T | F | T | T |
| Spty2d1 | 1.11890781 | 5.71858723 | 21.7483825 | 1.66E-05 | 9.72E-05 | F | F | T | T |
| Clcn7 | 1.07247619 | 5.9467942 | 21.7447631 | 1.66E-05 | 9.73E-05 | T | F | T | T |
| Mboat1 | -3.6666091 | 2.20479391 | 21.7290814 | 1.67E-05 | 9.78E-05 | F | F | F | F |
| Herc2 | 0.84226089 | 8.07427507 | 21.7290651 | 1.67E-05 | 9.78E-05 | T | F | T | T |
| Hnf1a | -8.5964388 | 1.10752928 | 21.7248769 | 1.68E-05 | 9.79E-05 | F | F | T | F |
| 4833439L1! | -1.0808731 | 6.49229103 | 21.6893257 | 1.70E-05 | 9.93E-05 | F | F | T | F |
| Chd4 | -0.7810961 | 10.1457568 | 21.6874864 | 1.70E-05 | 9.93E-05 | T | F | F | T |
| Icmt | 1.00783619 | 6.41943669 | 21.6741881 | 1.71E-05 | 9.98E-05 | T | F | T | F |
| Dmkn | 2.69687229 | 2.29965275 | 21.6718623 | 1.71E-05 | 9.99E-05 | F | F | F | F |
| Dpf2 | -1.0938876 | 6.41261003 | 21.6596525 | 1.72E-05 | 0.0001 | T | F | F | F |
| Samd9l | 3.32015857 | 1.98528538 | 21.5884322 | 1.77E-05 | 0.000103 | F | F | F | T |
| Dnaaf3 | -3.9289719 | 2.03943251 | 21.5876261 | 1.77E-05 | 0.000103 | F | F | T | F |
| Grasp | -2.8944646 | 2.83869164 | 21.587094 | 1.77E-05 | 0.000103 | F | F | T | F |
| Dkc1 | -0.9619269 | 7.2598145 | 21.5730764 | 1.78E-05 | 0.000104 | F | F | F | F |
| Slc19a1 | 1.10287239 | 5.7172666 | 21.5519097 | 1.79E-05 | 0.000104 | T | F | T | F |
| Olfr639 | 1.05840263 | 6.51223796 | 21.5397813 | 1.80E-05 | 0.000105 | F | F | F | F |
| Garem1 | -1.6084333 | 4.4001181 | 21.5319096 | 1.81E-05 | 0.000105 | T | F | T | T |
| D10Wsu10. | 0.77663905 | 9.8912039 | 21.516873 | 1.82E-05 | 0.000106 | T | F | T | T |
| Afg3l2 | 0.97796034 | 6.59599307 | 21.5100941 | 1.82E-05 | 0.000106 | F | F | F | F |
| Mbtd1 | -1.0250748 | 6.86924557 | 21.4843388 | 1.84E-05 | 0.000107 | T | F | T | F |
| Zfp746 | -1.3518345 | 5.11937057 | 21.4807214 | 1.84E-05 | 0.000107 | F | F | T | T |
| Erbb3 | -1.4383008 | 4.88418121 | 21.479817 | 1.84E-05 | 0.000107 | F | F | T | T |
| Hdac6 | -1.04188 | 6.86004294 | 21.477787 | 1.85E-05 | 0.000107 | F | F | T | F |
| Srprb | 1.02570254 | 6.35635957 | 21.4711151 | 1.85E-05 | 0.000108 | F | F | T | F |
| Sptlc2 | 1.03407615 | 6.16661842 | 21.4664979 | 1.85E-05 | 0.000108 | T | T | T | T |
| Ccne2 | 1.17109842 | 5.30004257 | 21.4589079 | 1.86E-05 | 0.000108 | T | F | T | T |
| Sp2 | 1.05934571 | 5.99671991 | 21.4411393 | 1.87E-05 | 0.000109 | T | F | T | T |

| | | | | | | | | | |
|---|---|---|---|---|---|---|---|---|---|
| Dram1 | 2.01123957 | 3.22549368 | 21.4282182 | 1.88E-05 | 0.000109 | T | F | T | T |
| Kif24 | -1.445725 | 4.87507201 | 21.4271123 | 1.88E-05 | 0.000109 | T | F | T | F |
| Fmn1 | -2.6990245 | 2.95717339 | 21.4081217 | 1.90E-05 | 0.00011 | T | F | F | T |
| Odf2 | -0.9602682 | 7.2431717 | 21.4043132 | 1.90E-05 | 0.00011 | T | F | T | F |
| Npas3 | -8.5769449 | 1.09196129 | 21.4039564 | 1.90E-05 | 0.00011 | T | T | T | T |
| Adam11 | -2.9283766 | 2.7473448 | 21.401153 | 1.90E-05 | 0.00011 | T | F | F | T |
| Aifm3 | -3.3261457 | 2.43136168 | 21.3998905 | 1.90E-05 | 0.00011 | F | F | F | F |
| Vwa5b2 | -8.6034971 | 1.09948548 | 21.387977 | 1.91E-05 | 0.000111 | T | F | F | F |
| Zswim4 | -1.1722519 | 5.9200305 | 21.3856683 | 1.91E-05 | 0.000111 | F | F | T | F |
| mt-Nd1 | 0.73175561 | 10.128978 | 21.368396 | 1.93E-05 | 0.000112 | F | F | F | F |
| Tmem132d | -8.601203 | 1.0974923 | 21.3626328 | 1.93E-05 | 0.000112 | F | F | F | F |
| Trim7 | 4.33043636 | 0.92729335 | 21.3413551 | 1.95E-05 | 0.000113 | T | F | T | T |
| Pitx2 | -2.790791 | 2.88728159 | 21.3412624 | 1.95E-05 | 0.000113 | T | F | F | F |
| A2m | 1.300897 | 4.91220777 | 21.331845 | 1.95E-05 | 0.000113 | F | F | F | F |
| Hook1 | -1.275865 | 5.49767172 | 21.3246399 | 1.96E-05 | 0.000113 | T | F | T | F |
| Nenf | 1.89937575 | 3.46887685 | 21.3237326 | 1.96E-05 | 0.000113 | F | F | F | F |
| Gnptg | 1.57299062 | 4.05952413 | 21.3041567 | 1.97E-05 | 0.000114 | F | F | F | F |
| Fastkd5 | 1.19628556 | 5.15807528 | 21.3008053 | 1.98E-05 | 0.000114 | F | F | F | T |
| Pigg | 1.36612826 | 4.53650599 | 21.3007019 | 1.98E-05 | 0.000114 | T | F | F | F |
| Pimreg | -1.3335212 | 5.16828251 | 21.2840043 | 1.99E-05 | 0.000115 | F | F | T | F |
| Rflna | -8.568292 | 1.08288187 | 21.2817407 | 1.99E-05 | 0.000115 | T | F | F | F |
| Bag3 | 1.04252407 | 6.19599072 | 21.2743704 | 2.00E-05 | 0.000115 | F | F | F | F |
| B3galt1 | -8.6195368 | 1.10978087 | 21.2700715 | 2.00E-05 | 0.000115 | T | F | T | T |
| Ttc19 | 1.37609311 | 4.52394732 | 21.2634069 | 2.01E-05 | 0.000116 | T | T | T | T |
| Per1 | 1.05952846 | 5.98724385 | 21.2633591 | 2.01E-05 | 0.000116 | F | F | F | T |
| Man2a1 | 0.90126888 | 7.31926798 | 21.2407898 | 2.02E-05 | 0.000117 | T | F | F | T |
| Sh3glb1 | 0.94719093 | 6.77543854 | 21.2405747 | 2.02E-05 | 0.000117 | T | F | T | T |
| Ttll12 | 0.87039289 | 7.51217938 | 21.2267356 | 2.04E-05 | 0.000117 | F | F | F | F |
| H3c10 | -0.9374553 | 7.48974746 | 21.2239251 | 2.04E-05 | 0.000117 | F | F | F | T |
| Olfml2b | -8.6072092 | 1.13283959 | 21.2220062 | 2.04E-05 | 0.000117 | F | F | F | F |
| Kcnj12 | -8.6072092 | 1.13283959 | 21.2220062 | 2.04E-05 | 0.000117 | F | F | T | F |
| Esrra | 1.26746522 | 4.97390115 | 21.2089903 | 2.05E-05 | 0.000118 | F | F | T | F |
| Duxbl1 | -8.5647277 | 1.07346937 | 21.1950209 | 2.06E-05 | 0.000119 | T | F | T | T |
| Fnbp1 | 1.05650112 | 5.9994634 | 21.1909575 | 2.06E-05 | 0.000119 | F | F | T | F |
| Klf10 | 1.14720039 | 5.37580638 | 21.1628346 | 2.09E-05 | 0.00012 | F | F | T | F |
| Olfr767 | 1.44646364 | 4.42402571 | 21.154901 | 2.09E-05 | 0.00012 | F | F | F | F |
| Enpp4 | 2.72676383 | 2.26106515 | 21.1509119 | 2.10E-05 | 0.000121 | T | F | F | T |
| Cdc42bpg | -1.31035 | 5.25818184 | 21.1482389 | 2.10E-05 | 0.000121 | F | F | T | F |
| Triml1 | 3.21658133 | 1.66122984 | 21.135417 | 2.11E-05 | 0.000121 | F | F | F | F |
| Olfr943 | -3.2705046 | 2.61090527 | 21.1163069 | 2.12E-05 | 0.000122 | F | F | F | F |
| Cttnbp2 | -3.7391136 | 2.11552965 | 21.1157878 | 2.13E-05 | 0.000122 | T | F | T | F |
| Erfe | 1.94515007 | 3.3204935 | 21.1024462 | 2.14E-05 | 0.000123 | T | F | T | F |

| | | | | | | | | | |
|---|---|---|---|---|---|---|---|---|---|
| Hcfc2 | 1.81578503 | 3.54640857 | 21.0986114 | 2.14E-05 | 0.000123 | F | F | F | T |
| C2cd2 | -2.1299329 | 3.5776658 | 21.0984937 | 2.14E-05 | 0.000123 | T | F | T | F |
| Zwint | 0.90116365 | 7.16031624 | 21.0772183 | 2.16E-05 | 0.000124 | T | F | T | T |
| Pcdhga8 | -8.5664156 | 1.06969013 | 21.0684254 | 2.17E-05 | 0.000124 | F | F | T | F |
| Lpin2 | -1.3671905 | 5.0212196 | 21.0611678 | 2.17E-05 | 0.000124 | F | F | T | F |
| Wscd1 | -5.5354345 | 1.43165627 | 21.0469228 | 2.18E-05 | 0.000125 | T | F | T | F |
| Clk3 | 1.09648314 | 5.77455319 | 21.0420848 | 2.19E-05 | 0.000125 | T | F | F | F |
| Adprh | 1.0808754 | 6.28572755 | 21.0363641 | 2.19E-05 | 0.000126 | F | F | F | F |
| E130309D0 | 1.04919694 | 6.10956765 | 21.0291955 | 2.20E-05 | 0.000126 | T | F | F | T |
| Lig1 | -0.916286 | 7.67888581 | 21.0287027 | 2.20E-05 | 0.000126 | F | F | F | F |
| Paip2 | -1.0546509 | 6.54859455 | 21.0263435 | 2.20E-05 | 0.000126 | F | F | T | T |
| Gpr63 | -5.522445 | 1.42283821 | 21.0218497 | 2.20E-05 | 0.000126 | F | F | F | T |
| Nap1l1 | -0.7884551 | 9.80355477 | 21.005092 | 2.22E-05 | 0.000127 | F | F | F | T |
| Sphkap | -8.6679943 | 1.14740551 | 21.0047687 | 2.22E-05 | 0.000127 | T | F | F | F |
| Kdr | 1.03020977 | 6.73832842 | 20.9963373 | 2.23E-05 | 0.000127 | T | F | F | F |
| Fam20b | 1.0389546 | 6.07118662 | 20.9902281 | 2.23E-05 | 0.000128 | T | F | T | F |
| Srebf1 | 0.99285335 | 6.4265388 | 20.9880937 | 2.23E-05 | 0.000128 | F | F | T | F |
| Emc1 | 0.87142095 | 7.42950549 | 20.9847747 | 2.24E-05 | 0.000128 | F | F | F | F |
| Rassf1 | 1.42121668 | 4.32405138 | 20.9746335 | 2.25E-05 | 0.000128 | T | F | T | F |
| Ctf2 | 1.68598065 | 3.89669749 | 20.9738518 | 2.25E-05 | 0.000128 | F | F | F | F |
| Sox6 | -8.5579778 | 1.08001884 | 20.9710247 | 2.25E-05 | 0.000128 | T | T | T | T |
| Psme1 | -1.5039874 | 4.61454866 | 20.9625688 | 2.26E-05 | 0.000129 | F | F | F | F |
| Plpp7 | -8.5458144 | 1.05794763 | 20.9337904 | 2.28E-05 | 0.00013 | F | F | T | F |
| Gpx3 | -1.0950439 | 6.71211824 | 20.932256 | 2.28E-05 | 0.00013 | F | F | F | F |
| Adamts7 | 1.45221686 | 4.30653846 | 20.917537 | 2.30E-05 | 0.000131 | F | F | F | T |
| Larp1 | 0.75078394 | 9.7873506 | 20.9142777 | 2.30E-05 | 0.000131 | F | F | T | F |
| Avpi1 | 1.57900939 | 4.00935301 | 20.874567 | 2.34E-05 | 0.000133 | F | F | T | T |
| Zyg11b | -1.0179741 | 6.79768116 | 20.8649986 | 2.34E-05 | 0.000134 | T | F | T | T |
| Mxra7 | -1.9615623 | 3.90369902 | 20.8647318 | 2.34E-05 | 0.000134 | T | F | F | F |
| Lypd6 | -3.221553 | 2.46719434 | 20.849271 | 2.36E-05 | 0.000134 | T | F | T | T |
| Aagab | -1.1841504 | 5.74417661 | 20.8432195 | 2.36E-05 | 0.000135 | F | F | T | F |
| Gatad1 | -1.0806559 | 6.35403218 | 20.8406212 | 2.37E-05 | 0.000135 | T | F | F | T |
| Popdc2 | -5.7881361 | 1.69881681 | 20.8299979 | 2.38E-05 | 0.000135 | F | F | F | F |
| Prss16 | 2.71386269 | 2.23559411 | 20.825073 | 2.38E-05 | 0.000135 | F | F | T | F |
| Cdc42 | 0.81420758 | 8.28024932 | 20.8144504 | 2.39E-05 | 0.000136 | T | F | T | F |
| Spry2 | -1.3159024 | 5.18879683 | 20.8085901 | 2.40E-05 | 0.000136 | F | F | T | F |
| Rgs9bp | -2.8716397 | 2.78722407 | 20.8015408 | 2.40E-05 | 0.000137 | T | F | T | T |
| Fam117b | -1.217801 | 5.56677991 | 20.7932938 | 2.41E-05 | 0.000137 | T | F | F | T |
| Tob2 | -1.2376041 | 5.55256275 | 20.7885223 | 2.42E-05 | 0.000137 | T | F | T | F |
| Epg5 | 1.03488787 | 6.10901802 | 20.7713797 | 2.43E-05 | 0.000138 | F | F | F | T |
| Gpatch11 | 1.62162205 | 3.94032698 | 20.7594605 | 2.44E-05 | 0.000139 | T | F | T | T |
| Nfatc2 | -2.9563217 | 2.67182288 | 20.7485125 | 2.45E-05 | 0.000139 | T | F | T | T |

| | | | | | | | | | |
|---|---|---|---|---|---|---|---|---|---|
| Unc5b | 0.87202091 | 7.43850526 | 20.7403507 | 2.46E-05 | 0.00014 | F | F | T | F |
| Gm10521 | 4.65572861 | 0.79428911 | 20.7347987 | 2.47E-05 | 0.00014 | F | F | F | T |
| Adrb1 | -8.5535901 | 1.05460245 | 20.7339426 | 2.47E-05 | 0.00014 | F | F | T | F |
| Nsun2 | 0.7997594 | 8.50860727 | 20.7240376 | 2.48E-05 | 0.000141 | T | F | T | F |
| Col7a1 | -2.1438139 | 3.5518429 | 20.7187747 | 2.48E-05 | 0.000141 | F | F | F | F |
| Ppif | 1.09886078 | 5.60798482 | 20.7141321 | 2.49E-05 | 0.000141 | F | F | T | T |
| Myg1 | -1.4656397 | 4.68392407 | 20.7083708 | 2.49E-05 | 0.000141 | T | F | F | T |
| Rpn2 | 0.84331474 | 7.79222125 | 20.7005538 | 2.50E-05 | 0.000142 | F | F | F | F |
| Dtx1 | 2.44195939 | 2.54176666 | 20.6996086 | 2.50E-05 | 0.000142 | F | F | T | F |
| Fgfbp3 | -5.6407945 | 1.5580357 | 20.6987835 | 2.50E-05 | 0.000142 | F | F | T | F |
| Trpm7 | 0.94811092 | 6.78475462 | 20.6849375 | 2.52E-05 | 0.000142 | T | F | F | F |
| Sumo3 | -0.9741282 | 6.98744792 | 20.6810396 | 2.52E-05 | 0.000143 | T | F | F | T |
| Plk4 | 1.1024068 | 5.71574416 | 20.6760333 | 2.52E-05 | 0.000143 | T | F | F | T |
| Kat2b | 2.18604403 | 2.88878032 | 20.6423918 | 2.56E-05 | 0.000145 | F | F | T | F |
| Snx18 | -1.035174 | 6.599937 | 20.6413718 | 2.56E-05 | 0.000145 | T | F | T | T |
| Yif1b | 1.11078762 | 5.56858171 | 20.626264 | 2.57E-05 | 0.000146 | T | F | F | F |
| Lcp1 | 1.11346134 | 5.49387014 | 20.6182645 | 2.58E-05 | 0.000146 | T | F | T | T |
| Klc2 | 1.04924577 | 5.90522212 | 20.5849234 | 2.62E-05 | 0.000148 | F | F | F | F |
| Zfp977 | 3.70563389 | 1.18793407 | 20.5679813 | 2.63E-05 | 0.000149 | F | F | F | F |
| Kcna2 | -2.9877994 | 2.66397202 | 20.5567299 | 2.65E-05 | 0.000149 | F | F | T | T |
| Adamts14 | -8.5285692 | 1.03589854 | 20.548689 | 2.65E-05 | 0.00015 | F | F | T | F |
| Rnf215 | 1.7653691 | 3.60203747 | 20.5427424 | 2.66E-05 | 0.00015 | T | F | F | T |
| Patj | -1.0434363 | 6.60767175 | 20.5413296 | 2.66E-05 | 0.00015 | T | F | T | F |
| 9530068E0' | 1.03343426 | 6.01533062 | 20.5367134 | 2.67E-05 | 0.00015 | T | F | T | F |
| Fli1 | -3.2071918 | 2.86241993 | 20.5328773 | 2.67E-05 | 0.000151 | T | F | F | T |
| Gpkow | 1.21470487 | 5.07166639 | 20.5326489 | 2.67E-05 | 0.000151 | T | F | F | F |
| Mcm3 | -0.8501638 | 8.51580053 | 20.5160369 | 2.69E-05 | 0.000152 | T | F | F | T |
| Slc45a1 | -3.9171447 | 1.97905812 | 20.5113366 | 2.69E-05 | 0.000152 | T | F | F | F |
| Usp53 | 1.22770044 | 4.98592309 | 20.4911387 | 2.72E-05 | 0.000153 | T | F | F | T |
| Rasl10a | 5.82272019 | 0.33023974 | 20.4743899 | 2.73E-05 | 0.000154 | F | F | F | F |
| Duxbl2 | -8.6456698 | 1.17688785 | 20.454849 | 2.75E-05 | 0.000155 | F | F | T | T |
| 1700066M2 | 1.99481014 | 3.17533304 | 20.43829 | 2.77E-05 | 0.000156 | F | F | F | T |
| Lurap1 | -4.0881504 | 1.88106265 | 20.4333941 | 2.78E-05 | 0.000156 | F | F | F | T |
| H4c11 | -0.8753 | 8.10667664 | 20.4102401 | 2.80E-05 | 0.000158 | F | F | F | T |
| Fdx1 | 1.7955942 | 3.53178411 | 20.4057198 | 2.81E-05 | 0.000158 | F | F | T | T |
| Sel1l | 0.89154431 | 7.02210344 | 20.3877396 | 2.83E-05 | 0.000159 | T | F | T | T |
| Lsg1 | -2.3082284 | 3.28442467 | 20.3798184 | 2.84E-05 | 0.00016 | F | F | F | T |
| Abat | -2.5585584 | 3.06308484 | 20.3744746 | 2.84E-05 | 0.00016 | T | F | T | T |
| Eno4 | -5.4752535 | 1.38628881 | 20.3603601 | 2.86E-05 | 0.000161 | T | F | F | T |
| Itk | 8.47527761 | -0.0599551 | 20.4090688 | 2.86E-05 | 0.000161 | T | T | T | F |
| Ifi30 | 1.15500937 | 5.36156448 | 20.3541538 | 2.87E-05 | 0.000161 | F | F | F | F |
| Entpd5 | 1.65238012 | 3.83322694 | 20.3500663 | 2.87E-05 | 0.000161 | T | F | T | T |

| | | | | | | | | | |
|---|---|---|---|---|---|---|---|---|---|
| Txndc11 | 0.99172953 | 6.30923943 | 20.3495905 | 2.87E-05 | 0.000161 | T | F | F | F |
| Utp20 | 0.78955476 | 8.6087951 | 20.3495433 | 2.87E-05 | 0.000161 | T | F | F | F |
| Celf3 | -8.525582 | 1.02896982 | 20.3417749 | 2.88E-05 | 0.000162 | T | T | F | T |
| Cpt1a | 2.75676669 | 2.09546833 | 20.3102697 | 2.92E-05 | 0.000164 | F | F | T | T |
| Nsg2 | -5.4721879 | 1.38351565 | 20.3101992 | 2.92E-05 | 0.000164 | F | F | T | F |
| Aadat | -4.3382612 | 1.73875121 | 20.2804313 | 2.95E-05 | 0.000165 | F | F | F | F |
| Usp18 | 3.45321052 | 1.37855495 | 20.2749463 | 2.96E-05 | 0.000166 | F | F | T | F |
| Pck2 | 1.1701483 | 5.35596467 | 20.2661688 | 2.97E-05 | 0.000166 | T | F | T | T |
| Zcwpw1 | -3.6933464 | 2.068487 | 20.2646947 | 2.97E-05 | 0.000166 | F | F | F | F |
| Tsnax | 0.96844786 | 6.43958829 | 20.2514311 | 2.98E-05 | 0.000167 | F | F | T | F |
| Fam163a | -5.4673811 | 1.37644714 | 20.2438007 | 2.99E-05 | 0.000168 | T | F | T | T |
| Nkx3-1 | -8.5127193 | 1.01880881 | 20.2303554 | 3.01E-05 | 0.000169 | T | F | T | T |
| Fmo4 | -8.5153912 | 1.04349788 | 20.2222597 | 3.02E-05 | 0.000169 | T | F | T | F |
| Scnn1a | 2.26471121 | 2.73533996 | 20.2056956 | 3.04E-05 | 0.00017 | F | F | T | F |
| Cadm4 | 2.55438422 | 2.32618439 | 20.2020172 | 3.04E-05 | 0.00017 | F | F | T | F |
| Nicn1 | -2.1503832 | 3.47433826 | 20.2002187 | 3.05E-05 | 0.00017 | T | F | T | F |
| Mmp25 | -3.8185934 | 2.07711452 | 20.1976131 | 3.05E-05 | 0.00017 | F | F | F | T |
| Mfsd14a | 0.99310234 | 6.24284115 | 20.1858881 | 3.06E-05 | 0.000171 | T | F | T | F |
| Pde10a | 2.36443798 | 2.74372148 | 20.1844729 | 3.06E-05 | 0.000171 | T | F | F | F |
| Rbpj | -1.0603851 | 6.38804687 | 20.1822678 | 3.07E-05 | 0.000171 | T | F | T | T |
| Hyal2 | -1.1633342 | 5.87291513 | 20.1714728 | 3.08E-05 | 0.000172 | F | F | F | F |
| Kifc3 | -1.910826 | 3.82010286 | 20.1662772 | 3.09E-05 | 0.000172 | F | F | T | T |
| Zfp974 | -4.7023878 | 1.58262562 | 20.1258771 | 3.14E-05 | 0.000175 | T | F | T | F |
| Col14a1 | -4.3377395 | 1.73095303 | 20.1205194 | 3.14E-05 | 0.000175 | T | F | T | T |
| Hpca | 2.73846384 | 2.13439529 | 20.1163856 | 3.15E-05 | 0.000176 | T | F | T | T |
| Smarca4 | -0.7969359 | 9.29811245 | 20.103605 | 3.16E-05 | 0.000177 | F | F | T | T |
| Setd1b | 0.82263038 | 7.94881021 | 20.1028256 | 3.17E-05 | 0.000177 | F | F | F | T |
| Wnt2 | -3.5505505 | 2.14711541 | 20.101035 | 3.17E-05 | 0.000177 | T | F | F | F |
| Prmt1 | -0.8597135 | 8.17717754 | 20.0745809 | 3.20E-05 | 0.000178 | F | F | F | F |
| Zdhhc21 | 1.0063315 | 6.26693349 | 20.0741292 | 3.20E-05 | 0.000178 | T | F | T | T |
| Olig1 | -8.6900306 | 1.1621545 | 20.0560071 | 3.22E-05 | 0.00018 | F | F | F | F |
| Rad21 | 0.7838875 | 8.57416109 | 20.0498587 | 3.23E-05 | 0.00018 | F | F | T | T |
| H3c4 | -0.8444465 | 8.4091159 | 20.0496156 | 3.23E-05 | 0.00018 | F | F | F | T |
| Serpini1 | -8.5330461 | 1.02919895 | 20.0426182 | 3.24E-05 | 0.00018 | T | F | T | T |
| Kdm3b | -0.9199988 | 7.33164072 | 20.040629 | 3.24E-05 | 0.000181 | T | F | T | F |
| Gadd45b | 2.10824715 | 2.95818753 | 20.0370968 | 3.25E-05 | 0.000181 | F | F | F | F |
| St3gal5 | -1.9726258 | 3.75794769 | 20.0253312 | 3.26E-05 | 0.000182 | F | F | F | F |
| Lrrk1 | 1.34120117 | 4.52177907 | 20.0202706 | 3.27E-05 | 0.000182 | F | F | T | T |
| Atrnl1 | -1.1419873 | 5.86276431 | 20.0072631 | 3.29E-05 | 0.000183 | F | F | T | F |
| Pgc | -8.4851212 | 1.00855634 | 20.0070131 | 3.29E-05 | 0.000183 | F | F | F | F |
| Esco1 | 1.16386245 | 5.22668566 | 20.0056427 | 3.29E-05 | 0.000183 | T | F | F | T |
| Tmem120b | 1.61134665 | 3.86477111 | 20.0038417 | 3.29E-05 | 0.000183 | F | F | F | F |

| | | | | | | | | | |
|---|---|---|---|---|---|---|---|---|---|
| Cyb561 | -2.1524606 | 3.47585739 | 20.0032966 | 3.29E-05 | 0.000183 | T | T | T | T |
| Pla1a | -8.4825391 | 1.00038808 | 19.9975361 | 3.30E-05 | 0.000183 | F | F | T | F |
| Dcaf5 | 1.01984964 | 6.02431433 | 19.9937608 | 3.31E-05 | 0.000183 | T | F | T | T |
| Tm9sf2 | 0.85488771 | 7.35327013 | 19.9914343 | 3.31E-05 | 0.000184 | T | F | F | F |
| H2bc9 | -0.9799805 | 7.18472459 | 19.9846568 | 3.32E-05 | 0.000184 | F | F | F | F |
| Padi3 | -2.3218049 | 3.2473812 | 19.9798343 | 3.32E-05 | 0.000184 | F | F | T | F |
| Micall1 | 1.09633393 | 5.49816089 | 19.9785927 | 3.33E-05 | 0.000184 | F | F | T | F |
| Phyhd1 | -8.4834642 | 0.99797783 | 19.9752669 | 3.33E-05 | 0.000185 | F | F | F | F |
| Zfp597 | 1.48163377 | 4.1509796 | 19.9665714 | 3.34E-05 | 0.000185 | T | F | T | T |
| Slc38a8 | -4.6728194 | 1.6297175 | 19.953366 | 3.36E-05 | 0.000186 | F | F | F | F |
| Chrdl2 | 4.81895443 | 0.5844935 | 19.943735 | 3.37E-05 | 0.000187 | F | F | F | F |
| Fyttd1 | 0.99558331 | 6.42212045 | 19.9365154 | 3.38E-05 | 0.000187 | T | F | T | T |
| Zbtb5 | -1.4662239 | 4.5989276 | 19.9222251 | 3.40E-05 | 0.000188 | T | F | T | T |
| Fam177a2 | -8.4825083 | 0.99500944 | 19.9199063 | 3.40E-05 | 0.000188 | F | F | F | T |
| Xlr3a | 1.51762512 | 4.31360278 | 19.9094409 | 3.42E-05 | 0.000189 | F | F | F | F |
| Gphn | 1.27987272 | 4.70008231 | 19.9089988 | 3.42E-05 | 0.000189 | F | F | F | F |
| Ttpa | -8.5141278 | 1.0134848 | 19.9085014 | 3.42E-05 | 0.000189 | T | F | F | T |
| mt-Atp8 | 1.08357136 | 6.2970013 | 19.9004595 | 3.43E-05 | 0.00019 | F | F | F | F |
| Pcdhga7 | -5.1363561 | 1.44530271 | 19.8924926 | 3.44E-05 | 0.00019 | F | F | F | F |
| Cyp4f16 | -2.6027394 | 2.93306578 | 19.8908545 | 3.44E-05 | 0.00019 | F | F | T | T |
| 4930402H2 | 1.30543255 | 4.59472516 | 19.8877473 | 3.45E-05 | 0.000191 | T | F | F | F |
| Srcin1 | -1.6858671 | 4.14749726 | 19.8757001 | 3.46E-05 | 0.000191 | T | F | T | T |
| Zfp982 | -2.0060993 | 3.76083164 | 19.8725441 | 3.47E-05 | 0.000192 | F | F | F | F |
| Exoc3l2 | -4.3168116 | 1.72032102 | 19.8605736 | 3.48E-05 | 0.000192 | T | F | F | T |
| Mindy4 | -2.5114179 | 3.02288322 | 19.8512278 | 3.50E-05 | 0.000193 | F | F | F | F |
| Ccm2 | 1.03333763 | 5.96391359 | 19.8487542 | 3.50E-05 | 0.000193 | F | F | F | F |
| Calm1 | 0.79430386 | 8.2767628 | 19.844799 | 3.51E-05 | 0.000193 | F | F | T | F |
| Shisa8 | -3.8320348 | 1.94905381 | 19.839356 | 3.51E-05 | 0.000194 | F | F | F | F |
| Ubac1 | 1.06702086 | 5.69500661 | 19.8369348 | 3.52E-05 | 0.000194 | F | F | T | F |
| Sema3b | -4.7907575 | 1.638933 | 19.8253358 | 3.53E-05 | 0.000195 | F | F | T | T |
| Pik3cd | 1.27475812 | 4.70394609 | 19.7958129 | 3.58E-05 | 0.000197 | F | F | T | F |
| Trak1 | 1.03325982 | 5.95046643 | 19.7896373 | 3.58E-05 | 0.000198 | T | F | T | F |
| Espl1 | -0.8960714 | 7.60044591 | 19.7741107 | 3.61E-05 | 0.000199 | F | F | F | F |
| Map9 | 1.21563344 | 4.90447626 | 19.7717299 | 3.61E-05 | 0.000199 | T | F | T | T |
| Hspa9 | 0.73382788 | 9.50051634 | 19.7666977 | 3.62E-05 | 0.000199 | T | F | F | F |
| Rpusd1 | -2.334718 | 3.20419218 | 19.7577104 | 3.63E-05 | 0.0002 | F | F | T | F |
| Cd248 | -3.0729778 | 2.56833826 | 19.7484495 | 3.64E-05 | 0.000201 | F | F | F | F |
| Stard13 | -2.5989165 | 2.92309231 | 19.7417296 | 3.65E-05 | 0.000201 | T | F | F | F |
| Rabgef1 | 1.27638939 | 4.68001295 | 19.7274766 | 3.67E-05 | 0.000202 | T | F | T | F |
| Hnrnpa1 | -0.7491553 | 10.0294102 | 19.7224198 | 3.68E-05 | 0.000202 | T | F | F | T |
| Mpeg1 | 6.42762937 | 0.16769978 | 19.7202446 | 3.69E-05 | 0.000203 | F | F | T | F |
| Acot1 | -2.0594017 | 3.57225667 | 19.7191743 | 3.69E-05 | 0.000203 | F | F | F | F |

| | | | | | | | | | |
|---|---|---|---|---|---|---|---|---|---|
| Slc28a2 | 1.52948982 | 4.0115398 | 19.7145733 | 3.69E-05 | 0.000203 | F | F | F | F |
| Lsm14b | -1.0402521 | 6.49608747 | 19.7145419 | 3.69E-05 | 0.000203 | T | F | T | T |
| Grcc10 | -1.6635684 | 4.1681653 | 19.7119218 | 3.70E-05 | 0.000203 | F | F | F | F |
| Sod2 | 1.08460579 | 5.59306199 | 19.7073389 | 3.70E-05 | 0.000203 | T | F | T | T |
| Prrt4 | -8.479795 | 1.01106656 | 19.6992559 | 3.72E-05 | 0.000204 | F | F | T | F |
| Lrrfip2 | 1.05384289 | 5.70647777 | 19.6937773 | 3.72E-05 | 0.000204 | F | F | T | F |
| Clcn3 | 0.90353398 | 6.83961348 | 19.6767342 | 3.75E-05 | 0.000206 | T | F | T | T |
| Caln1 | -3.6006003 | 2.11901017 | 19.6742492 | 3.75E-05 | 0.000206 | F | F | T | T |
| Polg2 | -8.4950191 | 0.99627309 | 19.6742004 | 3.75E-05 | 0.000206 | F | F | F | F |
| Hs3st3a1 | -4.6453714 | 1.58731938 | 19.673563 | 3.75E-05 | 0.000206 | T | T | T | T |
| Elovl6 | -1.0820806 | 6.16575402 | 19.6679167 | 3.76E-05 | 0.000206 | T | F | T | T |
| Gm20695 | -8.4592241 | 0.97945609 | 19.6613286 | 3.77E-05 | 0.000207 | F | F | F | F |
| Memo1 | 1.25910982 | 4.81108133 | 19.6411784 | 3.80E-05 | 0.000208 | T | F | T | T |
| Pam | 1.09789013 | 5.47823941 | 19.6367351 | 3.81E-05 | 0.000209 | T | F | T | T |
| Sacm1l | 1.02883571 | 5.91079021 | 19.5948966 | 3.87E-05 | 0.000212 | T | F | F | F |
| Lysmd2 | 1.71574666 | 3.64622171 | 19.5927642 | 3.88E-05 | 0.000212 | T | F | F | F |
| Cnot6 | 0.87012647 | 7.0628815 | 19.581287 | 3.89E-05 | 0.000213 | T | F | T | T |
| Cdc42se2 | 0.90633553 | 6.78695037 | 19.5633306 | 3.92E-05 | 0.000215 | T | F | T | T |
| Oas1g | 8.42720475 | -0.1035963 | 19.604808 | 3.93E-05 | 0.000215 | F | F | F | T |
| Pdzk1ip1 | -8.4788664 | 0.98315699 | 19.552187 | 3.94E-05 | 0.000215 | F | F | F | F |
| Nhs | -1.4536379 | 4.62830308 | 19.5364861 | 3.97E-05 | 0.000217 | T | T | F | T |
| Asap1 | -1.0617008 | 6.24841725 | 19.5317544 | 3.97E-05 | 0.000217 | F | F | T | F |
| Fam107a | -2.8482793 | 2.77655373 | 19.5233116 | 3.99E-05 | 0.000218 | T | F | T | T |
| Plpbp | 0.98870167 | 6.15157056 | 19.5102545 | 4.01E-05 | 0.000219 | T | F | T | F |
| Foxm1 | -0.9498983 | 6.94521405 | 19.5081722 | 4.01E-05 | 0.000219 | T | F | T | T |
| Crebrf | 1.77467686 | 3.50736613 | 19.5023719 | 4.02E-05 | 0.000219 | F | T | T | T |
| Mettl7a1 | -2.8741182 | 2.6497989 | 19.4941085 | 4.03E-05 | 0.00022 | T | F | T | T |
| Shb | -1.326858 | 4.96468986 | 19.4895892 | 4.04E-05 | 0.00022 | T | F | T | F |
| Rorc | -8.4579184 | 0.96925957 | 19.4834158 | 4.05E-05 | 0.000221 | F | F | F | F |
| Ago3 | 1.10904028 | 5.34989414 | 19.4561531 | 4.09E-05 | 0.000223 | F | F | T | T |
| Use1 | -1.5043825 | 4.44951908 | 19.4533279 | 4.10E-05 | 0.000224 | F | F | F | F |
| Mov10l1 | 3.73526409 | 1.08915926 | 19.4514776 | 4.10E-05 | 0.000224 | F | F | F | F |
| Etv1 | -1.7738329 | 4.13566494 | 19.4348175 | 4.13E-05 | 0.000225 | T | F | T | T |
| Hus1 | 1.09984907 | 5.40945519 | 19.4310891 | 4.14E-05 | 0.000225 | T | F | T | F |
| Hoxd9 | -5.0741277 | 1.91997854 | 19.4259078 | 4.14E-05 | 0.000226 | T | F | F | T |
| Sec61a1 | 0.76542219 | 8.84296307 | 19.4235504 | 4.15E-05 | 0.000226 | F | F | T | F |
| Wbp1 | -1.7286598 | 4.08201446 | 19.418796 | 4.16E-05 | 0.000226 | F | F | F | F |
| Sap30l | 1.11129246 | 5.29909243 | 19.4029377 | 4.18E-05 | 0.000228 | F | F | T | T |
| Klhl14 | -8.5155506 | 1.00809215 | 19.3946164 | 4.20E-05 | 0.000228 | F | F | T | T |
| Dkk3 | -3.8315299 | 1.95818787 | 19.3834785 | 4.22E-05 | 0.000229 | T | T | F | T |
| Ccdc73 | 5.7621853 | 0.27475705 | 19.3795923 | 4.22E-05 | 0.00023 | T | F | F | F |
| Rilpl1 | -1.8159463 | 3.89865889 | 19.3743542 | 4.23E-05 | 0.00023 | F | F | F | F |

| | | | | | | | | | |
|---|---|---|---|---|---|---|---|---|---|
| Ccdc32 | 1.52401762 | 4.01350371 | 19.3636229 | 4.25E-05 | 0.000231 | F | F | F | F |
| Spats2l | -2.6890982 | 2.7994826 | 19.3622967 | 4.25E-05 | 0.000231 | T | F | F | T |
| Wdtc1 | 0.96487223 | 6.30858453 | 19.3588641 | 4.26E-05 | 0.000231 | F | F | T | F |
| Lzic | 1.20593676 | 4.9152003 | 19.3539298 | 4.27E-05 | 0.000232 | F | F | F | T |
| Sdf2l1 | 1.12228942 | 5.30694914 | 19.3341224 | 4.30E-05 | 0.000234 | F | F | F | F |
| H3c1 | -0.8206259 | 8.71018194 | 19.3246487 | 4.32E-05 | 0.000234 | F | F | F | F |
| Gm45837 | -9.1093999 | 1.61439223 | 19.3143135 | 4.33E-05 | 0.000235 | F | F | F | F |
| Dpysl4 | -8.4380383 | 0.95527833 | 19.3084798 | 4.34E-05 | 0.000236 | T | F | F | F |
| Nck2 | -1.3545468 | 4.85908188 | 19.3003098 | 4.36E-05 | 0.000236 | T | F | F | F |
| Tor1b | 1.06516693 | 5.57397546 | 19.2918362 | 4.37E-05 | 0.000237 | T | F | T | F |
| Sgtb | -3.3982085 | 2.18859303 | 19.2913018 | 4.37E-05 | 0.000237 | T | F | T | F |
| Tnk1 | -1.9908554 | 3.64133096 | 19.2748505 | 4.40E-05 | 0.000239 | F | F | T | F |
| Rab17 | -5.4057585 | 1.32110447 | 19.2725924 | 4.41E-05 | 0.000239 | T | F | T | T |
| Rai14 | 0.87953567 | 7.1393055 | 19.2701228 | 4.41E-05 | 0.000239 | T | F | T | F |
| Chac1 | 2.93302368 | 1.77284605 | 19.2669737 | 4.42E-05 | 0.000239 | F | F | F | F |
| Olfr1160 | 2.90215399 | 1.78275436 | 19.2609868 | 4.43E-05 | 0.00024 | F | F | F | F |
| Six5 | -2.0709711 | 3.4958772 | 19.2483282 | 4.45E-05 | 0.000241 | T | F | T | F |
| Larp4b | 0.81035148 | 7.8205819 | 19.2398328 | 4.46E-05 | 0.000242 | T | F | T | T |
| Tcerg1l | -8.4286066 | 0.95347173 | 19.2235635 | 4.49E-05 | 0.000243 | T | F | F | F |
| Tnip1 | 1.02707515 | 5.81545742 | 19.2218757 | 4.50E-05 | 0.000243 | T | F | T | F |
| Pcdhga3 | -8.4281906 | 0.95460699 | 19.2085966 | 4.52E-05 | 0.000245 | F | F | F | F |
| Pcca | -1.4437982 | 4.58928527 | 19.1919808 | 4.55E-05 | 0.000246 | F | T | F | T |
| Map11 | 1.16552255 | 5.05746516 | 19.181764 | 4.57E-05 | 0.000247 | F | F | T | F |
| Dlec1 | -8.4245123 | 0.95138865 | 19.1553003 | 4.62E-05 | 0.00025 | F | F | F | T |
| Sp6 | -1.6773754 | 4.0948061 | 19.1509631 | 4.63E-05 | 0.00025 | F | F | T | F |
| Zdhhc2 | -1.4683183 | 4.53057356 | 19.1465568 | 4.64E-05 | 0.00025 | F | F | F | T |
| Ngef | -8.4274115 | 0.94479644 | 19.1457121 | 4.64E-05 | 0.00025 | T | F | T | T |
| Clgn | -8.4274115 | 0.94479644 | 19.1457121 | 4.64E-05 | 0.00025 | F | F | T | T |
| Arl6ip5 | 1.19361503 | 4.92601308 | 19.1383962 | 4.65E-05 | 0.000251 | T | F | T | T |
| Twsg1 | -1.0693021 | 6.21224048 | 19.1302519 | 4.67E-05 | 0.000252 | T | F | T | T |
| March7 | 0.94940621 | 6.39228086 | 19.1272037 | 4.67E-05 | 0.000252 | T | F | F | T |
| P4htm | -8.4360843 | 0.94746369 | 19.1220191 | 4.68E-05 | 0.000253 | T | F | T | F |
| Pllp | -5.4341094 | 1.33234493 | 19.1171303 | 4.69E-05 | 0.000253 | T | F | T | F |
| Cfap206 | -5.4156066 | 1.31878033 | 19.1140249 | 4.70E-05 | 0.000253 | F | F | F | F |
| Cblb | -1.0500396 | 6.23490639 | 19.100375 | 4.72E-05 | 0.000255 | T | F | T | T |
| 2010300C0: | 2.68338025 | 2.05904438 | 19.0963453 | 4.73E-05 | 0.000255 | F | F | T | F |
| Brap | 1.00903071 | 5.97276859 | 19.094872 | 4.73E-05 | 0.000255 | F | F | T | T |
| Hist1h4n | -0.968492 | 8.35151635 | 19.0858849 | 4.75E-05 | 0.000256 | F | F | T | F |
| Prkar1b | -8.4301761 | 0.94329201 | 19.0823359 | 4.76E-05 | 0.000256 | F | F | F | F |
| mt-Cytb | 0.6982755 | 10.0199082 | 19.0705031 | 4.78E-05 | 0.000257 | F | F | F | F |
| Crlf1 | -5.3919636 | 1.30518024 | 19.0524159 | 4.81E-05 | 0.000259 | F | F | F | F |
| Ndufv3 | -1.3247035 | 4.9314596 | 19.0104698 | 4.90E-05 | 0.000263 | F | F | F | F |

| | | | | | | | | | |
|---|---|---|---|---|---|---|---|---|---|
| Zbtb48 | -2.4930134 | 2.97361965 | 18.998338 | 4.92E-05 | 0.000265 | F | F | F | F |
| Fbxo25 | -2.1863103 | 3.37484265 | 18.9920028 | 4.93E-05 | 0.000265 | F | F | F | T |
| Sept7 | 0.85953409 | 7.03865811 | 18.9878759 | 4.94E-05 | 0.000266 | T | F | T | T |
| Kif6 | -8.4124265 | 0.93466436 | 18.9856767 | 4.94E-05 | 0.000266 | F | F | F | F |
| March8 | 1.03940581 | 5.67745114 | 18.9632443 | 4.99E-05 | 0.000268 | T | T | T | T |
| Tcp11l1 | -3.0605995 | 2.43861404 | 18.9534624 | 5.01E-05 | 0.000269 | T | F | T | T |
| Rbm4 | 1.01198959 | 5.89144393 | 18.9501672 | 5.02E-05 | 0.000269 | T | F | F | T |
| Xylt2 | 1.03154042 | 5.77760152 | 18.9451682 | 5.03E-05 | 0.00027 | F | F | T | F |
| AI467606 | -8.4136293 | 0.93161626 | 18.9431802 | 5.03E-05 | 0.00027 | F | F | F | F |
| Bcas1 | -5.3917793 | 1.30031171 | 18.934807 | 5.05E-05 | 0.000271 | F | F | T | F |
| Twf2 | 1.10682879 | 5.2461772 | 18.927139 | 5.06E-05 | 0.000272 | T | F | F | F |
| Sp1 | 0.78622828 | 8.13145066 | 18.9248144 | 5.07E-05 | 0.000272 | T | F | T | T |
| Rhox4a | 8.38746785 | -0.1395186 | 18.9624704 | 5.08E-05 | 0.000273 | F | F | F | F |
| Mreg | -1.6899791 | 4.05881744 | 18.9029043 | 5.11E-05 | 0.000274 | F | F | F | F |
| S100a10 | 1.11266307 | 5.27155088 | 18.886797 | 5.15E-05 | 0.000276 | F | F | T | F |
| Kif13b | 0.95821545 | 6.26632611 | 18.8711495 | 5.18E-05 | 0.000278 | F | F | T | T |
| Maml1 | -0.9990502 | 6.51845601 | 18.8471481 | 5.23E-05 | 0.00028 | T | F | T | F |
| Ptpn23 | 0.82091869 | 7.53229571 | 18.8453145 | 5.23E-05 | 0.00028 | F | F | F | F |
| Siah2 | 1.53092987 | 3.93810822 | 18.8411316 | 5.24E-05 | 0.000281 | T | F | T | T |
| Gm7694 | 1.55692625 | 3.92302558 | 18.8400655 | 5.24E-05 | 0.000281 | F | F | T | T |
| Eif4e3 | 1.65557883 | 3.67321823 | 18.8294414 | 5.27E-05 | 0.000282 | F | F | T | T |
| Mospd1 | 1.41743591 | 4.17281459 | 18.8206484 | 5.28E-05 | 0.000283 | T | F | F | T |
| Ndufa4l2 | -8.412718 | 0.9265065 | 18.8190612 | 5.29E-05 | 0.000283 | F | F | T | F |
| Mitd1 | 1.64819872 | 3.68897716 | 18.8176587 | 5.29E-05 | 0.000283 | F | F | T | F |
| Mpzl3 | 2.12323541 | 2.83038468 | 18.8095709 | 5.31E-05 | 0.000284 | F | F | T | F |
| Snx25 | 1.44815948 | 4.10588891 | 18.7946686 | 5.34E-05 | 0.000286 | T | F | F | F |
| Pqlc2 | -1.673579 | 4.08003692 | 18.7827772 | 5.37E-05 | 0.000287 | T | F | T | F |
| Bicc1 | -1.3120021 | 4.97378094 | 18.7783141 | 5.38E-05 | 0.000287 | T | F | T | T |
| 2410004P0: | -2.5197979 | 2.92832507 | 18.7772408 | 5.38E-05 | 0.000287 | T | F | F | F |
| Snap91 | 2.2402146 | 2.6519591 | 18.7759516 | 5.38E-05 | 0.000287 | T | F | T | T |
| Phldb1 | 0.82709248 | 7.39556551 | 18.756964 | 5.42E-05 | 0.00029 | F | F | T | F |
| Bsn | -3.2913905 | 2.20557026 | 18.7410897 | 5.46E-05 | 0.000291 | F | F | F | F |
| Rbm10 | -0.9015814 | 7.15105175 | 18.7255935 | 5.49E-05 | 0.000293 | T | F | F | F |
| Thoc7 | -1.3278765 | 4.87280752 | 18.7211952 | 5.50E-05 | 0.000294 | F | F | F | F |
| Rnf31 | 1.06778738 | 5.46277854 | 18.6989186 | 5.55E-05 | 0.000296 | T | F | F | F |
| Ankrd27 | -1.0378846 | 6.23186214 | 18.6715058 | 5.61E-05 | 0.000299 | F | F | F | F |
| Bcan | -8.3892999 | 0.92152372 | 18.6349417 | 5.70E-05 | 0.000304 | F | F | F | F |
| Prr7 | -3.4632951 | 2.06905001 | 18.6338256 | 5.70E-05 | 0.000304 | F | F | F | F |
| Proz | -8.388425 | 0.92034959 | 18.6286208 | 5.71E-05 | 0.000304 | F | F | T | T |
| Xylb | 2.2315088 | 2.85205144 | 18.6161331 | 5.74E-05 | 0.000306 | T | F | T | T |
| Nckap1 | 0.78145151 | 8.03166127 | 18.613302 | 5.75E-05 | 0.000306 | F | T | T | T |
| Qars | 1.086648 | 5.91259019 | 18.5890519 | 5.80E-05 | 0.000309 | T | F | F | F |

| | | | | | | | | | |
|---|---|---|---|---|---|---|---|---|---|
| Homez | -1.4719908 | 4.43088191 | 18.5866798 | 5.81E-05 | 0.000309 | T | F | F | T |
| Cmtm3 | 1.52140937 | 4.00115133 | 18.5850739 | 5.81E-05 | 0.000309 | F | F | F | F |
| Dse | -1.1842051 | 5.43373028 | 18.5818011 | 5.82E-05 | 0.00031 | T | T | T | T |
| Gm45623 | -8.4343765 | 0.93577371 | 18.5808475 | 5.82E-05 | 0.00031 | F | F | F | F |
| Dbp | -3.513569 | 2.10150414 | 18.5729408 | 5.84E-05 | 0.000311 | F | F | T | F |
| Sec24b | 0.8438439 | 7.12131035 | 18.5540395 | 5.89E-05 | 0.000313 | F | F | F | T |
| Pogz | -0.9154135 | 7.04975173 | 18.5530071 | 5.89E-05 | 0.000313 | F | F | F | F |
| Tigar | 1.15942773 | 5.00148775 | 18.5508315 | 5.89E-05 | 0.000313 | F | F | T | F |
| 1110032F0 | -8.3800107 | 0.90814878 | 18.5474895 | 5.90E-05 | 0.000314 | F | F | T | F |
| Kcnn1 | -8.3852412 | 0.90502989 | 18.5418136 | 5.91E-05 | 0.000314 | T | F | T | T |
| Derl2 | 1.08616689 | 5.30226856 | 18.5398801 | 5.92E-05 | 0.000314 | T | F | T | F |
| Ttll4 | -0.8645531 | 7.58321092 | 18.523339 | 5.96E-05 | 0.000316 | F | F | T | T |
| 4930452B0 | -2.8800042 | 2.58953924 | 18.522011 | 5.96E-05 | 0.000317 | F | F | T | F |
| Il17rc | 2.37559297 | 2.49969862 | 18.5159001 | 5.98E-05 | 0.000317 | F | F | F | F |
| Ccsap | 1.62378524 | 3.70862035 | 18.5135031 | 5.98E-05 | 0.000318 | T | F | T | F |
| Apmap | 1.29507453 | 4.51909955 | 18.5010715 | 6.01E-05 | 0.000319 | F | F | F | F |
| Ppp1r3d | 3.21094278 | 1.41507451 | 18.4983241 | 6.02E-05 | 0.000319 | F | F | T | F |
| Eif2s2 | 0.76873305 | 8.31889285 | 18.4878769 | 6.05E-05 | 0.000321 | T | F | F | F |
| Pou5f2 | 4.53011565 | 0.59051754 | 18.480643 | 6.06E-05 | 0.000321 | F | F | T | F |
| Zfp606 | -1.4201617 | 4.55899439 | 18.4663894 | 6.10E-05 | 0.000323 | F | F | T | F |
| Klk8 | -2.5120125 | 2.91140718 | 18.4632409 | 6.11E-05 | 0.000324 | F | F | F | F |
| Ntn1 | -0.9571819 | 6.72982764 | 18.4615107 | 6.11E-05 | 0.000324 | T | F | T | T |
| Smad5 | -1.0215247 | 6.32906052 | 18.4521982 | 6.13E-05 | 0.000325 | T | F | T | T |
| Plcxd1 | -5.3617216 | 1.27148719 | 18.4493213 | 6.14E-05 | 0.000325 | F | F | F | F |
| Zmat4 | -8.3712768 | 0.89874986 | 18.4265435 | 6.20E-05 | 0.000328 | T | F | T | T |
| Isg15 | 5.02244295 | 0.44989766 | 18.4206151 | 6.21E-05 | 0.000329 | F | F | F | F |
| Cercam | -2.1807194 | 3.27793581 | 18.4170545 | 6.22E-05 | 0.000329 | F | F | F | F |
| Pnpla8 | 1.01046924 | 5.83361127 | 18.4156189 | 6.23E-05 | 0.000329 | F | F | T | T |
| Cyp2j6 | -8.3700562 | 0.89763906 | 18.409893 | 6.24E-05 | 0.00033 | T | T | F | T |
| Git1 | 0.84670584 | 7.03787683 | 18.4088596 | 6.24E-05 | 0.00033 | F | F | T | F |
| Bivm | -1.3744416 | 4.68313234 | 18.4062903 | 6.25E-05 | 0.00033 | T | F | F | T |
| Klhdc4 | 1.01880935 | 5.72117993 | 18.4037423 | 6.26E-05 | 0.000331 | F | F | T | F |
| Coro1b | 0.87679158 | 6.87438237 | 18.3893568 | 6.29E-05 | 0.000332 | T | F | F | F |
| Adig | 8.35364473 | -0.1699841 | 18.4314566 | 6.29E-05 | 0.000332 | F | F | F | T |
| Dmrtc1b | 8.35364473 | -0.1699841 | 18.4314566 | 6.29E-05 | 0.000332 | T | F | F | F |
| Card19 | 1.97268076 | 3.03374725 | 18.3869031 | 6.30E-05 | 0.000332 | T | T | F | F |
| Map3k8 | -8.465761 | 0.95801993 | 18.3861923 | 6.30E-05 | 0.000332 | T | F | T | T |
| Acer3 | 1.08817235 | 5.29348171 | 18.3861736 | 6.30E-05 | 0.000332 | F | F | T | F |
| Dnajc18 | -1.3436169 | 4.76553061 | 18.3778648 | 6.32E-05 | 0.000333 | T | F | T | T |
| Wfs1 | 1.24442165 | 4.64509526 | 18.3575337 | 6.37E-05 | 0.000336 | F | F | F | F |
| Pdxk | 0.97857262 | 6.00977255 | 18.3572075 | 6.37E-05 | 0.000336 | T | F | T | T |
| Cc2d1a | 1.08862287 | 5.27002329 | 18.3447835 | 6.41E-05 | 0.000338 | F | F | F | F |

| | | | | | | | | | |
|---|---|---|---|---|---|---|---|---|---|
| Ddah1 | -1.637163 | 4.10564611 | 18.3412769 | 6.42E-05 | 0.000338 | F | F | F | T |
| Phka1 | 1.56605679 | 3.81826803 | 18.3279223 | 6.45E-05 | 0.00034 | T | F | F | T |
| Tmf1 | 1.05156071 | 5.48762836 | 18.3129324 | 6.49E-05 | 0.000342 | T | F | F | F |
| Ptpn21 | -1.7120291 | 3.97528046 | 18.3074767 | 6.50E-05 | 0.000343 | T | F | T | T |
| Pbx2 | -0.9139768 | 7.04173601 | 18.3055283 | 6.51E-05 | 0.000343 | T | F | T | T |
| Trpc3 | -8.3666121 | 0.88863879 | 18.3021524 | 6.52E-05 | 0.000343 | T | F | F | T |
| Slc35d3 | -3.665159 | 2.03370734 | 18.300635 | 6.52E-05 | 0.000343 | F | F | T | F |
| St3gal6 | -4.6441355 | 1.49843706 | 18.2931272 | 6.54E-05 | 0.000344 | T | F | T | F |
| Mnt | 1.00089487 | 5.82578069 | 18.2596183 | 6.63E-05 | 0.000349 | F | F | T | F |
| Rab33b | 1.11264169 | 5.18685956 | 18.2572417 | 6.64E-05 | 0.000349 | T | F | T | T |
| Abl1 | -0.8789014 | 7.56333156 | 18.2488919 | 6.66E-05 | 0.00035 | F | F | F | T |
| Fam49a | 1.30337584 | 4.40812619 | 18.2475729 | 6.67E-05 | 0.00035 | T | F | T | T |
| Foxo3 | -1.0562044 | 6.32801725 | 18.2385135 | 6.69E-05 | 0.000352 | F | F | T | T |
| Cfap77 | -8.3566556 | 0.88545806 | 18.227928 | 6.72E-05 | 0.000353 | F | F | F | T |
| Srsf7 | -0.8678547 | 7.43860313 | 18.227529 | 6.72E-05 | 0.000353 | T | F | T | T |
| Pdlim2 | 1.77880632 | 3.52234101 | 18.2127993 | 6.76E-05 | 0.000355 | F | F | F | F |
| Sardh | -4.7181857 | 1.43334055 | 18.2105995 | 6.77E-05 | 0.000355 | F | F | F | F |
| Maff | 1.8147075 | 3.43352819 | 18.2084178 | 6.77E-05 | 0.000355 | T | F | T | T |
| Ripk3 | -2.7616228 | 2.65558719 | 18.1955409 | 6.81E-05 | 0.000357 | F | F | F | F |
| Tmem9b | 1.47386909 | 4.04850946 | 18.1756293 | 6.86E-05 | 0.00036 | F | F | F | T |
| Flywch1 | -0.9817116 | 6.52781142 | 18.1716826 | 6.87E-05 | 0.000361 | F | F | T | F |
| Pifo | -8.3542615 | 0.87932179 | 18.1658275 | 6.89E-05 | 0.000361 | F | F | F | F |
| Slc25a43 | 3.04045059 | 1.53764797 | 18.1618436 | 6.90E-05 | 0.000362 | T | F | F | T |
| Tmem229b | 2.07756314 | 2.97624841 | 18.1537104 | 6.92E-05 | 0.000363 | F | F | T | F |
| Ermn | -8.375416 | 0.88754248 | 18.1461857 | 6.95E-05 | 0.000364 | T | F | T | T |
| Zfp326 | -1.2981386 | 4.88565317 | 18.1437607 | 6.95E-05 | 0.000364 | T | F | F | T |
| Rwdd3 | -8.3571149 | 0.87788889 | 18.1331822 | 6.98E-05 | 0.000366 | F | F | F | F |
| Abhd2 | 0.79965623 | 7.65090651 | 18.1304921 | 6.99E-05 | 0.000366 | F | F | T | T |
| Itga3 | -1.3933977 | 5.4149483 | 18.1125006 | 7.04E-05 | 0.000369 | T | F | F | T |
| Zfp217 | -0.9529857 | 6.73909279 | 18.1094022 | 7.05E-05 | 0.000369 | T | F | F | T |
| H3c11 | -0.8175626 | 8.54709008 | 18.1090839 | 7.05E-05 | 0.000369 | F | F | F | F |
| Yme1l1 | 0.82094406 | 7.28840286 | 18.0951503 | 7.09E-05 | 0.000371 | T | F | T | T |
| Inpp5k | 1.61313545 | 3.6989592 | 18.0905744 | 7.11E-05 | 0.000372 | T | F | T | T |
| Trappc10 | 0.95435446 | 6.13955806 | 18.0879776 | 7.11E-05 | 0.000372 | F | F | T | T |
| Mapk9 | 1.03883311 | 5.5253377 | 18.0740343 | 7.15E-05 | 0.000374 | T | F | T | T |
| Spef2 | -8.3642945 | 0.87878837 | 18.058615 | 7.20E-05 | 0.000376 | F | F | F | F |
| Rragc | 1.02929259 | 5.59622651 | 18.0535962 | 7.21E-05 | 0.000377 | T | F | T | T |
| Ywhah | 0.78372509 | 7.8141357 | 18.0507585 | 7.22E-05 | 0.000377 | T | F | F | F |
| Eps15 | 0.96760977 | 6.03451365 | 18.0276324 | 7.29E-05 | 0.000381 | F | F | F | T |
| Ncln | 0.80881883 | 7.48360364 | 18.0168497 | 7.32E-05 | 0.000382 | F | F | F | F |
| Tmem192 | 1.30364451 | 4.38469407 | 18.0076232 | 7.35E-05 | 0.000384 | F | F | F | F |
| Lrch3 | 0.93848758 | 6.29755138 | 18.0075809 | 7.35E-05 | 0.000384 | T | F | T | T |

| | | | | | | | | | |
|---|---|---|---|---|---|---|---|---|---|
| Rab35 | 0.89574808 | 6.60378596 | 18.0020926 | 7.37E-05 | 0.000384 | F | F | T | F |
| Prr16 | 5.27552414 | 0.32482046 | 18.0018714 | 7.37E-05 | 0.000384 | T | F | T | F |
| Cul4b | 0.80428981 | 7.58035797 | 17.9962633 | 7.38E-05 | 0.000385 | F | F | T | T |
| Myo1e | 1.0664741 | 5.35211398 | 17.9784183 | 7.44E-05 | 0.000388 | T | F | T | T |
| Josd1 | 0.97807133 | 5.9392382 | 17.971617 | 7.46E-05 | 0.000389 | F | F | T | F |
| Gem | -8.4358274 | 0.92968463 | 17.9710673 | 7.46E-05 | 0.000389 | F | F | T | F |
| Ppib | 0.78798058 | 7.78569912 | 17.9685862 | 7.47E-05 | 0.000389 | T | F | T | T |
| Zfp507 | -1.0480942 | 6.03192276 | 17.9361812 | 7.57E-05 | 0.000394 | T | F | T | T |
| Tap1 | 2.86993776 | 1.74084708 | 17.9329866 | 7.58E-05 | 0.000395 | T | F | F | F |
| Garnl3 | -2.4771617 | 2.90020635 | 17.9312464 | 7.58E-05 | 0.000395 | T | F | F | F |
| Ccdc39 | -8.3551906 | 0.8699532 | 17.9218621 | 7.61E-05 | 0.000396 | T | F | T | F |
| Esf1 | 0.88926234 | 6.61574612 | 17.9182919 | 7.62E-05 | 0.000397 | F | F | F | F |
| Hspa1a | 2.89980245 | 1.68997406 | 17.91724 | 7.63E-05 | 0.000397 | F | F | F | F |
| Lrig1 | -1.1343725 | 5.50880287 | 17.9098747 | 7.65E-05 | 0.000398 | F | F | T | F |
| Bglap3 | 1.89924227 | 3.11432583 | 17.9067312 | 7.66E-05 | 0.000398 | F | F | F | F |
| Svip | -8.3671839 | 0.87703165 | 17.9049371 | 7.66E-05 | 0.000398 | T | T | T | T |
| Zmat2 | -1.0785178 | 5.84250127 | 17.8962693 | 7.69E-05 | 0.0004 | F | F | T | F |
| Kctd5 | -1.018807 | 6.21020379 | 17.8592363 | 7.81E-05 | 0.000406 | F | F | F | F |
| Nckap5 | -3.9154666 | 1.72639742 | 17.8546862 | 7.82E-05 | 0.000406 | F | F | T | F |
| Atp9b | 1.08716782 | 5.39254949 | 17.8487491 | 7.84E-05 | 0.000407 | T | F | T | T |
| Whamm | 1.78585034 | 3.33364281 | 17.8423932 | 7.86E-05 | 0.000408 | F | F | T | T |
| Pip4p2 | -1.2010302 | 5.1952171 | 17.8408356 | 7.87E-05 | 0.000408 | T | F | F | T |
| Uhrf1bp1 | 0.84479538 | 6.94167171 | 17.8084304 | 7.97E-05 | 0.000414 | T | F | T | F |
| Ascc1 | -1.5868859 | 4.13937431 | 17.8062559 | 7.98E-05 | 0.000414 | F | F | F | F |
| Spout1 | -1.1386206 | 5.47028541 | 17.8044439 | 7.99E-05 | 0.000414 | F | F | F | F |
| Dennd6a | -1.0258722 | 6.17517155 | 17.8021432 | 7.99E-05 | 0.000415 | T | F | F | T |
| Rab14 | 0.84630075 | 6.97019782 | 17.7923562 | 8.03E-05 | 0.000416 | T | T | T | T |
| Ccdc141 | -3.1195091 | 2.29973087 | 17.7906335 | 8.03E-05 | 0.000416 | F | F | F | T |
| Abcd1 | 1.47017903 | 3.97539658 | 17.7838598 | 8.05E-05 | 0.000417 | T | F | T | T |
| Rbfox1 | -8.4522113 | 0.94156322 | 17.7742494 | 8.09E-05 | 0.000419 | T | T | F | T |
| Urah | -8.3252054 | 0.86312507 | 17.7686554 | 8.10E-05 | 0.00042 | F | F | F | T |
| Kcnn2 | -8.3232124 | 0.85968892 | 17.7620886 | 8.13E-05 | 0.000421 | F | F | F | F |
| Ccdc116 | 2.27185023 | 2.50467647 | 17.7181664 | 8.27E-05 | 0.000428 | T | F | T | F |
| Cnnm1 | -8.3206902 | 0.84853341 | 17.7175956 | 8.28E-05 | 0.000428 | F | F | F | F |
| Atp2b1 | 0.78463717 | 7.70033389 | 17.7013335 | 8.33E-05 | 0.000431 | T | F | T | T |
| Pigo | 0.98735988 | 5.8129197 | 17.6945508 | 8.35E-05 | 0.000432 | F | F | F | F |
| Galnt13 | -8.4385356 | 0.929229 | 17.6683762 | 8.44E-05 | 0.000437 | T | F | T | T |
| Gabpb2 | 0.81734325 | 7.24416837 | 17.6668377 | 8.45E-05 | 0.000437 | T | F | T | T |
| Rxrg | -2.5861339 | 2.76770772 | 17.6578916 | 8.48E-05 | 0.000438 | F | F | F | F |
| Sumf2 | 1.03149978 | 5.71087251 | 17.6345407 | 8.56E-05 | 0.000443 | F | F | T | F |
| Tmprss6 | 3.85783526 | 0.89269948 | 17.6296195 | 8.58E-05 | 0.000443 | F | F | F | F |
| Mccc2 | -1.3473792 | 4.66809993 | 17.6253255 | 8.59E-05 | 0.000444 | T | F | F | F |

| | | | | | | | | | |
|---|---|---|---|---|---|---|---|---|---|
| Zdhhc5 | 0.79817225 | 7.44152154 | 17.6214832 | 8.61E-05 | 0.000445 | T | F | F | T |
| Sergef | -8.3913971 | 0.94189812 | 17.6182328 | 8.62E-05 | 0.000445 | F | F | T | F |
| Tnip2 | 2.6538372 | 1.91928588 | 17.6046295 | 8.67E-05 | 0.000447 | F | F | T | T |
| H2bc3 | -0.7968315 | 8.5327006 | 17.5949075 | 8.70E-05 | 0.000449 | F | F | F | F |
| Pign | 1.068689 | 5.33080789 | 17.5718254 | 8.79E-05 | 0.000453 | T | F | T | T |
| D930048N1 | 2.08367779 | 2.78344095 | 17.5621474 | 8.82E-05 | 0.000455 | T | F | F | F |
| Kpna1 | 0.88896053 | 6.56116042 | 17.5604635 | 8.83E-05 | 0.000455 | T | F | T | T |
| Dazap2 | 1.02619623 | 5.51951549 | 17.5521047 | 8.86E-05 | 0.000457 | T | F | F | T |
| Srsf5 | -0.8545914 | 7.52235401 | 17.5497059 | 8.87E-05 | 0.000457 | T | F | F | T |
| 2410141K0 | 4.07431056 | 0.73924547 | 17.5470609 | 8.88E-05 | 0.000457 | F | F | F | F |
| Atp1a1 | 0.69230179 | 9.61800031 | 17.5442853 | 8.89E-05 | 0.000458 | F | F | T | F |
| Zfp174 | -8.3260395 | 0.84271847 | 17.5279398 | 8.95E-05 | 0.000461 | T | F | T | T |
| Sgk3 | -1.2461228 | 4.99807715 | 17.5253763 | 8.95E-05 | 0.000461 | T | F | F | F |
| Nln | 0.96602066 | 5.93159656 | 17.5166989 | 8.99E-05 | 0.000463 | T | F | T | T |
| Agpat2 | 1.33641025 | 4.2485906 | 17.5065499 | 9.02E-05 | 0.000465 | F | F | F | T |
| Gm2004 | 2.77517923 | 1.92377068 | 17.4849972 | 9.10E-05 | 0.000469 | F | F | T | F |
| Enox1 | -8.3015488 | 0.83152665 | 17.4719825 | 9.15E-05 | 0.000471 | T | F | F | F |
| Necap1 | 1.03908545 | 5.41144531 | 17.4709555 | 9.16E-05 | 0.000471 | F | F | T | F |
| H1f1 | -0.7374807 | 9.39173578 | 17.4703108 | 9.16E-05 | 0.000471 | F | F | F | F |
| Pde4a | -2.6054984 | 2.73765063 | 17.4493895 | 9.24E-05 | 0.000475 | T | F | F | T |
| Cad | -0.7263295 | 9.55813582 | 17.4442837 | 9.26E-05 | 0.000476 | F | F | F | F |
| Psors1c2 | -3.3525504 | 2.30047342 | 17.4371491 | 9.29E-05 | 0.000477 | F | F | F | F |
| Srd5a2 | -8.3686401 | 0.87039616 | 17.427156 | 9.32E-05 | 0.000479 | F | F | F | F |
| D130040H2 | 2.34544724 | 2.34955986 | 17.4167374 | 9.36E-05 | 0.000481 | F | F | F | T |
| Zfp553 | -0.9654421 | 6.46798864 | 17.41543 | 9.37E-05 | 0.000481 | F | F | F | F |
| Mcm4 | -0.7907034 | 8.28768536 | 17.4147979 | 9.37E-05 | 0.000481 | T | T | F | T |
| Calcoco1 | -1.1692336 | 5.27576509 | 17.4094957 | 9.39E-05 | 0.000482 | T | F | F | F |
| Slc25a10 | 0.96243265 | 6.10132924 | 17.4011521 | 9.42E-05 | 0.000484 | F | F | T | F |
| Efna3 | -2.9126254 | 2.41449732 | 17.3995821 | 9.43E-05 | 0.000484 | F | F | F | F |
| Hps3 | 1.49541981 | 3.96337808 | 17.3976173 | 9.44E-05 | 0.000484 | T | F | F | T |
| C2cd3 | -0.9722134 | 6.41938702 | 17.3970623 | 9.44E-05 | 0.000484 | T | F | F | F |
| Plppr3 | -8.3871622 | 0.93946809 | 17.3918276 | 9.46E-05 | 0.000485 | F | F | F | F |
| Lpin3 | 1.29388843 | 4.3245141 | 17.3872426 | 9.48E-05 | 0.000486 | F | F | F | F |
| Txnip | -1.2955913 | 4.78780609 | 17.3687375 | 9.55E-05 | 0.000489 | F | F | T | F |
| Urb2 | 1.02010094 | 5.56891191 | 17.3636031 | 9.57E-05 | 0.00049 | T | F | F | T |
| Stra6l | -8.2914211 | 0.82597204 | 17.3629068 | 9.57E-05 | 0.00049 | F | F | F | F |
| Rab3a | 3.20237922 | 1.3131922 | 17.3506375 | 9.62E-05 | 0.000493 | F | F | F | F |
| Tmc4 | 1.5278928 | 3.81083641 | 17.3421269 | 9.66E-05 | 0.000494 | F | F | F | F |
| Recql4 | -1.1428812 | 5.37580654 | 17.3284042 | 9.71E-05 | 0.000497 | F | F | F | F |
| Dgcr6 | -1.4726413 | 4.29670592 | 17.3279211 | 9.71E-05 | 0.000497 | F | F | F | F |
| Gm14412 | 3.29747105 | 1.24385301 | 17.3219782 | 9.74E-05 | 0.000498 | F | F | F | F |
| Slc5a7 | -8.3857462 | 0.88258221 | 17.3177594 | 9.75E-05 | 0.000499 | T | F | T | T |

| | | | | | | | | | |
|---|---|---|---|---|---|---|---|---|---|
| Slc22a17 | -3.0181026 | 2.29406818 | 17.3145208 | 9.77E-05 | 0.000499 | T | F | F | F |
| Lgals3bp | 2.55930814 | 2.18556016 | 17.3005681 | 9.82E-05 | 0.000502 | F | F | F | F |
| Gcsh | -0.9392276 | 6.64019603 | 17.2913055 | 9.86E-05 | 0.000504 | F | F | F | F |
| Shd | -5.2994126 | 1.23404513 | 17.2890815 | 9.87E-05 | 0.000504 | F | F | F | F |
| Snapc4 | 0.98729963 | 5.8845459 | 17.2850633 | 9.89E-05 | 0.000505 | F | F | F | F |
| Pkib | 4.98972501 | 0.34505585 | 17.2780573 | 9.91E-05 | 0.000506 | T | F | T | T |
| Slc39a1 | 0.83631018 | 6.91126314 | 17.2755308 | 9.93E-05 | 0.000507 | F | F | F | F |
| Lrrc3b | -8.2863199 | 0.81706932 | 17.2684785 | 9.95E-05 | 0.000508 | F | F | F | T |
| Edaradd | 3.06993094 | 1.42608317 | 17.2505341 | 0.0001 | 0.000512 | F | F | F | F |
| Cfb | -3.5162184 | 1.91041631 | 17.2416318 | 0.0001 | 0.000514 | F | F | F | F |
| H2bc7 | -0.7923063 | 9.02257578 | 17.2304988 | 0.0001 | 0.000516 | F | F | F | F |
| Ms4a6d | 5.20198714 | 0.27162183 | 17.2294474 | 0.0001 | 0.000516 | F | F | F | F |
| Dnah1 | -8.2804394 | 0.81618494 | 17.2211344 | 0.0001 | 0.000518 | F | F | F | F |
| G6pc3 | 1.16679403 | 4.80982748 | 17.217178 | 0.0001 | 0.000518 | F | F | F | F |
| Smoc2 | -3.2741662 | 2.24876577 | 17.2142556 | 0.0001 | 0.000519 | T | F | F | F |
| Tox2 | 1.8278004 | 3.19567326 | 17.2120392 | 0.0001 | 0.000519 | F | F | F | T |
| Nudt22 | 1.79070979 | 3.24126138 | 17.2030056 | 0.0001 | 0.000521 | F | F | F | F |
| Ube2c | -0.8638705 | 7.11460272 | 17.1848982 | 0.0001 | 0.000525 | T | F | F | F |
| Gna13 | 0.8297743 | 6.96509956 | 17.1773782 | 0.0001 | 0.000526 | T | F | T | F |
| Amhr2 | -3.5417872 | 1.92290768 | 17.1765473 | 0.0001 | 0.000526 | F | F | F | F |
| Asz1 | 6.02915529 | 0.08207168 | 17.1529632 | 0.0001 | 0.000531 | T | F | F | F |
| Prrc1 | -1.0964068 | 5.57253354 | 17.1425412 | 0.0001 | 0.000534 | T | F | F | T |
| Cacng8 | -3.0476238 | 2.3031544 | 17.1419958 | 0.0001 | 0.000534 | F | F | T | F |
| Ptgs1 | -8.4366165 | 0.98913866 | 17.1407926 | 0.0001 | 0.000534 | T | F | F | T |
| Tgfb2 | -2.1612263 | 3.28322786 | 17.1385493 | 0.00011 | 0.000534 | T | T | T | T |
| Osbpl5 | 1.74819136 | 3.31119661 | 17.131845 | 0.00011 | 0.000535 | T | F | T | T |
| Sncaip | -8.2756065 | 0.81780862 | 17.125631 | 0.00011 | 0.000537 | F | F | F | F |
| Lamp2 | 0.99886988 | 5.64071949 | 17.1250575 | 0.00011 | 0.000537 | T | F | T | T |
| BC003965 | 1.20646081 | 4.6132118 | 17.1242428 | 0.00011 | 0.000537 | F | F | T | F |
| 1700021F0: | 1.35109372 | 4.17642149 | 17.1186155 | 0.00011 | 0.000538 | F | F | F | F |
| Trappc8 | 0.89473753 | 6.42066428 | 17.1082895 | 0.00011 | 0.00054 | F | F | T | T |
| Mmp16 | -3.0395737 | 2.24515798 | 17.061802 | 0.00011 | 0.00055 | T | F | T | T |
| Ppp1r14c | -5.2243465 | 1.18774896 | 17.0614716 | 0.00011 | 0.00055 | T | F | F | F |
| Uhrf1 | -0.7430207 | 9.05007858 | 17.0601648 | 0.00011 | 0.000551 | F | F | T | F |
| Camk1d | -3.4851261 | 1.90117692 | 17.0476029 | 0.00011 | 0.000553 | T | F | T | T |
| Utf1 | 1.40638984 | 5.47009859 | 17.0455647 | 0.00011 | 0.000554 | F | F | F | F |
| Rbm38 | -1.1225868 | 5.41959781 | 17.0361488 | 0.00011 | 0.000556 | F | F | T | F |
| Mark1 | -1.5829801 | 4.07468119 | 17.0220936 | 0.00011 | 0.000559 | F | F | T | T |
| Gpd1l | -1.2984128 | 4.73612028 | 17.0059057 | 0.00011 | 0.000562 | F | F | F | F |
| Thy1 | -3.0172841 | 2.27751575 | 17.004287 | 0.00011 | 0.000563 | T | F | F | T |
| Zmym4 | -0.8488566 | 7.29901325 | 17.0002756 | 0.00011 | 0.000563 | F | F | T | F |
| Xrn2 | 0.78194383 | 7.49857901 | 16.9949967 | 0.00011 | 0.000565 | F | F | F | F |

| | | | | | | | | | |
|---|---|---|---|---|---|---|---|---|---|
| Snx1 | 0.92954775 | 6.12808664 | 16.9806918 | 0.00011 | 0.000568 | T | F | F | F |
| Sfrp5 | -2.739244 | 2.86642446 | 16.9782414 | 0.00011 | 0.000568 | F | F | T | F |
| Cacna1b | -2.16681 | 3.23144123 | 16.9648559 | 0.00011 | 0.000571 | T | F | T | T |
| Fam217b | 1.77559963 | 3.25079218 | 16.9145381 | 0.00012 | 0.000583 | F | F | T | T |
| Atp1a2 | -2.9343744 | 2.32803021 | 16.9144228 | 0.00012 | 0.000583 | T | F | T | T |
| Osbpl2 | 1.38695651 | 4.090317 | 16.9128665 | 0.00012 | 0.000583 | T | F | T | F |
| Pphln1 | -0.8710331 | 7.28614904 | 16.9028336 | 0.00012 | 0.000586 | T | F | T | T |
| Trafd1 | 1.32160854 | 4.24632828 | 16.902518 | 0.00012 | 0.000586 | T | F | T | T |
| Ubr2 | 0.79049204 | 7.42415631 | 16.881862 | 0.00012 | 0.00059 | F | F | T | F |
| Ankrd45 | -2.8079292 | 2.4583241 | 16.8801662 | 0.00012 | 0.000591 | F | F | F | F |
| Nbl1 | 1.78761939 | 3.21671628 | 16.8795883 | 0.00012 | 0.000591 | T | F | T | T |
| Hmgb1 | -0.775682 | 8.33830329 | 16.8776806 | 0.00012 | 0.000591 | T | F | F | F |
| Cbx2 | -0.8470026 | 7.24752676 | 16.8768122 | 0.00012 | 0.000591 | T | F | T | T |
| Tti1 | -1.0314357 | 6.03506566 | 16.8761576 | 0.00012 | 0.000591 | F | F | F | F |
| Rmnd5a | -0.8847285 | 6.99208816 | 16.8699492 | 0.00012 | 0.000593 | F | F | T | T |
| Znfx1 | 0.9763204 | 5.72877399 | 16.8666002 | 0.00012 | 0.000593 | F | F | T | F |
| Mcm5 | -0.7567485 | 8.71668009 | 16.8665515 | 0.00012 | 0.000593 | F | F | T | F |
| E2f8 | 0.9259 | 6.1202235 | 16.8602027 | 0.00012 | 0.000595 | T | F | F | F |
| Ankrd52 | -1.0329733 | 6.00257254 | 16.8593834 | 0.00012 | 0.000595 | T | F | T | T |
| Zfp493 | -2.8098268 | 2.45557873 | 16.8571582 | 0.00012 | 0.000595 | T | F | T | F |
| Chst3 | -2.1290864 | 3.24479499 | 16.8525691 | 0.00012 | 0.000596 | T | F | T | F |
| Plp2 | 0.84631118 | 6.75578924 | 16.8463212 | 0.00012 | 0.000597 | F | F | F | T |
| Adck1 | 1.28534627 | 4.32534848 | 16.8343913 | 0.00012 | 0.0006 | F | F | F | F |
| Bphl | -3.1712972 | 2.15005854 | 16.8279808 | 0.00012 | 0.000602 | F | F | F | F |
| Ccdc34 | -1.2445526 | 4.89241198 | 16.8273091 | 0.00012 | 0.000602 | T | F | F | F |
| Inafm1 | 2.26757609 | 2.40620503 | 16.8229077 | 0.00012 | 0.000603 | F | F | T | T |
| Bhlhe40 | 2.6125519 | 1.89386574 | 16.822129 | 0.00012 | 0.000603 | T | F | T | F |
| Foxo6 | -8.2511119 | 0.78489911 | 16.8185702 | 0.00012 | 0.000603 | T | F | F | F |
| Srsf12 | -8.2511119 | 0.78489911 | 16.8185702 | 0.00012 | 0.000603 | F | F | F | F |
| Mrpl11 | -1.0558527 | 5.76120686 | 16.8103161 | 0.00012 | 0.000605 | F | F | T | F |
| Hirip3 | -1.1046338 | 5.49407929 | 16.8101747 | 0.00012 | 0.000605 | F | F | F | F |
| Gsdma | 6.24769246 | 0.00741535 | 16.8011093 | 0.00012 | 0.000607 | T | F | T | F |
| Arxes2 | -1.4401321 | 4.30559879 | 16.7963768 | 0.00012 | 0.000608 | F | F | T | T |
| Cyp7b1 | -8.3072172 | 0.81513405 | 16.7903306 | 0.00012 | 0.00061 | F | F | F | T |
| Rassf9 | -8.258513 | 0.78436664 | 16.7812685 | 0.00012 | 0.000612 | F | F | F | T |
| Chmp1b | 1.03934321 | 5.27411612 | 16.7496786 | 0.00012 | 0.00062 | F | F | F | F |
| Klhl24 | -1.0220913 | 6.15154967 | 16.7376178 | 0.00012 | 0.000623 | T | F | T | T |
| Zfp607a | -1.5921487 | 4.06943246 | 16.7334686 | 0.00012 | 0.000624 | F | F | T | T |
| Tm6sf1 | -8.2694376 | 0.78822491 | 16.7218643 | 0.00012 | 0.000627 | T | F | T | F |
| Camkk1 | -8.2467626 | 0.79431328 | 16.7176667 | 0.00013 | 0.000628 | T | F | T | T |
| Minar1 | -2.3337477 | 2.95619675 | 16.7150412 | 0.00013 | 0.000628 | T | F | F | F |
| Atp8a2 | -2.0874994 | 3.24657833 | 16.712843 | 0.00013 | 0.000628 | T | F | F | T |

| | | | | | | | | | |
|---|---|---|---|---|---|---|---|---|---|
| Dlst | 0.80204608 | 7.19806809 | 16.7126295 | 0.00013 | 0.000628 | T | F | F | F |
| Erp44 | 0.91784081 | 6.16810926 | 16.7049897 | 0.00013 | 0.00063 | T | F | T | T |
| Ntm | -2.8335769 | 2.42164716 | 16.6852474 | 0.00013 | 0.000635 | T | F | T | T |
| Znrd1 | -1.5161322 | 4.17093705 | 16.6677127 | 0.00013 | 0.00064 | F | F | F | F |
| Vps37a | 0.95105365 | 5.98662676 | 16.6662952 | 0.00013 | 0.00064 | T | F | T | T |
| Tpst1 | -1.222131 | 4.94781602 | 16.6638212 | 0.00013 | 0.000641 | T | F | F | F |
| Cyb5rl | -4.2586009 | 1.48815715 | 16.6581662 | 0.00013 | 0.000642 | F | F | T | F |
| Lyz2 | 5.5986258 | 0.12642135 | 16.6567706 | 0.00013 | 0.000642 | F | F | F | T |
| Foxn4 | -8.2378749 | 0.7837098 | 16.6462355 | 0.00013 | 0.000645 | F | F | T | T |
| Scmh1 | -1.5908743 | 4.0486423 | 16.6448466 | 0.00013 | 0.000645 | T | F | T | T |
| Zfp729a | 1.14936851 | 4.77750597 | 16.6175057 | 0.00013 | 0.000652 | T | F | F | F |
| Ttn | -3.1353676 | 2.26560232 | 16.6130413 | 0.00013 | 0.000653 | T | F | F | T |
| Zfand1 | -1.8602808 | 3.56172083 | 16.6003781 | 0.00013 | 0.000657 | T | F | T | F |
| Tars | 0.74587962 | 7.94512364 | 16.5967531 | 0.00013 | 0.000658 | F | F | F | F |
| Ccn4 | -4.5924208 | 1.45844659 | 16.583464 | 0.00013 | 0.000661 | T | F | T | F |
| Cab39 | 0.77976136 | 7.35521853 | 16.5793735 | 0.00013 | 0.000662 | T | F | T | T |
| Ano5 | -8.3426482 | 0.84009711 | 16.578938 | 0.00013 | 0.000662 | T | F | T | T |
| Tmc6 | 1.32884482 | 4.24529336 | 16.5787912 | 0.00013 | 0.000662 | F | F | F | T |
| Dnajc2 | 0.788374 | 7.20959838 | 16.5780734 | 0.00013 | 0.000662 | F | F | F | F |
| Rflnb | -1.6216839 | 3.96480606 | 16.5622291 | 0.00013 | 0.000666 | F | F | T | T |
| Tshz3 | -3.6744882 | 1.81601584 | 16.5536463 | 0.00013 | 0.000668 | F | F | F | F |
| Pfn2 | -1.8623821 | 3.55699771 | 16.552201 | 0.00013 | 0.000669 | F | F | T | F |
| Adpgk | 0.95835042 | 5.79499955 | 16.5480406 | 0.00013 | 0.00067 | T | F | F | F |
| Glb1 | 1.07360009 | 5.07947012 | 16.5178488 | 0.00014 | 0.000678 | T | F | F | F |
| Bend7 | -3.3372036 | 1.95498964 | 16.5160615 | 0.00014 | 0.000678 | F | F | T | F |
| Bex1 | -0.8362694 | 7.33947004 | 16.5041197 | 0.00014 | 0.000682 | F | F | F | F |
| Arf5 | 0.90151975 | 6.25137379 | 16.5026824 | 0.00014 | 0.000682 | F | F | T | F |
| Slc35c2 | 1.0200626 | 5.40274306 | 16.4812337 | 0.00014 | 0.000688 | F | F | T | F |
| Ypel5 | 1.00651839 | 5.84362684 | 16.4762612 | 0.00014 | 0.000689 | T | F | T | T |
| Ube2f | 0.94863434 | 5.84181536 | 16.4662578 | 0.00014 | 0.000692 | F | F | F | T |
| Fus | -0.753826 | 8.67199929 | 16.4654805 | 0.00014 | 0.000692 | T | F | F | F |
| Azi2 | 1.00644928 | 5.47487603 | 16.4573538 | 0.00014 | 0.000694 | T | T | T | T |
| Nr2c1 | -1.5642074 | 4.05829759 | 16.4526205 | 0.00014 | 0.000695 | T | F | T | F |
| Eci2 | 1.16919049 | 4.72332286 | 16.4462227 | 0.00014 | 0.000697 | F | F | F | F |
| Rec8 | -4.0949761 | 1.5086615 | 16.4387603 | 0.00014 | 0.000699 | F | F | F | F |
| Tmem87a | 1.00634328 | 5.50535648 | 16.4312759 | 0.00014 | 0.000701 | T | F | F | T |
| H4c2 | -0.7832848 | 8.2078234 | 16.4119113 | 0.00014 | 0.000707 | F | F | F | F |
| Myom3 | 3.14005708 | 1.29379317 | 16.4081205 | 0.00014 | 0.000708 | T | F | F | F |
| Selenos | 1.01602666 | 5.42373433 | 16.3926093 | 0.00014 | 0.000712 | F | F | T | F |
| Cd46 | 3.95979918 | 0.70525077 | 16.3757032 | 0.00014 | 0.000717 | F | F | F | T |
| Rhod | -2.3267489 | 2.9104597 | 16.3569487 | 0.00015 | 0.000722 | F | F | F | F |
| Ankfy1 | 0.77241943 | 7.38689634 | 16.3475688 | 0.00015 | 0.000725 | T | T | T | T |

| | | | | | | | | | |
|---|---|---|---|---|---|---|---|---|---|
| Upf3b | -1.5441973 | 4.0959447 | 16.3453048 | 0.00015 | 0.000726 | T | F | F | F |
| Tpd52 | 0.91457638 | 6.11505368 | 16.3433214 | 0.00015 | 0.000726 | T | F | T | T |
| Tmem184c | -1.2739679 | 4.72363715 | 16.3425986 | 0.00015 | 0.000726 | F | F | F | T |
| Fam126a | 0.94268252 | 5.89033834 | 16.3273795 | 0.00015 | 0.00073 | T | F | T | T |
| Elf1 | -1.4630344 | 4.24819742 | 16.326418 | 0.00015 | 0.000731 | T | F | T | F |
| Kcnab3 | -8.21922 | 0.74962947 | 16.3224256 | 0.00015 | 0.000732 | F | F | F | F |
| Cul9 | -4.5339047 | 1.32851029 | 16.3147033 | 0.00015 | 0.000734 | F | F | F | F |
| Cmtr2 | -1.8126242 | 3.6104911 | 16.3145979 | 0.00015 | 0.000734 | F | F | F | F |
| Creb3l2 | 0.99059277 | 5.76303609 | 16.3069117 | 0.00015 | 0.000736 | T | F | F | T |
| Klf16 | 0.87435745 | 6.40815003 | 16.3008645 | 0.00015 | 0.000738 | F | F | T | T |
| Pdcd4 | 0.93387054 | 5.93136074 | 16.2935205 | 0.00015 | 0.00074 | T | F | F | T |
| Rhot2 | -0.9681234 | 6.23340601 | 16.2895559 | 0.00015 | 0.000741 | T | F | F | F |
| Chrnb4 | -3.065301 | 2.14148768 | 16.2879905 | 0.00015 | 0.000741 | F | F | F | F |
| Dop1b | 1.22652905 | 4.43393419 | 16.2660967 | 0.00015 | 0.000748 | F | F | F | F |
| Wnt9a | 2.833437 | 1.57026701 | 16.2538032 | 0.00015 | 0.000751 | T | F | T | T |
| Map3k5 | -1.3268908 | 4.8243579 | 16.2523556 | 0.00015 | 0.000752 | F | F | F | F |
| Tubgcp2 | -1.1449398 | 5.26920269 | 16.2521994 | 0.00015 | 0.000752 | F | F | F | F |
| Bdh2 | -8.2211968 | 0.74730674 | 16.2429273 | 0.00015 | 0.000754 | F | F | F | F |
| Mtf2 | -1.7594718 | 3.92750643 | 16.2406021 | 0.00015 | 0.000755 | F | F | F | T |
| Ccdc85c | -0.8304237 | 7.26923912 | 16.2352873 | 0.00015 | 0.000756 | F | F | T | T |
| Pak6 | -4.0795374 | 1.58114854 | 16.2260702 | 0.00015 | 0.000759 | F | F | F | T |
| Adgb | -8.2007257 | 0.74243109 | 16.2158904 | 0.00015 | 0.000762 | T | F | F | F |
| Vps37d | -4.480892 | 1.33298735 | 16.1950128 | 0.00016 | 0.000769 | F | F | T | F |
| Pitpna | 0.86481957 | 6.55842976 | 16.1902053 | 0.00016 | 0.00077 | T | F | T | T |
| Gal3st4 | -8.2078884 | 0.73930555 | 16.1846213 | 0.00016 | 0.000772 | F | F | F | F |
| Gpd2 | 0.84357704 | 6.73856305 | 16.1784735 | 0.00016 | 0.000774 | T | F | F | F |
| Lifr | 0.84213598 | 6.77733146 | 16.1725444 | 0.00016 | 0.000775 | T | F | T | T |
| Ttc23 | -1.3590807 | 4.44195525 | 16.1692343 | 0.00016 | 0.000776 | F | F | F | F |
| Zmynd8 | -0.7790389 | 8.1153164 | 16.164324 | 0.00016 | 0.000778 | F | F | T | F |
| Carmil3 | -8.2015404 | 0.75384032 | 16.1516115 | 0.00016 | 0.000782 | F | F | F | F |
| Polh | -1.5943618 | 3.97038922 | 16.144013 | 0.00016 | 0.000784 | F | F | T | T |
| Pgpep1 | -1.6133678 | 3.95214187 | 16.1354984 | 0.00016 | 0.000787 | T | F | T | T |
| Pknox1 | 1.07570218 | 5.01731138 | 16.1245559 | 0.00016 | 0.00079 | T | F | T | F |
| Doc2g | -8.2213514 | 0.7436887 | 16.1226821 | 0.00016 | 0.000791 | F | F | F | F |
| Nptx1 | -2.9532053 | 2.22110798 | 16.1217859 | 0.00016 | 0.000791 | T | F | T | T |
| Ptpn1 | -0.9855426 | 6.06238246 | 16.1035667 | 0.00016 | 0.000797 | T | F | T | T |
| Aifm2 | 2.82669122 | 1.72615452 | 16.0799728 | 0.00016 | 0.000804 | F | F | T | F |
| Ano10 | 1.25067194 | 4.354604 | 16.0780381 | 0.00016 | 0.000805 | T | F | F | T |
| Rnf125 | 3.01373517 | 1.37120599 | 16.0748324 | 0.00016 | 0.000806 | F | F | F | F |
| Olfr1307 | 0.96831876 | 6.06089793 | 16.0669546 | 0.00016 | 0.000808 | F | F | F | F |
| Med14 | 0.84075896 | 6.94822537 | 16.0646346 | 0.00016 | 0.000809 | T | F | T | T |
| Gulp1 | -2.284899 | 2.92505733 | 16.0637472 | 0.00016 | 0.000809 | T | F | T | T |

| | | | | | | | | | |
|---|---|---|---|---|---|---|---|---|---|
| Chaf1a | -0.8239649 | 7.26107732 | 16.0619528 | 0.00016 | 0.000809 | T | F | T | F |
| Ctps2 | -1.2228185 | 4.85470539 | 16.0462648 | 0.00017 | 0.000814 | F | F | T | F |
| Klhl9 | -0.832996 | 7.16835401 | 16.0392057 | 0.00017 | 0.000817 | T | F | F | F |
| Pde12 | 0.95107203 | 5.72865231 | 16.0312392 | 0.00017 | 0.000819 | F | F | F | T |
| Timd2 | -4.4687798 | 1.33217502 | 16.027638 | 0.00017 | 0.00082 | T | F | T | T |
| Csrnp1 | 1.37860551 | 4.01816791 | 16.0262427 | 0.00017 | 0.000821 | T | F | F | T |
| Zfp3 | 2.00434706 | 2.74854031 | 16.0164303 | 0.00017 | 0.000824 | T | F | F | T |
| Ube2h | 0.84143383 | 6.64304484 | 16.0158198 | 0.00017 | 0.000824 | T | T | T | T |
| Daglb | 1.16686035 | 4.65000842 | 16.000101 | 0.00017 | 0.000829 | T | T | F | T |
| Spg20 | -1.0325884 | 5.70661483 | 15.9933146 | 0.00017 | 0.000831 | T | F | T | F |
| Swap70 | -1.1137872 | 5.2632447 | 15.9895403 | 0.00017 | 0.000832 | T | F | T | T |
| Tesk2 | -3.0921885 | 2.15454486 | 15.9746355 | 0.00017 | 0.000837 | F | F | F | F |
| Oprk1 | -8.180702 | 0.72756468 | 15.9743287 | 0.00017 | 0.000837 | F | F | T | T |
| Naaa | -1.7428364 | 3.69417828 | 15.9721075 | 0.00017 | 0.000838 | T | F | F | F |
| Zfp865 | -1.3313315 | 4.53770852 | 15.9692706 | 0.00017 | 0.000839 | F | F | F | F |
| Lama1 | -0.854511 | 6.97642319 | 15.9684367 | 0.00017 | 0.000839 | F | F | F | F |
| Akap13 | -0.8191684 | 7.32297108 | 15.9669284 | 0.00017 | 0.000839 | T | F | T | F |
| Prrg3 | -1.5980405 | 3.95634226 | 15.9565089 | 0.00017 | 0.000843 | T | F | T | F |
| Ddx25 | -1.8551306 | 3.4999918 | 15.9504873 | 0.00017 | 0.000845 | F | F | T | T |
| Abcc3 | 6.19799597 | -0.0656643 | 15.947921 | 0.00017 | 0.000845 | F | F | F | F |
| Spg21 | 1.10821816 | 4.8494473 | 15.9450454 | 0.00017 | 0.000846 | T | F | T | T |
| Tmem248 | 0.76015094 | 7.43989308 | 15.9426208 | 0.00017 | 0.000847 | T | T | T | T |
| Gorasp2 | 0.78007905 | 7.33722889 | 15.9404039 | 0.00017 | 0.000847 | F | F | T | F |
| Vps35l | 0.88483174 | 6.26538648 | 15.9403763 | 0.00017 | 0.000847 | F | F | F | F |
| Gm10251 | 8.18240383 | -0.3225852 | 15.9662534 | 0.00017 | 0.000849 | F | F | F | F |
| Dzip3 | -1.270006 | 4.68699864 | 15.9214812 | 0.00017 | 0.000854 | T | F | F | F |
| Tmem147 | 1.06357634 | 5.11393531 | 15.9152458 | 0.00018 | 0.000856 | F | F | F | F |
| Hoxb5 | -8.3830546 | 0.94304086 | 15.9111551 | 0.00018 | 0.000857 | F | F | T | F |
| Cdt1 | 0.77946319 | 7.18091367 | 15.9044816 | 0.00018 | 0.000859 | T | F | F | F |
| Gng10 | -2.0414696 | 3.24546587 | 15.9000463 | 0.00018 | 0.000861 | T | F | F | T |
| Slc4a8 | 1.28326174 | 4.23894667 | 15.8988321 | 0.00018 | 0.000861 | T | F | T | T |
| Rhox4d | 8.17787151 | -0.3265854 | 15.9158552 | 0.00018 | 0.000866 | F | F | F | F |
| Ednra | -3.3159819 | 1.93966228 | 15.8794851 | 0.00018 | 0.000867 | T | F | T | T |
| Sertm1 | -8.1811545 | 0.7158073 | 15.8793793 | 0.00018 | 0.000867 | T | F | T | T |
| Tekt1 | -4.0685187 | 1.47931228 | 15.8747278 | 0.00018 | 0.000869 | F | F | F | F |
| Ebf3 | -3.7401331 | 1.63584827 | 15.8587164 | 0.00018 | 0.000875 | T | F | T | T |
| Foxr2 | -8.1987937 | 0.7230625 | 15.8561893 | 0.00018 | 0.000875 | F | F | F | F |
| Wdr44 | 1.45836485 | 3.9102835 | 15.855737 | 0.00018 | 0.000875 | T | T | F | T |
| Macrod2 | -8.1794481 | 0.73575775 | 15.8474948 | 0.00018 | 0.000878 | T | F | F | T |
| Map3k1 | 0.79730306 | 6.97821991 | 15.8431428 | 0.00018 | 0.00088 | T | T | T | T |
| Rnf135 | 2.01955448 | 2.71487136 | 15.8318481 | 0.00018 | 0.000884 | F | F | F | F |
| Zdhhc18 | 0.99263824 | 5.40915531 | 15.8280157 | 0.00018 | 0.000885 | F | F | F | T |

| | | | | | | | | | |
|---|---|---|---|---|---|---|---|---|---|
| Sall1 | -0.7907652 | 7.74560161 | 15.8217504 | 0.00018 | 0.000887 | F | F | F | T |
| Slc22a8 | -8.1984572 | 0.72170845 | 15.8150083 | 0.00018 | 0.000889 | F | F | T | F |
| Sfpq | -0.674362 | 9.92225753 | 15.8098276 | 0.00018 | 0.000891 | T | F | F | T |
| Olfr1031 | 2.71132641 | 2.17194137 | 15.8093244 | 0.00018 | 0.000891 | F | F | F | F |
| Lrp8 | 1.04812759 | 5.17844422 | 15.7881177 | 0.00018 | 0.000899 | T | F | T | F |
| Slc8a2 | -2.8664525 | 2.26303892 | 15.7789609 | 0.00019 | 0.000902 | T | F | T | T |
| Xlr | -8.1845018 | 0.71273066 | 15.7743737 | 0.00019 | 0.000904 | F | F | T | F |
| Isyna1 | 0.7720394 | 7.16319464 | 15.7736206 | 0.00019 | 0.000904 | F | F | F | F |
| Twf1 | 0.83647591 | 6.60823996 | 15.768322 | 0.00019 | 0.000906 | F | F | T | F |
| Cfap36 | -1.5173753 | 4.07445508 | 15.7649994 | 0.00019 | 0.000907 | T | F | F | F |
| Chaf1b | -0.9740432 | 6.05818518 | 15.7583587 | 0.00019 | 0.000909 | T | F | F | F |
| Rassf2 | -1.6439004 | 3.85752 | 15.7422877 | 0.00019 | 0.000915 | T | F | F | T |
| Zer1 | -1.1657543 | 5.02350478 | 15.72937 | 0.00019 | 0.00092 | T | F | T | T |
| Relb | -8.1633787 | 0.70369547 | 15.7277359 | 0.00019 | 0.00092 | F | F | F | F |
| Eya4 | -8.15969 | 0.70750925 | 15.7237162 | 0.00019 | 0.000922 | F | F | T | F |
| Ctla2a | 1.43156478 | 3.8760892 | 15.7223175 | 0.00019 | 0.000922 | F | F | T | F |
| Rars | 0.74082207 | 7.73965147 | 15.7115078 | 0.00019 | 0.000926 | F | F | F | F |
| Tmem100 | -8.15844 | 0.70535437 | 15.7084027 | 0.00019 | 0.000927 | F | F | F | T |
| Gnaq | 0.8676776 | 6.33127989 | 15.7007468 | 0.00019 | 0.00093 | F | F | T | F |
| Cuedc1 | 1.44762718 | 4.04009266 | 15.6940375 | 0.00019 | 0.000932 | F | F | T | T |
| Carmil1 | 0.83051306 | 6.63712649 | 15.6849677 | 0.00019 | 0.000936 | T | T | T | F |
| Me1 | 1.11813477 | 4.76397696 | 15.6812746 | 0.00019 | 0.000937 | F | F | T | F |
| Slc22a5 | 1.66616097 | 3.33644723 | 15.678818 | 0.00019 | 0.000938 | T | F | T | T |
| Commd1 | -1.2271615 | 4.80130972 | 15.6777502 | 0.00019 | 0.000938 | F | F | F | T |
| Ap3b1 | 0.81378376 | 6.77490126 | 15.669414 | 0.00019 | 0.000941 | F | F | F | T |
| 1700003E1( | -8.1566796 | 0.69953519 | 15.6670524 | 0.00019 | 0.000942 | F | F | F | F |
| Tecr | -0.8622579 | 6.90480313 | 15.6650702 | 0.00019 | 0.000942 | T | F | F | F |
| Insig1 | 1.10792142 | 4.80249947 | 15.661582 | 0.00019 | 0.000943 | T | F | F | T |
| Eno1 | -0.6106504 | 11.3993096 | 15.6481676 | 0.0002 | 0.000949 | T | T | F | T |
| Nusap1 | -1.0194167 | 5.77326076 | 15.6445268 | 0.0002 | 0.00095 | T | F | F | T |
| Fam81a | -4.2456482 | 1.37246746 | 15.6432089 | 0.0002 | 0.00095 | T | F | T | F |
| Nudt19 | -1.3989984 | 4.2768637 | 15.6396428 | 0.0002 | 0.000951 | F | F | T | F |
| Itsn1 | -0.8123694 | 7.26328789 | 15.6371347 | 0.0002 | 0.000952 | T | F | T | F |
| Cfap300 | -2.5537569 | 2.58409983 | 15.6368473 | 0.0002 | 0.000952 | T | F | F | T |
| Kdelr1 | 0.88051014 | 6.2497991 | 15.636407 | 0.0002 | 0.000952 | T | T | T | T |
| Sh3bgrl | -1.1462113 | 5.08264984 | 15.6272545 | 0.0002 | 0.000955 | T | T | F | T |
| Rtn4 | 0.71109498 | 8.24354685 | 15.6209114 | 0.0002 | 0.000958 | F | F | F | F |
| Tmem175 | 1.37092911 | 3.99267316 | 15.6169592 | 0.0002 | 0.000959 | T | F | T | F |
| Nono | -0.7109147 | 9.04652135 | 15.6168287 | 0.0002 | 0.000959 | T | F | T | F |
| Cep68 | -0.9929798 | 5.89465347 | 15.6162158 | 0.0002 | 0.000959 | T | F | T | F |
| Nectin3 | 0.88936642 | 6.12735498 | 15.6057935 | 0.0002 | 0.000963 | F | F | F | F |
| Mfsd10 | -1.0463492 | 5.56418227 | 15.5938993 | 0.0002 | 0.000968 | F | F | F | F |

| | | | | | | | | | |
|---|---|---|---|---|---|---|---|---|---|
| Ssr4 | 1.03653341 | 5.13188312 | 15.5923406 | 0.0002 | 0.000968 | F | F | F | F |
| Vmn2r29 | 8.14993608 | -0.3511954 | 15.6082342 | 0.0002 | 0.000974 | F | F | F | F |
| Lman2 | 0.74793798 | 7.73125311 | 15.575487 | 0.0002 | 0.000975 | T | F | F | F |
| Mmut | 0.93515373 | 5.81437999 | 15.5733025 | 0.0002 | 0.000975 | F | F | T | T |
| Gm20708 | 8.14949755 | -0.3515811 | 15.6034464 | 0.0002 | 0.000975 | F | F | F | T |
| Zfp786 | -5.0764676 | 1.11814883 | 15.5659414 | 0.0002 | 0.000978 | F | F | F | F |
| Pfkfb1 | 1.59965209 | 3.45143823 | 15.5600317 | 0.0002 | 0.00098 | T | F | F | F |
| Scai | -1.7314509 | 3.73490475 | 15.5513344 | 0.0002 | 0.000984 | T | F | T | T |
| Gadd45a | 2.08622211 | 2.66076304 | 15.5356383 | 0.00021 | 0.00099 | T | F | T | F |
| Zfyve26 | 0.8632786 | 6.33028488 | 15.5268808 | 0.00021 | 0.000993 | F | F | T | T |
| Dnmt3a | -0.7932383 | 7.62957035 | 15.4983263 | 0.00021 | 0.001005 | T | T | T | T |
| Arrdc1 | 1.71053859 | 3.21942841 | 15.4824354 | 0.00021 | 0.001012 | F | F | F | F |
| Efcab1 | -8.2474432 | 0.75134855 | 15.4820493 | 0.00021 | 0.001012 | T | F | F | T |
| Znhit1 | -1.2855951 | 4.57597759 | 15.4759814 | 0.00021 | 0.001014 | F | F | F | F |
| Slc6a1 | -8.1396452 | 0.68487625 | 15.4718186 | 0.00021 | 0.001016 | T | F | T | T |
| Sh3bp4 | -0.9436067 | 6.2434652 | 15.4587577 | 0.00021 | 0.001021 | F | F | T | F |
| Lypd6b | -5.1499268 | 1.07439477 | 15.458749 | 0.00021 | 0.001021 | T | F | F | F |
| Gprin2 | -4.4559518 | 1.30147921 | 15.4507294 | 0.00021 | 0.001024 | F | F | F | F |
| Tbc1d10a | 1.03659305 | 5.18047989 | 15.4485736 | 0.00021 | 0.001025 | F | F | F | T |
| Reck | -1.36114 | 4.32795933 | 15.4480762 | 0.00021 | 0.001025 | T | F | T | T |
| Sec24c | 0.72586734 | 7.85150448 | 15.4399526 | 0.00021 | 0.001028 | T | F | T | F |
| Ano8 | -1.750638 | 3.62057501 | 15.4305408 | 0.00021 | 0.001032 | F | F | F | F |
| Srrm3 | -8.1432298 | 0.68113379 | 15.4282194 | 0.00022 | 0.001033 | T | F | T | F |
| Thsd4 | -1.785359 | 3.81151686 | 15.4239406 | 0.00022 | 0.001035 | T | F | T | T |
| Lrrc27 | -3.5526485 | 1.69223957 | 15.4194472 | 0.00022 | 0.001036 | T | F | F | F |
| Nanog | -1.5253663 | 4.22414637 | 15.4190801 | 0.00022 | 0.001036 | F | F | F | F |
| Palld | -1.0319826 | 5.60944027 | 15.3980947 | 0.00022 | 0.001045 | T | F | T | T |
| Olfr624 | 0.93405238 | 5.78854333 | 15.3969644 | 0.00022 | 0.001046 | F | F | F | F |
| Srsf6 | -0.772182 | 7.98067963 | 15.3881888 | 0.00022 | 0.001049 | F | F | T | T |
| Rai2 | -8.1563617 | 0.68505788 | 15.3874955 | 0.00022 | 0.001049 | F | F | F | F |
| Ergic2 | 0.9023871 | 5.98110744 | 15.3856518 | 0.00022 | 0.00105 | T | F | T | T |
| Tmem254b | 1.24680622 | 4.62998896 | 15.381369 | 0.00022 | 0.001052 | F | F | T | F |
| Trak2 | 0.84368202 | 6.46069441 | 15.3762127 | 0.00022 | 0.001054 | T | T | T | T |
| Arhgef11 | 0.88638396 | 6.17411591 | 15.3631247 | 0.00022 | 0.001059 | F | F | T | F |
| Dpp9 | -0.9020848 | 6.47972057 | 15.362904 | 0.00022 | 0.001059 | T | F | F | F |
| Ociad2 | -2.8920813 | 2.19854162 | 15.3566673 | 0.00022 | 0.001061 | F | F | T | F |
| Tmem230 | 1.28189211 | 4.21368676 | 15.3563681 | 0.00022 | 0.001061 | F | F | F | F |
| Usp14 | 0.81976357 | 6.65544271 | 15.3563026 | 0.00022 | 0.001061 | T | F | T | F |
| Tarbp1 | -1.0664839 | 5.37894019 | 15.3530496 | 0.00022 | 0.001063 | F | F | F | T |
| Rapgef5 | -1.2008994 | 5.01654039 | 15.3440271 | 0.00022 | 0.001066 | F | F | T | T |
| Dnaja2 | 0.72921566 | 7.72085241 | 15.3439308 | 0.00022 | 0.001066 | T | F | T | T |
| Scaf11 | 0.74371237 | 7.49000099 | 15.3399221 | 0.00022 | 0.001068 | F | F | F | T |

| | | | | | | | | | |
|---|---|---|---|---|---|---|---|---|---|
| Draxin | -2.5712232 | 2.6442661 | 15.3376438 | 0.00022 | 0.001069 | T | F | T | T |
| Neurl4 | -0.8667908 | 6.7419105 | 15.321956 | 0.00022 | 0.001076 | F | F | T | F |
| Prr14l | 0.7433741 | 7.50875909 | 15.3160167 | 0.00023 | 0.001078 | T | F | T | T |
| Cfap65 | -4.4120731 | 1.27422144 | 15.3148604 | 0.00023 | 0.001078 | F | F | F | F |
| Fcna | 8.12492309 | -0.3731632 | 15.3371696 | 0.00023 | 0.001082 | F | F | F | F |
| Clip4 | -8.1661082 | 0.68854604 | 15.3047076 | 0.00023 | 0.001083 | F | F | T | T |
| Ntn3 | -3.498076 | 1.90414015 | 15.3035028 | 0.00023 | 0.001083 | T | F | F | T |
| Hecw1 | -8.1636764 | 0.68689208 | 15.300348 | 0.00023 | 0.001084 | T | F | T | T |
| Arf1 | 0.68152504 | 8.75955524 | 15.3003034 | 0.00023 | 0.001084 | F | F | F | F |
| Mmp1b | 3.80072494 | 0.86580901 | 15.2980585 | 0.00023 | 0.001085 | T | F | T | F |
| Ccdc166 | -4.0141661 | 1.43430145 | 15.2951632 | 0.00023 | 0.001086 | F | F | F | F |
| Strbp | -0.8145534 | 7.34360086 | 15.2885318 | 0.00023 | 0.001089 | T | F | T | T |
| Duxbl3 | -8.268283 | 0.76647633 | 15.2880298 | 0.00023 | 0.001089 | F | F | T | T |
| Dexi | -2.3842528 | 2.73556477 | 15.2874606 | 0.00023 | 0.001089 | F | F | F | F |
| Capsl | -5.1373124 | 1.06235341 | 15.2820804 | 0.00023 | 0.001091 | F | F | F | F |
| Ints9 | -0.9656042 | 6.01253017 | 15.2747622 | 0.00023 | 0.001094 | T | F | F | F |
| Suz12 | 0.76040475 | 7.27991818 | 15.2676165 | 0.00023 | 0.001097 | T | F | T | F |
| Rgs10 | -4.7329061 | 1.16399876 | 15.2658602 | 0.00023 | 0.001098 | T | F | F | F |
| Pitpnm1 | 1.22505373 | 4.29589785 | 15.2444861 | 0.00023 | 0.001108 | F | F | F | F |
| Tceal6 | -8.1178358 | 0.66709363 | 15.2276636 | 0.00023 | 0.001115 | F | F | F | F |
| Mprip | 0.70076785 | 8.40389986 | 15.20845 | 0.00024 | 0.001124 | T | F | T | F |
| Lmo7 | -1.1767782 | 4.89869581 | 15.2058079 | 0.00024 | 0.001125 | T | F | F | F |
| Ick | -1.068039 | 5.34216123 | 15.2040879 | 0.00024 | 0.001126 | T | F | F | T |
| Fsd1l | 1.33918619 | 4.06428089 | 15.1976709 | 0.00024 | 0.001128 | T | F | T | T |
| Lect2 | -8.1148927 | 0.66735459 | 15.1976705 | 0.00024 | 0.001128 | T | T | T | T |
| AB124611 | 8.11227708 | -0.3842451 | 15.2016873 | 0.00024 | 0.00114 | F | F | F | F |
| Atg2a | 0.83723383 | 6.49327714 | 15.1593815 | 0.00024 | 0.001146 | T | F | T | F |
| Tmem72 | 0.72498533 | 8.23871711 | 15.155166 | 0.00024 | 0.001148 | T | T | T | T |
| Echdc2 | -1.8743568 | 3.38836747 | 15.1520569 | 0.00024 | 0.00115 | F | F | T | T |
| Cetn4 | -8.1417651 | 0.66891069 | 15.1289924 | 0.00024 | 0.001161 | T | F | F | T |
| Hes5 | -8.10849 | 0.66136382 | 15.1239662 | 0.00024 | 0.001163 | F | F | F | F |
| Pkn1 | 0.95105982 | 5.53137293 | 15.1222016 | 0.00024 | 0.001163 | T | F | F | F |
| Cnot9 | -0.7987818 | 7.24960405 | 15.1105161 | 0.00025 | 0.001169 | T | F | F | F |
| Selenoh | -0.896285 | 6.59200143 | 15.0993485 | 0.00025 | 0.001174 | F | F | F | F |
| Gbp3 | 4.4291213 | 0.43956244 | 15.0984144 | 0.00025 | 0.001175 | T | T | F | T |
| Vps8 | 1.01899161 | 5.11743639 | 15.0967729 | 0.00025 | 0.001175 | T | F | F | F |
| Cdc26 | -1.3266119 | 4.42657815 | 15.0896941 | 0.00025 | 0.001178 | F | F | T | F |
| Ak8 | -8.1332415 | 0.66283742 | 15.0890416 | 0.00025 | 0.001179 | F | F | F | F |
| Wfdc2 | 1.31963144 | 4.08895514 | 15.0859156 | 0.00025 | 0.00118 | F | F | F | F |
| Hmgcl | 1.13716199 | 4.60793848 | 15.0805819 | 0.00025 | 0.001182 | T | F | T | T |
| Trpv6 | -8.1109054 | 0.65336892 | 15.0797394 | 0.00025 | 0.001182 | T | F | T | T |
| Ctsk | 3.77849973 | 0.71880377 | 15.0665597 | 0.00025 | 0.001189 | F | F | T | F |

| | | | | | | | | | |
|---|---|---|---|---|---|---|---|---|---|
| Oscp1 | -2.241163 | 2.87025477 | 15.0528231 | 0.00025 | 0.001196 | F | F | T | F |
| Zbtb6 | 1.114028 | 4.69080454 | 15.0514491 | 0.00025 | 0.001196 | T | F | F | T |
| Fam187b | -8.1038752 | 0.65153787 | 15.0497695 | 0.00025 | 0.001197 | F | F | F | F |
| Ipo7 | 0.68548215 | 8.49196938 | 15.0403705 | 0.00025 | 0.001201 | T | F | F | T |
| Sirt4 | 3.74889881 | 0.72008686 | 15.0373274 | 0.00025 | 0.001202 | F | F | F | F |
| Fdxr | 1.16316367 | 4.49176499 | 15.0301577 | 0.00025 | 0.001206 | F | F | F | F |
| Naprt | -3.043908 | 2.02759098 | 15.0263285 | 0.00026 | 0.001208 | F | F | F | F |
| Oasl1 | 8.09830791 | -0.396467 | 15.0532421 | 0.00026 | 0.001208 | F | F | F | F |
| Gart | -0.7301508 | 8.33620274 | 15.0206081 | 0.00026 | 0.00121 | T | F | F | F |
| Mageh1 | -2.0377302 | 3.13642709 | 15.0099424 | 0.00026 | 0.001215 | F | F | T | F |
| 2610528J11 | -1.5552564 | 3.95962518 | 15.0037295 | 0.00026 | 0.001218 | F | F | F | F |
| Dgkg | -8.1119826 | 0.67767704 | 15.0000661 | 0.00026 | 0.00122 | F | F | T | F |
| St3gal2 | -1.0607214 | 5.3336962 | 14.9949696 | 0.00026 | 0.001222 | F | F | T | T |
| Ttc26 | -1.2636802 | 4.61815654 | 14.9770878 | 0.00026 | 0.001231 | F | F | T | F |
| Dhx32 | -1.0588328 | 5.35923577 | 14.9605923 | 0.00026 | 0.00124 | F | F | F | F |
| Eme1 | -1.0795806 | 5.23483336 | 14.9583136 | 0.00026 | 0.001241 | F | F | F | F |
| Tmx1 | 0.95440087 | 5.4702051 | 14.9551462 | 0.00026 | 0.001242 | F | F | F | F |
| D17H6S53I | 1.08766939 | 4.78241915 | 14.9538709 | 0.00026 | 0.001243 | F | T | F | T |
| Tubb5 | -0.591525 | 11.3153576 | 14.9534506 | 0.00026 | 0.001243 | T | F | T | F |
| Mlf2 | -0.7669221 | 7.6873428 | 14.952473 | 0.00026 | 0.001243 | T | F | F | F |
| Acan | -2.8366483 | 2.23191424 | 14.9452342 | 0.00026 | 0.001246 | F | F | F | F |
| Sprr2a2 | -4.0149419 | 1.44895568 | 14.9406228 | 0.00026 | 0.001249 | F | F | T | T |
| Mypop | -2.2772615 | 2.81254309 | 14.9342748 | 0.00027 | 0.001252 | F | F | F | F |
| Grm8 | -8.1005835 | 0.64211512 | 14.9325637 | 0.00027 | 0.001252 | T | F | T | F |
| Cep135 | -1.1779887 | 4.84408104 | 14.9202085 | 0.00027 | 0.001259 | T | F | F | T |
| Lgi2 | -1.8272033 | 3.44751139 | 14.9179469 | 0.00027 | 0.00126 | F | F | T | F |
| Nat8l | -2.2129649 | 2.95775874 | 14.9134077 | 0.00027 | 0.001262 | T | F | T | F |
| Mlh1 | -1.0914293 | 5.18762631 | 14.9124074 | 0.00027 | 0.001262 | F | F | F | T |
| Mapk1ip1l | -0.7069348 | 8.79289295 | 14.9102502 | 0.00027 | 0.001263 | T | T | T | T |
| Slc25a39 | -0.8072158 | 7.0838233 | 14.9022864 | 0.00027 | 0.001267 | F | F | F | F |
| Dchs2 | -4.3808162 | 1.24785396 | 14.8953917 | 0.00027 | 0.001271 | F | F | F | F |
| Msra | -1.9203393 | 3.2787747 | 14.8910942 | 0.00027 | 0.001273 | T | F | F | T |
| Olig3 | -8.3407284 | 0.90660385 | 14.8583116 | 0.00027 | 0.00129 | T | F | T | T |
| Efnb1 | -0.990771 | 6.29008026 | 14.8463301 | 0.00028 | 0.001297 | F | F | F | F |
| Purb | 0.72980212 | 7.60254276 | 14.8430496 | 0.00028 | 0.001298 | F | F | T | F |
| Iqsec3 | -8.0903477 | 0.65428591 | 14.8404368 | 0.00028 | 0.001299 | T | F | T | T |
| Bdkrb2 | 4.16227025 | 0.4664657 | 14.836027 | 0.00028 | 0.001301 | F | F | F | F |
| Cyp17a1 | 3.40402129 | 0.93512193 | 14.8358322 | 0.00028 | 0.001301 | F | F | F | F |
| Orai1 | 1.21753237 | 4.30765561 | 14.8352047 | 0.00028 | 0.001301 | T | F | F | T |
| Mycbp | 1.157819 | 4.47905546 | 14.8318574 | 0.00028 | 0.001303 | T | F | F | F |
| H2bc6 | -0.7899443 | 7.493147 | 14.829195 | 0.00028 | 0.001304 | T | F | F | T |
| Zfp345 | 2.02551634 | 2.76557226 | 14.8181949 | 0.00028 | 0.00131 | T | F | F | T |

| | | | | | | | | | |
|---|---|---|---|---|---|---|---|---|---|
| Plxdc2 | -1.7131448 | 3.6150216 | 14.8147504 | 0.00028 | 0.001312 | F | T | T | T |
| Sorbs3 | -1.3572537 | 4.291997 | 14.8123383 | 0.00028 | 0.001313 | F | F | F | F |
| Plaat3 | -2.8670486 | 2.14886248 | 14.8100298 | 0.00028 | 0.001314 | T | F | T | F |
| Sod3 | -8.1054608 | 0.675683 | 14.8060756 | 0.00028 | 0.001316 | F | F | F | T |
| Papln | 2.94944573 | 1.31048544 | 14.8039353 | 0.00028 | 0.001317 | T | F | T | F |
| H2bc12 | -0.7062929 | 8.89054739 | 14.7968824 | 0.00028 | 0.00132 | F | F | F | T |
| Pde1c | -3.0303542 | 2.01207709 | 14.7769895 | 0.00028 | 0.001331 | T | T | T | T |
| 2610044O1 | -1.196611 | 4.84311785 | 14.7678838 | 0.00029 | 0.001336 | T | F | T | F |
| Stap1 | -8.1124114 | 0.64091696 | 14.7601431 | 0.00029 | 0.001341 | T | F | F | F |
| Tspan3 | -0.8643504 | 6.633232 | 14.7592223 | 0.00029 | 0.001341 | T | F | T | F |
| Ccdc85b | 1.04440734 | 4.96801244 | 14.7574369 | 0.00029 | 0.001342 | T | F | T | T |
| Armh4 | -1.4448189 | 4.16274442 | 14.7556904 | 0.00029 | 0.001342 | T | F | T | F |
| Cfdp1 | 0.74819152 | 7.2391215 | 14.7543367 | 0.00029 | 0.001343 | F | F | T | F |
| Snrk | 0.95510721 | 5.43271512 | 14.7490751 | 0.00029 | 0.001345 | T | F | T | T |
| Rabl3 | -1.5098968 | 4.00049502 | 14.7471378 | 0.00029 | 0.001346 | T | T | T | T |
| Deptor | -1.3794074 | 4.29970445 | 14.7383715 | 0.00029 | 0.001351 | F | F | T | T |
| E130308A1 | -1.0487004 | 5.34785958 | 14.7362671 | 0.00029 | 0.001352 | T | F | T | T |
| Pcbp3 | -1.5866733 | 3.83727479 | 14.7320414 | 0.00029 | 0.001354 | F | F | T | F |
| Lss | 1.04278617 | 4.97483527 | 14.7286138 | 0.00029 | 0.001356 | F | F | T | T |
| Zmiz1 | 0.71432362 | 7.99330753 | 14.7184478 | 0.00029 | 0.001362 | F | T | T | T |
| Fbxo41 | -8.0954572 | 0.62958486 | 14.700848 | 0.00029 | 0.001372 | T | F | T | T |
| Hnrnph2 | -0.8667191 | 6.62361323 | 14.6957466 | 0.00029 | 0.001374 | F | F | F | T |
| Prdx3 | 0.91732698 | 5.74467565 | 14.6832598 | 0.0003 | 0.001381 | F | F | T | T |
| Ypel2 | -2.7397794 | 2.25869602 | 14.6777934 | 0.0003 | 0.001384 | T | F | T | T |
| Kras | 0.84981932 | 6.25742929 | 14.6747133 | 0.0003 | 0.001386 | T | F | T | T |
| Chchd4 | 0.92729636 | 5.687161 | 14.6708565 | 0.0003 | 0.001388 | F | F | T | F |
| Cks1b | -0.9112976 | 6.3126893 | 14.6658728 | 0.0003 | 0.001391 | F | F | F | F |
| Tmem243 | -2.238361 | 2.82977994 | 14.6639624 | 0.0003 | 0.001391 | F | F | F | F |
| Smyd5 | -0.8591716 | 6.66187363 | 14.6607513 | 0.0003 | 0.001393 | F | F | T | F |
| Trmt9b | -2.6466436 | 2.38535833 | 14.6592582 | 0.0003 | 0.001394 | F | F | T | F |
| Zfp708 | -8.0727462 | 0.61961905 | 14.6588368 | 0.0003 | 0.001394 | F | F | T | F |
| Slco3a1 | -1.3273698 | 4.39464501 | 14.6563249 | 0.0003 | 0.001395 | T | F | T | F |
| Eif5a2 | -1.3260655 | 4.32032488 | 14.6531689 | 0.0003 | 0.001396 | T | F | T | T |
| Actr10 | 0.92580903 | 5.70294294 | 14.6522931 | 0.0003 | 0.001397 | F | F | T | T |
| Zbtb38 | 1.03693276 | 4.97795215 | 14.6435215 | 0.0003 | 0.001402 | F | F | F | T |
| Rnf6 | 0.90624738 | 5.757038 | 14.6369761 | 0.0003 | 0.001405 | T | F | F | F |
| Fam111a | -1.4082131 | 4.18255467 | 14.6292254 | 0.0003 | 0.00141 | T | F | T | T |
| Chst12 | -1.6729502 | 3.66548519 | 14.6275602 | 0.0003 | 0.00141 | F | F | F | T |
| Cep192 | 0.73344167 | 7.44934447 | 14.6194675 | 0.0003 | 0.001415 | T | F | F | T |
| Tspyl5 | -4.3629644 | 1.22540984 | 14.6190169 | 0.0003 | 0.001415 | T | F | T | T |
| Gm9803 | -2.6888433 | 2.30268275 | 14.6140146 | 0.0003 | 0.001417 | F | F | F | F |
| Lct | -8.085063 | 0.62117577 | 14.6136779 | 0.0003 | 0.001417 | F | F | F | F |

| | | | | | | | | | |
|---|---|---|---|---|---|---|---|---|---|
| Arl16 | -8.0704614 | 0.63671636 | 14.6136189 | 0.0003 | 0.001417 | T | F | F | F |
| Prmt9 | -1.2517577 | 4.55851358 | 14.6028311 | 0.00031 | 0.001424 | F | F | F | F |
| Mpped1 | -8.0622773 | 0.61876984 | 14.6000633 | 0.00031 | 0.001425 | F | F | T | F |
| Prdx2 | -0.7539976 | 7.82043283 | 14.5952092 | 0.00031 | 0.001428 | T | F | T | F |
| Ascc3 | 0.75964607 | 7.02751603 | 14.5933794 | 0.00031 | 0.001428 | F | F | T | T |
| Asxl2 | 0.85353654 | 6.19843049 | 14.5926604 | 0.00031 | 0.001429 | T | F | T | T |
| Acbd3 | 0.86975975 | 6.05070551 | 14.5881852 | 0.00031 | 0.001431 | F | F | T | T |
| Bicral | -1.235394 | 4.6444153 | 14.5842275 | 0.00031 | 0.001433 | F | F | T | T |
| Zkscan4 | -5.017367 | 1.02274472 | 14.5797363 | 0.00031 | 0.001436 | F | F | F | T |
| Slc25a34 | 3.82870499 | 0.63788758 | 14.5773355 | 0.00031 | 0.001437 | T | F | F | F |
| Dhx40 | -0.9740587 | 5.77024884 | 14.5721698 | 0.00031 | 0.001439 | T | F | T | F |
| Herc1 | 0.71979914 | 7.93073074 | 14.5720546 | 0.00031 | 0.001439 | T | F | F | T |
| Psmg2 | 0.97721506 | 5.22934347 | 14.5688637 | 0.00031 | 0.001441 | F | F | F | F |
| Erv3 | 2.10666012 | 2.44384595 | 14.5654462 | 0.00031 | 0.001443 | T | F | F | T |
| Cmip | 0.87432787 | 6.02837353 | 14.5614669 | 0.00031 | 0.001445 | F | F | T | F |
| Slc46a3 | -8.0806126 | 0.61691668 | 14.5591584 | 0.00031 | 0.001446 | F | F | F | F |
| Plcb2 | -8.0590383 | 0.61993946 | 14.5556976 | 0.00031 | 0.001448 | T | F | F | T |
| Cd302 | -8.0595378 | 0.62167187 | 14.5534519 | 0.00031 | 0.001449 | F | F | F | F |
| Rassf7 | -1.2657282 | 4.5050148 | 14.5528399 | 0.00031 | 0.001449 | F | F | F | F |
| Mras | -2.3330476 | 2.75270908 | 14.5511131 | 0.00031 | 0.00145 | T | F | T | F |
| Mrpl28 | -0.9795347 | 5.79867192 | 14.5500122 | 0.00031 | 0.00145 | F | F | F | F |
| Foxi2 | -5.0711239 | 1.00588495 | 14.5465934 | 0.00031 | 0.001452 | F | F | F | F |
| Uggt1 | 0.81689174 | 6.49189755 | 14.5362812 | 0.00031 | 0.001458 | F | F | T | T |
| Gm45844 | 2.89719783 | 1.32562695 | 14.5348657 | 0.00032 | 0.001459 | F | F | F | F |
| Tmem44 | -8.0680783 | 0.60931101 | 14.5081451 | 0.00032 | 0.001475 | F | F | T | F |
| Fkbp6 | 2.5661532 | 1.69187686 | 14.5030829 | 0.00032 | 0.001478 | F | F | T | F |
| Vwa5a | -1.8632609 | 3.3988647 | 14.5013965 | 0.00032 | 0.001479 | T | F | F | T |
| Sox7 | -2.2398577 | 2.84697646 | 14.500636 | 0.00032 | 0.001479 | T | F | T | T |
| Plat | -1.7498239 | 3.56576122 | 14.4984593 | 0.00032 | 0.00148 | T | F | F | F |
| Rtn4rl1 | -2.3092432 | 2.72964109 | 14.4965055 | 0.00032 | 0.001481 | F | F | T | T |
| Rfx7 | 0.77155562 | 6.98615432 | 14.4959243 | 0.00032 | 0.001481 | T | F | F | T |
| Lsm7 | -1.1028049 | 5.20508792 | 14.4880203 | 0.00032 | 0.001486 | F | F | F | F |
| Batf3 | 3.17603694 | 1.40647628 | 14.4867247 | 0.00032 | 0.001487 | F | F | F | F |
| Cdh17 | -4.4074813 | 1.25216315 | 14.4842899 | 0.00032 | 0.001488 | F | F | F | F |
| Ehmt2 | 0.68092214 | 8.29994643 | 14.4682427 | 0.00032 | 0.001498 | F | F | F | F |
| Gstt1 | -8.0899191 | 0.66572118 | 14.4671605 | 0.00032 | 0.001498 | F | F | F | F |
| Epm2aip1 | -0.9757189 | 5.73212109 | 14.4650613 | 0.00032 | 0.001499 | T | T | T | T |
| Pkmyt1 | 1.15634078 | 4.42897 | 14.4521631 | 0.00033 | 0.001507 | F | F | F | F |
| Minpp1 | -0.913432 | 6.22925379 | 14.4510402 | 0.00033 | 0.001508 | F | F | F | F |
| Mlycd | -2.7352212 | 2.28496889 | 14.4471794 | 0.00033 | 0.00151 | F | F | T | T |
| Itm2a | -2.1583095 | 2.90904482 | 14.4452031 | 0.00033 | 0.001511 | T | F | F | F |
| Tmem129 | 1.90594444 | 2.75070782 | 14.4404328 | 0.00033 | 0.001514 | F | F | T | F |

| | | | | | | | | | |
|---|---|---|---|---|---|---|---|---|---|
| Gtf2h2 | 1.67500373 | 3.24824995 | 14.4314895 | 0.00033 | 0.001519 | T | F | F | F |
| Cyp4f39 | 5.75591841 | -0.0888896 | 14.4237806 | 0.00033 | 0.001524 | T | F | F | T |
| Nptn | 0.90689343 | 5.70538264 | 14.417423 | 0.00033 | 0.001528 | T | F | T | T |
| Dennd5b | -0.8822439 | 6.49983172 | 14.4158747 | 0.00033 | 0.001529 | T | F | T | F |
| Hif1a | 0.68651358 | 8.10596104 | 14.4064152 | 0.00033 | 0.001534 | T | F | T | F |
| Mef2a | 0.93915805 | 5.45652938 | 14.3939211 | 0.00033 | 0.001542 | F | F | T | F |
| Kcna3 | -8.0570672 | 0.59917085 | 14.3868425 | 0.00034 | 0.001547 | F | F | F | F |
| 0610010F0 | 1.14814506 | 4.46557912 | 14.3851582 | 0.00034 | 0.001548 | F | F | T | F |
| Trrap | 0.6305126 | 9.44075504 | 14.3772907 | 0.00034 | 0.001553 | T | F | F | F |
| Lrif1 | 1.52526909 | 3.48537755 | 14.3673954 | 0.00034 | 0.001559 | T | F | F | F |
| Cep290 | 0.94024547 | 5.53134494 | 14.3646834 | 0.00034 | 0.00156 | F | F | T | F |
| Mlxipl | -3.9301867 | 1.41582809 | 14.3645947 | 0.00034 | 0.00156 | F | F | F | T |
| D430041D0 | -2.1020255 | 3.05120701 | 14.3607233 | 0.00034 | 0.001562 | T | F | T | F |
| Gm49909 | 0.79455134 | 6.72875135 | 14.3599045 | 0.00034 | 0.001563 | F | F | F | F |
| Adamts13 | 1.37661562 | 3.84152129 | 14.3400317 | 0.00034 | 0.001576 | F | F | F | F |
| Lrrc19 | -3.9369129 | 1.37690263 | 14.3351489 | 0.00034 | 0.001579 | T | F | F | T |
| Klhdc3 | -0.794414 | 7.04768166 | 14.3323046 | 0.00034 | 0.00158 | F | F | F | F |
| Ap1ar | -0.9361899 | 6.02180394 | 14.3317127 | 0.00034 | 0.00158 | T | F | T | F |
| Casc4 | -0.9278514 | 6.11747564 | 14.3310713 | 0.00034 | 0.001581 | T | F | F | T |
| Rbmxl2 | -8.0917627 | 0.6166703 | 14.3279041 | 0.00034 | 0.001582 | F | F | F | F |
| Setd6 | 1.19336509 | 4.36523803 | 14.3231226 | 0.00035 | 0.001585 | F | F | T | F |
| Srd5a1 | 1.42765136 | 3.7256607 | 14.322357 | 0.00035 | 0.001585 | T | F | T | T |
| Glb1l2 | -8.0813852 | 0.60978841 | 14.3219366 | 0.00035 | 0.001585 | T | F | T | T |
| Patl1 | 0.84998297 | 6.14759114 | 14.3127777 | 0.00035 | 0.001591 | T | T | F | T |
| Pi4k2b | 0.96137649 | 5.31457699 | 14.3081915 | 0.00035 | 0.001594 | F | F | F | F |
| Rab5b | 0.86516918 | 6.11561082 | 14.307388 | 0.00035 | 0.001594 | F | F | T | T |
| Rabep1 | 0.79869932 | 6.60430186 | 14.2947543 | 0.00035 | 0.001603 | T | F | F | T |
| Cdk5r2 | -8.04115 | 0.58885047 | 14.2861492 | 0.00035 | 0.001608 | T | F | T | F |
| Clec1a | 3.41090437 | 0.87033713 | 14.2677314 | 0.00035 | 0.001621 | T | F | T | T |
| Fez2 | 1.04065353 | 4.88085615 | 14.2602627 | 0.00035 | 0.001626 | T | F | T | F |
| Mus81 | -1.7313708 | 3.51514457 | 14.2539202 | 0.00036 | 0.00163 | F | F | F | T |
| Sh2d4b | -8.0378064 | 0.58568607 | 14.2484145 | 0.00036 | 0.001634 | F | F | T | T |
| Fyn | -1.0624947 | 5.18708931 | 14.2442747 | 0.00036 | 0.001636 | T | F | F | T |
| Ccnk | -0.8752397 | 6.45500639 | 14.2427184 | 0.00036 | 0.001637 | F | F | F | F |
| Aldh4a1 | 0.9327851 | 5.51814941 | 14.2420031 | 0.00036 | 0.001637 | F | F | T | T |
| Tnfaip8l3 | -8.0306033 | 0.58485261 | 14.2337081 | 0.00036 | 0.001643 | T | F | F | T |
| Atxn7l2 | -1.3168894 | 4.30607038 | 14.2246509 | 0.00036 | 0.001649 | F | F | F | F |
| Pop7 | 1.25120618 | 4.17187229 | 14.2212526 | 0.00036 | 0.001651 | F | F | F | F |
| Dnd1 | -8.0326694 | 0.6023043 | 14.2104229 | 0.00036 | 0.001658 | T | F | T | F |
| Esrrg | -8.0304515 | 0.58195752 | 14.2067687 | 0.00036 | 0.00166 | T | F | T | T |
| Rap1gds1 | 0.81243218 | 6.45049382 | 14.2054758 | 0.00036 | 0.001661 | T | F | T | T |
| Lrfn1 | -1.1008937 | 5.08102604 | 14.2015998 | 0.00036 | 0.001663 | F | F | F | F |

| | | | | | | | | | |
|---|---|---|---|---|---|---|---|---|---|
| Sac3d1 | -1.0638465 | 5.1728698 | 14.1916834 | 0.00037 | 0.00167 | F | F | F | F |
| Nmnat2 | -1.3760979 | 4.30500518 | 14.1896994 | 0.00037 | 0.001671 | T | F | F | T |
| Msl3l2 | -2.9857713 | 1.97977356 | 14.183914 | 0.00037 | 0.001675 | T | T | F | T |
| Oit3 | -8.0404038 | 0.61462877 | 14.1838526 | 0.00037 | 0.001675 | T | F | F | F |
| Grpel2 | -1.1989433 | 4.66206531 | 14.1835923 | 0.00037 | 0.001675 | T | F | T | T |
| 1700025G0 | 0.95391399 | 5.37958335 | 14.1730237 | 0.00037 | 0.001682 | F | F | T | T |
| Znrf2 | 0.89953692 | 6.23569324 | 14.1711267 | 0.00037 | 0.001683 | F | F | F | T |
| Gnl1 | 0.83751459 | 6.23374499 | 14.1700487 | 0.00037 | 0.001684 | T | F | F | T |
| Arhgap33 | -1.6844553 | 3.58743415 | 14.1637976 | 0.00037 | 0.001688 | F | F | F | F |
| Gm45902 | 1.07093355 | 4.73283071 | 14.1556249 | 0.00037 | 0.001693 | F | F | F | F |
| Selenoi | 0.80237122 | 6.5550578 | 14.1475097 | 0.00037 | 0.001699 | T | F | F | F |
| Pqbp1 | -1.0284898 | 5.33630988 | 14.1416934 | 0.00037 | 0.001703 | F | F | F | F |
| Ddrgk1 | 1.07188758 | 4.80777291 | 14.1399048 | 0.00037 | 0.001704 | F | F | F | T |
| Acacb | 2.01550692 | 2.51408534 | 14.1386019 | 0.00037 | 0.001704 | F | F | F | T |
| Agpat5 | 0.8540964 | 6.07632222 | 14.1383488 | 0.00037 | 0.001704 | F | F | F | T |
| Kat6a | 0.6968509 | 7.83058168 | 14.1374794 | 0.00037 | 0.001705 | T | F | T | T |
| Unc13d | 2.28123127 | 2.14903744 | 14.1350252 | 0.00037 | 0.001706 | F | F | F | F |
| Prss23 | -4.3221994 | 1.19077582 | 14.1295227 | 0.00038 | 0.00171 | T | F | F | T |
| Ccdc65 | -5.043016 | 0.97628078 | 14.1180894 | 0.00038 | 0.001718 | F | F | F | F |
| Slc35g2 | -8.0196823 | 0.57505453 | 14.1177084 | 0.00038 | 0.001718 | F | F | F | T |
| Tuba1b | -0.6313026 | 10.5535261 | 14.11507 | 0.00038 | 0.001719 | F | F | F | T |
| Stc1 | -3.9751585 | 1.38593732 | 14.1128308 | 0.00038 | 0.001721 | T | F | F | F |
| AW551984 | -3.3019758 | 1.71350515 | 14.1009501 | 0.00038 | 0.001729 | T | F | T | T |
| Pcdhga9 | -8.0360522 | 0.61207996 | 14.1008303 | 0.00038 | 0.001729 | F | F | F | F |
| Rnf10 | -0.6958865 | 8.54831683 | 14.0879762 | 0.00038 | 0.001738 | F | F | F | F |
| Zfp58 | -2.2910227 | 2.71081583 | 14.0832739 | 0.00038 | 0.001742 | F | F | F | F |
| Camk2a | -2.5947938 | 2.47094662 | 14.082032 | 0.00038 | 0.001742 | T | F | T | F |
| D830030K2 | -8.0219925 | 0.57111465 | 14.0764582 | 0.00038 | 0.001746 | F | F | T | F |
| Brsk2 | -3.5691475 | 1.55689975 | 14.0731671 | 0.00038 | 0.001748 | F | F | T | F |
| AU040320 | 0.98801044 | 5.1283514 | 14.0680686 | 0.00039 | 0.001752 | T | F | F | F |
| Cnksr2 | -1.3665099 | 4.18855942 | 14.0655774 | 0.00039 | 0.001753 | F | F | T | T |
| Tbl1x | -0.7421935 | 7.69027718 | 14.0647496 | 0.00039 | 0.001753 | F | F | T | T |
| Osbpl1a | 1.04374118 | 4.85412623 | 14.0576564 | 0.00039 | 0.001758 | T | T | T | T |
| Galnt4 | 1.3972926 | 3.77591437 | 14.0506864 | 0.00039 | 0.001763 | T | F | T | F |
| Gbf1 | -0.9358004 | 6.36093449 | 14.0443947 | 0.00039 | 0.001768 | T | F | T | F |
| Tpm3-rs7 | 2.95846146 | 1.27700891 | 14.0441917 | 0.00039 | 0.001768 | F | F | F | F |
| Ttc39b | -1.8626577 | 3.37672507 | 14.0392433 | 0.00039 | 0.001771 | F | F | T | F |
| Calcr | -8.1002504 | 0.61702398 | 14.0366071 | 0.00039 | 0.001773 | T | F | F | F |
| Tmsb10 | 0.64720762 | 9.28865683 | 14.0325201 | 0.00039 | 0.001775 | F | F | F | F |
| Sox15 | 3.07679751 | 1.09950879 | 14.0300878 | 0.00039 | 0.001777 | F | F | F | F |
| Bsg | 0.65615051 | 8.64552164 | 14.0263969 | 0.00039 | 0.001779 | T | F | F | F |
| Adamts12 | -3.1503704 | 1.87121414 | 14.0236155 | 0.00039 | 0.001781 | T | F | T | T |

| | | | | | | | | | |
|---|---|---|---|---|---|---|---|---|---|
| Glmp | 0.88432713 | 5.79769704 | 14.0211142 | 0.00039 | 0.001783 | F | F | F | F |
| Hrh3 | -8.0855806 | 0.60587929 | 14.0195175 | 0.00039 | 0.001783 | F | F | T | F |
| 2210016F1( | 1.00411546 | 5.01511118 | 14.0190925 | 0.00039 | 0.001783 | F | F | T | T |
| Dmrt3 | -8.0088417 | 0.56814164 | 14.0189191 | 0.00039 | 0.001783 | T | F | F | F |
| Srf | 0.7296694 | 7.15076571 | 14.0111471 | 0.0004 | 0.001789 | F | F | T | F |
| Lama2 | -8.0223612 | 0.59809684 | 14.004991 | 0.0004 | 0.001793 | F | F | F | T |
| Asrgl1 | -1.369337 | 4.18252508 | 13.9939481 | 0.0004 | 0.001802 | F | F | T | F |
| M6pr | 0.72858305 | 7.15218171 | 13.9916053 | 0.0004 | 0.001803 | T | F | T | T |
| Lrrc73 | 2.82009493 | 1.43821134 | 13.9884477 | 0.0004 | 0.001805 | F | F | T | T |
| Klf8 | -1.3727843 | 4.22522812 | 13.9867512 | 0.0004 | 0.001806 | T | F | T | T |
| Ttc30a1 | -1.7795563 | 3.42653333 | 13.9805803 | 0.0004 | 0.001811 | T | F | F | F |
| Zmynd10 | -8.0187192 | 0.59430748 | 13.9798636 | 0.0004 | 0.001811 | F | F | F | F |
| Csad | -1.7735743 | 3.39779203 | 13.969888 | 0.0004 | 0.001818 | F | F | F | T |
| Ttc8 | -1.9549832 | 3.12598644 | 13.9610811 | 0.0004 | 0.001825 | F | F | F | F |
| Csnk1e | -0.8659413 | 6.46455751 | 13.957857 | 0.0004 | 0.001827 | T | F | T | T |
| Lrrk2 | -3.4478921 | 1.59645544 | 13.9567414 | 0.0004 | 0.001827 | F | F | F | F |
| Wdr47 | 1.39502884 | 3.75099384 | 13.9567055 | 0.0004 | 0.001827 | T | F | F | T |
| Cxcr2 | 0.91169012 | 5.60593295 | 13.9558609 | 0.0004 | 0.001828 | T | F | T | T |
| Gm3373 | 3.51360444 | 0.95694836 | 13.9528314 | 0.00041 | 0.00183 | F | F | F | F |
| 4931414P1( | -2.0511084 | 2.99017649 | 13.944532 | 0.00041 | 0.001836 | F | F | T | F |
| Tnfsf9 | -4.3101368 | 1.17708194 | 13.9432341 | 0.00041 | 0.001837 | T | F | F | T |
| Nr1d2 | -1.7035903 | 3.52372413 | 13.9381487 | 0.00041 | 0.00184 | T | F | T | T |
| Mei4 | 2.60431741 | 1.58580675 | 13.937411 | 0.00041 | 0.00184 | F | F | T | F |
| Tigit | -3.0142437 | 1.91042925 | 13.9358694 | 0.00041 | 0.001841 | F | F | F | F |
| Stk32b | -8.0102336 | 0.55792399 | 13.9123839 | 0.00041 | 0.00186 | T | F | T | T |
| Rap2a | 1.03827012 | 4.83814143 | 13.9083202 | 0.00041 | 0.001863 | T | F | T | T |
| Hspa12b | -2.2286412 | 2.7543883 | 13.8991793 | 0.00041 | 0.00187 | F | F | F | T |
| Fosb | 2.47109386 | 1.8022791 | 13.8964888 | 0.00042 | 0.001872 | F | F | F | T |
| Hr | -2.7268321 | 2.17415601 | 13.8950115 | 0.00042 | 0.001872 | F | F | F | F |
| Ap5z1 | 1.25092758 | 4.11824532 | 13.8930959 | 0.00042 | 0.001874 | T | F | F | T |
| Cacna1h | -2.189329 | 2.83659184 | 13.8851853 | 0.00042 | 0.00188 | F | F | T | F |
| Dlgap2 | -8.0512986 | 0.57905507 | 13.8844598 | 0.00042 | 0.00188 | F | F | T | T |
| Lmbr1 | 1.45800287 | 3.59595347 | 13.8826036 | 0.00042 | 0.001881 | T | T | T | T |
| Drc7 | -8.0118447 | 0.55660308 | 13.877284 | 0.00042 | 0.001885 | F | F | F | T |
| Tmem176b | -2.5222968 | 2.44054798 | 13.873265 | 0.00042 | 0.001888 | F | F | F | T |
| Parp14 | 2.2653359 | 2.03907633 | 13.8633357 | 0.00042 | 0.001896 | T | F | T | F |
| Nhej1 | -2.6915735 | 2.21368568 | 13.8605019 | 0.00042 | 0.001898 | F | F | F | F |
| Adsl | -0.9127746 | 6.07488328 | 13.8535426 | 0.00042 | 0.001903 | F | F | T | T |
| Tkfc | -1.2151089 | 4.55248898 | 13.8410185 | 0.00043 | 0.001913 | F | F | T | F |
| Morf4l1 | -0.7294208 | 7.8402138 | 13.8403033 | 0.00043 | 0.001913 | F | F | F | T |
| Lancl2 | -1.2127992 | 4.56337327 | 13.8253307 | 0.00043 | 0.001925 | T | F | F | T |
| Sav1 | 0.8651454 | 5.88896921 | 13.8213824 | 0.00043 | 0.001928 | F | F | F | F |

| | | | | | | | | | |
|---|---|---|---|---|---|---|---|---|---|
| Pkp1 | -8.0187681 | 0.5998009 | 13.8198254 | 0.00043 | 0.001929 | F | F | T | F |
| Prkcg | 2.75101313 | 1.427674 | 13.8141132 | 0.00043 | 0.001933 | T | F | F | T |
| Cat | -0.9024277 | 6.16074058 | 13.8122574 | 0.00043 | 0.001934 | F | F | F | T |
| Mdm4 | 0.73619371 | 7.04162309 | 13.8121382 | 0.00043 | 0.001934 | T | F | T | T |
| Gpx7 | -2.052886 | 3.07519012 | 13.811733 | 0.00043 | 0.001934 | F | F | T | F |
| Rundc1 | 1.02007458 | 4.96111596 | 13.8038249 | 0.00043 | 0.00194 | F | F | T | T |
| Grb7 | -0.9473961 | 5.76106898 | 13.8034954 | 0.00043 | 0.00194 | F | F | T | F |
| Pdzrn4 | -7.9938425 | 0.54739859 | 13.8031521 | 0.00043 | 0.00194 | F | F | F | F |
| Galnt3 | 1.53992722 | 3.41520964 | 13.8026236 | 0.00043 | 0.00194 | F | F | T | F |
| Mbtps2 | 0.92910817 | 5.49044904 | 13.798891 | 0.00043 | 0.001943 | T | F | T | F |
| Ccdc180 | -7.997637 | 0.54760935 | 13.7983632 | 0.00043 | 0.001943 | F | F | F | F |
| Snapc5 | -1.8775624 | 3.23587804 | 13.7869748 | 0.00044 | 0.001952 | F | F | F | T |
| Ass1 | 0.90909354 | 5.76228195 | 13.7843753 | 0.00044 | 0.001954 | F | F | F | F |
| Olfr843 | 4.3718332 | 0.3401259 | 13.7831897 | 0.00044 | 0.001955 | F | F | F | F |
| Xntrpc | -7.9865563 | 0.55514285 | 13.7823181 | 0.00044 | 0.001955 | F | F | F | F |
| Plekho2 | 1.46482038 | 3.55481865 | 13.7745853 | 0.00044 | 0.001961 | T | F | T | F |
| Bche | -2.6261338 | 2.28422697 | 13.7718558 | 0.00044 | 0.001963 | T | F | T | T |
| Ano9 | -4.2930968 | 1.16723611 | 13.7717054 | 0.00044 | 0.001963 | T | F | F | F |
| Rab37 | -7.9848602 | 0.5473801 | 13.7699913 | 0.00044 | 0.001964 | T | F | T | T |
| Vmn2r112 | -8.499872 | 0.97248903 | 13.7693859 | 0.00044 | 0.001964 | F | F | F | F |
| Zfp763 | -1.6987619 | 3.51217689 | 13.7664403 | 0.00044 | 0.001966 | T | T | F | T |
| Crlf2 | 1.91299291 | 2.68168021 | 13.7550946 | 0.00044 | 0.001976 | F | F | F | T |
| Dbndd1 | -5.0046157 | 0.94597631 | 13.7494295 | 0.00044 | 0.00198 | F | F | F | F |
| Nphp4 | -1.6785942 | 3.54416631 | 13.7399208 | 0.00044 | 0.001988 | F | F | F | F |
| Sox5 | -8.0253541 | 0.60927803 | 13.7326703 | 0.00045 | 0.001994 | T | F | T | F |
| Tamm41 | -1.2414382 | 4.47999627 | 13.7291772 | 0.00045 | 0.001997 | F | F | F | F |
| Prelid3b | 0.85882576 | 5.91864144 | 13.727463 | 0.00045 | 0.001998 | T | F | T | F |
| Dynll1 | -0.7492576 | 7.43931694 | 13.7233359 | 0.00045 | 0.002001 | F | F | F | F |
| Agk | 1.07352358 | 4.65723095 | 13.7212279 | 0.00045 | 0.002002 | T | F | F | T |
| Wsb1 | -0.9560713 | 5.68219041 | 13.7163623 | 0.00045 | 0.002006 | F | F | F | F |
| Mtdh | 0.7126938 | 7.36393202 | 13.7118265 | 0.00045 | 0.00201 | T | T | T | T |
| Dstyk | -1.4152356 | 4.05059676 | 13.7107518 | 0.00045 | 0.00201 | T | F | F | T |
| Sept11 | 0.63630557 | 9.0950209 | 13.7082938 | 0.00045 | 0.002012 | T | F | T | T |
| Uaca | 0.7313147 | 7.31526144 | 13.7008584 | 0.00045 | 0.002018 | F | F | F | T |
| Ggt7 | -2.3921804 | 2.54507802 | 13.6951003 | 0.00045 | 0.002023 | F | F | F | T |
| Asphd2 | -2.2087527 | 2.75199712 | 13.6940347 | 0.00045 | 0.002023 | T | F | T | T |
| Olfr316 | 4.60907841 | 0.21896717 | 13.6930138 | 0.00045 | 0.002024 | F | F | T | F |
| Tomm40l | -1.4406056 | 3.9998986 | 13.6831652 | 0.00046 | 0.002032 | F | F | T | F |
| Ccdc47 | 0.76967322 | 6.71023315 | 13.6791279 | 0.00046 | 0.002035 | T | T | T | T |
| Loxl1 | -3.287399 | 1.70542752 | 13.6662702 | 0.00046 | 0.002046 | F | F | F | F |
| Rasal1 | 1.71821502 | 3.04951861 | 13.6592185 | 0.00046 | 0.002052 | F | F | F | F |
| Mgme1 | -1.1618846 | 4.70817568 | 13.6413125 | 0.00046 | 0.002068 | F | F | F | F |

| | | | | | | | | | |
|---|---|---|---|---|---|---|---|---|---|
| Ugt1a6a | -7.9867047 | 0.53482991 | 13.6357191 | 0.00047 | 0.002073 | F | F | F | T |
| St14 | -1.1071194 | 4.90554899 | 13.6306196 | 0.00047 | 0.002077 | T | F | T | T |
| Snrnp70 | -0.7410079 | 7.50145191 | 13.6193598 | 0.00047 | 0.002087 | F | F | F | F |
| Sra1 | -1.5384724 | 3.81119364 | 13.6168892 | 0.00047 | 0.002088 | F | F | T | F |
| Pnpla6 | 0.84372886 | 6.01996248 | 13.6122176 | 0.00047 | 0.002092 | F | F | F | F |
| Gorab | -2.1217633 | 2.85690926 | 13.6096902 | 0.00047 | 0.002094 | F | F | F | T |
| Podn | -7.9845581 | 0.53246591 | 13.6057274 | 0.00047 | 0.002097 | F | F | F | F |
| Pou4f1 | -7.9723175 | 0.54684813 | 13.6005549 | 0.00047 | 0.002102 | T | F | T | F |
| Pnmal2 | -7.9682027 | 0.53689973 | 13.598776 | 0.00047 | 0.002103 | F | F | T | F |
| Gtf3c6 | -0.9930043 | 5.42206428 | 13.5970598 | 0.00047 | 0.002104 | F | F | T | T |
| Wnt4 | 1.21527333 | 4.15757823 | 13.5935473 | 0.00047 | 0.002107 | F | F | T | F |
| Ung | -0.8634217 | 6.34547838 | 13.5907174 | 0.00047 | 0.002109 | T | F | T | T |
| Ppp1r16a | -1.0157562 | 5.30908758 | 13.5743882 | 0.00048 | 0.002124 | T | F | F | F |
| Akt2 | 0.75873697 | 6.78187377 | 13.5672266 | 0.00048 | 0.00213 | F | F | T | F |
| Ifit2 | -2.1546972 | 2.80760206 | 13.5614865 | 0.00048 | 0.002135 | T | F | T | F |
| Lrrc34 | -8.0652776 | 0.58275534 | 13.5597889 | 0.00048 | 0.002136 | T | F | F | F |
| Slc52a2 | 1.02801826 | 4.88615868 | 13.5589981 | 0.00048 | 0.002136 | F | F | T | T |
| Zfp444 | -1.1052803 | 4.90090875 | 13.5568867 | 0.00048 | 0.002138 | F | F | T | F |
| Ficd | 1.56660358 | 3.29044871 | 13.556427 | 0.00048 | 0.002138 | T | F | T | F |
| H2-Eb2 | -8.0648756 | 0.58235948 | 13.5542231 | 0.00048 | 0.002139 | T | F | F | F |
| Trio | 0.67191404 | 8.38293804 | 13.5537856 | 0.00048 | 0.002139 | T | F | T | T |
| Ehd4 | 0.88634157 | 5.64776871 | 13.5496548 | 0.00048 | 0.002143 | T | T | F | T |
| Doc2b | -7.9724481 | 0.55076535 | 13.5490216 | 0.00048 | 0.002143 | T | F | F | T |
| Hdac10 | -2.4769783 | 2.45128881 | 13.5445145 | 0.00048 | 0.002147 | F | F | F | F |
| Hoxa5 | -5.0815481 | 1.04669047 | 13.5407375 | 0.00049 | 0.00215 | F | F | T | F |
| Hdgfl2 | -0.7912366 | 6.92185779 | 13.5357351 | 0.00049 | 0.002154 | F | F | F | F |
| Ess2 | -1.0899715 | 4.95691142 | 13.5262816 | 0.00049 | 0.002163 | F | F | T | F |
| Stag1 | 0.70156711 | 7.43329931 | 13.5251293 | 0.00049 | 0.002163 | T | F | T | T |
| Ccdc30 | -4.9796733 | 0.93164881 | 13.5204741 | 0.00049 | 0.002167 | T | F | T | F |
| Tmem65 | 0.9252441 | 5.36808538 | 13.5070675 | 0.00049 | 0.00218 | T | F | T | T |
| Zfand3 | 0.85129495 | 5.99045454 | 13.4989478 | 0.00049 | 0.002187 | T | F | T | F |
| Med15 | 0.86114396 | 5.83833835 | 13.4960786 | 0.00049 | 0.002189 | T | F | T | F |
| B3gntl1 | -1.9035024 | 3.16085916 | 13.4923693 | 0.0005 | 0.002192 | F | F | T | F |
| Notum | -2.1903927 | 2.78628229 | 13.4913244 | 0.0005 | 0.002193 | F | F | F | T |
| Ppp1r9a | -1.1320571 | 4.80855903 | 13.4892386 | 0.0005 | 0.002194 | T | F | F | T |
| Kif16b | 0.95290549 | 5.2087973 | 13.4771608 | 0.0005 | 0.002206 | T | T | F | T |
| Ccdc134 | 1.37430926 | 3.87710418 | 13.4728214 | 0.0005 | 0.002209 | T | F | F | F |
| Acbd4 | 3.12250773 | 1.080716 | 13.4701056 | 0.0005 | 0.002212 | F | F | F | F |
| Cldn11 | -7.9548556 | 0.52280995 | 13.463832 | 0.0005 | 0.002217 | F | F | F | F |
| Itfg1 | 0.92998562 | 5.36545144 | 13.4613961 | 0.0005 | 0.002219 | T | F | F | T |
| Syngr3 | -4.9962618 | 0.92969126 | 13.4544895 | 0.0005 | 0.002225 | F | F | T | F |
| Rtca | -1.0168339 | 5.24058829 | 13.4465284 | 0.00051 | 0.002233 | F | F | F | T |

| | | | | | | | | | |
|---|---|---|---|---|---|---|---|---|---|
| BC048403 | 1.0844262 | 4.57585357 | 13.4431205 | 0.00051 | 0.002236 | F | F | F | F |
| Dhtkd1 | -1.3923521 | 4.06519668 | 13.4425831 | 0.00051 | 0.002236 | T | F | T | T |
| Nmrk2 | -7.9726259 | 0.51939835 | 13.438378 | 0.00051 | 0.002239 | T | F | T | T |
| Sae1 | -0.7101142 | 7.91240164 | 13.4370833 | 0.00051 | 0.00224 | T | F | F | F |
| Eci1 | -1.6489969 | 3.57289384 | 13.414515 | 0.00051 | 0.002262 | F | F | T | F |
| Gid4 | -0.8694956 | 6.27996063 | 13.4116751 | 0.00051 | 0.002264 | T | T | T | T |
| Ciapin1 | -0.8559588 | 6.39592636 | 13.4114066 | 0.00051 | 0.002264 | F | F | T | F |
| Mdc1 | -0.7354964 | 7.57912805 | 13.4110312 | 0.00051 | 0.002264 | F | F | T | F |
| Serinc5 | -1.0198719 | 5.28702462 | 13.4068792 | 0.00051 | 0.002268 | T | F | T | F |
| Ino80b | -1.316416 | 4.21285497 | 13.4035952 | 0.00052 | 0.002271 | F | F | F | F |
| Aig1 | -1.9733108 | 3.03467761 | 13.3951244 | 0.00052 | 0.002279 | T | F | T | F |
| Sap25 | 4.53495851 | 0.27262692 | 13.3927338 | 0.00052 | 0.002281 | F | F | F | F |
| Vwa3a | 2.46676943 | 1.80032432 | 13.3875046 | 0.00052 | 0.002285 | F | F | F | F |
| Gpr50 | -3.3605923 | 1.74843138 | 13.3872305 | 0.00052 | 0.002285 | F | F | T | T |
| Nktr | 0.70871981 | 7.26452765 | 13.3859006 | 0.00052 | 0.002286 | T | F | T | F |
| Nap1l2 | -4.9836216 | 0.9208492 | 13.38555 | 0.00052 | 0.002286 | T | F | F | F |
| Dand5 | -8.0006057 | 0.58930807 | 13.3806902 | 0.00052 | 0.00229 | F | F | F | F |
| Nudt4 | -0.8373184 | 6.51108236 | 13.373788 | 0.00052 | 0.002297 | T | T | T | T |
| Kremen1 | 0.97301076 | 5.07239663 | 13.3707249 | 0.00052 | 0.002299 | T | F | T | T |
| Sim1 | -7.9665928 | 0.5513699 | 13.3586339 | 0.00053 | 0.002311 | T | T | T | T |
| Hectd4 | 0.70642634 | 7.39891631 | 13.3544911 | 0.00053 | 0.002315 | F | F | F | F |
| Ube2q1 | -0.8647825 | 6.27841348 | 13.3390198 | 0.00053 | 0.00233 | T | F | T | T |
| Ssc5d | -2.6498977 | 2.17747602 | 13.3307574 | 0.00053 | 0.002338 | F | F | F | F |
| Ncald | -4.968049 | 0.91360862 | 13.3305176 | 0.00053 | 0.002338 | T | F | T | T |
| Atpaf1 | 1.03946041 | 4.81645746 | 13.3126659 | 0.00054 | 0.002356 | F | F | T | T |
| mt-Nd5 | 0.58916194 | 9.88451117 | 13.3073873 | 0.00054 | 0.002361 | F | F | F | F |
| Zfp967 | 1.18767249 | 4.19474391 | 13.2997547 | 0.00054 | 0.002368 | F | F | T | F |
| Donson | 0.88728032 | 5.5826272 | 13.2896012 | 0.00054 | 0.002379 | F | F | F | T |
| Pgbd5 | -2.396815 | 2.47816257 | 13.2795934 | 0.00054 | 0.002389 | T | F | T | T |
| Ift122 | -1.0862095 | 4.938553 | 13.2702739 | 0.00055 | 0.002398 | T | F | T | F |
| Lysmd1 | 1.77400757 | 2.9065888 | 13.2632254 | 0.00055 | 0.002405 | F | F | F | F |
| Amfr | -0.830979 | 6.51698316 | 13.2468295 | 0.00055 | 0.002422 | F | F | F | T |
| Pik3r2 | -0.9112546 | 5.93293121 | 13.2464707 | 0.00055 | 0.002422 | F | F | T | T |
| Gtf2a1 | 0.71882756 | 7.18014302 | 13.2461666 | 0.00055 | 0.002422 | F | F | T | T |
| Gm49496 | -8.3171718 | 0.88750691 | 13.2440627 | 0.00055 | 0.002423 | F | F | F | F |
| Ypel1 | -2.3632103 | 2.52909065 | 13.2435324 | 0.00055 | 0.002423 | T | F | T | F |
| Smarce1 | -0.7289412 | 7.56632167 | 13.2395889 | 0.00055 | 0.002427 | F | F | T | F |
| Irf2bp2 | -0.9144893 | 5.86557433 | 13.2335956 | 0.00056 | 0.002433 | F | F | T | F |
| Pank3 | 0.787605 | 6.44592992 | 13.2257601 | 0.00056 | 0.002441 | T | T | T | T |
| Gm14443 | 2.12445305 | 2.18248964 | 13.2133302 | 0.00056 | 0.002454 | F | F | F | T |
| Cdkl1 | -7.9299682 | 0.50143644 | 13.2108799 | 0.00056 | 0.002456 | F | F | T | F |
| Caml | 1.17597827 | 4.21092722 | 13.2070753 | 0.00056 | 0.002459 | T | T | T | T |

| | | | | | | | | | |
|---|---|---|---|---|---|---|---|---|---|
| Flrt1 | -7.9839948 | 0.51755773 | 13.2070029 | 0.00056 | 0.002459 | F | F | T | T |
| Bach2 | -1.3713218 | 4.08160641 | 13.2053597 | 0.00056 | 0.00246 | F | F | T | F |
| Crybg2 | 2.3237477 | 1.92911088 | 13.1998913 | 0.00056 | 0.002466 | F | F | F | F |
| Acvr1 | -1.3611881 | 4.10653434 | 13.1831214 | 0.00057 | 0.002484 | F | F | T | T |
| Scamp5 | -1.1944544 | 4.54892128 | 13.1768128 | 0.00057 | 0.00249 | F | F | T | F |
| Il5ra | 2.10179819 | 2.23504925 | 13.1746584 | 0.00057 | 0.002492 | T | T | T | T |
| Metap1 | 0.74904276 | 6.7890668 | 13.1714594 | 0.00057 | 0.002495 | T | F | T | F |
| Rims1 | -3.3940746 | 1.54553786 | 13.1640729 | 0.00057 | 0.002503 | T | F | T | T |
| Cggbp1 | 0.77735952 | 6.51919156 | 13.1585666 | 0.00057 | 0.002508 | T | F | T | T |
| Ebag9 | -1.5983185 | 3.62236143 | 13.1524405 | 0.00058 | 0.002514 | F | F | T | T |
| Nfxl1 | -0.8916746 | 6.01004073 | 13.1497437 | 0.00058 | 0.002517 | F | F | F | F |
| Ogfod1 | 0.88073019 | 5.61458684 | 13.1445825 | 0.00058 | 0.002522 | T | F | T | T |
| Tnrc6a | 0.70629725 | 7.36411445 | 13.1430391 | 0.00058 | 0.002523 | T | F | T | F |
| Ubr3 | 0.71887661 | 7.06855344 | 13.1392262 | 0.00058 | 0.002527 | T | F | T | F |
| Olfr365 | 0.95866922 | 5.31885642 | 13.1346279 | 0.00058 | 0.002532 | F | F | F | F |
| Rpn1 | 0.66847102 | 8.13003227 | 13.1317601 | 0.00058 | 0.002534 | F | F | T | F |
| Slc39a11 | 1.12232096 | 4.40591016 | 13.1230884 | 0.00058 | 0.002544 | F | F | F | T |
| Olfr1565 | 1.80258033 | 2.81167859 | 13.1167526 | 0.00058 | 0.00255 | F | F | F | F |
| Ehd3 | -1.5574762 | 3.70239736 | 13.1159984 | 0.00058 | 0.002551 | F | F | T | F |
| Rab11fip2 | -1.9884495 | 2.97609317 | 13.1072434 | 0.00059 | 0.00256 | T | F | T | T |
| Ppp4r3b | 0.78436815 | 6.48461723 | 13.1046371 | 0.00059 | 0.002562 | F | F | T | F |
| Gltp | 0.86342085 | 5.70471739 | 13.0949563 | 0.00059 | 0.002573 | F | F | F | T |
| Olfr735 | 1.09179886 | 4.47618514 | 13.0943883 | 0.00059 | 0.002573 | F | F | F | F |
| Adh5 | -0.7940037 | 6.77444304 | 13.0877571 | 0.00059 | 0.00258 | F | F | F | F |
| Slc41a2 | 2.89770491 | 1.23683759 | 13.0848981 | 0.00059 | 0.002583 | T | F | T | F |
| Tfpi | 1.06146056 | 4.60917301 | 13.0679236 | 0.0006 | 0.002602 | T | F | T | T |
| Zfp213 | -1.3387918 | 4.13681324 | 13.0641276 | 0.0006 | 0.002605 | F | F | T | F |
| Nthl1 | -2.0868339 | 2.83294358 | 13.0638597 | 0.0006 | 0.002605 | F | F | F | F |
| Gbe1 | 1.11613673 | 4.35697485 | 13.0555281 | 0.0006 | 0.002614 | T | F | F | F |
| Abcg2 | -0.9533737 | 5.67080512 | 13.0454973 | 0.0006 | 0.002625 | F | F | F | F |
| Prkcb | -3.8521512 | 1.29822686 | 13.0423676 | 0.0006 | 0.002629 | T | F | T | T |
| Pcdhgb5 | -7.9671326 | 0.55992289 | 13.0392636 | 0.00061 | 0.002632 | F | F | F | F |
| 1810030O0 | 0.87970759 | 5.56426777 | 13.0324978 | 0.00061 | 0.002639 | T | F | T | T |
| Dnajb13 | -7.9458157 | 0.48931285 | 13.032027 | 0.00061 | 0.002639 | F | F | F | T |
| Tmem14c | 0.83344138 | 6.09175566 | 13.0296365 | 0.00061 | 0.002641 | F | F | F | F |
| Ipo8 | -0.83781 | 6.50579141 | 13.0294431 | 0.00061 | 0.002641 | T | F | T | F |
| Zkscan16 | -4.943616 | 0.88950459 | 13.0254832 | 0.00061 | 0.002645 | F | F | T | T |
| Depdc1b | -1.3005635 | 4.2033149 | 13.0228885 | 0.00061 | 0.002648 | T | F | T | T |
| Nsmce4a | -0.8123276 | 6.59710548 | 13.0187503 | 0.00061 | 0.002652 | F | F | F | F |
| Lamb1 | -0.6878211 | 8.11226035 | 13.0175637 | 0.00061 | 0.002652 | F | F | F | T |
| Shisa5 | 1.05738128 | 4.66028632 | 13.0174456 | 0.00061 | 0.002652 | F | F | F | F |
| Pld4 | -7.9482064 | 0.48972433 | 13.0073318 | 0.00061 | 0.002664 | F | F | F | F |

| | | | | | | | | | |
|---|---|---|---|---|---|---|---|---|---|
| Tmed4 | -1.0990826 | 4.82888796 | 12.9952297 | 0.00062 | 0.002677 | T | F | T | T |
| Prph | -7.9149482 | 0.49808648 | 12.9950404 | 0.00062 | 0.002677 | T | F | T | T |
| Uvrag | 0.92408021 | 5.23337421 | 12.9917313 | 0.00062 | 0.00268 | T | F | F | F |
| Acox1 | 0.85029151 | 5.82643904 | 12.9727087 | 0.00062 | 0.002703 | T | F | T | T |
| Xpc | -1.1354368 | 4.70829389 | 12.9704008 | 0.00062 | 0.002705 | F | F | T | F |
| Vps13a | 0.88640576 | 5.86043007 | 12.9697382 | 0.00062 | 0.002705 | T | F | T | T |
| Cd44 | -1.3611168 | 4.16732575 | 12.9645612 | 0.00063 | 0.002711 | T | F | T | F |
| Timm9 | 1.04395635 | 4.65228236 | 12.9628689 | 0.00063 | 0.002712 | F | F | F | F |
| Zfp157 | -1.3383115 | 4.12138775 | 12.9589061 | 0.00063 | 0.002716 | T | F | T | T |
| Hddc3 | 1.07489745 | 4.56470811 | 12.9564403 | 0.00063 | 0.002719 | F | F | F | T |
| Zfp654 | 1.07175061 | 4.5340091 | 12.9561476 | 0.00063 | 0.002719 | T | F | T | T |
| Hnrnpl | -0.6286413 | 9.44824392 | 12.9555755 | 0.00063 | 0.002719 | F | F | T | F |
| Abi1 | 0.87143912 | 5.64295619 | 12.9538749 | 0.00063 | 0.00272 | T | T | F | T |
| Myo5a | -0.8269479 | 6.5517184 | 12.9515425 | 0.00063 | 0.002723 | T | F | T | T |
| Bccip | -0.888589 | 5.97495904 | 12.9499233 | 0.00063 | 0.002724 | F | F | T | F |
| Erh | -0.8313141 | 6.48989487 | 12.9443646 | 0.00063 | 0.00273 | F | F | F | F |
| Kbtbd7 | -1.6114059 | 3.5943591 | 12.943956 | 0.00063 | 0.00273 | T | F | T | T |
| Marcks | -0.6808745 | 8.24250186 | 12.9406497 | 0.00063 | 0.002734 | T | T | T | T |
| Pcbd1 | -1.6072517 | 3.70956811 | 12.9305038 | 0.00063 | 0.002745 | F | F | F | F |
| Wdr83os | 1.15482092 | 4.26225446 | 12.926302 | 0.00064 | 0.00275 | F | F | T | T |
| Pigx | -1.3050198 | 4.18462455 | 12.9234542 | 0.00064 | 0.002753 | F | F | F | T |
| Ly6e | -0.9261349 | 5.70055963 | 12.9199128 | 0.00064 | 0.002757 | F | F | T | F |
| Snap47 | -0.965358 | 5.42897498 | 12.9189309 | 0.00064 | 0.002757 | T | F | T | T |
| Stmn3 | -7.9091675 | 0.49454529 | 12.9154959 | 0.00064 | 0.002761 | F | F | F | F |
| Slc27a2 | -4.9601622 | 0.89264538 | 12.9032586 | 0.00064 | 0.002775 | F | F | F | T |
| Aard | -4.927659 | 0.88227733 | 12.8967668 | 0.00064 | 0.002783 | F | F | F | F |
| Hint2 | -2.8611856 | 1.91469921 | 12.8918496 | 0.00065 | 0.002788 | F | F | F | F |
| Map3k19 | -7.9033708 | 0.46599991 | 12.8832306 | 0.00065 | 0.002798 | F | F | T | T |
| Ptgds | -7.9094549 | 0.49709119 | 12.8788107 | 0.00065 | 0.002803 | F | F | F | F |
| Atn1 | -0.7991216 | 6.6921856 | 12.8773409 | 0.00065 | 0.002805 | F | F | F | F |
| Ccdc120 | -1.3872822 | 4.04949319 | 12.8765792 | 0.00065 | 0.002805 | T | F | T | T |
| Rxylt1 | 1.65408157 | 3.09661076 | 12.8713558 | 0.00065 | 0.002811 | F | F | F | F |
| Ftl1-ps1 | -2.6265359 | 2.15857962 | 12.8615055 | 0.00065 | 0.002823 | F | F | F | F |
| Myt1l | -7.9113873 | 0.46582492 | 12.8583941 | 0.00066 | 0.002826 | T | F | T | T |
| Dph6 | 0.92908267 | 5.18562761 | 12.8575721 | 0.00066 | 0.002826 | T | F | F | T |
| Pds5a | 0.64489931 | 8.43613484 | 12.8557543 | 0.00066 | 0.002828 | F | F | T | F |
| 9930012K1 | 1.77131457 | 2.80621499 | 12.8529139 | 0.00066 | 0.002831 | F | F | F | F |
| Igsf1 | -2.3678009 | 2.4523805 | 12.847289 | 0.00066 | 0.002838 | T | F | T | F |
| Eif2ak2 | 1.00021902 | 4.82056512 | 12.8421421 | 0.00066 | 0.002844 | T | T | T | T |
| Mrpl3 | 1.06566551 | 4.54179875 | 12.8400021 | 0.00066 | 0.002846 | T | F | T | T |
| Galnt1 | 0.71221011 | 7.29769746 | 12.8386428 | 0.00066 | 0.002847 | T | T | T | T |
| Ppp2r3a | 0.97864534 | 4.9424612 | 12.8372624 | 0.00066 | 0.002848 | T | T | T | T |

| | | | | | | | | | |
|---|---|---|---|---|---|---|---|---|---|
| Plscr3 | -1.2427701 | 4.31410271 | 12.8277359 | 0.00066 | 0.002859 | F | F | F | F |
| Dpy30 | -0.9528442 | 5.49749896 | 12.8276441 | 0.00066 | 0.002859 | T | F | F | T |
| Tek | -2.4193058 | 2.64675062 | 12.8268667 | 0.00066 | 0.002859 | F | F | F | T |
| Dnaic2 | -7.8942794 | 0.46099127 | 12.8223862 | 0.00067 | 0.002865 | T | F | F | F |
| Hpgd | 5.95121744 | -0.249959 | 12.8211585 | 0.00067 | 0.002866 | T | F | T | T |
| Ebp | -1.6447576 | 3.54273858 | 12.8207066 | 0.00067 | 0.002866 | F | F | F | T |
| Lemd2 | 0.98433686 | 4.93427692 | 12.8178103 | 0.00067 | 0.002869 | T | F | F | T |
| Plekha6 | 0.78366854 | 6.38427571 | 12.8163465 | 0.00067 | 0.00287 | T | F | T | F |
| Ptk6 | 3.37802115 | 0.7390432 | 12.8066062 | 0.00067 | 0.002882 | F | F | F | T |
| Rassf10 | -4.931942 | 0.87399249 | 12.8032434 | 0.00067 | 0.002886 | T | F | T | T |
| Map4k3 | 0.82956429 | 5.92386708 | 12.7905447 | 0.00068 | 0.002901 | T | F | F | T |
| Sgms1 | -1.1935805 | 4.44788058 | 12.7825251 | 0.00068 | 0.002911 | T | F | T | F |
| Samd1 | -0.8575615 | 6.17753946 | 12.7741085 | 0.00068 | 0.002921 | F | F | F | F |
| Fkrp | 0.87783698 | 5.56985462 | 12.7726972 | 0.00068 | 0.002923 | F | F | F | T |
| Tjp3 | 1.31765103 | 3.94088902 | 12.7627035 | 0.00068 | 0.002935 | F | F | F | F |
| Sbno2 | 0.87681989 | 5.52137526 | 12.7571773 | 0.00069 | 0.002942 | T | F | F | F |
| Muc4 | 7.86727439 | -0.5955185 | 12.7772092 | 0.00069 | 0.002943 | T | F | T | F |
| Vmn1r90 | 1.982232 | 2.38096766 | 12.7547782 | 0.00069 | 0.002944 | F | F | F | F |
| Sirt6 | 0.91938311 | 5.21223427 | 12.7530413 | 0.00069 | 0.002945 | F | F | F | F |
| Nup214 | 0.65218019 | 8.05486825 | 12.7523084 | 0.00069 | 0.002946 | F | F | T | F |
| Slx1b | 0.84768272 | 5.93341249 | 12.7516542 | 0.00069 | 0.002946 | F | F | T | T |
| Lyplal1 | -2.3272412 | 2.48819636 | 12.7494527 | 0.00069 | 0.002948 | F | F | F | F |
| Slc26a9 | -7.8912296 | 0.45383315 | 12.7492976 | 0.00069 | 0.002948 | T | F | T | T |
| Myrfl | -7.8975451 | 0.48745613 | 12.744944 | 0.00069 | 0.002953 | F | F | F | T |
| Phyh | -1.3414008 | 4.0893059 | 12.741739 | 0.00069 | 0.002957 | F | F | F | T |
| Nr1h3 | -7.8989112 | 0.48995809 | 12.7301722 | 0.00069 | 0.002971 | F | F | F | F |
| Ctf1 | -4.206737 | 1.08757061 | 12.726995 | 0.00069 | 0.002975 | T | F | F | F |
| Tmub1 | -3.2367237 | 1.66294945 | 12.7265051 | 0.00069 | 0.002975 | F | F | F | F |
| Bhlha15 | 2.02246054 | 2.29822632 | 12.724125 | 0.0007 | 0.002978 | T | F | T | T |
| Acrbp | -1.5891933 | 3.57843878 | 12.706839 | 0.0007 | 0.003 | F | F | F | F |
| Nus1 | 0.72738437 | 6.83562056 | 12.7037648 | 0.0007 | 0.003003 | F | F | T | F |
| Pheta2 | -7.8788855 | 0.45777356 | 12.7011704 | 0.0007 | 0.003006 | F | F | T | F |
| Lipc | -7.8812022 | 0.46538971 | 12.6938053 | 0.00071 | 0.003016 | F | F | F | F |
| Camk2n1 | -2.3219814 | 2.48608217 | 12.6858367 | 0.00071 | 0.003026 | F | F | T | T |
| Sec63 | 0.76765107 | 6.48045785 | 12.6788646 | 0.00071 | 0.003034 | T | T | T | T |
| Moxd1 | -2.4299366 | 2.35188659 | 12.676451 | 0.00071 | 0.003037 | T | F | T | F |
| Stx3 | -0.9191519 | 5.70233558 | 12.6647813 | 0.00071 | 0.003052 | T | F | T | F |
| Rln3 | -7.8956401 | 0.44913073 | 12.6580468 | 0.00072 | 0.003061 | F | F | F | F |
| Nkain1 | 0.91409435 | 5.29243176 | 12.6563903 | 0.00072 | 0.003063 | F | F | T | F |
| Anxa3 | 1.73038743 | 2.86047173 | 12.6548916 | 0.00072 | 0.003064 | T | F | T | T |
| Eif2ak4 | 0.81832934 | 5.98681439 | 12.6428214 | 0.00072 | 0.00308 | F | F | F | F |
| Tbc1d12 | 1.13342719 | 4.28510027 | 12.6360481 | 0.00072 | 0.003089 | T | F | F | F |

| | | | | | | | | | |
|---|---|---|---|---|---|---|---|---|---|
| Tpd52l1 | -2.5500345 | 2.19049004 | 12.6350516 | 0.00072 | 0.003089 | F | F | F | F |
| Fasn | -0.5980756 | 9.95455452 | 12.634559 | 0.00072 | 0.003089 | T | F | T | T |
| Abhd14a | -1.5852266 | 3.58126552 | 12.6306536 | 0.00073 | 0.003094 | F | F | F | F |
| Plpp4 | -7.871859 | 0.44466857 | 12.6249983 | 0.00073 | 0.003101 | F | F | F | F |
| Dynlt1a | 1.33345028 | 3.77094227 | 12.6027425 | 0.00073 | 0.003132 | F | F | T | F |
| Appbp2 | 0.64337544 | 8.17261772 | 12.602304 | 0.00073 | 0.003132 | T | T | F | T |
| Cstf2t | -0.8293297 | 6.41727452 | 12.6015612 | 0.00073 | 0.003132 | T | F | F | F |
| Nfe2l3 | -4.9128196 | 0.85701698 | 12.5998794 | 0.00074 | 0.003134 | T | F | F | F |
| Bcap31 | 0.82535004 | 6.00150244 | 12.5991925 | 0.00074 | 0.003134 | T | F | F | F |
| Rad54l2 | -0.7023948 | 7.82794367 | 12.5850355 | 0.00074 | 0.003153 | T | F | T | T |
| Mcm6 | -0.6565779 | 8.55299341 | 12.5849374 | 0.00074 | 0.003153 | T | F | T | T |
| Prkca | 1.43931844 | 3.47204378 | 12.5786479 | 0.00074 | 0.003161 | T | F | F | T |
| Paqr9 | -7.8682976 | 0.43859528 | 12.5741396 | 0.00074 | 0.003167 | F | F | F | F |
| Zbtb39 | -0.7995532 | 6.58315398 | 12.5649794 | 0.00075 | 0.003179 | T | F | T | T |
| Cd59a | -3.0680757 | 1.71356463 | 12.5607119 | 0.00075 | 0.003184 | T | F | T | T |
| Naaladl1 | -2.5950614 | 2.12832649 | 12.5589928 | 0.00075 | 0.003186 | F | F | T | F |
| Mapk1ip1 | -2.1675123 | 2.65986272 | 12.5585801 | 0.00075 | 0.003186 | F | F | F | F |
| Tmed9 | 0.7124039 | 7.02747644 | 12.5497706 | 0.00075 | 0.003198 | F | F | F | T |
| Zbtb8b | -3.0318219 | 1.71751695 | 12.5491142 | 0.00075 | 0.003198 | T | F | T | F |
| Loxl3 | -7.8628383 | 0.44411581 | 12.5436913 | 0.00075 | 0.003206 | T | F | T | T |
| Rrp7a | 0.68818057 | 7.21267342 | 12.5430102 | 0.00075 | 0.003206 | F | F | F | T |
| Rpgrip1 | 7.84276994 | -0.616274 | 12.5549446 | 0.00076 | 0.003218 | T | F | F | T |
| Rpl36a | -0.783319 | 6.75176577 | 12.5049181 | 0.00077 | 0.00326 | F | F | F | F |
| Rnf187 | -0.6930526 | 7.7651187 | 12.4997764 | 0.00077 | 0.003266 | F | F | T | F |
| Scamp4 | 0.82177006 | 5.98168428 | 12.4916139 | 0.00077 | 0.003278 | T | F | T | T |
| Acsf2 | -1.7137171 | 3.31562753 | 12.4839994 | 0.00077 | 0.003288 | F | F | T | T |
| Afg1l | 1.55984877 | 3.17284475 | 12.4835568 | 0.00077 | 0.003288 | T | F | F | F |
| Ptpn3 | 0.89172363 | 5.41040834 | 12.4797711 | 0.00078 | 0.003293 | F | T | F | T |
| Tm9sf1 | 0.8237848 | 5.90446793 | 12.4788299 | 0.00078 | 0.003294 | T | F | T | F |
| Rfx3 | -0.8705325 | 6.24877691 | 12.4770316 | 0.00078 | 0.003296 | T | T | T | T |
| Mios | 0.88452474 | 5.38935112 | 12.4700831 | 0.00078 | 0.003305 | T | F | T | T |
| Mapk10 | -7.8742983 | 0.43079063 | 12.4683012 | 0.00078 | 0.003307 | T | F | T | T |
| Scarf1 | -4.1852582 | 1.06565633 | 12.4644749 | 0.00078 | 0.003312 | T | F | F | F |
| Rnf214 | -1.0117069 | 5.0974692 | 12.4641716 | 0.00078 | 0.003312 | F | F | F | T |
| Oxct1 | 0.82117523 | 6.01975845 | 12.4566016 | 0.00078 | 0.003323 | T | F | T | T |
| Cep85 | -0.943599 | 5.46640043 | 12.4528736 | 0.00078 | 0.003328 | T | F | T | F |
| Xpo5 | -0.6826386 | 7.91242741 | 12.4512974 | 0.00079 | 0.003329 | F | F | T | T |
| Canx | 0.57698717 | 9.75731346 | 12.4392516 | 0.00079 | 0.003347 | F | F | T | F |
| Armc9 | -1.8631703 | 3.09292478 | 12.4326639 | 0.00079 | 0.003356 | T | F | T | F |
| Mydgf | 0.79147615 | 6.17832548 | 12.4302558 | 0.00079 | 0.003359 | F | F | F | F |
| Plekhd1 | 2.97141504 | 1.01606633 | 12.4219359 | 0.0008 | 0.003371 | F | F | T | T |
| Mtrr | 0.8692622 | 5.51343582 | 12.420251 | 0.0008 | 0.003373 | F | F | F | F |

| | | | | | | | | | |
|---|---|---|---|---|---|---|---|---|---|
| Snx13 | 1.30233395 | 3.82287093 | 12.4134007 | 0.0008 | 0.003382 | T | T | F | T |
| Kpna6 | 0.66108994 | 7.65156689 | 12.390414 | 0.00081 | 0.003416 | T | F | T | T |
| Gm14399 | 2.03909558 | 2.30580946 | 12.3853833 | 0.00081 | 0.003423 | F | F | F | F |
| Ap3m1 | 0.77424607 | 6.3307924 | 12.378478 | 0.00081 | 0.003433 | T | F | T | F |
| A930004D1 | -2.0492673 | 2.81103439 | 12.3772139 | 0.00081 | 0.003435 | T | F | F | F |
| Golt1b | 0.74254599 | 6.63212114 | 12.3740866 | 0.00081 | 0.003439 | T | F | T | F |
| Tmed7 | 0.76823923 | 6.37679035 | 12.3699108 | 0.00081 | 0.003445 | T | F | T | T |
| Primpol | 1.63564482 | 3.00481245 | 12.3665507 | 0.00082 | 0.003449 | T | F | T | T |
| Eif2d | -0.8556877 | 6.11536215 | 12.3661081 | 0.00082 | 0.003449 | F | F | T | T |
| Ldha | -0.5558444 | 10.8297792 | 12.3591097 | 0.00082 | 0.003459 | T | F | F | F |
| Gm45716 | -3.4616547 | 1.41252303 | 12.3577625 | 0.00082 | 0.003461 | F | F | F | F |
| Fgfr1op | -0.968859 | 5.25487339 | 12.3505313 | 0.00082 | 0.003471 | F | F | T | F |
| Cox17 | 0.95458969 | 4.97048226 | 12.3455602 | 0.00082 | 0.003478 | F | F | T | T |
| Bcl11b | -2.4015247 | 2.40529831 | 12.344312 | 0.00082 | 0.003479 | T | F | T | T |
| Tmem106b | 0.90290748 | 5.27297389 | 12.3381637 | 0.00083 | 0.003488 | T | F | T | F |
| Pde4d | -1.6231599 | 3.50105309 | 12.3211652 | 0.00083 | 0.003514 | F | F | T | T |
| Blnk | -7.8634808 | 0.41856213 | 12.317603 | 0.00083 | 0.003519 | F | F | F | F |
| Serbp1 | -0.5933358 | 9.88119352 | 12.3169045 | 0.00083 | 0.00352 | T | F | T | F |
| Ifit1 | 5.89552705 | -0.3047825 | 12.3150359 | 0.00083 | 0.003522 | T | F | F | T |
| Sp9 | -7.8477471 | 0.41359947 | 12.313307 | 0.00084 | 0.003524 | T | F | F | T |
| Eef1e1 | 0.87973605 | 5.37432118 | 12.3125951 | 0.00084 | 0.003524 | T | F | F | T |
| Mau2 | 0.78846806 | 6.29031765 | 12.3110612 | 0.00084 | 0.003526 | F | F | T | T |
| Pacsin1 | 1.73076572 | 2.88571618 | 12.3102512 | 0.00084 | 0.003526 | F | F | F | F |
| Adamts8 | -7.8381636 | 0.4202044 | 12.3102241 | 0.00084 | 0.003526 | T | F | F | F |
| Mpdu1 | 0.90115729 | 5.22221166 | 12.3005097 | 0.00084 | 0.003541 | F | F | T | F |
| Pdxdc1 | 0.71472164 | 6.84395833 | 12.2906778 | 0.00084 | 0.003555 | F | F | T | T |
| Nol8 | 0.73674761 | 6.63835946 | 12.2905794 | 0.00084 | 0.003555 | T | F | T | T |
| Zfp946 | -2.2295809 | 2.54839939 | 12.2862198 | 0.00085 | 0.003561 | T | F | F | F |
| Ccdc103 | -7.8386284 | 0.41159301 | 12.2849134 | 0.00085 | 0.003563 | F | F | T | T |
| Cmtm8 | 1.34829435 | 3.64848854 | 12.2680805 | 0.00085 | 0.003589 | T | F | T | F |
| Ahctf1 | 0.61042925 | 8.76188695 | 12.2606983 | 0.00086 | 0.0036 | F | F | T | T |
| Ilf2 | -0.7715867 | 6.73448054 | 12.2560008 | 0.00086 | 0.003607 | T | F | F | F |
| Lrrc3 | 1.04770464 | 4.5361621 | 12.2473686 | 0.00086 | 0.00362 | F | F | F | T |
| Slc25a12 | 1.6344498 | 2.98470948 | 12.2371864 | 0.00086 | 0.003636 | T | F | T | T |
| Fat2 | -7.8681281 | 0.46969448 | 12.2330926 | 0.00087 | 0.003642 | F | F | T | F |
| Klf14 | -7.8296474 | 0.40844613 | 12.2240434 | 0.00087 | 0.003656 | F | F | T | F |
| Leprot | -1.1750406 | 4.4137196 | 12.2222561 | 0.00087 | 0.003659 | T | F | F | F |
| Gcc1 | 0.95649165 | 4.9250796 | 12.2190144 | 0.00087 | 0.003663 | T | F | F | F |
| Tmem267 | -1.6759574 | 3.34509118 | 12.2179816 | 0.00087 | 0.003664 | F | F | F | F |
| Inafm2 | -1.583285 | 3.52563103 | 12.2175713 | 0.00087 | 0.003664 | F | F | F | F |
| Hnrnpd | -0.6603714 | 8.25939923 | 12.2136613 | 0.00087 | 0.003669 | T | F | T | T |
| 4930412O1 | 3.4596942 | 0.612865 | 12.2135712 | 0.00087 | 0.003669 | F | F | F | F |

| | | | | | | | | | |
|---|---|---|---|---|---|---|---|---|---|
| Laptm5 | 3.27348808 | 0.79697892 | 12.2130043 | 0.00087 | 0.003669 | T | F | T | F |
| Zfp950 | 1.24199309 | 3.94452007 | 12.2128558 | 0.00087 | 0.003669 | F | F | T | F |
| Kiss1r | 3.00781455 | 0.96621116 | 12.2068961 | 0.00088 | 0.003678 | F | F | F | T |
| Cldn7 | -0.9690769 | 5.25184413 | 12.2060287 | 0.00088 | 0.003679 | F | F | F | F |
| Ezh2 | -0.7307096 | 7.05129781 | 12.2023877 | 0.00088 | 0.003684 | F | F | F | F |
| Zdhhc7 | 0.85835243 | 5.54791202 | 12.2001115 | 0.00088 | 0.003687 | F | F | T | F |
| Hmgb2 | -0.6813298 | 7.8172998 | 12.1991835 | 0.00088 | 0.003688 | T | T | T | T |
| Cd74 | 3.1752559 | 0.82164269 | 12.1987085 | 0.00088 | 0.003688 | T | F | T | T |
| Rmdn1 | 1.82106732 | 2.60955164 | 12.195544 | 0.00088 | 0.003693 | F | F | F | T |
| Map7d2 | -2.2895546 | 2.45487629 | 12.1939278 | 0.00088 | 0.003695 | T | T | F | T |
| Hsd17b12 | 0.77921928 | 6.29193381 | 12.1928188 | 0.00088 | 0.003696 | F | F | F | F |
| Klf5 | -1.0900989 | 4.72245543 | 12.1891651 | 0.00088 | 0.003701 | T | F | F | F |
| Zfp202 | -3.716292 | 1.4310172 | 12.1819439 | 0.00089 | 0.003712 | T | T | T | T |
| Gm28778 | -7.826628 | 0.40215096 | 12.1790301 | 0.00089 | 0.003716 | F | F | F | F |
| Ankrd10 | -0.9213608 | 5.54678487 | 12.1780302 | 0.00089 | 0.003717 | T | F | F | F |
| Phyhip | -7.8272275 | 0.40027134 | 12.1679544 | 0.00089 | 0.003734 | T | F | F | T |
| Rft1 | 1.45931372 | 3.43821151 | 12.1645456 | 0.00089 | 0.003739 | T | F | F | T |
| Vmn1r174 | 4.27521304 | 0.21710031 | 12.150378 | 0.0009 | 0.003762 | F | F | F | F |
| Ssh3 | 1.55904068 | 3.13304423 | 12.1483989 | 0.0009 | 0.003764 | F | F | F | F |
| Krtcap2 | -0.9486891 | 5.37648174 | 12.1387098 | 0.0009 | 0.00378 | F | F | F | T |
| Hagh | 1.30879903 | 3.77682644 | 12.1338809 | 0.00091 | 0.003787 | F | F | F | F |
| Pnpla1 | 2.38511168 | 1.63753442 | 12.1321561 | 0.00091 | 0.00379 | T | F | F | F |
| Runx3 | -4.5708563 | 0.93122437 | 12.1288319 | 0.00091 | 0.003795 | T | T | T | T |
| Wnt9b | -7.8198154 | 0.40914206 | 12.1204191 | 0.00091 | 0.003808 | T | F | T | T |
| Pithd1 | 0.80339648 | 5.95526876 | 12.1028276 | 0.00092 | 0.003838 | T | F | T | T |
| Chrna9 | -7.8710412 | 0.41385798 | 12.0932143 | 0.00092 | 0.003854 | F | F | F | F |
| Ttll1 | -1.6693636 | 3.34055468 | 12.0915125 | 0.00092 | 0.003856 | F | F | T | T |
| Tmem128 | 1.04530135 | 4.48914484 | 12.0881969 | 0.00092 | 0.00386 | F | F | F | F |
| Tdp1 | -0.9401672 | 5.40159765 | 12.0879095 | 0.00092 | 0.00386 | T | F | F | T |
| Gars | 0.61532705 | 8.47196631 | 12.0869868 | 0.00092 | 0.003861 | F | F | F | F |
| Prm1 | 7.79185162 | -0.6591727 | 12.1043504 | 0.00093 | 0.003864 | F | F | F | F |
| Id1 | -0.8511839 | 6.22664656 | 12.0698012 | 0.00093 | 0.00389 | F | F | F | F |
| Fgf10 | -2.291215 | 2.4740321 | 12.0661213 | 0.00093 | 0.003895 | T | F | T | T |
| Spata18 | -7.8170081 | 0.41044936 | 12.0650615 | 0.00093 | 0.003897 | F | F | T | F |
| Sh3rf2 | 3.15448715 | 0.81737152 | 12.0636044 | 0.00093 | 0.003898 | T | F | T | F |
| Ctbp1 | -0.6938036 | 7.54697839 | 12.0635514 | 0.00093 | 0.003898 | F | F | F | T |
| Mcm8 | -0.9996021 | 5.10500191 | 12.0588595 | 0.00094 | 0.003905 | T | T | F | T |
| Ube2z | 0.64740718 | 7.72791717 | 12.0544705 | 0.00094 | 0.003912 | F | F | T | T |
| Dcc | -3.2746651 | 1.48312426 | 12.054172 | 0.00094 | 0.003912 | T | F | T | F |
| Stam2 | 0.87387095 | 5.37222963 | 12.0504522 | 0.00094 | 0.003918 | T | F | T | T |
| Hdac9 | -3.7960579 | 1.23047808 | 12.0476069 | 0.00094 | 0.003922 | T | T | T | T |
| Thnsl2 | -2.005452 | 2.82164437 | 12.0463621 | 0.00094 | 0.003923 | F | F | F | F |

| | | | | | | | | | |
|---|---|---|---|---|---|---|---|---|---|
| Ptpra | -0.893702 | 5.69537214 | 12.0407193 | 0.00094 | 0.003933 | F | F | F | T |
| Pcdhgc4 | -2.7019929 | 1.94040038 | 12.0402474 | 0.00094 | 0.003933 | F | F | F | F |
| Atp1a4 | 4.8623088 | -0.0445524 | 12.0397226 | 0.00094 | 0.003933 | F | F | T | T |
| Pcdhga12 | -8.1463491 | 0.73412737 | 12.0393758 | 0.00094 | 0.003933 | F | F | F | F |
| Akr1c13 | -7.8423846 | 0.39562971 | 12.0354419 | 0.00095 | 0.003939 | T | F | F | T |
| Anapc7 | 0.77793093 | 6.18524117 | 12.0253562 | 0.00095 | 0.003956 | F | F | F | T |
| Col5a3 | 3.01270994 | 0.99685885 | 12.017238 | 0.00095 | 0.003969 | T | F | F | T |
| Cnpy1 | -7.8346952 | 0.39118427 | 12.0172234 | 0.00095 | 0.003969 | T | F | T | T |
| Gfpt1 | 0.71276883 | 6.85129732 | 12.0125081 | 0.00096 | 0.003977 | T | F | T | F |
| Smim13 | 0.99795307 | 4.76071271 | 12.0063203 | 0.00096 | 0.003987 | F | F | T | T |
| Eif2ak3 | 0.87048612 | 5.37031365 | 12.0034932 | 0.00096 | 0.003991 | T | T | T | T |
| Rasgrp4 | 1.50643946 | 3.22057698 | 12.0012352 | 0.00096 | 0.003995 | T | F | T | F |
| Troap | -1.0652083 | 4.77979338 | 11.9989145 | 0.00096 | 0.003998 | F | F | F | F |
| Rpl22 | -0.7268858 | 7.18141523 | 11.9928698 | 0.00096 | 0.004008 | T | F | T | F |
| H2az1 | -0.6559446 | 8.18540882 | 11.989969 | 0.00097 | 0.004013 | F | F | F | F |
| Tubg1 | -0.8881026 | 5.77235222 | 11.9869375 | 0.00097 | 0.004017 | F | F | F | F |
| Nt5c | 1.1228264 | 4.24174865 | 11.9846667 | 0.00097 | 0.004021 | F | F | F | F |
| Vmn1r224 | 1.94858046 | 2.43164162 | 11.980346 | 0.00097 | 0.004028 | F | F | F | F |
| Hrct1 | 4.76640129 | -0.0351066 | 11.9752997 | 0.00097 | 0.004036 | T | F | T | T |
| Bptf | -0.6438831 | 8.58511529 | 11.9717248 | 0.00097 | 0.004042 | T | T | T | T |
| Rnpepl1 | -0.9461912 | 5.29836739 | 11.9652668 | 0.00098 | 0.004053 | F | F | T | T |
| Mpv17l2 | 1.53763994 | 3.1692782 | 11.9646147 | 0.00098 | 0.004053 | F | F | F | F |
| Rbm8a | -0.7934793 | 6.52664846 | 11.9637797 | 0.00098 | 0.004054 | F | F | T | F |
| Arhgef39 | 1.64577229 | 2.92249795 | 11.9571487 | 0.00098 | 0.004065 | T | F | F | F |
| Arrdc4 | -1.2097623 | 4.30724342 | 11.9569851 | 0.00098 | 0.004065 | T | F | F | T |
| Olfr718-ps1 | -1.1464474 | 4.49614991 | 11.9555604 | 0.00098 | 0.004067 | F | F | F | F |
| Tapbp | 1.0927894 | 4.2819006 | 11.9545665 | 0.00098 | 0.004068 | T | F | T | T |
| Lrrc75a | -2.6732998 | 1.97368658 | 11.9534569 | 0.00098 | 0.004069 | F | F | T | T |
| Fubp1 | -0.6433538 | 8.45009715 | 11.9496653 | 0.00098 | 0.004075 | T | F | T | T |
| Moap1 | -1.7750013 | 3.14577492 | 11.9482956 | 0.00098 | 0.004077 | T | F | T | T |
| Rnf13 | 1.21078373 | 3.98091298 | 11.9440437 | 0.00099 | 0.004084 | T | F | F | T |
| Zfyve19 | -1.4308086 | 3.8244955 | 11.9425476 | 0.00099 | 0.004086 | F | F | F | F |
| Aoc1 | 2.34599897 | 1.93253162 | 11.9352898 | 0.00099 | 0.004099 | F | F | F | F |
| Fbxo22 | 0.75588278 | 6.3577028 | 11.9331106 | 0.00099 | 0.004102 | F | F | F | F |
| Kctd20 | 0.68701312 | 7.02861596 | 11.9326023 | 0.00099 | 0.004102 | T | F | T | F |
| Fam214a | 1.82738331 | 2.69680368 | 11.9281401 | 0.00099 | 0.004109 | F | F | F | F |
| Mbl2 | -7.8186084 | 0.42145037 | 11.92157 | 0.001 | 0.004121 | F | F | F | F |
| Sos2 | 0.98635664 | 4.74053554 | 11.9122061 | 0.001 | 0.004137 | T | F | F | F |
| Pym1 | -0.9292043 | 5.40027988 | 11.9118358 | 0.001 | 0.004137 | F | F | T | F |
| Npm1 | -0.5413133 | 10.9433575 | 11.910584 | 0.001 | 0.004139 | T | F | F | F |
| Zfp39 | -1.5771367 | 3.48672006 | 11.9055953 | 0.001 | 0.004148 | F | F | F | F |
| Tspan15 | -2.2668576 | 2.44394422 | 11.8993695 | 0.00101 | 0.004158 | T | F | T | F |

| | | | | | | | | | |
|---|---|---|---|---|---|---|---|---|---|
| Mtbp | 0.90454982 | 5.12789234 | 11.8946113 | 0.00101 | 0.004167 | F | F | T | F |
| Tcf19 | -1.8616627 | 3.02014529 | 11.8939075 | 0.00101 | 0.004167 | F | F | F | F |
| Gsto1 | -1.0203862 | 5.03105901 | 11.8914515 | 0.00101 | 0.004171 | F | F | F | F |
| St6galnac2 | 1.3865514 | 3.57429214 | 11.8878067 | 0.00101 | 0.004177 | T | F | F | T |
| Ppef1 | 4.08092299 | 0.34766502 | 11.8767859 | 0.00102 | 0.004197 | T | F | T | F |
| Cpne8 | 1.50075712 | 3.39712165 | 11.8712044 | 0.00102 | 0.004207 | T | T | T | T |
| Zfp185 | -1.6885138 | 3.27736693 | 11.8582041 | 0.00103 | 0.004231 | T | F | T | T |
| Nat10 | 0.66274852 | 7.37223013 | 11.8518629 | 0.00103 | 0.004242 | T | F | F | T |
| Ing5 | 0.75318887 | 6.35577248 | 11.8456748 | 0.00103 | 0.004253 | F | F | T | F |
| Pfkfb3 | -1.043439 | 4.91994277 | 11.845279 | 0.00103 | 0.004253 | T | F | T | T |
| Pqlc3 | 1.75264723 | 2.69027071 | 11.8426404 | 0.00103 | 0.004257 | T | F | F | F |
| Sp7 | -7.8038275 | 0.40620454 | 11.8364229 | 0.00104 | 0.004268 | F | F | F | T |
| Eftud2 | -0.6415452 | 8.43250916 | 11.8302174 | 0.00104 | 0.00428 | T | F | F | F |
| Arpc3 | 0.81249053 | 5.9245118 | 11.8292225 | 0.00104 | 0.004281 | F | F | T | T |
| Taf7 | -0.8784307 | 5.71956044 | 11.8250152 | 0.00104 | 0.004288 | F | F | T | T |
| Dhdh | -0.9031674 | 5.69814565 | 11.8131422 | 0.00105 | 0.00431 | T | F | T | T |
| H4f16 | -0.6626293 | 7.95282246 | 11.811996 | 0.00105 | 0.004312 | F | F | F | F |
| Samd4 | -1.5859224 | 3.50571675 | 11.8078791 | 0.00105 | 0.004319 | F | F | T | T |
| Gng11 | -4.8276425 | 0.79143151 | 11.8046356 | 0.00105 | 0.004324 | F | F | F | F |
| Acvrl1 | 1.4650817 | 3.50801443 | 11.8012768 | 0.00105 | 0.00433 | T | F | T | F |
| Igsf23 | -2.4864584 | 2.13480413 | 11.7962177 | 0.00105 | 0.004339 | T | F | T | T |
| Top2a | -0.6068903 | 9.46286949 | 11.7935886 | 0.00106 | 0.004344 | T | F | T | F |
| Hecw2 | -2.474024 | 2.2138402 | 11.7897395 | 0.00106 | 0.00435 | F | T | T | T |
| Pde8b | -4.846131 | 0.7923374 | 11.7883512 | 0.00106 | 0.004352 | T | F | T | T |
| Sdr39u1 | 1.72900416 | 2.7279875 | 11.7813897 | 0.00106 | 0.004365 | F | F | F | F |
| Pcyt1a | 0.83113719 | 5.6485145 | 11.7785922 | 0.00106 | 0.004369 | F | F | T | F |
| Trank1 | -7.7996257 | 0.36449605 | 11.7784976 | 0.00106 | 0.004369 | F | F | T | F |
| 1700008O0 | -7.8283742 | 0.37714783 | 11.7593391 | 0.00107 | 0.004404 | T | F | F | T |
| Slc25a20 | 1.27516672 | 3.77603769 | 11.7592306 | 0.00107 | 0.004404 | T | F | T | T |
| Rnaseh2c | -1.1325171 | 4.57130247 | 11.7591672 | 0.00107 | 0.004404 | F | F | F | T |
| 6030458C1 | 0.77346663 | 6.13535956 | 11.7591465 | 0.00107 | 0.004404 | T | F | T | T |
| Trap1 | 0.6659118 | 7.21733841 | 11.7538315 | 0.00107 | 0.004414 | F | F | T | F |
| Ppp1r8 | -0.7872044 | 6.45059599 | 11.7465261 | 0.00108 | 0.004428 | F | F | T | F |
| Hadh | -0.7572891 | 6.70549932 | 11.7452022 | 0.00108 | 0.004429 | T | F | T | T |
| Efna1 | -1.6994056 | 3.25959098 | 11.7417678 | 0.00108 | 0.004436 | F | F | T | F |
| Kcp | -4.2010835 | 1.27125789 | 11.7397001 | 0.00108 | 0.004439 | F | F | F | F |
| Arhgef2 | -0.7287069 | 6.97070924 | 11.7365453 | 0.00108 | 0.004444 | T | F | T | T |
| Dusp18 | -1.6184835 | 3.40200747 | 11.7301954 | 0.00109 | 0.004456 | T | T | T | T |
| 1810009A1 | -0.9607639 | 5.28025031 | 11.7215649 | 0.00109 | 0.004473 | F | F | F | T |
| Stx1b | -1.6576868 | 3.34711775 | 11.7190058 | 0.00109 | 0.004477 | F | F | F | F |
| Gsr | 0.68011565 | 7.02149963 | 11.7184308 | 0.00109 | 0.004478 | T | F | F | F |
| Clcn2 | -1.7147938 | 3.22530799 | 11.7173246 | 0.00109 | 0.004479 | F | F | F | F |

| | | | | | | | | | |
|---|---|---|---|---|---|---|---|---|---|
| Runx1 | -1.7853918 | 3.10241969 | 11.7082353 | 0.0011 | 0.004497 | T | F | T | T |
| Sgsm1 | 1.11646072 | 4.18710766 | 11.6989349 | 0.0011 | 0.004515 | T | F | T | T |
| Etnk1 | -0.8236695 | 6.22113835 | 11.6966004 | 0.0011 | 0.004519 | T | F | T | F |
| B4gat1 | 1.06498064 | 4.34342258 | 11.6873201 | 0.00111 | 0.004537 | T | F | F | F |
| Timeless | -0.7388402 | 6.84870279 | 11.6843679 | 0.00111 | 0.004542 | T | F | F | T |
| Acaa1a | 0.85167421 | 5.42380893 | 11.6795212 | 0.00111 | 0.004551 | F | F | F | T |
| Ube2e2 | -2.1659543 | 2.53730611 | 11.6744705 | 0.00111 | 0.004561 | F | F | T | F |
| Slc30a3 | -7.7722627 | 0.35319501 | 11.6691029 | 0.00112 | 0.004571 | F | F | T | F |
| Kdm5a | 0.70298679 | 6.87671385 | 11.6565737 | 0.00112 | 0.004596 | T | T | T | T |
| Miga2 | 0.84120789 | 5.49923004 | 11.6519995 | 0.00113 | 0.004604 | F | F | T | F |
| Mon1a | 1.20657207 | 3.94485884 | 11.6518026 | 0.00113 | 0.004604 | F | F | F | F |
| Prx | -1.8711212 | 2.95267177 | 11.6498447 | 0.00113 | 0.004607 | T | F | T | F |
| Fam193b | -0.8391429 | 5.99449809 | 11.6394621 | 0.00113 | 0.004628 | F | F | F | F |
| Sept1 | -7.7658752 | 0.35722104 | 11.6356906 | 0.00113 | 0.004635 | F | F | T | F |
| Rgl1 | -0.9771302 | 5.10823335 | 11.6289505 | 0.00114 | 0.004648 | F | F | T | F |
| Vmn2r52 | 1.49438262 | 3.32263845 | 11.6289309 | 0.00114 | 0.004648 | F | F | F | F |
| Rprd1a | -0.840713 | 6.00317132 | 11.6191496 | 0.00114 | 0.004668 | T | T | T | T |
| Dppa2 | 3.75608274 | 0.38820385 | 11.6157351 | 0.00114 | 0.004674 | T | F | F | T |
| Fam184a | -2.9843677 | 1.63006863 | 11.6131427 | 0.00115 | 0.004678 | T | F | T | F |
| Ilrun | 0.63756587 | 7.76570542 | 11.6047317 | 0.00115 | 0.004695 | F | F | T | F |
| Cln5 | 0.98118495 | 4.69277271 | 11.6027535 | 0.00115 | 0.004699 | F | F | T | T |
| H6pd | -1.444947 | 3.71810491 | 11.5977031 | 0.00115 | 0.004709 | F | F | T | F |
| Ift46 | -0.9343957 | 5.31887196 | 11.5941954 | 0.00116 | 0.004715 | T | F | T | T |
| Tmem254c | -1.4189436 | 3.84064131 | 11.5908827 | 0.00116 | 0.004721 | F | F | T | F |
| Cep120 | 0.73381564 | 6.48406777 | 11.5732852 | 0.00117 | 0.004758 | T | F | T | F |
| Gm3298 | -7.7669593 | 0.34194863 | 11.5661725 | 0.00117 | 0.004773 | F | F | F | F |
| Crlf3 | 0.83868517 | 5.63063603 | 11.5653704 | 0.00117 | 0.004774 | F | F | F | F |
| Zfp251 | -1.0057215 | 4.97600495 | 11.5583727 | 0.00117 | 0.004788 | T | F | T | F |
| Isca1 | 0.78995924 | 5.92200964 | 11.5514408 | 0.00118 | 0.004802 | T | F | T | T |
| Scly | -1.0577338 | 4.72580136 | 11.54811 | 0.00118 | 0.004809 | T | F | F | T |
| B3galt5 | -3.7233556 | 1.18220289 | 11.5464134 | 0.00118 | 0.004811 | F | F | T | F |
| Cxcl1 | 7.72801717 | -0.7125061 | 11.5604881 | 0.00118 | 0.004818 | F | F | F | F |
| Asb13 | -1.3693617 | 3.871898 | 11.5394826 | 0.00118 | 0.004825 | F | F | F | F |
| Ipo9 | -0.7203 | 6.96624651 | 11.5388949 | 0.00118 | 0.004825 | T | F | F | T |
| Krr1 | 0.78370308 | 5.97184266 | 11.5366261 | 0.00119 | 0.004829 | T | F | T | T |
| Znrd2 | -0.9503175 | 5.22726201 | 11.5346731 | 0.00119 | 0.004833 | T | F | F | T |
| Olfr907 | 5.54497273 | -0.27503 | 11.528543 | 0.00119 | 0.004845 | F | F | F | F |
| Trim63 | -7.7582857 | 0.35901532 | 11.5248835 | 0.00119 | 0.004852 | F | F | F | F |
| Prkag2 | -1.4647172 | 3.66576677 | 11.5241233 | 0.00119 | 0.004853 | F | F | T | F |
| L2hgdh | -1.1176932 | 4.498865 | 11.5234477 | 0.00119 | 0.004853 | T | T | T | T |
| Sertad4 | -2.6328414 | 1.93468914 | 11.5233722 | 0.00119 | 0.004853 | F | F | T | F |
| Pex3 | 1.20410719 | 3.93627463 | 11.5227597 | 0.00119 | 0.004853 | T | F | T | T |

| | | | | | | | | | |
|---|---|---|---|---|---|---|---|---|---|
| Traf3 | 0.87048887 | 5.23148154 | 11.518618 | 0.0012 | 0.004862 | T | F | T | T |
| Steap1 | -7.7530612 | 0.34175221 | 11.5173505 | 0.0012 | 0.004863 | F | F | F | F |
| Plekhf1 | -2.2694178 | 2.3672769 | 11.5160158 | 0.0012 | 0.004864 | F | F | F | T |
| Tut7 | 0.75450794 | 6.26084372 | 11.5159141 | 0.0012 | 0.004864 | F | F | F | F |
| Bod1l | 0.63698799 | 7.67247406 | 11.5156923 | 0.0012 | 0.004864 | T | F | T | F |
| Cep85l | 1.01598349 | 4.50962254 | 11.5118843 | 0.0012 | 0.004872 | F | F | T | T |
| Ybx2 | -2.1661966 | 2.51557359 | 11.5090868 | 0.0012 | 0.004877 | T | F | F | F |
| Trpc4ap | 0.66131572 | 7.26710943 | 11.5088151 | 0.0012 | 0.004877 | T | F | F | F |
| Irx3 | 0.65198785 | 7.35332712 | 11.506327 | 0.0012 | 0.004881 | T | F | F | T |
| Rgs16 | 1.68497962 | 2.79114898 | 11.4963662 | 0.00121 | 0.004903 | T | F | F | T |
| Csgalnact2 | 1.52468732 | 3.14367469 | 11.4958805 | 0.00121 | 0.004903 | T | F | F | T |
| Bhlhb9 | -1.0211062 | 4.85840922 | 11.4920496 | 0.00121 | 0.00491 | F | F | F | F |
| Fbxo3 | -0.9849059 | 5.03062117 | 11.4792521 | 0.00122 | 0.004938 | T | F | T | T |
| Rtkn2 | -1.4364514 | 3.73924453 | 11.4790911 | 0.00122 | 0.004938 | F | F | T | F |
| Mdm1 | -1.0520083 | 4.74805322 | 11.4778055 | 0.00122 | 0.00494 | T | F | T | F |
| Osbpl3 | -1.382328 | 3.83486564 | 11.4727258 | 0.00122 | 0.00495 | F | T | T | T |
| Kcnma1 | -4.100943 | 0.98512323 | 11.4702701 | 0.00122 | 0.004955 | T | F | T | T |
| Dst | 0.56515235 | 9.52614779 | 11.466704 | 0.00122 | 0.004962 | T | F | F | F |
| Ilk | 0.72571058 | 6.55565561 | 11.4636444 | 0.00123 | 0.004968 | F | F | F | F |
| Kat7 | 0.6408822 | 7.57848566 | 11.4624807 | 0.00123 | 0.004969 | F | F | T | T |
| Zbtb44 | 0.75335034 | 6.4145931 | 11.4490799 | 0.00123 | 0.004999 | T | F | T | T |
| Pde6c | -7.7518737 | 0.35525146 | 11.4486723 | 0.00123 | 0.004999 | F | F | F | T |
| Exog | -1.4034251 | 3.78880068 | 11.4470182 | 0.00124 | 0.005002 | F | F | T | T |
| Zfp608 | -0.8623225 | 5.73181252 | 11.4443779 | 0.00124 | 0.005007 | T | F | T | F |
| Dvl2 | -0.8154512 | 6.15814074 | 11.4431946 | 0.00124 | 0.005009 | T | F | F | F |
| Mindy3 | 1.31583698 | 3.63739253 | 11.437579 | 0.00124 | 0.00502 | T | F | F | F |
| Psg16 | 0.7931349 | 5.87499562 | 11.4357898 | 0.00124 | 0.005024 | T | T | F | T |
| Pde7a | -0.830793 | 6.00039845 | 11.4331679 | 0.00124 | 0.005029 | T | F | T | T |
| Npcd | 7.71398182 | -0.7241646 | 11.4439856 | 0.00125 | 0.005042 | T | F | T | T |
| Coq2 | 0.8941383 | 5.11824444 | 11.4204891 | 0.00125 | 0.005055 | F | F | F | F |
| Cfap70 | -7.7543145 | 0.32789593 | 11.4202678 | 0.00125 | 0.005055 | F | F | F | F |
| Zdhhc14 | 2.48745859 | 1.42529311 | 11.4140989 | 0.00125 | 0.005069 | T | F | F | F |
| Lrrtm4 | -7.7401353 | 0.33411245 | 11.403644 | 0.00126 | 0.005092 | T | T | T | T |
| Usb1 | 1.00354294 | 4.54346078 | 11.3980101 | 0.00126 | 0.005104 | F | F | F | F |
| Smlr1 | -4.7960602 | 0.77472352 | 11.3888478 | 0.00127 | 0.005124 | F | F | F | F |
| Tlcd1 | -1.3161225 | 3.97659218 | 11.3844384 | 0.00127 | 0.005134 | T | F | F | T |
| Cdkl2 | 1.75467421 | 2.66134264 | 11.383194 | 0.00127 | 0.005136 | F | F | T | F |
| Sohlh2 | 5.20179379 | -0.2224272 | 11.3776039 | 0.00127 | 0.005147 | F | F | T | F |
| Myl6 | 0.64858014 | 7.67676789 | 11.3773477 | 0.00127 | 0.005147 | F | F | F | F |
| Nsd2 | 0.59010793 | 8.66141028 | 11.3757806 | 0.00128 | 0.00515 | T | F | T | F |
| Gng4 | -7.7792875 | 0.39337324 | 11.374597 | 0.00128 | 0.005152 | T | F | T | F |
| Pold3 | -0.798521 | 6.24451674 | 11.3704404 | 0.00128 | 0.005161 | F | F | T | F |

| | | | | | | | | | |
|---|---|---|---|---|---|---|---|---|---|
| Vmn1r65 | 2.51773875 | 1.35242282 | 11.3693913 | 0.00128 | 0.005162 | T | T | F | T |
| Kcnk16 | -7.7374895 | 0.32525059 | 11.3688181 | 0.00128 | 0.005163 | F | F | F | F |
| Hif3a | -1.1990243 | 4.28902329 | 11.3658606 | 0.00128 | 0.005169 | F | F | T | T |
| Slc9a5 | 1.48268222 | 3.21621658 | 11.3646851 | 0.00128 | 0.00517 | T | F | F | T |
| Rnpc3 | 1.12888367 | 4.12221227 | 11.3631158 | 0.00128 | 0.005173 | T | T | T | T |
| Armc7 | 1.80310342 | 2.56283997 | 11.3617672 | 0.00128 | 0.005175 | T | F | T | T |
| Acot3 | -7.734736 | 0.33108144 | 11.3529955 | 0.00129 | 0.005195 | T | F | F | F |
| Rab3gap1 | 0.8184282 | 5.60203855 | 11.348379 | 0.00129 | 0.005204 | F | F | F | F |
| Zfp961 | -1.3722706 | 3.84559317 | 11.3483519 | 0.00129 | 0.005204 | F | F | F | T |
| Rnf224 | -7.7336343 | 0.33028926 | 11.3427986 | 0.0013 | 0.005214 | T | F | T | T |
| Wdr93 | -7.7336343 | 0.33028926 | 11.3427986 | 0.0013 | 0.005214 | F | F | F | F |
| Ccdc149 | -7.7336343 | 0.33028926 | 11.3427986 | 0.0013 | 0.005214 | F | F | T | F |
| Galnt6 | 2.36153208 | 1.54682873 | 11.3401287 | 0.0013 | 0.00522 | T | F | T | T |
| Zfp991 | -0.9355063 | 5.41020608 | 11.3249789 | 0.00131 | 0.005255 | T | F | F | F |
| Lsm3 | -0.855609 | 5.85920799 | 11.3203964 | 0.00131 | 0.005265 | T | F | F | T |
| Slc38a10 | 0.64692172 | 7.37262025 | 11.3173529 | 0.00131 | 0.005271 | T | F | F | T |
| Bnip2 | 0.75743046 | 6.19691044 | 11.3091058 | 0.00132 | 0.00529 | T | F | T | T |
| Dpy19l4 | 0.833892 | 5.59146106 | 11.3042843 | 0.00132 | 0.005301 | F | F | F | T |
| Spats2 | -0.7312799 | 6.79804036 | 11.3034027 | 0.00132 | 0.005302 | T | F | F | T |
| Slc6a13 | -4.0272628 | 0.99619428 | 11.2922952 | 0.00133 | 0.005328 | T | F | F | F |
| Zfp433 | 1.63170985 | 2.92673167 | 11.2894339 | 0.00133 | 0.005334 | F | F | T | F |
| Ercc6l2 | -1.098362 | 4.52366934 | 11.2870158 | 0.00133 | 0.005339 | T | F | F | T |
| Smim10l2a | -1.2623754 | 4.07417758 | 11.2775539 | 0.00133 | 0.005361 | F | F | T | T |
| Snx14 | 0.93773109 | 4.82683692 | 11.2771975 | 0.00133 | 0.005361 | F | F | F | F |
| Smap1 | 0.74520399 | 6.25484683 | 11.2752292 | 0.00134 | 0.005365 | T | F | T | F |
| Nkx2-9 | -7.7639341 | 0.37912278 | 11.2716885 | 0.00134 | 0.005372 | F | F | F | F |
| Bzw2 | -0.6935071 | 7.13304029 | 11.2694566 | 0.00134 | 0.005377 | F | F | F | F |
| Pdcd10 | 0.98665286 | 4.59461691 | 11.2680322 | 0.00134 | 0.005379 | F | F | T | F |
| Larp1b | 1.29075632 | 3.65566005 | 11.2662321 | 0.00134 | 0.005383 | T | F | T | T |
| Antxr2 | 1.25344658 | 3.75982518 | 11.2655297 | 0.00134 | 0.005383 | F | F | F | T |
| Psme3 | 0.60147952 | 8.28228708 | 11.2653037 | 0.00134 | 0.005383 | F | F | F | F |
| Swsap1 | -1.4924479 | 3.57264665 | 11.2591139 | 0.00135 | 0.005397 | F | F | F | T |
| Ift52 | -1.1194418 | 4.42938452 | 11.2564478 | 0.00135 | 0.005403 | F | F | F | T |
| Gsdme | -2.4645052 | 2.12129087 | 11.2501963 | 0.00135 | 0.005417 | F | F | F | F |
| Ska2 | -0.8831601 | 5.52816862 | 11.2498766 | 0.00135 | 0.005417 | F | F | T | T |
| Olfr731 | -3.4221249 | 2.69486652 | 11.2478307 | 0.00135 | 0.005421 | F | F | F | F |
| Ntan1 | 1.07891796 | 4.263013 | 11.2458128 | 0.00135 | 0.005425 | T | F | F | T |
| Timm44 | 0.75203283 | 6.20749931 | 11.2413664 | 0.00136 | 0.005435 | F | F | T | F |
| Cops7a | 0.70902982 | 6.59995535 | 11.2391732 | 0.00136 | 0.00544 | F | F | F | F |
| Ext2 | 0.74178417 | 6.31949434 | 11.236841 | 0.00136 | 0.005444 | T | F | T | T |
| Lrtm2 | -4.7968899 | 0.74564851 | 11.2337763 | 0.00136 | 0.005451 | F | F | T | F |
| Asl | -0.9965859 | 4.97100667 | 11.2287695 | 0.00136 | 0.005462 | F | F | T | F |

| | | | | | | | | | |
|---|---|---|---|---|---|---|---|---|---|
| Ulk4 | -1.4482227 | 3.69643934 | 11.2281793 | 0.00136 | 0.005463 | T | F | T | F |
| Hpcal4 | 2.477022 | 1.39284806 | 11.223465 | 0.00137 | 0.005474 | T | F | T | T |
| Nrcam | -2.262699 | 2.33951807 | 11.2194447 | 0.00137 | 0.005483 | T | F | T | T |
| Lbr | -0.6564848 | 7.73322378 | 11.2013077 | 0.00138 | 0.005527 | T | F | T | T |
| Aph1b | 2.01857062 | 2.40349375 | 11.2001387 | 0.00138 | 0.005528 | T | F | T | T |
| Exoc3l | -3.4820662 | 1.26079155 | 11.200039 | 0.00138 | 0.005528 | F | F | F | F |
| H1f5 | -0.6103121 | 8.95537123 | 11.1945703 | 0.00139 | 0.005541 | F | F | F | F |
| Hgs | 0.7239289 | 6.4446075 | 11.1939245 | 0.00139 | 0.005542 | T | F | F | F |
| Pcolce | -1.7704155 | 3.11256947 | 11.1881077 | 0.00139 | 0.005556 | F | F | F | F |
| Hoxd4 | -8.0189616 | 0.62061176 | 11.1804342 | 0.00139 | 0.005574 | F | F | F | F |
| Enpp5 | -2.7088988 | 1.8153991 | 11.1748114 | 0.0014 | 0.005587 | T | F | F | F |
| Cyhr1 | -0.8174193 | 6.04051784 | 11.1726886 | 0.0014 | 0.005592 | T | F | T | F |
| D130043K2 | -7.7293187 | 0.30298605 | 11.1640296 | 0.00141 | 0.005611 | T | T | T | T |
| Prss50 | -4.784268 | 0.73654684 | 11.1638976 | 0.00141 | 0.005611 | F | F | F | F |
| Fam91a1 | 0.63162782 | 7.56841768 | 11.1635733 | 0.00141 | 0.005611 | T | F | T | T |
| Fam126b | 0.94554343 | 4.86126819 | 11.1633998 | 0.00141 | 0.005611 | F | F | T | F |
| Ubl3 | 0.90068926 | 5.04691117 | 11.1617441 | 0.00141 | 0.005615 | T | T | F | T |
| Cdh4 | -1.2173837 | 4.20813333 | 11.1566874 | 0.00141 | 0.005627 | F | F | T | F |
| Megf10 | -7.7139773 | 0.31880677 | 11.1427644 | 0.00142 | 0.005662 | T | T | T | T |
| Dok5 | -7.7214961 | 0.29987009 | 11.1418691 | 0.00142 | 0.005663 | F | F | F | F |
| Ankle1 | -2.2632346 | 2.37080492 | 11.1363838 | 0.00142 | 0.005676 | F | F | F | T |
| Yipf3 | 0.84325918 | 5.35254437 | 11.1310036 | 0.00143 | 0.005689 | T | T | T | T |
| Slc34a3 | -7.713793 | 0.29891277 | 11.1276906 | 0.00143 | 0.005697 | F | F | F | F |
| Cant1 | 0.75812181 | 6.14870344 | 11.1265682 | 0.00143 | 0.005698 | T | F | T | T |
| Thg1l | -0.9716083 | 5.01315482 | 11.12605 | 0.00143 | 0.005699 | F | F | T | T |
| Hccs | 0.9467778 | 4.75109761 | 11.1235676 | 0.00143 | 0.005704 | T | F | T | T |
| Ddost | 0.61934163 | 7.87319271 | 11.1207605 | 0.00143 | 0.00571 | T | F | F | F |
| Rnaseh2a | -0.9019251 | 5.3572432 | 11.1110203 | 0.00144 | 0.005735 | T | F | T | T |
| Zfp810 | -2.5093248 | 2.01322344 | 11.097404 | 0.00145 | 0.00577 | F | F | F | T |
| Cdyl2 | 1.54412574 | 3.01174435 | 11.0969651 | 0.00145 | 0.00577 | T | F | T | T |
| Cpsf1 | -0.6771281 | 7.3076644 | 11.0938588 | 0.00145 | 0.005777 | F | F | F | F |
| Rraga | -0.846596 | 5.74104151 | 11.0893322 | 0.00145 | 0.005787 | T | T | F | T |
| Hmces | -1.2042872 | 4.19143106 | 11.0893205 | 0.00145 | 0.005787 | T | F | T | T |
| Gabra4 | -2.542275 | 1.96951897 | 11.0852738 | 0.00146 | 0.005796 | T | T | T | T |
| Scn1a | -4.0991885 | 0.97034163 | 11.0638989 | 0.00147 | 0.005852 | T | T | T | T |
| Osgep | -1.2106329 | 4.23260063 | 11.0629953 | 0.00147 | 0.005854 | F | F | T | F |
| Fbxw4 | -1.5781763 | 3.35400255 | 11.061348 | 0.00147 | 0.005857 | F | F | T | F |
| P4ha1 | 0.71376664 | 6.5572097 | 11.0555025 | 0.00148 | 0.005872 | F | T | T | T |
| Tent2 | 0.80588634 | 5.62330378 | 11.0398316 | 0.00149 | 0.005913 | F | F | T | T |
| Mtor | 0.62049512 | 7.72117338 | 11.0350987 | 0.00149 | 0.005925 | T | F | F | T |
| Trim33 | -0.7219526 | 6.85515369 | 11.0343244 | 0.00149 | 0.005926 | T | F | F | T |
| Tctex1d1 | -7.7163913 | 0.29003889 | 11.0310223 | 0.00149 | 0.005934 | T | F | T | F |

| | | | | | | | | | |
|---|---|---|---|---|---|---|---|---|---|
| Eif2b3 | 0.84923433 | 5.26336321 | 11.0139599 | 0.00151 | 0.005979 | T | F | T | F |
| Cox4i2 | -7.6952114 | 0.29294926 | 11.0070277 | 0.00151 | 0.005997 | F | F | F | F |
| Slc17a9 | 2.35259536 | 1.53943574 | 11.0001545 | 0.00151 | 0.006015 | T | F | F | F |
| Fam90a1b | 7.66122697 | -0.7677622 | 11.015806 | 0.00151 | 0.006015 | F | F | T | F |
| Gm10719 | 5.79423859 | -0.4371879 | 10.9979716 | 0.00152 | 0.006019 | F | F | F | F |
| Ccdc136 | -1.0518818 | 4.64619622 | 10.993561 | 0.00152 | 0.00603 | F | F | F | F |
| 9030025P2( | 3.27238899 | 0.62749997 | 10.9867552 | 0.00152 | 0.006048 | F | F | F | F |
| Dnajc14 | 0.75101158 | 6.12477474 | 10.9857574 | 0.00152 | 0.006049 | T | T | F | T |
| Slc35c1 | 0.94107455 | 4.85683837 | 10.98167 | 0.00153 | 0.006059 | T | F | T | T |
| Prim2 | 0.81076803 | 5.54845955 | 10.9768122 | 0.00153 | 0.006072 | T | F | F | F |
| Thbs1 | -1.5221059 | 3.45255399 | 10.967487 | 0.00154 | 0.006097 | T | F | F | F |
| Rnase4 | -1.8623068 | 2.86747568 | 10.9636015 | 0.00154 | 0.006107 | F | F | F | F |
| Hnrnph1 | -0.6229598 | 8.28497452 | 10.9573277 | 0.00154 | 0.006123 | T | T | T | T |
| Prokr2 | -7.6920352 | 0.29891001 | 10.9566193 | 0.00155 | 0.006124 | T | F | F | T |
| Bsdc1 | 0.85689661 | 5.22078189 | 10.9536335 | 0.00155 | 0.006131 | F | F | T | F |
| Ap5b1 | 1.42169881 | 3.31351993 | 10.9528228 | 0.00155 | 0.006132 | F | F | F | T |
| Plekhb1 | -7.6912971 | 0.29824322 | 10.9503797 | 0.00155 | 0.006138 | F | F | F | F |
| Rhoq | 0.9894478 | 4.61327198 | 10.9455764 | 0.00155 | 0.006151 | T | F | T | T |
| Tvp23a | -2.2601985 | 2.28287609 | 10.9449651 | 0.00155 | 0.006151 | F | F | T | F |
| Rnf24 | -1.8508944 | 2.89551117 | 10.9446775 | 0.00155 | 0.006151 | T | T | F | T |
| Atad2 | -0.6472447 | 7.90456477 | 10.9431552 | 0.00155 | 0.006154 | T | F | F | T |
| Idh2 | -0.7530707 | 6.51004658 | 10.9396449 | 0.00156 | 0.006163 | F | F | T | T |
| Pptc7 | 0.80321043 | 5.60046262 | 10.9375575 | 0.00156 | 0.006168 | T | F | T | F |
| Gm11627 | -4.7517212 | 0.71233917 | 10.9365136 | 0.00156 | 0.006169 | F | F | F | F |
| Fam102b | -1.0984474 | 4.4491565 | 10.935702 | 0.00156 | 0.006171 | F | F | T | F |
| Tnnt3 | -7.6959706 | 0.30886286 | 10.9294872 | 0.00156 | 0.006187 | F | F | F | F |
| Slc45a3 | -1.7799638 | 3.01775671 | 10.9270443 | 0.00157 | 0.006193 | T | F | T | T |
| Cdyl | 0.72946457 | 6.31948341 | 10.9257538 | 0.00157 | 0.006195 | F | F | F | T |
| Tmc1 | 5.79671401 | -0.4487023 | 10.9144276 | 0.00158 | 0.006227 | T | F | T | T |
| Dynlrb2 | -7.6857314 | 0.27973778 | 10.9138152 | 0.00158 | 0.006227 | F | F | F | F |
| Ebf4 | -7.68606 | 0.29284366 | 10.9099416 | 0.00158 | 0.006237 | T | F | F | T |
| Klk1b22 | 4.27365277 | 0.07628305 | 10.9078541 | 0.00158 | 0.006242 | F | F | T | F |
| Atp7a | 0.8270878 | 5.39257571 | 10.9042421 | 0.00158 | 0.006251 | F | F | T | F |
| Cmtm6 | 0.73647132 | 6.21643132 | 10.89318 | 0.00159 | 0.006282 | T | F | T | T |
| Zfp945 | -1.9171085 | 2.86871468 | 10.877961 | 0.0016 | 0.006325 | T | F | T | T |
| Csk | 0.64622029 | 7.13063595 | 10.8737827 | 0.00161 | 0.006336 | F | F | F | F |
| Coq10a | -3.0462567 | 1.47668195 | 10.8652848 | 0.00161 | 0.00636 | F | F | T | F |
| Tmcc1 | 0.90201613 | 4.95506974 | 10.8616103 | 0.00161 | 0.006368 | T | F | T | F |
| Snapc1 | 1.0302068 | 4.35663907 | 10.8615774 | 0.00161 | 0.006368 | F | F | T | T |
| Rnf4 | 0.6225516 | 7.57711868 | 10.8590023 | 0.00162 | 0.006375 | T | F | T | F |
| Cln8 | 0.83350676 | 5.32316803 | 10.8562513 | 0.00162 | 0.006381 | T | F | T | T |
| Lman2l | 0.96438262 | 4.61705184 | 10.8478521 | 0.00162 | 0.006405 | T | F | F | F |

| | | | | | | | | | |
|---|---|---|---|---|---|---|---|---|---|
| Traf3ip2 | 1.20793901 | 3.82255519 | 10.8446312 | 0.00163 | 0.006413 | T | F | F | T |
| Gm3468 | -7.6772441 | 0.2730841 | 10.8434573 | 0.00163 | 0.006416 | F | F | F | F |
| Fbxl4 | 1.50332317 | 3.0626574 | 10.8306716 | 0.00164 | 0.006452 | T | F | F | F |
| Prkag1 | 0.82788626 | 5.3834268 | 10.8295598 | 0.00164 | 0.006454 | F | F | F | F |
| Abhd12b | -7.6746254 | 0.27727012 | 10.8287793 | 0.00164 | 0.006456 | F | F | F | F |
| Rest | 0.60366526 | 8.1891642 | 10.8258831 | 0.00164 | 0.006463 | F | F | T | F |
| Txndc9 | 0.76230486 | 5.93817568 | 10.8249285 | 0.00164 | 0.006465 | T | F | T | T |
| Zfp445 | -0.6911124 | 7.04922342 | 10.8206475 | 0.00164 | 0.006476 | T | F | T | T |
| Ankmy2 | -1.0965967 | 4.43003052 | 10.8180783 | 0.00165 | 0.006483 | T | F | F | F |
| Fkbp2 | 0.89372429 | 4.97134759 | 10.8108977 | 0.00165 | 0.006503 | F | F | T | F |
| Mgst3 | 1.21755784 | 3.86418898 | 10.7925421 | 0.00167 | 0.006557 | T | F | T | T |
| Tor2a | 0.96612138 | 4.6260824 | 10.7859779 | 0.00167 | 0.006576 | F | F | T | F |
| Vti1a | 0.90117835 | 4.92673167 | 10.7849566 | 0.00167 | 0.006578 | T | F | T | T |
| Fam98b | -0.8154652 | 5.91255404 | 10.7830232 | 0.00167 | 0.006582 | F | F | F | F |
| Tmem179b | 1.49649536 | 3.06798925 | 10.7811602 | 0.00167 | 0.006586 | F | F | F | F |
| Hilpda | 1.40258783 | 3.29057631 | 10.7809707 | 0.00168 | 0.006586 | F | F | F | T |
| Ppp1r10 | -0.6863884 | 7.07933486 | 10.780392 | 0.00168 | 0.006587 | F | F | F | F |
| Arfgef3 | -1.9096643 | 2.98224852 | 10.7791224 | 0.00168 | 0.006589 | T | F | F | T |
| Onecut2 | -2.3682766 | 2.12325381 | 10.7745853 | 0.00168 | 0.006602 | T | F | T | T |
| Ctnnbl1 | -0.8219182 | 5.88947705 | 10.7728052 | 0.00168 | 0.006606 | F | F | F | F |
| Aurka | -0.7057175 | 6.86271796 | 10.7654508 | 0.00169 | 0.006627 | T | F | F | F |
| Anapc4 | -1.0458664 | 4.61374943 | 10.7623269 | 0.00169 | 0.006636 | F | F | F | F |
| Jmjd7 | -1.6478779 | 3.18754868 | 10.7614951 | 0.00169 | 0.006637 | F | F | F | F |
| Aven | -1.0074791 | 4.76442319 | 10.7475949 | 0.0017 | 0.006678 | F | F | F | F |
| Exd2 | 1.05113072 | 4.24370784 | 10.7444232 | 0.0017 | 0.006687 | F | F | T | F |
| Ifrd1 | 0.84270856 | 5.22370199 | 10.7432003 | 0.0017 | 0.00669 | F | F | F | F |
| Larp7 | -0.7521223 | 6.46778484 | 10.7354306 | 0.00171 | 0.006712 | F | F | F | F |
| Tspo | 1.28339488 | 3.6148778 | 10.7315502 | 0.00171 | 0.006723 | T | F | F | T |
| Cblc | -7.6701464 | 0.25790428 | 10.73036 | 0.00171 | 0.006725 | T | F | F | T |
| Ccdc167 | 1.13472472 | 4.00791037 | 10.7300181 | 0.00171 | 0.006725 | F | F | T | F |
| Sdf2 | 0.99196973 | 4.50750351 | 10.7274188 | 0.00172 | 0.006732 | T | F | F | F |
| Manba | 1.01746832 | 4.35619173 | 10.7269924 | 0.00172 | 0.006732 | T | F | F | F |
| Phykpl | 1.8706963 | 2.30019776 | 10.7232545 | 0.00172 | 0.006743 | T | F | F | T |
| Rax | -7.7985853 | 0.33102588 | 10.7218804 | 0.00172 | 0.006746 | F | F | F | F |
| Gm20521 | 3.08646658 | 0.93238571 | 10.7164262 | 0.00173 | 0.006761 | F | F | F | F |
| Ctnnbip1 | 0.77516811 | 5.77568767 | 10.7161443 | 0.00173 | 0.006761 | T | F | F | F |
| Nacc2 | -1.9573328 | 2.68115708 | 10.7083103 | 0.00173 | 0.006785 | T | F | T | T |
| Atp8a1 | -1.3362547 | 3.8296255 | 10.7028512 | 0.00174 | 0.0068 | T | F | F | T |
| Mfsd4b5 | 1.58608976 | 2.89912832 | 10.6962825 | 0.00174 | 0.006819 | F | F | T | F |
| Tcea3 | -1.7162612 | 3.06144705 | 10.6961481 | 0.00174 | 0.006819 | F | F | F | F |
| Cisd3 | 1.48954913 | 3.09215309 | 10.6949356 | 0.00174 | 0.006822 | F | F | F | F |
| L1cam | -1.795294 | 3.03165172 | 10.6936699 | 0.00174 | 0.006824 | F | F | F | F |

| | | | | | | | | | |
|---|---|---|---|---|---|---|---|---|---|
| Ppil1 | -0.8177797 | 5.90051166 | 10.6905847 | 0.00175 | 0.006833 | F | F | F | F |
| Abcb10 | 0.83491364 | 5.27890048 | 10.6887939 | 0.00175 | 0.006837 | T | F | T | T |
| Marcksl1 | -0.5811123 | 9.39067825 | 10.6850871 | 0.00175 | 0.006848 | F | F | F | F |
| Vmn1r173 | -7.7176136 | 0.27326262 | 10.6822535 | 0.00175 | 0.006855 | F | F | F | F |
| Ccbe1 | -4.0355577 | 0.94671452 | 10.6789737 | 0.00176 | 0.006865 | T | F | T | T |
| Gm8159 | 7.61982747 | -0.8017248 | 10.6903582 | 0.00176 | 0.006876 | F | F | F | F |
| Atg7 | 0.99679312 | 4.42887315 | 10.6730286 | 0.00176 | 0.006881 | T | F | T | T |
| Parl | -0.9072849 | 5.2409294 | 10.6646579 | 0.00177 | 0.006906 | F | F | T | F |
| Bckdha | -0.9476295 | 5.03332128 | 10.6643495 | 0.00177 | 0.006906 | F | F | T | F |
| Gspt2 | -1.3916267 | 3.6904744 | 10.661079 | 0.00177 | 0.006915 | T | F | F | T |
| Ttc32 | -2.3120596 | 2.16489814 | 10.6603771 | 0.00177 | 0.006916 | T | F | T | T |
| Snx24 | -1.6099141 | 3.23694202 | 10.6524107 | 0.00178 | 0.006941 | F | F | T | T |
| Frat2 | -1.3441588 | 3.79910862 | 10.6479548 | 0.00178 | 0.006954 | T | F | T | T |
| Rnf5 | -0.9841226 | 4.84898902 | 10.6470369 | 0.00178 | 0.006955 | F | F | F | F |
| 2510002D2 | 1.2522689 | 3.6632546 | 10.6443863 | 0.00178 | 0.006963 | T | F | T | F |
| Ntng2 | -7.6602439 | 0.27641759 | 10.6419233 | 0.00179 | 0.006969 | F | F | F | F |
| Clec16a | 0.82138981 | 5.49070658 | 10.641572 | 0.00179 | 0.006969 | T | F | T | T |
| Adgrb3 | -7.6533517 | 0.26252503 | 10.6390011 | 0.00179 | 0.006976 | F | F | T | F |
| Rerg | -2.1440239 | 2.39651584 | 10.638364 | 0.00179 | 0.006977 | T | F | T | F |
| Acsl6 | -2.0373253 | 2.56959883 | 10.6328736 | 0.00179 | 0.006993 | T | F | F | T |
| Mfn1 | 0.7213929 | 6.28372626 | 10.631193 | 0.00179 | 0.006998 | F | F | T | F |
| Ckap4 | -0.6761286 | 7.23362196 | 10.6275356 | 0.0018 | 0.007008 | T | F | F | F |
| Zfp69 | -2.2727605 | 2.21370617 | 10.6270506 | 0.0018 | 0.007008 | F | F | F | T |
| Mettl17 | -1.4275663 | 3.6042801 | 10.6265304 | 0.0018 | 0.007009 | F | F | F | F |
| Ddb1 | 0.51185605 | 10.3435455 | 10.6262128 | 0.0018 | 0.007009 | F | F | F | F |
| Zfp942 | 1.2718398 | 3.61402354 | 10.620332 | 0.0018 | 0.007026 | F | F | T | T |
| Tmem63b | -0.7864896 | 6.0949616 | 10.6071989 | 0.00181 | 0.007068 | F | F | T | F |
| Spen | 0.58105037 | 8.35224669 | 10.6006459 | 0.00182 | 0.007088 | T | F | F | F |
| Cysltr1 | -7.65584 | 0.24377141 | 10.5959146 | 0.00182 | 0.007102 | T | F | F | T |
| Cbx7 | 0.76829491 | 6.15571601 | 10.593824 | 0.00183 | 0.007108 | T | F | F | F |
| Tbc1d20 | 0.76240267 | 5.86804275 | 10.5896638 | 0.00183 | 0.00712 | F | F | F | F |
| Stk31 | 4.36190329 | -0.0416298 | 10.588738 | 0.00183 | 0.007122 | F | F | F | F |
| Noc2l | 0.59845427 | 8.16156531 | 10.5841919 | 0.00183 | 0.007136 | F | F | F | F |
| Bcl3 | 3.07017711 | 0.73531158 | 10.5837271 | 0.00183 | 0.007136 | F | F | T | F |
| Tmem181a | 0.85889701 | 5.13448477 | 10.5817913 | 0.00184 | 0.007141 | F | F | T | F |
| 9130023H2 | -2.1701321 | 2.35243047 | 10.5805473 | 0.00184 | 0.007144 | F | F | F | F |
| Dennd4a | 1.15871219 | 3.93611867 | 10.5771835 | 0.00184 | 0.007154 | T | T | F | T |
| Sdha | 0.59660807 | 8.00182848 | 10.5768772 | 0.00184 | 0.007154 | F | F | F | F |
| Hinfp | 0.80677747 | 5.46917626 | 10.5695147 | 0.00185 | 0.007177 | F | F | T | T |
| Ttll9 | -7.6507875 | 0.23932658 | 10.5534556 | 0.00186 | 0.007229 | F | F | F | F |
| Akap5 | -1.5908068 | 3.28605274 | 10.5502829 | 0.00186 | 0.007238 | F | F | T | T |
| Ppp1r18 | 0.7997466 | 5.51695035 | 10.5490591 | 0.00186 | 0.007241 | F | F | F | T |

| | | | | | | | | | |
|---|---|---|---|---|---|---|---|---|---|
| Mroh1 | 0.85027679 | 5.14214988 | 10.5485286 | 0.00186 | 0.007242 | F | F | F | F |
| Zfp131 | 0.72493409 | 6.26977032 | 10.5470359 | 0.00187 | 0.007245 | T | F | F | F |
| Kcnq1 | 3.21857859 | 0.59935791 | 10.5431286 | 0.00187 | 0.007257 | T | F | F | T |
| Pkn3 | -2.0528679 | 2.54944842 | 10.5389335 | 0.00187 | 0.00727 | T | F | F | F |
| Samd8 | 0.74343024 | 6.06013372 | 10.5334738 | 0.00188 | 0.007287 | T | T | T | T |
| Pdgfc | -2.6406745 | 1.77376933 | 10.5321301 | 0.00188 | 0.00729 | T | F | T | F |
| Dyrk3 | -1.2817374 | 3.92151529 | 10.5291693 | 0.00188 | 0.007299 | F | F | F | T |
| Srsf11 | -0.6665953 | 7.17094168 | 10.5231477 | 0.00189 | 0.007317 | T | F | T | T |
| Apoc3 | -7.638504 | 0.24339592 | 10.5228414 | 0.00189 | 0.007317 | F | F | F | T |
| Dennd5a | 0.69724428 | 6.50505616 | 10.5226059 | 0.00189 | 0.007317 | F | F | F | T |
| Sncb | -7.6578169 | 0.23734055 | 10.5215 | 0.00189 | 0.00732 | F | F | T | F |
| Gvin1 | 5.34770769 | -0.3821665 | 10.5107407 | 0.0019 | 0.007355 | T | T | T | T |
| Mad2l1bp | 1.05212566 | 4.20789771 | 10.5071475 | 0.0019 | 0.007366 | F | F | T | F |
| Rhobtb1 | -1.8529126 | 2.80786734 | 10.5012738 | 0.00191 | 0.007384 | F | F | T | F |
| Nrg1 | -2.566944 | 1.86532728 | 10.4936149 | 0.00191 | 0.007409 | T | F | T | T |
| Zfp229 | 0.83055632 | 5.27104824 | 10.4851233 | 0.00192 | 0.007437 | T | F | T | F |
| Zcchc7 | 0.88464897 | 4.94033198 | 10.4832817 | 0.00192 | 0.007442 | T | F | T | F |
| Ankrd50 | 0.64569509 | 7.28398589 | 10.4825617 | 0.00192 | 0.007443 | F | T | F | T |
| Rab36 | -2.6715083 | 1.74638391 | 10.4771078 | 0.00193 | 0.007461 | F | F | F | T |
| Tgif2 | 0.69319499 | 6.52179438 | 10.4717971 | 0.00193 | 0.007478 | T | F | T | F |
| Tk1 | 0.80605128 | 5.48860511 | 10.4650476 | 0.00194 | 0.007499 | F | F | F | F |
| Arntl | 1.25393461 | 3.6285978 | 10.4647668 | 0.00194 | 0.007499 | F | F | F | F |
| Mcfd2 | 0.77571674 | 5.80137755 | 10.4647246 | 0.00194 | 0.007499 | F | F | T | F |
| Mgrn1 | -0.7893353 | 6.02370573 | 10.460596 | 0.00194 | 0.007512 | T | F | T | F |
| Pdap1 | -0.6564721 | 7.31241562 | 10.4565749 | 0.00195 | 0.007522 | F | F | T | F |
| BC053393 | 7.59119396 | -0.8250835 | 10.4705935 | 0.00195 | 0.007522 | F | F | T | F |
| Tecta | 7.59119396 | -0.8250835 | 10.4705935 | 0.00195 | 0.007522 | F | F | F | F |
| Osm | 7.59119396 | -0.8250835 | 10.4705935 | 0.00195 | 0.007522 | F | F | T | F |
| Crym | -4.0196083 | 0.90356502 | 10.4486314 | 0.00195 | 0.007547 | T | F | F | F |
| Dph7 | -1.5369229 | 3.34097054 | 10.4428494 | 0.00196 | 0.007565 | F | F | F | T |
| Thap3 | 1.26163335 | 3.60431707 | 10.4425426 | 0.00196 | 0.007565 | F | F | F | F |
| Wdsub1 | -2.1057322 | 2.4188478 | 10.4380795 | 0.00196 | 0.00758 | F | F | T | F |
| Kif23 | -0.7159205 | 6.68621006 | 10.4362515 | 0.00196 | 0.007585 | F | F | F | F |
| Armc3 | -7.6361147 | 0.2268093 | 10.4342523 | 0.00197 | 0.00759 | F | F | F | T |
| Hhat | -3.9978538 | 0.89527074 | 10.4315712 | 0.00197 | 0.007598 | T | F | F | T |
| Camk2d | 1.29567227 | 3.50103193 | 10.4276202 | 0.00197 | 0.007611 | T | F | T | T |
| Fbxw17 | -2.2242844 | 2.24633903 | 10.4196452 | 0.00198 | 0.007638 | F | F | F | F |
| Trim26 | 0.80125939 | 5.45375862 | 10.4175421 | 0.00198 | 0.007644 | T | F | T | T |
| Ganc | 2.25054981 | 1.5613763 | 10.4141752 | 0.00198 | 0.007655 | T | F | F | T |
| Kcnab1 | -7.6290836 | 0.2253719 | 10.4104117 | 0.00199 | 0.007667 | T | T | T | T |
| Ints14 | 0.75000344 | 5.91510046 | 10.4026286 | 0.002 | 0.007693 | F | F | F | F |
| Ccdc13 | -7.6265646 | 0.22527732 | 10.4011581 | 0.002 | 0.007697 | F | F | F | F |

| | | | | | | | | | |
|---|---|---|---|---|---|---|---|---|---|
| Xpot | 0.60072792 | 7.75375226 | 10.3980743 | 0.002 | 0.007706 | T | F | T | F |
| Thop1 | -0.6474855 | 7.45024325 | 10.3950723 | 0.002 | 0.007716 | T | F | F | T |
| Raet1e | -2.1070003 | 2.41627767 | 10.3941229 | 0.002 | 0.007718 | F | F | F | F |
| Uqcrc1 | 0.60735292 | 7.72617024 | 10.3807648 | 0.00202 | 0.007764 | F | F | F | F |
| Stard7 | 0.61704724 | 7.40758599 | 10.3748804 | 0.00202 | 0.007784 | T | F | F | F |
| Hspa5 | 0.53093428 | 9.93566862 | 10.3745151 | 0.00202 | 0.007784 | F | F | F | T |
| Ikzf2 | -2.2568817 | 2.20422015 | 10.372669 | 0.00202 | 0.00779 | F | F | T | T |
| Sestd1 | -0.8501689 | 5.50075341 | 10.3701161 | 0.00203 | 0.007797 | T | T | T | T |
| Zfp68 | -1.232963 | 4.00747437 | 10.3665771 | 0.00203 | 0.007809 | T | T | T | T |
| Mki67 | 0.52270108 | 9.85783817 | 10.3654149 | 0.00203 | 0.007812 | F | F | F | F |
| Rasa2 | 0.88278593 | 4.99451742 | 10.36198 | 0.00203 | 0.007823 | T | F | T | T |
| Wac | 0.64311072 | 6.98494301 | 10.3553003 | 0.00204 | 0.007846 | T | F | F | T |
| Iqsec1 | 0.64475925 | 6.96836169 | 10.352591 | 0.00204 | 0.007854 | F | F | T | T |
| Sh3gl2 | -2.1234214 | 2.37339666 | 10.3473683 | 0.00205 | 0.007872 | T | F | T | T |
| Retreg3 | 0.74156008 | 5.97905259 | 10.3435364 | 0.00205 | 0.007884 | F | F | T | F |
| Vwa8 | 0.85861269 | 5.06395595 | 10.3425532 | 0.00205 | 0.007887 | F | F | T | F |
| Tmem82 | -7.8867292 | 0.50355375 | 10.3402983 | 0.00205 | 0.007893 | T | F | F | F |
| Flvcr1 | 0.8433591 | 5.14554386 | 10.3401767 | 0.00205 | 0.007893 | T | T | T | T |
| Lrrc39 | 2.3523382 | 1.41774327 | 10.337777 | 0.00206 | 0.0079 | T | T | T | T |
| Rnf111 | 0.66599442 | 6.75261781 | 10.3355914 | 0.00206 | 0.007907 | T | F | T | T |
| Hoxc4 | -7.7927688 | 0.41906478 | 10.33519 | 0.00206 | 0.007907 | T | F | F | F |
| Ndufa2 | -0.9632297 | 4.92487345 | 10.3290069 | 0.00206 | 0.007928 | F | F | F | F |
| Slc29a1 | 0.80973657 | 5.38043693 | 10.3246429 | 0.00207 | 0.007943 | T | F | F | F |
| Zfp87 | -1.2547834 | 3.98735908 | 10.3161744 | 0.00208 | 0.007973 | T | F | T | F |
| Sertad2 | 0.76182144 | 5.7916448 | 10.3087601 | 0.00208 | 0.007999 | T | F | T | T |
| Olfm2 | -7.6150688 | 0.21469953 | 10.3034937 | 0.00209 | 0.008016 | F | F | F | F |
| Nim1k | -7.6173565 | 0.23740195 | 10.3030725 | 0.00209 | 0.008016 | T | F | F | T |
| Rnf14 | 0.76163089 | 5.7614558 | 10.3029813 | 0.00209 | 0.008016 | T | F | T | T |
| Ggnbp2 | 0.72174529 | 6.16285209 | 10.2962983 | 0.0021 | 0.00804 | T | F | F | T |
| Dyrk1b | -1.05186 | 4.51141179 | 10.2958059 | 0.0021 | 0.00804 | F | F | T | T |
| Coq4 | -1.3408948 | 3.74623497 | 10.293252 | 0.0021 | 0.008048 | F | F | T | T |
| Nebl | -3.5951018 | 1.05923114 | 10.2862758 | 0.00211 | 0.008073 | F | F | T | T |
| H13 | 0.69603669 | 6.46295437 | 10.2813827 | 0.00211 | 0.00809 | F | F | F | F |
| Rnh1 | 0.82726732 | 5.24154695 | 10.2759078 | 0.00212 | 0.008109 | F | F | F | F |
| Slc35a1 | 0.82863814 | 5.19883055 | 10.2632194 | 0.00213 | 0.008156 | F | F | T | F |
| Naa30 | 0.74965856 | 5.90772509 | 10.2604992 | 0.00213 | 0.008164 | T | T | T | T |
| Dtx3l | -1.0051173 | 4.68620052 | 10.2599199 | 0.00213 | 0.008165 | T | F | T | F |
| Nup188 | -0.6097112 | 8.06414549 | 10.2594824 | 0.00213 | 0.008165 | F | F | T | F |
| Fahd1 | -2.4156148 | 1.96921287 | 10.2591401 | 0.00213 | 0.008165 | T | F | T | T |
| Gm14444 | 4.62764135 | -0.1531025 | 10.2511953 | 0.00214 | 0.008194 | F | F | F | F |
| Olfr1423 | -1.4651405 | 3.48579255 | 10.2509482 | 0.00214 | 0.008194 | F | F | F | F |
| Pax3 | -4.5901689 | 0.70619509 | 10.2461455 | 0.00215 | 0.008211 | T | F | T | T |

| | | | | | | | | | |
|---|---|---|---|---|---|---|---|---|---|
| Phkb | -0.9233832 | 5.0427705 | 10.2416958 | 0.00215 | 0.008226 | T | F | F | T |
| Ccdc158 | -7.6348729 | 0.2107834 | 10.2402917 | 0.00215 | 0.00823 | T | F | T | T |
| Spag5 | -0.7359199 | 6.42876471 | 10.236453 | 0.00216 | 0.008243 | T | F | F | F |
| Bach1 | -0.8085225 | 5.85117404 | 10.2316843 | 0.00216 | 0.00826 | F | F | T | T |
| Tmem59l | -1.8492377 | 2.77217198 | 10.229493 | 0.00216 | 0.008267 | T | F | F | T |
| Samd11 | 3.59498405 | 0.28721123 | 10.2255539 | 0.00217 | 0.008281 | F | F | F | F |
| Gtsf1l | 3.02553395 | 0.69096042 | 10.2113595 | 0.00218 | 0.008334 | F | F | T | F |
| Tomm7 | -0.8480493 | 5.51454841 | 10.2110431 | 0.00218 | 0.008334 | T | F | T | F |
| Mnx1 | -7.663299 | 0.29717718 | 10.2099292 | 0.00218 | 0.008337 | T | F | F | F |
| Ppp2r3d | 0.81279933 | 5.30151704 | 10.2016276 | 0.00219 | 0.008368 | F | F | T | F |
| Tma7 | -0.7573848 | 6.2797525 | 10.1998799 | 0.00219 | 0.008373 | F | F | F | F |
| Plin3 | -0.714264 | 6.60489556 | 10.1929503 | 0.0022 | 0.008399 | T | T | F | T |
| Dnm1l | 0.64107978 | 7.02281904 | 10.1819046 | 0.00221 | 0.008441 | T | F | F | T |
| Pomgnt1 | 0.74722987 | 5.86136531 | 10.177115 | 0.00222 | 0.008458 | F | F | F | F |
| Lins1 | -1.7501321 | 2.92407993 | 10.1738962 | 0.00222 | 0.008469 | F | F | F | F |
| Gm14305 | 2.37939988 | 2.03431548 | 10.1664115 | 0.00223 | 0.008497 | T | F | F | T |
| Tsc22d2 | -0.7266264 | 6.48476437 | 10.1646163 | 0.00223 | 0.008503 | T | F | T | F |
| Kctd2 | -1.1588106 | 4.15067372 | 10.1596484 | 0.00223 | 0.008521 | F | F | F | T |
| Pex1 | 0.85972097 | 5.00200322 | 10.1581685 | 0.00224 | 0.008524 | T | F | F | T |
| Adcy2 | -3.5825703 | 1.04645714 | 10.1580484 | 0.00224 | 0.008524 | F | F | F | F |
| Gm17018 | -1.6287963 | 3.1409388 | 10.1569003 | 0.00224 | 0.008528 | F | F | F | F |
| 4930590J08 | 3.63805101 | 0.25668294 | 10.1564248 | 0.00224 | 0.008528 | F | F | F | F |
| Pdss1 | 1.01154739 | 4.27330943 | 10.1549792 | 0.00224 | 0.008532 | T | F | F | F |
| Fam162a | -0.9105366 | 5.07846272 | 10.1542487 | 0.00224 | 0.008534 | F | F | F | F |
| Tmc3 | -3.5868321 | 1.04409175 | 10.1518765 | 0.00224 | 0.008542 | T | F | T | T |
| Gm14419 | 3.03197752 | 0.89856773 | 10.1512198 | 0.00224 | 0.008543 | F | F | F | T |
| Zfp148 | 0.733251 | 6.00886564 | 10.1465241 | 0.00225 | 0.00856 | T | T | T | T |
| Lrriq1 | -7.597876 | 0.19637308 | 10.1434327 | 0.00225 | 0.008571 | F | F | F | F |
| Ccdc187 | -2.1528831 | 2.38035864 | 10.1402319 | 0.00225 | 0.008582 | F | F | F | F |
| Sowahd | -4.7394708 | 0.67823626 | 10.130559 | 0.00226 | 0.008619 | T | F | T | F |
| Nupr1 | 7.54726006 | -0.8607122 | 10.1417063 | 0.00227 | 0.008629 | F | F | F | F |
| Bak1 | 1.21388141 | 3.71322882 | 10.1223453 | 0.00227 | 0.008649 | T | F | F | T |
| Obsl1 | 0.66372203 | 6.71436936 | 10.1203731 | 0.00228 | 0.008656 | F | F | F | F |
| Bcl6b | -2.9770519 | 1.42240878 | 10.1182802 | 0.00228 | 0.008663 | T | F | F | T |
| Wars2 | 1.05259061 | 4.15425599 | 10.1177818 | 0.00228 | 0.008663 | T | F | T | T |
| Tirap | 0.83506019 | 5.14516824 | 10.113734 | 0.00228 | 0.008678 | F | F | T | T |
| Apol8 | 2.8043126 | 0.90638482 | 10.1130169 | 0.00228 | 0.00868 | F | F | T | F |
| Pax7 | -7.5896543 | 0.20867827 | 10.1071031 | 0.00229 | 0.008702 | T | F | T | T |
| Pigh | -1.1969251 | 4.06144798 | 10.1065641 | 0.00229 | 0.008703 | F | F | T | F |
| Fig4 | 0.86654794 | 4.95016409 | 10.1025845 | 0.00229 | 0.008717 | F | F | F | F |
| Gemin4 | -0.6811237 | 6.87675373 | 10.0999149 | 0.0023 | 0.008727 | F | F | F | F |
| Sod1 | -0.6560689 | 7.37390783 | 10.0952233 | 0.0023 | 0.008744 | F | F | F | T |

| | | | | | | | | | |
|---|---|---|---|---|---|---|---|---|---|
| Xpnpep3 | 0.7756248 | 5.6865428 | 10.0912203 | 0.00231 | 0.008758 | T | F | F | T |
| Ino80c | 0.80661142 | 5.32082895 | 10.0910158 | 0.00231 | 0.008758 | F | F | T | F |
| Deaf1 | -0.9616475 | 4.8297218 | 10.0905335 | 0.00231 | 0.008759 | F | F | T | F |
| Hspd1 | 0.49870889 | 10.3405891 | 10.0872061 | 0.00231 | 0.00877 | T | F | F | F |
| Dppa4 | 3.43248189 | 0.41609656 | 10.0869508 | 0.00231 | 0.00877 | T | F | F | T |
| Sec22b | 0.82984074 | 5.28201918 | 10.0853944 | 0.00231 | 0.008775 | F | F | T | F |
| Ppp1r12a | 0.6134442 | 7.30388694 | 10.0787269 | 0.00232 | 0.008801 | F | F | T | F |
| Brd1 | 0.62043534 | 7.18407717 | 10.0772905 | 0.00232 | 0.008805 | F | F | T | T |
| Pbdc1 | -0.7717689 | 6.04894112 | 10.0723051 | 0.00233 | 0.008824 | T | F | F | F |
| Ibtk | -0.7455664 | 6.3581125 | 10.0697852 | 0.00233 | 0.008833 | T | F | F | T |
| Bud13 | -0.9510901 | 4.94323634 | 10.0655651 | 0.00233 | 0.008849 | F | F | F | F |
| Agfg2 | 1.21084939 | 3.68389081 | 10.0641338 | 0.00234 | 0.008854 | F | F | F | F |
| Dnah6 | -2.9749755 | 1.41630197 | 10.059215 | 0.00234 | 0.008872 | F | F | F | T |
| Stxbp1 | -1.5021868 | 3.34577172 | 10.0498336 | 0.00235 | 0.00891 | T | F | F | T |
| Ubac2 | 0.74799292 | 5.80449348 | 10.0425671 | 0.00236 | 0.008939 | T | F | T | F |
| Rtf1 | -0.6970699 | 6.73310348 | 10.0377255 | 0.00236 | 0.008957 | F | F | F | F |
| Gltpd2 | -7.5793889 | 0.19447542 | 10.0373979 | 0.00237 | 0.008957 | F | F | F | T |
| Olfr374 | 0.90488743 | 4.92297084 | 10.0346966 | 0.00237 | 0.008967 | F | F | F | F |
| Vhl | -1.2784166 | 3.84846717 | 10.0327502 | 0.00237 | 0.008973 | T | F | T | F |
| Cdc25c | -1.0628328 | 4.41460137 | 10.0315799 | 0.00237 | 0.008977 | F | F | F | F |
| Gtf2h3 | -0.8480164 | 5.43571527 | 10.0302847 | 0.00237 | 0.008981 | F | F | F | T |
| Slc46a1 | 1.269092 | 3.53148961 | 10.028926 | 0.00237 | 0.008984 | F | F | F | F |
| Zfp773 | 2.07507206 | 1.76750378 | 10.0288111 | 0.00237 | 0.008984 | T | F | T | T |
| Usp7 | 0.58601542 | 7.88611291 | 10.0218453 | 0.00238 | 0.009011 | F | F | T | F |
| Uqcrc2 | 0.64673748 | 6.88892685 | 10.020508 | 0.00238 | 0.009015 | F | F | F | F |
| Magi1 | -0.7159816 | 6.53020833 | 10.0189337 | 0.00239 | 0.009021 | T | F | T | T |
| Zfp276 | 1.13135302 | 3.91426818 | 10.0155897 | 0.00239 | 0.009033 | T | F | T | F |
| Paqr8 | -2.0989835 | 2.35499825 | 10.0149631 | 0.00239 | 0.009034 | T | T | T | T |
| Foxl2 | -7.6230089 | 0.25948709 | 10.0131413 | 0.00239 | 0.00904 | F | T | F | T |
| Dlg2 | -7.6239312 | 0.19328733 | 10.0096103 | 0.0024 | 0.009054 | T | F | T | T |
| Kank1 | 0.64177076 | 6.90763664 | 10.0088096 | 0.0024 | 0.009055 | F | F | T | F |
| Hfm1 | -7.6210785 | 0.19170298 | 10.0062899 | 0.0024 | 0.009065 | T | F | F | F |
| Rit1 | 1.18843144 | 3.75106718 | 10.0005794 | 0.00241 | 0.009087 | F | F | T | T |
| 1500009L1( | 1.32977225 | 3.33501109 | 9.9993047 | 0.00241 | 0.009091 | F | F | F | F |
| Dnajc9 | -0.8042721 | 5.80536406 | 9.99828886 | 0.00241 | 0.009094 | T | T | T | T |
| Vars2 | 0.78311723 | 5.50767897 | 9.98394374 | 0.00243 | 0.009153 | F | F | T | T |
| Grin3a | -7.5724089 | 0.18869543 | 9.98118384 | 0.00243 | 0.009164 | F | F | T | F |
| Slc27a6 | -7.5804677 | 0.17774218 | 9.97807309 | 0.00243 | 0.009175 | T | F | F | T |
| Kmt5c | -0.8127626 | 5.70907268 | 9.97770331 | 0.00243 | 0.009175 | F | F | F | F |
| Ptgr1 | 1.29541859 | 3.42263284 | 9.97036051 | 0.00244 | 0.009205 | F | F | T | F |
| Usp37 | -0.7127873 | 6.73432332 | 9.97018961 | 0.00244 | 0.009205 | F | F | T | F |
| Ablim3 | -7.5752797 | 0.2006448 | 9.96606941 | 0.00245 | 0.009221 | T | F | T | F |

| | | | | | | | | | |
|---|---|---|---|---|---|---|---|---|---|
| Akr1b3 | -0.6841702 | 6.95452226 | 9.96056369 | 0.00245 | 0.009243 | F | F | F | F |
| Pcgf5 | 0.84344628 | 5.04737623 | 9.95944596 | 0.00245 | 0.009246 | F | F | T | T |
| Ncdn | -0.7846647 | 5.88600533 | 9.9567635 | 0.00246 | 0.009256 | T | F | T | T |
| AU041133 | 1.71462842 | 2.47138026 | 9.95249791 | 0.00246 | 0.009273 | F | F | F | T |
| Gpt | -2.4142841 | 1.93906783 | 9.95144097 | 0.00246 | 0.009276 | T | F | F | F |
| Gm14418 | 1.86635269 | 2.19275462 | 9.94804288 | 0.00247 | 0.009289 | F | F | F | T |
| Xlr4c | 2.15037818 | 1.68756948 | 9.94565176 | 0.00247 | 0.009298 | T | T | F | T |
| Clec11a | -4.6469117 | 0.61732969 | 9.93168149 | 0.00249 | 0.009357 | F | F | F | F |
| 2310039H0 | -2.2610625 | 2.12466622 | 9.93029003 | 0.00249 | 0.009362 | F | F | F | T |
| Snx19 | -0.787964 | 5.84033094 | 9.92744735 | 0.00249 | 0.009373 | T | F | T | T |
| Gm8281 | -7.5787395 | 0.21005487 | 9.92590126 | 0.00249 | 0.009378 | F | F | F | F |
| Mat2b | -0.9198708 | 4.98496754 | 9.91899024 | 0.0025 | 0.009407 | T | T | T | T |
| Slco2a1 | 1.36125582 | 3.24473446 | 9.91400142 | 0.00251 | 0.009424 | T | F | T | T |
| 4930486L2 | 5.05605469 | -0.3457768 | 9.91392222 | 0.00251 | 0.009424 | F | F | T | F |
| Ube2i | -0.5905608 | 8.2943072 | 9.91387333 | 0.00251 | 0.009424 | T | F | T | F |
| Ccng1 | 0.73709972 | 5.87416539 | 9.91343173 | 0.00251 | 0.009425 | T | F | F | T |
| Cd79b | 7.51758709 | -0.8846289 | 9.92516903 | 0.00251 | 0.009425 | F | F | F | F |
| Tex43 | 7.51758709 | -0.8846289 | 9.92516903 | 0.00251 | 0.009425 | F | F | F | F |
| 1700029P1 | 7.51758709 | -0.8846289 | 9.92516903 | 0.00251 | 0.009425 | T | F | T | T |
| Cyp21a1 | 7.51758709 | -0.8846289 | 9.92516903 | 0.00251 | 0.009425 | F | F | F | F |
| H2ac13 | -0.6984235 | 7.52055918 | 9.90268896 | 0.00252 | 0.009464 | F | F | F | F |
| Ddx4 | -2.9545001 | 1.40027866 | 9.89639134 | 0.00253 | 0.00949 | T | F | F | T |
| Asic4 | -7.5617609 | 0.17596001 | 9.89592365 | 0.00253 | 0.009491 | F | F | F | F |
| Anks4b | -3.8514406 | 0.88482995 | 9.89256683 | 0.00253 | 0.009504 | F | F | F | F |
| Cog1 | 0.74449972 | 5.78448812 | 9.89078336 | 0.00253 | 0.00951 | T | F | F | T |
| Gdf2 | -7.568046 | 0.19662526 | 9.88995372 | 0.00253 | 0.009513 | T | F | T | T |
| Ash1l | 0.62575923 | 7.02352512 | 9.88935015 | 0.00254 | 0.009514 | T | T | F | T |
| Ppp1r16b | -1.1651166 | 4.14302738 | 9.88667421 | 0.00254 | 0.009524 | T | F | T | F |
| Hnrnpm | -0.5645018 | 8.92215345 | 9.87032495 | 0.00256 | 0.009596 | T | F | F | F |
| Olfr635 | 0.84024861 | 5.24822849 | 9.86654808 | 0.00256 | 0.009611 | F | F | F | F |
| Pcdhga5 | -4.322102 | 0.70466551 | 9.86198837 | 0.00257 | 0.00963 | F | F | T | F |
| Btbd1 | -0.7723223 | 5.95742249 | 9.86159512 | 0.00257 | 0.00963 | T | T | T | T |
| Cpd | 0.62306216 | 7.0455809 | 9.85881083 | 0.00257 | 0.009641 | T | F | T | T |
| Add1 | 0.63064358 | 6.96058772 | 9.85455988 | 0.00258 | 0.009658 | T | F | F | T |
| Vmn2r76 | 0.95472517 | 4.51885978 | 9.85331353 | 0.00258 | 0.009662 | F | F | F | F |
| Casq1 | 3.16423339 | 0.60970349 | 9.85192997 | 0.00258 | 0.009667 | F | F | T | F |
| Mettl9 | -0.7812694 | 5.87622985 | 9.84960129 | 0.00258 | 0.009676 | T | F | F | F |
| Ebpl | 0.91592764 | 4.63836341 | 9.84392965 | 0.00259 | 0.0097 | F | F | T | T |
| Ddx23 | -0.6370092 | 7.30250659 | 9.84152759 | 0.00259 | 0.009709 | F | F | T | F |
| Col9a2 | -4.6255279 | 0.61551522 | 9.82962951 | 0.00261 | 0.009762 | F | F | F | F |
| 5031439G0 | 0.67853976 | 6.45111734 | 9.82247533 | 0.00262 | 0.009793 | T | F | T | F |
| Dcn | 3.99323053 | 0.02596064 | 9.81655305 | 0.00262 | 0.009819 | T | F | F | T |

| | | | | | | | | | |
|---|---|---|---|---|---|---|---|---|---|
| Sh3bp5 | 0.97202035 | 4.35656877 | 9.81459309 | 0.00263 | 0.009826 | F | F | T | F |
| Stau1 | 0.66467477 | 6.58656482 | 9.81262609 | 0.00263 | 0.009833 | F | F | T | F |
| Pofut2 | 0.70112384 | 6.35318545 | 9.81050604 | 0.00263 | 0.009842 | F | F | F | F |
| Sirt5 | 0.7810532 | 5.95115342 | 9.80626576 | 0.00264 | 0.009859 | T | F | T | F |
| Xiap | 0.68293408 | 6.47636869 | 9.79704867 | 0.00265 | 0.009899 | T | F | T | T |
| P3h2 | -0.8480202 | 5.37069981 | 9.7964561 | 0.00265 | 0.009899 | T | F | T | T |
| Map3k13 | -2.0845515 | 2.33871893 | 9.79623844 | 0.00265 | 0.009899 | F | F | T | T |
| Pfdn6 | -0.7961724 | 5.71694225 | 9.796238 | 0.00265 | 0.009899 | F | F | F | F |
| Hmmr | 0.63569349 | 6.94471178 | 9.78683375 | 0.00266 | 0.009941 | T | F | T | T |
| Kctd21 | 1.81459943 | 2.20555908 | 9.78366433 | 0.00266 | 0.009954 | T | F | T | F |
| Mfap3l | -1.0880139 | 4.26579616 | 9.77221233 | 0.00268 | 0.010005 | F | F | T | T |
| Acvr2b | -0.6245752 | 7.49517019 | 9.77204963 | 0.00268 | 0.010005 | F | F | T | F |
| Trim67 | -3.1513632 | 1.25828764 | 9.76942422 | 0.00268 | 0.010016 | T | F | T | F |
| Mindy1 | 1.00553272 | 4.27364409 | 9.76915623 | 0.00268 | 0.010016 | F | F | F | F |
| Grb14 | -3.9278215 | 0.8405704 | 9.76588712 | 0.00269 | 0.010029 | T | F | F | T |
| Prom1 | -0.735396 | 6.26354549 | 9.75835464 | 0.0027 | 0.010063 | T | F | T | T |
| Sh3glb2 | -1.1718509 | 4.07214463 | 9.75751437 | 0.0027 | 0.010065 | F | F | F | F |
| Pianp | -1.8284088 | 2.75276036 | 9.75070399 | 0.00271 | 0.010096 | F | F | T | F |
| Ep400 | 0.5463535 | 8.73415946 | 9.7472567 | 0.00271 | 0.010111 | T | F | T | F |
| Fblim1 | -0.8232348 | 5.49706354 | 9.73353151 | 0.00273 | 0.010174 | T | F | T | T |
| Bdnf | -7.567078 | 0.15210414 | 9.71925076 | 0.00275 | 0.010241 | T | T | F | T |
| Ap3m2 | 2.07167933 | 1.72997863 | 9.71552749 | 0.00275 | 0.010257 | F | F | F | F |
| Mrpl38 | 0.74886107 | 5.78319267 | 9.71116366 | 0.00276 | 0.010277 | T | F | F | F |
| Tprn | -1.3475426 | 3.62827749 | 9.70490889 | 0.00276 | 0.010305 | T | T | F | T |
| Mapk15 | -7.5504336 | 0.14714211 | 9.70265851 | 0.00277 | 0.010314 | F | F | F | F |
| P3h1 | 0.8296997 | 5.07968331 | 9.70052042 | 0.00277 | 0.010323 | F | F | T | F |
| Dnaaf4 | -7.559681 | 0.14785702 | 9.69210565 | 0.00278 | 0.010362 | T | F | T | T |
| mt-Nd2 | 0.57342072 | 8.03855421 | 9.6918422 | 0.00278 | 0.010362 | F | F | F | F |
| Stk17b | 1.10703198 | 3.96243669 | 9.68766553 | 0.00279 | 0.01038 | T | F | T | F |
| Atxn2l | 0.54177789 | 8.77573499 | 9.68685689 | 0.00279 | 0.010382 | F | F | T | F |
| Rhox4e | 4.10641503 | 0.05350375 | 9.68152329 | 0.0028 | 0.010407 | F | F | F | F |
| Fan1 | -1.4842161 | 3.31534168 | 9.67000983 | 0.00281 | 0.010461 | F | F | T | T |
| Notch2 | 0.54891993 | 8.57744925 | 9.66977127 | 0.00281 | 0.010461 | T | F | F | T |
| Mphosph9 | -0.8527277 | 5.26472611 | 9.66832659 | 0.00281 | 0.010466 | F | F | T | T |
| Pole | -0.6270942 | 7.38324517 | 9.66468165 | 0.00282 | 0.010482 | F | F | F | F |
| Sf3a3 | -0.6607752 | 6.9413614 | 9.66322911 | 0.00282 | 0.010488 | F | F | F | F |
| Ubiad1 | 0.77616208 | 5.41873605 | 9.66182291 | 0.00282 | 0.010493 | T | F | T | T |
| 1110032A0 | -1.9923459 | 2.45210978 | 9.64753209 | 0.00284 | 0.010562 | T | F | T | T |
| Arhgef5 | 0.73252513 | 5.85282461 | 9.64451822 | 0.00284 | 0.010575 | T | F | F | F |
| Vmn1r183 | 1.24513003 | 3.51034181 | 9.63588281 | 0.00286 | 0.010617 | F | F | F | F |
| Plagl1 | -1.564806 | 3.27867275 | 9.63074406 | 0.00286 | 0.010641 | T | F | F | F |
| Cd38 | -1.7774658 | 2.78375951 | 9.6281927 | 0.00287 | 0.010652 | T | T | F | T |

| | | | | | | | | | |
|---|---|---|---|---|---|---|---|---|---|
| Timm29 | 0.730173 | 5.83475618 | 9.62776846 | 0.00287 | 0.010652 | T | F | T | F |
| Raly | -0.6061471 | 7.8641718 | 9.62338386 | 0.00287 | 0.010672 | F | F | F | T |
| Ppp5c | -0.6634826 | 6.90953115 | 9.62052877 | 0.00288 | 0.010685 | F | F | T | F |
| Jph1 | -1.2907173 | 3.79958514 | 9.61968109 | 0.00288 | 0.010687 | F | F | F | F |
| Ranbp2 | 0.53660935 | 8.9481238 | 9.61746834 | 0.00288 | 0.010696 | F | F | F | F |
| Gmps | -0.6040821 | 7.79061281 | 9.61518411 | 0.00288 | 0.010706 | F | F | T | T |
| Kcnh5 | -7.5566875 | 0.19874476 | 9.60977579 | 0.00289 | 0.010732 | F | F | F | T |
| Tmem107 | -2.1804758 | 2.15251654 | 9.60724264 | 0.00289 | 0.010743 | F | F | T | F |
| Fam98c | 1.49238825 | 2.8899295 | 9.60391745 | 0.0029 | 0.010758 | T | F | T | F |
| Nos1ap | 1.8757992 | 2.10524553 | 9.59675958 | 0.00291 | 0.010792 | T | F | T | F |
| Syt1 | -7.5229402 | 0.14023568 | 9.58824584 | 0.00292 | 0.010834 | T | F | T | T |
| Pxdc1 | 1.53080597 | 2.79624633 | 9.58777521 | 0.00292 | 0.010834 | T | F | T | T |
| Efnb2 | 0.632532 | 6.84415509 | 9.57945548 | 0.00293 | 0.010875 | T | F | F | T |
| Nos3 | -4.6196208 | 0.62560291 | 9.57787186 | 0.00294 | 0.010881 | T | F | F | F |
| Pola2 | -0.7235951 | 6.29926536 | 9.57085953 | 0.00294 | 0.010915 | T | F | T | F |
| Chtf8 | -0.607349 | 7.69801474 | 9.56904709 | 0.00295 | 0.010923 | F | F | T | T |
| Lrrc38 | -4.7309312 | 0.66249171 | 9.56877052 | 0.00295 | 0.010923 | F | F | F | F |
| Stil | 0.77425631 | 5.42162276 | 9.55487593 | 0.00297 | 0.010993 | F | F | F | T |
| Mctp2 | 1.88907323 | 2.11534522 | 9.54703346 | 0.00298 | 0.011031 | T | F | T | T |
| Samd4b | 0.58351882 | 7.66628436 | 9.54579506 | 0.00298 | 0.011036 | T | F | T | T |
| Bmi1 | -1.0007604 | 4.53306838 | 9.54531324 | 0.00298 | 0.011037 | T | F | T | T |
| Usp34 | 0.55785754 | 8.42576268 | 9.53607841 | 0.00299 | 0.011083 | T | F | F | F |
| Kctd6 | 0.94080789 | 4.4940114 | 9.52438748 | 0.00301 | 0.011142 | F | F | F | T |
| Rnf180 | -7.5225649 | 0.1268997 | 9.5216926 | 0.00301 | 0.011155 | F | F | T | T |
| Dlx2 | -7.5487133 | 0.13209216 | 9.52125357 | 0.00301 | 0.011155 | F | F | T | F |
| Scn1b | -7.515697 | 0.12956476 | 9.51895816 | 0.00302 | 0.011165 | F | F | T | F |
| Sass6 | 0.9896364 | 4.24794014 | 9.51795398 | 0.00302 | 0.011169 | F | F | T | F |
| Tsr1 | 0.58013459 | 7.64786571 | 9.51688593 | 0.00302 | 0.011173 | T | F | F | F |
| Gosr1 | 0.78140987 | 5.32103816 | 9.51594916 | 0.00302 | 0.011176 | T | T | T | T |
| Ino80e | 0.68714652 | 6.25016181 | 9.51441458 | 0.00302 | 0.011182 | T | F | T | F |
| Vcam1 | -7.5320365 | 0.17314556 | 9.51388202 | 0.00303 | 0.011183 | T | F | F | F |
| Rmdn2 | 1.47103696 | 2.92640798 | 9.51312314 | 0.00303 | 0.011185 | F | F | T | T |
| Slfn1 | 5.00526872 | -0.3878653 | 9.50925535 | 0.00303 | 0.011203 | T | F | T | T |
| Atp5d | 0.60071166 | 7.44232765 | 9.50611054 | 0.00304 | 0.011218 | F | F | T | F |
| Plekhg2 | -0.7084143 | 6.47710923 | 9.50505174 | 0.00304 | 0.011222 | F | F | F | F |
| Mob3a | 0.78399201 | 5.3719898 | 9.5034358 | 0.00304 | 0.011228 | T | F | T | F |
| Lrrc18 | -3.5599608 | 1.00589359 | 9.49342979 | 0.00305 | 0.01128 | T | F | F | T |
| Lrrc56 | -1.6547052 | 2.96899177 | 9.48872775 | 0.00306 | 0.011303 | F | F | T | F |
| Foxred2 | -1.3136012 | 3.67661788 | 9.4837834 | 0.00307 | 0.011327 | T | F | T | T |
| Pak1ip1 | 0.69762585 | 6.11400156 | 9.4812183 | 0.00307 | 0.011339 | F | F | F | T |
| Cnbd2 | 2.46225582 | 1.180199 | 9.47879177 | 0.00308 | 0.01135 | F | F | F | F |
| Insr | 0.6636006 | 6.53023903 | 9.4784824 | 0.00308 | 0.01135 | T | F | T | T |

| | | | | | | | | | |
|---|---|---|---|---|---|---|---|---|---|
| Tmx3 | 0.73715477 | 5.72371308 | 9.47774753 | 0.00308 | 0.011352 | T | F | T | T |
| Sost | -7.6190633 | 0.26498234 | 9.47505429 | 0.00308 | 0.011365 | F | F | T | F |
| Slc16a11 | -7.5304894 | 0.12281395 | 9.47464164 | 0.00308 | 0.011365 | T | F | F | F |
| Gpsm2 | 0.778437 | 5.3723963 | 9.47394 | 0.00308 | 0.011367 | F | F | F | F |
| Sin3a | -0.6063902 | 7.7147064 | 9.46767591 | 0.00309 | 0.011399 | F | F | F | F |
| Scg5 | 2.39635921 | 1.28703963 | 9.46472889 | 0.0031 | 0.011413 | F | F | F | F |
| Cnppd1 | 0.79013564 | 5.23464776 | 9.46242063 | 0.0031 | 0.011423 | F | F | F | F |
| Rpgr | -1.1287916 | 4.15327117 | 9.45977094 | 0.0031 | 0.011435 | T | F | T | F |
| Angpt1 | -7.5184822 | 0.1582774 | 9.45904372 | 0.0031 | 0.011437 | T | F | F | T |
| Cdcp2 | -7.5065259 | 0.13648106 | 9.45638294 | 0.00311 | 0.01145 | T | F | F | F |
| Lztr1 | 0.77460667 | 5.38635024 | 9.4496483 | 0.00312 | 0.011485 | T | F | F | F |
| Nhlrc2 | -0.725953 | 6.23220951 | 9.44452882 | 0.00313 | 0.01151 | T | F | T | T |
| Mospd3 | 1.01456242 | 4.16927077 | 9.44416312 | 0.00313 | 0.01151 | F | F | F | F |
| Adtrp | 3.41553117 | 0.34495304 | 9.4426297 | 0.00313 | 0.011517 | T | F | T | T |
| Dusp10 | -2.3693264 | 1.92276569 | 9.43982591 | 0.00313 | 0.01153 | F | F | T | F |
| Efhb | -7.5327804 | 0.12053698 | 9.43634226 | 0.00314 | 0.011547 | F | F | F | F |
| Olfr1167 | 1.53685326 | 2.82065878 | 9.43437116 | 0.00314 | 0.011555 | F | F | F | F |
| Cd300e | -7.5100602 | 0.14748619 | 9.43428337 | 0.00314 | 0.011555 | F | F | T | F |
| Dennd2a | 0.65149291 | 6.60637917 | 9.42343099 | 0.00316 | 0.011612 | F | F | F | F |
| Pdx1 | -7.5193709 | 0.11567146 | 9.41969996 | 0.00316 | 0.011631 | F | F | F | F |
| Vps13d | 0.6560761 | 6.68185134 | 9.41888149 | 0.00316 | 0.011633 | T | F | F | T |
| Frmpd2 | -7.4999934 | 0.12257852 | 9.41558114 | 0.00317 | 0.011648 | F | F | T | F |
| Emid1 | -1.849992 | 2.64129527 | 9.41548555 | 0.00317 | 0.011648 | F | F | F | F |
| Mettl27 | -1.9966877 | 2.39324855 | 9.41205148 | 0.00317 | 0.011665 | T | F | T | F |
| Lrpprc | 0.5620107 | 7.96544956 | 9.40949088 | 0.00318 | 0.011677 | T | F | F | T |
| Akna | -0.9949078 | 4.55866174 | 9.40741432 | 0.00318 | 0.011687 | T | F | F | F |
| Wrap53 | -0.9032314 | 4.93934586 | 9.40710333 | 0.00318 | 0.011687 | F | F | F | F |
| Dpp7 | -1.3761996 | 3.50522305 | 9.40142382 | 0.00319 | 0.011716 | F | F | F | F |
| Fam174b | -1.8568682 | 2.64811306 | 9.39983283 | 0.00319 | 0.011723 | T | F | F | T |
| Slc12a9 | 0.8727051 | 4.7909917 | 9.39065141 | 0.00321 | 0.011772 | F | F | F | F |
| Oma1 | 0.91579876 | 4.52707091 | 9.3898783 | 0.00321 | 0.011773 | F | F | F | F |
| Tnfaip8l1 | 2.33839787 | 1.2780534 | 9.38970357 | 0.00321 | 0.011773 | F | F | T | F |
| Cyb5r1 | -0.9690272 | 4.67537369 | 9.38566922 | 0.00321 | 0.011794 | T | F | T | T |
| Gm21874 | -7.5349336 | 0.11755133 | 9.37890437 | 0.00322 | 0.01183 | F | F | F | F |
| Gm4631 | 0.71124458 | 5.93578973 | 9.37395457 | 0.00323 | 0.011855 | F | F | F | F |
| Stk39 | -0.8912648 | 4.99149827 | 9.3735311 | 0.00323 | 0.011856 | F | F | F | F |
| Gm14139 | 2.40253936 | 1.26991135 | 9.37051408 | 0.00324 | 0.011871 | F | F | F | F |
| Slc7a11 | 2.71759766 | 0.85008264 | 9.37006799 | 0.00324 | 0.011871 | T | T | T | T |
| Ltbp2 | 7.44002192 | -0.9465767 | 9.37988206 | 0.00324 | 0.011876 | F | F | F | F |
| Best3 | 7.44002192 | -0.9465767 | 9.37988206 | 0.00324 | 0.011876 | F | F | T | T |
| Rnf183 | 7.44002192 | -0.9465767 | 9.37988206 | 0.00324 | 0.011876 | T | F | T | T |
| Slc25a19 | 1.05574733 | 4.06055387 | 9.3677454 | 0.00324 | 0.011876 | T | F | T | T |

| | | | | | | | | | |
|---|---|---|---|---|---|---|---|---|---|
| Trat1 | 1.55752385 | 2.70085619 | 9.36372238 | 0.00325 | 0.011897 | F | F | T | F |
| Ermp1 | 0.68311415 | 6.29072234 | 9.34659437 | 0.00327 | 0.011992 | T | F | T | T |
| Tnik | -4.5908099 | 0.56061223 | 9.34378857 | 0.00328 | 0.012006 | T | F | T | T |
| Ybx3 | 0.53434452 | 8.70546547 | 9.34198432 | 0.00328 | 0.012014 | F | F | F | F |
| Zfat | 0.87026529 | 4.75366615 | 9.33305754 | 0.0033 | 0.012063 | F | F | F | F |
| Olfr1380 | 7.43467291 | -0.9508178 | 9.34336199 | 0.0033 | 0.01207 | F | F | F | F |
| Gm20390 | 0.71297729 | 5.95511123 | 9.32993061 | 0.0033 | 0.012077 | F | F | F | T |
| Srxn1 | 1.3653672 | 3.21694891 | 9.3261931 | 0.00331 | 0.012096 | T | F | F | F |
| Rab2b | -1.5817041 | 3.0759002 | 9.32101484 | 0.00331 | 0.012124 | T | F | F | T |
| Spryd3 | 0.84233892 | 4.91034035 | 9.32015872 | 0.00332 | 0.012127 | T | F | T | F |
| Tent5a | -1.7705212 | 2.74843287 | 9.31453353 | 0.00332 | 0.012157 | F | F | T | F |
| Hcrtr1 | 4.54500612 | -0.2640669 | 9.30722246 | 0.00334 | 0.012198 | F | F | F | F |
| Plekhm1 | 0.75870582 | 5.45595343 | 9.30660717 | 0.00334 | 0.012199 | T | F | T | T |
| Dlat | 0.648611 | 6.57974079 | 9.30456141 | 0.00334 | 0.012209 | T | T | T | T |
| Hs6st3 | -7.5231474 | 0.10802035 | 9.30416613 | 0.00334 | 0.012209 | F | F | F | T |
| Tmem214 | 0.82904866 | 5.00474485 | 9.2999159 | 0.00335 | 0.012232 | F | F | T | T |
| Cct6b | 5.59227015 | -0.6155328 | 9.29910256 | 0.00335 | 0.012234 | F | F | F | T |
| Gdap2 | 0.89354435 | 4.61967635 | 9.2958781 | 0.00335 | 0.012251 | T | F | T | T |
| Edil3 | -7.4886812 | 0.12635446 | 9.29538554 | 0.00336 | 0.012252 | T | F | F | F |
| Zfp449 | -1.4649279 | 3.28893567 | 9.29316289 | 0.00336 | 0.012263 | F | F | T | F |
| Mlec | -0.5631533 | 8.59959667 | 9.28929608 | 0.00336 | 0.012283 | T | F | T | T |
| Adar | -0.7593033 | 5.88801935 | 9.27152281 | 0.00339 | 0.012385 | T | F | F | F |
| Nr4a1 | -2.3295221 | 1.89657358 | 9.26405033 | 0.00341 | 0.012427 | F | F | F | F |
| Zfp52 | 1.73772481 | 2.26995526 | 9.2636469 | 0.00341 | 0.012428 | F | F | F | T |
| Dppa5a | -7.5085719 | 0.09994765 | 9.26324984 | 0.00341 | 0.012428 | T | T | F | T |
| Ripply2 | -7.4899114 | 0.09702503 | 9.26261214 | 0.00341 | 0.012429 | F | F | F | F |
| Sdccag8 | -2.4074629 | 1.82838091 | 9.25738476 | 0.00342 | 0.012458 | F | F | F | T |
| Rufy3 | -0.9010169 | 4.93188401 | 9.25611202 | 0.00342 | 0.012463 | T | F | T | T |
| Has3 | -4.5737683 | 0.5488703 | 9.25579761 | 0.00342 | 0.012463 | T | F | T | F |
| Mapk11 | -2.1155851 | 2.17271533 | 9.24733698 | 0.00343 | 0.012512 | F | F | F | F |
| Padi2 | 1.19334768 | 3.65635996 | 9.24687682 | 0.00343 | 0.012512 | F | F | T | F |
| Rabggtb | 0.78342432 | 5.25114635 | 9.24586855 | 0.00344 | 0.012516 | F | F | F | F |
| Txndc16 | -1.3705165 | 3.52587263 | 9.23228493 | 0.00346 | 0.012595 | T | F | T | T |
| Tia1 | -0.8151867 | 5.40985038 | 9.22506117 | 0.00347 | 0.012636 | T | F | T | T |
| Bloc1s2 | 0.9141289 | 4.49413594 | 9.216962 | 0.00348 | 0.012683 | T | F | F | T |
| Plxnc1 | -1.8742885 | 2.71152559 | 9.2073162 | 0.0035 | 0.012739 | F | F | T | F |
| Ubr7 | -0.7155714 | 6.30404694 | 9.19723917 | 0.00352 | 0.012798 | T | F | T | T |
| Bri3bp | 0.61764629 | 6.87493674 | 9.18739907 | 0.00353 | 0.012856 | T | F | F | T |
| Eef1a2 | -7.4734856 | 0.09019276 | 9.18683844 | 0.00353 | 0.012857 | F | F | F | T |
| Slc22a18 | 1.77690028 | 2.1692495 | 9.18646535 | 0.00353 | 0.012858 | F | F | F | F |
| Nat9 | 1.80862136 | 2.29016455 | 9.1856606 | 0.00353 | 0.01286 | F | F | T | F |
| Acy3 | -3.0343333 | 1.22666873 | 9.1780868 | 0.00355 | 0.012905 | T | F | F | T |

| | | | | | | | | | |
|---|---|---|---|---|---|---|---|---|---|
| Ndfip2 | 0.77661974 | 5.24392958 | 9.17644612 | 0.00355 | 0.012913 | T | F | F | T |
| Pla2g15 | 0.88724644 | 4.61729278 | 9.16702023 | 0.00357 | 0.012968 | T | F | T | F |
| Ube2j1 | -0.7937083 | 5.528695 | 9.16613216 | 0.00357 | 0.012972 | T | T | F | T |
| Pcdh9 | -7.4778315 | 0.08564506 | 9.16551416 | 0.00357 | 0.012973 | T | T | F | T |
| Mllt1 | 0.61719262 | 6.86520904 | 9.16485618 | 0.00357 | 0.012975 | F | F | T | T |
| Epha2 | -0.9995581 | 4.76843209 | 9.16409197 | 0.00357 | 0.012978 | F | F | F | F |
| Tlx2 | -7.5703215 | 0.22255313 | 9.16358557 | 0.00357 | 0.012979 | F | F | F | F |
| Slc2a6 | 3.51031862 | 0.28232844 | 9.16036497 | 0.00358 | 0.012997 | T | F | T | T |
| Haghl | -1.8400829 | 2.59564695 | 9.15891547 | 0.00358 | 0.013003 | F | F | T | F |
| Eif3j2 | 0.63576718 | 6.659117 | 9.15523255 | 0.00359 | 0.013022 | F | F | T | T |
| Pigu | 0.85016186 | 4.81008116 | 9.15489883 | 0.00359 | 0.013022 | T | F | F | F |
| Rimbp3 | 1.79357506 | 2.13895698 | 9.15488914 | 0.00359 | 0.013022 | F | F | F | F |
| Gm17571 | 1.65523286 | 2.63329275 | 9.15148416 | 0.00359 | 0.013041 | F | F | F | F |
| Zfp397 | 0.75674945 | 5.53919522 | 9.15068666 | 0.00359 | 0.013043 | T | T | F | T |
| Nes | -0.6827623 | 6.54047267 | 9.1450721 | 0.0036 | 0.013076 | F | F | F | F |
| Sar1b | 0.84879521 | 4.82152285 | 9.14029503 | 0.00361 | 0.013104 | F | F | F | F |
| Coro2a | 1.3036659 | 3.25574046 | 9.13850798 | 0.00362 | 0.013113 | F | F | T | F |
| Tbc1d22b | 0.72404027 | 5.72426493 | 9.13667294 | 0.00362 | 0.013122 | F | F | T | F |
| Skil | -0.7781676 | 5.68494989 | 9.13015766 | 0.00363 | 0.013161 | T | T | T | T |
| Fam131c | -7.461362 | 0.09324179 | 9.12622481 | 0.00364 | 0.013183 | F | F | T | F |
| Edem3 | 0.62721787 | 6.77200671 | 9.12489973 | 0.00364 | 0.013189 | T | F | T | T |
| Zfp646 | -0.6182984 | 7.23188042 | 9.1135424 | 0.00366 | 0.013259 | T | F | T | F |
| Osbpl10 | 0.93366512 | 4.37950027 | 9.10701448 | 0.00367 | 0.013298 | F | F | F | F |
| Cpt1c | -1.8023966 | 2.66107294 | 9.10413264 | 0.00367 | 0.013314 | T | F | F | T |
| Rcan3 | -1.4915771 | 3.21466396 | 9.08954283 | 0.0037 | 0.013405 | F | F | T | F |
| Hspa13 | 0.75505542 | 5.4086067 | 9.08753235 | 0.0037 | 0.013415 | T | F | T | F |
| Cyp4f13 | 2.18894365 | 1.44600806 | 9.08463172 | 0.00371 | 0.01343 | F | F | F | F |
| Ints12 | 0.86649374 | 4.70471674 | 9.08455453 | 0.00371 | 0.01343 | T | F | F | F |
| Nucks1 | -0.5335678 | 9.3246431 | 9.08245061 | 0.00371 | 0.013441 | T | F | T | T |
| Mbd6 | 0.67647403 | 6.18959623 | 9.0803881 | 0.00372 | 0.013449 | F | F | F | F |
| Tlr6 | 4.17882483 | -0.0658654 | 9.08024274 | 0.00372 | 0.013449 | F | F | F | T |
| Timp2 | -1.0197681 | 4.44443207 | 9.08009304 | 0.00372 | 0.013449 | T | F | T | F |
| Klhl20 | 0.94779624 | 4.3505007 | 9.07919625 | 0.00372 | 0.013453 | T | F | T | T |
| Esd | -0.7416457 | 5.94074295 | 9.06841521 | 0.00374 | 0.01352 | F | F | T | F |
| Rrm2 | 0.52598283 | 8.73846995 | 9.05962508 | 0.00375 | 0.013575 | T | F | T | T |
| Sec24d | 0.82314784 | 5.01735064 | 9.05843659 | 0.00376 | 0.01358 | T | F | F | F |
| Scamp2 | 0.72117696 | 5.73810626 | 9.05751097 | 0.00376 | 0.013584 | F | F | F | F |
| Mxi1 | -1.1767357 | 3.95406407 | 9.0528831 | 0.00377 | 0.013612 | T | F | T | F |
| Txn1 | 0.59302969 | 7.11676449 | 9.04896861 | 0.00377 | 0.013635 | T | F | F | F |
| Cox7a2l | 0.63638864 | 6.62039449 | 9.04853657 | 0.00377 | 0.013635 | T | F | F | F |
| Actn3 | -1.7060476 | 2.79894135 | 9.04603389 | 0.00378 | 0.013649 | F | F | T | F |
| H2bc8 | -0.5394992 | 9.1747673 | 9.04399845 | 0.00378 | 0.01366 | F | F | F | F |

| | | | | | | | | | |
|---|---|---|---|---|---|---|---|---|---|
| Rbm12 | -0.6455432 | 7.00471469 | 9.03938728 | 0.00379 | 0.013688 | F | F | T | T |
| C1s1 | 2.31799166 | 1.24983246 | 9.03825573 | 0.00379 | 0.013693 | F | F | T | F |
| Cyb5r4 | -0.7808652 | 5.5798629 | 9.03355782 | 0.0038 | 0.013722 | T | F | F | T |
| Klhl5 | -1.3562425 | 3.55721036 | 9.02789587 | 0.00381 | 0.013757 | F | F | F | T |
| Trpm5 | 5.51804606 | -0.6107188 | 9.0245678 | 0.00382 | 0.013776 | F | F | F | T |
| Fdx2 | 1.11608634 | 3.77767959 | 9.00353635 | 0.00386 | 0.013913 | F | F | F | F |
| Zfyve16 | 0.75728436 | 5.38498938 | 9.00234385 | 0.00386 | 0.013919 | T | F | F | T |
| Polrmt | 0.70648674 | 5.84646848 | 8.99913153 | 0.00386 | 0.013938 | F | F | F | F |
| Kcne1 | -4.5583535 | 0.52754729 | 8.99789602 | 0.00387 | 0.013944 | T | F | T | F |
| Dhcr7 | 0.7966208 | 5.08199151 | 8.99452359 | 0.00387 | 0.013964 | T | F | T | T |
| Cit | -0.612053 | 7.28564692 | 8.99021624 | 0.00388 | 0.01399 | T | F | T | F |
| Ttc6 | -3.8508864 | 0.75861009 | 8.98383042 | 0.00389 | 0.014031 | F | F | F | F |
| Gm3739 | -4.0414197 | 0.68103606 | 8.98282266 | 0.00389 | 0.014035 | F | F | F | F |
| Trappc6b | 1.14925478 | 3.7086064 | 8.97820495 | 0.0039 | 0.014062 | T | T | F | T |
| Panx1 | 1.02669586 | 4.08703911 | 8.97807247 | 0.0039 | 0.014062 | F | F | T | F |
| Zfp512 | -0.7489012 | 5.84730462 | 8.97452932 | 0.00391 | 0.014084 | T | T | T | T |
| Cyyr1 | -1.4958798 | 3.17313357 | 8.97045477 | 0.00392 | 0.014109 | T | F | T | T |
| Ears2 | -1.1025583 | 4.11264614 | 8.96700739 | 0.00392 | 0.01413 | F | F | T | F |
| Pxmp4 | 1.09274335 | 3.84399443 | 8.96469011 | 0.00393 | 0.014143 | T | F | F | T |
| Hmgxb4 | -1.9429524 | 2.38905294 | 8.96390167 | 0.00393 | 0.014146 | F | F | T | F |
| Rabl6 | -0.6754816 | 6.54883234 | 8.95729105 | 0.00394 | 0.014189 | F | F | F | F |
| Sccpdh | -0.8270635 | 5.28520532 | 8.95457184 | 0.00395 | 0.014205 | F | F | T | F |
| Rnf220 | -0.6524494 | 6.75509999 | 8.95272933 | 0.00395 | 0.014215 | F | F | F | F |
| Rexo2 | 0.7612744 | 5.30550281 | 8.94525987 | 0.00396 | 0.014263 | T | F | T | F |
| Eprs | 0.51246865 | 9.00957227 | 8.94451101 | 0.00397 | 0.014266 | T | F | F | F |
| Ptcra | 1.77512922 | 2.11916307 | 8.93618574 | 0.00398 | 0.014321 | F | F | T | F |
| Keap1 | -0.9177127 | 4.76365652 | 8.93198951 | 0.00399 | 0.014347 | T | F | F | F |
| Gm11100 | 2.63519311 | 0.86063207 | 8.93167251 | 0.00399 | 0.014347 | F | F | F | F |
| Ager | -7.4405432 | 0.08663131 | 8.93096885 | 0.00399 | 0.014349 | F | F | F | F |
| Matr3 | -0.5695343 | 8.23178259 | 8.92120401 | 0.00401 | 0.014414 | F | F | F | F |
| Dap3 | 0.66354517 | 6.32473952 | 8.91372625 | 0.00402 | 0.014464 | F | F | T | F |
| Arfrp1 | 1.31513789 | 3.23583692 | 8.89794522 | 0.00406 | 0.014571 | T | F | T | T |
| Btbd11 | -1.9358957 | 2.39361147 | 8.89677355 | 0.00406 | 0.014577 | T | F | T | T |
| Hjv | -7.4431118 | 0.05362985 | 8.89580336 | 0.00406 | 0.01458 | F | F | F | F |
| Kcnc2 | -7.4430846 | 0.05360473 | 8.89559402 | 0.00406 | 0.01458 | T | F | T | T |
| Nop10 | -0.8053247 | 5.33130782 | 8.89365016 | 0.00406 | 0.014591 | F | F | F | F |
| Efcab6 | -7.4326626 | 0.05451304 | 8.88507184 | 0.00408 | 0.014647 | F | F | F | F |
| Trp53i13 | 1.39981684 | 2.97683806 | 8.88496667 | 0.00408 | 0.014647 | T | F | T | T |
| Abca4 | -2.0442457 | 2.20543059 | 8.87885226 | 0.00409 | 0.014688 | T | F | F | F |
| Rfx1 | -0.8109314 | 5.30017254 | 8.87336107 | 0.0041 | 0.014724 | T | F | T | F |
| Nr6a1 | 0.53463735 | 9.11856896 | 8.87160484 | 0.00411 | 0.014734 | T | F | T | T |
| Mfsd1 | 0.58353947 | 7.14082349 | 8.86653345 | 0.00412 | 0.014767 | F | F | T | F |

| | | | | | | | | | |
|---|---|---|---|---|---|---|---|---|---|
| Lpcat2 | -3.8727816 | 0.76348725 | 8.85442004 | 0.00414 | 0.014851 | F | F | F | F |
| Metrnl | 2.03087681 | 1.64990812 | 8.85386228 | 0.00414 | 0.014852 | T | F | T | T |
| Alkbh7 | -1.266776 | 3.68688785 | 8.84514127 | 0.00416 | 0.014912 | T | F | F | T |
| Hoxb7 | -7.4640533 | 0.12267092 | 8.84447743 | 0.00416 | 0.014915 | F | F | T | F |
| Vmn2r102 | 1.37964698 | 3.03994255 | 8.83816255 | 0.00417 | 0.014957 | F | F | F | F |
| Nat2 | -3.0063259 | 1.19530525 | 8.83481268 | 0.00418 | 0.014979 | T | F | F | T |
| Samd12 | -3.4008901 | 0.96799074 | 8.83267627 | 0.00418 | 0.014992 | T | F | T | T |
| Entr1 | 0.71149563 | 5.71635583 | 8.8282425 | 0.00419 | 0.015021 | T | F | T | F |
| Stkld1 | 7.35804839 | -1.0111275 | 8.83511635 | 0.0042 | 0.015046 | F | F | F | F |
| Phospho1 | 1.70349741 | 2.27381341 | 8.82188521 | 0.00421 | 0.01506 | F | F | F | F |
| Nfs1 | 0.79745454 | 5.05735356 | 8.82179728 | 0.00421 | 0.01506 | F | F | T | F |
| Plod2 | -0.829641 | 5.17707561 | 8.8192218 | 0.00421 | 0.015075 | T | F | F | T |
| Itch | -0.7127349 | 6.12751112 | 8.81913044 | 0.00421 | 0.015075 | T | F | F | T |
| Clu | -1.8987317 | 2.4847848 | 8.81823632 | 0.00421 | 0.015079 | F | F | F | F |
| 2010315B0 | -2.2452116 | 1.91158644 | 8.80747869 | 0.00424 | 0.015154 | T | F | F | F |
| Tmem56 | -7.4183118 | 0.06038684 | 8.80614348 | 0.00424 | 0.015161 | T | F | T | F |
| Vac14 | 0.6472864 | 6.42983985 | 8.80568504 | 0.00424 | 0.015162 | T | F | F | F |
| Naa10 | -0.724956 | 6.01530215 | 8.80219431 | 0.00425 | 0.015185 | F | F | F | T |
| Pcdhac2 | -4.5071142 | 0.51664773 | 8.8009205 | 0.00425 | 0.015192 | T | F | T | F |
| Taf4b | 0.8393821 | 4.79669881 | 8.79397422 | 0.00426 | 0.01524 | F | F | T | F |
| Nupl2 | -1.5562134 | 3.03290601 | 8.79333801 | 0.00426 | 0.015242 | T | F | F | T |
| Pdlim3 | -3.4472173 | 0.91813605 | 8.79099104 | 0.00427 | 0.015257 | T | F | F | T |
| Th | 3.64008754 | 0.14854823 | 8.79016842 | 0.00427 | 0.01526 | T | T | F | T |
| Sf3b6 | -0.7887999 | 5.42834262 | 8.78826101 | 0.00427 | 0.015272 | T | F | T | F |
| Wipi2 | -0.6065329 | 7.18381711 | 8.77974002 | 0.00429 | 0.015332 | T | F | T | F |
| Mtmr14 | -0.7453575 | 5.79374679 | 8.77171513 | 0.00431 | 0.015389 | F | F | F | F |
| Drc1 | -2.8543385 | 1.30208053 | 8.76976267 | 0.00431 | 0.015401 | T | F | F | F |
| Ankfn1 | -7.4122436 | 0.05663559 | 8.75855196 | 0.00434 | 0.015481 | T | F | T | F |
| Mettl25 | 1.20078746 | 3.46560763 | 8.75785178 | 0.00434 | 0.015484 | T | F | F | F |
| Zfp37 | -1.172031 | 3.89842223 | 8.7568011 | 0.00434 | 0.015489 | F | F | F | F |
| Grem1 | -7.4152628 | 0.06426249 | 8.75547915 | 0.00434 | 0.015496 | T | F | F | T |
| Gm14685 | 1.22677391 | 3.38548134 | 8.7477451 | 0.00436 | 0.015552 | F | F | F | F |
| Farsb | 0.60636221 | 6.83621397 | 8.74457349 | 0.00437 | 0.015573 | T | T | T | F |
| Pcnx4 | -1.1220983 | 4.01924795 | 8.74380378 | 0.00437 | 0.015576 | T | F | T | F |
| Trim35 | -0.6610813 | 6.66414079 | 8.73820215 | 0.00438 | 0.015615 | T | F | T | T |
| Ufl1 | 0.78227326 | 5.10008352 | 8.73590368 | 0.00438 | 0.01563 | F | F | T | F |
| Slc16a1 | -0.5013327 | 9.78678483 | 8.73450406 | 0.00439 | 0.015638 | T | F | T | T |
| Dcakd | -0.5731699 | 7.82273132 | 8.72826303 | 0.0044 | 0.015682 | F | F | F | F |
| Cryz | -1.108287 | 4.05756099 | 8.72607185 | 0.0044 | 0.015696 | T | F | F | T |
| Ahr | -1.4308582 | 3.26196899 | 8.72447592 | 0.00441 | 0.015706 | T | F | T | T |
| Samsn1 | 4.65234453 | -0.3644346 | 8.72374121 | 0.00441 | 0.015709 | T | F | F | F |
| ccdc198 | -7.4067468 | 0.05102236 | 8.72236624 | 0.00441 | 0.015717 | F | F | F | F |

| | | | | | | | | | |
|---|---|---|---|---|---|---|---|---|---|
| Bbs2 | -1.7742051 | 2.62107546 | 8.72095535 | 0.00442 | 0.015725 | F | F | T | F |
| Rab12 | 0.89920803 | 4.44296736 | 8.71919987 | 0.00442 | 0.015735 | F | F | T | T |
| Tgs1 | 0.64510659 | 6.3887405 | 8.71838402 | 0.00442 | 0.015739 | T | F | T | F |
| Hes7 | -1.3755663 | 3.52673184 | 8.71386009 | 0.00443 | 0.015771 | T | F | F | T |
| Lsm8 | -0.863446 | 4.96107026 | 8.71091604 | 0.00444 | 0.01579 | T | F | F | T |
| Armc4 | -7.4263067 | 0.03204687 | 8.7106204 | 0.00444 | 0.01579 | F | F | F | T |
| Prr23a3 | 2.17785938 | 1.47602801 | 8.7073772 | 0.00444 | 0.015812 | F | F | F | F |
| Kpna4 | 0.56641716 | 7.4917079 | 8.70199675 | 0.00446 | 0.015851 | T | F | T | T |
| Mief1 | 0.65823264 | 6.2800048 | 8.70007501 | 0.00446 | 0.015863 | T | F | T | F |
| Tmed10 | 0.58152117 | 7.19546082 | 8.69820743 | 0.00446 | 0.015874 | T | F | T | F |
| Ywhaz | 0.48243156 | 9.75066118 | 8.69265177 | 0.00448 | 0.015913 | T | F | T | T |
| Gm21750 | 1.42137736 | 2.88390616 | 8.69246503 | 0.00448 | 0.015913 | F | F | F | F |
| Kcnh4 | -7.4156038 | 0.02826677 | 8.68628291 | 0.00449 | 0.015958 | F | F | F | F |
| Abca5 | 1.71171466 | 2.25864416 | 8.68129559 | 0.0045 | 0.015994 | T | F | T | T |
| Cep70 | 1.08642266 | 3.8077761 | 8.67807921 | 0.00451 | 0.016016 | F | F | F | F |
| Dcps | -0.7879677 | 5.39485039 | 8.67209028 | 0.00452 | 0.016059 | F | F | F | T |
| Mboat7 | -0.7579543 | 5.63170394 | 8.66779713 | 0.00453 | 0.01609 | T | F | F | F |
| Ccdc82 | 0.6894107 | 6.12396324 | 8.66343791 | 0.00454 | 0.016121 | T | T | T | T |
| Usp31 | 0.61195204 | 6.8255727 | 8.66179304 | 0.00454 | 0.016131 | F | F | T | T |
| Rsrp1 | -0.767984 | 5.60518971 | 8.66117486 | 0.00454 | 0.016133 | F | F | F | F |
| Fam110b | -1.6067591 | 2.90507932 | 8.65661427 | 0.00455 | 0.016166 | T | F | T | F |
| Dnajc7 | -0.6449709 | 6.71564323 | 8.65173473 | 0.00456 | 0.016202 | F | F | F | F |
| Fgf14 | -7.4441757 | 0.03338927 | 8.65054107 | 0.00457 | 0.016208 | T | F | T | T |
| Dmrt2 | -7.4075098 | 0.02388189 | 8.64991154 | 0.00457 | 0.01621 | F | F | F | T |
| Tdrp | -1.0914958 | 4.08153072 | 8.64460247 | 0.00458 | 0.016249 | F | F | F | F |
| Lrrc23 | -7.4158389 | 0.02369992 | 8.64424377 | 0.00458 | 0.016249 | F | F | T | F |
| Cnbp | 0.5116354 | 8.85664651 | 8.64337457 | 0.00458 | 0.016252 | F | F | F | T |
| 2810021J22 | -1.3080953 | 3.52821201 | 8.64319447 | 0.00458 | 0.016252 | T | F | F | T |
| 1810058I24 | -0.8972 | 4.8119227 | 8.64029521 | 0.00459 | 0.016272 | T | F | F | F |
| Zfp451 | 0.60491368 | 6.89154055 | 8.63787814 | 0.0046 | 0.016289 | T | F | T | T |
| Dda1 | 0.69460092 | 5.88870411 | 8.63649657 | 0.0046 | 0.016297 | F | F | T | F |
| Cradd | -2.1753245 | 1.96544416 | 8.63345858 | 0.00461 | 0.016318 | F | F | F | F |
| Stk36 | -1.5688445 | 2.99417894 | 8.63225677 | 0.00461 | 0.016323 | F | F | T | F |
| Rnaset2a | 0.7662605 | 5.46210319 | 8.63221799 | 0.00461 | 0.016323 | T | F | F | F |
| Dnah7b | -1.5803763 | 2.9791131 | 8.63004478 | 0.00461 | 0.016337 | F | F | F | F |
| Unc79 | -3.0011284 | 1.18452522 | 8.62622042 | 0.00462 | 0.016364 | F | F | F | F |
| Gapvd1 | 0.56545832 | 7.3567642 | 8.61701648 | 0.00464 | 0.016435 | F | F | F | T |
| 5730455P1( | -0.9747443 | 4.44100599 | 8.61501447 | 0.00465 | 0.016448 | T | F | T | T |
| Olfr509 | 0.89100142 | 4.51049249 | 8.61153424 | 0.00465 | 0.016473 | F | F | F | F |
| Tex9 | -1.1249589 | 4.05615992 | 8.60356407 | 0.00467 | 0.016533 | F | F | T | T |
| Hoxd3 | -7.4560479 | 0.12097116 | 8.60131963 | 0.00468 | 0.016549 | T | T | T | T |
| Npnt | -2.2528845 | 1.93779133 | 8.59726534 | 0.00469 | 0.016578 | F | F | T | T |

| | | | | | | | | | |
|---|---|---|---|---|---|---|---|---|---|
| Rpl11 | -0.5534338 | 8.49688668 | 8.59040814 | 0.0047 | 0.01663 | F | F | T | F |
| Plin5 | -2.9759375 | 1.17130423 | 8.58856736 | 0.00471 | 0.016643 | F | F | T | F |
| Fbxo5 | -0.7137666 | 6.01025304 | 8.58765799 | 0.00471 | 0.016647 | F | F | T | F |
| Senp8 | -1.2052486 | 3.77400561 | 8.58424218 | 0.00472 | 0.016672 | F | F | T | T |
| Ntng1 | -7.5124027 | 0.17476478 | 8.58378349 | 0.00472 | 0.016673 | T | T | T | T |
| Ssna1 | -1.1985907 | 3.82928048 | 8.58235615 | 0.00472 | 0.01668 | F | F | T | F |
| Gm10715 | 7.31993041 | -1.0408152 | 8.59242807 | 0.00472 | 0.01668 | F | F | F | F |
| Ppp4r4 | -1.9694983 | 2.30475664 | 8.58054549 | 0.00472 | 0.016691 | T | F | T | T |
| Grk1 | -3.8668398 | 0.69417907 | 8.57920569 | 0.00473 | 0.016699 | T | F | T | T |
| Ltv1 | 0.66607485 | 6.11865248 | 8.575315 | 0.00474 | 0.016727 | T | F | F | F |
| Atxn7 | 0.82382675 | 4.80112879 | 8.56860113 | 0.00475 | 0.016779 | F | F | T | T |
| Prr19 | -7.4120577 | 0.01569557 | 8.56748889 | 0.00475 | 0.016785 | F | F | F | F |
| Mxra8 | -1.1703456 | 3.86641903 | 8.55988405 | 0.00477 | 0.016844 | F | F | F | T |
| Cep41 | -1.7401638 | 2.68339146 | 8.54810881 | 0.0048 | 0.016938 | T | F | T | T |
| Hyal1 | -2.3428207 | 1.75644277 | 8.54659748 | 0.0048 | 0.016947 | T | F | F | F |
| C77080 | -0.7250835 | 5.8827764 | 8.54430066 | 0.00481 | 0.016963 | F | F | F | F |
| Hnrnpc | -0.5210736 | 9.01180349 | 8.54383839 | 0.00481 | 0.016964 | F | F | T | F |
| Elavl4 | -7.399673 | 0.01045627 | 8.53859486 | 0.00482 | 0.017004 | T | F | T | T |
| Pdgfrl | -7.3788927 | 0.02641154 | 8.52900327 | 0.00484 | 0.017081 | F | F | F | F |
| Dgkb | -7.4596622 | 0.03518383 | 8.524936 | 0.00485 | 0.017111 | T | T | T | T |
| Aebp2 | 0.6202747 | 6.64473895 | 8.51589256 | 0.00487 | 0.017184 | T | F | T | T |
| Ece2 | 1.40366714 | 2.89610988 | 8.51514339 | 0.00488 | 0.017184 | T | F | T | F |
| Srrm4 | -1.5065276 | 3.12135888 | 8.5151165 | 0.00488 | 0.017184 | T | F | T | T |
| Gpatch8 | -0.680668 | 6.33177237 | 8.50690582 | 0.0049 | 0.01725 | F | F | F | F |
| Usp33 | 0.69245541 | 5.7678002 | 8.50159497 | 0.00491 | 0.017291 | T | F | T | F |
| Tbc1d8 | 1.94489296 | 1.82319381 | 8.49599099 | 0.00492 | 0.017336 | T | F | F | T |
| Gm11032 | 0.77335543 | 5.69319119 | 8.48967326 | 0.00494 | 0.017386 | F | F | F | F |
| Ulk2 | 0.98900792 | 4.08524008 | 8.48746996 | 0.00494 | 0.017401 | T | F | T | T |
| A630010A( | 2.63572199 | 0.78966786 | 8.48287634 | 0.00495 | 0.017437 | F | F | F | F |
| Ntrk3 | -1.3267877 | 3.49103982 | 8.48262127 | 0.00495 | 0.017437 | T | F | T | T |
| Cutal | -7.37405 | 0.02695307 | 8.4793825 | 0.00496 | 0.017461 | T | F | T | T |
| Cbx3 | -0.5567793 | 8.07300577 | 8.47521814 | 0.00497 | 0.017494 | T | F | T | F |
| Dnajc17 | -1.2697599 | 3.58322299 | 8.47447022 | 0.00497 | 0.017497 | F | F | F | F |
| Cyb561d2 | 1.20959438 | 3.38413742 | 8.47278329 | 0.00498 | 0.017508 | F | F | T | F |
| Cd2bp2 | 0.59618904 | 6.84508769 | 8.47173771 | 0.00498 | 0.017514 | F | F | T | F |
| Snx5 | -0.545093 | 8.28209336 | 8.471474 | 0.00498 | 0.017514 | T | F | F | T |
| Ntn4 | -7.3706281 | 0.00747775 | 8.46862075 | 0.00499 | 0.017535 | F | F | F | F |
| Pla2g12a | 0.83093486 | 4.73723804 | 8.46365423 | 0.005 | 0.017575 | F | F | T | F |
| Olfr393 | 1.0404437 | 4.03600088 | 8.46086194 | 0.00501 | 0.017595 | F | F | F | F |
| Zfp947 | 2.32009621 | 1.22696556 | 8.46038283 | 0.00501 | 0.017597 | T | F | F | F |
| Cep170 | 0.66356543 | 6.0784312 | 8.45572377 | 0.00502 | 0.017634 | T | F | T | T |
| Zfp60 | -0.7567114 | 5.65267087 | 8.45533606 | 0.00502 | 0.017634 | T | T | F | T |

| | | | | | | | | | |
|---|---|---|---|---|---|---|---|---|---|
| Zfp850 | -0.9155401 | 4.65907545 | 8.45305636 | 0.00502 | 0.017651 | F | F | T | T |
| Cd93 | 1.30001732 | 3.14027817 | 8.45186425 | 0.00503 | 0.017658 | T | F | F | T |
| Gin1 | 1.38868686 | 2.9159787 | 8.44838811 | 0.00504 | 0.017685 | T | T | T | T |
| Tbc1d24 | 0.86318444 | 4.57269647 | 8.44416741 | 0.00505 | 0.017718 | T | F | T | T |
| Otop3 | -7.4247271 | 0.01182761 | 8.44026702 | 0.00506 | 0.017749 | F | F | F | F |
| BC035947 | -7.499437 | 0.05932949 | 8.43925184 | 0.00506 | 0.017754 | T | F | F | F |
| Vnn1 | -7.3777688 | 0.03905 | 8.43591422 | 0.00507 | 0.01778 | F | F | F | F |
| Slc24a4 | -3.782995 | 0.71832155 | 8.43386883 | 0.00507 | 0.017795 | F | F | T | T |
| Ptger2 | 4.50439655 | -0.3441523 | 8.43085058 | 0.00508 | 0.017818 | T | F | F | T |
| Rrp1b | -0.6613129 | 6.59394341 | 8.42917693 | 0.00508 | 0.01783 | F | F | T | F |
| Ufm1 | 0.72115195 | 5.49003618 | 8.42497391 | 0.00509 | 0.017863 | T | F | F | T |
| Vmn1r18 | 0.95399385 | 4.29176355 | 8.42444261 | 0.00509 | 0.017865 | F | F | F | T |
| Rela | -0.8577577 | 4.90723241 | 8.42150749 | 0.0051 | 0.017887 | F | F | F | T |
| Pcsk1n | -1.1468152 | 3.91706434 | 8.41974198 | 0.00511 | 0.0179 | F | F | T | F |
| Rbm20 | 1.10163301 | 3.76205322 | 8.41900563 | 0.00511 | 0.017903 | T | F | T | T |
| Dbt | 0.72042282 | 5.46306296 | 8.41041955 | 0.00513 | 0.017975 | T | F | F | T |
| Hesx1 | -2.3066067 | 1.88564232 | 8.40788631 | 0.00514 | 0.017994 | F | F | F | F |
| Fam161b | 1.1935584 | 3.45073345 | 8.4067751 | 0.00514 | 0.018001 | T | F | F | T |
| Gcat | -0.7085393 | 6.00240959 | 8.40504839 | 0.00514 | 0.018013 | F | F | T | T |
| Pafah1b3 | -0.8662397 | 4.8605537 | 8.40329283 | 0.00515 | 0.018024 | F | F | F | F |
| Ermap | 1.71964986 | 2.16098392 | 8.40314361 | 0.00515 | 0.018024 | T | F | T | T |
| Pabpc4l | -2.0529382 | 2.0998439 | 8.4022575 | 0.00515 | 0.018027 | F | T | T | T |
| Ccnh | 0.76872825 | 5.10100677 | 8.40214805 | 0.00515 | 0.018027 | T | F | F | F |
| Zfp760 | -1.3369969 | 3.42289571 | 8.40179003 | 0.00515 | 0.018027 | F | F | T | F |
| Sgsh | 1.26149708 | 3.74591767 | 8.39734883 | 0.00516 | 0.018061 | F | F | F | F |
| Abcg1 | -1.5477433 | 3.01626287 | 8.39718212 | 0.00516 | 0.018061 | F | F | T | F |
| Hist1h2ap | -0.5080945 | 9.64958237 | 8.38802863 | 0.00518 | 0.018139 | F | F | F | F |
| H3c7 | -0.5232521 | 9.09278085 | 8.38745436 | 0.00519 | 0.018141 | F | F | F | T |
| Ahi1 | -1.1827659 | 3.81291338 | 8.38667111 | 0.00519 | 0.018145 | T | F | T | F |
| Snrpd3 | -0.6930075 | 6.32464822 | 8.386375 | 0.00519 | 0.018145 | F | F | F | T |
| Pde1b | -1.9081527 | 2.33169409 | 8.38566746 | 0.00519 | 0.018148 | T | F | T | T |
| Kcnq5 | -3.776126 | 0.69628503 | 8.3788031 | 0.00521 | 0.018205 | F | F | T | T |
| Rgn | -7.3921612 | -0.0060577 | 8.3771464 | 0.00521 | 0.018217 | T | F | T | T |
| Fuca1 | 0.99711089 | 4.05002359 | 8.37665007 | 0.00521 | 0.018217 | T | F | F | T |
| Gm2956 | -7.5538787 | 0.21298605 | 8.37641472 | 0.00521 | 0.018217 | F | F | F | F |
| C330021F2 | -7.3697148 | -0.0101921 | 8.37613218 | 0.00521 | 0.018217 | F | F | T | F |
| Gigyf1 | -0.6261326 | 6.93705884 | 8.37403139 | 0.00522 | 0.018233 | T | F | T | T |
| Srr | 1.83323174 | 1.91548723 | 8.3718992 | 0.00523 | 0.018249 | T | F | T | F |
| Chtf18 | -0.7364284 | 5.74979925 | 8.36799628 | 0.00524 | 0.018278 | F | F | F | F |
| Riox2 | 0.68851052 | 5.75019015 | 8.36797841 | 0.00524 | 0.018278 | F | F | T | F |
| Mfsd5 | 0.70798578 | 5.64069728 | 8.36595265 | 0.00524 | 0.018293 | F | F | F | T |
| Zfp385b | -3.4026993 | 0.89866171 | 8.36384071 | 0.00525 | 0.018306 | F | F | T | T |

| | | | | | | | | | |
|---|---|---|---|---|---|---|---|---|---|
| Cep55 | -1.0877209 | 4.03693048 | 8.36383191 | 0.00525 | 0.018306 | T | T | F | T |
| Ppp1cc | -0.5400977 | 8.32431839 | 8.36208041 | 0.00525 | 0.018318 | F | F | T | F |
| Tceal3 | -7.3538289 | -0.0030203 | 8.35998398 | 0.00526 | 0.018334 | F | F | T | F |
| Sdhaf2 | 0.79083127 | 4.99191535 | 8.34946248 | 0.00528 | 0.018425 | T | F | F | T |
| Mlx | 0.87187032 | 4.48928953 | 8.34608838 | 0.00529 | 0.018452 | T | T | T | T |
| H3c6 | -0.512553 | 9.34161375 | 8.34015688 | 0.00531 | 0.018502 | F | F | F | F |
| Tmem198 | 1.95193262 | 1.70801926 | 8.33723579 | 0.00531 | 0.018525 | F | F | F | F |
| Nyap1 | -1.3188449 | 3.46669858 | 8.33584452 | 0.00532 | 0.018534 | T | F | F | F |
| Hsf2 | -0.8255973 | 5.07083309 | 8.33565543 | 0.00532 | 0.018534 | T | F | F | T |
| Hdac3 | -0.6937629 | 6.10266653 | 8.33111081 | 0.00533 | 0.018572 | T | F | F | T |
| Rbm11 | 2.1414912 | 1.36348901 | 8.32765097 | 0.00534 | 0.0186 | T | F | F | T |
| Nr3c1 | 1.11550411 | 3.67884809 | 8.32080911 | 0.00536 | 0.018659 | T | F | T | T |
| Immp1l | -1.4740309 | 3.09621394 | 8.31643163 | 0.00537 | 0.018695 | T | T | F | T |
| Cbfa2t3 | 1.23714599 | 3.47394141 | 8.31521933 | 0.00537 | 0.018703 | F | F | T | F |
| Cage1 | 1.58573593 | 2.44886436 | 8.31266227 | 0.00538 | 0.018724 | F | F | F | F |
| Sh3rf1 | -0.7130038 | 5.89475355 | 8.31058417 | 0.00538 | 0.018739 | T | F | T | F |
| Pde4b | -2.2923297 | 1.75185943 | 8.30831474 | 0.00539 | 0.018757 | T | F | F | F |
| Brdt | 3.07648299 | 0.41520977 | 8.30606762 | 0.0054 | 0.018775 | T | F | T | T |
| Vmn1r53 | 5.20096651 | -0.6524495 | 8.29928646 | 0.00541 | 0.018833 | F | F | F | F |
| Net1 | 0.5597963 | 7.25401155 | 8.29843608 | 0.00542 | 0.018838 | T | F | T | F |
| Zbtb1 | 0.7025025 | 5.69895199 | 8.29796697 | 0.00542 | 0.018839 | T | F | F | T |
| Epb41l2 | -0.6181527 | 6.84692256 | 8.2919067 | 0.00543 | 0.018892 | F | F | F | F |
| Prps1l3 | 0.64702379 | 6.22607553 | 8.28937242 | 0.00544 | 0.018912 | F | F | F | F |
| Dipk1c | -7.3433151 | -0.0131378 | 8.28681325 | 0.00545 | 0.018933 | F | F | F | F |
| Tex26 | -7.3503534 | -0.0206182 | 8.28515348 | 0.00545 | 0.018945 | F | F | F | F |
| Nlrp3 | 7.27113506 | -1.0785085 | 8.29129238 | 0.00546 | 0.018973 | F | F | F | F |
| Cspg4 | -7.4118851 | -0.0036047 | 8.27577693 | 0.00548 | 0.019025 | T | F | F | F |
| H3c2 | -0.5123012 | 9.27000556 | 8.27304576 | 0.00548 | 0.019048 | F | F | F | T |
| Ttc16 | -4.4643352 | 0.4443468 | 8.26408763 | 0.00551 | 0.019128 | T | F | F | F |
| Actr5 | -0.9615815 | 4.37530276 | 8.26181442 | 0.00551 | 0.019146 | T | F | F | T |
| Csrp2 | -1.0252496 | 4.36608851 | 8.2612773 | 0.00551 | 0.019148 | T | F | F | T |
| Fam122a | -1.3150738 | 3.42551875 | 8.25894711 | 0.00552 | 0.019166 | T | T | T | T |
| Slc36a1 | 0.73933188 | 5.3287165 | 8.25854024 | 0.00552 | 0.019167 | T | F | T | T |
| Pdpr | 0.65653783 | 6.08241482 | 8.25644649 | 0.00553 | 0.019183 | F | F | F | T |
| Gdpd2 | -7.338358 | -0.0092943 | 8.25369997 | 0.00553 | 0.019206 | F | F | F | F |
| Nudcd2 | -0.7346625 | 5.68576153 | 8.25327204 | 0.00554 | 0.019207 | T | T | F | T |
| Tmem87b | 0.78556043 | 4.94934481 | 8.25043374 | 0.00554 | 0.01923 | T | F | F | F |
| Glrx | -0.882849 | 4.73038752 | 8.24509537 | 0.00556 | 0.019277 | T | F | F | T |
| Ccdc24 | -7.3736395 | -0.023298 | 8.23771684 | 0.00558 | 0.019343 | F | F | T | F |
| Hnrnpll | -0.6871005 | 6.26855732 | 8.2281213 | 0.0056 | 0.019431 | T | F | F | F |
| Gja4 | 2.10057509 | 1.44965246 | 8.22561411 | 0.00561 | 0.019451 | F | F | F | F |
| Retreg1 | -1.6411055 | 2.79138686 | 8.22481514 | 0.00561 | 0.019456 | T | F | T | F |

| | | | | | | | | | |
|---|---|---|---|---|---|---|---|---|---|
| Dhrs4 | -1.0860948 | 4.01343241 | 8.22444064 | 0.00561 | 0.019456 | F | F | F | F |
| Slc1a6 | -7.3328526 | -0.0156526 | 8.21858855 | 0.00563 | 0.019509 | F | F | F | F |
| Ptchd1 | -7.4131593 | -0.0056474 | 8.21809065 | 0.00563 | 0.01951 | T | T | T | T |
| Chn1 | -1.3056321 | 3.43284232 | 8.21547461 | 0.00564 | 0.019532 | F | F | T | F |
| Tcf25 | 0.53439325 | 7.76577655 | 8.21330278 | 0.00564 | 0.019549 | T | F | T | F |
| Birc3 | 0.8996314 | 4.4488146 | 8.21303601 | 0.00564 | 0.019549 | F | F | F | T |
| Ier5l | -0.9085963 | 4.61921994 | 8.21188716 | 0.00565 | 0.019557 | F | F | F | F |
| Nemf | -0.7600771 | 5.50776028 | 8.2064834 | 0.00566 | 0.019605 | F | F | F | F |
| Ccdc126 | 1.73846259 | 2.06781417 | 8.20192903 | 0.00568 | 0.019645 | T | F | T | F |
| Crhr2 | -7.353973 | -0.0317762 | 8.20058876 | 0.00568 | 0.019655 | F | F | T | F |
| Abi3 | -7.3310004 | -0.0102751 | 8.18990678 | 0.00571 | 0.019753 | F | F | T | F |
| Tlk1 | -0.6127144 | 6.90271197 | 8.18977969 | 0.00571 | 0.019753 | T | F | F | T |
| H2ac8 | -0.5519474 | 7.90411744 | 8.18244644 | 0.00573 | 0.01982 | F | F | F | F |
| Rnf121 | 0.73435588 | 5.24607065 | 8.17605228 | 0.00575 | 0.019879 | T | F | T | F |
| Cpne1 | -0.6699845 | 6.27372349 | 8.17522378 | 0.00575 | 0.019884 | T | F | T | F |
| Ybx1 | -0.4846235 | 9.84013668 | 8.16853721 | 0.00577 | 0.019945 | T | F | F | T |
| Rps27 | -1.783225 | 2.56591391 | 8.16693892 | 0.00577 | 0.019958 | T | F | F | F |
| Psmb6 | -0.5932637 | 7.10818427 | 8.15891791 | 0.0058 | 0.020033 | F | F | T | F |
| Defa30 | -7.3243501 | -0.0200524 | 8.15770851 | 0.0058 | 0.020041 | F | F | F | F |
| Grina | 1.0087361 | 3.96490833 | 8.15740182 | 0.0058 | 0.020041 | F | F | F | F |
| Gpr1 | 3.62915639 | 0.00266112 | 8.15702006 | 0.0058 | 0.020041 | F | F | T | T |
| Srsf4 | -0.6280935 | 6.69611439 | 8.15641656 | 0.0058 | 0.020044 | F | F | F | T |
| Gfod2 | -0.9181144 | 4.54487332 | 8.15473859 | 0.00581 | 0.020057 | F | F | F | T |
| Neurl2 | 1.89381181 | 1.72901528 | 8.15447674 | 0.00581 | 0.020057 | F | F | F | F |
| Tfcp2 | -1.2833741 | 3.48936883 | 8.15273525 | 0.00581 | 0.020071 | F | F | T | T |
| Pif1 | 0.86701181 | 4.46057085 | 8.14665224 | 0.00583 | 0.020127 | F | F | T | F |
| Zbbx | -7.3671525 | -0.0327792 | 8.14372756 | 0.00584 | 0.020152 | F | F | F | F |
| Vmn2r117 | 1.6640586 | 2.21468173 | 8.1432706 | 0.00584 | 0.020154 | F | F | F | F |
| Pgd | 0.5312078 | 7.79342554 | 8.14209993 | 0.00584 | 0.020162 | F | F | F | F |
| Asic2 | -3.4092717 | 0.91394433 | 8.14035257 | 0.00585 | 0.020176 | F | F | F | F |
| Hexb | -1.2367444 | 3.61391831 | 8.13989751 | 0.00585 | 0.020177 | F | F | F | F |
| Zfp595 | 1.55018661 | 2.48482754 | 8.1373097 | 0.00586 | 0.0202 | T | F | T | T |
| Rhoc | 0.73737729 | 5.21692952 | 8.13598564 | 0.00586 | 0.020209 | F | F | F | F |
| Magi2 | -2.9070757 | 1.14809921 | 8.13518191 | 0.00586 | 0.020214 | F | F | T | T |
| Mak | -2.6682315 | 1.3440339 | 8.1312534 | 0.00587 | 0.02025 | F | F | T | F |
| Lmbrd2 | 0.67535672 | 5.90670037 | 8.13030329 | 0.00588 | 0.020256 | T | F | F | T |
| Xlr5c | 2.81465654 | 0.76879707 | 8.1298288 | 0.00588 | 0.020256 | F | F | F | F |
| Vezf1 | -0.6833465 | 6.13824904 | 8.12963113 | 0.00588 | 0.020256 | T | F | T | T |
| Neil2 | -3.7790718 | 0.68148909 | 8.12809045 | 0.00588 | 0.020268 | F | F | F | T |
| Dhx30 | -0.6254798 | 6.81032207 | 8.12728918 | 0.00589 | 0.020273 | F | F | F | F |
| 3110009E1 | -2.6672145 | 1.34306638 | 8.1199406 | 0.00591 | 0.020342 | F | F | F | F |
| Ccdc66 | 0.96840777 | 4.12313856 | 8.11837686 | 0.00591 | 0.020354 | T | F | T | F |

| | | | | | | | | | |
|---|---|---|---|---|---|---|---|---|---|
| H60b | -7.3251114 | -0.0418552 | 8.11751953 | 0.00591 | 0.02036 | F | F | F | F |
| Bloc1s5 | 1.01900902 | 3.93807426 | 8.11327109 | 0.00593 | 0.020399 | F | F | T | F |
| Ccdc85a | -7.4579747 | 0.02096737 | 8.11097723 | 0.00593 | 0.020418 | F | T | T | T |
| Rpap1 | -0.6598308 | 6.34105563 | 8.10635471 | 0.00595 | 0.020461 | F | F | F | F |
| Trim29 | 3.56510972 | 0.08954522 | 8.10550078 | 0.00595 | 0.020466 | F | F | T | F |
| Pcdhgb6 | -2.4394186 | 1.54838357 | 8.10485366 | 0.00595 | 0.02047 | F | F | F | F |
| Slc23a3 | -3.7388645 | 0.67209279 | 8.10149133 | 0.00596 | 0.0205 | F | F | F | F |
| Gbgt1 | -7.3382787 | -0.0463559 | 8.08610987 | 0.006 | 0.020651 | F | F | F | F |
| Ap4s1 | 1.14520554 | 3.51174775 | 8.07636746 | 0.00603 | 0.020746 | F | F | F | T |
| Tcp11 | -7.4024901 | -0.0181841 | 8.07591072 | 0.00603 | 0.020748 | F | F | F | F |
| Prr11 | -0.7816483 | 5.21765294 | 8.07456883 | 0.00604 | 0.020758 | T | F | T | F |
| Gca | 1.46375538 | 2.77457446 | 8.07371849 | 0.00604 | 0.020763 | F | F | T | F |
| Rfc3 | -0.7677501 | 5.32350763 | 8.07205963 | 0.00605 | 0.020777 | F | F | F | F |
| Hes6 | -1.3783093 | 3.2424395 | 8.0707395 | 0.00605 | 0.020787 | F | F | T | F |
| Rab22a | 0.68932631 | 5.62450342 | 8.05919579 | 0.00608 | 0.020901 | F | F | T | F |
| Fbxw7 | 0.65597506 | 6.01620026 | 8.0559924 | 0.00609 | 0.020931 | F | F | T | T |
| Hmgcs1 | 0.58148913 | 6.89955771 | 8.05542806 | 0.0061 | 0.020933 | T | F | F | F |
| Ppat | -0.5978949 | 7.03648868 | 8.05380643 | 0.0061 | 0.020946 | T | F | F | F |
| Cdc42ep2 | 1.07419998 | 3.75865061 | 8.03647318 | 0.00615 | 0.021121 | F | F | F | F |
| Mip | -7.3078469 | -0.0508865 | 8.02914661 | 0.00617 | 0.021194 | F | F | F | F |
| Rps18 | -0.5916277 | 7.06381559 | 8.0269285 | 0.00618 | 0.021213 | F | F | F | F |
| 2610301B2( | -1.4150849 | 3.15142646 | 8.01897162 | 0.00621 | 0.021292 | T | F | T | T |
| Flii | 0.57703376 | 6.89199003 | 8.01801598 | 0.00621 | 0.021299 | F | F | F | F |
| Ostm1 | 0.95811604 | 4.10071548 | 8.01472515 | 0.00622 | 0.02133 | T | F | T | T |
| Sike1 | 0.70528798 | 5.44645887 | 8.00972143 | 0.00623 | 0.021379 | T | F | T | T |
| Arhgap11a | 0.57938829 | 6.84933287 | 8.00870481 | 0.00624 | 0.021386 | T | F | F | T |
| Necab3 | -3.1337245 | 0.96657745 | 8.00787981 | 0.00624 | 0.021391 | T | F | F | T |
| Khsrp | -0.4886419 | 9.53670684 | 8.00762186 | 0.00624 | 0.021391 | F | F | T | F |
| Fbxo33 | 0.80849327 | 4.73792658 | 8.00613301 | 0.00624 | 0.021403 | T | F | T | F |
| Acsl4 | 0.61762536 | 6.64085811 | 8.00479842 | 0.00625 | 0.021413 | T | F | F | F |
| Foxa3 | -1.5976009 | 2.79419831 | 8.00405735 | 0.00625 | 0.021418 | T | F | T | F |
| Lrrc71 | 5.40658413 | -0.7648994 | 8.00180022 | 0.00626 | 0.021438 | F | F | T | F |
| Pgam1 | -0.4770766 | 9.84244567 | 8.00067404 | 0.00626 | 0.021446 | F | F | F | F |
| Gm4724 | -7.3621533 | 0.04082218 | 8.00006816 | 0.00626 | 0.021449 | T | F | T | T |
| Aunip | -1.0842814 | 3.97697642 | 7.99815919 | 0.00627 | 0.021466 | F | F | F | F |
| Pisd | 0.67700122 | 5.72710223 | 7.99176106 | 0.00629 | 0.021527 | F | F | F | F |
| Lamc2 | -2.286222 | 1.75313857 | 7.99170456 | 0.00629 | 0.021527 | T | F | F | F |
| Sec62 | 0.61636165 | 6.40722059 | 7.99031566 | 0.00629 | 0.021538 | T | F | F | T |
| Dnaaf1 | -7.3125506 | -0.0599544 | 7.98829521 | 0.0063 | 0.021556 | T | F | T | F |
| Foxf2 | -2.6117082 | 1.36989551 | 7.98032826 | 0.00632 | 0.021637 | F | F | F | F |
| Vmn2r124 | -7.3109992 | -0.0159643 | 7.97468797 | 0.00634 | 0.021693 | F | F | F | F |
| Trp53rkb | 1.05643981 | 3.7620523 | 7.97185424 | 0.00635 | 0.02172 | T | F | T | T |

| | | | | | | | | | |
|---|---|---|---|---|---|---|---|---|---|
| Pgr | -1.934719 | 2.21879053 | 7.96294368 | 0.00638 | 0.021811 | T | F | T | T |
| Insyn2b | -7.2945193 | -0.0464373 | 7.9626654 | 0.00638 | 0.021811 | T | F | F | F |
| Appl1 | 0.64622965 | 6.10797646 | 7.96027684 | 0.00639 | 0.021833 | T | F | T | T |
| Tom1l2 | 0.91999844 | 4.32519614 | 7.95811056 | 0.00639 | 0.021852 | T | F | T | F |
| Lrrc4 | 0.97221086 | 4.03279522 | 7.95783683 | 0.00639 | 0.021852 | T | F | T | T |
| Gsc | -7.2985847 | -0.0332595 | 7.95369504 | 0.00641 | 0.021893 | F | F | F | F |
| U2af2 | -0.527997 | 8.42800334 | 7.95032548 | 0.00642 | 0.021926 | F | F | F | F |
| Atp2b2 | -3.7408557 | 0.65363646 | 7.94804759 | 0.00642 | 0.021947 | T | F | T | F |
| Hoxa4 | -7.3679069 | 0.04775045 | 7.94245857 | 0.00644 | 0.022004 | F | F | F | F |
| Ank1 | -7.2907369 | -0.059626 | 7.9338163 | 0.00647 | 0.022089 | T | F | T | T |
| 9330182L0( | -7.3073565 | -0.0671715 | 7.93366409 | 0.00647 | 0.022089 | T | F | T | F |
| Klk14 | -7.2908661 | -0.0468114 | 7.9335864 | 0.00647 | 0.022089 | F | F | F | T |
| Wwc1 | 0.60099047 | 6.56261239 | 7.9277136 | 0.00649 | 0.022149 | F | F | F | F |
| Ptpru | -1.2325119 | 3.55886868 | 7.92531901 | 0.0065 | 0.022172 | F | F | F | F |
| Sgsm3 | 0.91330853 | 4.2342819 | 7.91839394 | 0.00652 | 0.022244 | F | F | F | F |
| Erlec1 | -1.2645869 | 3.48323215 | 7.91610374 | 0.00653 | 0.022265 | F | F | T | T |
| Gpr37 | 1.10388618 | 3.65339235 | 7.91332827 | 0.00653 | 0.022292 | F | F | T | F |
| Aim2 | -2.4523327 | 1.49368077 | 7.91232185 | 0.00654 | 0.022297 | T | F | T | T |
| 6820408C1: | -7.2951768 | -0.0690272 | 7.91170208 | 0.00654 | 0.022297 | F | F | F | F |
| Spdef | -7.2951768 | -0.0690272 | 7.91170208 | 0.00654 | 0.022297 | F | F | F | F |
| Hsf2bp | 2.30846364 | 1.08770793 | 7.91160166 | 0.00654 | 0.022297 | F | F | T | F |
| Pcnx3 | 0.52743071 | 7.71431527 | 7.91058422 | 0.00654 | 0.022304 | F | T | F | F |
| Ptprn2 | -7.4681634 | 0.13917231 | 7.90994624 | 0.00655 | 0.022308 | T | F | F | F |
| Mafg | 0.58484375 | 6.76608297 | 7.90384325 | 0.00656 | 0.022371 | F | F | T | T |
| Dcx | -2.836722 | 1.20225946 | 7.90062792 | 0.00658 | 0.022403 | T | F | T | T |
| Sgip1 | -7.2859451 | -0.0651884 | 7.89924314 | 0.00658 | 0.022415 | T | F | T | F |
| Tmem106c | -1.5540361 | 2.86003664 | 7.89750597 | 0.00659 | 0.02243 | F | F | T | F |
| Tead3 | 0.91528277 | 4.21631736 | 7.89576101 | 0.00659 | 0.022445 | T | F | T | F |
| Asf1b | 0.82114862 | 4.64648983 | 7.89554035 | 0.00659 | 0.022445 | F | F | F | F |
| Zup1 | 1.21065163 | 3.26168544 | 7.89522126 | 0.00659 | 0.022445 | F | F | T | F |
| Mapt | 2.43534908 | 0.91087875 | 7.88787267 | 0.00662 | 0.022522 | T | F | T | T |
| Zfp473 | 0.81715862 | 4.71256125 | 7.88723201 | 0.00662 | 0.022526 | T | F | T | F |
| Polq | 0.65186571 | 5.96817036 | 7.88289521 | 0.00663 | 0.02257 | F | F | F | F |
| Ppp6r3 | -0.5668483 | 7.35558337 | 7.8809787 | 0.00664 | 0.022588 | T | T | T | T |
| Ewsr1 | -0.511643 | 8.69693303 | 7.8782749 | 0.00665 | 0.022614 | T | F | T | T |
| D6Wsu163e | 0.82784538 | 4.60113415 | 7.87767346 | 0.00665 | 0.022618 | T | F | F | T |
| Cytip | 3.03570461 | 0.35973777 | 7.87720388 | 0.00665 | 0.022619 | T | F | T | F |
| Lias | 0.73770776 | 5.1377778 | 7.87552001 | 0.00666 | 0.022634 | F | F | T | T |
| Dpp8 | 0.63959608 | 6.09812833 | 7.87367074 | 0.00666 | 0.022651 | T | F | T | T |
| Ubtd2 | -0.8288167 | 4.90822069 | 7.87295135 | 0.00667 | 0.022656 | T | F | F | T |
| Cda | 2.31121108 | 1.09154207 | 7.86595487 | 0.00669 | 0.02273 | F | F | F | F |
| Crat | 1.02845319 | 3.84255374 | 7.86278954 | 0.0067 | 0.022762 | T | F | F | F |

| | | | | | | | | | |
|---|---|---|---|---|---|---|---|---|---|
| Rasa1 | 0.62221936 | 6.28853551 | 7.86126859 | 0.0067 | 0.022775 | F | F | F | T |
| Knl1 | 0.58823716 | 6.77711838 | 7.85996731 | 0.00671 | 0.022786 | F | F | F | F |
| Chrnb1 | 2.18901822 | 1.19561686 | 7.85729655 | 0.00672 | 0.022813 | F | F | F | F |
| Cand2 | -0.5768193 | 7.14317298 | 7.85215973 | 0.00673 | 0.022867 | F | F | F | F |
| Cbx8 | 1.41319866 | 2.75079334 | 7.84552823 | 0.00676 | 0.022937 | T | T | F | T |
| Ripply1 | -7.2999393 | -0.0783658 | 7.84526625 | 0.00676 | 0.022937 | F | F | F | F |
| Kat2a | -0.5747084 | 7.11880141 | 7.84142227 | 0.00677 | 0.022977 | F | F | F | F |
| Smarcad1 | 0.59290879 | 6.65870995 | 7.8338206 | 0.00679 | 0.023059 | T | F | T | F |
| Phtf2 | 0.78234756 | 4.88558029 | 7.83293389 | 0.0068 | 0.023065 | T | F | T | T |
| Dnaic1 | -7.2859147 | -0.0808626 | 7.82807011 | 0.00681 | 0.023117 | F | F | F | F |
| Rbm22 | -0.6344443 | 6.48326936 | 7.82505101 | 0.00682 | 0.023148 | T | F | T | F |
| Mob2 | 0.797728 | 4.75678648 | 7.8234391 | 0.00683 | 0.023162 | F | F | T | T |
| Atat1 | -1.1469115 | 3.79288094 | 7.82090246 | 0.00684 | 0.023188 | T | F | T | F |
| Cnp | -1.8826727 | 2.23627125 | 7.81579694 | 0.00685 | 0.023242 | T | F | T | F |
| Nfx1 | 0.52608506 | 7.66078995 | 7.81060576 | 0.00687 | 0.023298 | T | F | T | T |
| Fndc5 | -2.7408468 | 1.21188701 | 7.80802584 | 0.00688 | 0.023324 | F | F | T | F |
| Hspa2 | -2.2993846 | 1.63866003 | 7.79894434 | 0.00691 | 0.023425 | T | F | T | T |
| Klhdc8a | -7.2811897 | -0.04207 | 7.79426007 | 0.00693 | 0.023475 | F | F | T | F |
| Heatr3 | 0.55535421 | 7.04905825 | 7.79382581 | 0.00693 | 0.023476 | T | F | F | F |
| Mesd | 0.60585691 | 6.45121004 | 7.79260542 | 0.00693 | 0.023487 | T | F | T | F |
| Ubr1 | 0.58645033 | 6.78331789 | 7.79053265 | 0.00694 | 0.023507 | T | F | F | T |
| Ppil2 | 0.60943221 | 6.4120525 | 7.78629007 | 0.00696 | 0.023552 | T | F | T | F |
| Gask1a | -7.2870755 | -0.0867208 | 7.78558774 | 0.00696 | 0.023557 | T | F | T | F |
| Cpm | 0.63047816 | 6.22107088 | 7.77885453 | 0.00698 | 0.023631 | T | F | T | T |
| Zfp986 | -7.2700953 | -0.058157 | 7.77568694 | 0.00699 | 0.023662 | F | F | F | F |
| Wdr91 | 0.8669896 | 4.51017044 | 7.77554581 | 0.00699 | 0.023662 | F | F | T | F |
| Lix1l | -1.3512719 | 3.23862851 | 7.77380682 | 0.007 | 0.023679 | F | F | T | F |
| Meltf | 2.90311497 | 0.41927174 | 7.76916457 | 0.00701 | 0.023729 | F | F | T | F |
| Gm49027 | 2.26210395 | 1.09525428 | 7.76768031 | 0.00702 | 0.023743 | F | F | F | F |
| Selenot | 0.5376512 | 7.39128987 | 7.76529874 | 0.00703 | 0.023767 | T | F | T | T |
| Gpi1 | 0.49035695 | 8.62783313 | 7.76464646 | 0.00703 | 0.023771 | F | F | T | F |
| Oas2 | 5.3309562 | -0.7593616 | 7.75909151 | 0.00705 | 0.023832 | T | F | T | T |
| Slc48a1 | -1.9592294 | 2.09439789 | 7.75535443 | 0.00706 | 0.023872 | F | F | T | T |
| 2310022A1 | 0.6931932 | 5.45406337 | 7.75399301 | 0.00707 | 0.023882 | T | F | F | T |
| Fmn2 | -1.2028141 | 3.61259897 | 7.75386066 | 0.00707 | 0.023882 | F | F | F | T |
| Pdcd5 | 0.71350159 | 5.40168443 | 7.74775522 | 0.00709 | 0.023951 | F | F | F | F |
| Hivep2 | -1.9803686 | 2.05531809 | 7.74573777 | 0.0071 | 0.023971 | T | F | F | F |
| Tspan10 | 7.17864819 | -1.1489782 | 7.74887004 | 0.00711 | 0.024025 | T | F | F | T |
| Ifi47 | 7.17864819 | -1.1489782 | 7.74887004 | 0.00711 | 0.024025 | F | F | F | F |
| Gtf3c1 | -0.5233006 | 8.19164696 | 7.73852425 | 0.00712 | 0.024045 | T | F | F | F |
| Magoh | -0.6869688 | 5.90922462 | 7.7348613 | 0.00713 | 0.024084 | F | F | F | F |
| Pcdhgb1 | -7.4344616 | 0.11026978 | 7.73131747 | 0.00715 | 0.024123 | F | F | F | F |

| | | | | | | | | | |
|---|---|---|---|---|---|---|---|---|---|
| Nutf2-ps1 | -0.5938113 | 6.88101633 | 7.72938167 | 0.00715 | 0.024142 | F | F | F | F |
| Tmem18 | 0.91599343 | 4.17086572 | 7.72693113 | 0.00716 | 0.024167 | T | T | T | T |
| Rpl41 | -0.4792961 | 10.0234051 | 7.72419259 | 0.00717 | 0.024196 | F | F | F | F |
| Aen | -0.6219509 | 6.64130047 | 7.72386561 | 0.00717 | 0.024196 | F | F | T | F |
| Tmem51 | -1.1024415 | 3.86844923 | 7.71916616 | 0.00719 | 0.024247 | T | F | F | F |
| Aida | 0.77143344 | 4.87313245 | 7.71894875 | 0.00719 | 0.024247 | T | F | T | T |
| Spata33 | -7.2646624 | -0.0967127 | 7.70614655 | 0.00723 | 0.024397 | F | F | T | F |
| Xrcc3 | -1.0979422 | 3.93892528 | 7.70531337 | 0.00724 | 0.024403 | T | F | F | T |
| Eif2s3x | 0.46887793 | 9.25551973 | 7.69835769 | 0.00726 | 0.024483 | T | F | T | T |
| Bok | -1.2532957 | 3.487761 | 7.6926378 | 0.00728 | 0.024548 | F | F | T | F |
| Mark2 | 0.53270943 | 7.43490107 | 7.69158382 | 0.00729 | 0.024557 | F | F | T | F |
| Cbr1 | -0.8318558 | 4.87919724 | 7.68951188 | 0.00729 | 0.024579 | F | F | F | F |
| Pou6f1 | -7.2521592 | -0.0921375 | 7.68626222 | 0.00731 | 0.024614 | T | F | T | T |
| Kmt5b | 0.63833942 | 6.17215101 | 7.68424497 | 0.00731 | 0.024635 | F | F | T | F |
| Syt2 | -7.348438 | -0.0679785 | 7.68053261 | 0.00733 | 0.024676 | F | F | T | T |
| Cacng4 | -1.2269228 | 3.56830257 | 7.67783759 | 0.00734 | 0.024705 | F | F | F | F |
| Prcp | -0.8011061 | 4.99958235 | 7.671319 | 0.00736 | 0.024778 | F | F | T | T |
| Ndufaf6 | -1.3115563 | 3.30335064 | 7.67120973 | 0.00736 | 0.024778 | F | F | F | T |
| Mtmr3 | -0.6930469 | 5.8781396 | 7.67056364 | 0.00736 | 0.024782 | F | F | T | F |
| Zbtb24 | -1.0848841 | 3.90978679 | 7.67014715 | 0.00736 | 0.024784 | T | F | F | F |
| Carmil2 | 1.54900792 | 2.39331179 | 7.66898276 | 0.00737 | 0.024794 | F | F | F | F |
| Nrbp1 | -0.6651218 | 6.15969643 | 7.66764748 | 0.00737 | 0.024806 | F | F | T | F |
| Zfp867 | -1.3265717 | 3.2888298 | 7.66597979 | 0.00738 | 0.024823 | F | F | F | T |
| Syap1 | 0.96526245 | 3.99868767 | 7.66520667 | 0.00738 | 0.024829 | F | F | F | F |
| Zbtb4 | -1.9776908 | 2.03758004 | 7.66448597 | 0.00738 | 0.024834 | T | F | T | T |
| Bicra | -0.6852573 | 5.87191653 | 7.65311374 | 0.00743 | 0.024969 | F | F | F | F |
| Vmn2r31 | 2.55341559 | 0.73552565 | 7.64917446 | 0.00744 | 0.025014 | F | F | F | F |
| Plp1 | -2.3227948 | 1.57903154 | 7.6464474 | 0.00745 | 0.02504 | F | F | T | T |
| Trmt11 | -0.81094 | 5.00268469 | 7.64642971 | 0.00745 | 0.02504 | F | F | F | F |
| 1700007K1 | -7.2462149 | -0.0823471 | 7.64236042 | 0.00747 | 0.025084 | F | F | T | F |
| Il3ra | 1.24848329 | 3.10995383 | 7.64227637 | 0.00747 | 0.025084 | F | F | F | T |
| Zfpm2 | -7.2933285 | -0.019121 | 7.63516755 | 0.00749 | 0.025168 | T | T | T | T |
| 1700017B0. | 0.57275118 | 6.76403181 | 7.63337761 | 0.0075 | 0.025186 | T | F | F | F |
| Aldh1l1 | -7.3313014 | -0.0805147 | 7.63196296 | 0.0075 | 0.0252 | F | F | F | T |
| Fam124a | -3.8574121 | 0.56159664 | 7.62921675 | 0.00751 | 0.02523 | F | F | T | F |
| Timm17a | 0.62054847 | 6.18889351 | 7.62723791 | 0.00752 | 0.025251 | F | F | T | T |
| Cenpm | -1.3113177 | 3.29887343 | 7.62666978 | 0.00752 | 0.025254 | T | F | T | F |
| Rbfox2 | -0.5382386 | 7.74589567 | 7.62596613 | 0.00753 | 0.025259 | F | F | T | T |
| Egln2 | -0.6963992 | 5.74606376 | 7.61996373 | 0.00755 | 0.02533 | F | F | F | F |
| Nfatc3 | 0.54330322 | 7.09833732 | 7.6155313 | 0.00757 | 0.025382 | T | F | T | T |
| Ltk | -4.3500797 | 0.36620026 | 7.61497339 | 0.00757 | 0.025383 | F | F | F | F |
| Fahd2a | -2.8610196 | 1.08152677 | 7.61477742 | 0.00757 | 0.025383 | F | F | F | F |

| | | | | | | | | | |
|---|---|---|---|---|---|---|---|---|---|
| Mef2d | 0.63307061 | 6.10080426 | 7.60896736 | 0.00759 | 0.025452 | T | F | T | T |
| Ackr1 | -7.2425449 | -0.0822071 | 7.60659714 | 0.0076 | 0.025478 | F | F | F | F |
| Swi5 | -0.6392622 | 6.54443356 | 7.60512254 | 0.0076 | 0.025493 | F | F | F | F |
| Zp1 | 4.38474462 | -0.439628 | 7.60407352 | 0.00761 | 0.025502 | F | F | F | F |
| Pdgfra | -0.556888 | 7.4590902 | 7.600162 | 0.00762 | 0.025547 | T | F | T | T |
| Cnih1 | 0.63621998 | 6.00901862 | 7.59840543 | 0.00763 | 0.025566 | T | T | T | T |
| Sart1 | -0.593565 | 6.83197356 | 7.59076577 | 0.00766 | 0.025658 | T | F | T | T |
| Plek | 2.04913844 | 1.35039758 | 7.59029622 | 0.00766 | 0.02566 | T | F | T | T |
| Jakmip1 | -2.2435347 | 1.70141219 | 7.57720662 | 0.00771 | 0.025823 | T | F | T | F |
| Wnk1 | 0.45962731 | 9.51319576 | 7.57610806 | 0.00771 | 0.025833 | T | F | F | F |
| Gm3752 | -7.2956024 | -0.1026395 | 7.5750374 | 0.00772 | 0.025843 | F | F | F | F |
| Gpm6b | -7.2329228 | -0.1008206 | 7.57164871 | 0.00773 | 0.025882 | T | F | T | T |
| Cdc25b | -1.1618664 | 3.68029792 | 7.56706252 | 0.00775 | 0.025937 | F | F | F | F |
| Cbx4 | 1.22895422 | 3.14976646 | 7.56655042 | 0.00775 | 0.025939 | T | F | F | T |
| Ptpn11 | 0.50462851 | 7.99214731 | 7.5630162 | 0.00776 | 0.025981 | T | F | T | T |
| Mgat1 | 0.6705826 | 5.594668 | 7.56153885 | 0.00777 | 0.025996 | T | F | F | T |
| Rplp0 | -0.4373498 | 10.8935248 | 7.56089515 | 0.00777 | 0.025997 | F | F | F | F |
| Sec14l4 | -1.6331467 | 2.63507577 | 7.56080101 | 0.00777 | 0.025997 | F | F | F | F |
| Atxn2 | -0.5624619 | 7.16344277 | 7.56011225 | 0.00778 | 0.026 | T | F | F | F |
| Fbxo10 | 0.91364452 | 4.14758129 | 7.55996184 | 0.00778 | 0.026 | F | F | T | F |
| Jmjd1c | 0.53199142 | 7.61544284 | 7.5595839 | 0.00778 | 0.026001 | T | T | T | T |
| Gm14409 | 2.23324928 | 1.10647356 | 7.55439089 | 0.0078 | 0.026063 | F | F | F | F |
| Pip4k2b | 0.56916372 | 6.76836037 | 7.5540532 | 0.0078 | 0.026064 | T | F | F | F |
| Mat2a | -0.5211995 | 8.12129764 | 7.54924158 | 0.00782 | 0.026122 | T | T | F | T |
| 5430403G1 | -1.7567391 | 2.39790891 | 7.54661601 | 0.00783 | 0.026152 | F | F | F | T |
| Pip5k1a | 0.62511241 | 6.09742984 | 7.54327278 | 0.00784 | 0.026191 | F | F | T | F |
| Dhx8 | -0.5596985 | 7.21442188 | 7.53446413 | 0.00787 | 0.026301 | T | F | F | T |
| Kcne1l | -7.2286323 | -0.1150791 | 7.53153118 | 0.00789 | 0.026336 | F | F | T | F |
| Nars2 | 0.83062651 | 4.49451417 | 7.5233936 | 0.00792 | 0.026438 | T | F | T | T |
| Babam2 | 0.89413491 | 4.20622235 | 7.52233868 | 0.00792 | 0.026447 | F | F | F | F |
| Zfp442 | -1.610751 | 2.69152863 | 7.5187685 | 0.00794 | 0.02649 | F | F | F | T |
| Cep350 | 0.53525193 | 7.25195067 | 7.51524163 | 0.00795 | 0.026532 | T | F | F | T |
| Egr1 | 1.50913111 | 2.41829056 | 7.5108941 | 0.00797 | 0.026582 | T | F | T | F |
| Abcg8 | -7.2743837 | -0.1163502 | 7.51056693 | 0.00797 | 0.026582 | T | F | T | F |
| Dusp26 | -7.2743837 | -0.1163502 | 7.51056693 | 0.00797 | 0.026582 | F | F | F | F |
| Srsf2 | -0.4833906 | 9.16063718 | 7.51013672 | 0.00797 | 0.026583 | T | F | T | T |
| Olfr986 | 0.69579965 | 5.38169101 | 7.50888495 | 0.00798 | 0.026596 | F | F | F | F |
| Bspry | 0.93486338 | 4.09117473 | 7.50834673 | 0.00798 | 0.026599 | T | F | F | F |
| Purg | -1.2524643 | 3.41249365 | 7.50515793 | 0.00799 | 0.026636 | T | F | F | T |
| Htr3a | -7.2337656 | -0.1247579 | 7.50220673 | 0.008 | 0.026671 | F | F | F | T |
| Txlnb | -7.2334812 | -0.1249904 | 7.5005011 | 0.00801 | 0.02669 | T | T | T | T |
| Rtl1 | -3.6701146 | 0.59885547 | 7.49662567 | 0.00802 | 0.026737 | F | F | F | F |

| | | | | | | | | | |
|---|---|---|---|---|---|---|---|---|---|
| P4ha2 | 0.9699723 | 3.96103673 | 7.49176117 | 0.00804 | 0.026797 | T | F | F | F |
| Gm14137 | -7.2465855 | -0.1263555 | 7.48943858 | 0.00805 | 0.026824 | T | F | F | T |
| Arhgap12 | -0.7488903 | 5.23609108 | 7.48572233 | 0.00807 | 0.026869 | T | F | T | T |
| Plod3 | 0.62333667 | 6.24668375 | 7.48477154 | 0.00807 | 0.026876 | T | F | F | T |
| Eif1 | 0.48569441 | 8.58414423 | 7.48460008 | 0.00807 | 0.026876 | F | F | T | F |
| Serinc3 | -0.656989 | 6.08428849 | 7.47776263 | 0.0081 | 0.026963 | T | F | T | T |
| Tmem35a | -2.8822584 | 1.07569719 | 7.46694735 | 0.00814 | 0.027101 | F | F | T | F |
| Itpk1 | 0.68071677 | 5.46565536 | 7.46683459 | 0.00814 | 0.027101 | T | F | T | F |
| Tars2 | 0.62383907 | 6.13482788 | 7.46568063 | 0.00815 | 0.027112 | T | F | T | F |
| Slc1a5 | 0.66992565 | 5.55823412 | 7.46194089 | 0.00816 | 0.027159 | F | F | F | T |
| Pacs1 | 0.64120334 | 5.88529268 | 7.46124784 | 0.00817 | 0.027164 | F | F | T | F |
| Sik1 | 0.60921841 | 6.26364506 | 7.45333328 | 0.0082 | 0.027266 | T | F | F | F |
| Sec61b | -0.7009179 | 5.72244487 | 7.44542218 | 0.00823 | 0.027369 | F | F | F | F |
| Ahrr | -4.3452734 | 0.34342874 | 7.44363813 | 0.00824 | 0.027389 | T | F | T | F |
| Zfp94 | -1.8185566 | 2.24825041 | 7.44320936 | 0.00824 | 0.027391 | F | F | T | F |
| Usp48 | 0.51355648 | 7.7312343 | 7.44274125 | 0.00824 | 0.027393 | F | F | T | F |
| Npbwr1 | -7.2570302 | -0.0503836 | 7.43715355 | 0.00826 | 0.027465 | F | F | F | T |
| Srsf3 | -0.5031312 | 8.48224781 | 7.43504794 | 0.00827 | 0.027489 | F | F | T | F |
| Ikbip | -1.3133957 | 3.24757825 | 7.42959779 | 0.00829 | 0.02756 | T | F | T | F |
| Ccdc191 | -1.4371688 | 2.99272332 | 7.42831305 | 0.0083 | 0.027573 | F | F | F | F |
| Kcnj4 | -7.2325492 | -0.0775984 | 7.42669434 | 0.00831 | 0.027591 | T | F | F | T |
| Hps5 | 1.02428125 | 3.88082829 | 7.42638503 | 0.00831 | 0.027591 | F | F | F | F |
| 5330417C2 | -7.2382937 | -0.1351959 | 7.42298251 | 0.00832 | 0.027633 | F | F | T | F |
| Cbln1 | -1.9646473 | 2.03098573 | 7.42175185 | 0.00833 | 0.027646 | T | F | T | F |
| Rgs4 | -7.2485382 | -0.1331128 | 7.41851776 | 0.00834 | 0.027686 | F | F | T | F |
| Slc25a16 | 0.89699863 | 4.17453361 | 7.41630638 | 0.00835 | 0.027712 | T | F | T | T |
| Pld2 | 1.00724465 | 3.82883714 | 7.41586208 | 0.00835 | 0.027714 | T | F | F | F |
| Pirb | 2.14417102 | 1.17426493 | 7.41150908 | 0.00837 | 0.02777 | F | F | F | F |
| Cnmd | -7.2077899 | -0.130077 | 7.40913109 | 0.00838 | 0.027798 | F | F | F | F |
| Cela1 | 4.65102019 | -0.6176519 | 7.40386285 | 0.0084 | 0.027867 | T | F | F | F |
| Brca1 | -0.5915374 | 6.78099365 | 7.40104131 | 0.00841 | 0.027902 | T | F | F | T |
| Pigc | 1.17711648 | 3.26621911 | 7.40013166 | 0.00842 | 0.02791 | T | F | T | T |
| Kif1b | -0.5623502 | 7.06659095 | 7.39774607 | 0.00843 | 0.027939 | T | F | T | T |
| Dmtn | -2.2982841 | 1.55300069 | 7.39713559 | 0.00843 | 0.027943 | T | F | T | T |
| Mettl2 | 0.83997375 | 4.50770368 | 7.39426154 | 0.00844 | 0.027979 | F | F | T | T |
| Plekha8 | -0.6653882 | 6.22992436 | 7.38691612 | 0.00847 | 0.028077 | F | F | T | T |
| Mtmr4 | -0.554762 | 7.23069949 | 7.38527115 | 0.00848 | 0.028095 | T | F | T | F |
| Scrn2 | -1.1781988 | 3.61663802 | 7.38401074 | 0.00848 | 0.028109 | F | F | F | F |
| Traf7 | 0.61835039 | 6.16585736 | 7.38309598 | 0.00849 | 0.028117 | F | F | F | F |
| Atrip | -0.9836463 | 4.16865147 | 7.3819004 | 0.00849 | 0.028129 | F | F | F | F |
| Xpnpep1 | 0.58643732 | 6.53622057 | 7.38058859 | 0.0085 | 0.028143 | F | F | T | F |
| Spopl | 1.13263621 | 3.45044542 | 7.37227676 | 0.00853 | 0.028256 | F | F | F | F |

| | | | | | | | | | |
|---|---|---|---|---|---|---|---|---|---|
| Maoa | -0.7695723 | 5.08035846 | 7.3711903 | 0.00854 | 0.028267 | F | F | T | T |
| Adgrl3 | 2.12379052 | 1.19747322 | 7.36882893 | 0.00855 | 0.028295 | T | F | T | F |
| Gm2916 | 2.54994423 | 0.68397803 | 7.36817739 | 0.00855 | 0.0283 | F | F | F | F |
| Cd180 | 4.77034183 | -0.625948 | 7.36670201 | 0.00856 | 0.028317 | T | F | F | F |
| Chchd6 | -1.1352167 | 3.69691815 | 7.36199134 | 0.00858 | 0.028379 | T | F | T | F |
| Psmd3 | 0.49101631 | 8.2392211 | 7.36062146 | 0.00858 | 0.028394 | T | F | F | F |
| Raver1 | -0.5152783 | 8.16249289 | 7.35955939 | 0.00859 | 0.028404 | F | F | T | F |
| Gria1 | -4.3308528 | 0.33221166 | 7.35765339 | 0.0086 | 0.028427 | T | F | T | T |
| Reps2 | 2.72224328 | 0.71186835 | 7.35590419 | 0.0086 | 0.028447 | T | T | T | T |
| Gm5891 | 3.24975792 | 0.10554034 | 7.35494656 | 0.00861 | 0.028457 | F | F | F | F |
| Tmem209 | 0.61663738 | 6.25606027 | 7.35365055 | 0.00861 | 0.028467 | T | F | F | F |
| Olfr853 | 1.45461769 | 2.52472614 | 7.35361992 | 0.00861 | 0.028467 | F | F | F | F |
| Top1 | -0.5063673 | 8.34150083 | 7.34972565 | 0.00863 | 0.028517 | T | F | T | F |
| Hcn2 | 1.33821023 | 2.80480291 | 7.33782787 | 0.00868 | 0.028682 | F | F | T | F |
| Slc35a4 | 0.56778047 | 6.68758372 | 7.33721587 | 0.00868 | 0.028687 | F | F | T | F |
| Mcrip1 | -0.8792617 | 4.52583012 | 7.33588594 | 0.00869 | 0.028699 | F | F | T | F |
| Tmem200c | -3.2691574 | 0.77299465 | 7.33556931 | 0.00869 | 0.028699 | F | F | F | F |
| Serpinc1 | -2.6623269 | 1.19228943 | 7.33543622 | 0.00869 | 0.028699 | F | F | F | F |
| Gck | -3.6421573 | 0.58124988 | 7.33471232 | 0.00869 | 0.028705 | F | F | T | F |
| Gtf2e1 | -0.6892525 | 5.67596963 | 7.33084951 | 0.00871 | 0.028756 | F | F | T | F |
| Ak5 | -3.6455844 | 0.57797622 | 7.32014799 | 0.00876 | 0.028905 | T | F | T | T |
| Tada3 | -0.8079937 | 4.88239369 | 7.31834532 | 0.00877 | 0.028926 | T | F | F | F |
| Ten1 | 1.14641616 | 3.35649442 | 7.31715525 | 0.00877 | 0.028939 | F | F | T | T |
| Vill | 1.24571483 | 3.21393053 | 7.31613424 | 0.00878 | 0.028947 | F | F | F | F |
| Hcn1 | -7.2177716 | -0.150733 | 7.31596006 | 0.00878 | 0.028947 | T | T | T | T |
| Mylk3 | -7.1904231 | -0.1418668 | 7.30823283 | 0.00881 | 0.029055 | T | F | T | F |
| Il23r | 3.12147192 | 0.19964074 | 7.30721647 | 0.00881 | 0.029065 | F | F | F | T |
| Klhl2 | 0.63279865 | 5.93330864 | 7.30516668 | 0.00882 | 0.02909 | T | F | F | F |
| Arhgef3 | 0.93252347 | 4.07830748 | 7.30429886 | 0.00883 | 0.029098 | T | F | T | T |
| Cc2d1b | 0.59263212 | 6.39144125 | 7.30208155 | 0.00884 | 0.029126 | T | F | T | T |
| AI854703 | -3.6545467 | 0.57815556 | 7.30051273 | 0.00884 | 0.029144 | T | F | F | F |
| Gm773 | -7.2443645 | -0.1445891 | 7.29853302 | 0.00885 | 0.029168 | T | T | F | T |
| Dlx6 | -1.9480741 | 1.99786987 | 7.28778447 | 0.0089 | 0.029321 | F | F | F | F |
| Ifi27l2b | -4.3037051 | 0.32675574 | 7.2860643 | 0.00891 | 0.029341 | F | F | F | F |
| Il33 | -7.1873568 | -0.1482143 | 7.28313954 | 0.00892 | 0.02938 | F | F | T | T |
| Wtip | -0.7514997 | 5.16225355 | 7.28245497 | 0.00892 | 0.029385 | T | F | F | F |
| Stx1a | -3.2406783 | 0.88780092 | 7.28131921 | 0.00893 | 0.029397 | F | F | T | F |
| Cited4 | -2.4664835 | 1.35959874 | 7.27929535 | 0.00894 | 0.029422 | F | F | F | F |
| Ndufs8 | 0.66950091 | 5.5228409 | 7.27879978 | 0.00894 | 0.029422 | F | F | F | F |
| Itga2b | 2.1722309 | 1.1276171 | 7.27871695 | 0.00894 | 0.029422 | F | F | F | F |
| Snx17 | 0.62953504 | 6.10093427 | 7.27507288 | 0.00896 | 0.029471 | T | F | F | T |
| Zcchc2 | 0.65610158 | 5.64800667 | 7.27450642 | 0.00896 | 0.029475 | T | F | T | F |

| | | | | | | | | | |
|---|---|---|---|---|---|---|---|---|---|
| Dbi | 0.72342037 | 5.09765139 | 7.27111552 | 0.00897 | 0.02952 | F | F | F | F |
| Cept1 | 0.81603921 | 4.4856638 | 7.26930769 | 0.00898 | 0.029542 | F | F | T | F |
| Soga1 | 0.51453001 | 7.49049664 | 7.26873043 | 0.00899 | 0.029543 | F | F | T | T |
| Sptlc1 | 0.67542772 | 5.41416936 | 7.26864197 | 0.00899 | 0.029543 | T | F | T | F |
| Dusp12 | -1.0552768 | 3.9209864 | 7.2606958 | 0.00902 | 0.029656 | F | F | F | T |
| Maged2 | 0.66988794 | 5.46969091 | 7.25786476 | 0.00903 | 0.029693 | F | F | F | F |
| Spata5 | 0.59796305 | 6.3042819 | 7.25556792 | 0.00904 | 0.029723 | F | F | F | F |
| Immt | 0.50186881 | 7.78521939 | 7.25042437 | 0.00907 | 0.029794 | T | F | T | T |
| Zkscan17 | 0.54587618 | 6.91438075 | 7.24997211 | 0.00907 | 0.029796 | T | F | F | T |
| Mthfr | -0.6500457 | 6.21596014 | 7.24969633 | 0.00907 | 0.029796 | T | F | T | T |
| Atoh8 | -1.5064583 | 2.82284798 | 7.2475912 | 0.00908 | 0.029823 | F | F | T | F |
| Ctnna2 | -2.2747167 | 1.55005822 | 7.24640793 | 0.00909 | 0.029836 | T | F | T | F |
| 9430015G1 | 0.90365403 | 4.11702084 | 7.24479527 | 0.00909 | 0.029855 | T | F | T | T |
| Fnip1 | 0.73142584 | 5.02539131 | 7.24338982 | 0.0091 | 0.029872 | T | T | T | T |
| Rims4 | -2.7328624 | 1.11158273 | 7.24071147 | 0.00911 | 0.029907 | F | F | F | F |
| Hmg20a | -0.6746415 | 5.80236215 | 7.23378336 | 0.00914 | 0.030006 | T | F | T | F |
| Lhfp | 0.88976539 | 4.36506742 | 7.2329051 | 0.00915 | 0.030015 | F | F | F | F |
| Slc11a2 | -0.6943404 | 5.5873862 | 7.23025858 | 0.00916 | 0.03005 | T | F | T | T |
| Usp25 | 0.63124356 | 5.89384994 | 7.22761026 | 0.00917 | 0.030085 | F | F | F | F |
| Olfr1191-ps | 1.09570908 | 3.78258153 | 7.22291279 | 0.00919 | 0.030151 | F | F | F | F |
| Ankrd17 | 0.47496636 | 8.60168225 | 7.22237415 | 0.0092 | 0.030155 | F | F | T | F |
| Ermard | -1.2339768 | 3.3874621 | 7.22190315 | 0.0092 | 0.030157 | F | F | F | F |
| Apol9a | 7.08278695 | -1.2206401 | 7.22399423 | 0.00922 | 0.030232 | T | F | T | T |
| Serpina1a | -7.1868141 | -0.1205781 | 7.21042106 | 0.00925 | 0.030321 | T | F | F | F |
| Pmm1 | 0.73018006 | 4.99439235 | 7.20684814 | 0.00927 | 0.030371 | T | F | T | T |
| Rpl29 | -0.4851046 | 8.88399494 | 7.2045819 | 0.00928 | 0.0304 | F | F | F | F |
| Jrk | -0.9945136 | 4.08258388 | 7.20344536 | 0.00928 | 0.030413 | F | F | T | T |
| Slamf9 | 7.07982365 | -1.2228325 | 7.2083509 | 0.00929 | 0.030445 | F | F | F | F |
| Ythdf3 | 0.51943271 | 7.54287228 | 7.19685807 | 0.00931 | 0.030502 | T | F | T | F |
| Pcdh11x | -7.3302641 | 0.02134997 | 7.19656921 | 0.00931 | 0.030502 | T | T | T | T |
| Ube2j2 | 0.57688474 | 6.53017626 | 7.19636629 | 0.00932 | 0.030502 | F | F | T | F |
| Ccdc117 | 0.64156717 | 5.72621821 | 7.19568657 | 0.00932 | 0.030508 | F | F | T | T |
| Nt5c2 | -0.857469 | 4.57965216 | 7.19488483 | 0.00932 | 0.030516 | F | F | T | T |
| Larp4 | 0.49290773 | 8.03661754 | 7.1926927 | 0.00933 | 0.030544 | T | T | T | T |
| Gnptab | 0.59304103 | 6.31305798 | 7.19172075 | 0.00934 | 0.030555 | T | T | F | T |
| Vmn2r62 | 7.07757156 | -1.2244979 | 7.19648493 | 0.00935 | 0.030589 | F | F | F | F |
| Atp5j | -0.7517975 | 5.26043549 | 7.18700064 | 0.00936 | 0.030617 | F | F | F | T |
| Rad23b | 0.48056757 | 8.33247176 | 7.18413966 | 0.00937 | 0.030657 | F | F | T | F |
| Ube2m | -0.5883904 | 6.77887952 | 7.17275969 | 0.00943 | 0.030827 | F | F | F | F |
| Arl5c | 1.46172316 | 2.49902132 | 7.16981378 | 0.00944 | 0.030867 | F | F | F | F |
| Nfu1 | 0.79866723 | 4.56625167 | 7.16663067 | 0.00945 | 0.030912 | F | F | F | T |
| Gm6710 | 2.46612539 | 1.01858513 | 7.16499482 | 0.00946 | 0.030932 | F | F | F | T |

| | | | | | | | | | |
|---|---|---|---|---|---|---|---|---|---|
| Ccdc173 | -2.5500339 | 1.24621656 | 7.16456714 | 0.00946 | 0.030934 | F | F | F | F |
| Fam169b | -1.6999639 | 2.41932128 | 7.15882176 | 0.00949 | 0.031019 | T | F | T | F |
| Tmem159 | 1.46512756 | 2.44764291 | 7.15828837 | 0.00949 | 0.031022 | F | F | T | T |
| Atp8b1 | 2.09754734 | 1.28651378 | 7.15780928 | 0.0095 | 0.031025 | T | F | T | T |
| Suv39h1 | -0.6348028 | 6.19716189 | 7.1568482 | 0.0095 | 0.031035 | F | F | T | T |
| Dynlt3 | 1.03166455 | 3.65937813 | 7.15518895 | 0.00951 | 0.031056 | T | F | T | F |
| Alkbh5 | 0.50084994 | 7.84317984 | 7.15185062 | 0.00952 | 0.031103 | T | F | T | F |
| Cavin2 | -4.3081555 | 0.35772735 | 7.1509359 | 0.00953 | 0.031113 | T | F | F | T |
| Slc25a3 | 0.4696966 | 8.67885218 | 7.1476101 | 0.00955 | 0.03116 | T | T | T | T |
| Fkbp1a | 0.48298632 | 8.21431011 | 7.14418071 | 0.00956 | 0.031209 | T | F | F | T |
| Adamtsl2 | -7.1688247 | -0.139416 | 7.13528195 | 0.0096 | 0.031343 | F | F | F | F |
| Kdm5d | 1.11892441 | 3.37972937 | 7.1345242 | 0.00961 | 0.03135 | T | F | F | T |
| Erich2 | -2.6651714 | 1.14011037 | 7.13202682 | 0.00962 | 0.031385 | F | F | F | F |
| Mars | 0.50468393 | 7.62202792 | 7.12932993 | 0.00963 | 0.031422 | F | F | F | F |
| Cramp1l | 0.52559597 | 7.16139819 | 7.12900275 | 0.00963 | 0.031423 | F | F | F | F |
| Irak1bp1 | -1.3674897 | 3.06063978 | 7.12703396 | 0.00964 | 0.031449 | T | F | F | F |
| Narf | -0.7991317 | 4.82687938 | 7.12667296 | 0.00965 | 0.03145 | T | F | T | T |
| Marc1 | -7.1636878 | -0.147258 | 7.12514769 | 0.00965 | 0.031469 | F | F | T | F |
| Ugt2b34 | -7.1590714 | -0.1621595 | 7.12402868 | 0.00966 | 0.031482 | T | F | T | T |
| Tent4b | 0.58290344 | 6.41680402 | 7.12152809 | 0.00967 | 0.031517 | F | F | T | F |
| Rpl7l1 | -0.5350328 | 7.39878172 | 7.11878033 | 0.00968 | 0.031555 | T | F | T | T |
| Pik3c3 | 0.65412209 | 5.56714126 | 7.11220669 | 0.00972 | 0.031655 | T | F | F | T |
| Esco2 | -0.866714 | 4.5791462 | 7.10578648 | 0.00975 | 0.031752 | T | T | F | T |
| Retreg2 | 0.67708277 | 5.37211275 | 7.10466322 | 0.00975 | 0.031765 | F | F | T | F |
| Tpcn1 | -0.5514866 | 7.11938068 | 7.10373121 | 0.00976 | 0.031775 | F | F | T | F |
| Gdpd1 | -1.8458844 | 2.142156 | 7.10216508 | 0.00976 | 0.031795 | T | F | T | T |
| Pi16 | 0.9773757 | 3.92145513 | 7.09917148 | 0.00978 | 0.031838 | F | F | F | T |
| Dlgap1 | -7.2638634 | -0.1453399 | 7.09875445 | 0.00978 | 0.03184 | T | F | T | F |
| Tfb2m | 0.78998614 | 4.57588583 | 7.09827599 | 0.00978 | 0.031842 | T | F | T | F |
| Vmn2r4 | 2.11337787 | 1.8851931 | 7.09525788 | 0.0098 | 0.031886 | F | F | F | F |
| Fam76b | 0.70327005 | 5.21976249 | 7.08728301 | 0.00984 | 0.032009 | T | T | T | T |
| Abcg4 | -7.1544694 | -0.1773808 | 7.08406407 | 0.00985 | 0.032055 | F | F | F | F |
| Mast2 | 0.55150499 | 6.76667729 | 7.08304688 | 0.00986 | 0.032067 | F | F | F | F |
| Olfr976 | 2.93526467 | 0.33871122 | 7.07898777 | 0.00988 | 0.032127 | T | F | T | T |
| Irf9 | 1.00078893 | 3.81268538 | 7.07751356 | 0.00989 | 0.032146 | T | F | F | T |
| Atf3 | 1.7646231 | 1.743142 | 7.07689765 | 0.00989 | 0.032151 | T | F | T | F |
| Cebpg | 0.66784755 | 5.47150785 | 7.07001905 | 0.00992 | 0.032257 | T | T | T | T |
| Ifrd2 | 0.61261624 | 6.2910386 | 7.0681024 | 0.00993 | 0.032283 | F | F | F | F |
| 3110040N1 | -2.3561143 | 1.41595965 | 7.0667338 | 0.00994 | 0.032295 | F | F | F | F |
| Rhoa | 0.47339835 | 8.43591441 | 7.06640458 | 0.00994 | 0.032295 | F | F | T | T |
| Bard1 | -0.6656403 | 5.80234913 | 7.06638518 | 0.00994 | 0.032295 | F | F | T | T |
| Platr25 | -1.0369086 | 3.95138241 | 7.06622177 | 0.00994 | 0.032295 | F | F | F | F |

| | | | | | | | | | |
|---|---|---|---|---|---|---|---|---|---|
| Cyp2j9 | -7.1929506 | -0.1848064 | 7.04472108 | 0.01005 | 0.032639 | F | F | F | T |
| Gramd1a | 0.78553805 | 4.62502373 | 7.0398842 | 0.01007 | 0.032714 | T | F | F | T |
| Rras2 | -0.6145272 | 6.35625331 | 7.03581128 | 0.01009 | 0.032776 | T | T | T | T |
| Zfp866 | -0.6928507 | 5.52291802 | 7.03383752 | 0.0101 | 0.032803 | T | F | T | T |
| Arhgap1 | 0.55796239 | 6.75489491 | 7.03004163 | 0.01012 | 0.032861 | T | F | T | T |
| Iqcd | -7.1728856 | -0.1915567 | 7.02881641 | 0.01013 | 0.032876 | T | F | F | F |
| Ly6a | 1.89338216 | 1.48075718 | 7.02615553 | 0.01014 | 0.032915 | F | F | F | T |
| Kbtbd2 | 0.58638423 | 6.34249351 | 7.02566304 | 0.01015 | 0.032916 | F | F | T | T |
| Gm43518 | 3.75106561 | -0.1610869 | 7.02537045 | 0.01015 | 0.032916 | F | F | F | F |
| Parp12 | 1.54047496 | 2.20767348 | 7.02517107 | 0.01015 | 0.032916 | T | F | T | F |
| Dsel | -1.2123131 | 3.41741804 | 7.02087921 | 0.01017 | 0.032982 | F | F | F | F |
| Spr | -0.7459968 | 5.1388901 | 7.01849374 | 0.01018 | 0.033017 | F | F | T | F |
| Tmem17 | 1.61833319 | 2.01715487 | 7.01718612 | 0.01019 | 0.033034 | T | F | F | F |
| Cpz | -7.1498839 | -0.192936 | 7.01395097 | 0.01021 | 0.033077 | F | F | F | F |
| Gm13420 | -7.1498839 | -0.192936 | 7.01395097 | 0.01021 | 0.033077 | F | F | T | F |
| Ccny | 0.60549256 | 6.06578056 | 7.01204181 | 0.01022 | 0.033104 | T | F | T | T |
| B3gat1 | -7.1500071 | -0.193616 | 7.01083119 | 0.01022 | 0.033119 | T | F | T | T |
| Anks1b | -7.1584062 | -0.1950838 | 7.01020913 | 0.01022 | 0.033125 | T | F | T | T |
| Dad1 | 0.62393254 | 5.9035299 | 7.00891774 | 0.01023 | 0.033141 | F | F | T | F |
| Zfp457 | 2.51801384 | 0.6489694 | 7.00828196 | 0.01023 | 0.033147 | F | F | F | T |
| Actn4 | 0.46109433 | 8.78417609 | 7.00598528 | 0.01025 | 0.03318 | T | F | F | T |
| Rbm19 | 0.51969917 | 7.14630953 | 7.00397144 | 0.01026 | 0.033205 | F | F | T | T |
| E030030I06 | -3.1907514 | 0.87216952 | 7.00386316 | 0.01026 | 0.033205 | F | F | T | F |
| Tbxa2r | -4.2645511 | 0.28911264 | 7.00344036 | 0.01026 | 0.033208 | F | F | T | F |
| Gnat2 | -3.2271536 | 0.72865275 | 6.99666957 | 0.01029 | 0.033312 | T | F | T | F |
| Abhd3 | 1.86423879 | 1.54688477 | 6.99658875 | 0.01029 | 0.033312 | F | F | F | F |
| Dhrs7 | -2.2122784 | 1.56325042 | 6.99404446 | 0.01031 | 0.03335 | F | F | F | F |
| Calr | 0.44126961 | 9.74168679 | 6.99289043 | 0.01031 | 0.033364 | F | F | F | T |
| Ssrp1 | -0.4797332 | 8.74418394 | 6.99168239 | 0.01032 | 0.033379 | F | F | T | F |
| Klk9 | 5.24202987 | -0.8931489 | 6.9898294 | 0.01033 | 0.0334 | F | F | F | T |
| Rasgef1a | 5.24202987 | -0.8931489 | 6.9898294 | 0.01033 | 0.0334 | F | F | F | F |
| Ncf2 | 2.81933622 | 0.35533138 | 6.98932328 | 0.01033 | 0.033404 | F | F | F | F |
| Mtss1 | -0.6689607 | 5.74058692 | 6.98595031 | 0.01035 | 0.033455 | T | F | F | F |
| Nck1 | 0.74330115 | 4.81794758 | 6.98510725 | 0.01035 | 0.033464 | T | F | T | T |
| Bnip3l | -0.5826471 | 6.6691832 | 6.98237558 | 0.01037 | 0.033505 | T | F | T | T |
| G6pdx | -0.664976 | 5.7432222 | 6.9795768 | 0.01038 | 0.033548 | F | F | F | F |
| Abcb9 | 2.06313293 | 1.19804121 | 6.97693806 | 0.0104 | 0.033587 | F | F | F | F |
| Cpq | -7.1434552 | -0.1987065 | 6.97495547 | 0.01041 | 0.033615 | F | F | T | F |
| Tmem41a | 0.94082484 | 3.93231626 | 6.97301735 | 0.01042 | 0.033643 | F | F | T | F |
| Gmfb | 0.52292142 | 7.06640446 | 6.96967425 | 0.01043 | 0.033695 | T | F | T | F |
| Nde1 | -0.5832779 | 6.65117788 | 6.96555953 | 0.01046 | 0.033759 | T | F | T | F |
| Ipmk | 0.58074961 | 6.40839248 | 6.9641635 | 0.01046 | 0.033778 | F | F | T | F |

| | | | | | | | | | |
|---|---|---|---|---|---|---|---|---|---|
| Cdk15 | 1.71168563 | 1.91940415 | 6.95844058 | 0.01049 | 0.03387 | F | F | F | F |
| Traf5 | 2.0943663 | 1.14597618 | 6.95541902 | 0.01051 | 0.033917 | T | F | F | T |
| Efhc1 | -2.4259555 | 1.32175515 | 6.95226344 | 0.01053 | 0.033965 | T | T | F | T |
| Cdk17 | 0.70990604 | 5.08687815 | 6.95042033 | 0.01054 | 0.033992 | T | F | T | T |
| Srsf10 | -0.5194626 | 7.63962469 | 6.94575433 | 0.01056 | 0.034066 | T | F | T | T |
| Rcan2 | -1.7699434 | 2.20933854 | 6.94352907 | 0.01057 | 0.034099 | T | F | T | T |
| Sec24a | 0.5736894 | 6.42438799 | 6.94143646 | 0.01058 | 0.03413 | T | F | T | F |
| B230307C2 | 1.22840223 | 3.00411568 | 6.94078687 | 0.01059 | 0.034136 | F | F | F | F |
| Dglucy | 2.7519267 | 0.41374585 | 6.93272828 | 0.01063 | 0.03427 | T | F | T | F |
| Crem | 1.64983305 | 1.92810576 | 6.92864358 | 0.01065 | 0.034335 | T | F | T | T |
| Lztfl1 | -0.9842238 | 4.03588819 | 6.92625475 | 0.01067 | 0.034371 | F | F | T | F |
| Pak7 | -7.1602984 | -0.205811 | 6.92553057 | 0.01067 | 0.034379 | T | F | F | F |
| Pald1 | -0.6444578 | 6.08608179 | 6.92110963 | 0.01069 | 0.03445 | F | F | T | T |
| Iars2 | 0.54049826 | 6.82368624 | 6.91769438 | 0.01071 | 0.034504 | T | F | T | F |
| Zfp488 | 0.57174251 | 6.63214009 | 6.91644814 | 0.01072 | 0.034517 | T | F | T | T |
| Tubb3 | -0.8526849 | 4.5271554 | 6.91638042 | 0.01072 | 0.034517 | F | F | F | T |
| Map1s | -0.5978436 | 6.45154902 | 6.90618101 | 0.01077 | 0.034689 | F | F | F | F |
| Psmg1 | -0.7008397 | 5.39170816 | 6.90498762 | 0.01078 | 0.034705 | F | F | F | F |
| Hmbs | 0.59370081 | 6.16809243 | 6.90463843 | 0.01078 | 0.034706 | F | F | F | F |
| Ercc6l | -0.6303391 | 6.17009279 | 6.90412661 | 0.01078 | 0.034709 | F | F | F | F |
| Colec12 | -0.7673922 | 5.04199784 | 6.90387066 | 0.01079 | 0.034709 | F | F | F | T |
| Skp1a | -0.5138965 | 7.68899674 | 6.90312717 | 0.01079 | 0.034717 | F | F | T | F |
| Fto | -0.5673787 | 6.7966219 | 6.9021859 | 0.01079 | 0.034728 | T | F | T | T |
| Pcm1 | -0.5173215 | 7.6041061 | 6.90099777 | 0.0108 | 0.034743 | T | T | T | T |
| Cdkn2a | -3.6038789 | 0.57386805 | 6.89661667 | 0.01083 | 0.034815 | T | F | T | F |
| Naf1 | -0.5802483 | 6.74225883 | 6.89596244 | 0.01083 | 0.034821 | T | F | F | T |
| Chmp5 | 0.66201924 | 5.38458749 | 6.89495614 | 0.01083 | 0.034834 | F | F | F | F |
| Trim50 | 3.67312255 | -0.2049979 | 6.89116574 | 0.01085 | 0.034895 | F | F | F | F |
| Dhrs3 | -2.04458 | 1.94453112 | 6.88920592 | 0.01087 | 0.034924 | T | F | T | F |
| Phpt1 | -0.9698531 | 4.13411137 | 6.88455984 | 0.01089 | 0.035001 | F | F | T | F |
| Prrt2 | -7.1516881 | -0.2122795 | 6.88399475 | 0.01089 | 0.035003 | T | F | F | T |
| Nkx2-2 | -7.1356269 | -0.2136496 | 6.88354626 | 0.0109 | 0.035003 | T | F | F | T |
| Trnp1 | -7.1356269 | -0.2136496 | 6.88354626 | 0.0109 | 0.035003 | T | F | T | T |
| Fads3 | 1.82325493 | 1.61430103 | 6.88049096 | 0.01091 | 0.035052 | F | F | F | F |
| Tasor | 0.61622154 | 6.17932508 | 6.87904562 | 0.01092 | 0.035072 | F | F | T | F |
| Ip6k2 | -0.8602912 | 4.45771106 | 6.87735187 | 0.01093 | 0.035097 | T | F | T | F |
| Rabif | 0.7046844 | 5.04293914 | 6.87551818 | 0.01094 | 0.035124 | F | F | F | F |
| Stk3 | 0.80022955 | 4.44359207 | 6.87494849 | 0.01094 | 0.035129 | T | F | F | F |
| Pgls | -0.7934129 | 4.85985333 | 6.87375573 | 0.01095 | 0.035142 | F | F | F | F |
| Dck | -0.6583976 | 5.77430572 | 6.87364784 | 0.01095 | 0.035142 | T | F | T | T |
| Pfdn2 | -0.7600553 | 4.97041739 | 6.87169425 | 0.01096 | 0.035171 | T | T | T | T |
| Gnb1l | -1.5428765 | 2.6344205 | 6.87039054 | 0.01097 | 0.035189 | T | F | T | F |

| | | | | | | | | | |
|---|---|---|---|---|---|---|---|---|---|
| Rpl39 | -0.543199 | 7.39985306 | 6.86905172 | 0.01098 | 0.035207 | F | F | F | F |
| Mtr | 0.62644034 | 5.87689499 | 6.86812637 | 0.01098 | 0.035215 | F | F | F | T |
| Eml6 | 1.7183649 | 1.7799202 | 6.86800582 | 0.01098 | 0.035215 | T | F | T | T |
| Fam183b | -7.1206198 | -0.180763 | 6.86536374 | 0.011 | 0.035257 | T | F | F | F |
| Zfp691 | -1.0199452 | 3.91334509 | 6.86046557 | 0.01102 | 0.035339 | F | F | T | F |
| Syn3 | -7.1706374 | -0.2090807 | 6.85755067 | 0.01104 | 0.035386 | T | F | T | T |
| Cyb5a | -0.7023023 | 5.37408073 | 6.85588684 | 0.01105 | 0.03541 | F | F | F | F |
| Unc50 | 0.79461809 | 4.51914647 | 6.85384029 | 0.01106 | 0.035441 | F | F | F | F |
| Gm49342 | 0.8485588 | 4.22657356 | 6.85111304 | 0.01108 | 0.035485 | F | F | F | F |
| Ifi207 | 4.2657775 | -0.5324943 | 6.850108 | 0.01108 | 0.035498 | F | F | F | F |
| Cdh22 | -7.1258811 | -0.2184597 | 6.8482799 | 0.01109 | 0.035525 | F | F | T | F |
| Clcnka | 2.72123019 | 0.425484 | 6.84522331 | 0.01111 | 0.035575 | F | F | F | F |
| Phka2 | -0.7868544 | 4.79810918 | 6.84322786 | 0.01112 | 0.035605 | T | F | F | F |
| Icosl | -2.0647213 | 1.75860929 | 6.8402519 | 0.01114 | 0.035653 | T | F | F | T |
| Stbd1 | -1.6999131 | 2.31187815 | 6.83922637 | 0.01114 | 0.035666 | F | F | F | F |
| FO907130.1 | 7.00433783 | -1.2782144 | 6.8414784 | 0.01117 | 0.035741 | F | F | F | F |
| Per3 | -1.052446 | 3.85883184 | 6.83146236 | 0.01119 | 0.035796 | T | F | F | F |
| Prelid2 | -1.5991209 | 2.51611503 | 6.83053246 | 0.01119 | 0.035807 | F | F | F | F |
| Npr2 | 1.84216263 | 1.52616199 | 6.82945072 | 0.0112 | 0.035816 | F | F | T | F |
| Tm4sf5 | -2.77299 | 1.00437554 | 6.82943566 | 0.0112 | 0.035816 | F | F | F | F |
| Commd7 | 0.7311304 | 4.8375782 | 6.8235014 | 0.01123 | 0.035918 | T | F | F | F |
| Jmjd4 | -1.0941112 | 3.73152037 | 6.82193666 | 0.01124 | 0.035934 | T | F | T | T |
| Ndufa9 | 0.57695866 | 6.32481825 | 6.82183079 | 0.01124 | 0.035934 | F | F | F | T |
| Rem2 | -2.7752101 | 1.00866577 | 6.82177715 | 0.01124 | 0.035934 | F | F | F | F |
| Eif3j1 | 0.6684225 | 5.26943319 | 6.81891183 | 0.01126 | 0.035977 | T | F | F | T |
| Elp5 | -0.8584741 | 4.45711444 | 6.81881589 | 0.01126 | 0.035977 | F | F | F | F |
| Slc26a1 | -7.1504583 | -0.2202441 | 6.81188666 | 0.0113 | 0.036097 | T | T | T | T |
| Nf2 | -0.5388354 | 7.1318865 | 6.80949695 | 0.01131 | 0.036136 | T | F | F | T |
| Timm8b | -0.7720284 | 4.9168126 | 6.80817942 | 0.01132 | 0.036153 | F | F | T | T |
| Hbb-bt | 3.91852187 | -0.3626536 | 6.80790934 | 0.01132 | 0.036153 | F | F | F | F |
| Wbp4 | -0.9176926 | 4.27706796 | 6.8076862 | 0.01132 | 0.036153 | T | F | T | T |
| Emc4 | 0.73546073 | 4.80166495 | 6.79966399 | 0.01137 | 0.036294 | T | F | F | T |
| Kifc2 | -7.1164474 | -0.2257594 | 6.79840175 | 0.01137 | 0.036312 | F | F | F | F |
| Stap2 | -2.0371246 | 1.74398404 | 6.79611651 | 0.01139 | 0.036348 | F | F | T | F |
| Mex3b | 0.5962007 | 6.10755112 | 6.7947503 | 0.0114 | 0.036368 | T | F | T | F |
| Frg2f1 | -7.1285237 | -0.2269801 | 6.79291033 | 0.01141 | 0.036396 | T | F | F | F |
| Mpi | -0.77249 | 4.87822641 | 6.79130491 | 0.01142 | 0.03642 | F | F | F | T |
| Hsdl1 | -0.8424444 | 4.51153081 | 6.78929885 | 0.01143 | 0.036451 | F | F | T | T |
| Odf3b | -7.1462265 | -0.2237625 | 6.78907652 | 0.01143 | 0.036451 | F | F | F | F |
| Ctsa | 0.52827482 | 6.97085398 | 6.78845435 | 0.01143 | 0.036452 | T | F | T | T |
| Tep1 | 0.73937201 | 4.77166217 | 6.78844142 | 0.01143 | 0.036452 | F | F | F | F |
| Ubxn4 | 0.54973773 | 6.64505474 | 6.78510488 | 0.01145 | 0.036508 | F | F | F | F |

| | | | | | | | | | |
|---|---|---|---|---|---|---|---|---|---|
| Arl2bp | -0.6432342 | 5.92590973 | 6.78380531 | 0.01146 | 0.036526 | T | F | T | T |
| Ifi27 | 4.60519834 | -0.7070678 | 6.78207239 | 0.01147 | 0.036553 | F | F | F | T |
| Atxn7l1 | 0.83825358 | 4.24632162 | 6.77953182 | 0.01148 | 0.036592 | T | F | T | T |
| Alg3 | -0.9086332 | 4.2292761 | 6.7793671 | 0.01148 | 0.036592 | F | F | F | F |
| Polr2g | -0.739917 | 5.13640161 | 6.77852839 | 0.01149 | 0.036602 | T | F | T | F |
| 1110051M2 | -1.0912618 | 3.69686363 | 6.77070349 | 0.01153 | 0.036742 | T | F | T | F |
| Mogat1 | -7.1310283 | -0.2299758 | 6.76615435 | 0.01156 | 0.036821 | F | F | F | F |
| Tnfrsf25 | -7.0973376 | -0.2214931 | 6.7637839 | 0.01157 | 0.036859 | F | F | F | F |
| Brwd3 | -0.7458599 | 5.37327538 | 6.75902701 | 0.0116 | 0.036943 | T | F | T | T |
| Mapk14 | -0.5326978 | 7.31951843 | 6.75643274 | 0.01162 | 0.036986 | F | F | T | F |
| March11 | -4.2491547 | 0.25655164 | 6.75474094 | 0.01163 | 0.037012 | F | F | F | F |
| Gsk3a | -0.5700849 | 6.68657648 | 6.75222855 | 0.01164 | 0.037053 | F | F | F | F |
| Gm21244 | 0.62135193 | 5.99068198 | 6.75178762 | 0.01165 | 0.037056 | F | F | F | F |
| Galnt11 | -1.0360582 | 3.85331517 | 6.73841303 | 0.01172 | 0.037302 | F | F | F | F |
| Parp11 | -1.1288257 | 3.55912652 | 6.73609102 | 0.01174 | 0.03734 | T | F | T | T |
| Gm3488 | -7.1100766 | -0.1808655 | 6.73526392 | 0.01174 | 0.037348 | F | F | F | F |
| Herc6 | -2.9600901 | 0.85491993 | 6.73512728 | 0.01174 | 0.037348 | F | F | T | T |
| Pnp2 | -7.1716173 | -0.2179407 | 6.72815981 | 0.01178 | 0.037474 | T | F | F | F |
| Helb | -0.8195539 | 4.65450298 | 6.72429291 | 0.01181 | 0.037542 | T | F | T | T |
| H2bc14 | -0.5059759 | 7.79158646 | 6.71926843 | 0.01184 | 0.037632 | F | F | F | F |
| Mast1 | -7.1056868 | -0.2382616 | 6.71842561 | 0.01184 | 0.03764 | T | F | F | F |
| Prepl | -0.7184696 | 5.21301062 | 6.7182414 | 0.01184 | 0.03764 | T | F | F | T |
| Car10 | -3.5741032 | 0.50656178 | 6.71723674 | 0.01185 | 0.037654 | T | F | T | F |
| Myo9b | 0.50380259 | 7.28138712 | 6.71676943 | 0.01185 | 0.037657 | F | F | F | F |
| Csdc2 | -3.5652009 | 0.50626775 | 6.71504733 | 0.01186 | 0.037684 | F | F | F | F |
| Slc35b3 | 0.83975271 | 4.23492182 | 6.71266546 | 0.01188 | 0.037724 | F | F | F | F |
| Prss35 | -1.7299168 | 2.21902857 | 6.71026525 | 0.01189 | 0.037764 | F | F | F | F |
| Eya2 | -1.4521616 | 2.84402728 | 6.70528144 | 0.01192 | 0.037854 | F | F | F | F |
| Sowahc | 0.87686573 | 4.09897029 | 6.70461034 | 0.01193 | 0.037862 | F | F | T | T |
| Pus1 | 0.59163342 | 6.07541517 | 6.70401443 | 0.01193 | 0.037867 | T | F | F | F |
| Socs5 | -1.0251301 | 3.91113654 | 6.70256745 | 0.01194 | 0.03789 | T | F | T | T |
| Atp8b2 | 0.5921935 | 6.13804814 | 6.69824485 | 0.01196 | 0.037957 | T | F | T | F |
| Il17f | 6.97372855 | -1.3004125 | 6.70403071 | 0.01197 | 0.037957 | F | F | T | F |
| Olfr453 | 6.97372855 | -1.3004125 | 6.70403071 | 0.01197 | 0.037957 | F | F | F | F |
| Zap70 | 6.97372855 | -1.3004125 | 6.70403071 | 0.01197 | 0.037957 | F | F | F | T |
| Clec4a3 | 6.97372855 | -1.3004125 | 6.70403071 | 0.01197 | 0.037957 | F | F | F | T |
| Kcnn3 | -7.0862161 | -0.2141518 | 6.69350503 | 0.01199 | 0.03803 | F | F | T | T |
| Lpar3 | -1.9732351 | 1.83459197 | 6.69258153 | 0.012 | 0.038038 | F | F | T | F |
| Pced1b | 0.71364433 | 4.91727765 | 6.69253383 | 0.012 | 0.038038 | F | F | F | F |
| Sec16a | 0.47635579 | 7.93600989 | 6.69125325 | 0.01201 | 0.038057 | T | F | T | F |
| Rad54l | -0.6755377 | 5.62242797 | 6.69089004 | 0.01201 | 0.038058 | T | F | T | F |
| Sycp3 | -7.0831353 | -0.2307736 | 6.6879518 | 0.01203 | 0.038109 | F | F | F | F |

| | | | | | | | | | |
|---|---|---|---|---|---|---|---|---|---|
| Pard3 | 0.48983452 | 7.6541228 | 6.68487619 | 0.01205 | 0.038163 | T | F | T | F |
| Setd2 | 0.4756686 | 8.05752005 | 6.68233473 | 0.01206 | 0.038207 | T | F | F | F |
| Vmn1r13 | 0.91724088 | 3.95923326 | 6.68020774 | 0.01207 | 0.038242 | F | F | F | F |
| Tsga10ip | -7.0812542 | -0.2262302 | 6.67952264 | 0.01208 | 0.038244 | F | F | F | F |
| Kctd8 | -7.0812542 | -0.2262302 | 6.67952264 | 0.01208 | 0.038244 | F | F | T | F |
| Mtfr1 | 0.63079183 | 5.64291185 | 6.67843718 | 0.01209 | 0.03826 | T | T | T | T |
| Bace2 | -2.963189 | 0.85349687 | 6.67665074 | 0.0121 | 0.038289 | T | F | T | T |
| Srp9 | 0.58836696 | 6.10428893 | 6.67180283 | 0.01213 | 0.038377 | F | F | F | F |
| Gabarap | -0.6693506 | 5.63268689 | 6.66992248 | 0.01214 | 0.038408 | F | F | F | F |
| Tuft1 | 1.02412838 | 3.57861733 | 6.66904576 | 0.01214 | 0.03842 | F | F | T | T |
| Dicer1 | -0.5142484 | 7.47893823 | 6.667851 | 0.01215 | 0.038437 | T | F | T | T |
| Clcn1 | -7.1419984 | -0.2363175 | 6.66529804 | 0.01217 | 0.038481 | F | F | F | F |
| H2ac20 | -0.5052794 | 7.68085171 | 6.66390714 | 0.01217 | 0.038503 | F | F | F | F |
| Rhbdd2 | -0.852904 | 4.45748977 | 6.663523 | 0.01218 | 0.038505 | T | F | T | F |
| Mob4 | 0.61116958 | 5.80780629 | 6.66094799 | 0.01219 | 0.038549 | F | F | T | T |
| Rhno1 | -0.6365994 | 5.8834414 | 6.65449612 | 0.01223 | 0.03867 | F | F | T | F |
| Mcmdc2 | -7.1268306 | -0.2431301 | 6.65240136 | 0.01225 | 0.038705 | F | F | T | F |
| Gpr107 | 0.58808713 | 6.10398348 | 6.64681913 | 0.01228 | 0.038809 | T | F | T | T |
| Neurog1 | -4.4029794 | 0.35722469 | 6.64641256 | 0.01228 | 0.038812 | F | F | T | F |
| H2-K1 | -1.4654442 | 2.78264921 | 6.64597697 | 0.01229 | 0.038815 | F | F | F | F |
| Ap3b2 | -3.2134623 | 0.74402348 | 6.64408308 | 0.0123 | 0.038846 | F | F | F | F |
| Sirt3 | -1.7434986 | 2.17615796 | 6.6420464 | 0.01231 | 0.038881 | F | F | F | F |
| Snurf | -1.9605823 | 1.84711713 | 6.63980703 | 0.01232 | 0.038919 | T | F | T | F |
| Phxr2 | 2.95166309 | 0.18047798 | 6.63680717 | 0.01234 | 0.038973 | F | F | F | T |
| Pold1 | -0.5306082 | 7.14276111 | 6.63629613 | 0.01235 | 0.038977 | F | F | F | F |
| Lingo1 | -1.1903322 | 3.3817514 | 6.63277008 | 0.01237 | 0.039041 | T | F | F | T |
| Gpsm3 | -1.650816 | 2.36286858 | 6.6263262 | 0.01241 | 0.039163 | F | F | T | F |
| Cyb561d1 | 1.26846605 | 2.95802076 | 6.62567963 | 0.01241 | 0.03917 | T | F | T | T |
| Olfr362 | 2.12280485 | 1.08288085 | 6.62346345 | 0.01243 | 0.039209 | F | F | F | F |
| Fam174a | -1.8208694 | 2.05344397 | 6.6124064 | 0.0125 | 0.039423 | T | F | T | T |
| Hoxd13 | -7.0814539 | -0.2543176 | 6.60786932 | 0.01253 | 0.039508 | F | F | T | F |
| Vmn2r69 | 1.63238377 | 1.98537806 | 6.60628796 | 0.01254 | 0.039534 | F | F | F | F |
| Synpo | -1.7786728 | 2.15080791 | 6.60435746 | 0.01255 | 0.039567 | T | F | T | T |
| Pcgf6 | -0.6994003 | 5.45533269 | 6.60380502 | 0.01255 | 0.039573 | T | F | F | F |
| Pop4 | -0.745961 | 5.07289991 | 6.60235526 | 0.01256 | 0.039596 | T | F | F | T |
| Pus3 | 0.92874317 | 3.8841922 | 6.59744387 | 0.01259 | 0.039689 | F | F | F | F |
| Ccdc186 | 0.61296088 | 5.82972111 | 6.59602472 | 0.0126 | 0.039712 | F | F | T | F |
| Nabp2 | -0.6441193 | 5.78350562 | 6.59382567 | 0.01261 | 0.03975 | T | F | F | F |
| Gdf3 | -7.1157346 | -0.2527189 | 6.59035865 | 0.01264 | 0.039815 | F | F | F | F |
| Vamp1 | 1.49566033 | 2.20459556 | 6.58936928 | 0.01264 | 0.039829 | F | F | T | F |
| Thoc2 | 0.5423532 | 6.7036472 | 6.5855617 | 0.01267 | 0.0399 | T | T | T | T |
| Cox6b2 | -4.0230977 | 0.36938263 | 6.5824827 | 0.01269 | 0.039957 | F | F | F | T |

| | | | | | | | | | |
|---|---|---|---|---|---|---|---|---|---|
| Golga3 | 0.55405289 | 6.48829187 | 6.58142161 | 0.01269 | 0.039972 | F | F | T | T |
| Gripap1 | -0.8528728 | 4.37423549 | 6.58037465 | 0.0127 | 0.039988 | T | F | F | T |
| Vmn2r84 | -7.3170862 | -0.1139675 | 6.57753794 | 0.01272 | 0.04004 | F | F | F | F |
| Ggnbp1 | 0.99415151 | 3.64344787 | 6.57662543 | 0.01273 | 0.040052 | T | F | F | F |
| Rel | 0.92210365 | 3.90293175 | 6.57607616 | 0.01273 | 0.040058 | F | F | T | T |
| Birc2 | -0.8288804 | 4.52974863 | 6.56379287 | 0.01281 | 0.040302 | F | F | F | F |
| Dcaf15 | -0.6298062 | 5.92916629 | 6.56175691 | 0.01282 | 0.040338 | T | F | F | F |
| Rbm33 | 0.49914983 | 7.41711016 | 6.56082438 | 0.01283 | 0.040351 | T | F | T | F |
| Ccdc146 | -7.0604321 | -0.2523912 | 6.5593306 | 0.01284 | 0.040376 | F | F | T | F |
| Fignl1 | -0.6320378 | 5.93040283 | 6.55898726 | 0.01284 | 0.040377 | T | F | F | T |
| Ccr2 | 2.43677937 | 0.6370678 | 6.55499106 | 0.01287 | 0.040454 | T | F | F | T |
| Qtrt1 | 0.75267033 | 4.65520247 | 6.55295431 | 0.01288 | 0.04049 | F | F | F | F |
| Ggact | -1.6761282 | 2.28909479 | 6.55055411 | 0.01289 | 0.040533 | F | F | T | F |
| Pml | -0.6950112 | 5.30180064 | 6.55016871 | 0.0129 | 0.040535 | T | F | T | F |
| Rps19 | -0.4837702 | 8.58111878 | 6.54405274 | 0.01294 | 0.040655 | T | F | F | T |
| Fam78a | -7.0589491 | -0.257426 | 6.54358644 | 0.01294 | 0.040659 | T | F | T | F |
| Triqk | -7.0833765 | -0.2651627 | 6.54286915 | 0.01295 | 0.040664 | F | F | F | F |
| Zfp566 | -1.2575054 | 3.17191041 | 6.54277689 | 0.01295 | 0.040664 | F | F | F | T |
| Akr1b10 | -1.3305708 | 3.00460862 | 6.53951415 | 0.01297 | 0.040721 | F | F | F | F |
| Brox | 0.67054622 | 5.15537615 | 6.53927589 | 0.01297 | 0.040721 | T | F | F | T |
| Ndufs2 | 0.50043409 | 7.25172876 | 6.53918197 | 0.01297 | 0.040721 | F | F | F | F |
| Rgmb | 0.70570264 | 4.91010207 | 6.53722268 | 0.01298 | 0.040755 | T | F | T | T |
| Olfr901 | 1.71389311 | 1.68978637 | 6.53523763 | 0.013 | 0.040791 | F | F | F | F |
| Eed | -0.6829161 | 5.36059776 | 6.53462864 | 0.013 | 0.040797 | F | F | F | F |
| Degs2 | 5.16043708 | -0.9552307 | 6.532302 | 0.01301 | 0.04084 | F | F | T | F |
| Col11a2 | 1.27525951 | 2.77395365 | 6.53092604 | 0.01302 | 0.040862 | F | T | F | T |
| Hsp90b1 | 0.40959838 | 10.1212757 | 6.52914367 | 0.01304 | 0.040893 | F | F | F | F |
| Gm10184 | -7.3062887 | -0.1234791 | 6.5265311 | 0.01305 | 0.040942 | F | F | F | F |
| Atp8b5 | 3.9290939 | -0.408855 | 6.52583888 | 0.01306 | 0.04095 | F | F | F | F |
| A630089N( | 0.84164005 | 4.18079582 | 6.52466725 | 0.01307 | 0.040969 | F | F | F | F |
| Olfr1028 | 1.19855822 | 3.02659361 | 6.52310088 | 0.01308 | 0.040993 | F | F | F | F |
| Cplx1 | -7.0662238 | -0.2197134 | 6.52273839 | 0.01308 | 0.040993 | T | F | T | T |
| Tmem254a | -2.0575305 | 1.64607094 | 6.52263813 | 0.01308 | 0.040993 | F | F | T | F |
| Vezt | 0.66451175 | 5.20480706 | 6.51077284 | 0.01316 | 0.041235 | T | T | T | T |
| D230025D1 | 0.71606396 | 4.82484921 | 6.50620138 | 0.01319 | 0.041325 | F | F | T | F |
| Cntf | 2.04918226 | 1.12358815 | 6.50557114 | 0.01319 | 0.041333 | T | F | F | T |
| Plb1 | -7.0536873 | -0.2372506 | 6.50188448 | 0.01322 | 0.041404 | T | F | F | F |
| Cdkl3 | -1.7162654 | 2.20940275 | 6.49804729 | 0.01324 | 0.041479 | T | F | F | T |
| Svbp | -1.2994046 | 3.08197136 | 6.495691 | 0.01326 | 0.041516 | F | F | F | F |
| Fbxl6 | -0.8257808 | 4.5069436 | 6.49560279 | 0.01326 | 0.041516 | F | F | F | F |
| Thrap3 | -0.4526344 | 9.0419355 | 6.49547834 | 0.01326 | 0.041516 | T | F | T | F |
| Snx4 | 0.73717798 | 4.7784785 | 6.49193421 | 0.01328 | 0.041582 | T | F | T | T |

| | | | | | | | | | |
|---|---|---|---|---|---|---|---|---|---|
| Cdc37l1 | 0.59308622 | 5.94401432 | 6.491778 | 0.01329 | 0.041582 | T | F | T | T |
| Ttc9 | 1.44396963 | 2.33974286 | 6.48930747 | 0.0133 | 0.041628 | T | F | T | F |
| Slc22a15 | 1.47389428 | 2.22610415 | 6.48859421 | 0.01331 | 0.041637 | F | T | T | T |
| Virma | 0.49459633 | 7.3110573 | 6.47694167 | 0.01339 | 0.041879 | F | F | F | F |
| H2bc21 | -2.3110532 | 1.36681917 | 6.47650038 | 0.01339 | 0.041882 | T | F | F | T |
| Pcdhb4 | 3.88237881 | -0.3381907 | 6.47529793 | 0.0134 | 0.041902 | T | F | T | T |
| Tmdd1 | 4.20237555 | -0.5811142 | 6.47465162 | 0.0134 | 0.04191 | F | F | F | F |
| Ctsz | 0.51345287 | 7.28976766 | 6.47407462 | 0.01341 | 0.041916 | F | F | F | F |
| Rnaset2b | 1.02810397 | 3.49961312 | 6.47302448 | 0.01341 | 0.04193 | T | F | F | T |
| Rpl31 | -0.4815668 | 8.48788012 | 6.4727368 | 0.01342 | 0.04193 | F | F | F | F |
| Fam189a1 | -7.0664493 | -0.2762527 | 6.47254116 | 0.01342 | 0.04193 | F | F | T | T |
| Pcbp2 | -0.443407 | 9.35206845 | 6.4687223 | 0.01344 | 0.042006 | T | F | T | T |
| Fnip2 | 0.52370212 | 6.88967034 | 6.46780518 | 0.01345 | 0.04202 | T | F | T | T |
| Gm11549 | -7.1840512 | -0.2220616 | 6.46625002 | 0.01346 | 0.042047 | F | F | F | F |
| Ncoa5 | -0.5829612 | 6.39248367 | 6.46225524 | 0.01349 | 0.042126 | F | F | T | T |
| Atg2b | 0.54597806 | 6.53562716 | 6.46114629 | 0.01349 | 0.042144 | T | F | T | F |
| Rpa2 | -0.6030784 | 6.17487687 | 6.46079234 | 0.0135 | 0.042146 | F | F | F | F |
| Usp15 | 0.63443835 | 5.52877458 | 6.45967699 | 0.0135 | 0.042163 | T | T | T | T |
| Dnaaf2 | -0.9526475 | 4.03989231 | 6.45418827 | 0.01354 | 0.042275 | F | F | F | F |
| Inka2 | -2.2062907 | 1.44045026 | 6.45226454 | 0.01356 | 0.042311 | F | F | T | T |
| Mvb12a | 0.71153497 | 4.86655058 | 6.44310649 | 0.01362 | 0.042503 | F | F | F | F |
| Ccdc190 | -7.0562389 | -0.2811823 | 6.43985875 | 0.01364 | 0.042567 | F | F | T | F |
| Tyk2 | 0.70328406 | 4.94535325 | 6.43764948 | 0.01366 | 0.042601 | F | F | T | T |
| Zfp410 | 0.79355237 | 4.47197549 | 6.43756631 | 0.01366 | 0.042601 | T | F | T | F |
| Tmem138 | -1.1563549 | 3.40198226 | 6.43745543 | 0.01366 | 0.042601 | T | F | T | F |
| Mark3 | 0.51374366 | 6.93967557 | 6.43435775 | 0.01368 | 0.042662 | T | T | F | T |
| Slc25a48 | 2.67441762 | 0.37769554 | 6.43384087 | 0.01368 | 0.042667 | F | F | F | T |
| Gngt2 | -4.1745999 | 0.22227034 | 6.43069552 | 0.01371 | 0.042729 | F | F | F | F |
| Dcdc2a | -7.060405 | -0.2832631 | 6.42751008 | 0.01373 | 0.042787 | F | F | T | T |
| Fam192a | -0.6538617 | 5.58128258 | 6.42747464 | 0.01373 | 0.042787 | T | F | F | F |
| Zbtb17 | -0.7515896 | 4.88148017 | 6.42371604 | 0.01375 | 0.042851 | F | F | F | F |
| Zmym5 | -0.8276 | 4.55043623 | 6.42356557 | 0.01376 | 0.042851 | T | F | T | T |
| Pdk2 | -1.7210741 | 2.15700665 | 6.42341515 | 0.01376 | 0.042851 | T | F | T | T |
| Fam185a | -0.9086437 | 4.16098406 | 6.42339609 | 0.01376 | 0.042851 | T | T | T | T |
| Prpf39 | -0.7953661 | 4.64739089 | 6.42226361 | 0.01376 | 0.04287 | T | F | T | T |
| Dnajc27 | -0.8142296 | 4.54075352 | 6.42182074 | 0.01377 | 0.042874 | T | F | T | T |
| Olfr6 | 6.90991474 | -1.3462032 | 6.42606242 | 0.01378 | 0.042905 | F | F | F | F |
| Sorcs3 | -7.0533964 | -0.2265045 | 6.41549221 | 0.01381 | 0.043 | T | F | T | T |
| Tox | -7.0387727 | -0.2820142 | 6.41115701 | 0.01384 | 0.043089 | T | F | F | F |
| Gm17535 | 6.90680443 | -1.3484181 | 6.41280518 | 0.01387 | 0.043177 | F | F | F | F |
| Slc4a5 | -1.6399822 | 2.33788267 | 6.40399801 | 0.01389 | 0.043232 | F | F | F | T |
| Aqp7 | 1.36939181 | 2.56410383 | 6.40380445 | 0.01389 | 0.043232 | F | F | T | F |

| | | | | | | | | | |
|---|---|---|---|---|---|---|---|---|---|
| Itgb6 | 6.90588156 | -1.349075 | 6.40887675 | 0.0139 | 0.043244 | T | F | T | F |
| D3Ertd751e | 1.61377661 | 2.13245239 | 6.40162483 | 0.01391 | 0.043268 | F | F | T | T |
| Klhl8 | -1.5076016 | 2.59115229 | 6.39950528 | 0.01393 | 0.043308 | F | F | F | F |
| Cdk2 | -0.6094993 | 6.04270227 | 6.39570211 | 0.01395 | 0.043384 | F | F | T | F |
| Zbtb8a | -1.0453739 | 3.73430148 | 6.39554188 | 0.01395 | 0.043384 | T | F | T | F |
| Thtpa | -1.3092908 | 3.01263102 | 6.39385561 | 0.01397 | 0.043415 | F | F | T | F |
| Smchd1 | 0.46313321 | 8.07651311 | 6.39315263 | 0.01397 | 0.043424 | T | F | F | F |
| Mia3 | 0.47827105 | 7.61196213 | 6.39249662 | 0.01398 | 0.043432 | T | F | F | F |
| Lyrm7 | -2.0186695 | 1.66960549 | 6.39204696 | 0.01398 | 0.043436 | F | F | T | T |
| N4bp1 | -0.5520891 | 6.74140305 | 6.38824061 | 0.01401 | 0.043514 | T | T | T | T |
| Fam149a | -1.3136025 | 3.01711474 | 6.38726539 | 0.01401 | 0.04353 | T | F | T | T |
| Hps4 | -0.7719645 | 4.73414801 | 6.38581419 | 0.01402 | 0.043556 | F | F | F | F |
| Sp3 | 0.58291517 | 6.02067556 | 6.38508432 | 0.01403 | 0.043566 | F | F | F | F |
| 1700010I14 | -2.5557096 | 1.07055424 | 6.37412862 | 0.01411 | 0.043804 | F | F | F | F |
| Rab9b | 4.88219214 | -0.8966963 | 6.37242911 | 0.01412 | 0.043836 | T | F | T | F |
| Rgs17 | -2.1345029 | 1.61140002 | 6.36625402 | 0.01416 | 0.043968 | F | F | T | T |
| St6galnac5 | 1.80868856 | 1.45906049 | 6.36587958 | 0.01417 | 0.04397 | F | F | T | T |
| Rasgrf1 | -2.1958573 | 1.46033761 | 6.36381994 | 0.01418 | 0.044007 | T | F | F | F |
| Pappa2 | 2.4650324 | 0.57333706 | 6.36366315 | 0.01418 | 0.044007 | T | F | T | T |
| Slc6a2 | 1.79053216 | 1.50582375 | 6.36340604 | 0.01418 | 0.044007 | F | F | T | F |
| Polr2d | -0.6614818 | 5.52313678 | 6.36081974 | 0.0142 | 0.044058 | F | F | T | F |
| Cdr2 | -0.7557377 | 4.81618781 | 6.35583477 | 0.01424 | 0.044164 | T | F | T | T |
| Lair1 | 2.23531773 | 0.82689226 | 6.35495866 | 0.01425 | 0.044178 | T | F | T | T |
| Prss42 | 4.91356172 | -0.8852566 | 6.34985642 | 0.01428 | 0.044285 | F | F | T | F |
| Olfr1198 | 1.41671773 | 2.3351655 | 6.34948855 | 0.01429 | 0.044285 | F | F | F | F |
| Gpbp1 | 0.51705804 | 6.87980409 | 6.34936008 | 0.01429 | 0.044285 | T | F | F | T |
| Lhx8 | -7.0615601 | -0.2937341 | 6.3468766 | 0.0143 | 0.044335 | T | F | F | F |
| Olfr887 | 1.69285249 | 1.7734633 | 6.34393803 | 0.01433 | 0.044396 | F | F | F | F |
| Zfp54 | 1.70938914 | 1.64244576 | 6.3412965 | 0.01435 | 0.044449 | F | F | F | F |
| Rcn2 | -0.6304343 | 5.81353593 | 6.33523987 | 0.01439 | 0.044581 | T | F | F | T |
| Ccl28 | -7.0663274 | -0.2939217 | 6.33390392 | 0.0144 | 0.044602 | T | F | T | T |
| Mrps24 | -1.069692 | 3.64790574 | 6.33376098 | 0.0144 | 0.044602 | T | F | T | T |
| Coil | -0.7157096 | 5.05410826 | 6.33075472 | 0.01442 | 0.044664 | T | F | T | F |
| Cmah | 0.89389981 | 3.95961042 | 6.32619396 | 0.01446 | 0.044762 | T | F | T | T |
| Olfr1247 | 0.62662368 | 5.75040206 | 6.32475391 | 0.01447 | 0.044788 | F | F | F | F |
| Olfr1336 | 0.719956 | 4.82555349 | 6.32429987 | 0.01447 | 0.044792 | F | F | F | F |
| Slc6a11 | -7.0512893 | -0.298725 | 6.32293305 | 0.01448 | 0.044817 | T | F | F | T |
| Polr3gl | 1.31153848 | 2.61887962 | 6.32209848 | 0.01449 | 0.04483 | T | F | T | T |
| Acot13 | -1.2867463 | 3.04555674 | 6.31927817 | 0.01451 | 0.044888 | F | F | F | T |
| Rspo4 | -4.1725225 | 0.19578251 | 6.31810769 | 0.01452 | 0.044909 | T | F | F | T |
| Copg2 | -0.6476185 | 5.58796446 | 6.31783314 | 0.01452 | 0.044909 | F | T | T | T |
| Rhof | -1.8086343 | 1.96691591 | 6.31553908 | 0.01454 | 0.044955 | T | F | T | F |

| | | | | | | | | | |
|---|---|---|---|---|---|---|---|---|---|
| Magee1 | 1.25347544 | 2.84318108 | 6.3149748 | 0.01454 | 0.044962 | F | F | F | F |
| Plppr2 | -1.5343621 | 2.50608811 | 6.31044699 | 0.01457 | 0.045055 | F | F | F | F |
| Thbs2 | -2.9092985 | 0.80764973 | 6.31036808 | 0.01457 | 0.045055 | F | F | T | F |
| Dym | 0.62484822 | 5.47427975 | 6.30713677 | 0.0146 | 0.045121 | T | F | F | F |
| C3 | 1.4873033 | 2.15438791 | 6.30692302 | 0.0146 | 0.045121 | F | F | F | F |
| Otx1 | -2.103185 | 1.55463319 | 6.30279697 | 0.01463 | 0.04521 | F | F | F | F |
| Cd8a | -7.04663 | -0.3024123 | 6.30034252 | 0.01465 | 0.045261 | T | F | T | F |
| Kmt2e | -0.6825222 | 5.24386357 | 6.29997326 | 0.01465 | 0.045263 | F | F | T | T |
| Ric8a | 0.92776328 | 3.84385684 | 6.29783035 | 0.01467 | 0.045306 | F | F | F | F |
| Rdh12 | -7.0116183 | -0.295752 | 6.29287334 | 0.01471 | 0.045415 | T | F | F | F |
| Gm3002 | 4.42168287 | -0.7301015 | 6.29038217 | 0.01472 | 0.045466 | T | F | F | F |
| Hck | -7.0147765 | -0.3009042 | 6.28788035 | 0.01474 | 0.045518 | F | F | F | F |
| Brd2 | -0.4947333 | 7.70914869 | 6.28637274 | 0.01475 | 0.045546 | F | F | T | F |
| Slc41a1 | -0.7550028 | 4.78536504 | 6.28505064 | 0.01476 | 0.045571 | T | F | T | F |
| Tpt1 | -0.4543607 | 9.27002206 | 6.27978058 | 0.0148 | 0.045687 | T | F | F | T |
| Usf2 | 0.54639854 | 6.43188442 | 6.27624651 | 0.01483 | 0.045764 | F | F | T | T |
| Art4 | 3.23340687 | -0.0147162 | 6.27343498 | 0.01485 | 0.045823 | T | F | T | T |
| Zfp407 | 0.54882985 | 6.4229267 | 6.2723338 | 0.01486 | 0.045842 | T | F | F | F |
| Zrsr1 | 1.04691231 | 3.36836443 | 6.26661846 | 0.0149 | 0.04597 | F | F | T | T |
| Plcb4 | -1.2752018 | 3.14394429 | 6.26634503 | 0.01491 | 0.04597 | F | F | F | T |
| Senp6 | -0.5567157 | 6.72848798 | 6.26601482 | 0.01491 | 0.045971 | T | F | F | F |
| Efhc2 | 1.66949604 | 1.71035122 | 6.2614166 | 0.01494 | 0.046073 | F | F | F | F |
| Tnfsf10 | 2.38099659 | 0.63493942 | 6.26058414 | 0.01495 | 0.046086 | T | T | T | T |
| Klhl41 | 1.4667067 | 2.21999518 | 6.25564626 | 0.01499 | 0.046196 | F | F | F | F |
| Scn3a | -7.0773652 | -0.2968622 | 6.25466157 | 0.015 | 0.046213 | F | F | T | T |
| H2ac12 | -0.4829682 | 8.46980727 | 6.25413761 | 0.015 | 0.046219 | F | F | F | T |
| Smarcc2 | 0.53318624 | 6.59223487 | 6.25076199 | 0.01503 | 0.046292 | F | F | F | F |
| Tnfaip1 | 0.5436998 | 6.4796475 | 6.24955751 | 0.01504 | 0.046308 | T | F | T | T |
| Srp54b | -7.2464925 | -0.1759836 | 6.24954658 | 0.01504 | 0.046308 | F | F | F | T |
| Dhrs7b | 0.6402061 | 5.27006821 | 6.24799247 | 0.01505 | 0.046338 | T | F | T | T |
| Endov | -0.8127914 | 4.4685585 | 6.24463664 | 0.01507 | 0.046412 | T | F | T | T |
| Pih1d2 | -2.4490496 | 1.15119112 | 6.24425432 | 0.01508 | 0.046414 | F | F | F | F |
| Wapl | 0.46760682 | 8.07574687 | 6.242607 | 0.01509 | 0.046447 | T | F | T | F |
| Rtl5 | -7.1293689 | -0.148135 | 6.24231427 | 0.01509 | 0.046447 | F | F | T | F |
| Plk1 | -0.4870027 | 7.75308463 | 6.23797935 | 0.01512 | 0.046544 | F | F | F | F |
| Pcmtd2 | -0.9708869 | 3.91085565 | 6.23585288 | 0.01514 | 0.046588 | T | F | F | T |
| Hoxa11 | -7.0043991 | -0.2762197 | 6.23494276 | 0.01515 | 0.046598 | F | F | T | F |
| Ednrb | 1.58866909 | 1.96737879 | 6.234874 | 0.01515 | 0.046598 | T | F | T | T |
| Ttc13 | 0.58142697 | 5.94791181 | 6.2288925 | 0.0152 | 0.046735 | T | F | T | F |
| Abhd18 | 1.35556172 | 2.57043038 | 6.227839 | 0.0152 | 0.046747 | F | F | F | F |
| Slc33a1 | 0.7159317 | 4.74784405 | 6.22782599 | 0.0152 | 0.046747 | F | F | T | T |
| Dennd4c | 0.57442111 | 6.04536994 | 6.22168482 | 0.01525 | 0.046888 | T | F | F | F |

| | | | | | | | | | |
|---|---|---|---|---|---|---|---|---|---|
| Rab2a | 0.51756381 | 6.76899079 | 6.22055143 | 0.01526 | 0.046908 | T | F | T | F |
| Slx4 | 0.68201139 | 5.05989579 | 6.21658773 | 0.01529 | 0.046997 | F | F | F | F |
| Slc8a1 | -1.2013626 | 3.23477069 | 6.2131271 | 0.01532 | 0.047074 | T | F | T | T |
| Fanca | -0.6461718 | 5.60684714 | 6.21243242 | 0.01532 | 0.047084 | T | F | T | F |
| Pcnx | 0.56057314 | 6.21224249 | 6.21185802 | 0.01533 | 0.047091 | T | F | F | T |
| Gm614 | 6.85920753 | -1.3821138 | 6.21323222 | 0.01536 | 0.04716 | F | F | F | T |
| Clvs1 | 6.85920753 | -1.3821138 | 6.21323222 | 0.01536 | 0.04716 | T | F | T | T |
| Ddx60 | 6.85920753 | -1.3821138 | 6.21323222 | 0.01536 | 0.04716 | T | T | F | T |
| Fyb | 6.85920753 | -1.3821138 | 6.21323222 | 0.01536 | 0.04716 | T | T | T | T |
| Fcmr | 6.85920753 | -1.3821138 | 6.21323222 | 0.01536 | 0.04716 | F | F | F | F |
| Olfr695 | 2.6952506 | 0.32623529 | 6.2065199 | 0.01537 | 0.04718 | F | F | F | F |
| Nol6 | 0.45773276 | 8.02141034 | 6.20594676 | 0.01538 | 0.047188 | T | F | T | T |
| Aco2 | 0.46082605 | 7.95016136 | 6.20480693 | 0.01538 | 0.047209 | T | F | F | T |
| Sntb1 | -1.2346897 | 3.22001057 | 6.19978173 | 0.01542 | 0.047324 | F | F | F | T |
| Gm266 | -6.9936646 | -0.3107663 | 6.1990666 | 0.01543 | 0.047334 | T | T | T | T |
| Rc3h1 | -0.5152881 | 7.12469477 | 6.18940487 | 0.01551 | 0.047563 | F | F | T | F |
| Srcap | 0.42700208 | 9.17725508 | 6.18697932 | 0.01553 | 0.047616 | T | F | F | T |
| Zcchc14 | -0.5281857 | 6.90015397 | 6.18592626 | 0.01553 | 0.047635 | F | F | F | F |
| Tmem26 | -7.0999357 | -0.2890299 | 6.18431046 | 0.01555 | 0.047667 | T | F | T | T |
| Rfc5 | -0.5742839 | 6.34405217 | 6.18094471 | 0.01557 | 0.047743 | T | F | F | T |
| Tmem38a | -1.5978594 | 2.32810074 | 6.17906687 | 0.01559 | 0.047782 | T | F | T | T |
| Slc12a6 | -0.6577643 | 5.41428865 | 6.17816018 | 0.0156 | 0.047798 | T | F | F | T |
| Ankhd1 | -0.5285845 | 6.9816822 | 6.16711869 | 0.01568 | 0.048063 | F | F | F | F |
| Dhx9 | -0.4444752 | 9.04783377 | 6.16315361 | 0.01572 | 0.048154 | T | F | F | F |
| Usp43 | 0.91553708 | 3.82629929 | 6.16276371 | 0.01572 | 0.048157 | F | F | F | F |
| Pstpip2 | 1.58575539 | 1.89317976 | 6.16159382 | 0.01573 | 0.048179 | T | F | T | F |
| Pex5l | -7.0162491 | -0.3273633 | 6.15622192 | 0.01577 | 0.048305 | F | F | F | T |
| Otud7a | -4.1303257 | 0.18542943 | 6.1547006 | 0.01579 | 0.048325 | F | F | T | T |
| Tsc22d1 | -0.4947633 | 7.48874007 | 6.15441948 | 0.01579 | 0.048325 | F | F | F | F |
| Ceacam1 | -1.6079597 | 2.3401622 | 6.15436379 | 0.01579 | 0.048325 | T | F | T | T |
| Adam4 | 1.61308321 | 1.79540745 | 6.15433422 | 0.01579 | 0.048325 | F | F | F | F |
| Rab30 | -1.3653275 | 2.82204832 | 6.15287109 | 0.0158 | 0.048348 | F | F | T | T |
| Igfbp1 | -4.1474783 | 0.17286042 | 6.15286301 | 0.0158 | 0.048348 | F | F | T | F |
| P2rx1 | -6.9830113 | -0.3156975 | 6.15116013 | 0.01581 | 0.048383 | T | F | T | T |
| Map1lc3b | -0.6122693 | 5.88996168 | 6.1498582 | 0.01582 | 0.048405 | F | F | T | F |
| Asprv1 | -3.5153722 | 0.44365635 | 6.14976383 | 0.01583 | 0.048405 | F | F | T | F |
| Slc25a46 | 0.63672127 | 5.33133635 | 6.14911702 | 0.01583 | 0.048407 | T | F | T | T |
| Olfml2a | -1.164559 | 3.3774667 | 6.14894055 | 0.01583 | 0.048407 | F | F | F | T |
| Hmox2 | 0.63849583 | 5.25219579 | 6.1488571 | 0.01583 | 0.048407 | F | F | F | T |
| Bves | -2.4932426 | 1.08043589 | 6.14513143 | 0.01586 | 0.048486 | T | F | T | T |
| Zfp40 | -1.6791509 | 2.15174199 | 6.14510746 | 0.01586 | 0.048486 | F | F | T | F |
| Noa1 | 0.65141193 | 5.19354387 | 6.1448136 | 0.01587 | 0.048487 | F | F | F | F |

| | | | | | | | | | |
|---|---|---|---|---|---|---|---|---|---|
| Slc6a15 | -6.9892994 | -0.3278165 | 6.14337092 | 0.01588 | 0.048516 | T | F | F | F |
| Gm15128 | 6.84326765 | -1.3933146 | 6.14776457 | 0.01589 | 0.048536 | F | F | F | F |
| Heatr5a | 0.49782739 | 6.97987838 | 6.13947101 | 0.01591 | 0.048597 | F | F | T | F |
| Gm10160 | 0.99848921 | 3.57865435 | 6.13929491 | 0.01591 | 0.048597 | F | F | F | F |
| Hgf | -6.9962951 | -0.3328156 | 6.1312393 | 0.01598 | 0.048792 | T | F | T | T |
| Stx19 | 1.77325288 | 1.58357719 | 6.13061413 | 0.01598 | 0.048801 | F | F | F | F |
| Supt20 | -0.6382068 | 5.58996589 | 6.13009114 | 0.01599 | 0.048807 | F | F | F | F |
| Mrpl51 | 0.62312321 | 5.39999665 | 6.12928718 | 0.01599 | 0.04882 | T | F | T | F |
| Slc15a4 | 0.70844169 | 4.86215808 | 6.12821543 | 0.016 | 0.04884 | F | F | T | T |
| Intu | -0.735085 | 4.91745858 | 6.12513661 | 0.01603 | 0.048907 | T | F | T | T |
| Zc2hc1c | -3.4563344 | 0.43441072 | 6.12497703 | 0.01603 | 0.048907 | T | F | T | T |
| Rab3ip | 0.58869923 | 5.78678334 | 6.12475354 | 0.01603 | 0.048907 | F | F | T | F |
| Nup50 | -0.4773755 | 7.83211246 | 6.12397884 | 0.01604 | 0.048915 | F | F | T | F |
| Spag16 | -1.8139988 | 1.9026671 | 6.12386938 | 0.01604 | 0.048915 | T | F | T | T |
| Zfp106 | -0.4333934 | 9.26432473 | 6.12293076 | 0.01605 | 0.048932 | T | F | F | T |
| Prkab2 | 0.76319828 | 4.45030111 | 6.1172128 | 0.01609 | 0.049069 | T | F | T | T |
| Gp5 | 3.71856952 | -0.366546 | 6.11530207 | 0.01611 | 0.049111 | F | F | F | F |
| Inpp4a | -0.8158208 | 4.40300135 | 6.11500935 | 0.01611 | 0.049111 | T | F | T | T |
| Tmem39b | -0.7815226 | 4.59061038 | 6.11438973 | 0.01612 | 0.04912 | F | F | F | F |
| Plvap | -1.7955982 | 1.96183014 | 6.11407102 | 0.01612 | 0.049121 | F | F | F | T |
| Ppp1r2 | -0.5721304 | 6.32784954 | 6.11103656 | 0.01614 | 0.049191 | T | F | T | T |
| Rab6a | 0.51513186 | 6.744932 | 6.1088567 | 0.01616 | 0.049239 | T | F | T | T |
| Tmem114 | -7.0043586 | -0.3369293 | 6.10696032 | 0.01618 | 0.04928 | F | F | F | F |
| Erich4 | -6.9722978 | -0.3206286 | 6.10335257 | 0.01621 | 0.049361 | F | F | F | F |
| Sox18 | -2.4815912 | 1.55046997 | 6.10322789 | 0.01621 | 0.049361 | F | F | F | F |
| Sox8 | 2.06189706 | 1.02439095 | 6.10271931 | 0.01621 | 0.049362 | F | F | T | F |
| Il17re | 2.3205987 | 0.73383032 | 6.10264217 | 0.01621 | 0.049362 | T | F | F | T |
| Far2 | -2.9386984 | 0.73854301 | 6.10079325 | 0.01623 | 0.049402 | T | F | F | T |
| Olfr344 | 0.70699272 | 5.01038694 | 6.09993948 | 0.01624 | 0.049417 | F | F | F | F |
| Usp1 | 0.47141351 | 7.50759569 | 6.09937662 | 0.01624 | 0.049424 | T | F | F | T |
| 4930447C0[illegible] | -2.5093457 | 1.09607808 | 6.09850124 | 0.01625 | 0.04944 | F | F | F | T |
| Otub2 | -1.5936553 | 2.31798487 | 6.09655701 | 0.01626 | 0.049482 | T | F | F | T |
| Gstk1 | -2.1217584 | 1.4527325 | 6.09373868 | 0.01629 | 0.049547 | F | F | T | F |
| 1700028P1[illegible] | -6.9937001 | -0.3419854 | 6.08375767 | 0.01637 | 0.049789 | F | F | F | T |
| Kcnmb2 | -6.9937001 | -0.3419854 | 6.08375767 | 0.01637 | 0.049789 | F | F | F | F |
| Smurf1 | 0.5894055 | 5.74697703 | 6.0827921 | 0.01638 | 0.049806 | T | F | T | T |

**Table S3. Oligos used in this study.**

| Name | Sequence | Usage |
|---|---|---|
| miR465 up gRNA1 S91 | AGGACTACGATGTACGGAGA | gRNA |
| miR465 up gRNA2 S87 | AGATGAAGGACTACGATGTA | gRNA |
| miR465 down gRNA1 S88 | TTCAGCAGTATCGCCTTACC | gRNA |
| miR465 down gRNA2 S77 | TAAGGCGATACTGCTGAAAA | gRNA |
| miR465 Ext F | GGCCTGATCTATTCTGAAGGGA | genotyping |
| miR465 Ext R | ATCTCACAAATGCCTTTCCGA | genotyping |
| miR465 Int R | GATGAGCTTGCACATATCCACA | genotyping |
| DyzEms3 F | TAGGATGGTAAGCCCAATGC | Sex determination |
| DyzEms3 R | TTGGTTGGTTAATTGTTTGGG | Sex determination |
| Rn18s F | AGAAACGGCTACCACATCCAA | Sex determination |
| Rn18s R | CCTGTATTGTTATTTTTCGTCACTACCT | Sex determination |
| mmu-miR-465a-5p | DIG-ucacaucagugccauucuaaaua-DIG | miRNA-ISH |
| mmu-miR-465a/b/c-3p | DIG-ucuacuuagaaaggcccugauc-DIG | miRNA-ISH |
| mmu-miR-465d-3p | DIG-uuacucagaaaggcccugauca-DIG | miRNA-ISH |
| mmu-miR-NC | DIG-uucuccgaacgugucacgutt-DIG | miRNA-ISH |
| LD_CDS | CAAGCAGAAGACGGCATACGAGATCGTGATGTGACTGGAGTTCAGACGT GTGCTCTTCCGATCTTTTTTTTTTTTTTTTTTTTTVN | miRNA cDNA RT primer |
| Illu lib quant probe | /56-FAM/AGCACACGT/ZEN/CTGAACTCCAGTCAC/3IABkFQ/ | qPCR probe |
| Truseq anti | AGCAGAAGACGGCATACGA | qPCR reverse primer |
| miR-465a-5p | TATTTAGAATGGCACTGATGTGA | qPCR forward primer |
| miR-465b-5p | TATTTAGAATGGTGCTGATCTG | qPCR forward primer |
| miR-465c-5p | TATTTAGAATGGCGCTGATCTG | qPCR forward primer |
| miR-465-3p | GATCAGGGCCTTTCTAAGTAGA | qPCR forward primer |
| miR-465d-5p | TATTTAGAATGGTACTGATGTGA | qPCR forward primer |
| miR-465d-3p | TGATCAGGGCCTTTCTGAGTAA | qPCR forward primer |
| U6 qPCR F | ATACAGAGAAGATTAGCATGGCCC | qPCR forward primer |
| XhoI Rlim 3'UTR F | CGATCGCTCGAGTGAGACCAGAACTCTGAGCTGTGTA | Luciferase construct |

| | | |
|---|---|---|
| NotI Rlim 3'UTR R | CGGCCAGCGGCCGCGATCACTTCATGCATTTTATTGATAGTCTATATTTTAAGTCTGC | Luciferase construct |
| XhoI Alkbh1 3'UTR F | CGATCGCTCGAGAACTTAGGGGCTAGATTCTCTTACTCAGTCA | Luciferase construct |
| NotI Alkbh1 3'UTR R | CGGCCAGCGGCCGCTTTAATTTAAAATAGAGTAACTTTATTCATGGAGAAAACAGCTAA | Luciferase construct |
